# Inferring the Hoxa1 Gene Regulatory Network in Mouse Embryonic Stem Cells: Time-Series RNA-seq Data and Computational Modeling Approach

**DOI:** 10.1101/2023.08.16.553596

**Authors:** Ugochi Emelogu, Bryan Rogers, Yaser Banad, Candice Cavalier, Anna Wilson, Xiaoping Yi, Oswald D’Auverne, Samire Almeida de Oliveira, Nana Akwaboa, Eduardo Martinez-Ceballos

## Abstract

The homeotic gene *Hoxa1* plays a pivotal role in regulating embryonic pattern formation and morphogenesis during mouse embryogenesis. However, despite the identification of a number of putative Hoxa1 target genes, the intricate regulatory relationships between these targets remain largely elusive. Leveraging the advancements in high-throughput technologies and sophisticated computational methods, we aimed to infer the Gene Regulatory Networks (GRNs) governed by Hoxa1 that direct cellular function, morphology, and/or differentiation. To achieve this, we generated time-series RNA-seq data from Retinoic Acid (RA)-treated Wild Type versus *Hoxa1*-null mouse ES cells, enabling the construction of the Hoxa1 GRN. To create this GRN, we employed NARROMI, a published technique known for its noise reduction capabilities and improved accuracy in gene-regulation inference. Using this technique, we identified putative direct and indirect connections between Hoxa1 and a set of genes with known relevance in embryonic development. Validation through qPCR confirmed the Hoxa1-dependence on mRNA expression for selected genes, both within the immediate vicinity (direct) and in secondary interactions (indirect). Furthermore, by mapping the candidate genes to relevant Gene Ontology (GO) networks, we verified their involvement in processes likely regulated by Hoxa1. Our findings provide compelling evidence supporting the accuracy of the NARROMI analysis in generating a hierarchical network of genes under the transcriptional control of Hoxa1 Transcription Factor (TF), specifically in mouse ES cells. This network reveals a pool of promising candidate genes that may function as direct targets of Hoxa1. However, further investigations, including the characterization of Hoxa1 protein interactions with target loci DNA, are necessary to confirm their direct regulatory relationship with this TF. Moreover, the time-series RNA-seq data from Wild Type ES cells, coupled with the methodology employed in this study, hold the potential for constructing GRNs for additional TFs activated by RA. This comprehensive approach can shed further light on the intricate regulatory networks governing cellular function, morphology, and differentiation, advancing our understanding of embryonic development and gene regulation processes.

## 1. Introduction

The homeobox (Hox) family of transcription factors comprises important regulators of embryonic patterning and organogenesis^1-6^. In mammals, the Hox genes are located in four separate chromosome clusters, *Hox*a, *Hoxb, Hoxc* and *Hoxd*. The expression of *Hox* genes during development depends upon their position in the chromosomal cluster; genes positioned at the 3′ end are expressed earlier and more anteriorly, whereas 5′ end genes are expressed at later times and more posteriorly^1-2^. Treatment of ES cells or teratocarcinoma cells with Retinoic Acid (RA, a Vitamin A derivative) results in the sequential activation of several *Hox* genes in a manner that resembles their position in the clusters, e.g. 3′ genes are activated before 5′ genes^7-9^. In fact, a retinoic acid response element (RARE) was discovered in the 3’ enhancer of *Hoxa1*, the most 3’ end gene in the *Hoxa* cluster^9-12^. The spatially restricted patterns of Hox gene expression within the developing central nervous system (CNS) in vertebrates, both along the anteroposterior and dorsoventral axes of the embryo, suggest a regulatory role in patterning of the CNS, as well as in cell specification^13-14^. Inactivation of both alleles of the *Hoxa1* gene in mice results in numerous developmental defects, including hindbrain deficiencies and abnormal skull ossification, and ultimately, in neonatal death^15-18^. The abnormalities of the mouse *Hoxa1* null embryos emphasize the importance of this gene in the proper development of the early brain and neural crest-derived structures^19-21^. Furthermore, although endoderm is relatively normal in *Hoxa1* null embryos, the presence of Hoxa1 expression in a number of endodermal-derived structures in wild type embryos suggests a role for Hoxa1 during the normal endodermal development of vertebrate embryos^22-27^. In humans, mutations in this gene have been described in association with various Central Nervous System (CNS) disorders^28-29^ and its overexpression has been associated with cancer development^30-32^. Thus, these observations highlight the importance of Hoxa1 for proper embryonic development and the onset of human disorders and cancer, which suggest a complex regulatory mechanism that orchestrates the actions of the Hoxa1 signaling pathway in cells. Although the relevance of Hoxa1 during embryonic development has been recognized for over two decades, little is known about the mechanism of action of this important TF. An important barrier to the complete understanding of the mechanism of Hoxa1 action may be the lack of information about its downstream pathway. In this regard, different approaches employing cDNA microarray analyses have been taken in order to elucidate the Hoxa1 pathway through the identification of Hoxa1 targets genes. For instance, Hoxa1 target genes identified from microarray analyses were first reported in a study that employed teratocarcinoma cells^33^, then in mouse Embryonic Stem (ES) cells^34^, and also by researchers that compared embryonic tissues from Wt vs. *Hoxa1*-null embryos^35^. These studies identified different sets of putative Hoxa1 target genes but, due to the nature of the technologies employed, the findings did not provide information about the regulatory interactions between the putative Hoxa1 targets identified.

In this work, we have performed time-series RNA-seq analyses in an attempt to construct the Hoxa1 Gene Regulatory Pathway (GRN) activated during the RA-induced differentiation of mouse ES cells in vitro. To construct the Hoxa1 GRN, we used NARROMI, a noise and redundancy reduction technique shown to improve accuracy of GRN inference^36^, which was shown to significantly outperform other methods^37^such as LP^38^, LASSO^39,40^, ARACNE^41^, and GENIE3^42^.

## 2. Materials and Methods

### 2.1. Cells and drug treatments

For this work, we used mouse J1 Wild-type ES cells (American Type Culture Collection or ATCC), and a Hoxa1 null ES cell line, EMC-ES-Hoxa1–15, whose generation and characterization we previously described^34^. Undifferentiated J1 and Hoxa1–15 ES cells were maintained on gelatin-coated dishes in ES medium (Dulbecco’s modified Eagle’s medium (Invitrogen) supplemented with 10% ES qualified fetal bovine serum (Atlanta Biologicals), 100 mM MEM nonessential Amino acids, 0.1 mM β-mercaptoethanol, Penicillin and Streptomycin (Invitrogen), 1 mM sodium pyruvate, and 1×10^3^ U/ml leukemia inhibitory factor (LIF, Esgro, Millipore), as previously described^43^. To avoid spontaneous differentiation, the ES cells were passaged every two days for up to three passages before drug treatments. For all experiments, ES cells were treated 24 hours after plating with 1 μM all-*trans*-Retinoic Acid (RA) for different periods of time. Control cells were treated with 0.1% ethanol (vehicle). After various times of drug treatment, cells were harvested for total RNA extraction for either RNA sequencing (RNA-seq) or qPCR analyses.

### 2.2. RNA-seq

For time-series RNA-seq, ES cells cultured as monolayers were treated with RA for 12, 24, 36, 48, and 60 hours (RA_12, RA_24, RA_36, RA_48, and RA_60, respectively). Control cells were treated with vehicle only for 48 hours (Ctl_48). To identify Hoxa1 target genes, J1 (Wild type or Wt) and Hoxa1-null (knockout or KO), were treated with RA or vehicle only for 48 hours. Total RNA was extracted from four biological replicates for each treatment and time point combination. Library preparation and sequencing was performed by Novogene (Novogene Inc.).

### 2.3. Data Analysis

Downstream analysis was performed using a combination of programs including STAR, HTseq, Cufflink and Novogene’s wrapped scripts. Alignments were parsed using STAR program and differential expressions were determined through DESeq2/edgeR.

### 2.4. Reads mapping to the reference genome

Reference genome and gene model annotation files were downloaded from genome website browser (NCBI/UCSC/Ensembl) directly. Indexes of the reference genome was built using STAR and paired-end clean reads were aligned to the reference genome using STAR (v2.5). STAR used the method of Maximal Mappable Prefix(MMP) which can generate a precise mapping result for junction reads.

### 2.5. Quantification of gene expression level

STAR will count number reads per gene while mapping. The counts coincide with those produced by htseq-count with default parameters. And then FPKM of each gene was calculated based on the length of the gene and reads count mapped to this gene. FPKM, Reads Per Kilobase of exon model per Million mapped reads, considers the effect of sequencing depth and gene length for the reads count at the same time, and is currently the most commonly used method for estimating gene expression levels^44^.

### 2.6. Network construction

NARROMI, a noise and redundancy reduction technique^36^, was employed to construct the Hoxa1 GRN. This method requires two data sets: a Target Gene (TG) set, and a Transcription Factor (TF) set, where TF is comprised of relevant Transcription Factors present within TG. To obtain the TG set, time-series RNA-seq was performed and data was filtered, based on levels of expression and significant differential expression among treatment, using MATLAB & Simulink software (MathWorks, Natick, Mass.). To construct the TF set for NARROMI, we first obtained a list of all Transcription Factors that were upregulated in RA-treated Wt versus KO cells by at least 2-fold. This list was then compared to the TG set in order to obtain a time-series subset comprised of Hoxa1 dependent Transcription Factors. Subsequently, the Hoxa1 GRN was identified by implementing NARROMI on MATLAB using threshold default conditions as indicated by Zhang et al. (2013)^36^. The resulting network was visualized using Cytoscape^45^.

### 2.7. qPCR analyses

Total RNA was isolated using TRIzol reagent (Thermo Fisher Scientific), according to standard procedures. The concentration of RNA was determined using a NanoDrop (Thermo Scientific). Real-time PCR with SYBR green detection was performed on a QuantStudio 3 thermocycler (Applied Biosystems) using the following primers: *Rplp0* (internal control), forward 5′-AGAACAACCCAGCTCTGGAGAAA-3′, reverse 5′-ACACCCTCCAGAAAGCGAGAGT-3′; *Nanog*, forward 5′-GCGGACTGTGTTCTCTCAGGC-3′, reverse 5′-TTCCAGATGCGTTCACCAGATAG-3′; *Nes*, forward 5′-CAGAGAGGCGCTGGAACAGAGATT-3′, reverse 5′-AGACATAGGTGGGATGGGAGTGCT-3′; *Tubb3*, forward 5′-TCAGCGATGAGCACGGCATA-3′, reverse 5′-CACTCTTTCCGCACGACATC-3′; *Mest*, forward 5′-GCTGCCGCGGTCCACAGTGTCG-3′, reverse 5′-GTCACCCTTAGAGATGAGGTGGAC-3′; *Sox17*, forward 5′-GCTGATCAGCCAGGATGAGCAGCCCGGATGCGGGATACGCC-3′, reverse 5′-ACGAAGGGCCGCTTCTCTGCC-3′; *Zscan4f*, forward 5′-GAGGTACCAGGGTTGCAGTC-3′, reverse 5′-GGATGATTGGCGAAAGCGAC-3′; *Lgals4*, forward 5′-CCCGCCCTTTACTCTGCTTCTGTT-3′, reverse 5′-CTTCCTTGCCCCATTGTCCACTTT-3′, *Col13a1*, forward 5′-GGAGAACCAGGTGACGAAGG-3′, reverse 5′-ACTTGTTCCAGCAGCCTTG-3′; *Coro6*, forward 5′-AGCGCCTTCTGTGCCGTCAA-3′, reverse 5′-CGTCAGGGTGTATGTCGTCCAG-3′.

Reactions were run in triplicate from three independent experiments. Expression data were normalized to the geometric mean of the housekeeping gene Rplp0 (36B4) to control the variability in expression levels and were analyzed using the standard 2 ^-ΔΔCT^ method. Expression levels were analyzed by one-way ANOVA with Tukey’s post-hoc analysis using GraphPad Prism 5.0 software.

## 3. Results

### 3.1 Time-series transcriptomic analysis of Wt ES cells treated with RA

Embryonic Stem Cell (ES) differentiation requires a precise sequence of molecular events that involve specific gene activation and their coordinated regulatory action. This regulatory relationship between genes has traditionally been achieved through biological experiments^46^. New high-throughput technologies, such as time-series RNA-seq and ChIP-seq, provide ample opportunities to identify novel regulatory gene relationships or Gene Regulatory Networks (GRNs). These technologies, together with the availability of null or transgenic animals and/or ES cells, can allow us to reconstruct GRNs for specific TFs and potentially afford us with a dynamic insight into the transcriptional reprogramming that takes place in differentiating ES cells. In previous work, we had generated mouse Hoxa1-null ES cells and used cDNA microarrays to identify a number of putative Hoxa1 target genes^34^. Thus, for this work, we aimed to construct the Hoxa1 GRN on differentiating mouse ES cells using the NARROMI algorithm. NARROMI requires the time-series RNA-seq data to be input as two separate sets: one is the total number of genes or TG, and the other is the Transcription Factor (TF) data set extracted from the TG set. Our strategy was to first generate TG by performing time-series RNA-seq analyses on D3 (Wt) mouse ES cells. For this purpose, ES cells were treated with 1 μM RA for 0, 12, 23, 36, 48, and 60 hours as described in Materials and Methods. The experiments were done by quadruplicate and whole transcriptome analyses were performed by Novogene, Inc. Filtering for low expression and significant differential expression yielded a preliminary list comprised of 16133 genes. The expression profile for the genes in this preliminary set is shown as a clustered heatmap in Figure 1a, where the average values from four repetitions were used for constructing the heatmap. The Venn diagram in Figure 1b shows the overlaps between differentially-expressed genes (DEGs) among different time-point treatments. The differential expression between RA-treated versus control samples at 48 hours is depicted as a volcano plot in figure 1c. To further trim the TG set for optimal NARROMI analysis, genes were selected if they showed a differential expression of 2-fold or higher between any of the RA-treatment time-points and the untreated control. This produced a working TG set comprised of 4060 genes (Supplementary Data 1).

**Fig. 1.**
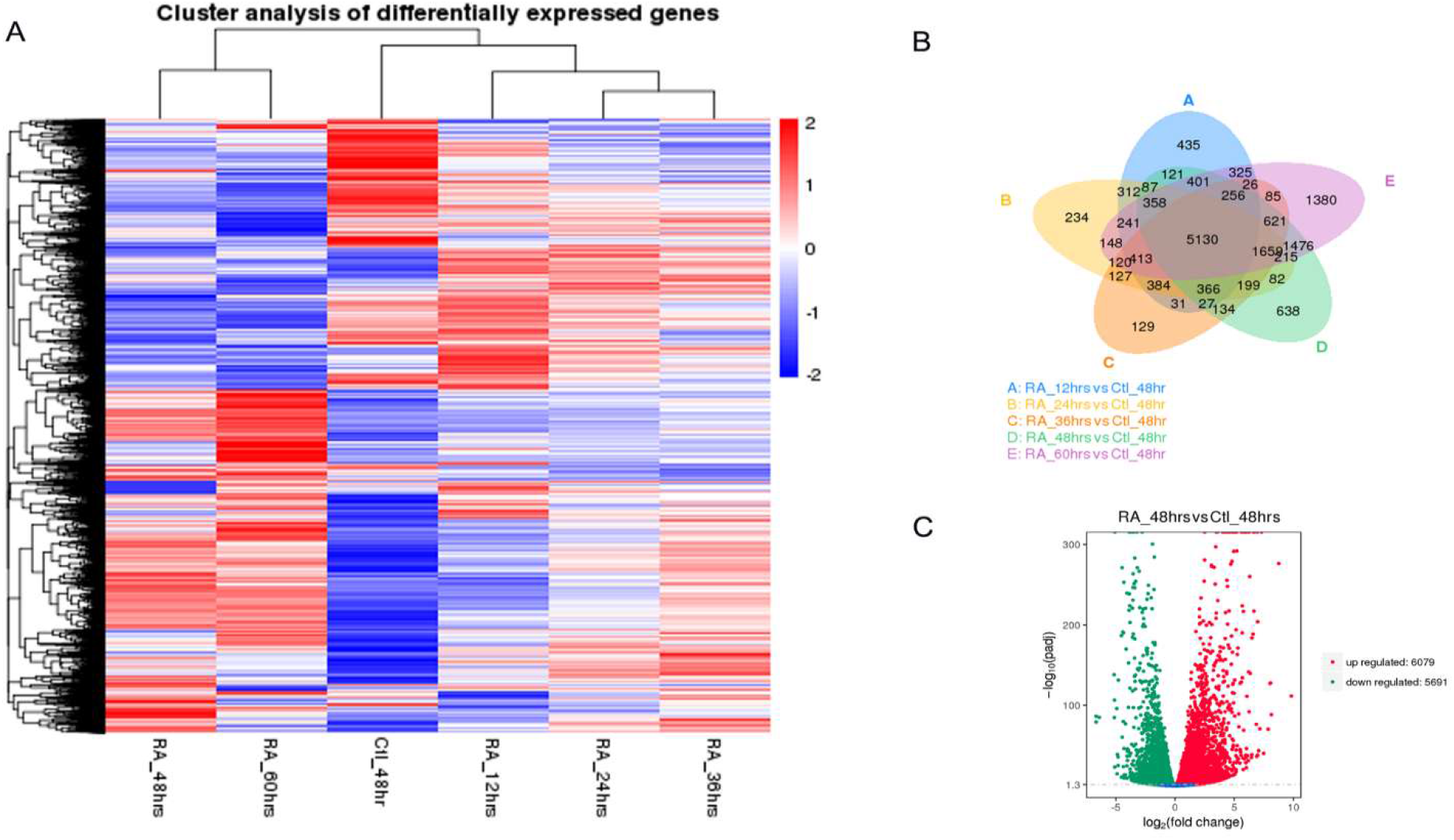
Time series transcriptomic analysis of Wt ES cells treated with RA. The NARROMI algorithm for GRN construction requires a TG (Total Genes) and a TF (Transcription Factor) data sets. Time series RNA-seq data was used to extract the TG set. A) Clustering heatmap for TG set. B) Venn diagram showing overlap between differentially expressed genes (DEGs) in Wt cells at different times of RA treatment. C) Volcano plot for DEGs at 48 hours of treatment.

**Fig. 2.**
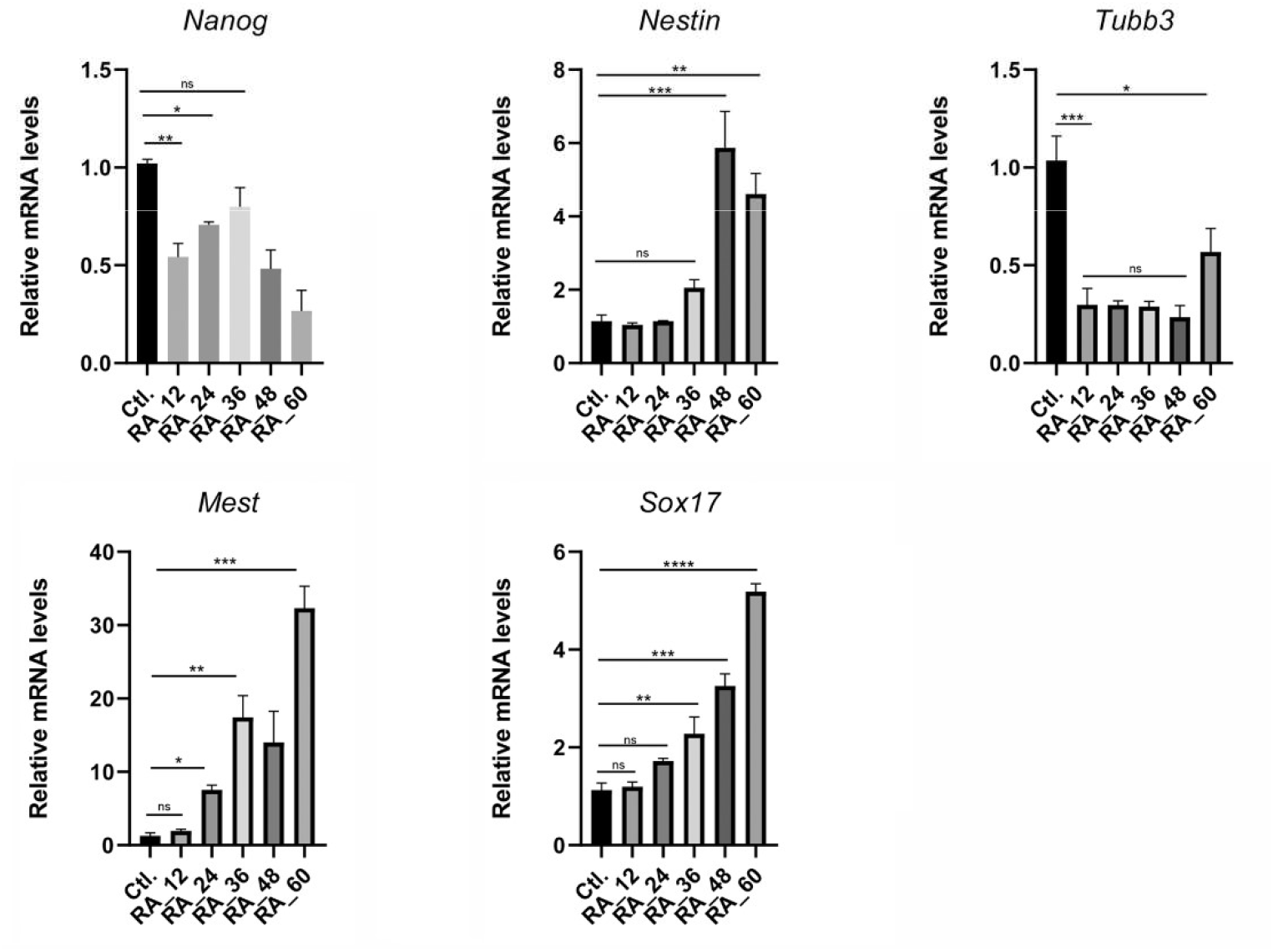
Verification of RNA-seq data by qPCR analysis of ES differentiation markers. The table indicates the genes tested and their FPKM values from RNA-seq. Results from qPCR are shown below. In general, qPCR analyses verified the RNA-seq data for the genes tested. * p<0.05, ** p<0.01, *** p<0.001, **** p<0.0001 by one-way ANOVA. Means ± SEM are shown.

### 3.2. Time-series expression of differentiation markers by qPCR

Our RNA-seq results indicate that markers for all three lineages are upregulated in mouse ES cells after RA treatment. Thus, to begin the validation of our time-series RNA-seq results, we performed qPCR analyses for a group of known differentiation markers, including the stem cell marker *Nanog*, the neuronal precursor *Nestin* (*Nes*), *Mest* (mesodermal marker), and the endodermal marker *Sox17*.

As shown in Table 1 for average RPKM values, RNA-seq results indicate that *Nanog* expression quickly decreased by more than two-fold after 12 hours of RA treatment, with slight increases at subsequent time-points, but always staying close to the two-fold mark decrease as compared to controls. On the other hand, *Nes* expression slowly increased after RA treatment followed by a dramatic increase of about 8-fold by 60 hours of treatment. Expression of *Tubb3*, a neuronal marker also found to be expressed in undifferentiated ES cells, initially decreased after treatment but then increased to reach initial levels of expression at 60 hours of RA treatment. Contrarily, *Mest* expression was found to increase after RA treatment in a time-dependent manner, reaching a maximum of about 13-fold at 48 hours of treatment versus control. *Sox17* expression levels were low in untreated control cells (1.36 FPKM) and increased after RA treatment, reaching a maximum of 5.04 FPKM (about 3.6-fold increase) at 60 hours.

**Table 1.** Expression of differentiation markers in Wt ES cels (in FPKM)

| Table 1. Expression of differentiation markers in Wt ES cells (in FPKM) |  |  |  |  |  |  |
| --- | --- | --- | --- | --- | --- | --- |
| Sample ID | Ctl-48hrs | RA_12hrs | RA_24hrs | RA_36hrs | RA_48hrs | RA_60hrs |
| Nanog | 205.56 | 97.34 | 94.38 | 102.82 | 126.41 | 104.43 |
| Nes | 4.87 | 6.39 | 9.15 | 13.52 | 22.41 | 83.11 |
| Mest | 48.48 | 77.26 | 207.63 | 459.74 | 650.76 | 502.84 |
| Sox17 | 1.36 | 0.94 | 1.26 | 2.24 | 4.01 | 5.04 |
| Tubb3 | 23.12 | 12.43 | 12.15 | 12.09 | 10.17 | 25.39 |

With regards to the qPCR results, *Nanog* gene expression declined, as expected, after treatment of cells with RA. Nanog mRNA expression it decreased by about two-fold after 60 hours of RA treatment. Conversely, *Nes* expression did not significantly change during the first 36 hours of treatment but increased to about 6-fold at 48 hours as compared to controls. *Nes* mRNA expression then decreased to about 5-fold, with respect to controls, at 60 hours. As it was the case with the RNA-seq results, *Tubb3* mRNA expression was detected by qPCR in untreated controls and levels quickly decreased after RA treatment for 12 to 48 hours, and then presented a small increase at 60 hours when*Tubb3* mRNA expression levels were about half of those of controls. In the case of *Mest*, mRNA expression was found to significantly increase with RA treatment at 24 hours, with a steady increase afterwards, reaching a maximum of about 32-fold with respect to controls at 60 hours of RA treatment. In the case of *Sox17*, mRNA expression levels also progressively increased with RA treatment, reaching the highest peak of about 5-fold at 60 hours. Thus, the patterns of marker expression obtained from qPCR analyses were similar to those obtained by RNA-seq, which validates our time-series transcriptome analyses.

### 3.3 Hoxa1 Gene Regulatory Network

Hoxa1 plays important roles during early embryogenesis and, in vitro, it has been found to be required for the neuronal differentiation of mouse ES cells into neurons. Furthermore, in human cells, HOXA1 has been identified as an important oncogene that drives tumor growth and metastasis when overexpressed. Although we and others have reported the identification of multiple putative Hoxa1-related genes, little is known about the hierarchical relationship between Hoxa1 and its target genes, including those that code for transcription factors.

Thus, to infer a Hoxa1-dependent GRN, we next sought to identify transcription factors within the TG set that were differentially expressed in Wt versus Hoxa1-null mouse ES cells as determined by RNA-seq analyses (see Materials and Methods). This analysis identified 124 transcription factors that were differentially expressed in Wt vs Hoxa1 KO Es cells. The resulting TF set used for Narromi is depicted as a heatmap in Fig. 3. The list of genes that constitute the TF set is provided as Supplemental Materials (Supplementary Data 2).

**Fig. 3.**
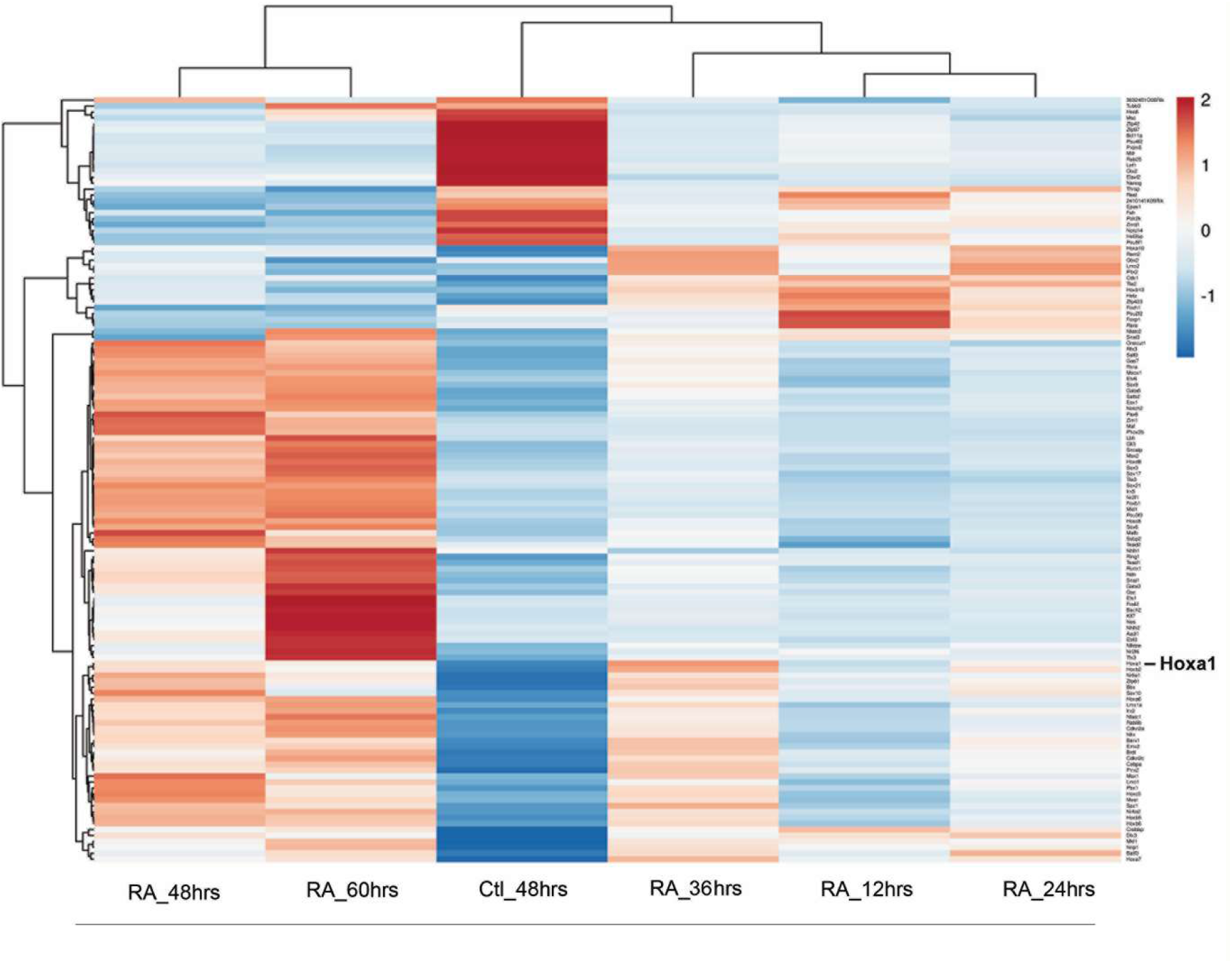
Clustering heatmap for TF data set. Transcription Factors that were differentially expressed between Wild Type versus Hoxa1 null mouse ES cells treated with Retinoic Acid were selected for the TF set for NARROMI.

### 3.4.

To verify the relationship between the selected genes for the TF set and the Hoxa1 signaling pathway, we employed ClueGo, a Cytoscape plugin^47^, to perform pathway analysis of TF members and to attain their complete Gene Ontological terms (GO). The resulting analysis, depicted in Figure 4, indicated that the genes selected for the TF set are involved in processes found to be defective in *Hoxa1* knockout mice. These processes include hindbrain and cranial nerve development, ear morphogenesis, cardiovascular development, and respiration, among others^15-18,48^. Thus, pathway analyses using ClueGo verified the relationship between genes that integrate the TF set and the Hoxa1 signaling pathway.

**Fig. 4.**
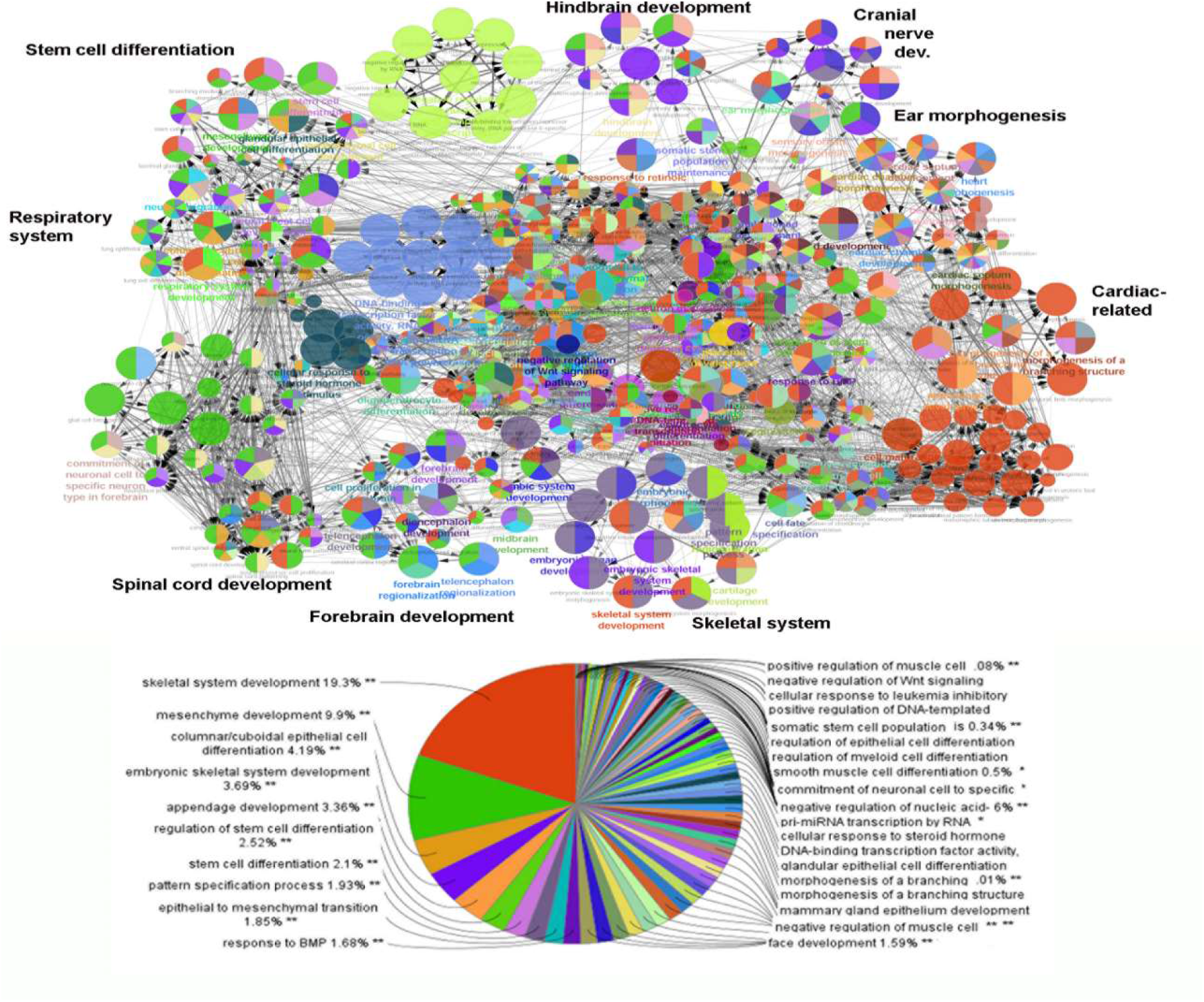
Pathway analysis using ClueGo. Functional analysis indicated that selected TFs interconnected to pathways related to observed deficiencies in *Hoxa1* knockout mice. A) Representation of GO terms enriched in the TF gene set (p<0.005). Processes with observed deficiencies in Hoxa1 knockout mice are shown in bold. B) Pie-chart indicating the prevalence of GO terms among TF genes.

### 3.5. Construction of the Hoxa1 GRN

Following the verification of the Hoxa1-related TF set by Gene Ontology, we then conducted the NARROMI analysis on MATLAB as described in Materials and Methods. The analysis from NARROMI resulted in a network containing a total of 27497 connections among genes or nodes. Those connections were visualized using Cytoscape (Figure 5, top right). An amplified partial view of the GRN is also shown for Hoxa1 and its first-and second-level connections (first-and second-neighbors). In this amplified view, the nodes are colored based on the differential expression of genes at 48 hours of RA treatment versus untreated control. Red color indicates increase of expression, and green indicates decrease of expression after treatment. In this view, the size of the nodes indicates whether the genes are first-Hoxa1 neighbors (larger nodes) or second neighbors (smaller nodes). In this network, NARROMI identified 281 direct connections between Hoxa1 and its putative targets (see Supplementary Data 3). Thus, using NARROMI, we were able to construct the Hoxa1 GRN and identified a set of putative direct targets of this TF, along with its indirect second-tier neighbors. A high-resolution image for figure 5 is provided as Supplementary Data 4.

**Fig. 5.**
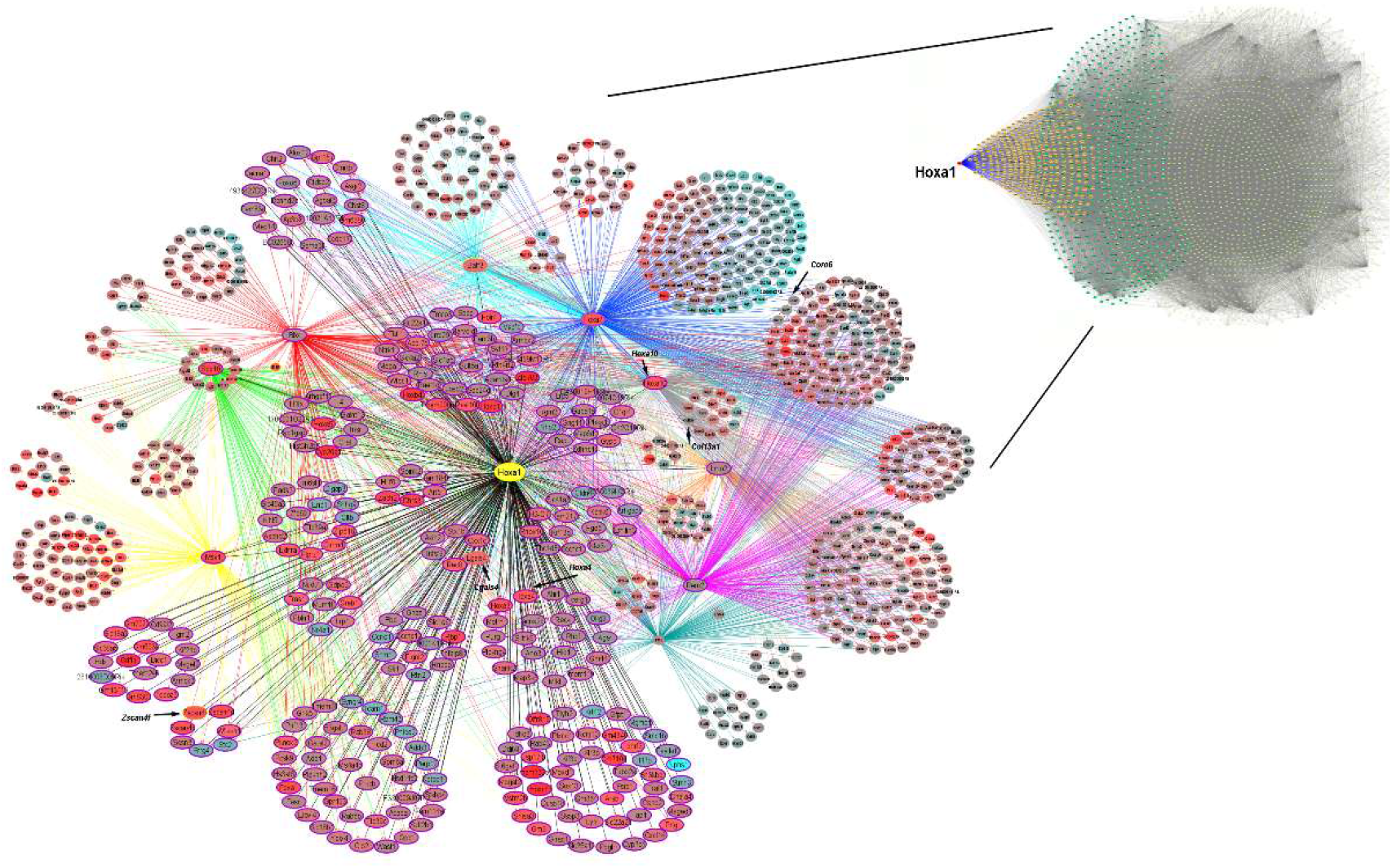
Hoxa1 Gene Regulatory Network. The full network, top right panel (A), contains 27497 connections among genes or nodes, where Hoxa1 is displayed as a red dot towards the left of the network. In the full network, the nodes for the Hoxa1-first neighbors are colored yellow, while the second neighbor nodes are shown in green. All other nodes are gray. (B) Close-up view of the Hoxa1 GRN showing the first- and second-level neighbors of Hoxa1 only. Bigger ovals indicate putative Hoxa1 direct targets and smaller ovals indicate indirect targets or second Hoxa1 neighbors. Node colors indicate whether target genes are upregulated (red) or downregulated (blue) by Hoxa1. The indicated genes were selected for verification of the network by qPCR.

### 3.6.

To verify the gene connections identified by NARROMI, and to test the Hoxa1-dependence of first and second-level neighbors identified in the GRN, we examined the expression of selected genes by qPCR in Wt vs *Hoxa1*-null mouse ES cells cultured in the presence or absence of 1 μM RA for 48 hours. To accomplish this goal, we examined two genes that are directly connected to *Hoxa1, Lgals* and *Zscan4f*, and two genes that are indirectly connected to *Hoxa1* through *Hoxa10, Col13a1* and *Coro6*. As observed in figure 6, the mRNA expression of both *Zscan4f* and *Lgals4* significantly increased in Wt ES cells after RA treatment, but no increase by RA was observed in Hoxa1-null cells (panels A and B). Thus, these results indicate that Hoxa1 is required for the expression of both *Zscan4f* and *Lgals4*. With respect to the qPCR analysis of indirect Hoxa1 target genes expression, we found that both *Col13a1* and *Coro6* mRNA levels were significantly upregulated by treatment with RA (figure 6, panels c and d) after 48 hours. As it was the case with *Zscan4f* and *Lgals4*, RA treatment failed to upregulate the mRNA expression levels of these two genes in Hoxa1-null cells. Interestingly, the levels of *Coro6* mRNA expression were found to be significantly higher in untreated Hoxa1-null than in untreated Wt ES cells, which indicates that small basal differences exist between these two cell lines even in the absence of RA treatment. All together, our qPCR results indicate that the examined genes, both first- and second-level neighbors, are all dependent on Hoxa1 for their expression, which constitutes an initial validation of the Hoxa1 GRN constructed using NARROMI with our time-series RNA-seq data.

**Fig. 6.**
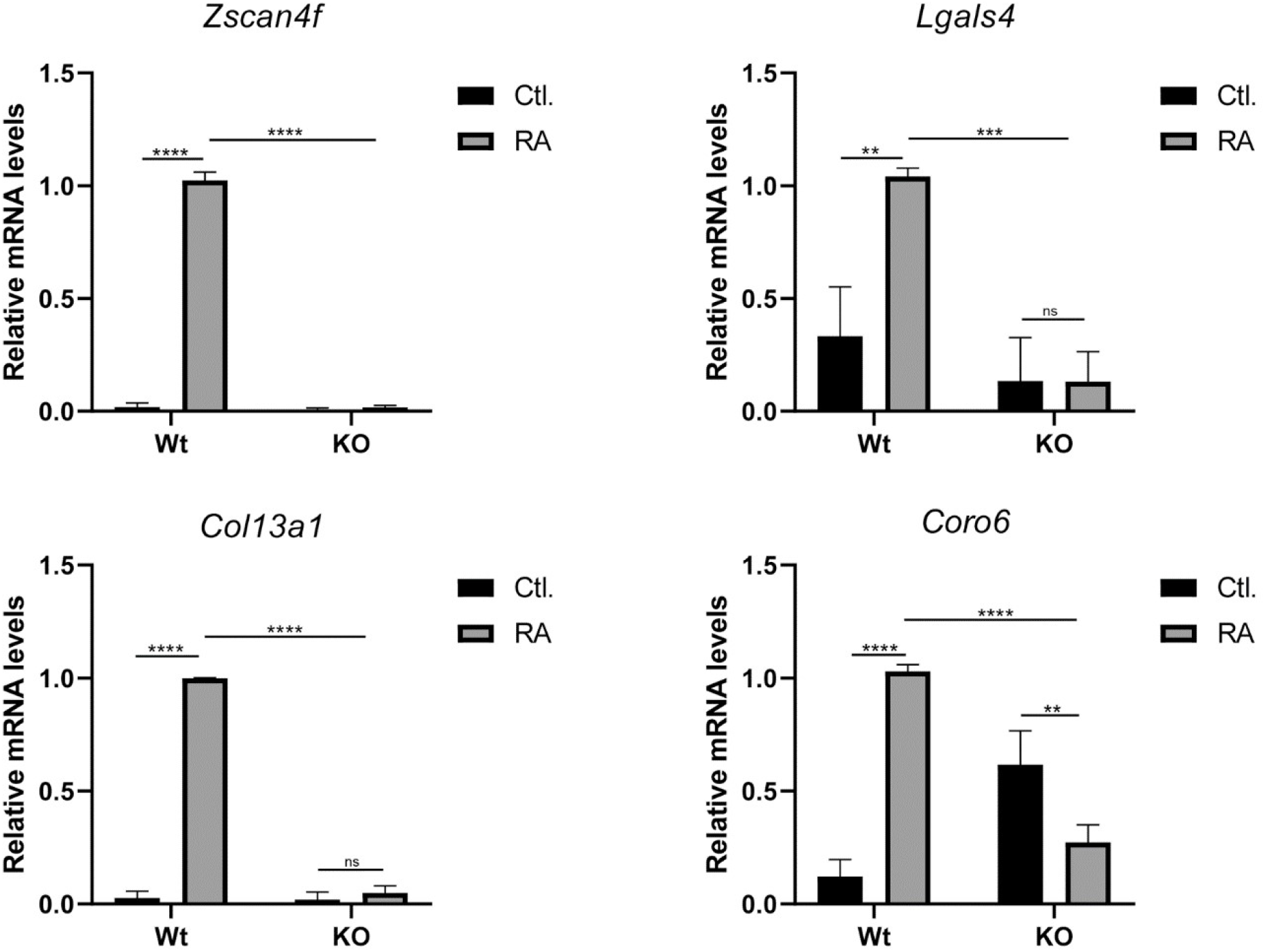
Verification of *Hoxa1* GRN. Expression of selected putative direct (*Zscan4f* and *Lgals4*) and indirect (*Col12a1* and *Coro6*) target genes was examined by qPCR on Wild Type versus Hoxa1-null mouse ES cells grown in the presence (RA) or absence (Ctl.) of Retinoic Acid treatment. These results demonstrated that the genes tested depend on Hoxa 1 for transcriptional activation, which correlates with the findings from the constructed GRN. * p<0.05, ** p<0.01, *** p<0.001, **** p<0.0001 by one-way ANOVA. Means ± SEM are shown.

## 4. Conclusions

Embryonic stem cells are critical in regenerative medicine due to their ability to differentiate into all cell types of the body^49^. Retinoic acid, a potent inducer of ES cell differentiation, acts by activating the expression of genes that code for important developmental proteins, including the members of the Hox family of Transcription Factors^50, 51^. *Hoxa1*, the most 3’ member of the Hoxa cluster, is a direct RA target and a master regulator of mouse embryogenesis^52^. In humans, its overexpression correlates with tumor cell proliferation in different types of cancer ^30-32,53^.

Although, extensive research has shed light into the molecular mechanism of *Hoxa1* action through the identification of putative *Hoxa1* target genes^33-35^, the regulatory relationship between these targets remains largely unknown. In this study, we sought to shed light on this gap of knowledge by constructing a *Hoxa1* Gene Regulatory Network (GRN) using RNA-seq time series data analyzed by the NARROMI algorithm. Our base information for NARROMI consisted in filtered RNA-seq data from Wild-type ES cells treated with 1 μM RA for different times: 0, 12, 24, 36, 48, and 60 hours. These data constituted the TG set required for running the algorithm. To generate a GRN specific for *Hoxa1*, RNA-seq from mouse *Hoxa1*-null ES cells treated with RA for 48 hours was employed to identify TF-coding genes differentially-expressed between Wt and *Hoxa1*-null ES cells treated with RA for 48 hours. This time point was selected for the differential expression analysis because this is when we observed the highest expression levels of *Hoxa1* mRNA in Wt ES cells treated with RA^34^. From the NARROMI analysis, we found a group of 281 genes that were directly connected to *Hoxa1* (Figure 5). This group included Hoxa1 itself, which suggest an autoregulatory feed-back loop for this gene (see Supplementary Data 3). Among the primary *Hoxa1* neighbors, *Zscan4f* and *Lgals* were selected for qPCR verification, and were confirmed to be differentially expressed between Wt and *Hoxa1*-null ES cells treated with RA for 48 hours. Importantly, these genes have been found to be key regulators of various developmental processes. *Zscan4f* plays a role in the regulation of telomere length and genomic stability in ES cells, as well as in the regulation of pluripotency and early embryonic development^54^. Similarly, *Lgals* is involved in a wide range of biological processes, including cell adhesion, signaling, differentiation, apoptosis, and immune modulation^55^. Our qPCR analyses also verified the differential expression of the second-level *Hoxa1* neighbors *Col13a1* and *Coro6*, who were indirectly connected to *Hoxa1* through *Hoxa10*. Research suggests that *Col13a1* plays a role in the development and maintenance of neuromuscular junctions^56^, while *Coro6* was found to be differentially regulated in muscle after denervation^57.^ Similarly, the protein expression of Coronin 6 was upregulated in skeletal muscle during development and after denervation^58^. Interestingly, *Hoxa10*, which connects *Hoxa1* to *Col13a1* and *Coro6* in our GRN, has been found to play a role during murine smooth muscle development^58^ and during the regeneration of skeletal muscle in adult mice^59, 60^, which hints to a role for *Hoxa1* during the development and regeneration of murine muscles.

Although qPCR was used as one validation method for the *Hoxa1* GRN, there are limitations that must be considered when using GRNs. One limitation of using a gene regulatory network (GRN) is that they are often constructed based on data from a single cell type or developmental stage, which may not accurately reflect the complex interactions that occur between genes and signaling pathways in vivo^61^. The GRN constructed by our team addresses this issue by employing RNA-seq data from both Wt and Hoxa1-null mouse ES cells. Nevertheless, as the accuracy of GRN predictions are known to vary widely depending on the algorithm and modeling approach used, which highlights the challenges of constructing reliable GRNs^61^, these limitations underscore the need for rigorous experimental validation of GRN predictions and the use of complementary approaches, such as genetic perturbations and single-cell transcriptomics, to refine and improve GRN models. To refine our GRN model, we are currently performing proteomic analyses on Wt vs Hoxa1-null mouse ES cells treated with RA for various periods of time. Taken together, our approach is the first step on the elucidation of the regulatory interactions among Hoxa1 and its targets. Furthermore, our TG data can be used to construct a GRN for other TFs if a list of differentially expressed transcription factors is generated or obtained from published data for mouse ES cells.

## Contributions

EMC conceptualized the study. EMC and UE conducted most of the experiments and performed NARROMI analyses. YB handed bioinformatics data. BR wrote sections of the manuscript and provided advice on the interpretation of genomic data., XY, AW, and OD wrote sections of the manuscript and provided editorial advice. CC submitted the RNA-seq data to the database. SAO and NA performed some qPCR experiments and performed the ClueGo analyses.

## Data Availability

Transcriptome data has been deposited in NCBI’s BioProject accession number PRJNA770639.

