## Supplementary Data 1 for "Inferring the Hoxa1 Gene Regulatory Network in Mouse Embryonic Stem Cells: Time-Series RNA-seq Data and Computational Modeling Approach"

TG list for NARROMI

RNA-seq was performed by quadruplicate. As NARROMI required duplicate values per condition, averages from each two of the quadruplicates were used for the TG set.

| Sample ID | Ctl1 | Ctl2 | 12RA1 | 12RA2 | 24RA1 | 24RA2 | 36RA1 | 36RA2 | 48RA1 | 48RA2 | 60RA1 | 60RA2 |
|---|---|---|---|---|---|---|---|---|---|---|---|---|
| Dbx1 | 0.125242 | 0.060324 | 0.023445 | 0.1 | 0.1 | 0.020577 | 0.043354 | 0.022414 | 10.65552 | 9.30744 | 20.46944 | 19.64139 |
| Sftpc | 41.34681 | 7.865645 | 0.029162 | 5.170108 | 3.802124 | 2.292986 | 4.378901 | 8.97703 | 0.075396 | 0.887734 | 0.1 | 0.028076 |
| Foxb1 | 0.006708 | 0.021136 | 0.008571 | 0.0077 | 0.033786 | 0.051517 | 0.112409 | 0.20418 | 2.245609 | 1.905847 | 2.110749 | 2.387489 |
| Ccl2 | 0.073306 | 0.078971 | 0.132243 | 0.057526 | 0.036758 | 0.021058 | 0.04199 | 0.085524 | 0.213928 | 0.050644 | 5.06255 | 4.826338 |
| Actc1 | 0.301031 | 0.129076 | 0.137462 | 0.272785 | 0.205467 | 0.270335 | 0.7253 | 0.636349 | 2.173328 | 2.115489 | 55.31904 | 51.48332 |
| Ascl1 | 0.040676 | 0.017071 | 0.189002 | 0.116441 | 0.364986 | 0.296252 | 0.765381 | 0.850068 | 8.2191 | 7.905828 | 21.645 | 20.33531 |
| Igfbp5 | 0.718667 | 0.639873 | 0.282487 | 0.176773 | 0.576961 | 0.442947 | 1.51026 | 1.654316 | 24.81852 | 26.44575 | 24.67375 | 24.07505 |
| Ednrb | 0.162106 | 0.100982 | 0.164574 | 0.152924 | 0.4801 | 0.42666 | 1.770424 | 1.398141 | 16.04976 | 17.1195 | 20.43317 | 22.54291 |
| Phox2b | 0.067121 | 0.091092 | 0.015595 | 0.021619 | 0.060255 | 0.037335 | 0.221712 | 0.249376 | 2.933853 | 2.903862 | 2.336901 | 2.115423 |
| Maf | 0.047514 | 0.039811 | 0.03988 | 0.031589 | 0.082314 | 0.108062 | 0.232233 | 0.140012 | 3.924189 | 3.801457 | 3.114939 | 3.049156 |
| Vit | 0.01482 | 0.038419 | 0.00862 | 0.039353 | 0.066389 | 0.032789 | 0.105889 | 0.017231 | 1.042869 | 0.975114 | 1.778304 | 1.975181 |
| Prdm8 | 0.038951 | 0.034096 | 0.118906 | 0.062528 | 0.13585 | 0.122782 | 0.572823 | 0.555974 | 7.524739 | 6.556677 | 5.438663 | 5.457116 |
| Itgb8 | 0.115364 | 0.131266 | 0.079394 | 0.121747 | 0.167703 | 0.18406 | 0.337923 | 0.356955 | 4.992316 | 5.300084 | 9.686783 | 11.03209 |
| Ccdc42 | 0.1 | 0.1 | 0.052362 | 0.062156 | 0.174359 | 0.082249 | 0.52958 | 0.541816 | 2.766538 | 2.036726 | 5.417237 | 5.42282 |
| Fam181a | 0.071314 | 0.089542 | 0.067594 | 0.13882 | 0.170772 | 0.094277 | 0.596619 | 0.56148 | 6.775994 | 4.865648 | 7.678256 | 7.826988 |
| 1810011O1 | 0.211777 | 0.175217 | 0.081479 | 0.156792 | 0.234521 | 0.490634 | 1.086767 | 1.246327 | 6.527839 | 5.445023 | 9.756064 | 11.65035 |
| Fezf2 | 0.069984 | 0.064539 | 0.062907 | 0.036486 | 0.061331 | 0.10543 | 0.416782 | 0.305124 | 0.811862 | 0.668496 | 2.642809 | 2.298171 |
| Acta2 | 3.023409 | 2.3791 | 4.722426 | 4.711584 | 2.702127 | 2.966588 | 5.365958 | 5.315638 | 18.01959 | 16.37697 | 192.271 | 180.6245 |
| Col12a1 | 0.06644 | 0.049632 | 0.075547 | 0.134055 | 0.12944 | 0.137593 | 0.252022 | 0.199504 | 1.68417 | 1.743357 | 7.228326 | 8.064523 |
| Cd248 | 0.094261 | 0.057364 | 0.084093 | 0.034369 | 0.05496 | 0.087096 | 0.15009 | 0.091537 | 0.277892 | 0.264815 | 2.39455 | 1.908464 |
| Kirrel2 | 4.011877 | 4.569632 | 1.768086 | 1.213918 | 0.402448 | 0.418644 | 0.14695 | 0.168198 | 0.015283 | 0.054366 | 0.06302 | 0.02187 |
| Pyy | 0.1 | 0.113637 | 0.046084 | 0.041399 | 0.290717 | 0.241044 | 0.908508 | 0.591116 | 3.65617 | 2.150177 | 2.714902 | 2.288162 |
| Vgf | 0.703941 | 0.621996 | 0.967944 | 0.852884 | 0.752913 | 0.665145 | 0.698599 | 0.628423 | 0.4928 | 0.295159 | 18.4787 | 15.74552 |
| Nedd9 | 0.247543 | 0.22317 | 0.563337 | 0.704231 | 1.800919 | 1.536448 | 4.020809 | 3.77977 | 29.24162 | 26.86992 | 33.11327 | 37.30902 |
| Thbs1 | 1.135079 | 1.275515 | 1.333605 | 1.461738 | 1.436561 | 1.490962 | 3.516495 | 2.997314 | 29.7635 | 27.64539 | 64.25166 | 76.24687 |
| Ptn | 2.424193 | 2.729839 | 0.993473 | 0.947377 | 2.791739 | 2.750562 | 6.306687 | 7.420816 | 51.42859 | 49.07814 | 39.5591 | 40.83568 |
| Ankrd1 | 0.251386 | 0.190401 | 0.280164 | 0.176695 | 0.228136 | 0.226499 | 0.316478 | 0.424857 | 1.169735 | 0.894016 | 8.054149 | 9.079339 |
| Nphs1 | 15.42895 | 17.50793 | 6.514561 | 5.775978 | 1.581854 | 1.436231 | 0.375927 | 0.488199 | 0.067553 | 0.197017 | 0.14249 | 0.114905 |
| 4930526L0 | 2.641868 | 2.936937 | 1.103255 | 1.103834 | 0.611293 | 0.412215 | 0.135092 | 0.321007 | 0.213564 | 0.223044 | 0.078791 | 0.022087 |
| Hoxd4 | 0.015746 | 0.1 | 0.037467 | 0.050022 | 0.117749 | 0.111694 | 0.436937 | 0.164767 | 2.709517 | 2.497147 | 1.97791 | 2.254497 |
| Tagln | 5.171499 | 4.628546 | 8.431665 | 8.488157 | 6.796286 | 7.846075 | 14.0498 | 12.08615 | 33.07456 | 29.65451 | 377.9503 | 377.4998 |
| Fam181b | 0.131735 | 0.116346 | 0.105644 | 0.107291 | 0.241808 | 0.303175 | 0.459598 | 0.561588 | 4.979412 | 4.934143 | 3.819993 | 3.79669 |
| Robo2 | 0.064745 | 0.080637 | 0.062967 | 0.048253 | 0.123998 | 0.103135 | 0.209618 | 0.213452 | 1.853429 | 1.795774 | 2.143687 | 2.203156 |
| Tmem171 | 0.1 | 0.1 | 0.1 | 0.1 | 0.120838 | 0.150938 | 0.095506 | 0.242207 | 0.323779 | 0.047809 | 2.312546 | 2.176489 |
| 2810055G2 | 0.05305 | 0.012377 | 0.04111 | 0.052569 | 0.135238 | 0.1366 | 0.568361 | 0.616312 | 2.2934 | 2.391136 | 1.790234 | 2.322435 |
| A830082K1 | 0.024166 | 0.010142 | 0.017864 | 0.035697 | 0.173551 | 0.123365 | 0.31996 | 0.269351 | 1.527878 | 1.421661 | 1.291037 | 1.621859 |
| Hes3 | 0.518164 | 0.495091 | 0.175967 | 0.072158 | 0.055232 | 0.143717 | 0.257361 | 0.256244 | 3.076873 | 2.641163 | 4.371631 | 3.266633 |
| Gcnt4 | 0.292962 | 0.260595 | 0.15066 | 0.173141 | 0.367523 | 0.23372 | 0.5358 | 0.533731 | 6.72957 | 6.590721 | 6.620616 | 7.837206 |
| Hoxc8 | 0.018134 | 0.019052 | 0.144779 | 0.24693 | 1.331137 | 1.436746 | 3.750926 | 4.469556 | 12.09654 | 10.86555 | 10.15163 | 10.87219 |
| Nppb | 0.307905 | 0.271148 | 0.060042 | 0.112657 | 0.155268 | 0.16728 | 0.171438 | 0.420064 | 0.302775 | 0.091388 | 3.954729 | 3.984416 |
| Olfr985 | 3.175887 | 3.158574 | 1.793386 | 1.55093 | 0.920792 | 0.916225 | 0.924801 | 0.797916 | 0.33235 | 0.364145 | 0.056545 | 0.035993 |
| Nr2f2 | 0.076419 | 0.123814 | 0.183345 | 0.214868 | 0.921598 | 0.902828 | 3.048076 | 2.154553 | 9.716306 | 9.216296 | 9.427342 | 9.06865 |
| Edn1 | 0.276227 | 0.276721 | 0.372821 | 0.272555 | 0.309538 | 0.249322 | 0.513503 | 0.39989 | 1.92324 | 1.808059 | 9.57276 | 10.51894 |
| Tgfb2 | 0.081137 | 0.111948 | 0.135779 | 0.088243 | 0.106385 | 0.12658 | 0.193491 | 0.119028 | 0.575104 | 0.472806 | 3.717655 | 3.72129 |
| Neurog1 | 0.036364 | 0.039128 | 0.116915 | 0.156741 | 0.218673 | 0.066726 | 0.145868 | 0.254252 | 2.608217 | 2.664542 | 3.227137 | 2.76796 |
| Srms | 0.188918 | 0.223353 | 4.412616 | 4.302082 | 3.159447 | 3.653781 | 1.760849 | 1.774437 | 0.194045 | 0.206578 | 0.073893 | 0.105796 |
| Agtr1a | 0.256893 | 0.335166 | 0.623243 | 0.444581 | 1.090655 | 1.001306 | 2.118099 | 2.088665 | 18.3671 | 17.9637 | 16.62329 | 17.08431 |
| Sytl2 | 0.074297 | 0.10715 | 0.095591 | 0.050138 | 0.083482 | 0.066123 | 0.19599 | 0.075453 | 0.479022 | 0.549344 | 1.862264 | 1.950014 |
| Krt13 | 0.39163 | 0.115509 | 3.286057 | 0.948552 | 0.194855 | 0.107843 | 0.83612 | 0.481705 | 3.708854 | 4.140023 | 6.694049 | 4.006469 |
| Cnn1 | 1.634104 | 1.173442 | 1.38705 | 1.07586 | 1.097795 | 1.186386 | 1.615024 | 1.563556 | 2.971028 | 2.708822 | 43.04941 | 41.08815 |
| Myl9 | 2.073705 | 1.782268 | 3.926538 | 3.545555 | 3.806588 | 3.875608 | 7.46768 | 6.481619 | 18.03578 | 15.20105 | 136.6328 | 134.0772 |
| Hoxc9 | 0.01698 | 0.1 | 0.120311 | 0.128065 | 0.560344 | 0.493484 | 1.657064 | 1.223993 | 5.203411 | 3.982927 | 5.376167 | 4.666712 |
| Cyp1b1 | 0.696316 | 0.693235 | 1.831756 | 1.736813 | 5.718637 | 5.588304 | 15.40879 | 14.78763 | 59.88879 | 61.0325 | 49.02951 | 52.22303 |
| Pou3f4 | 0.082604 | 0.100939 | 0.133942 | 0.059429 | 0.179213 | 0.144416 | 0.266513 | 0.288117 | 1.881337 | 2.017494 | 1.97764 | 2.007934 |
| Adamts5 | 0.032505 | 0.01303 | 0.032403 | 0.070199 | 0.089474 | 0.087602 | 0.119981 | 0.081095 | 0.48188 | 0.584553 | 2.3501 | 2.305461 |
| Hp | 1.970055 | 0.864158 | 0.073005 | 0.544545 | 0.46639 | 0.446505 | 0.373838 | 1.012022 | 0.044391 | 0.243566 | 0.046988 | 0.031763 |
| Smoc2 | 0.09257 | 0.083394 | 0.10311 | 0.068865 | 0.20659 | 0.222106 | 0.174828 | 0.529572 | 1.028044 | 1.068122 | 2.330523 | 2.171769 |

| | | | | | | | | | | | | |
|---|---|---|---|---|---|---|---|---|---|---|---|---|
| Pcdh18 | 0.421366 | 0.585696 | 0.235498 | 0.225011 | 0.623548 | 0.6168 | 1.610073 | 1.568887 | 6.324617 | 7.039378 | 4.027643 | 4.576925 |
| Lbh | 0.916586 | 0.7867 | 0.685899 | 0.67208 | 1.262432 | 1.242566 | 3.029384 | 2.936841 | 13.49905 | 11.86053 | 21.12797 | 20.98653 |
| Nsg2 | 1.276291 | 1.288773 | 0.255122 | 0.15631 | 0.279208 | 0.404628 | 1.044337 | 1.151268 | 4.539198 | 4.624109 | 4.947983 | 4.815838 |
| Nr2f1 | 0.119715 | 0.106803 | 0.311043 | 0.409083 | 1.161762 | 1.033229 | 2.299305 | 2.572064 | 12.58986 | 11.04753 | 12.85605 | 12.14948 |
| Tox3 | 0.859697 | 0.976002 | 0.785634 | 1.029736 | 2.256378 | 2.154668 | 6.461801 | 6.032951 | 27.4857 | 27.45563 | 28.58321 | 30.44612 |
| Plat | 0.939463 | 0.919267 | 2.088491 | 2.01547 | 7.306848 | 7.698441 | 19.86047 | 19.8875 | 39.25209 | 38.44307 | 57.29797 | 59.13358 |
| Csf1 | 0.352633 | 0.470612 | 0.583393 | 0.630328 | 0.882734 | 0.851647 | 1.356651 | 1.380424 | 1.921948 | 1.978905 | 19.39942 | 18.48798 |
| 2610528J1 | 3.031365 | 2.361872 | 11.59379 | 10.71655 | 6.34908 | 6.718837 | 2.499781 | 3.033152 | 0.578036 | 0.370387 | 0.511612 | 0.453247 |
| Ccnd2 | 1.220914 | 1.50674 | 2.53372 | 2.372589 | 3.365869 | 3.411405 | 5.586539 | 4.315948 | 51.24174 | 50.93404 | 63.13787 | 68.03819 |
| Zbtb16 | 0.264978 | 0.248594 | 0.400377 | 0.580907 | 1.90293 | 1.68109 | 5.091078 | 4.741847 | 16.5686 | 16.64607 | 14.02057 | 15.25924 |
| Pou3f2 | 0.288505 | 0.288727 | 0.279258 | 0.312857 | 0.435868 | 0.446127 | 0.700104 | 0.612368 | 7.378672 | 6.993678 | 8.997336 | 8.899758 |
| Fosl2 | 0.323398 | 0.292256 | 0.648715 | 0.701354 | 0.972593 | 0.902658 | 1.601619 | 1.416819 | 2.512885 | 2.550337 | 18.37507 | 18.94237 |
| Vstm4 | 0.095542 | 0.030904 | 0.175742 | 0.087916 | 0.477637 | 0.39274 | 0.953584 | 0.82073 | 1.639501 | 1.612003 | 2.598154 | 2.368372 |
| Sost | 0.009688 | 0.092027 | 0.118642 | 0.102672 | 0.730011 | 0.785375 | 1.856184 | 1.451836 | 1.443877 | 1.077526 | 3.156182 | 2.751151 |
| Megf10 | 0.142555 | 0.144163 | 0.192039 | 0.201075 | 0.46874 | 0.68061 | 0.992223 | 0.698578 | 4.873317 | 5.008201 | 5.214407 | 5.330961 |
| Acta1 | 0.632836 | 0.677206 | 1.67629 | 1.267873 | 0.955144 | 0.914174 | 1.819137 | 2.093157 | 2.292186 | 2.262819 | 27.07338 | 24.12016 |
| Vrtn | 4.693445 | 4.169749 | 0.737943 | 0.602198 | 0.292837 | 0.349093 | 0.135218 | 0.081745 | 0.067577 | 0.023045 | 0.025772 | 0.03281 |
| Id3 | 6.847031 | 6.400738 | 2.299866 | 1.73145 | 2.726336 | 2.248419 | 5.32141 | 5.684193 | 55.91305 | 45.20853 | 38.57631 | 33.01636 |
| Cbr2 | 10.42222 | 1.483102 | 0.08833 | 0.743414 | 0.619917 | 0.379535 | 0.476798 | 1.393346 | 0.1 | 0.419735 | 0.1 | 0.056524 |
| Kdr | 0.276411 | 0.114547 | 0.093908 | 0.07814 | 0.146092 | 0.090066 | 0.191529 | 0.158228 | 0.27908 | 0.292774 | 1.90305 | 1.874324 |
| Slitrk2 | 0.049167 | 0.09001 | 0.122909 | 0.101421 | 0.212945 | 0.210762 | 0.304473 | 0.361955 | 2.349844 | 2.397317 | 1.666729 | 1.699597 |
| Irx3 | 0.896968 | 0.95195 | 1.205359 | 1.613394 | 5.217927 | 5.534165 | 12.64297 | 13.43164 | 43.13268 | 36.44417 | 42.74259 | 37.28265 |
| Clca6 | 1.723652 | 2.147652 | 1.304119 | 1.56987 | 0.743124 | 0.63974 | 0.400683 | 0.624062 | 0.112016 | 0.128005 | 0.064825 | 0.069343 |
| Syt6 | 0.045453 | 0.033881 | 0.055024 | 0.148099 | 0.141289 | 0.212769 | 0.417381 | 0.488199 | 2.94585 | 3.305932 | 2.890347 | 2.947041 |
| Sox21 | 2.270073 | 2.118928 | 1.086753 | 0.838404 | 1.658449 | 1.706863 | 4.114446 | 4.145128 | 21.26084 | 17.82123 | 18.89777 | 18.56398 |
| Ntn1 | 3.25083 | 4.266685 | 1.001852 | 0.758573 | 1.080734 | 1.034858 | 2.188877 | 1.938833 | 13.42806 | 14.25207 | 17.79462 | 16.78425 |
| Meox1 | 0.162461 | 0.192581 | 1.108767 | 1.276401 | 4.566431 | 4.351658 | 15.18094 | 14.53553 | 28.48094 | 28.113 | 27.28132 | 24.5506 |
| Cxcl10 | 0.073694 | 0.305496 | 0.204569 | 0.225648 | 0.165804 | 0.436919 | 0.541789 | 0.17136 | 0.197679 | 0.248209 | 4.003102 | 3.771001 |
| Cdh11 | 0.152784 | 0.126339 | 0.120829 | 0.131595 | 0.343856 | 0.418624 | 0.729124 | 0.922668 | 2.929099 | 2.870853 | 2.028655 | 2.487201 |
| Stmn4 | 0.11827 | 0.138342 | 0.161306 | 0.246028 | 0.148003 | 0.245161 | 0.362765 | 0.482101 | 0.706885 | 0.770435 | 5.983548 | 5.258485 |
| Daam2 | 0.093727 | 0.070216 | 0.148795 | 0.125897 | 0.226321 | 0.298114 | 0.423956 | 0.341133 | 2.707033 | 2.661031 | 2.563612 | 2.679127 |
| Sema3c | 0.584212 | 0.412742 | 0.450666 | 0.522547 | 0.70776 | 0.736816 | 1.258836 | 1.083977 | 4.943448 | 4.830812 | 10.01155 | 11.07478 |
| Calcrl | 0.545705 | 0.1157 | 0.135343 | 0.13597 | 0.210862 | 0.243821 | 0.428092 | 0.393599 | 2.545697 | 2.847437 | 2.000676 | 2.271315 |
| Prss23 | 0.112374 | 0.105602 | 0.099105 | 0.277428 | 0.317785 | 0.254026 | 0.488959 | 0.624701 | 4.476597 | 4.514491 | 4.950322 | 5.212525 |
| Cdh6 | 0.32184 | 0.610432 | 0.697251 | 0.766629 | 2.095891 | 1.902774 | 4.084554 | 3.899655 | 15.6548 | 15.72741 | 9.337372 | 9.565316 |
| Nhlh1 | 0.433142 | 0.571351 | 0.141863 | 0.277136 | 0.163582 | 0.098584 | 0.017549 | 0.076083 | 0.616815 | 0.654054 | 1.266548 | 1.539913 |
| Hist1h2af | 274.1791 | 154.1994 | 5.235164 | 23.43865 | 63.63392 | 91.64197 | 33.06165 | 201.6812 | 73.01245 | 117.8048 | 13.14054 | 10.03 |
| Slc25a21 | 0.015451 | 0.1 | 0.20424 | 0.133391 | 0.678562 | 0.604664 | 1.245053 | 1.221526 | 2.255447 | 1.889578 | 2.330739 | 2.652652 |
| Maml3 | 0.315543 | 0.407125 | 0.388385 | 0.406758 | 1.02307 | 0.93112 | 1.75415 | 1.589336 | 7.868077 | 8.017823 | 5.061868 | 5.474746 |
| Igf2os | 0.084717 | 0.053379 | 0.164608 | 0.152536 | 0.282523 | 0.289979 | 0.425029 | 0.559109 | 1.158386 | 1.273696 | 3.284003 | 2.959911 |
| Tarm1 | 1.045968 | 1.588171 | 2.849287 | 2.637263 | 1.603905 | 1.424367 | 0.98449 | 1.135189 | 0.424337 | 0.994565 | 0.151823 | 0.141861 |
| Tgfb3 | 0.113125 | 0.187464 | 0.327524 | 0.330055 | 0.791468 | 0.692056 | 1.746131 | 1.454209 | 5.091034 | 6.124255 | 2.613114 | 2.876426 |
| Cxcr4 | 0.547078 | 0.470822 | 0.419202 | 0.325228 | 0.62494 | 0.425832 | 0.797061 | 0.90878 | 6.137141 | 6.02909 | 5.470655 | 5.734782 |
| Olfml3 | 0.233592 | 0.158656 | 0.45777 | 0.483409 | 1.268921 | 1.334443 | 3.064492 | 2.966719 | 9.660853 | 8.914473 | 8.568243 | 8.500796 |
| Pou3f3 | 0.073432 | 0.090841 | 0.199075 | 0.205512 | 0.190476 | 0.306746 | 0.408688 | 0.392193 | 3.381224 | 3.111004 | 4.16966 | 3.777257 |
| Cdh2 | 2.43598 | 2.742098 | 2.705337 | 2.351643 | 4.802925 | 4.509351 | 8.63291 | 8.506333 | 36.11365 | 35.29015 | 41.16936 | 42.81228 |
| Fas | 0.311868 | 0.285016 | 0.229496 | 0.376944 | 0.487717 | 0.723374 | 0.659822 | 0.977712 | 0.571899 | 0.656309 | 6.529605 | 6.767933 |
| Plagl1 | 0.960968 | 1.114758 | 1.407312 | 1.240954 | 2.07878 | 1.993089 | 3.577858 | 3.572947 | 12.53328 | 10.49526 | 21.18774 | 22.15254 |
| Pitpnm3 | 0.073668 | 0.08287 | 0.104702 | 0.090752 | 0.319432 | 0.347123 | 0.766299 | 0.80793 | 1.680356 | 1.61031 | 0.837838 | 0.894655 |
| Ptprcap | 1.477297 | 1.394995 | 7.914111 | 7.15962 | 3.408345 | 2.889207 | 1.773896 | 2.123428 | 0.612685 | 0.404153 | 0.789318 | 0.887738 |
| Eya4 | 0.044375 | 0.066729 | 0.098472 | 0.102397 | 0.142616 | 0.136062 | 0.311433 | 0.274947 | 1.325037 | 1.580056 | 1.617664 | 1.811188 |
| Ltb | 0.354913 | 0.239115 | 0.192534 | 0.198341 | 0.127438 | 0.142168 | 0.207183 | 0.132299 | 0.222505 | 0.131501 | 3.470368 | 2.314607 |
| AU015228 | 0.030538 | 0.095559 | 0.130792 | 0.084061 | 0.244313 | 0.217807 | 0.374033 | 0.532581 | 0.75794 | 0.872526 | 2.105833 | 1.427565 |
| Timp3 | 1.276341 | 0.972101 | 0.721768 | 0.876579 | 0.947224 | 0.876935 | 1.111316 | 1.161256 | 1.747209 | 2.002794 | 13.30511 | 14.48658 |
| C430049B0 | 0.811596 | 0.611521 | 0.68019 | 0.806402 | 0.883476 | 1.424532 | 1.701612 | 1.835808 | 9.421591 | 9.585082 | 13.87346 | 13.26153 |
| Nhlh2 | 0.294588 | 0.283245 | 0.237106 | 0.204826 | 0.210474 | 0.113746 | 0.179903 | 0.153754 | 0.58979 | 0.60836 | 1.867949 | 1.859014 |
| Tnc | 0.084935 | 0.079619 | 0.138276 | 0.222596 | 0.133101 | 0.185014 | 0.248775 | 0.23084 | 0.847861 | 0.876725 | 3.320152 | 3.020697 |
| Dclk1 | 0.12329 | 0.163621 | 0.061388 | 0.067525 | 0.195478 | 0.159319 | 0.300738 | 0.286685 | 0.947202 | 1.03342 | 1.066563 | 1.101312 |
| Slc24a3 | 0.04986 | 0.067894 | 0.121104 | 0.115321 | 0.134429 | 0.185725 | 0.220166 | 0.347335 | 0.537502 | 0.536703 | 1.721399 | 1.842894 |
| St8sia2 | 0.135021 | 0.181886 | 0.104669 | 0.098197 | 0.217469 | 0.190198 | 0.31599 | 0.226664 | 1.052119 | 1.221206 | 1.546493 | 1.55941 |
| Nuak1 | 1.654829 | 1.834578 | 3.342114 | 3.397065 | 4.84517 | 4.709272 | 7.562131 | 7.454211 | 16.65878 | 17.27652 | 51.54018 | 53.72529 |

| | | | | | | | | | | | | |
|---|---|---|---|---|---|---|---|---|---|---|---|---|
| Atf3 | 0.48721 | 0.329723 | 1.241898 | 1.461203 | 2.050252 | 2.052241 | 2.835211 | 3.307154 | 4.46699 | 3.534975 | 21.40478 | 22.62492 |
| Hoxd1 | 0.043448 | 0.067743 | 0.208337 | 0.17983 | 0.649407 | 0.52772 | 1.092889 | 1.183004 | 3.334995 | 2.778586 | 1.554194 | 1.969051 |
| Zfp503 | 0.115016 | 0.10053 | 0.675905 | 0.83583 | 1.610851 | 1.746264 | 3.029347 | 2.336124 | 12.81454 | 11.39044 | 14.0848 | 12.85051 |
| Slc44a5 | 0.011626 | 0.040811 | 0.075169 | 0.115398 | 0.202015 | 0.199166 | 0.349551 | 0.261928 | 1.495856 | 1.270165 | 1.654749 | 1.768989 |
| Phlda1 | 1.066664 | 0.855763 | 1.06845 | 0.964551 | 1.137798 | 1.141587 | 1.684184 | 1.803505 | 7.538311 | 6.52757 | 15.5593 | 14.77499 |
| Zfp385b | 0.198967 | 0.078971 | 0.142355 | 0.154317 | 0.198454 | 0.34562 | 0.416124 | 0.496039 | 0.986626 | 0.912248 | 3.152116 | 2.351851 |
| Dpysl3 | 6.197385 | 6.76827 | 3.475828 | 3.699407 | 4.930079 | 4.426933 | 8.455856 | 8.580335 | 36.78891 | 39.40967 | 56.2603 | 56.29286 |
| Tbx2 | 0.23369 | 0.080745 | 0.504938 | 0.448902 | 1.338998 | 1.455817 | 2.967574 | 2.727236 | 4.749221 | 4.486574 | 6.301629 | 6.803319 |
| Dio3 | 0.158723 | 0.176205 | 0.284544 | 0.283656 | 0.237846 | 0.205213 | 0.437661 | 0.484832 | 0.477871 | 0.297852 | 3.669488 | 3.095354 |
| Serpine1 | 1.813104 | 1.88789 | 2.426762 | 2.400378 | 2.177273 | 2.162537 | 1.978876 | 1.937297 | 1.244797 | 1.289414 | 20.79175 | 19.30736 |
| Cyr61 | 5.626186 | 6.484092 | 9.584099 | 8.873197 | 8.359582 | 8.916382 | 12.14271 | 12.22519 | 36.24822 | 34.4874 | 120.6874 | 132.8223 |
| Pla2g10 | 12.72964 | 12.19716 | 5.422195 | 5.510407 | 3.040779 | 2.541629 | 1.73729 | 1.086869 | 0.433154 | 0.514024 | 0.276412 | 0.36859 |
| Cp | 0.185479 | 0.222717 | 0.127494 | 0.140839 | 0.123844 | 0.237241 | 0.209013 | 0.18484 | 1.221158 | 1.330766 | 1.994175 | 2.098044 |
| Gtsf1l | 23.76914 | 26.29669 | 20.28487 | 20.74656 | 9.507955 | 8.847352 | 4.042501 | 4.561274 | 2.539891 | 2.804601 | 1.450613 | 1.395093 |
| Nrcam | 0.37273 | 0.423095 | 0.355241 | 0.426638 | 0.46269 | 0.460806 | 0.844813 | 0.730364 | 6.389604 | 6.335884 | 4.937795 | 6.058127 |
| Dach1 | 0.154797 | 0.202173 | 0.243814 | 0.312634 | 0.726144 | 0.799568 | 1.813927 | 1.571516 | 4.734237 | 4.61731 | 4.006456 | 3.705499 |
| Igf2 | 1.842893 | 2.161616 | 4.263764 | 5.089949 | 9.782887 | 9.691986 | 22.13692 | 18.70912 | 70.73484 | 72.21152 | 92.82604 | 74.40239 |
| Rln3 | 5.438211 | 4.217343 | 6.155619 | 5.291222 | 4.035142 | 3.764214 | 2.037103 | 2.291608 | 0.431982 | 0.36266 | 0.370657 | 0.783567 |
| Rasgrp2 | 4.773495 | 5.24787 | 3.067178 | 2.769674 | 1.910273 | 1.764314 | 1.073778 | 1.164367 | 0.33028 | 0.322032 | 0.25653 | 0.189876 |
| Irx5 | 0.628896 | 0.546765 | 0.731766 | 0.96947 | 1.540862 | 1.426124 | 3.553945 | 3.101068 | 13.69567 | 12.26868 | 13.96527 | 14.01373 |
| Id2 | 16.53979 | 13.37839 | 4.95267 | 4.320697 | 7.346793 | 6.110168 | 10.48893 | 12.10426 | 63.96613 | 61.57616 | 37.68747 | 37.89085 |
| Sox9 | 0.048676 | 0.046427 | 0.084019 | 0.146102 | 0.559253 | 0.44064 | 1.307023 | 1.212764 | 1.804089 | 2.078544 | 2.388875 | 1.918241 |
| Dcn | 0.438087 | 0.135719 | 0.268974 | 0.284067 | 0.202873 | 0.104322 | 0.307638 | 0.1491 | 0.61841 | 0.822909 | 1.789979 | 1.478168 |
| Efna5 | 0.370192 | 0.53366 | 0.471006 | 0.490986 | 0.659046 | 0.729169 | 1.196357 | 1.174002 | 6.009328 | 5.871835 | 6.355242 | 6.954969 |
| Hoxd8 | 0.106917 | 0.040024 | 0.139991 | 0.188721 | 0.42072 | 0.486205 | 0.49455 | 0.533794 | 1.950133 | 2.003355 | 2.288624 | 2.661884 |
| Chrna3 | 0.161302 | 0.169079 | 0.505731 | 0.677236 | 1.25725 | 1.237384 | 4.378669 | 4.216988 | 10.63286 | 9.147449 | 8.829087 | 9.507663 |
| Irs1 | 0.154448 | 0.240552 | 0.211185 | 0.278399 | 0.398732 | 0.470739 | 0.752658 | 0.656057 | 1.937118 | 2.140697 | 3.880935 | 3.8835 |
| Gbp5 | 0.500705 | 0.490993 | 0.345011 | 0.222145 | 0.319916 | 0.495651 | 0.835861 | 0.791202 | 0.622489 | 0.752593 | 3.506464 | 3.059915 |
| Plac1 | 0.319209 | 0.261031 | 0.26798 | 0.223043 | 0.619298 | 0.449473 | 0.764708 | 0.936561 | 3.089638 | 2.79273 | 3.312806 | 3.071279 |
| H19 | 8.441229 | 8.374322 | 18.17776 | 16.91842 | 38.76017 | 36.02953 | 71.89036 | 79.38293 | 203.9285 | 211.2626 | 278.6131 | 228.9163 |
| Cldn2 | 0.178731 | 0.112825 | 0.248574 | 0.320987 | 0.807913 | 0.70472 | 3.162222 | 2.494241 | 2.987116 | 2.589834 | 4.087921 | 4.340166 |
| Anxa1 | 1.492328 | 1.156968 | 3.351221 | 2.456763 | 1.397031 | 2.081304 | 3.119382 | 3.107674 | 5.6348 | 5.461358 | 28.94655 | 28.13892 |
| Efemp1 | 0.396059 | 0.259221 | 0.454121 | 0.865791 | 1.415218 | 1.003165 | 2.650833 | 2.323798 | 11.58335 | 11.61061 | 6.370684 | 6.735902 |
| Rrad | 0.269357 | 0.161346 | 0.188106 | 0.325571 | 0.55039 | 0.696157 | 0.68676 | 0.535624 | 0.864245 | 0.577709 | 5.132679 | 4.320897 |
| Nes | 4.352064 | 5.390426 | 6.500411 | 6.27574 | 9.752323 | 8.545517 | 13.81996 | 12.92173 | 22.55673 | 22.26194 | 83.63139 | 82.59738 |
| Rfx4 | 0.49064 | 0.425581 | 0.577191 | 0.732787 | 1.281742 | 1.328792 | 2.674393 | 2.599307 | 10.41195 | 9.586961 | 6.011386 | 6.20942 |
| Plxdc1 | 4.66241 | 5.628062 | 1.859924 | 1.91009 | 1.453856 | 1.437853 | 0.657667 | 0.650706 | 0.286578 | 0.424521 | 0.1673 | 0.146336 |
| Sfrp5 | 0.562156 | 0.602349 | 4.656616 | 4.163572 | 18.01983 | 17.38628 | 53.01689 | 54.11916 | 43.41079 | 39.95056 | 48.87794 | 51.39407 |
| Gpc3 | 17.8251 | 18.43566 | 7.894782 | 8.839802 | 12.88148 | 11.8504 | 18.27467 | 22.33378 | 117.3921 | 113.7821 | 111.9172 | 111.431 |
| Asb4 | 0.306432 | 0.313956 | 0.224429 | 0.260168 | 0.380059 | 0.323311 | 0.751909 | 0.841497 | 3.583058 | 3.34855 | 1.339099 | 1.497366 |
| Fbn2 | 0.456375 | 0.503848 | 1.402111 | 1.617711 | 3.671473 | 3.480572 | 7.35011 | 5.715281 | 15.54378 | 14.94043 | 19.74108 | 20.35258 |
| Foxe3 | 0.046174 | 0.1 | 0.0295 | 0.1 | 0.069583 | 0.1 | 0.131099 | 0.16217 | 1.064197 | 1.239182 | 1.536525 | 1.25606 |
| Crabp1 | 20.49683 | 19.69621 | 14.18648 | 12.83989 | 19.23214 | 19.77684 | 25.76371 | 24.77761 | 127.8178 | 108.927 | 184.1095 | 161.2219 |
| Crlf1 | 0.417178 | 0.549681 | 1.485782 | 1.407514 | 1.041279 | 1.047638 | 0.686775 | 0.456127 | 0.121081 | 0.112142 | 0.301427 | 0.346399 |
| Ncald | 0.189235 | 0.118759 | 0.182141 | 0.190656 | 0.248377 | 0.310167 | 0.425736 | 0.552411 | 1.520012 | 1.549982 | 2.268529 | 2.391773 |
| Ets1 | 0.746128 | 0.679215 | 1.021242 | 0.937093 | 1.530203 | 1.426831 | 2.141238 | 2.020451 | 2.383622 | 2.308091 | 11.62523 | 11.74368 |
| Camk2n1 | 4.089383 | 4.3868 | 27.59303 | 26.38021 | 35.37124 | 36.69456 | 27.25399 | 25.96462 | 6.342974 | 6.117202 | 3.040975 | 2.928358 |
| Arhgap36 | 0.132816 | 0.202937 | 0.152497 | 0.123612 | 0.245762 | 0.272773 | 0.758978 | 0.556286 | 1.471635 | 1.43979 | 1.359152 | 1.537176 |
| Gm14005 | 0.166474 | 0.100245 | 0.320344 | 0.158506 | 0.142789 | 0.211091 | 0.321843 | 0.332784 | 0.486763 | 0.290881 | 2.717339 | 1.947973 |
| Mid1 | 0.09626 | 0.096151 | 0.249402 | 0.164078 | 0.298775 | 0.222847 | 0.396226 | 0.353803 | 1.838816 | 1.589676 | 1.897832 | 2.012867 |
| Pdzrn3 | 0.790934 | 0.778599 | 1.21633 | 1.489728 | 3.774558 | 3.612364 | 7.153135 | 7.491426 | 19.26968 | 18.19702 | 11.66925 | 11.24116 |
| Arl4c | 5.862876 | 6.013942 | 3.401664 | 3.688599 | 4.404985 | 4.440128 | 5.703858 | 6.789103 | 11.46475 | 11.33768 | 47.12173 | 44.87079 |
| Ildr2 | 0.276254 | 0.298056 | 0.555655 | 0.658525 | 0.937016 | 0.868287 | 1.670089 | 1.300759 | 7.60226 | 8.008609 | 6.481056 | 6.882132 |
| Dusp8 | 0.184836 | 0.132052 | 0.37136 | 0.325702 | 0.444795 | 0.337017 | 0.592271 | 0.497548 | 1.117178 | 1.059187 | 3.80513 | 3.959089 |
| Spta1 | 3.824248 | 4.100332 | 2.190512 | 2.205251 | 0.967929 | 1.132369 | 0.720339 | 0.691354 | 0.235993 | 0.18275 | 0.274769 | 0.254343 |
| Nhsl2 | 0.244723 | 0.348045 | 0.468317 | 0.524982 | 0.643083 | 0.663421 | 0.972868 | 0.729777 | 2.289802 | 2.108592 | 5.992448 | 6.300243 |
| Synpo | 0.246212 | 0.311249 | 0.726731 | 0.770665 | 0.995816 | 1.0882 | 2.662149 | 2.12578 | 3.706881 | 3.458062 | 9.849714 | 9.233675 |
| Foxr2 | 1.124754 | 1.323155 | 1.267571 | 1.421023 | 0.740906 | 0.749675 | 0.529643 | 0.584014 | 0.365834 | 0.32854 | 0.12169 | 0.118824 |
| Bach2 | 0.342796 | 0.45948 | 0.575214 | 0.640956 | 0.905162 | 0.888248 | 1.411284 | 1.102817 | 2.628305 | 2.59211 | 7.702146 | 7.631688 |
| Fhl2 | 0.335635 | 0.336062 | 0.769115 | 0.614198 | 0.979319 | 0.937494 | 1.025304 | 0.659678 | 1.765055 | 2.060869 | 7.089984 | 7.301704 |
| Plaur | 5.068424 | 4.575249 | 3.012407 | 2.276981 | 2.668746 | 2.308605 | 2.55268 | 1.880055 | 3.466738 | 2.871177 | 24.12168 | 22.2873 |

| | | | | | | | | | | | | |
|---|---|---|---|---|---|---|---|---|---|---|---|---|
| Rgs16 | 0.732275 | 0.86109 | 0.536729 | 0.829003 | 0.757237 | 0.563423 | 1.214071 | 1.182197 | 1.726012 | 1.667877 | 6.670504 | 6.659135 |
| Nkx2-5 | 0.09279 | 0.123464 | 0.331249 | 0.204503 | 0.581312 | 0.670496 | 1.014294 | 1.106371 | 1.35645 | 1.189372 | 3.16215 | 2.408852 |
| Boc | 0.851377 | 0.97181 | 2.032335 | 2.205087 | 3.292937 | 3.246088 | 6.410358 | 5.463616 | 24.5613 | 25.97253 | 17.38814 | 17.17707 |
| Cacnb2 | 0.163491 | 0.172945 | 0.161546 | 0.149302 | 0.214349 | 0.28606 | 0.374398 | 0.249483 | 1.160682 | 1.066218 | 1.418452 | 1.757512 |
| Cd24a | 35.2219 | 28.7545 | 15.52901 | 14.71417 | 38.13199 | 36.61774 | 71.53035 | 75.25102 | 170.2845 | 172.7546 | 109.145 | 117.4161 |
| Tgfb1i1 | 0.419949 | 0.390986 | 1.053042 | 1.245563 | 2.237719 | 1.805357 | 2.933765 | 2.447125 | 3.023594 | 2.651087 | 14.59266 | 14.59211 |
| Gm715 | 0.436579 | 0.252613 | 0.394987 | 0.63291 | 1.309184 | 1.036037 | 2.195327 | 2.030343 | 7.069889 | 5.548452 | 7.44099 | 7.397857 |
| Fbxo32 | 0.372117 | 0.349716 | 0.389656 | 0.371467 | 0.691157 | 0.734677 | 2.10395 | 1.709541 | 3.339435 | 3.029911 | 4.12603 | 4.32435 |
| Foxi3 | 3.013468 | 3.138618 | 1.742838 | 2.295592 | 1.291964 | 1.228275 | 0.629696 | 0.403394 | 0.294137 | 0.265278 | 0.213261 | 0.198321 |
| Sdk2 | 0.169828 | 0.166287 | 0.306299 | 0.292273 | 0.585216 | 0.572417 | 1.069374 | 0.86368 | 3.121957 | 3.370303 | 2.29351 | 2.065982 |
| Chrna9 | 2.963331 | 2.764411 | 1.361199 | 0.877871 | 0.633693 | 0.507264 | 0.220212 | 0.076289 | 0.128849 | 0.148442 | 0.161748 | 0.223259 |
| Sox6 | 0.046565 | 0.064519 | 0.128761 | 0.113895 | 0.210441 | 0.20954 | 0.512067 | 0.307624 | 1.081064 | 1.137711 | 1.16073 | 1.309955 |
| Flt1 | 0.218937 | 0.174061 | 0.324276 | 0.261104 | 0.242521 | 0.305727 | 0.346203 | 0.360255 | 0.521481 | 0.44302 | 3.174195 | 2.999921 |
| Irs2 | 2.717258 | 3.661157 | 4.60769 | 4.706412 | 6.759321 | 6.447046 | 10.86641 | 9.107869 | 38.61925 | 37.93835 | 53.98157 | 53.86095 |
| Pla2g7 | 0.355561 | 0.170135 | 0.279516 | 0.144981 | 0.114882 | 0.150351 | 0.179914 | 0.156447 | 0.338293 | 0.414027 | 1.848769 | 1.658469 |
| Lmod1 | 0.337156 | 0.230063 | 0.321223 | 0.331309 | 0.571129 | 0.526611 | 0.859355 | 0.878461 | 1.00469 | 0.877164 | 3.294071 | 3.768558 |
| Alppl2 | 0.019269 | 0.060295 | 0.166071 | 0.095098 | 0.221507 | 0.168281 | 0.349024 | 0.403254 | 0.959832 | 1.079334 | 1.338903 | 0.92964 |
| Sftpa1 | 2.721125 | 0.746995 | 0.1 | 0.394479 | 0.290827 | 0.21066 | 0.314442 | 1.133087 | 0.00708 | 0.162554 | 0.1 | 0.1 |
| Hoxc5 | 0.013791 | 0.029295 | 1.100351 | 1.089291 | 4.328332 | 4.159795 | 10.05521 | 8.610204 | 13.81588 | 12.33187 | 8.16885 | 8.447574 |
| Zfp608 | 1.732077 | 2.122989 | 0.943665 | 1.246717 | 2.371524 | 2.223224 | 5.466608 | 4.501142 | 14.37126 | 14.06623 | 12.28939 | 11.46476 |
| Trabd2b | 0.070331 | 0.108812 | 0.141408 | 0.197674 | 0.380259 | 0.354334 | 0.639087 | 0.524129 | 1.026132 | 1.037119 | 2.170168 | 2.223406 |
| Olfm2 | 1.580634 | 2.266901 | 2.029739 | 2.029945 | 2.922201 | 2.655063 | 4.181498 | 3.890275 | 22.62104 | 20.98463 | 23.00904 | 22.69088 |
| Lefty1 | 64.74421 | 56.678 | 140.3824 | 129.2476 | 126.8189 | 135.1446 | 86.02575 | 86.44374 | 21.90027 | 19.53629 | 11.33477 | 12.16123 |
| Stk32a | 0.192721 | 0.218992 | 0.853708 | 0.858034 | 2.055301 | 1.811204 | 5.821073 | 4.717829 | 7.676267 | 7.085904 | 8.456019 | 9.450855 |
| Eef1a2 | 0.231982 | 0.315229 | 1.811315 | 1.424427 | 2.290799 | 1.989245 | 1.925448 | 1.506763 | 0.45147 | 0.361445 | 0.328627 | 0.181831 |
| Scube1 | 0.299148 | 0.262012 | 0.394195 | 0.345423 | 0.558971 | 0.602801 | 0.866455 | 1.07224 | 3.632799 | 3.766376 | 3.21913 | 2.91414 |
| Prox1 | 0.160838 | 0.206241 | 0.327809 | 0.342034 | 0.771143 | 0.702843 | 1.513612 | 1.391857 | 3.461457 | 3.696127 | 3.243397 | 3.172609 |
| Jph2 | 0.248825 | 0.210118 | 0.387213 | 0.284267 | 0.483394 | 0.593765 | 0.679638 | 0.756818 | 0.760226 | 0.661845 | 3.325163 | 3.062322 |
| Ankrd34a | 1.263671 | 1.476797 | 1.619823 | 1.697838 | 0.958452 | 1.240749 | 0.805645 | 0.707633 | 0.251949 | 0.244817 | 0.164872 | 0.1584 |
| Prdm12 | 0.065142 | 0.127244 | 0.453718 | 0.506209 | 0.235706 | 0.187904 | 0.179495 | 0.265536 | 2.239889 | 2.003901 | 2.265684 | 2.005478 |
| Mmp25 | 3.121857 | 3.665699 | 3.474248 | 4.009374 | 5.217919 | 5.30712 | 7.720643 | 7.169706 | 1.837459 | 1.84347 | 0.634948 | 0.673964 |
| Miat | 0.314778 | 0.383644 | 0.371483 | 0.3684 | 0.246211 | 0.284271 | 0.26995 | 0.281315 | 1.177544 | 1.212347 | 2.663366 | 2.992001 |
| Accsl | 2.99981 | 3.225033 | 8.073381 | 8.168215 | 3.849251 | 4.389724 | 3.08359 | 2.992413 | 1.742128 | 1.428754 | 1.015557 | 0.770651 |
| Antxr2 | 0.208697 | 0.14845 | 0.275318 | 0.308036 | 0.701558 | 0.719988 | 1.358252 | 1.364474 | 3.59014 | 3.262764 | 2.906477 | 3.049909 |
| Trim63 | 2.586513 | 2.567133 | 2.222972 | 2.141521 | 1.218097 | 1.084312 | 0.922914 | 0.82005 | 0.417934 | 0.378374 | 0.089704 | 0.20356 |
| Arhgef38 | 1.71589 | 2.009164 | 2.541366 | 2.101228 | 1.138462 | 1.167399 | 0.892246 | 0.902806 | 0.372494 | 0.441548 | 0.19437 | 0.199884 |
| Fam19a4 | 2.705903 | 2.478601 | 1.990781 | 2.160622 | 1.494761 | 1.29594 | 1.522885 | 1.503527 | 0.411296 | 0.316152 | 0.312625 | 0.205919 |
| Dpysl4 | 1.047623 | 1.028462 | 0.773125 | 0.660515 | 1.47283 | 1.282195 | 2.333425 | 1.999335 | 6.048522 | 5.89859 | 6.870722 | 6.927123 |
| Hsd17b14 | 55.41485 | 58.30446 | 43.22238 | 41.16133 | 21.92392 | 20.9791 | 12.70608 | 12.05154 | 5.858233 | 4.395056 | 3.68065 | 3.932702 |
| Gm6792 | 4.313876 | 4.263506 | 2.106928 | 2.208563 | 1.093872 | 0.936846 | 0.997601 | 0.930486 | 0.28932 | 0.21129 | 0.437709 | 0.621434 |
| Pkdcc | 7.993058 | 6.97465 | 4.039791 | 3.774537 | 6.710186 | 6.297976 | 6.765564 | 7.434354 | 12.54289 | 12.56627 | 39.87562 | 39.36138 |
| Add2 | 0.045369 | 0.054021 | 0.176995 | 0.141046 | 0.316614 | 0.386528 | 0.405813 | 0.405711 | 0.548856 | 0.657149 | 1.4967 | 1.468182 |
| Trim9 | 0.024856 | 0.030513 | 0.080979 | 0.110624 | 0.187395 | 0.240372 | 0.495346 | 0.435478 | 1.012014 | 0.936876 | 1.009073 | 1.148232 |
| Stk39 | 2.378301 | 2.176036 | 1.12057 | 1.222972 | 2.831921 | 2.853547 | 4.375879 | 5.076384 | 12.19954 | 12.01109 | 13.05794 | 12.5785 |
| Sox3 | 2.979171 | 3.486916 | 6.786512 | 6.423044 | 8.671842 | 8.140588 | 14.70507 | 13.70834 | 58.4307 | 44.05547 | 73.06397 | 65.55717 |
| Zfp819 | 5.165932 | 4.684321 | 1.919378 | 1.361289 | 0.929252 | 0.753848 | 0.986422 | 0.873053 | 1.870048 | 2.002351 | 6.79552 | 7.692345 |
| Pgm5 | 0.087118 | 0.07263 | 0.08423 | 0.126911 | 0.317072 | 0.339482 | 0.661406 | 0.578785 | 1.250491 | 1.292539 | 1.178645 | 1.176835 |
| Pcdh8 | 0.181332 | 0.2154 | 0.431826 | 0.264599 | 0.42724 | 0.499836 | 0.509748 | 0.491736 | 1.56817 | 1.411272 | 2.880328 | 2.662383 |
| Galr2 | 0.685269 | 0.54579 | 1.543646 | 1.05193 | 1.059072 | 1.150921 | 0.467676 | 0.652274 | 0.193767 | 0.114445 | 0.067851 | 0.245808 |
| Lrrc2 | 10.52011 | 9.901512 | 6.330762 | 5.949931 | 2.595231 | 2.541668 | 1.536393 | 1.717307 | 0.954441 | 0.906308 | 0.768889 | 0.59353 |
| B630019K0 | 0.117639 | 0.114421 | 0.234352 | 0.241477 | 0.319928 | 0.366628 | 0.449787 | 0.417322 | 1.993675 | 1.661462 | 2.537653 | 2.415185 |
| Col1a2 | 1.037257 | 1.11298 | 2.598218 | 2.418207 | 5.604322 | 5.065689 | 15.04671 | 12.71819 | 24.77221 | 24.12875 | 19.16911 | 18.83731 |
| Cxcl12 | 1.159599 | 0.995088 | 0.584467 | 0.773796 | 0.372184 | 0.336109 | 0.327803 | 0.174908 | 0.659827 | 0.611571 | 1.999294 | 1.741526 |
| Apln | 0.65277 | 0.542811 | 0.153648 | 0.21685 | 0.447382 | 0.411167 | 0.417445 | 0.476261 | 0.674563 | 0.805096 | 1.986998 | 2.153966 |
| Fam65b | 0.147529 | 0.241838 | 0.099959 | 0.17458 | 0.348175 | 0.392351 | 0.619522 | 0.702003 | 1.644151 | 1.726341 | 0.825652 | 1.017586 |
| Spsb4 | 2.190899 | 2.51527 | 3.514983 | 3.36781 | 7.193214 | 6.378094 | 14.25578 | 15.34642 | 29.16811 | 28.13454 | 33.5493 | 33.24277 |
| Grifin | 0.033107 | 0.067587 | 0.952716 | 0.645891 | 2.103697 | 2.430588 | 6.193195 | 6.345789 | 5.551994 | 4.922136 | 6.313037 | 3.880665 |
| Fat3 | 0.108274 | 0.167572 | 0.246912 | 0.249326 | 0.604204 | 0.570852 | 1.176096 | 0.901486 | 2.216066 | 2.437678 | 1.830739 | 1.985842 |
| Kcnk4 | 0.270062 | 0.186564 | 2.623591 | 2.296064 | 1.784927 | 1.627149 | 0.975495 | 1.912942 | 0.417147 | 0.565218 | 0.307683 | 0.235568 |
| Bambi-ps1 | 0.717557 | 0.712311 | 1.569773 | 1.140237 | 1.258433 | 2.020572 | 2.195717 | 1.82085 | 3.932874 | 3.103992 | 11.42797 | 11.08374 |
| Foxd1 | 0.074075 | 0.08489 | 0.187865 | 0.156836 | 0.16355 | 0.3354 | 0.768464 | 0.590122 | 1.214152 | 1.237547 | 1.731305 | 1.520666 |

| | | | | | | | | | | | | |
|---|---|---|---|---|---|---|---|---|---|---|---|---|
| Gas1 | 2.570117 | 2.003285 | 8.337148 | 8.973634 | 18.58461 | 19.61692 | 30.54747 | 25.97396 | 79.03687 | 69.02095 | 91.34613 | 86.9424 |
| Lsp1 | 0.3552 | 0.166005 | 0.352704 | 0.513938 | 0.831196 | 1.039201 | 1.152945 | 1.37172 | 1.822428 | 1.77182 | 6.169339 | 4.949925 |
| Inhbb | 12.4687 | 13.24515 | 14.70612 | 13.26818 | 6.080539 | 5.636383 | 3.980667 | 4.141487 | 1.295313 | 1.382236 | 1.245323 | 1.379968 |
| Tshz1 | 0.317623 | 0.219352 | 5.419413 | 5.689643 | 17.35682 | 17.90186 | 40.92607 | 37.94279 | 51.65748 | 50.45203 | 51.75516 | 54.10344 |
| Crabp2 | 6.340376 | 5.277105 | 39.58597 | 40.6312 | 118.2178 | 108.3969 | 268.3469 | 258.0962 | 421.2728 | 385.4221 | 336.9503 | 329.8855 |
| Olfr986 | 0.794889 | 1.307217 | 1.059423 | 0.959769 | 0.495349 | 0.423902 | 0.354163 | 0.322903 | 0.21384 | 0.169729 | 0.101739 | 0.101289 |
| Ak7 | 2.991729 | 3.084732 | 9.390543 | 8.894499 | 3.777573 | 3.791273 | 2.695461 | 2.651711 | 1.347037 | 1.23947 | 0.967801 | 0.940239 |
| Apba2 | 0.675336 | 0.838604 | 1.030651 | 0.929344 | 1.454442 | 1.276938 | 2.056759 | 1.881474 | 9.127962 | 7.613637 | 8.549567 | 8.76807 |
| Adamts1 | 0.487364 | 0.580636 | 0.710478 | 0.75502 | 1.282143 | 1.130163 | 1.84507 | 1.928805 | 3.610974 | 3.317136 | 5.849827 | 7.115214 |
| Ndnf | 1.358416 | 1.409659 | 1.179198 | 1.296568 | 2.028911 | 2.128086 | 3.56949 | 3.297132 | 13.00146 | 12.21058 | 9.898487 | 10.14732 |
| Pygl | 0.97562 | 0.777012 | 0.383972 | 0.472289 | 1.002407 | 1.070335 | 1.650192 | 1.723055 | 3.791915 | 4.417768 | 3.069083 | 2.698033 |
| 1700123I0 | 1.574772 | 1.606271 | 2.190121 | 2.397717 | 1.521308 | 1.296458 | 1.188898 | 1.3202 | 0.414713 | 0.256336 | 0.350709 | 0.401563 |
| Pcdhb22 | 0.170683 | 0.091115 | 0.143856 | 0.14152 | 0.196833 | 0.287727 | 0.338324 | 0.259778 | 0.769046 | 0.796243 | 1.567515 | 1.321414 |
| Crispld2 | 0.294359 | 0.26464 | 0.625873 | 0.487983 | 0.732515 | 0.658864 | 0.773236 | 0.941376 | 1.306585 | 1.591235 | 4.915116 | 4.518973 |
| Zfp521 | 0.668533 | 0.728705 | 0.420952 | 0.612199 | 0.926825 | 0.763066 | 1.586177 | 1.485547 | 5.418242 | 5.669175 | 2.802456 | 3.283367 |
| Gpr50 | 0.02627 | 0.048452 | 0.177097 | 0.128371 | 0.268869 | 0.342225 | 0.593528 | 0.442863 | 0.911434 | 0.638141 | 1.029989 | 1.185559 |
| Ly6h | 0.101138 | 0.099109 | 0.272421 | 0.257399 | 0.352632 | 0.39801 | 0.447145 | 0.418307 | 1.090991 | 0.963205 | 2.462061 | 2.373467 |
| Bcl11a | 2.319437 | 2.021471 | 0.475826 | 0.507824 | 0.262443 | 0.310441 | 0.161963 | 0.14086 | 0.072387 | 0.073474 | 0.067034 | 0.055734 |
| Dtx4 | 1.876021 | 2.107755 | 2.373238 | 2.822578 | 3.610975 | 3.535155 | 5.11627 | 4.853366 | 16.77021 | 16.24645 | 24.47262 | 25.61663 |
| Dpys | 3.157749 | 2.2259 | 1.130489 | 1.14319 | 0.652774 | 0.492122 | 0.325511 | 0.626667 | 0.176874 | 0.149402 | 0.153414 | 0.127109 |
| Hs6st2 | 1.278851 | 1.423827 | 1.766975 | 1.665914 | 3.248077 | 2.88568 | 4.932378 | 5.041094 | 15.19505 | 12.82489 | 12.89627 | 14.92456 |
| Cass4 | 1.237874 | 1.453381 | 0.786648 | 0.67314 | 0.562078 | 0.49244 | 0.450839 | 0.376846 | 0.12649 | 0.075973 | 0.118394 | 0.108949 |
| Irf2 | 0.21175 | 0.179215 | 0.225672 | 0.342589 | 0.51866 | 0.518719 | 0.791128 | 0.792439 | 1.999596 | 1.852979 | 2.743521 | 3.029633 |
| Cpne5 | 1.626648 | 1.724911 | 3.212881 | 3.09394 | 2.24829 | 1.869659 | 1.413749 | 1.421537 | 0.590322 | 0.694595 | 0.476397 | 0.350511 |
| Hist1h2bl | 4.548053 | 5.067511 | 1.767823 | 3.822094 | 3.18225 | 6.780219 | 3.4701 | 2.952258 | 2.076709 | 2.216717 | 0.955161 | 0.770662 |
| Slc2a7 | 0.269198 | 0.3638 | 1.104885 | 1.749243 | 0.765899 | 0.903302 | 0.641221 | 0.831609 | 0.24912 | 0.44847 | 0.243376 | 0.200377 |
| Tcstv3 | 0.832383 | 0.667984 | 1.287224 | 0.945011 | 2.411065 | 2.692125 | 7.470015 | 6.984115 | 6.487325 | 6.678062 | 8.525525 | 8.236927 |
| Cryge | 0.232879 | 0.092741 | 0.781525 | 0.726103 | 1.545906 | 1.421808 | 3.768834 | 2.784856 | 4.040573 | 2.513701 | 6.305985 | 6.310199 |
| Car3 | 4.051834 | 3.852139 | 4.589587 | 4.537772 | 5.895449 | 6.878258 | 9.994585 | 10.41836 | 13.38853 | 9.598144 | 35.76506 | 39.31591 |
| 9430069I0 | 2.763511 | 1.756483 | 0.570927 | 0.865629 | 0.179548 | 0.1 | 0.1 | 0.1 | 0.060137 | 0.117971 | 0.135964 | 0.1 |
| Bmp2 | 0.158905 | 0.17236 | 0.263805 | 0.235672 | 0.373282 | 0.287625 | 0.524335 | 0.474603 | 1.155825 | 1.00529 | 2.103976 | 2.037815 |
| Shisa4 | 1.189764 | 1.292032 | 1.844906 | 1.886431 | 3.646972 | 3.914086 | 5.322201 | 4.917987 | 6.692081 | 6.4607 | 16.81061 | 16.31109 |
| Cryab | 2.023927 | 1.548902 | 1.532471 | 1.746037 | 1.243022 | 1.538915 | 1.638945 | 2.135715 | 4.391794 | 4.016351 | 14.82653 | 13.21549 |
| Arg1 | 0.082121 | 0.099494 | 6.595016 | 6.160579 | 11.21628 | 11.57732 | 24.13381 | 21.33786 | 42.28719 | 37.87182 | 54.14963 | 52.75269 |
| Gdf15 | 1.526061 | 1.683221 | 3.44201 | 3.238071 | 2.2827 | 2.042161 | 0.991669 | 0.940375 | 0.574921 | 0.378524 | 0.665789 | 0.615693 |
| Mex3b | 1.373934 | 1.683447 | 2.076312 | 2.10645 | 4.284868 | 4.464423 | 8.248118 | 7.216678 | 18.13102 | 17.98294 | 15.77261 | 14.63516 |
| Gm4340 | 1.104065 | 0.860347 | 2.817616 | 2.644138 | 7.116107 | 7.746493 | 14.86229 | 22.50216 | 15.50807 | 17.23008 | 17.86706 | 15.35446 |
| Fbxl7 | 0.232236 | 0.286947 | 0.568967 | 0.725577 | 1.350584 | 1.357421 | 1.616315 | 1.746402 | 3.508852 | 3.468925 | 5.068046 | 6.171344 |
| Zim1 | 0.291989 | 0.330062 | 0.361599 | 0.395186 | 0.441073 | 0.422956 | 0.657713 | 0.506095 | 2.509887 | 3.360663 | 2.177214 | 2.477979 |
| Gm11544 | 0.421374 | 0.224161 | 0.334771 | 0.790747 | 1.211481 | 1.453044 | 3.937645 | 4.416794 | 6.314859 | 5.640986 | 8.548578 | 6.687501 |
| S1pr1 | 0.359416 | 0.221894 | 0.083032 | 0.234321 | 0.339255 | 0.416216 | 0.622526 | 0.696276 | 1.058932 | 1.167491 | 1.711667 | 1.96005 |
| Plk5 | 2.013566 | 2.085143 | 2.269695 | 2.014914 | 1.800232 | 1.763943 | 1.085345 | 1.242688 | 0.561239 | 0.423277 | 0.312172 | 0.241192 |
| Tnfaip8l2 | 0.178646 | 0.227311 | 0.516715 | 0.79592 | 0.628622 | 0.730381 | 2.844762 | 3.25814 | 5.827149 | 6.099027 | 2.599243 | 2.256579 |
| Foxn4 | 0.521445 | 0.499899 | 4.060034 | 3.858046 | 2.542626 | 2.442584 | 1.44699 | 1.50733 | 0.372254 | 0.475183 | 0.496896 | 0.462614 |
| Mafb | 0.442841 | 0.328265 | 0.557736 | 0.558998 | 1.179807 | 1.031011 | 2.000834 | 2.146159 | 4.960544 | 4.658807 | 2.624833 | 2.729593 |
| Jph4 | 2.163734 | 2.557177 | 1.254145 | 1.585526 | 1.11274 | 1.005262 | 1.022494 | 0.626984 | 0.441863 | 0.289051 | 0.172467 | 0.190777 |
| Scgb3a1 | 0.909794 | 1.169383 | 0.620179 | 1.218754 | 0.337181 | 0.699248 | 0.574867 | 0.705358 | 0.461335 | 0.371278 | 0.388438 | 0.146829 |
| Liph | 12.22594 | 13.41001 | 5.7329 | 4.938667 | 3.147776 | 3.150788 | 2.09651 | 2.063486 | 0.989522 | 1.003449 | 0.818378 | 0.595443 |
| Stk32c | 0.420033 | 0.528186 | 0.96137 | 0.748915 | 1.670481 | 1.511111 | 2.227096 | 2.464453 | 3.075663 | 2.683114 | 5.61694 | 6.172169 |
| Sstr2 | 0.197834 | 0.167344 | 0.147202 | 0.190244 | 0.206978 | 0.186547 | 0.304255 | 0.153089 | 1.025118 | 1.108128 | 1.154746 | 1.261088 |
| Lrtm2 | 0.387929 | 0.334487 | 1.194373 | 1.214878 | 1.509953 | 1.542775 | 1.018771 | 0.878538 | 0.208608 | 0.204114 | 0.188977 | 0.187554 |
| Gngt2 | 1.42226 | 1.061692 | 0.734656 | 0.676063 | 0.533216 | 0.59304 | 0.397226 | 0.531574 | 0.137532 | 0.187366 | 0.052448 | 0.082206 |
| Khdc3 | 4.303545 | 3.073828 | 1.624246 | 1.497215 | 0.826394 | 0.616662 | 0.334774 | 0.334566 | 0.234451 | 0.240577 | 0.390598 | 0.182075 |
| Tuba4a | 73.70396 | 75.1467 | 53.51845 | 47.41534 | 29.40372 | 28.66784 | 20.74399 | 23.51606 | 10.19322 | 9.925451 | 7.092236 | 5.785447 |
| Osr2 | 0.070349 | 0.098325 | 0.08412 | 0.154332 | 0.420984 | 0.325787 | 0.762933 | 0.924829 | 0.737763 | 0.601688 | 1.762265 | 1.264178 |
| Ubash3a | 4.391453 | 5.152141 | 5.450535 | 5.980442 | 4.938869 | 4.838366 | 4.037859 | 3.755733 | 1.900704 | 2.314138 | 1.02774 | 0.731051 |
| C130071C0 | 0.264144 | 0.194789 | 0.553341 | 0.530205 | 1.156072 | 1.33378 | 2.291161 | 2.075729 | 4.694698 | 4.325065 | 2.78389 | 3.464949 |
| Ccdc171 | 1.206636 | 1.386889 | 4.254863 | 4.372836 | 4.028874 | 3.994131 | 2.617583 | 2.775775 | 1.510792 | 1.794309 | 0.459353 | 0.536239 |
| Id4 | 3.979161 | 3.185921 | 0.738297 | 0.787107 | 0.800994 | 0.951287 | 0.817099 | 1.016395 | 2.472148 | 2.124369 | 7.004303 | 6.400391 |
| Gab2 | 0.174953 | 0.182392 | 0.317032 | 0.332305 | 0.668229 | 0.618676 | 1.011322 | 1.015962 | 2.674759 | 2.699461 | 2.683155 | 2.471206 |
| Mest | 47.44881 | 49.51181 | 75.70185 | 78.81197 | 205.7119 | 209.5556 | 455.0403 | 469.148 | 661.8067 | 639.7094 | 489.3141 | 516.3675 |

| | | | | | | | | | | | | |
|---|---|---|---|---|---|---|---|---|---|---|---|---|
| Samd4 | 0.502069 | 0.500302 | 0.733127 | 0.843351 | 1.09072 | 1.046355 | 1.18927 | 1.330289 | 2.179684 | 2.12678 | 6.618835 | 6.763421 |
| Fst | 2.000998 | 2.037449 | 0.914779 | 0.886185 | 1.069218 | 1.108058 | 1.324013 | 1.78484 | 6.460009 | 7.086715 | 3.117696 | 3.29953 |
| Dchs1 | 0.138629 | 0.159467 | 0.332843 | 0.388871 | 0.831558 | 0.759743 | 1.325004 | 1.140022 | 3.089538 | 3.104406 | 2.722308 | 2.682888 |
| Plxna2 | 1.207672 | 1.551944 | 1.648989 | 1.803161 | 2.833862 | 2.746845 | 6.481387 | 5.537757 | 13.54828 | 14.37376 | 9.564867 | 9.675688 |
| Atp1a3 | 7.458549 | 9.751637 | 8.206151 | 7.880616 | 5.851841 | 5.235905 | 4.137727 | 2.996125 | 1.377949 | 1.668728 | 1.125948 | 0.992779 |
| Cd44 | 0.226208 | 0.237725 | 0.2688 | 0.309355 | 0.64606 | 0.544731 | 0.940326 | 1.101969 | 1.230912 | 1.089986 | 2.342936 | 2.447178 |
| Pax6 | 1.920291 | 1.57034 | 3.950403 | 3.955803 | 5.460033 | 5.833994 | 9.662897 | 9.485636 | 33.60089 | 31.28339 | 21.37152 | 22.48527 |
| Cdc42ep1 | 1.159379 | 1.294708 | 2.24379 | 2.02461 | 2.7173 | 3.045616 | 5.73455 | 5.254571 | 7.610027 | 6.763193 | 16.35847 | 15.95799 |
| Zfhx4 | 0.085842 | 0.101967 | 0.221687 | 0.21893 | 0.32656 | 0.314226 | 0.505097 | 0.401988 | 1.586061 | 1.723891 | 1.463725 | 1.504038 |
| Fgf4 | 74.63988 | 70.88407 | 23.4422 | 22.25989 | 13.68706 | 13.71589 | 10.25079 | 10.20867 | 6.057621 | 7.496755 | 2.844562 | 2.828972 |
| Krt4 | 0.152043 | 0.038866 | 1.090695 | 0.303078 | 0.170444 | 0.191184 | 0.377823 | 0.395688 | 1.712935 | 1.502249 | 2.623232 | 1.348185 |
| Tox2 | 0.273855 | 0.328044 | 0.368135 | 0.333548 | 0.53369 | 0.415855 | 0.584247 | 0.4771 | 1.216632 | 0.876433 | 2.883299 | 2.611607 |
| Atoh8 | 0.059733 | 0.071154 | 0.374697 | 0.336247 | 1.078324 | 1.061052 | 1.418158 | 1.154264 | 1.764491 | 1.546249 | 2.621901 | 2.630681 |
| Cd82 | 0.365188 | 0.309676 | 0.350564 | 0.218142 | 0.386463 | 0.22839 | 0.427824 | 0.359725 | 0.636583 | 0.510758 | 2.026779 | 1.703317 |
| Shisa3 | 0.900726 | 0.77719 | 1.314356 | 1.458208 | 5.13482 | 4.710416 | 11.65372 | 11.38325 | 10.6327 | 10.4433 | 7.723736 | 6.882473 |
| Lhfp | 0.669297 | 0.429564 | 1.290183 | 1.12069 | 2.512394 | 2.490993 | 3.060747 | 2.979217 | 6.554609 | 5.646779 | 9.272009 | 8.741525 |
| Gbp2 | 0.888932 | 0.812579 | 0.960678 | 0.64517 | 0.607894 | 0.678941 | 1.42022 | 1.508837 | 1.263232 | 1.125833 | 5.711764 | 5.020814 |
| Aass | 11.04893 | 9.998147 | 3.012002 | 2.974918 | 1.323351 | 1.52059 | 0.766324 | 1.063722 | 0.49833 | 0.479372 | 0.384635 | 0.382417 |
| Bhlhe22 | 0.138837 | 0.101876 | 0.135766 | 0.153341 | 0.367171 | 0.365079 | 0.414993 | 0.350382 | 0.999983 | 0.802054 | 1.264935 | 1.18942 |
| Crct1 | 0.81427 | 1.150907 | 3.835403 | 3.183347 | 2.383082 | 3.1266 | 2.815712 | 4.50394 | 0.657288 | 0.585258 | 1.170594 | 1.010227 |
| Cyp2d22 | 0.640386 | 0.601906 | 0.72996 | 0.95539 | 1.308761 | 1.352358 | 1.040125 | 1.522208 | 0.576201 | 0.623056 | 0.233039 | 0.199226 |
| Glis3 | 0.161091 | 0.244885 | 0.321106 | 0.274128 | 0.484959 | 0.582431 | 0.859226 | 0.679866 | 1.757085 | 1.528486 | 2.137952 | 2.077132 |
| Igsf21 | 3.391977 | 3.660101 | 11.01041 | 10.73821 | 10.79744 | 10.73341 | 8.642551 | 9.378154 | 3.059275 | 3.011939 | 1.614311 | 1.417595 |
| Samd14 | 0.352744 | 0.39376 | 0.837667 | 0.796113 | 1.438582 | 1.377337 | 1.842211 | 1.850388 | 4.316291 | 3.671765 | 6.442018 | 6.025312 |
| Gpc4 | 9.724853 | 12.87969 | 9.081306 | 8.423745 | 8.513703 | 7.959233 | 13.19373 | 11.68583 | 30.93743 | 30.75571 | 57.67978 | 60.03051 |
| Ly6a | 4.473706 | 3.816915 | 2.800989 | 2.052411 | 2.385444 | 2.590713 | 2.952989 | 3.778812 | 12.5158 | 14.24604 | 17.60628 | 15.47176 |
| Insm1 | 3.64322 | 4.090538 | 3.195079 | 2.935305 | 1.299427 | 1.429893 | 1.163478 | 1.23971 | 6.890597 | 5.370651 | 10.04665 | 9.331957 |
| Lrrn1 | 1.374485 | 1.497766 | 1.181484 | 1.000641 | 1.442707 | 1.295657 | 2.096933 | 2.246027 | 7.687965 | 7.515664 | 7.135069 | 7.482424 |
| Cnn2 | 25.39434 | 23.95357 | 20.81093 | 20.70931 | 20.04579 | 19.03855 | 22.51369 | 21.18545 | 38.01619 | 34.42291 | 149.3341 | 142.5152 |
| Nuak2 | 2.648973 | 2.718892 | 3.943873 | 3.706358 | 4.984856 | 4.438805 | 7.290854 | 6.441019 | 17.33229 | 17.85621 | 26.7763 | 27.69665 |
| BC033916 | 24.74249 | 21.61122 | 25.7066 | 24.13529 | 16.42503 | 18.76048 | 12.33569 | 12.48628 | 5.087692 | 4.907911 | 3.799011 | 3.230569 |
| Capn6 | 0.394683 | 0.418674 | 0.470051 | 0.543189 | 1.034117 | 0.867563 | 1.244923 | 1.373281 | 3.090011 | 3.331738 | 4.038201 | 4.056768 |
| Pard3b | 0.045815 | 0.08064 | 0.229602 | 0.203511 | 0.275996 | 0.336583 | 0.442504 | 0.290406 | 1.071586 | 1.023936 | 1.397593 | 1.517765 |
| Gata3 | 0.069091 | 0.098659 | 0.385847 | 0.222424 | 0.325547 | 0.3133 | 0.418433 | 0.291553 | 0.950814 | 0.752829 | 1.721742 | 1.658577 |
| Dtx1 | 3.927286 | 4.374062 | 3.106025 | 2.982344 | 2.146755 | 1.940027 | 1.451095 | 1.451892 | 0.874796 | 0.845474 | 0.627298 | 0.400495 |
| Sfrp2 | 10.34131 | 12.61867 | 7.555252 | 8.08181 | 9.250854 | 8.85483 | 14.48601 | 13.38391 | 57.51849 | 49.29273 | 61.17634 | 59.57611 |
| Jun | 1.7811 | 1.933828 | 3.140666 | 2.531697 | 4.283248 | 4.431622 | 5.89431 | 6.319488 | 10.6431 | 10.43591 | 19.48206 | 18.65588 |
| 4930591A1 | 2.539013 | 3.111992 | 6.602509 | 6.246291 | 2.488657 | 2.597514 | 2.216119 | 1.84416 | 0.533393 | 0.849978 | 1.182444 | 1.171332 |
| Hoxc6 | 0.012837 | 0.105405 | 0.404445 | 0.614645 | 1.159473 | 1.150483 | 2.388972 | 2.40116 | 4.446799 | 4.508763 | 3.762471 | 3.156689 |
| Clca4 | 2.141826 | 2.68123 | 6.533834 | 6.062085 | 3.821913 | 3.775046 | 3.239922 | 3.179321 | 0.963377 | 0.827463 | 1.179402 | 0.974308 |
| Nsg1 | 0.706446 | 0.721303 | 1.05874 | 0.900852 | 1.710837 | 1.485149 | 2.292825 | 2.299553 | 5.389467 | 4.59259 | 6.221279 | 6.574653 |
| Igdcc3 | 5.344453 | 7.262683 | 11.46169 | 11.66753 | 29.50133 | 27.72754 | 59.04585 | 53.39532 | 94.41888 | 84.81647 | 77.22407 | 77.96675 |
| Meis1 | 0.635478 | 0.67302 | 2.552474 | 2.869211 | 5.622294 | 6.109926 | 11.07859 | 9.791641 | 21.52154 | 20.82055 | 15.96422 | 15.76955 |
| Thsd4 | 0.084862 | 0.064636 | 0.3155 | 0.295306 | 0.810119 | 0.686677 | 1.557988 | 1.364026 | 1.862078 | 1.876551 | 2.13044 | 2.134417 |
| Tmtc2 | 0.178834 | 0.245436 | 0.250666 | 0.328243 | 0.418662 | 0.465451 | 0.79052 | 0.616124 | 2.221127 | 2.362898 | 1.821212 | 1.863393 |
| Zic1 | 0.129382 | 0.101746 | 0.491227 | 0.33194 | 0.488631 | 0.462405 | 0.887204 | 0.702254 | 2.440542 | 2.379652 | 1.83994 | 1.800779 |
| Dlk1 | 0.40706 | 0.340446 | 1.324756 | 1.301436 | 1.44978 | 1.572986 | 1.988843 | 1.913765 | 5.450547 | 5.278836 | 11.26146 | 9.315655 |
| Fabp7 | 1.553669 | 1.207413 | 0.201029 | 0.354395 | 0.572824 | 0.58364 | 0.505592 | 0.692355 | 1.882125 | 1.306473 | 2.438575 | 2.534991 |
| Rin1 | 0.748538 | 0.796927 | 0.606861 | 0.587251 | 0.539704 | 0.483155 | 0.523023 | 0.538702 | 0.53358 | 0.411554 | 2.71317 | 2.935362 |
| Emilin2 | 0.387217 | 0.281869 | 0.881466 | 0.768956 | 1.283058 | 1.272245 | 2.038435 | 1.989754 | 5.424703 | 5.059978 | 5.175069 | 5.475565 |
| Nrg1 | 1.189357 | 1.16809 | 0.884803 | 0.853945 | 1.207346 | 0.897838 | 1.507467 | 1.158863 | 2.903818 | 3.319677 | 5.943226 | 6.04826 |
| Krt18 | 5.5762 | 5.681514 | 7.059624 | 6.415528 | 7.684906 | 7.017553 | 9.983867 | 9.942518 | 16.17529 | 15.19575 | 54.40591 | 44.9266 |
| Bcl2 | 0.13736 | 0.081944 | 0.197997 | 0.182441 | 0.322204 | 0.326545 | 0.411864 | 0.306321 | 0.686078 | 0.557652 | 1.067596 | 1.277165 |
| 1700018B0 | 0.72579 | 0.709242 | 1.167966 | 1.43438 | 1.041539 | 0.82275 | 0.71988 | 0.432353 | 0.181786 | 0.206642 | 0.428572 | 0.335637 |
| Il6ra | 0.320501 | 0.309155 | 0.390193 | 0.271237 | 0.391383 | 0.345414 | 0.604335 | 0.897246 | 0.788377 | 1.011604 | 1.853729 | 1.881058 |
| Gm8994 | 2.39521 | 2.324556 | 9.431046 | 8.738318 | 23.61843 | 22.48258 | 48.63029 | 60.5177 | 54.89402 | 53.4649 | 61.49481 | 46.32276 |
| Chrnb4 | 0.364853 | 0.46789 | 0.892071 | 0.872696 | 1.485109 | 1.320605 | 3.830343 | 3.119095 | 6.45409 | 6.04155 | 4.363319 | 4.186486 |
| Hspa1a | 0.258141 | 0.298838 | 0.742087 | 0.737397 | 2.062372 | 1.68174 | 4.532274 | 5.102147 | 3.064301 | 2.995953 | 2.732715 | 2.021259 |
| Tmem125 | 1.145131 | 1.215861 | 4.458404 | 3.621185 | 2.462774 | 2.875841 | 1.407795 | 1.483454 | 0.394105 | 0.524276 | 1.015007 | 0.706849 |
| Fam107b | 3.047227 | 3.117358 | 4.859205 | 4.326694 | 5.768745 | 5.352641 | 7.373392 | 7.038974 | 9.422303 | 9.475818 | 28.36105 | 29.85822 |
| Olfr815 | 0.063435 | 0.022753 | 0.257744 | 0.213319 | 0.57242 | 0.855121 | 1.344105 | 1.379893 | 1.22727 | 1.471679 | 1.139449 | 1.237621 |

| | | | | | | | | | | | | |
|---|---|---|---|---|---|---|---|---|---|---|---|---|
| Foxc1 | 0.015402 | 0.027136 | 0.419494 | 0.323094 | 0.504496 | 0.367221 | 0.721182 | 0.533524 | 1.419577 | 1.292222 | 2.260384 | 2.22427 |
| Akr1c13 | 0.329295 | 0.225109 | 0.379308 | 0.359121 | 0.427859 | 0.605118 | 1.394182 | 1.147765 | 2.354303 | 1.94666 | 2.060874 | 2.470485 |
| Cpe | 3.891011 | 3.631919 | 5.047222 | 5.303252 | 8.99783 | 9.632934 | 13.29677 | 14.77649 | 30.20183 | 28.55956 | 34.14559 | 36.44842 |
| Tmem92 | 0.591663 | 0.573037 | 2.210719 | 2.297607 | 5.523155 | 5.782892 | 13.5158 | 15.73706 | 15.21507 | 15.30581 | 14.39444 | 13.12968 |
| AF067061 | 0.394512 | 0.416054 | 0.826058 | 0.804818 | 2.642286 | 2.50827 | 4.154436 | 5.509079 | 4.419142 | 3.611906 | 5.537428 | 5.005741 |
| Lrrc4 | 0.590309 | 0.744604 | 1.082521 | 1.196788 | 1.125172 | 1.19915 | 1.259302 | 1.439072 | 4.045015 | 4.10653 | 8.28115 | 8.190168 |
| Pde10a | 0.109732 | 0.128916 | 0.53859 | 0.545165 | 1.37521 | 1.429277 | 2.332645 | 2.313899 | 3.88061 | 3.683169 | 3.1214 | 3.727236 |
| Gstm3 | 1.292271 | 1.802105 | 1.004751 | 1.202047 | 0.79912 | 1.06847 | 0.565431 | 0.857979 | 0.163432 | 0.35232 | 0.229452 | 0.176111 |
| Rgs11 | 1.729126 | 1.727158 | 0.840594 | 1.140003 | 1.25915 | 0.790147 | 0.582539 | 0.624201 | 0.351044 | 0.213413 | 0.237472 | 0.167086 |
| Col5a3 | 0.721524 | 1.109636 | 0.503683 | 0.418912 | 0.510063 | 0.335231 | 0.518319 | 0.536149 | 0.263418 | 0.262073 | 2.103638 | 1.786034 |
| Fgf17 | 8.374773 | 8.515209 | 1.661426 | 1.707156 | 0.596891 | 0.460262 | 0.513006 | 0.694533 | 0.244408 | 0.251432 | 0.526216 | 0.352526 |
| B020004J0 | 0.101438 | 0.204322 | 0.465976 | 0.509081 | 1.18321 | 1.203784 | 2.596685 | 3.320543 | 2.961567 | 3.456176 | 3.052278 | 2.746759 |
| Ano9 | 7.946877 | 8.263613 | 5.930781 | 6.056447 | 4.714745 | 4.434637 | 2.615314 | 3.267626 | 1.790612 | 1.8803 | 0.805136 | 0.892289 |
| Prss35 | 1.140436 | 0.948263 | 0.71729 | 0.793832 | 0.970012 | 0.72691 | 1.012425 | 0.884541 | 1.887348 | 1.716888 | 4.297832 | 4.929808 |
| Rbm24 | 0.125384 | 0.163953 | 0.222755 | 0.186251 | 0.357157 | 0.204643 | 0.347054 | 0.30613 | 1.311974 | 1.257173 | 0.873035 | 0.846067 |
| Zscan4f | 3.342168 | 2.90271 | 11.34424 | 10.98944 | 28.57861 | 26.92375 | 60.83024 | 74.15991 | 61.59283 | 60.07281 | 51.05257 | 44.32238 |
| Lix1 | 0.659772 | 0.757181 | 0.37416 | 0.307455 | 0.645662 | 0.445149 | 1.067303 | 0.736999 | 2.054497 | 2.047787 | 1.979831 | 2.06726 |
| Pou4f2 | 3.364092 | 3.710346 | 0.876124 | 0.606416 | 0.446842 | 0.306195 | 0.332146 | 0.160904 | 0.110393 | 0.09032 | 0.142945 | 0.133914 |
| Foxf1 | 0.176013 | 0.130577 | 0.11915 | 0.182975 | 0.458956 | 0.34063 | 0.667929 | 0.64042 | 0.401274 | 0.531901 | 1.282787 | 1.228125 |
| Rnf217 | 0.245265 | 0.281867 | 0.26031 | 0.301146 | 0.552092 | 0.508445 | 0.726297 | 0.748988 | 1.434384 | 1.499314 | 2.016929 | 2.011626 |
| Gm11627 | 8.542037 | 6.412412 | 7.040479 | 6.91716 | 3.914735 | 4.970418 | 3.230252 | 3.208505 | 1.44295 | 1.063879 | 1.28875 | 1.042964 |
| Scara3 | 0.153903 | 0.098967 | 0.460094 | 0.508898 | 1.62597 | 1.564447 | 3.292525 | 2.890481 | 3.724931 | 3.358489 | 3.243154 | 3.31836 |
| Hoxb1 | 0.406747 | 0.397468 | 24.91409 | 27.77148 | 62.39087 | 59.54038 | 87.80173 | 80.07397 | 23.93046 | 24.61603 | 13.85017 | 12.13606 |
| Tdpoz4 | 0.198825 | 0.210537 | 0.861763 | 0.556886 | 1.140269 | 1.163311 | 2.87526 | 2.189657 | 2.573059 | 3.295069 | 4.706248 | 3.67262 |
| Dpysl5 | 3.205286 | 3.785949 | 3.396906 | 3.219224 | 4.756977 | 4.483672 | 5.797915 | 5.796315 | 19.49805 | 19.74618 | 21.09893 | 21.22764 |
| Unc5a | 0.257477 | 0.311707 | 0.404412 | 0.37244 | 0.62983 | 0.487216 | 0.539916 | 0.457526 | 1.314113 | 1.256831 | 2.499921 | 2.450791 |
| Gm10863 | 4.876924 | 4.724225 | 5.242604 | 3.024674 | 1.646616 | 2.247564 | 1.261403 | 0.932163 | 0.702302 | 0.819224 | 0.592054 | 0.460348 |
| Folr4 | 0.156338 | 0.097705 | 1.510651 | 1.839021 | 3.307625 | 2.938126 | 5.108375 | 4.392655 | 1.692345 | 1.86101 | 0.948516 | 0.668553 |
| Slc25a24 | 1.62679 | 1.623949 | 2.553252 | 2.580085 | 4.638207 | 4.271023 | 6.487672 | 6.898245 | 9.277349 | 9.38254 | 17.12013 | 16.94549 |
| S100a13 | 3.328681 | 2.815618 | 1.571003 | 0.987187 | 1.859367 | 1.87542 | 2.559269 | 2.228645 | 5.445732 | 4.472104 | 6.32696 | 6.475615 |
| Mfsd2a | 0.21571 | 0.372824 | 0.741195 | 0.48199 | 0.842056 | 0.76196 | 1.115277 | 1.02406 | 2.59414 | 2.710272 | 2.649354 | 3.15542 |
| Pla2g5 | 2.270074 | 2.15719 | 3.41831 | 3.221707 | 2.287285 | 2.336221 | 1.507762 | 1.474194 | 0.609258 | 0.769009 | 0.586044 | 0.493378 |
| Glipr1 | 0.954808 | 0.723052 | 1.264805 | 0.889177 | 1.091109 | 0.788944 | 0.551156 | 1.030046 | 1.997183 | 2.230277 | 4.098889 | 5.149912 |
| Havcr1 | 0.069433 | 0.144682 | 0.121795 | 0.1753 | 0.178908 | 0.253779 | 0.410504 | 0.392483 | 1.109084 | 0.822202 | 0.910268 | 1.142165 |
| Nlrp4a | 1.13637 | 1.391689 | 0.496953 | 0.602523 | 0.269608 | 0.258079 | 0.246752 | 0.264102 | 0.202405 | 0.380251 | 0.131441 | 0.09252 |
| Limch1 | 0.857797 | 0.634791 | 0.739 | 0.602558 | 1.095095 | 1.027742 | 1.629786 | 1.659082 | 3.833707 | 3.664316 | 4.270593 | 3.91524 |
| Kcnip2 | 1.353336 | 1.496696 | 0.940471 | 1.043355 | 0.834479 | 0.799415 | 0.693113 | 0.707371 | 0.252801 | 0.439887 | 0.170323 | 0.161185 |
| Pxdc1 | 2.106367 | 1.645665 | 0.551 | 0.936139 | 0.894567 | 1.237176 | 1.519951 | 1.826591 | 2.610807 | 2.171994 | 6.392379 | 6.042147 |
| Anxa3 | 2.145883 | 1.818102 | 5.390637 | 5.124605 | 6.838083 | 6.776463 | 9.843002 | 10.09535 | 9.996571 | 9.118165 | 32.20938 | 32.90873 |
| Hpca | 0.96321 | 0.85325 | 4.152926 | 3.918931 | 2.417344 | 2.950101 | 1.972081 | 2.111818 | 0.77959 | 0.837237 | 0.94117 | 0.611663 |
| Tmem132c | 0.208532 | 0.222918 | 0.389421 | 0.362564 | 0.588995 | 0.413035 | 0.722894 | 0.675529 | 2.243525 | 2.322729 | 1.875779 | 1.84998 |
| Peg3 | 6.507331 | 8.336508 | 11.31151 | 11.46278 | 10.79941 | 10.01099 | 15.71819 | 13.97819 | 61.41254 | 64.01536 | 55.67365 | 61.48359 |
| Itgb7 | 3.793728 | 4.398928 | 2.808162 | 2.812865 | 1.468511 | 1.448943 | 0.970432 | 1.086671 | 0.640944 | 0.623422 | 0.607214 | 0.440093 |
| Zscan4d | 2.68255 | 1.834523 | 10.17168 | 10.73262 | 24.09384 | 24.40475 | 59.39077 | 68.15524 | 54.50689 | 54.45804 | 48.35091 | 43.01444 |
| Amer2 | 0.170711 | 0.069612 | 0.143507 | 0.167424 | 0.30456 | 0.218895 | 0.443356 | 0.310378 | 1.218005 | 0.905798 | 1.049334 | 1.057932 |
| S1pr3 | 0.355567 | 0.402768 | 0.206774 | 0.245274 | 0.265199 | 0.237281 | 0.296742 | 0.219494 | 0.462219 | 0.717041 | 1.283348 | 1.385674 |
| Cybrd1 | 0.128406 | 0.102263 | 0.24592 | 0.259003 | 0.354722 | 0.313359 | 0.492427 | 0.4038 | 2.102299 | 1.57306 | 1.549453 | 1.635036 |
| Dkk1 | 0.489803 | 0.40023 | 0.474237 | 0.263234 | 0.448596 | 0.366184 | 0.812315 | 1.042778 | 1.416473 | 1.189988 | 1.722452 | 1.66043 |
| Hoxb9 | 0.383418 | 0.329086 | 10.52468 | 9.750312 | 26.36447 | 24.97231 | 48.49363 | 44.1177 | 54.85507 | 50.4686 | 58.36367 | 61.42012 |
| Mecom | 0.081419 | 0.093254 | 0.547464 | 0.54205 | 1.150216 | 1.29129 | 1.885646 | 1.893786 | 3.301672 | 3.404516 | 2.272402 | 2.185228 |
| Gm5039 | 4.172002 | 3.859664 | 16.08119 | 14.32594 | 38.48989 | 36.90168 | 71.64282 | 89.5323 | 66.55403 | 73.91351 | 61.54255 | 52.13614 |
| Slc35f1 | 0.437011 | 0.337559 | 0.29377 | 0.397882 | 0.599669 | 0.554251 | 0.989797 | 0.739857 | 1.978376 | 1.9473 | 1.935494 | 2.485457 |
| Scg3 | 0.295806 | 0.433958 | 0.397093 | 0.353839 | 0.411818 | 0.31446 | 0.515349 | 0.479299 | 0.960697 | 0.759736 | 2.508952 | 1.962627 |
| Fam84b | 0.528735 | 0.555247 | 0.4305 | 0.449891 | 0.4596 | 0.405572 | 0.882947 | 0.741115 | 2.390682 | 2.483389 | 2.552097 | 2.527394 |
| Ndufa4l2 | 7.288961 | 6.370732 | 2.371236 | 3.414442 | 4.907781 | 3.793861 | 2.43425 | 3.173293 | 1.454785 | 1.684976 | 0.439165 | 0.60912 |
| Myh14 | 1.939709 | 2.458652 | 3.21928 | 2.842853 | 2.68332 | 2.310164 | 2.198014 | 1.947086 | 0.937456 | 1.016609 | 0.557277 | 0.457898 |
| Gm7104 | 0.066531 | 0.028452 | 0.533907 | 0.373425 | 0.927168 | 1.184814 | 1.847714 | 2.311 | 1.843654 | 1.85851 | 2.120509 | 1.562366 |
| Oas3 | 0.985741 | 0.914285 | 1.431947 | 1.673522 | 1.386438 | 1.368344 | 1.010882 | 1.052524 | 0.545111 | 0.499338 | 0.280271 | 0.270563 |
| Zbbx | 1.061086 | 1.232183 | 2.694749 | 3.299555 | 1.796472 | 1.832644 | 1.52161 | 1.728782 | 0.588248 | 0.533528 | 0.792492 | 0.612391 |
| Hoxc4 | 0.190187 | 0.239188 | 3.771462 | 3.970987 | 10.35297 | 11.57103 | 22.27782 | 21.16863 | 27.1682 | 24.55285 | 18.80838 | 16.81447 |
| 2210417A0 | 1.318757 | 1.235452 | 2.177939 | 2.048558 | 1.390235 | 1.489978 | 0.929405 | 0.340852 | 0.422661 | 0.331611 | 0.571575 | 0.415258 |

| | | | | | | | | | | | | |
|---|---|---|---|---|---|---|---|---|---|---|---|---|
| Cyp11a1 | 0.591175 | 0.491819 | 2.395458 | 2.14795 | 2.296343 | 2.190945 | 2.063716 | 1.962449 | 0.700228 | 0.673255 | 0.730495 | 0.356096 |
| Slc24a1 | 0.796419 | 0.911977 | 1.011131 | 0.958283 | 1.093149 | 1.134703 | 0.927724 | 1.033899 | 0.608306 | 0.755933 | 0.180836 | 0.184831 |
| Scarf2 | 0.519303 | 0.429271 | 1.165772 | 1.442275 | 1.77503 | 1.576273 | 2.163466 | 2.086451 | 2.95202 | 3.073054 | 10.44985 | 8.84789 |
| Slco5a1 | 0.44081 | 0.615797 | 0.440616 | 0.482994 | 0.664811 | 0.672551 | 0.961663 | 0.754588 | 1.282276 | 1.332947 | 2.835044 | 2.960222 |
| Cpb2 | 0.136832 | 0.087512 | 0.314 | 0.386925 | 0.656808 | 0.710898 | 1.523469 | 1.813594 | 2.354503 | 2.361913 | 2.370174 | 2.049244 |
| Zfp352 | 0.097473 | 0.083536 | 0.455128 | 0.435329 | 0.957428 | 0.706027 | 1.90334 | 2.20333 | 2.174777 | 2.18183 | 2.934538 | 2.653586 |
| Cmtm3 | 2.580152 | 2.984941 | 3.468854 | 3.608529 | 5.971461 | 6.113683 | 9.24916 | 9.483625 | 17.25167 | 15.19837 | 21.68778 | 21.98311 |
| Gm4027 | 0.348484 | 0.205544 | 1.357199 | 1.315003 | 3.103954 | 3.021904 | 7.385085 | 8.005406 | 6.512327 | 6.309295 | 5.481111 | 4.78566 |
| Zyx | 9.61674 | 11.32898 | 10.07863 | 10.58643 | 12.45824 | 11.71398 | 15.36806 | 12.73661 | 20.07629 | 17.72063 | 65.26339 | 64.42834 |
| Peg12 | 4.229686 | 3.540943 | 5.570792 | 5.731925 | 10.45015 | 9.887144 | 13.18875 | 13.68732 | 27.13951 | 26.5244 | 35.29793 | 34.81975 |
| Hmx3 | 0.055334 | 0.1 | 0.219433 | 0.406944 | 0.482696 | 0.549834 | 0.696908 | 0.707671 | 1.215168 | 1.357814 | 2.606108 | 2.471381 |
| C430002E0 | 11.1019 | 10.25962 | 26.86989 | 25.43687 | 14.31929 | 15.15237 | 10.86342 | 9.316379 | 5.152167 | 4.995899 | 4.841667 | 4.189101 |
| Slc47a1 | 2.395669 | 2.465948 | 4.370624 | 4.514444 | 3.600677 | 4.158001 | 3.252658 | 2.902716 | 1.666557 | 1.564135 | 0.860867 | 0.7435 |
| Scg5 | 0.700307 | 1.002667 | 0.448164 | 0.625589 | 0.582695 | 1.066638 | 1.005607 | 0.969931 | 2.886082 | 2.779818 | 3.144914 | 3.79518 |
| Ednra | 0.167034 | 0.152397 | 0.204029 | 0.251067 | 0.600071 | 0.506092 | 0.921869 | 1.047237 | 1.452436 | 1.522922 | 0.615376 | 0.618871 |
| Zp3 | 1.049822 | 1.333077 | 1.092299 | 1.182668 | 0.731276 | 0.668199 | 0.472287 | 0.779347 | 0.418562 | 0.521423 | 0.501264 | 0.195338 |
| Gm20767 | 0.315072 | 0.503702 | 1.06571 | 1.199946 | 2.763553 | 2.604688 | 8.204327 | 7.259804 | 6.102904 | 6.678062 | 8.380529 | 6.740806 |
| Tnnt3 | 0.13166 | 0.038397 | 0.772372 | 0.648709 | 0.447102 | 0.477452 | 1.454874 | 0.738406 | 2.764064 | 2.840012 | 4.024649 | 2.888607 |
| F3 | 7.163234 | 6.658562 | 7.572501 | 6.63759 | 6.454024 | 6.67672 | 6.932365 | 7.12 | 11.05345 | 10.40164 | 35.24613 | 40.15667 |
| Sema6d | 0.222353 | 0.210464 | 0.232858 | 0.278685 | 0.321824 | 0.366278 | 0.561438 | 0.458198 | 1.523677 | 1.497117 | 1.713912 | 1.683822 |
| Zscan4c | 3.541396 | 3.280408 | 14.16441 | 12.6021 | 28.10488 | 29.11523 | 65.92883 | 75.89655 | 64.76474 | 65.24183 | 55.10448 | 48.40923 |
| Dnajc22 | 4.237627 | 3.875998 | 5.851912 | 5.594287 | 5.485523 | 4.759531 | 4.150734 | 4.075399 | 1.940319 | 1.96832 | 0.904989 | 0.928923 |
| Wnt5b | 0.381539 | 0.451422 | 0.852217 | 0.73083 | 1.918902 | 1.740043 | 2.460637 | 2.767352 | 4.698164 | 4.398494 | 2.620045 | 2.234063 |
| Fam69c | 0.496226 | 0.48303 | 0.64547 | 0.577334 | 1.057822 | 0.945673 | 0.90013 | 1.219023 | 0.732209 | 0.800064 | 0.248866 | 0.202833 |
| Tet1 | 32.68842 | 45.46316 | 27.80643 | 28.81946 | 14.48744 | 14.78091 | 11.5827 | 9.080519 | 6.919261 | 7.627658 | 4.778567 | 4.806705 |
| Prdm1 | 2.325119 | 2.880129 | 1.290721 | 1.546468 | 0.911169 | 0.686855 | 0.879195 | 0.825344 | 3.815658 | 4.107082 | 2.747549 | 2.621982 |
| Smpd3 | 0.285944 | 0.420301 | 0.32874 | 0.308781 | 0.222971 | 0.306443 | 0.241399 | 0.309534 | 0.523924 | 0.455125 | 1.875563 | 1.827827 |
| Gpc6 | 0.756011 | 0.913626 | 0.81658 | 0.950005 | 1.754511 | 1.66578 | 2.60866 | 2.24962 | 5.363765 | 5.531983 | 5.278481 | 5.664322 |
| Ptrf | 8.070527 | 8.648225 | 11.35748 | 9.992732 | 10.81105 | 9.78712 | 9.940773 | 9.411008 | 6.146162 | 6.373726 | 44.61487 | 37.97036 |
| Dsp | 2.819523 | 4.002563 | 6.976223 | 7.049733 | 10.07555 | 9.553713 | 17.74179 | 15.5765 | 35.45609 | 35.82708 | 42.98949 | 41.99659 |
| Slit3 | 0.888151 | 0.988176 | 0.627444 | 0.54573 | 0.762222 | 0.817349 | 1.069539 | 1.08472 | 2.195648 | 2.086116 | 3.190397 | 3.238829 |
| Pdzd4 | 31.48151 | 31.79544 | 12.79772 | 12.11537 | 7.30128 | 6.656141 | 4.038946 | 4.474213 | 2.808449 | 2.926443 | 1.969269 | 2.042285 |
| Gm11545 | 1.610948 | 1.470709 | 1.51189 | 2.279762 | 4.78577 | 5.207183 | 9.705399 | 10.37325 | 13.9538 | 13.49547 | 12.03449 | 12.29192 |
| Tmem40 | 2.958238 | 2.869104 | 2.086166 | 2.328917 | 1.373464 | 1.303132 | 0.998049 | 1.20636 | 0.573523 | 0.394171 | 0.726705 | 0.519947 |
| Artn | 0.500334 | 0.767409 | 0.560029 | 0.41381 | 0.583246 | 0.522501 | 0.437903 | 0.731962 | 0.642442 | 0.539965 | 3.051453 | 2.433305 |
| Airn | 0.219574 | 0.249775 | 0.368935 | 0.283673 | 0.526647 | 0.466266 | 0.899195 | 0.715788 | 1.347873 | 1.667929 | 1.669088 | 1.545741 |
| Pfkfb3 | 4.030149 | 3.751321 | 2.023864 | 2.67511 | 3.625185 | 3.430628 | 3.494382 | 4.306006 | 5.951451 | 5.813565 | 14.66091 | 15.72584 |
| Filip1l | 0.32153 | 0.421789 | 0.347581 | 0.516797 | 0.397839 | 0.517077 | 0.452685 | 0.494778 | 1.316052 | 1.377599 | 2.563574 | 2.895914 |
| Mycl | 6.106194 | 6.276108 | 3.841583 | 4.570944 | 6.035202 | 6.473406 | 10.77405 | 11.24489 | 27.06752 | 26.69074 | 17.18876 | 17.61719 |
| Zscan4e | 3.106333 | 3.314637 | 12.87165 | 12.59228 | 31.40556 | 31.02634 | 76.96574 | 73.34901 | 61.70512 | 63.10735 | 54.62754 | 48.30009 |
| Actn3 | 16.33118 | 15.92362 | 33.47165 | 31.09497 | 27.86412 | 24.80049 | 21.05768 | 23.80783 | 9.697331 | 10.58346 | 5.72168 | 5.341956 |
| Fgfbp1 | 9.56324 | 9.475336 | 4.18898 | 3.836696 | 3.523992 | 2.700878 | 4.489935 | 5.239042 | 5.599159 | 5.847294 | 16.98507 | 15.66424 |
| Tspan12 | 0.508112 | 0.424493 | 0.675384 | 0.834882 | 1.988988 | 1.796379 | 2.445252 | 2.578631 | 4.560963 | 4.838558 | 3.637607 | 3.968559 |
| Nos1ap | 0.352639 | 0.446909 | 0.702882 | 0.77892 | 1.166969 | 1.115636 | 2.15215 | 1.772953 | 4.293465 | 4.509994 | 4.221731 | 3.605821 |
| Lgals3 | 14.4557 | 12.37167 | 5.183616 | 5.491566 | 3.120969 | 3.359534 | 1.531816 | 1.981695 | 1.112367 | 0.948537 | 4.027785 | 3.363616 |
| Gm2022 | 0.926046 | 1.451975 | 4.130592 | 4.211487 | 8.847021 | 10.61547 | 17.38564 | 24.34934 | 18.45213 | 23.11406 | 19.06705 | 17.2304 |
| 4933413J0 | 0.33932 | 0.375413 | 2.303871 | 2.443014 | 1.569521 | 1.520898 | 0.880183 | 1.1485 | 0.608379 | 0.64201 | 0.37983 | 0.423065 |
| Krt19 | 2.802813 | 2.470646 | 1.489029 | 1.543626 | 1.509465 | 1.402824 | 2.155792 | 1.724804 | 3.611301 | 3.604931 | 9.972411 | 8.09781 |
| Pdlim3 | 0.766365 | 0.804027 | 0.926585 | 0.935448 | 2.183389 | 2.105809 | 3.470044 | 4.900551 | 4.416838 | 4.992137 | 6.570165 | 5.395558 |
| Spon1 | 0.464456 | 0.456775 | 0.299253 | 0.349972 | 0.40883 | 0.403037 | 0.464178 | 0.266247 | 1.198684 | 1.217802 | 1.635611 | 1.533979 |
| Ptprd | 0.232369 | 0.252341 | 0.322546 | 0.35839 | 0.49209 | 0.415392 | 0.682632 | 0.620223 | 2.028354 | 1.981217 | 1.990401 | 2.057825 |
| Tpm1 | 44.20798 | 40.799 | 46.32952 | 45.75477 | 50.64915 | 49.00021 | 54.34727 | 56.98649 | 90.07889 | 86.39365 | 264.6796 | 262.3777 |
| Fam149a | 0.368285 | 0.365667 | 0.716323 | 0.616517 | 1.395413 | 1.282073 | 1.997097 | 2.039563 | 3.052021 | 3.107654 | 3.237885 | 3.533288 |
| Tmem238 | 5.305393 | 5.027811 | 5.825159 | 6.779413 | 4.747054 | 4.906101 | 3.131628 | 3.365176 | 1.37631 | 1.183117 | 1.325046 | 2.047644 |
| Sftpb | 7.127468 | 1.173935 | 0.253185 | 0.984955 | 0.73123 | 0.642173 | 1.132182 | 1.391792 | 0.089373 | 0.499117 | 0.22019 | 0.243026 |
| Cdh10 | 0.604449 | 0.662234 | 0.29042 | 0.183975 | 0.241266 | 0.471267 | 0.431387 | 0.510372 | 1.042208 | 1.053499 | 0.847398 | 0.939108 |
| Zscan4b | 1.055437 | 1.039886 | 3.97728 | 3.717289 | 8.865969 | 9.092729 | 21.51163 | 21.26922 | 18.74999 | 18.5587 | 16.74984 | 14.54779 |
| Gm1322 | 0.716339 | 0.514645 | 2.500802 | 1.955878 | 1.829756 | 1.281827 | 1.408547 | 1.398024 | 0.279909 | 0.351946 | 0.509225 | 0.342099 |
| Tm4sf5 | 1.390862 | 1.160564 | 2.01633 | 1.719075 | 1.348071 | 1.320979 | 0.914244 | 0.984209 | 0.681188 | 0.301301 | 0.386758 | 0.338769 |
| Tnnt1 | 3.312704 | 3.694624 | 8.517703 | 8.733255 | 6.142127 | 7.27 | 4.672213 | 3.894527 | 2.173796 | 1.639688 | 1.637049 | 1.530826 |
| AF067063 | 0.170573 | 0.071785 | 0.202309 | 0.22369 | 0.30744 | 0.635803 | 1.044906 | 1.191536 | 1.332579 | 1.270106 | 0.723763 | 1.267366 |

| | | | | | | | | | | | | |
|---|---|---|---|---|---|---|---|---|---|---|---|---|
| Lrmp | 5.845546 | 5.81716 | 5.181065 | 4.637606 | 2.960705 | 2.291681 | 1.626575 | 1.980584 | 0.80661 | 1.065907 | 0.75442 | 0.818026 |
| Lhx1 | 0.052551 | 0.027411 | 0.185101 | 0.155184 | 0.361514 | 0.319464 | 0.822279 | 0.767714 | 0.598557 | 0.592713 | 1.221027 | 0.879451 |
| Trim43a | 0.081535 | 0.17197 | 0.296959 | 0.288693 | 0.274592 | 0.315936 | 0.398128 | 0.82804 | 1.449273 | 1.584066 | 1.971877 | 1.635437 |
| Rgl1 | 0.605038 | 0.667122 | 0.863123 | 0.652368 | 1.321847 | 1.295798 | 1.680565 | 1.665935 | 3.211473 | 3.362255 | 3.431312 | 3.686013 |
| Mcf2 | 11.26353 | 11.58745 | 3.439942 | 3.76346 | 2.483443 | 2.372713 | 2.124907 | 2.394912 | 1.509014 | 1.680703 | 0.795445 | 0.666675 |
| Vopp1 | 1.014569 | 0.918217 | 1.149768 | 1.180777 | 2.329689 | 2.294167 | 3.410443 | 3.451215 | 6.882555 | 6.456389 | 6.301192 | 6.662773 |
| Pmp22 | 1.439008 | 1.121355 | 5.891276 | 5.133826 | 9.175518 | 8.406658 | 14.44651 | 11.96114 | 20.9764 | 17.52607 | 26.55451 | 28.94401 |
| Capsl | 8.665187 | 8.536182 | 9.535113 | 7.995962 | 5.747386 | 5.918817 | 3.70868 | 4.16156 | 1.624773 | 1.651238 | 1.182301 | 1.419269 |
| Cacng8 | 1.783807 | 2.200107 | 3.453882 | 3.760416 | 2.973371 | 3.751232 | 3.448243 | 2.915296 | 1.669528 | 1.530855 | 0.822449 | 0.668039 |
| Glod5 | 4.10151 | 2.861355 | 0.611248 | 0.759009 | 0.360067 | 0.134848 | 0.201016 | 0.626752 | 0.557068 | 0.412378 | 0.289353 | 0.391398 |
| Kcne1l | 0.392596 | 0.206762 | 0.299755 | 0.349399 | 0.836015 | 0.564062 | 1.255982 | 0.971968 | 1.536406 | 1.175498 | 1.986161 | 1.966067 |
| Lpp | 2.228969 | 3.267484 | 4.553975 | 4.863622 | 4.560498 | 4.571609 | 5.970253 | 3.929803 | 6.449338 | 6.49525 | 20.89689 | 22.09941 |
| 1110032F0 | 0.203706 | 0.263576 | 0.250511 | 0.204774 | 0.476504 | 0.437137 | 0.446345 | 0.618107 | 0.953721 | 1.151063 | 0.959003 | 1.117381 |
| Tmem59l | 0.924809 | 0.954578 | 2.716792 | 2.880716 | 2.242179 | 2.514083 | 1.373117 | 2.055574 | 0.795095 | 0.513641 | 0.553546 | 0.704357 |
| Nrp2 | 1.178433 | 1.136078 | 0.839193 | 0.92823 | 0.973464 | 0.917842 | 1.267761 | 1.254808 | 4.791238 | 5.147336 | 5.494134 | 5.106295 |
| Col13a1 | 1.701531 | 1.889919 | 3.589469 | 3.431956 | 5.82483 | 5.505409 | 4.603003 | 5.335912 | 1.382238 | 1.521523 | 1.050262 | 0.984759 |
| Tchh | 0.052087 | 0.072576 | 2.921338 | 3.053406 | 2.070988 | 2.296816 | 1.754509 | 1.867279 | 0.667612 | 0.546389 | 1.428242 | 1.303168 |
| Utf1 | 248.0913 | 281.3352 | 151.2642 | 132.3069 | 74.70037 | 73.51936 | 43.41451 | 47.8602 | 26.75963 | 24.3463 | 26.08193 | 23.68262 |
| Ahnak | 3.664089 | 4.951147 | 3.992483 | 4.188439 | 2.999764 | 3.196121 | 3.154755 | 2.478137 | 2.959758 | 3.357761 | 12.89106 | 13.84311 |
| Sh3pxd2b | 2.454499 | 2.852722 | 3.501506 | 3.692322 | 5.654274 | 5.613692 | 8.325273 | 7.049346 | 13.96404 | 14.40944 | 21.06497 | 20.57023 |
| Nrarp | 2.413223 | 2.293603 | 1.928352 | 1.629584 | 1.895929 | 2.318527 | 1.817966 | 1.942207 | 8.606391 | 9.036154 | 7.373141 | 6.740898 |
| Mfap3l | 1.324422 | 1.314152 | 0.638591 | 0.608317 | 0.603911 | 0.740281 | 0.889107 | 0.86558 | 1.182656 | 1.171518 | 3.236552 | 3.371112 |
| Prkcdbp | 0.487518 | 0.741037 | 1.256205 | 1.3106 | 1.401653 | 1.640261 | 1.633861 | 1.766006 | 1.203822 | 0.984411 | 5.645207 | 5.454772 |
| Stbd1 | 0.852784 | 0.642425 | 0.900892 | 1.045307 | 1.896034 | 1.904214 | 2.351037 | 3.211045 | 2.624921 | 2.563425 | 5.312562 | 5.790956 |
| Bspry | 3.404317 | 3.324299 | 2.892729 | 3.138268 | 2.312266 | 1.865585 | 1.528294 | 1.297658 | 0.707444 | 0.56716 | 0.799298 | 0.741056 |
| Clu | 6.613188 | 6.602643 | 18.24136 | 18.57116 | 29.54626 | 28.8059 | 58.22122 | 62.36868 | 77.233 | 69.45865 | 111.1204 | 102.6345 |
| Spats2l | 1.307878 | 1.205209 | 1.224938 | 1.03446 | 2.313392 | 2.265159 | 3.133979 | 3.02984 | 5.677359 | 5.709987 | 4.16657 | 4.652679 |
| Rgmb | 2.09122 | 2.499029 | 2.638192 | 2.460139 | 3.328056 | 3.413975 | 6.699035 | 5.517963 | 14.86467 | 13.57143 | 13.64254 | 13.51423 |
| 4930461G1 | 9.504935 | 8.123043 | 5.02255 | 4.60976 | 2.892089 | 3.250688 | 2.724161 | 3.845505 | 2.962895 | 2.302499 | 0.843784 | 0.836506 |
| Sox17 | 1.45451 | 1.256653 | 1.009196 | 0.871415 | 1.21589 | 1.303214 | 2.164727 | 2.395762 | 4.481528 | 3.545229 | 5.282368 | 4.794325 |
| Echdc3 | 0.228893 | 0.241002 | 0.292481 | 0.21602 | 0.52235 | 0.580731 | 0.826228 | 0.886196 | 0.617463 | 0.711117 | 1.24551 | 1.187516 |
| Ier5 | 1.145119 | 1.145836 | 1.800918 | 1.700656 | 3.086462 | 3.464026 | 4.864778 | 5.050086 | 4.805883 | 4.596929 | 9.835213 | 9.336567 |
| Errfi1 | 6.252338 | 6.297703 | 8.444482 | 8.919373 | 10.83562 | 9.893954 | 11.51833 | 10.33448 | 18.88066 | 18.32891 | 47.01539 | 48.95724 |
| Vgll3 | 0.303195 | 0.344232 | 0.524196 | 0.540592 | 0.418172 | 0.462877 | 0.495003 | 0.564324 | 0.645516 | 0.521517 | 2.355728 | 2.534991 |
| AA543401 | 0.828542 | 0.94938 | 3.041789 | 2.681715 | 2.506056 | 1.960449 | 1.807929 | 2.25287 | 0.494544 | 0.492038 | 0.694049 | 0.506184 |
| Hif3a | 1.593256 | 1.910413 | 1.651015 | 1.950548 | 1.560226 | 1.414841 | 1.357911 | 1.200594 | 0.915847 | 1.149353 | 0.488739 | 0.358226 |
| Frem2 | 1.948991 | 2.808323 | 4.149287 | 4.508614 | 3.996584 | 3.807812 | 4.530577 | 3.283391 | 1.687834 | 1.906723 | 0.928496 | 0.829356 |
| Pkd2l1 | 0.051284 | 0.101621 | 0.276018 | 0.248208 | 0.44533 | 0.473635 | 0.388904 | 0.424978 | 1.025115 | 1.295845 | 1.276623 | 1.347562 |
| Foxa2 | 0.48259 | 0.348517 | 0.542873 | 0.460999 | 0.577649 | 0.686374 | 0.877612 | 0.925008 | 2.53672 | 2.401436 | 3.136816 | 2.501112 |
| Vsig2 | 2.359958 | 2.549299 | 1.424598 | 1.698248 | 1.067761 | 1.329963 | 0.720027 | 1.20172 | 0.65557 | 0.376151 | 0.371926 | 0.313094 |
| Msx1 | 0.218238 | 0.243629 | 0.811816 | 0.879293 | 1.884138 | 1.69808 | 3.591382 | 4.626157 | 5.712412 | 4.762455 | 1.974544 | 2.449665 |
| Peg10 | 61.19212 | 81.299 | 129.3332 | 144.1433 | 212.4274 | 207.6161 | 391.1575 | 319.88 | 751.8877 | 779.6676 | 442.4306 | 466.0411 |
| Esrp1 | 5.364271 | 6.030308 | 2.578795 | 3.031728 | 1.813505 | 1.936801 | 1.002735 | 1.128012 | 0.576147 | 0.578671 | 0.568491 | 0.56109 |
| Pianp | 3.405508 | 4.075721 | 5.190276 | 5.420307 | 4.107265 | 3.538948 | 2.889546 | 2.853685 | 1.425113 | 1.423714 | 1.286873 | 1.003784 |
| Sox11 | 7.5763 | 8.55121 | 4.505485 | 5.284176 | 6.657059 | 6.893821 | 10.1621 | 8.441828 | 27.89263 | 28.4121 | 28.10068 | 28.53291 |
| Tmprss12 | 1.149735 | 1.200575 | 0.818444 | 0.893172 | 0.492828 | 0.446842 | 0.346873 | 0.277937 | 0.409753 | 0.452607 | 0.17822 | 0.165488 |
| Epha7 | 0.741582 | 0.807922 | 1.586415 | 1.401988 | 1.397119 | 1.430084 | 1.55184 | 1.632637 | 5.040081 | 5.365204 | 7.387657 | 7.564858 |
| Cyp26b1 | 0.152725 | 0.103392 | 1.152079 | 1.369893 | 2.846439 | 2.862893 | 6.155245 | 5.277603 | 7.918258 | 6.320027 | 6.683652 | 7.390776 |
| Slc7a7 | 43.66179 | 43.66082 | 28.74747 | 26.76668 | 23.71369 | 24.27256 | 21.21218 | 19.95462 | 5.293461 | 4.963999 | 5.923023 | 5.55519 |
| Col26a1 | 0.515226 | 0.601069 | 0.59551 | 0.607266 | 0.941388 | 0.99191 | 1.089805 | 1.247093 | 2.794748 | 2.517588 | 3.642164 | 3.270035 |
| Rasgrp4 | 2.001628 | 2.235649 | 2.619436 | 2.50567 | 2.201076 | 2.409809 | 1.618222 | 1.734316 | 0.733036 | 0.856223 | 0.53443 | 0.465366 |
| Crb2 | 0.297595 | 0.384934 | 0.528845 | 0.513888 | 0.710681 | 0.631254 | 0.998773 | 0.838488 | 2.318139 | 1.85125 | 2.681456 | 2.76665 |
| Bcl6b | 1.652588 | 1.716029 | 3.291502 | 3.795691 | 3.2065 | 3.125152 | 2.699136 | 2.548033 | 0.873332 | 0.784999 | 0.661459 | 0.705249 |
| Eml1 | 1.779329 | 1.776416 | 2.187166 | 2.329906 | 4.280063 | 4.271021 | 6.595646 | 6.216719 | 12.59857 | 12.5366 | 12.12222 | 12.53947 |
| Spata31d1l | 0.867071 | 1.224951 | 0.857353 | 1.027491 | 0.582882 | 0.501456 | 0.426143 | 0.481699 | 0.24833 | 0.277817 | 0.077502 | 0.191066 |
| BB287469 | 0.141481 | 0.137786 | 0.692051 | 0.466101 | 1.131212 | 1.215343 | 1.777108 | 2.02559 | 2.232397 | 2.506036 | 1.992197 | 1.890984 |
| Tnfaip3 | 0.285943 | 0.325054 | 0.428074 | 0.491253 | 0.467589 | 0.515664 | 0.451341 | 0.385582 | 0.374456 | 0.465898 | 2.083924 | 2.06766 |
| Rhbg | 0.530895 | 0.723464 | 1.067363 | 0.942026 | 0.790858 | 0.83099 | 0.683668 | 0.798458 | 0.418912 | 0.415491 | 0.341774 | 0.176179 |
| Aadat | 0.237554 | 0.226302 | 0.259844 | 0.219129 | 0.373156 | 0.674651 | 0.689136 | 0.552926 | 1.22855 | 1.118049 | 1.391984 | 1.169981 |
| Nell2 | 0.622828 | 0.653477 | 0.923241 | 1.036252 | 1.160412 | 1.190466 | 1.623885 | 1.698688 | 3.900805 | 3.51274 | 4.249518 | 5.524576 |
| Gm7325 | 76.27105 | 66.53032 | 39.39898 | 37.0455 | 30.56328 | 32.46725 | 23.11056 | 25.57887 | 14.54357 | 13.82512 | 6.14524 | 6.994995 |

| | | | | | | | | | | | | |
|---|---|---|---|---|---|---|---|---|---|---|---|---|
| Tmem54 | 1.532944 | 0.937575 | 1.265662 | 1.071146 | 1.416535 | 1.458245 | 0.908636 | 0.527584 | 0.388666 | 0.303091 | 0.272015 | 0.275443 |
| Cysltr2 | 0.714915 | 0.953172 | 3.582741 | 3.383932 | 1.944784 | 2.142739 | 2.250763 | 1.664536 | 0.759923 | 0.6413 | 0.582006 | 0.81176 |
| Dbn1 | 9.675192 | 10.84782 | 11.95002 | 11.97043 | 18.62573 | 19.00358 | 29.41978 | 30.05706 | 52.62094 | 47.52696 | 68.10543 | 63.14552 |
| Notch4 | 4.203111 | 4.742261 | 2.115909 | 2.14273 | 1.246605 | 1.367093 | 0.903166 | 0.874625 | 0.459665 | 0.472174 | 0.443945 | 0.406254 |
| A930011O1 | 2.299594 | 2.306072 | 1.103894 | 1.254474 | 0.682427 | 0.734572 | 0.357421 | 0.479751 | 0.196058 | 0.238676 | 0.163332 | 0.247092 |
| Fzd3 | 2.13423 | 2.083424 | 3.44249 | 3.466889 | 4.500706 | 4.534773 | 6.007994 | 4.881754 | 14.07316 | 16.15533 | 16.16184 | 18.14181 |
| Gm16880 | 7.791377 | 9.217745 | 3.213623 | 3.009652 | 2.30588 | 2.201367 | 1.254565 | 1.130375 | 0.837465 | 0.75148 | 0.33878 | 0.575752 |
| Cited1 | 0.646043 | 0.545681 | 0.608142 | 0.610795 | 1.479139 | 1.447312 | 2.101705 | 1.891883 | 2.894399 | 2.256923 | 3.309895 | 3.181672 |
| Tpm4 | 52.24113 | 48.55099 | 36.94756 | 37.34581 | 52.13647 | 48.65608 | 62.94845 | 66.33631 | 119.6813 | 118.17 | 186.9071 | 194.3098 |
| BC147527 | 0.399909 | 0.545308 | 0.647032 | 0.930122 | 2.676352 | 2.215066 | 4.505214 | 4.091571 | 5.016508 | 4.838048 | 6.223487 | 4.620482 |
| Slc7a3 | 68.17158 | 63.65339 | 26.12825 | 24.30065 | 14.6536 | 14.54286 | 9.072244 | 9.390461 | 5.265422 | 4.682508 | 8.603626 | 8.944831 |
| Ackr3 | 1.281637 | 1.134311 | 1.575109 | 1.396374 | 2.261821 | 1.924809 | 2.502069 | 3.058211 | 5.920114 | 5.804466 | 6.929451 | 7.240642 |
| Csrp1 | 20.67993 | 18.14447 | 27.91147 | 25.29476 | 28.89185 | 27.73174 | 32.38698 | 30.35593 | 33.87128 | 33.18027 | 138.2478 | 131.12 |
| Nlrp9b | 1.596081 | 1.741302 | 2.319355 | 2.343065 | 1.443121 | 1.477328 | 0.967862 | 1.050518 | 0.394134 | 0.629155 | 0.359608 | 0.452208 |
| Lfng | 1.138945 | 1.438986 | 1.661622 | 1.925222 | 2.276381 | 2.237813 | 3.99015 | 3.470876 | 6.483874 | 5.667974 | 9.747333 | 9.970315 |
| Flnc | 2.916968 | 3.813843 | 6.574285 | 5.907847 | 6.257771 | 5.871571 | 7.870919 | 6.140887 | 4.785901 | 5.100362 | 28.99259 | 26.40559 |
| Vstm2b | 0.375644 | 0.333672 | 1.127355 | 0.837615 | 2.509489 | 2.123296 | 3.935376 | 4.328211 | 4.588428 | 4.25956 | 3.452155 | 2.925966 |
| Dnaic1 | 1.027956 | 1.203354 | 1.403683 | 1.310827 | 1.195546 | 1.085608 | 1.08829 | 1.792526 | 0.362334 | 0.409883 | 0.192029 | 0.346908 |
| Fgfbp3 | 1.90964 | 1.675703 | 1.167962 | 0.844894 | 2.059282 | 1.857817 | 2.447483 | 2.618421 | 5.123917 | 4.361691 | 2.533712 | 2.839297 |
| Nexn | 2.163604 | 2.020949 | 1.658905 | 2.078469 | 2.036289 | 1.955374 | 1.802145 | 2.048835 | 4.472928 | 4.536248 | 9.273054 | 10.09112 |
| Zfp36l1 | 12.83094 | 15.39696 | 10.59502 | 9.693979 | 12.49655 | 12.52493 | 20.05403 | 19.7269 | 45.09565 | 40.40427 | 53.04479 | 50.02518 |
| Irx1 | 0.246609 | 0.159466 | 0.4985 | 0.343379 | 0.661674 | 0.716553 | 0.988294 | 0.790006 | 1.920562 | 1.54118 | 1.977817 | 1.77124 |
| Adra2a | 0.090109 | 0.111184 | 0.190659 | 0.241505 | 0.360361 | 0.292474 | 0.674815 | 0.505538 | 1.351921 | 1.244828 | 1.314214 | 1.23379 |
| Glipr2 | 18.11493 | 15.97356 | 13.9517 | 15.06957 | 17.46626 | 17.99364 | 21.62561 | 22.96891 | 27.36414 | 26.20857 | 75.94645 | 77.5704 |
| Plekho1 | 6.3148 | 4.910684 | 4.942822 | 4.599685 | 8.137375 | 7.97728 | 11.15612 | 12.0869 | 18.8995 | 17.58481 | 22.45182 | 23.59895 |
| Lats2 | 1.002395 | 1.258164 | 2.202057 | 2.390149 | 3.172565 | 3.061058 | 3.897507 | 3.819067 | 7.282545 | 7.76309 | 11.88456 | 12.25535 |
| Penk | 0.958917 | 0.627476 | 0.529545 | 0.621714 | 0.890249 | 1.22884 | 1.580993 | 1.617222 | 2.716433 | 2.633513 | 3.863426 | 3.177938 |
| Pde5a | 1.956204 | 2.052746 | 2.270506 | 2.724436 | 3.838154 | 3.884104 | 6.954951 | 5.753897 | 12.99065 | 13.89681 | 10.26544 | 10.80767 |
| Has2 | 1.3315 | 1.692917 | 0.691195 | 0.895577 | 1.060333 | 0.972608 | 1.450365 | 1.924681 | 2.577123 | 2.506719 | 0.431002 | 0.492147 |
| D930048N1 | 6.823794 | 7.642461 | 5.518944 | 5.243944 | 3.809852 | 3.841323 | 2.238242 | 3.036922 | 1.604946 | 1.434951 | 0.772596 | 1.029803 |
| Camk2d | 1.257142 | 1.114057 | 1.540408 | 1.619863 | 2.106858 | 2.397695 | 2.595908 | 2.808682 | 6.163072 | 6.839036 | 7.805418 | 8.239666 |
| Lpar3 | 0.299297 | 0.186063 | 0.264934 | 0.429411 | 0.621942 | 0.56291 | 0.92022 | 0.751677 | 1.144269 | 1.175523 | 1.930915 | 2.181251 |
| Pecam1 | 17.48559 | 19.46475 | 26.24617 | 24.34776 | 13.49871 | 11.98039 | 7.314722 | 9.002949 | 6.890932 | 8.111662 | 4.789406 | 4.794452 |
| Lemd1 | 2.361704 | 2.843667 | 1.501023 | 1.618938 | 0.966398 | 0.967204 | 0.554213 | 0.551273 | 0.578991 | 0.536292 | 0.401247 | 0.318958 |
| Cacna1s | 0.186968 | 0.206577 | 0.591588 | 0.507025 | 1.324838 | 1.135953 | 2.029372 | 2.1293 | 2.281209 | 2.075887 | 2.985119 | 2.565903 |
| Tgm1 | 1.845638 | 2.207209 | 8.242475 | 6.849541 | 3.276506 | 3.622288 | 3.02644 | 2.448526 | 1.384752 | 1.353971 | 2.191473 | 2.054399 |
| Speer7-ps1 | 0.018482 | 0.1 | 0.348345 | 0.243725 | 0.428067 | 0.722741 | 1.480309 | 0.865505 | 1.209212 | 1.231875 | 1.101019 | 0.732126 |
| Tuft1 | 1.37851 | 1.310744 | 1.092859 | 1.246071 | 1.309151 | 1.074183 | 1.332961 | 1.246939 | 1.988403 | 1.88131 | 5.713136 | 5.426859 |
| Nefm | 0.317969 | 0.373167 | 0.452933 | 0.438053 | 0.498573 | 0.550716 | 0.447226 | 0.573103 | 0.540969 | 0.496637 | 2.379151 | 2.212743 |
| Klk13 | 1.188183 | 1.476011 | 0.97802 | 1.644344 | 1.403583 | 1.107622 | 0.739875 | 1.088877 | 0.346276 | 0.784834 | 0.369154 | 0.325537 |
| A230050P2 | 7.652426 | 8.267003 | 6.765473 | 6.460893 | 5.037503 | 4.841934 | 2.899156 | 3.343478 | 1.796957 | 1.965644 | 1.148896 | 1.280489 |
| Prickle1 | 2.438521 | 3.051323 | 1.93547 | 2.095462 | 2.600208 | 2.793566 | 4.163689 | 3.901821 | 6.463505 | 6.249938 | 10.09745 | 10.56669 |
| Inpp5d | 30.27164 | 33.20206 | 36.12258 | 34.36156 | 24.67208 | 24.49971 | 19.66451 | 19.72652 | 9.805243 | 9.707288 | 7.119955 | 6.825585 |
| 2410088K1 | 1.55718 | 1.947173 | 3.105904 | 4.692134 | 3.324895 | 3.396374 | 2.368724 | 2.250403 | 1.151892 | 1.038786 | 0.658708 | 0.932477 |
| Prss50 | 1.315128 | 1.58104 | 6.59499 | 6.276819 | 5.257689 | 4.530542 | 4.209599 | 3.543216 | 1.869671 | 1.440008 | 1.178459 | 1.248685 |
| Parp8 | 0.55457 | 0.534305 | 0.643984 | 0.903684 | 1.3248 | 1.107372 | 1.707266 | 1.639134 | 2.392254 | 2.229387 | 4.517488 | 4.535675 |
| Tppp3 | 3.033966 | 2.075912 | 1.945162 | 1.800463 | 3.373527 | 2.738158 | 5.101376 | 5.237297 | 7.972752 | 7.758696 | 8.740983 | 9.012947 |
| Il1r1 | 0.233389 | 0.149101 | 0.374292 | 0.357321 | 0.608295 | 0.547664 | 0.665735 | 0.766722 | 0.725324 | 0.667871 | 1.897882 | 1.788092 |
| Krt8 | 9.039575 | 8.321468 | 9.216515 | 9.009922 | 9.149329 | 9.232836 | 11.39818 | 11.78818 | 18.40477 | 18.91561 | 51.61435 | 45.04085 |
| Sncaip | 0.189767 | 0.25577 | 0.723451 | 0.736483 | 0.69817 | 0.623936 | 1.008211 | 0.918716 | 2.55464 | 2.366022 | 3.160572 | 3.11505 |
| Hoxb6 | 0.010333 | 0.021095 | 9.780268 | 9.574677 | 23.14258 | 24.62771 | 45.82128 | 45.22054 | 59.07142 | 46.76128 | 48.7356 | 47.80025 |
| BC080695 | 0.095887 | 0.121333 | 0.350844 | 0.390186 | 0.786853 | 0.7987 | 1.729474 | 1.382378 | 1.91053 | 1.946546 | 1.631454 | 1.502832 |
| Oprd1 | 0.351154 | 0.50204 | 2.19823 | 2.099143 | 2.404654 | 2.080641 | 2.038854 | 2.121316 | 0.778584 | 0.765102 | 0.422768 | 0.425583 |
| Ebf3 | 0.257187 | 0.300462 | 0.157914 | 0.224008 | 0.260866 | 0.226005 | 0.270696 | 0.308029 | 0.633199 | 0.59772 | 1.22898 | 1.116483 |
| Snca | 0.819659 | 0.464686 | 0.371912 | 0.438293 | 1.165714 | 0.948237 | 0.858325 | 1.3202 | 0.570114 | 0.752305 | 0.199631 | 0.265077 |
| Tmem132e | 0.023009 | 0.009446 | 0.489851 | 0.538054 | 1.634365 | 1.603453 | 3.247071 | 2.671403 | 1.988387 | 1.910382 | 2.740556 | 2.554923 |
| Fgfrl1 | 3.363178 | 3.707493 | 4.541328 | 4.212594 | 6.689558 | 6.515368 | 9.845388 | 10.81684 | 22.29842 | 20.61436 | 21.73639 | 20.8738 |
| Elovl2 | 3.16342 | 3.458112 | 3.277705 | 3.574938 | 5.644002 | 4.720679 | 7.91216 | 7.714493 | 17.84423 | 17.70004 | 14.48069 | 16.74996 |
| Rasa4 | 2.388241 | 2.488639 | 2.133579 | 1.970763 | 1.508351 | 1.272102 | 0.9658 | 0.968768 | 0.63532 | 0.711581 | 0.538507 | 0.398489 |
| Steap3 | 3.434052 | 3.88095 | 27.17638 | 25.29635 | 19.36574 | 17.68297 | 13.1826 | 12.43005 | 6.780711 | 6.738118 | 5.846344 | 5.12171 |
| Pcyt1b | 8.499304 | 9.024521 | 5.45374 | 5.916388 | 5.870326 | 5.670872 | 5.857711 | 5.426918 | 2.780247 | 2.828987 | 1.028294 | 1.198205 |

| | | | | | | | | | | | | |
|---|---|---|---|---|---|---|---|---|---|---|---|---|
| 1700019D0 | 14.83462 | 14.46 | 13.69364 | 13.37442 | 8.870991 | 8.78657 | 7.225653 | 6.835052 | 3.07722 | 3.086858 | 2.540419 | 2.710592 |
| Hdgfrp3 | 1.687395 | 1.667372 | 2.629628 | 2.848523 | 3.938383 | 4.004938 | 4.899519 | 5.341909 | 9.038966 | 8.832335 | 13.23954 | 14.03662 |
| Il23a | 1.098931 | 1.149713 | 0.941027 | 1.273611 | 0.830183 | 0.62345 | 0.663984 | 0.655243 | 0.450607 | 0.474082 | 0.541549 | 0.258561 |
| Bhlha15 | 0.876107 | 0.837984 | 1.980753 | 2.034666 | 2.499836 | 2.428389 | 1.982345 | 1.475495 | 0.867675 | 1.017222 | 0.694182 | 0.493384 |
| Gm8300 | 1.483388 | 1.264333 | 6.642368 | 6.055336 | 13.80681 | 13.17584 | 26.86923 | 29.67665 | 27.38521 | 29.00881 | 24.99507 | 23.63425 |
| Phf11a | 0.08952 | 0.170337 | 0.36206 | 0.261373 | 0.610172 | 0.679204 | 1.225771 | 1.280711 | 1.182168 | 1.152259 | 1.310116 | 1.269376 |
| Dusp5 | 4.076442 | 4.731875 | 8.224923 | 8.560763 | 7.435239 | 8.02308 | 8.502988 | 8.14915 | 15.76047 | 14.69261 | 42.93363 | 39.30883 |
| Rhbdf1 | 2.064201 | 2.121881 | 1.725177 | 1.583503 | 2.129656 | 1.98019 | 2.419188 | 2.735218 | 4.926607 | 4.784823 | 8.225061 | 7.71063 |
| Nlrp4f | 5.816282 | 5.886166 | 8.282794 | 7.339145 | 5.87448 | 6.107501 | 5.835707 | 6.471299 | 2.659781 | 2.602754 | 1.444735 | 1.508939 |
| Mical2 | 0.344662 | 0.376833 | 0.307778 | 0.380624 | 0.422797 | 0.394917 | 0.451211 | 0.358408 | 0.587002 | 0.639925 | 1.838371 | 1.741273 |
| Epcam | 57.69132 | 54.27975 | 47.77859 | 44.6692 | 30.86441 | 31.2289 | 23.01241 | 23.2038 | 9.870148 | 9.223489 | 9.930646 | 10.21299 |
| Ldlrad4 | 1.043106 | 1.37277 | 0.761778 | 1.101718 | 0.605625 | 0.50273 | 0.426316 | 0.432388 | 0.221322 | 0.303732 | 0.224088 | 0.227501 |
| Cyp2b23 | 2.968274 | 2.628643 | 2.540386 | 2.23883 | 1.914568 | 2.274442 | 2.588747 | 3.340003 | 1.415386 | 1.666214 | 0.871029 | 0.690086 |
| Prrg4 | 6.119419 | 7.383096 | 5.707088 | 6.010483 | 4.129088 | 3.928686 | 3.643643 | 2.333563 | 1.994017 | 1.971872 | 8.643633 | 9.524109 |
| Plau | 0.128976 | 0.161507 | 0.758694 | 0.604784 | 0.980595 | 0.884675 | 1.54176 | 1.500227 | 1.165843 | 1.142976 | 3.016357 | 2.920581 |
| Fam167a | 1.096385 | 1.17653 | 0.822649 | 0.828567 | 0.776233 | 0.852226 | 0.891703 | 0.919397 | 2.687263 | 2.494761 | 4.023809 | 3.998543 |
| B3gnt5 | 10.08301 | 10.29822 | 7.680963 | 7.625792 | 8.951108 | 9.039702 | 11.0278 | 10.8991 | 19.20485 | 19.64737 | 32.98406 | 36.7784 |
| Hand1 | 0.243547 | 0.343092 | 0.280983 | 0.297279 | 0.465812 | 0.361787 | 0.462973 | 0.319996 | 1.219672 | 1.433619 | 1.813715 | 1.212252 |
| Foxq1 | 0.888716 | 0.853221 | 1.139231 | 0.85361 | 1.100964 | 1.124241 | 1.715936 | 1.430921 | 3.636087 | 2.708476 | 3.74829 | 4.111498 |
| Leprel1 | 0.282324 | 0.320834 | 0.551848 | 0.703204 | 0.975034 | 0.973811 | 1.353724 | 1.427569 | 3.192619 | 3.381211 | 3.244567 | 3.385307 |
| Trp73 | 1.272088 | 1.441624 | 1.06487 | 0.950828 | 0.619135 | 0.76581 | 0.510651 | 0.436881 | 0.160339 | 0.197741 | 0.305445 | 0.296113 |
| BC055111 | 1.18006 | 1.170409 | 1.770076 | 1.421566 | 1.128858 | 1.077014 | 0.838702 | 1.164956 | 0.362626 | 0.540724 | 0.438222 | 0.29589 |
| Greb1 | 0.803381 | 0.903813 | 1.086085 | 1.310204 | 1.958587 | 2.091699 | 2.517443 | 2.489342 | 6.371345 | 6.289369 | 5.770524 | 5.794253 |
| Ddr2 | 0.326134 | 0.309742 | 0.581909 | 0.615886 | 0.670294 | 0.777285 | 0.991901 | 1.193022 | 1.803018 | 1.760572 | 2.803938 | 2.953262 |
| Foxd2 | 0.007722 | 0.033036 | 0.124168 | 0.227236 | 0.225093 | 0.390254 | 0.284379 | 0.343547 | 0.979752 | 0.98425 | 1.10139 | 1.0881 |
| Hs3st3b1 | 1.86475 | 2.334521 | 1.965206 | 2.099988 | 2.530442 | 2.600605 | 3.449095 | 2.729381 | 9.878083 | 10.03944 | 10.13009 | 9.94298 |
| Hsd17b11 | 25.98393 | 25.22267 | 20.68558 | 20.90448 | 12.32487 | 11.97506 | 8.904991 | 9.356635 | 4.390051 | 4.644186 | 4.5149 | 4.376224 |
| Vcl | 15.20711 | 17.32435 | 17.5231 | 16.63555 | 17.34185 | 17.49725 | 22.9519 | 20.50715 | 30.68843 | 29.701 | 77.5029 | 79.37189 |
| AU019990 | 0.0751 | 0.118371 | 0.672177 | 0.511805 | 0.786696 | 0.927025 | 1.812056 | 2.439179 | 1.327388 | 1.327681 | 2.463426 | 1.798749 |
| Kcnh6 | 0.916065 | 1.031989 | 0.985342 | 1.018663 | 0.687334 | 0.983372 | 0.579765 | 0.612278 | 0.173742 | 0.2831 | 0.137937 | 0.214523 |
| Tdgf1 | 568.4683 | 492.5929 | 222.2028 | 219.0617 | 158.8021 | 160.3549 | 98.02087 | 103.9354 | 52.13225 | 58.45358 | 44.72857 | 46.14308 |
| C330021F2 | 6.203416 | 5.394869 | 4.427225 | 4.912166 | 4.960682 | 4.777265 | 3.832247 | 4.279043 | 2.079059 | 2.594951 | 1.230219 | 1.035618 |
| E130309D1 | 2.851329 | 3.157538 | 3.017672 | 2.965141 | 2.575313 | 2.247014 | 1.492156 | 2.002567 | 1.000934 | 1.156558 | 0.582168 | 0.62577 |
| Nrep | 6.811568 | 5.563051 | 4.469955 | 4.412844 | 6.135418 | 6.603351 | 9.042496 | 9.993253 | 20.45194 | 20.90449 | 18.36806 | 20.68044 |
| Klhdc8a | 0.04259 | 0.054458 | 0.584659 | 0.521095 | 0.59429 | 0.670732 | 0.817275 | 0.889256 | 1.298187 | 1.357252 | 2.518662 | 2.465811 |
| Agpat9 | 0.434048 | 0.451034 | 1.001259 | 0.846574 | 0.90911 | 0.774152 | 1.27484 | 1.179721 | 1.702343 | 1.359513 | 3.506067 | 3.654414 |
| 2900041M1 | 1.963802 | 1.895398 | 0.522105 | 0.561672 | 0.485456 | 0.540253 | 0.485113 | 0.604228 | 0.286067 | 0.244476 | 0.188146 | 0.128022 |
| Gm5662 | 1.456053 | 1.52772 | 7.044124 | 6.206726 | 13.75901 | 12.49035 | 23.01809 | 25.67501 | 30.04068 | 29.28536 | 28.21926 | 23.46259 |
| Ror1 | 0.507087 | 0.561589 | 0.447965 | 0.407277 | 0.943812 | 0.94445 | 1.66451 | 1.358348 | 2.354187 | 1.921639 | 1.313954 | 1.260331 |
| Mob3a | 1.401803 | 1.560946 | 1.355924 | 1.090731 | 1.645888 | 1.397487 | 1.871595 | 1.949841 | 2.871844 | 2.561729 | 4.840189 | 5.13532 |
| Frk | 0.234963 | 0.219867 | 0.515647 | 0.474347 | 1.183741 | 1.005185 | 1.649303 | 1.474668 | 2.148721 | 2.230042 | 1.802134 | 2.025049 |
| Gm13078 | 0.124746 | 0.121541 | 0.516156 | 0.554026 | 1.20476 | 1.174801 | 2.13977 | 2.603727 | 2.184192 | 2.24121 | 1.96237 | 1.361819 |
| Aplp1 | 30.5935 | 30.20309 | 23.62225 | 20.74887 | 14.30449 | 13.75153 | 9.554686 | 9.820331 | 6.054332 | 6.261139 | 4.797455 | 4.418472 |
| Scn2b | 0.282759 | 0.369244 | 1.452568 | 1.751199 | 1.681142 | 1.593122 | 1.559529 | 1.345811 | 0.602976 | 0.507063 | 0.352351 | 0.372931 |
| Hdac7 | 1.851771 | 1.948848 | 2.656475 | 2.869431 | 4.179172 | 3.812064 | 4.995939 | 4.746596 | 6.875541 | 6.457056 | 14.42083 | 13.4546 |
| Crygd | 0.160184 | 0.204612 | 0.923627 | 0.841062 | 1.758367 | 1.357305 | 3.12151 | 3.942787 | 3.915398 | 3.646792 | 3.843182 | 3.680581 |
| Mfap2 | 1.107876 | 1.393942 | 1.847375 | 2.057279 | 4.036232 | 3.440655 | 6.211287 | 5.590855 | 10.24204 | 9.543819 | 9.671397 | 9.630649 |
| Slc22a23 | 2.261938 | 2.420296 | 2.466957 | 2.713712 | 4.02766 | 4.158271 | 6.318584 | 5.698126 | 10.83832 | 11.20362 | 12.60065 | 12.69218 |
| Hmgb3 | 22.13214 | 18.31559 | 19.83099 | 20.56035 | 32.11362 | 33.70945 | 46.19248 | 45.68398 | 80.29941 | 70.43111 | 95.30336 | 96.14595 |
| Pcdh1 | 4.192986 | 5.393073 | 3.72164 | 3.876171 | 4.452526 | 3.950877 | 5.620046 | 4.691036 | 6.310662 | 6.283457 | 19.60675 | 18.10706 |
| Slco4a1 | 3.413526 | 3.652462 | 1.553383 | 1.86124 | 1.273482 | 1.236563 | 0.969925 | 0.906812 | 0.508044 | 0.398904 | 0.503625 | 0.578498 |
| Calml4 | 1.848526 | 1.553263 | 3.738044 | 3.236533 | 1.542427 | 1.760934 | 1.531596 | 1.9596 | 0.890844 | 0.893286 | 0.650181 | 0.693811 |
| Sdc3 | 0.860292 | 0.910911 | 0.92363 | 0.973875 | 1.081708 | 1.045132 | 1.415515 | 1.121493 | 2.518587 | 2.323038 | 4.75177 | 4.536808 |
| Thrsp | 1.704389 | 1.334599 | 1.109949 | 1.546065 | 1.654742 | 1.394353 | 0.849199 | 1.064327 | 0.573696 | 0.731137 | 0.316506 | 0.332093 |
| Col4a2 | 9.066352 | 10.25246 | 12.28467 | 12.45668 | 14.23363 | 13.61578 | 19.02293 | 17.40259 | 41.30759 | 39.11542 | 56.24761 | 57.98451 |
| Hmga2 | 15.29082 | 15.7269 | 15.08228 | 16.28886 | 18.26971 | 17.18512 | 23.24408 | 22.59707 | 77.38791 | 75.69894 | 59.83822 | 62.98014 |
| Kif5c | 2.859493 | 3.094209 | 4.154368 | 3.961071 | 4.800093 | 4.379453 | 5.689603 | 5.796863 | 10.12394 | 10.65005 | 17.64255 | 18.407 |
| Krt20 | 0.114982 | 0.033208 | 1.602721 | 1.588174 | 2.622101 | 3.167089 | 2.638582 | 2.86991 | 1.032051 | 0.707741 | 2.876561 | 3.286904 |
| Bmp1 | 0.900391 | 1.217012 | 2.263626 | 2.354204 | 3.274553 | 3.438656 | 5.689116 | 4.802723 | 8.625862 | 7.7205 | 10.36295 | 10.92757 |
| Olfr376 | 0.066629 | 0.035003 | 0.288885 | 0.349445 | 0.600639 | 0.405391 | 1.364754 | 1.045795 | 1.234397 | 1.002669 | 1.555938 | 1.62168 |
| Chst1 | 0.466697 | 0.458301 | 0.615574 | 0.463806 | 1.404943 | 1.353464 | 2.128525 | 2.15099 | 1.605531 | 1.568037 | 1.016109 | 0.958855 |

| | | | | | | | | | | | | |
|---|---|---|---|---|---|---|---|---|---|---|---|---|
| Relb | 2.369714 | 3.036595 | 2.310513 | 2.212973 | 2.363327 | 2.122424 | 1.806154 | 2.434282 | 1.847517 | 1.710967 | 8.988306 | 7.933491 |
| Rarg | 54.53645 | 55.55833 | 63.65929 | 57.5085 | 45.54035 | 44.74784 | 34.03899 | 33.80525 | 15.66246 | 14.54839 | 14.14005 | 12.40809 |
| Arsi | 0.190072 | 0.212306 | 0.278699 | 0.29377 | 0.413053 | 0.403657 | 0.604546 | 0.488363 | 0.808767 | 0.912826 | 1.630688 | 1.357923 |
| Erbb2ip | 6.172519 | 6.693055 | 7.063294 | 6.817435 | 8.629264 | 8.415812 | 10.17622 | 9.772255 | 21.14052 | 21.31249 | 28.62877 | 31.43963 |
| Cox8c | 0.506366 | 0.405245 | 1.795601 | 1.953196 | 2.411035 | 1.941895 | 3.02962 | 1.969205 | 1.199499 | 1.128607 | 0.818212 | 0.427162 |
| Pygo1 | 0.315548 | 0.325205 | 0.629518 | 0.546289 | 0.875397 | 0.784754 | 1.062517 | 1.012414 | 2.04497 | 1.994999 | 2.361846 | 2.517612 |
| Casq1 | 0.621938 | 0.553338 | 1.634066 | 1.600071 | 1.102535 | 1.256311 | 0.957118 | 0.989614 | 0.593669 | 0.522684 | 0.583129 | 0.347779 |
| Itga7 | 0.301698 | 0.311459 | 0.44987 | 0.424742 | 0.652531 | 0.389316 | 0.649057 | 0.572262 | 0.627538 | 0.688593 | 2.027568 | 1.790847 |
| Epo | 0.211349 | 0.074844 | 0.790258 | 1.402349 | 1.55254 | 1.929264 | 1.964095 | 1.808905 | 1.034209 | 1.342363 | 0.628497 | 0.419711 |
| Rnf122 | 2.121617 | 2.278195 | 1.415454 | 1.28189 | 2.21179 | 2.369498 | 3.175685 | 3.305783 | 6.285971 | 5.88823 | 4.724951 | 4.279814 |
| Zadh2 | 1.268814 | 0.977828 | 8.791445 | 8.623879 | 23.04773 | 24.55347 | 42.03196 | 39.59923 | 31.1081 | 31.30237 | 26.44723 | 28.04758 |
| Pdgfc | 0.710213 | 0.849994 | 0.501833 | 0.476297 | 0.383614 | 0.427166 | 0.652996 | 0.572362 | 2.563521 | 1.961415 | 1.218941 | 1.308575 |
| BC048502 | 0.504901 | 0.882248 | 2.555939 | 4.171106 | 2.552538 | 2.139429 | 2.307679 | 1.958235 | 0.68543 | 1.239568 | 1.258483 | 0.908739 |
| Pogk | 0.558011 | 0.665924 | 1.041707 | 1.16274 | 1.418561 | 1.189531 | 1.628694 | 1.730164 | 3.632838 | 3.80736 | 5.283624 | 5.334611 |
| Pxdn | 5.33545 | 6.157383 | 7.624068 | 8.363423 | 13.09254 | 11.90311 | 19.26964 | 15.80896 | 32.68315 | 32.50162 | 38.03041 | 38.34836 |
| Pou5f1 | 261.0595 | 206.5594 | 162.0745 | 143.9489 | 101.2333 | 102.6432 | 63.48587 | 74.70518 | 35.22717 | 31.42673 | 37.46669 | 33.98454 |
| Ndn | 7.6088 | 6.872247 | 13.30597 | 11.94746 | 18.52335 | 18.05565 | 25.94492 | 29.90188 | 40.64554 | 41.15742 | 59.4743 | 54.61378 |
| Gadd45g | 4.686899 | 4.790573 | 14.02647 | 10.69206 | 12.03661 | 12.84491 | 18.19021 | 20.43598 | 22.09123 | 17.92211 | 51.14757 | 48.8011 |
| Slc18a3 | 0.12512 | 0.209861 | 0.212878 | 0.353481 | 0.609596 | 0.650058 | 1.741835 | 1.612088 | 1.567124 | 1.366049 | 1.094692 | 1.01295 |
| Myh10 | 19.36781 | 24.08926 | 29.89891 | 30.66041 | 32.66212 | 32.00342 | 38.97373 | 36.08122 | 91.46281 | 94.67611 | 136.7746 | 139.7747 |
| L1td1 | 259.1093 | 267.359 | 202.4406 | 191.1941 | 131.786 | 129.2765 | 80.06061 | 92.55533 | 41.89101 | 43.04083 | 42.56477 | 41.94034 |
| Ms4a4d | 2.435532 | 2.099471 | 1.466508 | 1.749396 | 3.621507 | 3.978612 | 7.478317 | 7.9609 | 6.440022 | 6.295269 | 3.370187 | 3.847857 |
| Hck | 9.49431 | 10.0573 | 1.654608 | 1.375048 | 1.020056 | 0.726242 | 0.60437 | 0.67209 | 0.26566 | 0.302276 | 0.619393 | 0.555307 |
| Gfra3 | 0.334109 | 0.457358 | 2.065093 | 2.147553 | 2.4466 | 2.283379 | 5.335999 | 5.905442 | 12.71646 | 9.756833 | 9.199563 | 8.999163 |
| Afap1l2 | 0.930072 | 0.866491 | 0.813165 | 0.858041 | 1.065327 | 1.211364 | 1.829654 | 1.380323 | 3.203086 | 3.021212 | 3.602337 | 3.895515 |
| Plekhg6 | 0.341797 | 0.623913 | 1.086071 | 1.092715 | 0.818366 | 0.664587 | 0.572652 | 0.497226 | 0.308786 | 0.240941 | 0.50884 | 0.376489 |
| Hpn | 3.778544 | 4.097993 | 4.669462 | 4.99885 | 5.116568 | 4.482943 | 4.171435 | 4.054595 | 2.147625 | 2.260366 | 1.681924 | 1.103007 |
| Spry1 | 9.372684 | 8.717006 | 5.43042 | 5.108892 | 5.970736 | 5.912064 | 6.851967 | 7.233723 | 16.0059 | 15.84456 | 23.03255 | 23.13702 |
| Hoxb3 | 0.144929 | 0.145467 | 1.92832 | 2.060962 | 3.617331 | 3.940476 | 4.91804 | 4.436947 | 10.01789 | 8.574696 | 8.892456 | 9.332855 |
| Pld4 | 0.86578 | 0.632826 | 1.726271 | 1.713258 | 2.076633 | 1.96512 | 1.789955 | 2.416806 | 0.969134 | 1.097371 | 0.48814 | 0.534112 |
| Atg9b | 1.374211 | 1.234784 | 1.44754 | 1.294034 | 0.882617 | 0.907543 | 0.71256 | 0.636585 | 0.384579 | 0.286526 | 0.414728 | 0.387178 |
| Cpt1a | 24.42001 | 26.60139 | 20.36564 | 19.383 | 14.5935 | 14.69365 | 11.34612 | 10.55054 | 4.955259 | 4.830615 | 4.562731 | 4.292589 |
| Serpinb6c | 5.834767 | 5.404334 | 1.341184 | 1.062333 | 1.245021 | 0.918808 | 1.557131 | 1.448041 | 2.659986 | 2.978688 | 3.702823 | 4.145165 |
| Chchd10 | 287.159 | 307.2378 | 297.4945 | 261.89 | 190.8644 | 192.5315 | 137.9008 | 107.8093 | 70.5397 | 58.05715 | 72.86566 | 58.19597 |
| Hist2h4 | 2.875562 | 2.27637 | 1.101039 | 2.399103 | 2.628294 | 3.865828 | 1.261618 | 2.875282 | 1.120199 | 1.593208 | 1.180677 | 0.857608 |
| Gli1 | 7.510928 | 7.891459 | 5.804007 | 5.994585 | 4.357757 | 4.351806 | 2.810386 | 3.069059 | 1.975568 | 1.776736 | 1.594952 | 1.330977 |
| Dll1 | 1.846513 | 1.615061 | 2.71592 | 2.251123 | 2.305969 | 2.394617 | 3.427239 | 3.507506 | 6.302977 | 5.806172 | 9.701541 | 10.13851 |
| Lonrf2 | 0.471464 | 0.520843 | 0.440731 | 0.529928 | 0.615188 | 0.590388 | 0.700021 | 0.670914 | 1.505975 | 1.54397 | 2.78163 | 2.386337 |
| Palld | 6.783007 | 7.539242 | 10.70324 | 10.16318 | 8.724445 | 7.82969 | 8.642083 | 7.036263 | 13.52331 | 13.07646 | 31.26997 | 31.67413 |
| G730013B( | 0.999721 | 1.336453 | 0.979511 | 0.7707 | 0.499783 | 0.36475 | 0.39098 | 0.258458 | 0.163466 | 0.266007 | 0.154252 | 0.171778 |
| Spz1 | 0.112625 | 0.040521 | 0.401803 | 0.450046 | 0.867399 | 0.977189 | 1.724216 | 2.01862 | 1.703225 | 1.776951 | 1.658053 | 1.229526 |
| Gm867 | 1.832195 | 2.084267 | 1.942904 | 1.658976 | 1.005917 | 0.928097 | 0.825288 | 0.92122 | 0.321138 | 0.409135 | 0.27934 | 0.369889 |
| Tmtc1 | 1.235621 | 1.195961 | 7.932556 | 8.039625 | 6.374594 | 6.135634 | 4.960648 | 4.578808 | 1.830388 | 1.792592 | 1.918453 | 1.871989 |
| Peli3 | 1.079831 | 1.301272 | 1.303266 | 1.037645 | 1.058154 | 0.988516 | 1.089921 | 1.411659 | 0.594547 | 0.575184 | 0.147835 | 0.314909 |
| Zc3h12c | 0.957178 | 0.980843 | 1.405437 | 1.401125 | 1.857344 | 1.908884 | 2.104725 | 1.536 | 4.031826 | 4.074322 | 5.781862 | 6.276042 |
| Osbpl5 | 0.629932 | 0.614815 | 0.982468 | 1.053339 | 1.291402 | 1.309731 | 1.880301 | 1.912703 | 2.607895 | 2.643293 | 4.454401 | 4.706328 |
| Sned1 | 1.431455 | 1.713457 | 1.732507 | 1.957067 | 1.270546 | 1.363502 | 1.065652 | 1.045248 | 0.615261 | 0.63343 | 0.374785 | 0.43808 |
| Aif1l | 4.143823 | 4.516636 | 6.419896 | 6.615807 | 9.453631 | 8.729179 | 10.64886 | 10.59046 | 19.49363 | 19.03055 | 29.36098 | 29.53629 |
| Tmem123 | 18.82423 | 16.8001 | 13.31722 | 13.49185 | 16.2551 | 16.92402 | 20.49056 | 21.16838 | 39.58001 | 40.072 | 55.58027 | 60.22795 |
| Pramel6 | 0.486549 | 0.465536 | 1.412172 | 1.04924 | 1.515884 | 1.239831 | 2.184089 | 1.789423 | 3.370969 | 4.683722 | 2.674137 | 2.373876 |
| Mn1 | 0.176489 | 0.324872 | 0.45456 | 0.533332 | 0.490055 | 0.499164 | 0.850462 | 0.676362 | 2.167669 | 2.044314 | 2.667181 | 2.224395 |
| Ggt1 | 1.347691 | 1.536599 | 3.292055 | 2.527427 | 2.081984 | 1.698595 | 1.407927 | 1.513935 | 0.702277 | 1.073916 | 0.596705 | 0.567338 |
| Ptch2 | 5.698737 | 6.014109 | 7.012527 | 6.335861 | 4.801775 | 4.53556 | 3.757037 | 3.572252 | 1.618712 | 1.497784 | 1.501668 | 1.424745 |
| Lrrc4b | 0.162646 | 0.27254 | 0.251584 | 0.251035 | 0.435758 | 0.370799 | 0.607373 | 0.755921 | 1.015755 | 1.115337 | 0.792599 | 0.587812 |
| Flrt3 | 1.804205 | 2.023881 | 2.529109 | 2.925928 | 3.636553 | 3.701461 | 6.461101 | 6.05725 | 13.37955 | 12.98826 | 8.757144 | 10.31731 |
| Spaca1 | 0.185105 | 0.140761 | 0.30983 | 0.33077 | 0.521753 | 0.522865 | 0.827257 | 1.213768 | 1.002957 | 1.054874 | 1.585125 | 1.467347 |
| Col4a5 | 0.231643 | 0.230675 | 0.38774 | 0.415809 | 0.705641 | 0.660187 | 1.002823 | 1.082762 | 1.879172 | 1.844578 | 1.349201 | 1.442461 |
| 1700095A2 | 0.755219 | 1.594667 | 1.469485 | 1.760402 | 1.034753 | 1.464413 | 0.990888 | 0.745314 | 0.277909 | 0.396847 | 0.344043 | 0.521079 |
| Ctnnd2 | 0.399146 | 0.477256 | 0.391401 | 0.383497 | 0.605722 | 0.498645 | 0.849591 | 0.824176 | 1.308385 | 1.205304 | 1.590479 | 1.701016 |
| Fam171a1 | 1.982557 | 2.069894 | 3.099805 | 3.042021 | 4.723587 | 4.513335 | 6.622345 | 6.68723 | 11.14825 | 11.30234 | 13.10263 | 13.49102 |
| Wfdc1 | 0.314675 | 0.070593 | 0.261817 | 0.331255 | 0.997365 | 0.90133 | 0.785914 | 1.468219 | 0.616497 | 0.555626 | 0.39512 | 0.415844 |

| | | | | | | | | | | | | |
|---|---|---|---|---|---|---|---|---|---|---|---|---|
| Bcas1 | 1.978151 | 1.646676 | 1.611569 | 1.653335 | 1.957915 | 1.790863 | 1.664969 | 2.048054 | 0.777454 | 0.817244 | 0.568653 | 0.462467 |
| Fos | 0.68908 | 0.462985 | 0.721936 | 0.669676 | 0.621641 | 0.60178 | 1.092385 | 0.97825 | 1.02733 | 1.140674 | 3.267957 | 2.664131 |
| Gm13119 | 0.558932 | 0.680877 | 2.359438 | 2.457765 | 3.93605 | 4.646736 | 8.491237 | 9.288368 | 8.248854 | 10.8775 | 8.550807 | 7.1395 |
| Kis2 | 1.101334 | 1.458285 | 0.752273 | 1.050807 | 0.893724 | 1.165067 | 0.948728 | 1.364796 | 0.426097 | 0.601526 | 0.182184 | 0.308376 |
| Plch1 | 2.965548 | 3.030522 | 2.822857 | 3.056047 | 3.493973 | 3.69503 | 4.715387 | 4.358068 | 12.84786 | 13.50531 | 8.152647 | 8.672632 |
| Gipc2 | 0.946717 | 0.689966 | 0.954869 | 1.167177 | 1.349751 | 1.166629 | 0.787358 | 0.646038 | 0.454639 | 0.445888 | 0.314614 | 0.264157 |
| Fam115a | 2.557005 | 2.494593 | 3.508993 | 3.190288 | 5.570665 | 5.277539 | 7.760435 | 7.695262 | 12.17837 | 14.04554 | 9.334563 | 9.795904 |
| Sgms2 | 0.500372 | 0.586222 | 0.423643 | 0.404539 | 0.694297 | 0.514228 | 0.458135 | 0.678648 | 0.599617 | 0.536147 | 1.588956 | 1.780387 |
| Stc2 | 2.79332 | 2.657762 | 2.403353 | 2.151094 | 1.532003 | 1.381657 | 0.649728 | 0.647335 | 0.545359 | 0.73974 | 2.617943 | 2.846403 |
| Crygc | 0.129239 | 0.054091 | 2.645904 | 2.301056 | 4.398947 | 4.958786 | 8.602683 | 10.11563 | 5.747212 | 3.8582 | 3.622256 | 4.413034 |
| Chrm4 | 0.673393 | 0.722837 | 1.749552 | 1.435862 | 3.062803 | 3.133942 | 4.665591 | 4.881989 | 5.011413 | 6.296346 | 3.754501 | 3.351219 |
| Msrb3 | 0.598693 | 0.572126 | 2.084839 | 2.109241 | 3.002149 | 3.192541 | 3.531465 | 2.975688 | 4.277084 | 4.728341 | 8.48878 | 9.248023 |
| Gbp9 | 0.677835 | 0.641187 | 2.115006 | 1.865191 | 1.25846 | 1.379366 | 1.337105 | 1.302939 | 0.40887 | 0.57398 | 0.330025 | 0.425485 |
| Gadd45b | 8.734112 | 7.711613 | 6.890955 | 6.486812 | 6.284547 | 6.180453 | 5.104702 | 6.737607 | 5.363926 | 5.037285 | 22.70086 | 22.0791 |
| Alpk3 | 2.527631 | 3.507469 | 4.861908 | 4.477854 | 2.947145 | 2.764261 | 1.954575 | 2.045698 | 1.712338 | 1.947347 | 1.129523 | 1.021611 |
| Lmx1a | 0.108228 | 0.101233 | 0.285931 | 0.305822 | 0.59596 | 0.531527 | 1.287419 | 1.036201 | 1.196216 | 1.186895 | 1.660544 | 1.339061 |
| Megf9 | 0.602954 | 0.674278 | 0.973994 | 0.96803 | 1.10041 | 1.167125 | 1.243454 | 1.378096 | 2.776202 | 2.754699 | 3.547622 | 4.23479 |
| F11r | 12.63987 | 13.05502 | 15.01062 | 14.17447 | 19.06612 | 19.17617 | 36.86053 | 32.9447 | 49.42422 | 43.2832 | 56.02384 | 61.98745 |
| Hs3st1 | 1.649895 | 1.506688 | 1.341317 | 1.361149 | 1.745266 | 1.263977 | 2.163252 | 1.973644 | 4.874691 | 3.966864 | 5.893496 | 5.526222 |
| Gm2016 | 1.722658 | 1.374324 | 6.455828 | 6.197404 | 10.72573 | 11.03641 | 22.3532 | 22.87624 | 26.30358 | 27.0191 | 24.87431 | 21.96144 |
| Rprm | 4.320701 | 3.66365 | 2.75626 | 2.70246 | 4.427552 | 4.012673 | 5.959631 | 6.773257 | 13.4534 | 11.77402 | 9.701173 | 9.706068 |
| Ap1m2 | 4.542781 | 4.287643 | 4.017571 | 3.597382 | 2.167824 | 2.130489 | 1.242661 | 1.644932 | 1.109404 | 0.923855 | 0.6736 | 0.8258 |
| Nt5e | 0.580437 | 0.451838 | 3.810433 | 3.833815 | 4.055765 | 4.306655 | 4.181716 | 4.070809 | 1.139547 | 1.15809 | 0.793284 | 0.989906 |
| Lhfpl2 | 1.117265 | 1.067156 | 1.569657 | 1.459422 | 2.222503 | 2.144486 | 2.942148 | 2.953446 | 6.029438 | 6.346098 | 5.309535 | 5.42723 |
| Maff | 5.137419 | 3.44735 | 2.590141 | 2.883974 | 3.115307 | 2.880427 | 2.335988 | 2.594298 | 2.945279 | 2.950357 | 10.88192 | 11.27842 |
| Syt3 | 1.680521 | 1.788172 | 2.527379 | 2.549489 | 2.454986 | 2.09133 | 2.300741 | 2.069502 | 1.332851 | 1.231478 | 0.666121 | 0.586528 |
| Bmpr1b | 0.301602 | 0.19795 | 0.278931 | 0.264911 | 0.485653 | 0.358186 | 0.671808 | 0.604567 | 1.221208 | 1.016496 | 1.176201 | 1.151259 |
| Cbr3 | 26.54026 | 24.25081 | 10.94137 | 8.957993 | 7.296793 | 7.877849 | 6.10234 | 6.92992 | 3.225877 | 2.985302 | 2.271245 | 2.061298 |
| Slc9b2 | 2.768737 | 3.31178 | 1.339607 | 1.301217 | 0.922277 | 0.991345 | 0.783247 | 0.667826 | 0.358471 | 0.299597 | 0.231697 | 0.369045 |
| Hipk3 | 2.646948 | 3.206042 | 4.501927 | 4.646386 | 5.408835 | 5.316127 | 6.67626 | 5.973133 | 10.38163 | 10.68967 | 19.41667 | 20.14836 |
| Pdcl2 | 2.029133 | 1.984495 | 0.89336 | 1.300434 | 1.347339 | 1.215882 | 1.417751 | 1.547685 | 0.799209 | 0.929893 | 0.727741 | 0.35698 |
| Cdh1 | 68.95344 | 84.70376 | 97.29344 | 93.78426 | 53.6104 | 52.89024 | 33.34505 | 33.24072 | 20.37325 | 21.63562 | 25.7452 | 22.94668 |
| Sema3b | 0.763645 | 0.813712 | 0.717723 | 0.705783 | 0.973534 | 0.810582 | 0.993941 | 0.84642 | 1.170623 | 1.319385 | 3.229103 | 3.058982 |
| Zfp9 | 0.106564 | 0.138526 | 0.248165 | 0.295035 | 0.527945 | 0.611493 | 0.910719 | 0.720487 | 1.211482 | 1.277877 | 1.010599 | 1.029563 |
| Setbp1 | 1.314111 | 1.73085 | 1.360711 | 1.624256 | 1.868042 | 2.036404 | 2.888797 | 2.368547 | 5.401225 | 5.607478 | 6.935396 | 7.028457 |
| Plcg2 | 13.67379 | 16.58323 | 23.13966 | 21.49783 | 16.7319 | 15.71652 | 12.99368 | 13.41865 | 8.253267 | 9.156271 | 4.605999 | 4.970055 |
| Mmel1 | 0.941999 | 1.090284 | 0.949881 | 1.0897 | 1.119407 | 0.973455 | 0.800948 | 1.101816 | 0.47339 | 0.574757 | 0.414977 | 0.254769 |
| Shd | 0.585624 | 0.490376 | 0.985217 | 1.049553 | 2.171597 | 2.600788 | 3.031635 | 3.418968 | 5.085888 | 4.536617 | 4.184613 | 4.114714 |
| Gbx2 | 37.56435 | 33.89538 | 41.53944 | 40.51272 | 57.5635 | 62.80793 | 65.45296 | 58.36983 | 35.05087 | 30.91703 | 14.08229 | 14.53423 |
| BC028528 | 1.395182 | 0.86553 | 0.545372 | 0.688215 | 0.099595 | 0.359256 | 0.30944 | 0.347593 | 0.196101 | 0.159277 | 0.312011 | 0.198606 |
| Lcp1 | 0.620067 | 0.496919 | 0.569897 | 0.625995 | 0.603431 | 0.627632 | 0.652873 | 0.719906 | 1.349927 | 1.53326 | 2.960587 | 2.702588 |
| Zcwpw1 | 8.162502 | 7.642819 | 11.99437 | 12.48293 | 7.431466 | 7.661377 | 5.011151 | 6.833905 | 3.788467 | 4.024284 | 3.212673 | 2.891974 |
| Hivep2 | 0.307625 | 0.296809 | 0.7081 | 0.920505 | 0.852747 | 0.800847 | 0.916293 | 0.775882 | 1.056887 | 1.204092 | 3.094016 | 3.347191 |
| Dusp10 | 0.650008 | 0.76193 | 0.626755 | 0.575466 | 0.908676 | 0.660301 | 0.915424 | 1.057531 | 1.667876 | 1.350361 | 2.597283 | 2.481521 |
| Mylk | 0.180434 | 0.11609 | 0.208676 | 0.26452 | 0.435471 | 0.330572 | 0.475419 | 0.646938 | 0.989645 | 1.140576 | 0.955049 | 0.961204 |
| Pramel7 | 0.585291 | 0.754993 | 1.153327 | 1.341109 | 1.320616 | 1.405559 | 2.613329 | 3.022058 | 5.372944 | 5.00644 | 5.337221 | 5.773586 |
| Msx2 | 0.279366 | 0.242099 | 0.315416 | 0.334355 | 0.467774 | 0.419394 | 0.685724 | 0.363015 | 1.235082 | 0.915583 | 1.443627 | 1.438212 |
| Jak2 | 1.84694 | 1.761094 | 2.839097 | 2.851757 | 3.996108 | 3.818731 | 5.237553 | 5.562555 | 7.833592 | 8.462459 | 11.27384 | 12.25837 |
| Wls | 2.464186 | 2.327031 | 4.914418 | 5.249554 | 7.973484 | 7.919582 | 14.05793 | 13.98637 | 23.61502 | 22.56422 | 17.57573 | 19.30856 |
| Uba1y | 0.582688 | 0.650593 | 1.18538 | 1.145814 | 0.829607 | 0.919701 | 0.848547 | 1.024224 | 0.400817 | 0.44167 | 0.382185 | 0.26663 |
| Klb | 0.686346 | 0.83713 | 0.66146 | 0.612266 | 0.685877 | 0.557952 | 0.922451 | 1.014732 | 2.164353 | 2.055449 | 1.909236 | 2.394751 |
| Bin1 | 3.589115 | 3.587674 | 4.924632 | 5.115298 | 8.245263 | 7.886357 | 12.64964 | 13.86317 | 19.02383 | 18.99377 | 22.82101 | 21.9462 |
| Pramef6 | 0.132854 | 0.140661 | 0.308826 | 0.250004 | 0.583762 | 0.511283 | 1.17242 | 1.072201 | 1.091717 | 1.047102 | 1.025433 | 0.958086 |
| Agap2 | 0.193316 | 0.222214 | 1.68418 | 1.6894 | 1.755865 | 1.94708 | 1.615337 | 1.431056 | 0.635758 | 0.596813 | 0.450787 | 0.455248 |
| Nfatc1 | 0.11135 | 0.09478 | 0.510224 | 0.501067 | 0.975128 | 0.809017 | 1.368669 | 1.119986 | 1.549254 | 1.518251 | 2.173657 | 2.138198 |
| Tnfrsf21 | 5.678569 | 6.909115 | 7.475297 | 7.382583 | 12.66903 | 12.60867 | 23.11856 | 18.61241 | 27.23362 | 27.12307 | 30.40476 | 31.49502 |
| Dnase2a | 4.420105 | 4.063015 | 5.846916 | 5.332795 | 4.703223 | 4.62658 | 3.411139 | 3.544086 | 2.331227 | 2.290379 | 1.449629 | 1.250141 |
| Dock6 | 30.65097 | 35.73824 | 62.9544 | 61.61058 | 48.39732 | 47.26293 | 38.97293 | 40.84786 | 21.0902 | 20.7938 | 15.83692 | 14.44431 |
| Itgal | 0.864202 | 0.91879 | 1.736326 | 1.908006 | 1.300028 | 1.09657 | 0.927722 | 1.068017 | 0.692804 | 0.747418 | 0.439707 | 0.447819 |
| Dok5 | 0.775523 | 0.944937 | 0.534068 | 0.57119 | 1.184661 | 1.085207 | 1.603379 | 1.882106 | 2.12911 | 2.432974 | 1.089173 | 1.209185 |
| Gcm1 | 0.298098 | 0.439091 | 3.209734 | 3.065195 | 2.266392 | 1.976018 | 1.692004 | 1.935514 | 1.181455 | 1.339078 | 0.817305 | 0.720394 |

| | | | | | | | | | | | | |
|---|---|---|---|---|---|---|---|---|---|---|---|---|
| Triml1 | 45.52903 | 42.0061 | 22.36276 | 21.75026 | 16.191 | 15.93399 | 10.97986 | 10.74143 | 7.183213 | 8.382419 | 4.813989 | 5.113577 |
| Tbc1d9 | 3.519424 | 4.01956 | 1.958815 | 1.766076 | 1.714433 | 1.625609 | 1.447659 | 1.502488 | 3.687509 | 3.763756 | 5.922294 | 6.385683 |
| Arl5c | 2.103711 | 2.28013 | 2.2644 | 2.171739 | 2.74246 | 2.159534 | 1.929184 | 1.909289 | 1.070445 | 1.352624 | 0.702576 | 0.511152 |
| Thbs2 | 0.388701 | 0.361714 | 0.383673 | 0.372094 | 0.27536 | 0.338059 | 0.44807 | 0.46693 | 0.627803 | 0.54962 | 1.372068 | 1.436012 |
| Pitx2 | 6.230139 | 5.321587 | 9.712549 | 10.47604 | 17.78499 | 17.90942 | 16.7614 | 18.52923 | 9.345037 | 9.708402 | 4.562136 | 4.362749 |
| Pou2f2 | 1.048977 | 1.44669 | 2.500039 | 2.626283 | 1.742727 | 1.486486 | 1.069866 | 1.274891 | 0.806713 | 0.631682 | 0.539602 | 0.618565 |
| Fam159a | 2.582216 | 2.422128 | 3.626675 | 2.790245 | 3.907197 | 4.615496 | 4.093646 | 3.793626 | 2.389208 | 2.350046 | 0.903556 | 1.087415 |
| Gm4858 | 0.137679 | 0.028184 | 0.289997 | 0.291266 | 0.520845 | 0.53751 | 0.544612 | 1.020236 | 0.932233 | 1.086995 | 1.352114 | 1.235711 |
| Lpar6 | 1.448409 | 1.543403 | 2.022493 | 1.915108 | 2.790761 | 2.851824 | 3.777518 | 3.957665 | 6.543713 | 5.960109 | 7.142972 | 8.100547 |
| Bmyc | 0.410646 | 0.362944 | 0.830451 | 0.842891 | 1.208277 | 1.610064 | 0.678217 | 0.381421 | 0.605464 | 0.564659 | 0.85778 | 0.694319 |
| Igsf11 | 0.210987 | 0.160453 | 0.477915 | 0.452172 | 0.631497 | 0.772557 | 1.009489 | 0.939248 | 1.81858 | 1.907956 | 1.195056 | 1.430731 |
| Plekha6 | 0.496632 | 0.508657 | 3.230338 | 3.522644 | 2.592473 | 2.715027 | 2.427658 | 2.607251 | 1.205619 | 1.130643 | 1.012247 | 0.835187 |
| Pkia | 1.416371 | 1.053348 | 1.012179 | 1.188084 | 1.262318 | 1.361801 | 1.667945 | 1.539755 | 3.765929 | 3.949991 | 4.420157 | 4.999685 |
| Chad | 0.23978 | 0.544493 | 0.511289 | 0.725038 | 0.706865 | 0.709213 | 0.762515 | 0.334765 | 1.241541 | 1.407605 | 1.16789 | 1.075422 |
| Il10rb | 0.332093 | 0.433974 | 0.40475 | 0.283018 | 0.591809 | 0.61429 | 0.820802 | 0.839611 | 0.993186 | 0.867219 | 1.085419 | 1.18762 |
| Lars2 | 926.9473 | 1317.234 | 423.7109 | 981.8392 | 713.1387 | 1296.655 | 677.1098 | 694.0822 | 463.9249 | 560.8119 | 264.7824 | 309.1292 |
| Hspa1b | 1.417303 | 1.691842 | 2.777017 | 3.149162 | 5.93724 | 5.082209 | 11.82407 | 13.20906 | 8.239433 | 8.211548 | 7.75753 | 5.625426 |
| Ube2ql1 | 0.180875 | 0.180602 | 0.298355 | 0.302694 | 0.578313 | 0.598065 | 0.632547 | 0.559816 | 0.482853 | 0.52756 | 1.545832 | 1.266265 |
| Cald1 | 13.58916 | 14.08283 | 16.03774 | 17.09212 | 19.08382 | 19.34704 | 25.99131 | 25.51838 | 40.18392 | 40.22166 | 69.25669 | 71.38712 |
| Ncam1 | 3.254332 | 4.620987 | 2.944305 | 3.044896 | 4.232627 | 3.682191 | 7.117367 | 5.488706 | 11.95145 | 12.66748 | 11.09675 | 11.19723 |
| Klf15 | 7.297115 | 6.696752 | 3.446736 | 2.8311 | 2.060507 | 2.467777 | 1.685557 | 1.678055 | 1.587308 | 1.470415 | 0.884693 | 0.680634 |
| Sult6b1 | 3.622171 | 2.903868 | 3.445464 | 3.296364 | 2.610737 | 2.812399 | 1.733179 | 1.501567 | 0.961926 | 0.952654 | 1.010893 | 0.793088 |
| Faxc | 0.949205 | 1.072496 | 0.995787 | 1.049095 | 1.321392 | 1.344513 | 1.706165 | 1.438027 | 3.628568 | 3.932883 | 4.189229 | 4.359866 |
| Asap3 | 3.578211 | 4.112746 | 3.58384 | 3.502482 | 2.787348 | 2.552522 | 2.387552 | 2.262932 | 1.254791 | 1.0134 | 0.867756 | 0.843343 |
| Junb | 11.36257 | 9.999338 | 12.95771 | 12.48168 | 10.66416 | 10.28129 | 9.426698 | 8.863993 | 7.937582 | 6.618065 | 29.63025 | 27.46625 |
| Ctgf | 6.261046 | 5.748592 | 12.59515 | 11.60819 | 12.68029 | 13.27567 | 12.37306 | 12.23438 | 25.39192 | 24.45541 | 43.08726 | 48.12713 |
| Slc28a1 | 3.858898 | 4.705113 | 3.216628 | 2.909844 | 1.31948 | 1.065551 | 0.802477 | 0.792165 | 0.583538 | 0.758303 | 0.774957 | 0.702001 |
| Flrt2 | 0.181865 | 0.209983 | 0.443161 | 0.309573 | 0.428328 | 0.440977 | 0.507961 | 0.435727 | 0.961898 | 0.902463 | 1.252454 | 1.283125 |
| Nodal | 16.88031 | 15.34619 | 23.34602 | 22.90537 | 23.62908 | 23.26277 | 15.1191 | 16.06232 | 6.046957 | 6.233578 | 6.171763 | 5.614143 |
| Gpsm3 | 1.786233 | 2.032754 | 4.510455 | 4.419248 | 3.581577 | 3.557501 | 2.877764 | 3.210553 | 1.282124 | 1.067175 | 1.328921 | 1.152722 |
| Ddr1 | 5.823119 | 6.50663 | 6.850951 | 6.369168 | 6.783951 | 6.427988 | 9.196134 | 8.819336 | 15.18453 | 13.51235 | 28.23719 | 26.36199 |
| Insl3 | 0.777792 | 1.497104 | 2.340115 | 1.64841 | 1.419378 | 1.100656 | 1.074586 | 2.106386 | 0.313088 | 0.509098 | 0.704685 | 0.668725 |
| Sh3bgrl | 12.00411 | 11.54418 | 12.48391 | 12.23935 | 18.11147 | 18.18097 | 22.10023 | 25.98668 | 49.68141 | 50.59999 | 37.26798 | 45.83306 |
| Naprt1 | 5.722351 | 5.817375 | 10.20719 | 8.490406 | 6.626225 | 5.731735 | 5.34575 | 5.207454 | 2.791483 | 2.949304 | 2.904612 | 2.053867 |
| Asb13 | 1.800504 | 1.663446 | 1.781364 | 1.293258 | 2.043544 | 1.861347 | 2.536877 | 3.02054 | 4.995514 | 5.340946 | 5.378989 | 5.182419 |
| Aire | 11.80211 | 10.32382 | 1.164173 | 1.390739 | 0.930381 | 0.998876 | 0.505735 | 0.616256 | 0.637807 | 0.71915 | 0.385843 | 0.336777 |
| Lama1 | 12.53833 | 17.08879 | 21.36745 | 22.31111 | 20.57335 | 20.17606 | 31.94054 | 25.7183 | 81.14489 | 79.0485 | 76.48212 | 83.24452 |
| Ikzf1 | 0.179737 | 0.253049 | 0.386818 | 0.383052 | 0.647678 | 0.705506 | 1.016124 | 0.898672 | 1.37579 | 1.579854 | 0.814933 | 0.779114 |
| Ssbp2 | 1.532226 | 1.452863 | 0.931864 | 1.014675 | 1.778405 | 1.725301 | 2.554527 | 2.013496 | 4.491897 | 4.183472 | 3.534596 | 3.731377 |
| Nupr1 | 1.541122 | 1.096482 | 3.660591 | 3.482852 | 2.753613 | 2.696631 | 1.379365 | 1.263058 | 1.350157 | 0.84505 | 2.521101 | 2.476506 |
| Dmrt1 | 9.462886 | 9.315258 | 34.49363 | 31.23686 | 24.25058 | 22.37458 | 14.9827 | 14.36056 | 8.523259 | 8.768621 | 9.672988 | 7.58288 |
| Xkrx | 0.603087 | 0.795871 | 1.866125 | 1.850671 | 1.112806 | 1.108205 | 0.910273 | 0.619245 | 0.432877 | 0.506602 | 0.510314 | 0.449292 |
| Cbx4 | 0.081593 | 0.090313 | 0.303314 | 0.326738 | 0.528683 | 0.445318 | 0.776668 | 0.760154 | 1.214826 | 1.051043 | 1.388389 | 1.345129 |
| Ptprv | 1.406186 | 1.377849 | 1.657547 | 1.565313 | 1.189041 | 1.301519 | 1.059743 | 1.204671 | 0.572101 | 0.380348 | 0.457754 | 0.560772 |
| Tdrd12 | 6.322123 | 6.098058 | 7.192056 | 6.448687 | 3.918556 | 3.853522 | 2.612997 | 3.703894 | 1.425441 | 1.605902 | 1.876548 | 1.568078 |
| Grb7 | 19.5599 | 19.6406 | 16.17825 | 16.24531 | 15.08877 | 13.31745 | 10.37787 | 11.53591 | 7.378981 | 9.129178 | 5.154182 | 3.960362 |
| Jakmip1 | 2.253338 | 2.564255 | 0.857359 | 1.004414 | 0.855844 | 0.706641 | 0.706024 | 0.638414 | 0.353865 | 0.245409 | 0.439571 | 0.472475 |
| Gm13051 | 6.90026 | 7.278258 | 9.293606 | 8.829625 | 5.642518 | 6.068107 | 3.930925 | 4.606308 | 2.154362 | 2.205476 | 1.955645 | 2.159478 |
| Osbpl6 | 0.40073 | 0.358275 | 0.30331 | 0.463222 | 0.724354 | 0.688107 | 0.979453 | 1.095476 | 1.705437 | 1.892711 | 1.45471 | 1.400599 |
| Ly6g6e | 3.916907 | 3.665954 | 0.330348 | 0.2277 | 0.440696 | 0.373016 | 0.453699 | 0.615592 | 0.777804 | 0.683919 | 0.858362 | 0.929319 |
| Ccbl2 | 5.148918 | 4.402134 | 8.757296 | 7.787138 | 5.722323 | 5.815068 | 4.577294 | 4.987193 | 3.089656 | 3.119067 | 1.668208 | 1.908195 |
| 4930486L2 | 0.69481 | 0.845137 | 3.231527 | 2.609576 | 1.843661 | 1.973897 | 1.766923 | 1.614922 | 0.621584 | 0.639464 | 0.955941 | 0.781965 |
| Pip5kl1 | 0.562365 | 0.759298 | 1.201167 | 1.121759 | 1.279773 | 0.994036 | 1.026626 | 1.253969 | 0.748039 | 0.695027 | 0.429014 | 0.307723 |
| Rnf128 | 1.691991 | 1.567903 | 1.650114 | 1.575371 | 1.780695 | 1.782583 | 2.19834 | 2.607514 | 2.994521 | 2.550324 | 6.573301 | 6.415639 |
| Col4a1 | 19.8352 | 21.3464 | 25.57948 | 25.91097 | 29.51719 | 26.81901 | 37.26405 | 34.37628 | 81.40479 | 74.48429 | 101.9328 | 105.4348 |
| Meis3 | 1.308431 | 1.366024 | 2.01739 | 2.011814 | 3.776072 | 3.125083 | 5.083694 | 4.57715 | 8.912139 | 7.713957 | 8.420444 | 8.174836 |
| Aurkc | 5.782691 | 5.033845 | 36.44571 | 35.85165 | 27.54344 | 24.51563 | 17.6386 | 17.8965 | 13.03048 | 13.22997 | 10.26382 | 8.824858 |
| Frmd4a | 5.423641 | 5.759053 | 3.984815 | 4.119712 | 4.977729 | 5.069071 | 7.138407 | 6.557649 | 13.26323 | 12.26734 | 15.93938 | 16.71296 |
| Ptprn2 | 0.387151 | 0.365734 | 0.283989 | 0.331426 | 0.453698 | 0.431598 | 0.709924 | 0.500005 | 1.368432 | 1.34405 | 1.302688 | 1.210076 |
| Hbegf | 6.7409 | 7.298668 | 5.122428 | 5.133937 | 4.311289 | 3.986335 | 3.324373 | 3.574776 | 4.29603 | 3.848477 | 14.24083 | 14.48271 |
| Pcgf5 | 5.163165 | 5.850352 | 9.264785 | 8.30152 | 10.70908 | 9.002076 | 10.80163 | 11.25289 | 14.78452 | 14.28072 | 31.82028 | 33.60967 |

| | | | | | | | | | | | | |
|---|---|---|---|---|---|---|---|---|---|---|---|---|
| B4galnt3 | 9.348463 | 10.89684 | 6.784418 | 5.820788 | 3.991778 | 3.836146 | 2.974418 | 3.186503 | 1.658341 | 1.824266 | 1.838861 | 1.437905 |
| Cd80 | 3.78176 | 3.658292 | 4.028547 | 4.017609 | 2.15559 | 2.531454 | 2.14581 | 2.314424 | 1.041447 | 0.99288 | 1.844911 | 2.090232 |
| Nxf7 | 1.713723 | 1.771142 | 1.352546 | 1.521368 | 1.722266 | 1.586052 | 2.627367 | 2.728492 | 4.485727 | 4.640986 | 5.464762 | 6.141014 |
| Gdf3 | 43.98531 | 41.14917 | 37.19823 | 33.70179 | 22.4619 | 23.11182 | 18.75412 | 17.02654 | 9.844209 | 8.552289 | 8.594981 | 8.371174 |
| Bmp6 | 0.853974 | 0.949378 | 1.229332 | 1.024442 | 1.3452 | 1.115226 | 1.245139 | 1.419626 | 2.806102 | 2.10727 | 3.73636 | 4.124122 |
| Tspan2 | 2.768647 | 2.629103 | 3.350218 | 2.833691 | 3.660645 | 3.740502 | 4.722777 | 4.149472 | 6.631489 | 6.349008 | 10.26296 | 11.40509 |
| Mycn | 74.08218 | 71.65317 | 14.90884 | 14.30345 | 13.08441 | 12.9382 | 20.41434 | 20.37166 | 55.60503 | 52.06517 | 47.94815 | 45.52383 |
| Wdr64 | 0.743554 | 1.035 | 2.208934 | 1.975008 | 1.459363 | 1.551666 | 1.141343 | 1.716294 | 0.467094 | 0.491028 | 0.499554 | 0.717415 |
| Fabp5 | 7.864429 | 5.973767 | 4.60318 | 4.859142 | 6.307638 | 6.883829 | 8.597741 | 7.512051 | 18.5435 | 16.66051 | 19.88783 | 19.54413 |
| Tead4 | 5.120149 | 5.585698 | 2.653895 | 2.600559 | 1.98317 | 1.865541 | 1.996682 | 1.889311 | 2.962081 | 2.981798 | 7.773989 | 7.497879 |
| Celf4 | 1.751192 | 2.210314 | 1.062784 | 1.367787 | 1.053822 | 0.966512 | 1.150691 | 0.891589 | 0.479883 | 0.528941 | 0.383849 | 0.340439 |
| Serpinb1a | 0.272639 | 0.266085 | 0.204582 | 0.193529 | 0.252078 | 0.311175 | 0.336498 | 0.464515 | 0.65527 | 0.690789 | 1.35389 | 0.777524 |
| Dapp1 | 3.418665 | 3.837142 | 2.985942 | 2.844152 | 1.965337 | 2.160616 | 1.902086 | 1.656994 | 1.248876 | 1.148344 | 0.830674 | 0.708127 |
| Cpq | 0.325569 | 0.176197 | 0.11638 | 0.262575 | 0.345301 | 0.356583 | 0.345245 | 0.511809 | 1.009019 | 1.05459 | 0.796031 | 0.725467 |
| Nostrin | 1.548866 | 1.320224 | 1.236661 | 1.293633 | 1.360001 | 1.203351 | 1.760351 | 2.047843 | 4.526831 | 4.343755 | 3.744039 | 4.831181 |
| Chst7 | 0.310141 | 0.404437 | 0.360388 | 0.453337 | 0.793531 | 0.534617 | 1.058068 | 0.637358 | 1.091153 | 1.244551 | 1.836046 | 1.818875 |
| Vgll4 | 2.182179 | 1.91412 | 4.108682 | 3.887461 | 4.84741 | 4.589176 | 5.767766 | 5.415942 | 9.15928 | 9.085164 | 15.22848 | 15.59659 |
| Grhl2 | 5.23046 | 5.573873 | 3.530413 | 3.161456 | 1.737651 | 1.648008 | 1.669607 | 1.411602 | 0.742924 | 0.78818 | 1.137015 | 1.118889 |
| Adamts8 | 0.627599 | 0.672861 | 0.433212 | 0.692997 | 0.460307 | 0.639917 | 0.644357 | 0.529207 | 1.058135 | 0.923697 | 2.239386 | 2.121009 |
| Axl | 0.916603 | 0.884025 | 0.585707 | 0.763291 | 0.92602 | 0.853317 | 1.400435 | 1.527439 | 2.043581 | 1.959051 | 3.278155 | 3.058847 |
| 2510009E0 | 0.058788 | 0.052864 | 0.395824 | 0.292199 | 0.869402 | 0.906025 | 1.246858 | 1.17077 | 1.118014 | 1.042929 | 0.86943 | 0.862499 |
| Aifm2 | 2.842126 | 2.819904 | 4.962535 | 4.588178 | 3.724738 | 3.412629 | 2.53528 | 2.688099 | 1.299394 | 1.145266 | 1.197566 | 1.324626 |
| Chd5 | 0.939962 | 1.133871 | 1.516673 | 1.487995 | 1.201067 | 1.209086 | 0.795804 | 0.785565 | 0.311705 | 0.371538 | 0.471666 | 0.413992 |
| Ydjc | 8.469516 | 8.026979 | 6.286236 | 6.567344 | 5.145088 | 5.183195 | 3.491819 | 3.586535 | 1.815559 | 1.8703 | 1.641713 | 1.639932 |
| Cbx8 | 0.084644 | 0.141709 | 0.611446 | 0.480889 | 0.703294 | 0.792818 | 1.196813 | 1.296045 | 1.32557 | 1.312809 | 2.047649 | 1.924927 |
| Nfatc4 | 6.778385 | 7.106343 | 3.669415 | 3.973728 | 3.837008 | 3.471351 | 3.527216 | 3.976531 | 2.371858 | 2.267353 | 1.27077 | 0.99399 |
| Vstm2l | 0.591093 | 0.511699 | 0.484772 | 0.473038 | 0.69526 | 0.87071 | 1.380381 | 1.891189 | 1.912797 | 1.179438 | 2.009564 | 1.851977 |
| Tgfbi | 6.711586 | 7.123233 | 3.597492 | 3.538376 | 5.270034 | 5.137162 | 10.87876 | 9.677932 | 13.83406 | 14.13249 | 10.98185 | 9.263676 |
| Rhbdl3 | 0.26476 | 0.317917 | 1.005245 | 0.946975 | 1.536805 | 1.509587 | 3.000944 | 2.960555 | 3.517749 | 3.100168 | 4.027199 | 3.781365 |
| Tmprss2 | 0.12127 | 0.146895 | 0.326563 | 0.294929 | 0.370089 | 0.495555 | 0.619003 | 0.385737 | 0.88325 | 1.127636 | 1.434729 | 1.176425 |
| Olig3 | 1.987052 | 2.165284 | 1.325509 | 1.121103 | 2.521201 | 2.589864 | 4.898624 | 4.166981 | 2.190365 | 1.594853 | 1.389526 | 1.045676 |
| Zfhx2 | 2.23255 | 3.293727 | 9.267651 | 8.620293 | 6.298786 | 6.276812 | 5.600538 | 4.48972 | 3.987085 | 3.959457 | 2.306711 | 2.163492 |
| Cacna1g | 0.202228 | 0.201651 | 0.550466 | 0.634919 | 1.238931 | 1.206447 | 1.759593 | 1.800215 | 2.54816 | 2.527523 | 1.971979 | 2.078581 |
| Wfdc15a | 0.70299 | 0.559405 | 2.336696 | 2.841681 | 1.278547 | 2.539354 | 2.09782 | 2.593056 | 1.176156 | 0.714967 | 1.629515 | 1.810155 |
| Stra8 | 16.24027 | 14.41273 | 400.4022 | 385.2561 | 525.8809 | 493.3814 | 443.7656 | 572.0488 | 193.8062 | 193.163 | 151.6613 | 143.9423 |
| Ankrd35 | 2.139002 | 1.618138 | 1.734233 | 1.512937 | 1.207465 | 1.073915 | 0.860716 | 0.980877 | 0.485286 | 0.380771 | 0.645649 | 0.657599 |
| Ltbp1 | 1.984795 | 2.169931 | 1.742278 | 1.950441 | 2.423921 | 2.372676 | 3.607659 | 3.077168 | 7.325406 | 7.217815 | 7.591603 | 7.739522 |
| Lgals4 | 2.185076 | 1.619245 | 3.585246 | 3.466134 | 6.054147 | 7.131728 | 11.8255 | 13.74858 | 9.90771 | 9.658245 | 10.61488 | 8.674471 |
| Maob | 0.466993 | 0.350569 | 0.4712 | 0.476036 | 0.660398 | 0.586662 | 1.002244 | 1.088576 | 1.503869 | 1.8877 | 1.23819 | 1.320691 |
| BC048679 | 0.03171 | 0.1 | 1.092346 | 2.04989 | 2.046895 | 1.679584 | 2.083396 | 1.903976 | 0.768967 | 0.517282 | 0.675691 | 1.113636 |
| Lamc2 | 1.223301 | 1.462833 | 1.845807 | 1.676355 | 1.299066 | 1.147069 | 0.995801 | 1.007406 | 0.66417 | 0.730872 | 3.127745 | 2.893301 |
| Plekhg2 | 3.253641 | 3.735601 | 3.993303 | 3.742497 | 5.345267 | 5.136981 | 6.340719 | 5.791415 | 7.547549 | 7.790472 | 15.87484 | 14.806 |
| Phc1 | 125.737 | 144.1355 | 93.20128 | 88.34636 | 48.5425 | 48.1555 | 39.43422 | 37.70059 | 29.30839 | 29.35323 | 23.20996 | 22.33824 |
| Fam124a | 0.402084 | 0.362155 | 1.416159 | 1.323934 | 2.870113 | 2.705727 | 4.458551 | 4.913273 | 5.182243 | 5.235432 | 4.861718 | 4.040328 |
| Mt2 | 129.4716 | 121.018 | 260.7088 | 251.5099 | 201.7624 | 221.9994 | 183.1465 | 185.361 | 112.955 | 94.72172 | 442.1656 | 374.5539 |
| Acap1 | 3.022713 | 3.542005 | 3.602565 | 3.105455 | 3.352338 | 2.90211 | 1.922024 | 2.222758 | 0.988777 | 0.976137 | 0.784189 | 0.78586 |
| Tdh | 69.58355 | 74.47787 | 10.37046 | 9.446863 | 3.426905 | 3.216893 | 2.483981 | 2.940094 | 2.24455 | 2.391162 | 2.678308 | 2.664179 |
| Gna15 | 3.081866 | 2.752765 | 1.111343 | 1.141255 | 1.026147 | 0.984292 | 0.846628 | 0.74051 | 0.492392 | 0.311196 | 0.366231 | 0.289073 |
| Gng12 | 2.484182 | 2.280782 | 1.661296 | 2.165996 | 2.806856 | 2.752708 | 3.329545 | 3.078135 | 7.890833 | 8.537507 | 6.872911 | 7.688322 |
| Lbx1 | 0.170452 | 0.153083 | 0.377547 | 0.577337 | 0.381008 | 0.484707 | 0.590128 | 0.358484 | 1.057306 | 0.833093 | 1.47175 | 1.411244 |
| Podxl | 6.907349 | 7.240568 | 3.048762 | 3.195988 | 4.892707 | 4.143941 | 6.01611 | 5.82898 | 12.65239 | 12.57982 | 12.65133 | 12.44716 |
| Kifc2 | 4.136649 | 4.132425 | 4.324566 | 4.278697 | 4.472374 | 4.970171 | 3.494128 | 3.608602 | 1.958049 | 1.926808 | 1.574447 | 1.263055 |
| Kctd11 | 0.346049 | 0.370661 | 0.932249 | 0.890139 | 1.119746 | 1.223492 | 1.588263 | 1.191789 | 2.269486 | 2.061014 | 3.773738 | 3.500808 |
| Ptprm | 0.163189 | 0.169709 | 0.324174 | 0.338623 | 0.683488 | 0.583131 | 0.881701 | 0.892065 | 1.227749 | 1.331238 | 1.136406 | 1.089413 |
| Meis2 | 1.82534 | 1.970226 | 8.420301 | 8.92237 | 15.10227 | 15.3635 | 27.16825 | 25.36858 | 38.21443 | 35.05441 | 25.91379 | 26.81164 |
| 2810459M | 0.744214 | 0.679831 | 3.710652 | 3.574378 | 6.162953 | 6.105761 | 8.954191 | 8.287216 | 2.717913 | 2.486266 | 1.88633 | 2.109378 |
| Hoxb13 | 0.456579 | 0.307605 | 1.845429 | 1.393284 | 1.133079 | 1.301864 | 1.458921 | 1.210981 | 0.876768 | 0.621813 | 0.328704 | 0.354912 |
| Capn2 | 6.558065 | 5.694948 | 7.049061 | 6.498164 | 8.22636 | 8.041401 | 11.43726 | 10.72085 | 15.94463 | 15.59589 | 25.42478 | 25.49455 |
| Adamts15 | 0.45265 | 0.48964 | 0.378613 | 0.396203 | 0.635163 | 0.497996 | 0.427973 | 0.60086 | 1.055561 | 1.201426 | 1.514816 | 1.554341 |
| Gsc | 0.158735 | 0.214564 | 0.389976 | 0.422083 | 0.650278 | 0.664485 | 0.745246 | 0.739229 | 1.151589 | 1.052145 | 2.235545 | 1.650389 |
| Gm44 | 0.588487 | 0.51461 | 1.76654 | 2.100855 | 2.151652 | 2.159864 | 2.547136 | 3.256626 | 1.422566 | 1.491258 | 0.808595 | 0.83392 |

| | | | | | | | | | | | | |
|---|---|---|---|---|---|---|---|---|---|---|---|---|
| Hoxa9 | 0.062564 | 0.098609 | 1.183982 | 1.005478 | 2.210062 | 2.671952 | 3.191298 | 3.021809 | 5.012581 | 3.925128 | 2.619294 | 2.994104 |
| Mgp | 3.101296 | 0.632796 | 0.488307 | 0.569774 | 0.740375 | 0.596563 | 0.782495 | 1.097223 | 0.196041 | 0.384493 | 0.325643 | 0.28117 |
| Ccdc88b | 0.374038 | 0.316378 | 5.663619 | 5.282061 | 6.029749 | 6.649975 | 6.804362 | 6.228153 | 2.255245 | 2.343217 | 1.935647 | 1.704904 |
| C330013F1 | 1.396046 | 1.547866 | 1.934132 | 1.755299 | 1.244875 | 1.317776 | 0.936673 | 0.961102 | 0.703704 | 0.712538 | 0.344252 | 0.450244 |
| Trh | 189.5555 | 156.1862 | 42.26676 | 41.88604 | 41.00171 | 39.28379 | 50.48116 | 52.11593 | 15.8578 | 16.358 | 14.39597 | 13.37171 |
| Ccl3 | 0.052988 | 0.162261 | 0.634424 | 0.694294 | 1.499828 | 0.658228 | 2.069323 | 1.713157 | 2.419485 | 2.56471 | 2.195005 | 1.824715 |
| Tax1bp3 | 2.135302 | 2.109926 | 3.160865 | 2.490595 | 3.351115 | 3.175936 | 4.5018 | 4.183295 | 5.521306 | 4.84924 | 9.813344 | 9.704209 |
| Ccser1 | 0.056908 | 0.067382 | 0.273164 | 0.273125 | 0.434304 | 0.409704 | 0.719067 | 0.654499 | 0.94462 | 0.910042 | 1.111886 | 1.063696 |
| Cpm | 0.443044 | 0.480122 | 1.451593 | 1.267319 | 2.047818 | 1.847254 | 2.318842 | 2.435826 | 3.753938 | 3.73443 | 5.369129 | 4.929317 |
| Psd3 | 0.436719 | 0.55354 | 0.68261 | 0.756376 | 0.96112 | 0.98761 | 1.278266 | 1.24217 | 2.615633 | 2.507592 | 2.444019 | 2.941503 |
| Ppm1j | 16.16729 | 14.8169 | 6.357145 | 5.786969 | 4.729018 | 4.819225 | 3.716173 | 4.193801 | 1.896642 | 1.488229 | 1.853166 | 1.558842 |
| 1190002F1 | 3.369794 | 2.456684 | 7.099501 | 5.945625 | 5.406436 | 5.404427 | 4.718725 | 4.135331 | 2.320144 | 1.530497 | 2.962671 | 2.648086 |
| Nphp4 | 1.631589 | 1.888605 | 6.710607 | 6.396097 | 4.571069 | 4.313945 | 3.417893 | 3.561414 | 2.207429 | 2.269332 | 2.048466 | 1.64674 |
| Lmo2 | 0.648733 | 0.322739 | 0.648424 | 0.818285 | 1.204338 | 1.43305 | 1.329336 | 1.208764 | 0.910456 | 0.729684 | 0.60228 | 0.369127 |
| Actg1 | 217.6389 | 246.5357 | 317.4809 | 299.0918 | 332.1867 | 318.9068 | 412.3405 | 402.716 | 583.0506 | 546.8582 | 1182.156 | 1161.055 |
| Lrp3 | 1.97191 | 2.333654 | 2.106512 | 2.102003 | 3.302415 | 3.095661 | 3.980328 | 4.046499 | 8.199378 | 8.148458 | 8.091719 | 7.764133 |
| Adcy2 | 0.25853 | 0.361204 | 0.567405 | 0.640396 | 1.033227 | 0.987614 | 1.743983 | 1.825268 | 2.448043 | 2.481685 | 2.291791 | 2.214402 |
| Dnaaf3 | 2.022292 | 2.107032 | 2.005877 | 1.652768 | 1.47691 | 1.411118 | 1.01659 | 1.095813 | 0.859236 | 0.924297 | 0.558433 | 0.426523 |
| Exoc3l | 3.276724 | 3.531892 | 2.650424 | 2.386639 | 1.500126 | 1.725556 | 1.067673 | 1.238778 | 0.704967 | 0.735117 | 0.862666 | 0.615999 |
| Ppp4r4 | 0.762271 | 0.723985 | 0.577489 | 0.570309 | 0.771829 | 0.887921 | 1.12151 | 1.217788 | 1.957665 | 2.207289 | 1.092089 | 1.209645 |
| Ybx2 | 5.996642 | 5.776661 | 3.704853 | 3.522098 | 3.730746 | 3.018951 | 2.163728 | 2.661311 | 1.533012 | 1.605512 | 1.164674 | 0.910367 |
| Mmp9 | 0.190182 | 0.31986 | 1.711188 | 1.516119 | 1.077127 | 1.261227 | 0.860349 | 0.9152 | 0.595507 | 0.392095 | 1.046326 | 0.837749 |
| Vim | 91.76086 | 99.21307 | 89.81761 | 85.17699 | 86.28521 | 83.34449 | 101.2721 | 99.99595 | 137.0754 | 115.7073 | 334.8919 | 322.174 |
| Rab3a | 2.628903 | 3.021349 | 5.823094 | 5.054168 | 5.606102 | 6.05103 | 3.864692 | 4.698757 | 2.743962 | 2.697044 | 1.716524 | 1.565945 |
| Rnf180 | 0.267548 | 0.477145 | 0.437882 | 0.403116 | 0.524033 | 0.571363 | 0.586808 | 0.515321 | 1.059675 | 0.741082 | 1.067786 | 1.556652 |
| Pipox | 42.62528 | 39.31053 | 23.84577 | 25.97211 | 22.38715 | 23.47351 | 21.3079 | 21.79155 | 13.80631 | 14.24199 | 5.622326 | 6.735461 |
| Gm19705 | 16.63604 | 16.96891 | 12.08439 | 10.5214 | 5.820701 | 6.211314 | 3.848762 | 4.009279 | 2.71405 | 2.73076 | 2.473609 | 2.731852 |
| Themis2 | 0.491634 | 0.578033 | 3.00089 | 3.050602 | 1.719949 | 1.773909 | 1.775905 | 1.461267 | 1.102749 | 1.200027 | 0.970441 | 0.792173 |
| Acbd7 | 1.970119 | 1.747801 | 1.677839 | 1.274917 | 1.93675 | 1.583806 | 1.209578 | 1.853501 | 0.810772 | 1.045311 | 0.738569 | 0.482003 |
| Ntn4 | 0.265653 | 0.214523 | 0.56998 | 0.318403 | 0.57979 | 0.622905 | 0.771191 | 0.874098 | 1.012191 | 1.224322 | 1.139134 | 1.1163 |
| Ccdc155 | 1.666928 | 1.371919 | 1.599996 | 1.563433 | 1.379964 | 1.183346 | 0.765476 | 0.870211 | 0.489618 | 0.602926 | 0.528855 | 0.407177 |
| Pcolce | 32.96749 | 34.27611 | 29.67023 | 28.00344 | 19.20649 | 18.7909 | 14.12837 | 14.82698 | 8.054515 | 7.311967 | 9.880382 | 10.19739 |
| Itga8 | 0.347579 | 0.20443 | 0.631359 | 0.586068 | 0.765906 | 0.894495 | 0.991907 | 0.782463 | 2.211366 | 2.169489 | 2.116436 | 2.239409 |
| Fam78a | 1.973522 | 2.084569 | 2.417913 | 2.511355 | 2.044301 | 1.956975 | 1.550271 | 1.814431 | 1.128196 | 1.168496 | 0.653913 | 0.657461 |
| Gm6484 | 3.152663 | 4.891325 | 6.219155 | 6.041619 | 4.965492 | 5.415668 | 3.690855 | 4.339545 | 1.444395 | 1.794161 | 1.279286 | 1.582778 |
| Srgap3 | 2.114964 | 2.711142 | 1.579808 | 1.547461 | 1.792379 | 1.940825 | 2.515832 | 2.352056 | 5.666327 | 5.898475 | 4.442038 | 4.658576 |
| Mvb12b | 2.469623 | 2.535671 | 1.858935 | 1.812254 | 2.864265 | 2.821519 | 3.309494 | 3.160106 | 6.767214 | 6.787434 | 6.821289 | 6.898861 |
| Klhdc7a | 0.872938 | 0.900439 | 1.498858 | 1.506732 | 3.256341 | 3.613793 | 5.937944 | 5.661031 | 4.96992 | 4.771376 | 5.73215 | 5.735178 |
| Wdr86 | 2.522317 | 2.446187 | 2.194471 | 2.126089 | 2.658452 | 2.507071 | 3.023373 | 2.643123 | 2.490206 | 2.643042 | 0.71243 | 0.695509 |
| Gm15706 | 2.394431 | 1.856289 | 1.081132 | 1.491421 | 1.768848 | 1.426286 | 0.943261 | 1.303312 | 0.747488 | 0.392497 | 0.525811 | 0.591736 |
| Fhl1 | 8.350301 | 7.687059 | 6.29966 | 6.292618 | 8.991276 | 8.944434 | 11.00952 | 11.48836 | 22.72478 | 21.06695 | 23.04592 | 23.90163 |
| 4930500J0: | 44.45271 | 43.14429 | 47.53925 | 48.43938 | 33.31985 | 32.4119 | 25.62368 | 27.99852 | 13.29881 | 15.33871 | 14.93322 | 12.76139 |
| Tmod4 | 0.288155 | 0.306185 | 0.568379 | 0.344478 | 0.606688 | 0.687022 | 0.782584 | 0.85551 | 1.62967 | 1.307361 | 1.099987 | 1.047528 |
| Eno3 | 14.96501 | 13.7352 | 22.4727 | 21.72749 | 18.70831 | 17.99647 | 16.34362 | 16.84099 | 7.555327 | 8.062098 | 7.482736 | 5.726259 |
| Adad2 | 2.109986 | 1.882288 | 5.401543 | 5.513283 | 4.187999 | 4.278679 | 3.159867 | 3.913694 | 1.542553 | 1.535095 | 1.582517 | 1.453802 |
| Il33 | 0.578304 | 0.840559 | 5.80781 | 5.50324 | 6.564139 | 6.554508 | 7.480227 | 7.700646 | 2.561879 | 2.035153 | 3.170359 | 3.266848 |
| Glis1 | 0.15297 | 0.204042 | 0.379931 | 0.263237 | 0.499212 | 0.575174 | 0.715356 | 0.62898 | 0.815442 | 0.875839 | 1.218053 | 0.994747 |
| Nrp1 | 0.28676 | 0.186264 | 0.601145 | 0.605478 | 1.039635 | 1.0004 | 1.78645 | 1.785186 | 2.35868 | 2.28791 | 1.755031 | 1.808797 |
| Pde2a | 1.093906 | 1.112622 | 2.554684 | 2.32505 | 2.147053 | 2.044443 | 1.592379 | 1.64368 | 0.753861 | 0.893414 | 0.526957 | 0.616019 |
| Acp6 | 90.88592 | 81.71971 | 85.61337 | 79.59451 | 59.01865 | 58.73454 | 42.28092 | 41.3093 | 26.98335 | 25.80158 | 21.80349 | 21.10758 |
| Rhob | 9.094021 | 8.087692 | 17.38226 | 16.14814 | 26.09282 | 24.7203 | 27.99716 | 27.57632 | 31.97801 | 28.94188 | 64.39335 | 60.88096 |
| Nxph4 | 0.601155 | 0.630487 | 2.929275 | 2.971543 | 3.643502 | 3.408391 | 3.218749 | 2.462271 | 1.015932 | 0.90416 | 1.343562 | 0.994801 |
| Sp110 | 0.327622 | 0.392455 | 1.472114 | 1.310667 | 2.371483 | 2.431035 | 3.784541 | 4.935288 | 4.210333 | 4.265622 | 5.328747 | 3.985736 |
| Pycard | 16.02103 | 13.13814 | 15.36599 | 16.12515 | 12.79818 | 13.38517 | 9.592872 | 9.440133 | 4.052116 | 4.689884 | 3.9191 | 4.28481 |
| Lpar5 | 2.613024 | 2.56102 | 3.537762 | 3.191249 | 1.987223 | 1.808819 | 1.95583 | 1.285791 | 1.073688 | 0.84838 | 1.002909 | 1.104834 |
| Usp17lc | 0.444485 | 0.404995 | 2.289312 | 1.898246 | 3.811887 | 3.900582 | 7.59119 | 7.138308 | 6.482566 | 6.477458 | 7.229439 | 5.321282 |
| Zfp534 | 19.09942 | 19.36422 | 21.13481 | 20.40488 | 14.17056 | 14.22328 | 9.204327 | 9.747055 | 5.853573 | 5.430166 | 5.080799 | 5.685479 |
| Nnat | 11.13135 | 11.01369 | 11.13113 | 11.22837 | 15.09589 | 16.15479 | 25.61754 | 25.18135 | 40.89915 | 34.59544 | 44.15271 | 42.1794 |
| Nxpe4 | 0.419506 | 0.533921 | 0.505147 | 0.489413 | 0.831595 | 0.874666 | 0.950107 | 0.98963 | 1.811602 | 1.837897 | 0.852014 | 1.016238 |
| Enox1 | 24.66597 | 22.06752 | 3.663377 | 3.779939 | 1.516445 | 1.394777 | 1.019611 | 1.180058 | 1.303485 | 1.243759 | 1.213386 | 1.006574 |
| Tex40 | 3.716589 | 2.914222 | 14.29775 | 12.03628 | 13.86751 | 17.30853 | 12.57038 | 13.02197 | 6.997378 | 5.886199 | 4.895074 | 4.617312 |

| | | | | | | | | | | | | |
|---|---|---|---|---|---|---|---|---|---|---|---|---|
| Myrf | 17.68212 | 21.84158 | 24.93298 | 23.27147 | 16.07004 | 15.00478 | 12.95397 | 11.09314 | 6.834678 | 7.176774 | 7.018415 | 6.212694 |
| Gas7 | 0.554045 | 0.735266 | 1.08487 | 1.370741 | 2.313821 | 1.993913 | 3.713996 | 3.296016 | 5.263347 | 5.001609 | 4.840543 | 5.132932 |
| Necab3 | 3.622125 | 3.924782 | 2.937871 | 3.196565 | 3.525331 | 3.548737 | 2.330766 | 2.511195 | 1.682927 | 1.80996 | 1.084385 | 0.948158 |
| Rassf8 | 1.792642 | 1.806195 | 2.787773 | 2.631762 | 3.824013 | 3.697967 | 4.331696 | 3.881208 | 6.551504 | 6.386048 | 9.686731 | 9.849201 |
| Arg2 | 0.338746 | 0.487761 | 1.637388 | 1.790924 | 3.385969 | 3.642956 | 7.221256 | 6.695299 | 5.962754 | 5.681859 | 5.879289 | 5.250501 |
| Sgce | 5.861865 | 4.567807 | 5.455775 | 5.914481 | 8.880522 | 8.749788 | 12.40721 | 11.31421 | 20.72839 | 20.01284 | 20.43624 | 22.10263 |
| Gm13154 | 30.07704 | 28.51098 | 23.78532 | 24.73326 | 16.46999 | 17.10556 | 11.00664 | 11.57254 | 8.303216 | 9.11183 | 6.342334 | 6.621294 |
| Gm1673 | 2.316993 | 1.9013 | 2.489354 | 2.554925 | 4.689891 | 6.399505 | 8.58049 | 6.026782 | 12.35542 | 9.531055 | 12.86537 | 8.317919 |
| Anxa2 | 11.56539 | 8.608711 | 3.580451 | 3.63236 | 3.4078 | 3.655668 | 3.82967 | 3.905591 | 6.014949 | 6.07898 | 15.51041 | 13.54701 |
| Adamts7 | 0.756336 | 1.002721 | 1.395206 | 1.398793 | 2.260978 | 2.085851 | 3.055874 | 3.39806 | 4.763422 | 5.214443 | 4.623218 | 4.219923 |
| Pla2g1b | 71.95804 | 77.82948 | 63.02077 | 63.68144 | 56.63842 | 52.90007 | 44.26098 | 39.04188 | 31.02027 | 26.93745 | 15.59145 | 17.08298 |
| Ooep | 18.41923 | 17.05042 | 17.30661 | 17.463 | 11.42244 | 10.7179 | 7.56498 | 7.803875 | 5.56875 | 5.4704 | 4.81829 | 4.709979 |
| Tle6 | 3.609656 | 3.347061 | 18.68142 | 16.90544 | 15.15343 | 14.5221 | 10.58137 | 11.02481 | 5.958817 | 5.907524 | 5.084555 | 4.561652 |
| Fgf10 | 0.490125 | 0.580654 | 0.448149 | 0.49176 | 0.508668 | 0.418267 | 1.093157 | 0.865799 | 1.760974 | 1.550001 | 1.049452 | 1.249179 |
| Ptpn9 | 3.574229 | 3.264437 | 4.814391 | 4.810159 | 6.532427 | 6.622424 | 8.463186 | 9.086713 | 12.12307 | 11.50769 | 17.03944 | 17.82057 |
| Gm13242 | 23.3823 | 23.59083 | 17.24624 | 16.91338 | 10.55956 | 10.26604 | 7.138008 | 8.376644 | 5.300593 | 5.290233 | 3.970603 | 4.567966 |
| Mfsd7c | 0.539581 | 0.549622 | 1.07968 | 0.92097 | 1.29558 | 1.365597 | 1.767056 | 1.719142 | 3.147502 | 3.084454 | 2.838949 | 3.407041 |
| Arhgap29 | 7.250291 | 7.216507 | 9.081094 | 9.810971 | 11.46478 | 10.91887 | 14.27609 | 15.2394 | 19.42236 | 21.28926 | 32.70111 | 36.26706 |
| Heg1 | 1.067227 | 1.41403 | 2.836867 | 3.098747 | 4.792761 | 4.621185 | 6.616755 | 5.461414 | 7.730374 | 7.887326 | 11.49723 | 11.42309 |
| Arc | 6.905729 | 7.479265 | 4.283798 | 4.022127 | 2.477896 | 2.186947 | 1.64265 | 1.779569 | 1.137214 | 1.091367 | 2.229982 | 2.207038 |
| Celsr3 | 1.982101 | 2.668274 | 2.360994 | 2.406022 | 1.599745 | 1.626287 | 1.155203 | 1.242642 | 0.635168 | 0.653193 | 0.912462 | 0.847339 |
| Ifitm7 | 0.94153 | 0.964919 | 2.161832 | 1.770229 | 1.646645 | 1.368117 | 0.982341 | 1.131186 | 0.431279 | 0.480591 | 0.318976 | 0.741956 |
| Cx3cl1 | 0.395387 | 0.422126 | 0.854892 | 1.007423 | 1.582069 | 1.338553 | 1.835985 | 1.443806 | 1.91912 | 1.717045 | 4.111505 | 3.710547 |
| Etv1 | 7.804426 | 8.247371 | 7.298452 | 7.201109 | 7.928766 | 8.109503 | 8.13081 | 7.105691 | 4.083951 | 3.856726 | 2.239182 | 2.202231 |
| Tle2 | 0.192602 | 0.1836 | 1.054052 | 1.078298 | 1.151917 | 1.159596 | 1.032359 | 0.907863 | 0.629395 | 0.4766 | 0.401263 | 0.315093 |
| Fyn | 5.615293 | 5.790444 | 7.547985 | 7.495658 | 9.494788 | 9.349776 | 11.50673 | 11.34178 | 19.25019 | 17.80008 | 27.88021 | 27.55797 |
| 1700025G0 | 1.126302 | 1.37757 | 1.327379 | 1.29325 | 1.991335 | 1.871649 | 2.354219 | 2.441658 | 4.288209 | 4.187812 | 4.159571 | 4.750589 |
| Klk1 | 4.987593 | 4.501681 | 0.632606 | 0.451977 | 0.252592 | 0.261681 | 0.154433 | 0.321007 | 0.133827 | 0.153861 | 0.219205 | 0.123159 |
| Sh3bgr | 0.441736 | 0.546443 | 0.535683 | 0.318598 | 0.64028 | 0.536554 | 0.493579 | 0.922441 | 1.09009 | 0.802832 | 1.348126 | 1.16828 |
| Fam102b | 1.071311 | 1.090916 | 1.671279 | 1.841443 | 2.950202 | 2.603837 | 3.63982 | 3.230229 | 5.042531 | 5.459969 | 6.672172 | 6.743705 |
| 5430402O1 | 0.171239 | 0.090113 | 0.671178 | 0.748806 | 1.327513 | 1.779812 | 2.433309 | 2.742223 | 2.285541 | 2.347246 | 2.889994 | 2.551874 |
| Gpr20 | 0.367217 | 0.355148 | 2.066821 | 1.732214 | 2.273336 | 1.958563 | 1.508892 | 1.792491 | 1.003924 | 1.185307 | 0.412061 | 0.534873 |
| Gpr56 | 0.475965 | 0.632883 | 0.41328 | 0.414907 | 0.446667 | 0.362953 | 0.408551 | 0.585026 | 0.569316 | 0.675787 | 1.555239 | 1.327345 |
| Lphn2 | 10.56831 | 11.95423 | 7.720639 | 7.701464 | 9.755694 | 9.719378 | 13.79611 | 10.57065 | 24.39079 | 24.38662 | 29.06262 | 28.15164 |
| Pdgfa | 15.46615 | 16.73439 | 5.318231 | 4.720942 | 4.860711 | 4.581377 | 6.529889 | 7.121068 | 7.453957 | 7.816498 | 18.40741 | 16.71361 |
| 1700019A0 | 2.662494 | 3.38487 | 4.603436 | 4.232833 | 2.912846 | 3.769957 | 2.565592 | 1.742991 | 1.574275 | 1.160425 | 1.270753 | 1.479199 |
| Gm4285 | 0.490978 | 0.562436 | 0.947568 | 0.992828 | 1.78743 | 1.812303 | 2.986617 | 2.909708 | 3.608563 | 2.918504 | 3.852551 | 3.616256 |
| Tcf15 | 92.83428 | 91.68093 | 94.73777 | 87.28189 | 98.18411 | 99.146 | 74.42105 | 74.46128 | 33.75061 | 31.76288 | 31.97105 | 27.22894 |
| Tcf12 | 6.378152 | 6.58195 | 7.912278 | 8.156459 | 11.24152 | 10.52459 | 14.15655 | 13.296 | 26.98688 | 26.9559 | 28.65131 | 29.69628 |
| Chsy3 | 0.123874 | 0.071621 | 0.305428 | 0.306875 | 0.569964 | 0.364608 | 0.733281 | 0.559655 | 0.913667 | 0.836581 | 1.127624 | 1.115606 |
| Lef1 | 15.09048 | 15.41041 | 2.158845 | 2.360448 | 2.627503 | 2.658995 | 2.484826 | 2.294641 | 0.814838 | 0.73197 | 0.965006 | 0.816741 |
| Rsph1 | 2.73611 | 3.306371 | 2.252273 | 1.879602 | 1.818167 | 1.607682 | 1.40728 | 0.885804 | 0.744025 | 0.517699 | 0.81087 | 0.668348 |
| Tnfaip2 | 0.160699 | 0.232842 | 0.461471 | 0.355572 | 0.621386 | 0.672492 | 0.520203 | 0.723072 | 0.724196 | 0.634597 | 1.652208 | 1.290552 |
| Rab33a | 1.913868 | 1.313496 | 0.970006 | 0.946363 | 1.269731 | 1.144436 | 1.325483 | 1.335591 | 1.193777 | 0.814855 | 0.381459 | 0.36837 |
| Cebpd | 6.472581 | 5.79897 | 5.870459 | 5.708616 | 5.75077 | 6.749934 | 3.671839 | 4.766145 | 2.757562 | 1.862984 | 4.963003 | 4.790955 |
| Flnb | 19.39378 | 25.82258 | 25.76789 | 25.36018 | 22.76235 | 21.50003 | 26.82676 | 20.36916 | 30.2648 | 31.73275 | 75.75156 | 73.77317 |
| Amotl2 | 10.14862 | 11.86757 | 23.08566 | 21.33154 | 21.44425 | 21.28655 | 27.13788 | 24.94214 | 40.10368 | 39.18225 | 77.75782 | 77.08719 |
| Myof | 5.313106 | 6.277246 | 3.750593 | 3.818296 | 3.594416 | 3.520275 | 4.616492 | 4.525508 | 7.195178 | 7.173942 | 11.95462 | 12.74667 |
| C530008M | 1.564842 | 1.819223 | 2.748293 | 3.157695 | 4.882515 | 5.085214 | 6.979677 | 6.609201 | 11.9076 | 11.43075 | 9.91822 | 9.788807 |
| Patl2 | 3.180503 | 3.18867 | 1.622654 | 1.502567 | 0.950753 | 0.971007 | 0.658973 | 0.847228 | 0.438827 | 0.513925 | 0.35915 | 0.4155 |
| Grb10 | 21.19627 | 22.34584 | 36.70401 | 38.95321 | 57.42194 | 57.83102 | 83.34095 | 75.79671 | 135.0874 | 140.7943 | 115.0025 | 118.8461 |
| Lonrf3 | 0.539776 | 0.437433 | 1.298188 | 1.306218 | 2.019916 | 2.219074 | 4.16163 | 4.717299 | 3.543637 | 3.806578 | 4.264968 | 3.382581 |
| Neto2 | 0.727028 | 0.928767 | 2.316844 | 2.270482 | 2.911507 | 2.8544 | 3.520966 | 3.191306 | 4.28303 | 4.368356 | 8.010983 | 8.195639 |
| 1700029P1 | 7.33429 | 7.166208 | 21.24141 | 19.62566 | 12.29469 | 14.32726 | 12.48374 | 11.25909 | 6.169724 | 5.538506 | 6.071434 | 5.438011 |
| 4632428N0 | 0.187479 | 0.152281 | 1.234611 | 1.51078 | 1.941178 | 1.694468 | 1.689299 | 1.731162 | 0.915241 | 1.06968 | 0.548444 | 0.480545 |
| Tpbg | 1.393423 | 1.353875 | 1.221005 | 1.314624 | 1.464696 | 1.592791 | 2.246757 | 2.133279 | 2.857483 | 2.880655 | 4.319624 | 4.735887 |
| Pde9a | 4.255167 | 4.779095 | 2.938599 | 2.709382 | 3.004303 | 2.740805 | 2.351794 | 2.973166 | 2.000958 | 1.546479 | 0.701543 | 0.826208 |
| Fam178a | 12.04448 | 14.39713 | 20.30284 | 20.50842 | 20.92693 | 19.08895 | 25.91646 | 21.50573 | 47.85803 | 45.90852 | 61.72712 | 68.68947 |
| Grin1 | 1.555492 | 1.212907 | 0.158513 | 0.191478 | 0.268441 | 0.282307 | 0.338555 | 0.197289 | 0.551839 | 0.688879 | 0.648262 | 0.417866 |
| Caprin2 | 21.53998 | 22.33175 | 31.99285 | 31.8779 | 39.82998 | 41.87088 | 53.16112 | 55.39875 | 22.64434 | 23.21536 | 14.9288 | 15.46353 |
| Tinagl1 | 3.046729 | 2.356547 | 6.451527 | 5.793364 | 8.384175 | 7.749405 | 8.906963 | 8.429333 | 11.35757 | 12.31232 | 26.63142 | 20.7524 |

| | | | | | | | | | | | | |
|---|---|---|---|---|---|---|---|---|---|---|---|---|
| Slfn2 | 0.121795 | 0.101256 | 0.441913 | 0.403305 | 0.82267 | 0.757643 | 1.283634 | 1.211109 | 1.131334 | 1.443886 | 1.51717 | 1.076787 |
| Lmo1 | 0.452181 | 0.312057 | 0.479622 | 0.426722 | 0.996718 | 0.811187 | 0.819751 | 0.918963 | 1.275622 | 1.526945 | 1.424226 | 0.974816 |
| Myh6 | 0.614503 | 0.583199 | 1.892306 | 1.801635 | 1.842074 | 1.596581 | 1.302379 | 1.318354 | 0.458823 | 0.503777 | 0.512924 | 0.518859 |
| B3gat1 | 0.191262 | 0.199112 | 0.571968 | 0.409068 | 0.763171 | 0.864453 | 1.074773 | 1.146709 | 1.287027 | 1.003351 | 1.67111 | 1.460907 |
| Syce3 | 1.97384 | 2.284527 | 1.92274 | 1.582569 | 1.132419 | 2.078875 | 1.24801 | 1.269317 | 1.357887 | 0.582304 | 0.489577 | 1.092715 |
| Ebf4 | 0.2299 | 0.256582 | 0.665447 | 0.769583 | 0.830888 | 0.938429 | 1.15592 | 1.445933 | 2.381629 | 2.746792 | 1.57792 | 1.473462 |
| 0610040J0: | 0.593624 | 0.405705 | 1.501116 | 0.977874 | 2.693692 | 2.130722 | 3.248543 | 3.35839 | 3.439152 | 3.486538 | 4.073498 | 2.999027 |
| Stat4 | 1.392079 | 1.442377 | 1.924863 | 1.985979 | 1.750303 | 1.854689 | 1.278034 | 1.5088 | 0.629623 | 0.68734 | 0.576434 | 0.557891 |
| Gng3 | 6.931161 | 6.975357 | 11.65702 | 11.34644 | 8.111449 | 7.809513 | 7.556943 | 7.061786 | 3.490806 | 3.187541 | 3.804911 | 3.828124 |
| Celf2 | 0.840233 | 1.011747 | 1.213011 | 1.165376 | 1.530465 | 1.384013 | 1.81893 | 1.529425 | 3.03462 | 2.845726 | 3.781399 | 4.147346 |
| Arhgap28 | 3.951891 | 4.498943 | 5.734284 | 5.169504 | 7.946843 | 7.565119 | 14.49296 | 13.30631 | 17.95651 | 18.39682 | 14.42627 | 15.06692 |
| Gdpd1 | 1.552118 | 1.841254 | 2.198954 | 2.21406 | 3.649099 | 3.311021 | 4.629617 | 4.93549 | 7.415543 | 7.878941 | 7.368019 | 7.059616 |
| Acacb | 2.927696 | 3.744969 | 3.844003 | 3.799149 | 3.122833 | 2.646377 | 2.311996 | 2.0959 | 1.307833 | 1.31569 | 1.082708 | 1.068151 |
| Usp17lb | 1.501731 | 1.124629 | 7.078302 | 6.6789 | 11.90571 | 12.13646 | 23.04463 | 23.73436 | 17.27086 | 16.96699 | 17.87822 | 14.72889 |
| Cap2 | 1.330972 | 1.11331 | 1.146851 | 1.379607 | 2.240051 | 2.018446 | 2.289919 | 2.500391 | 4.652509 | 4.900167 | 4.21338 | 4.157366 |
| Tmem253 | 3.651688 | 4.49408 | 8.966964 | 8.537952 | 6.087992 | 5.851593 | 4.480318 | 5.080133 | 2.80764 | 2.408842 | 3.893846 | 3.435834 |
| Prom1 | 3.586443 | 4.232899 | 3.208825 | 3.454259 | 5.191816 | 5.218097 | 6.795713 | 6.460972 | 11.86465 | 12.23499 | 9.579122 | 10.49453 |
| Zscan10 | 42.07035 | 46.26484 | 32.00629 | 29.419 | 16.92708 | 16.24558 | 12.64719 | 12.80246 | 8.958067 | 8.311887 | 12.64204 | 11.29793 |
| Tmem145 | 1.140331 | 1.230428 | 1.498869 | 1.451753 | 1.349624 | 1.368903 | 0.828751 | 1.611184 | 0.645964 | 0.58139 | 0.465146 | 0.455329 |
| Gm5134 | 1.299122 | 1.621654 | 1.467513 | 1.301409 | 0.965566 | 0.953112 | 0.771077 | 0.84222 | 0.429085 | 0.372975 | 0.329304 | 0.367814 |
| Gpr27 | 1.37881 | 1.867679 | 3.683272 | 2.90703 | 4.081903 | 3.876308 | 4.778209 | 4.193371 | 5.644763 | 5.132974 | 10.7616 | 10.27454 |
| Rhobtb3 | 5.030843 | 6.284757 | 5.435164 | 5.735569 | 7.858064 | 7.42926 | 9.526353 | 8.648766 | 19.57777 | 20.25369 | 18.69426 | 20.19249 |
| Adam19 | 10.08813 | 11.89121 | 3.703209 | 3.749324 | 3.205352 | 2.944258 | 3.256979 | 2.973707 | 5.099325 | 5.904512 | 9.75614 | 10.39391 |
| Nptx2 | 6.62364 | 7.037673 | 7.283295 | 6.411477 | 5.954192 | 6.025734 | 5.405092 | 5.38301 | 4.303685 | 3.734503 | 15.02655 | 13.17194 |
| Gstt2 | 11.67928 | 9.878907 | 7.018162 | 6.555613 | 6.423521 | 5.555393 | 4.294139 | 4.241985 | 2.915129 | 2.510509 | 2.193436 | 1.859239 |
| Dmrtc2 | 1.394836 | 1.48942 | 0.959733 | 1.032557 | 0.821153 | 0.683044 | 0.233032 | 0.550015 | 0.207224 | 0.33304 | 0.257058 | 0.292902 |
| Pomc | 0.438679 | 0.399101 | 0.994795 | 1.223224 | 0.93904 | 0.975784 | 0.742411 | 0.69812 | 0.570968 | 0.356288 | 1.036101 | 1.255821 |
| Nr0b1 | 64.12468 | 55.09039 | 19.13336 | 15.93026 | 9.232457 | 9.319406 | 6.227369 | 7.258178 | 4.457353 | 4.520376 | 5.843565 | 6.134618 |
| Slc25a35 | 0.982926 | 0.985586 | 1.093178 | 1.043399 | 0.916324 | 1.066376 | 0.741135 | 0.680909 | 0.429945 | 0.478178 | 0.366664 | 0.302662 |
| Sp8 | 0.630167 | 0.781376 | 3.432177 | 3.097468 | 4.030841 | 4.301955 | 6.241265 | 5.303932 | 11.81508 | 10.90879 | 10.07346 | 10.34829 |
| Mogat2 | 0.701573 | 0.595887 | 1.440516 | 1.454562 | 2.727644 | 1.998901 | 3.393969 | 3.264898 | 2.561058 | 2.761842 | 1.080307 | 0.927632 |
| Hist1h2an | 1.024304 | 1.135099 | 0.618281 | 0.554711 | 1.12709 | 1.298906 | 1.318382 | 1.550311 | 1.409054 | 1.952019 | 2.166234 | 0.766132 |
| Stat6 | 7.325944 | 8.386042 | 9.874609 | 9.409159 | 7.00648 | 7.086999 | 5.467173 | 5.723372 | 3.738381 | 4.032836 | 2.649908 | 2.678832 |
| Net1 | 20.11203 | 21.1813 | 22.01475 | 22.15989 | 23.47729 | 22.1551 | 25.00112 | 23.3248 | 42.99109 | 41.45803 | 73.81537 | 77.79421 |
| Egr1 | 0.427549 | 0.606247 | 1.649914 | 1.518199 | 1.639098 | 1.817476 | 1.86005 | 1.745496 | 1.547891 | 1.318253 | 4.818946 | 4.628088 |
| Olfml2b | 0.429628 | 0.565668 | 1.029566 | 0.870678 | 1.562054 | 1.402511 | 2.427817 | 1.968569 | 2.786766 | 2.755944 | 3.002841 | 3.056581 |
| S100g | 1.091751 | 0.796047 | 1.705165 | 1.210738 | 1.637263 | 2.216596 | 1.952613 | 3.222112 | 3.743148 | 3.873923 | 2.688227 | 4.249811 |
| Piwil4 | 1.627981 | 1.866113 | 5.523402 | 5.904523 | 5.374068 | 5.073076 | 4.960273 | 5.276803 | 1.9382 | 2.357816 | 1.497605 | 1.682272 |
| Rinl | 14.60183 | 13.66549 | 14.6431 | 13.40005 | 7.825424 | 7.398257 | 5.892766 | 7.035777 | 3.719758 | 3.818882 | 5.262127 | 4.688182 |
| Rin3 | 0.326864 | 0.410203 | 1.072838 | 0.990156 | 1.029435 | 1.073099 | 1.259469 | 1.360735 | 2.020048 | 1.963939 | 4.444306 | 3.472519 |
| Acsf2 | 9.103319 | 11.49517 | 8.121614 | 7.786334 | 9.799322 | 8.743102 | 10.56602 | 9.406109 | 26.75708 | 27.30629 | 19.59368 | 20.6577 |
| Usp17la | 0.818944 | 0.740981 | 4.855098 | 4.201523 | 8.183418 | 7.981368 | 14.22487 | 14.71655 | 10.22029 | 11.34102 | 10.29386 | 8.875946 |
| Plcl2 | 0.875683 | 1.045078 | 2.134598 | 2.138075 | 3.347018 | 3.299603 | 4.476917 | 4.513127 | 5.465741 | 6.082187 | 7.08303 | 7.486423 |
| Rhox6 | 3.41121 | 3.223435 | 4.9403 | 4.776264 | 6.442117 | 6.362058 | 7.474868 | 8.49176 | 8.712574 | 8.688927 | 20.99588 | 16.71779 |
| Sema6a | 1.151113 | 1.461793 | 0.41783 | 0.428564 | 0.399543 | 0.360642 | 0.605005 | 0.512694 | 0.848083 | 0.953235 | 1.393632 | 1.260839 |
| Rnf130 | 9.446636 | 6.646116 | 7.425961 | 7.055722 | 12.54113 | 12.41394 | 14.37326 | 16.75608 | 24.90572 | 24.66356 | 22.21213 | 23.03177 |
| Ggnbp1 | 1.318307 | 1.250104 | 1.128659 | 1.134805 | 0.911723 | 0.763601 | 0.621523 | 0.67327 | 0.476233 | 0.465134 | 0.309747 | 0.324977 |
| Cidea | 0.224745 | 0.367031 | 2.9298 | 2.964785 | 3.585898 | 2.824185 | 2.421576 | 2.697676 | 1.850173 | 2.024913 | 1.221632 | 0.849299 |
| Clgn | 6.735115 | 6.297705 | 26.86853 | 26.02713 | 23.86521 | 23.1761 | 19.90397 | 21.39167 | 10.20797 | 10.07635 | 6.810562 | 7.456317 |
| Cpa2 | 0.334087 | 0.333659 | 0.565106 | 0.490563 | 0.434933 | 0.373418 | 0.691834 | 0.91168 | 0.615884 | 0.651226 | 1.119315 | 1.303457 |
| Rragd | 10.14553 | 12.23377 | 7.105188 | 7.248069 | 8.579244 | 7.564428 | 11.93084 | 12.1363 | 22.28293 | 25.26585 | 17.12434 | 18.14213 |
| Syde1 | 1.392311 | 1.252969 | 3.73066 | 3.515874 | 3.49283 | 3.377377 | 3.530117 | 4.019623 | 5.756613 | 5.48014 | 11.64897 | 11.76364 |
| Arhgef15 | 0.96043 | 1.301584 | 1.103742 | 1.161523 | 0.952948 | 0.886298 | 0.716036 | 0.589591 | 0.424824 | 0.44479 | 0.363323 | 0.333482 |
| Hsf2bp | 23.99653 | 21.8525 | 17.56738 | 16.86136 | 11.84362 | 11.01118 | 8.818408 | 8.74569 | 5.143345 | 4.848174 | 4.98102 | 5.201585 |
| Cubn | 1.418018 | 1.981741 | 2.074467 | 2.40695 | 1.855012 | 1.858773 | 2.146651 | 1.475285 | 5.292952 | 5.120858 | 3.8112 | 5.129677 |
| Dusp27 | 10.78462 | 12.93615 | 10.87241 | 9.913394 | 6.161436 | 5.849308 | 5.495162 | 5.083097 | 3.577677 | 3.544815 | 3.137703 | 2.851992 |
| Slc9a7 | 0.281105 | 0.229268 | 0.523526 | 0.39365 | 0.664872 | 0.784207 | 0.69292 | 0.690319 | 0.794494 | 0.849886 | 1.269166 | 1.366438 |
| Nol3 | 0.168994 | 0.192233 | 0.356602 | 0.381293 | 0.821205 | 0.761064 | 1.112788 | 1.323191 | 0.741793 | 0.71028 | 0.91274 | 0.950587 |
| Cpne2 | 1.13602 | 1.072398 | 2.034871 | 2.125859 | 3.236676 | 2.868882 | 5.428511 | 4.66668 | 6.628625 | 6.244905 | 7.805442 | 7.37319 |
| Slfn9 | 9.798995 | 11.50407 | 14.27404 | 14.02403 | 9.156751 | 9.040104 | 8.260367 | 8.34917 | 3.945964 | 4.731052 | 3.908451 | 4.044158 |
| Lipg | 0.553515 | 0.562471 | 0.481217 | 0.57077 | 0.83522 | 0.763739 | 1.199537 | 0.965311 | 1.733948 | 1.978057 | 1.412716 | 1.131148 |

| | | | | | | | | | | | | |
|---|---|---|---|---|---|---|---|---|---|---|---|---|
| Celf5 | 2.692116 | 3.397501 | 3.298359 | 3.548177 | 3.306825 | 3.319797 | 2.623497 | 2.433545 | 1.321028 | 1.23242 | 1.03544 | 1.024128 |
| Tspan18 | 0.311654 | 0.395438 | 0.698355 | 0.888728 | 1.66375 | 1.699193 | 2.459443 | 1.911667 | 2.243878 | 2.568883 | 3.036266 | 3.076037 |
| Ccno | 1.0852 | 1.195074 | 1.719001 | 1.753532 | 1.643614 | 1.264001 | 2.235243 | 2.273404 | 0.995483 | 0.718523 | 0.70722 | 0.657996 |
| Wnt9a | 0.412319 | 0.439978 | 1.412397 | 1.083016 | 1.832542 | 1.60343 | 1.763373 | 1.595175 | 1.314648 | 1.30521 | 3.532228 | 3.732235 |
| Myo1f | 3.229057 | 3.363701 | 6.134768 | 5.573618 | 3.678337 | 3.657026 | 2.907254 | 3.644809 | 1.624445 | 1.617367 | 1.929441 | 1.935016 |
| Cacng6 | 2.414583 | 2.545606 | 5.554451 | 5.811207 | 5.927388 | 6.254696 | 5.251897 | 4.741233 | 3.067844 | 2.458496 | 1.996232 | 1.816396 |
| Fam57a | 2.170026 | 2.035107 | 2.474178 | 2.463658 | 3.490082 | 3.659864 | 4.089425 | 4.127085 | 6.05747 | 6.063001 | 7.952186 | 8.483167 |
| Bcam | 32.03131 | 34.70347 | 86.97896 | 79.06192 | 64.71794 | 62.24637 | 57.68815 | 54.08335 | 24.66361 | 22.96868 | 26.39519 | 23.77061 |
| Reep1 | 4.163705 | 4.032372 | 6.053728 | 5.93797 | 9.768214 | 9.273358 | 12.98477 | 13.59193 | 19.81718 | 20.43469 | 15.93886 | 16.86146 |
| Itpka | 5.885061 | 5.115621 | 4.136244 | 3.640389 | 3.123511 | 3.137941 | 2.665199 | 2.96464 | 1.270108 | 1.185187 | 1.561572 | 1.058704 |
| Bdh2 | 10.74805 | 10.86369 | 4.781176 | 5.298821 | 5.103755 | 4.985714 | 3.625485 | 3.678987 | 2.661679 | 2.286663 | 1.221559 | 1.541706 |
| Cntln | 0.524932 | 0.623335 | 0.695189 | 0.730372 | 1.221034 | 1.162595 | 1.609285 | 1.763451 | 2.273398 | 2.505775 | 2.34444 | 2.285488 |
| Hic1 | 0.14421 | 0.162843 | 2.320381 | 1.947102 | 2.244535 | 2.269252 | 1.489082 | 1.568044 | 0.685466 | 0.661488 | 1.538067 | 1.189732 |
| Wnt8a | 1.419803 | 1.494205 | 3.532856 | 3.044236 | 2.410656 | 2.325946 | 2.331354 | 2.872153 | 1.578912 | 1.544079 | 1.241255 | 0.888182 |
| Gpr1 | 0.967575 | 1.526786 | 0.926189 | 1.034024 | 0.653872 | 0.839397 | 0.557154 | 0.603957 | 0.347419 | 0.30244 | 0.363397 | 0.441048 |
| Mt1 | 102.5367 | 104.2208 | 108.2412 | 115.8183 | 105.3547 | 117.3247 | 131.2744 | 142.4239 | 116.0451 | 99.61641 | 397.9058 | 340.364 |
| Tsc22d3 | 1.697614 | 1.58101 | 2.745805 | 2.835392 | 3.675379 | 3.495466 | 4.684215 | 3.666584 | 6.851102 | 6.545826 | 10.15299 | 9.685171 |
| Lrig1 | 4.157225 | 4.149491 | 6.010719 | 6.319055 | 9.107263 | 9.60177 | 14.42178 | 14.45487 | 19.47782 | 19.74835 | 22.21911 | 21.55525 |
| 1700061G1 | 7.341237 | 8.174573 | 7.226323 | 6.70875 | 3.550363 | 3.547378 | 2.403949 | 2.468532 | 1.922618 | 1.966724 | 1.962914 | 2.058431 |
| Tusc1 | 0.717472 | 0.728749 | 1.082783 | 0.799017 | 1.173702 | 1.434551 | 1.676956 | 1.262069 | 1.628268 | 1.533618 | 2.60237 | 2.724803 |
| Fam171a2 | 1.80338 | 2.065578 | 3.428893 | 2.960511 | 4.88163 | 4.571651 | 6.449079 | 7.169853 | 10.04813 | 9.419147 | 10.94696 | 10.09341 |
| Mpped1 | 0.215474 | 0.232075 | 0.661324 | 0.528131 | 1.099527 | 1.105193 | 1.60792 | 1.799833 | 1.660755 | 1.539972 | 1.536073 | 1.603711 |
| Cdh3 | 5.234107 | 6.619087 | 2.205999 | 2.288623 | 1.110153 | 1.180783 | 0.899932 | 0.672395 | 0.796902 | 0.747415 | 2.443307 | 1.783559 |
| Snai1 | 1.207035 | 1.283113 | 2.12091 | 2.159029 | 2.9571 | 2.655865 | 3.49975 | 3.961006 | 5.250847 | 5.488213 | 7.532476 | 7.343631 |
| Pcbd1 | 2.218547 | 3.016552 | 3.786011 | 2.836708 | 4.229043 | 4.848729 | 5.964102 | 4.518706 | 10.19152 | 8.69615 | 8.991913 | 9.64663 |
| St3gal1 | 0.299532 | 0.212219 | 0.50496 | 0.505665 | 0.969902 | 0.96062 | 1.066254 | 1.278621 | 1.640786 | 1.717838 | 1.626553 | 1.535924 |
| Cplx2 | 1.273253 | 1.399496 | 1.103345 | 1.215555 | 1.471701 | 1.434644 | 1.650849 | 1.635783 | 3.305235 | 3.466455 | 3.964389 | 4.128084 |
| Tes | 3.18148 | 3.143159 | 6.601488 | 6.607977 | 13.47008 | 12.66358 | 19.90518 | 22.01095 | 22.10392 | 22.43538 | 14.45976 | 15.35308 |
| Dll3 | 2.849398 | 3.551349 | 1.112123 | 1.124897 | 1.077715 | 0.93966 | 0.864366 | 0.895093 | 1.630411 | 1.175903 | 3.749292 | 3.038453 |
| Sox7 | 1.138577 | 1.255171 | 1.107082 | 1.342233 | 1.727891 | 1.708299 | 2.699625 | 2.316354 | 4.059611 | 3.419319 | 4.553473 | 4.553722 |
| Tnfrsf25 | 1.310023 | 1.454536 | 2.08395 | 1.746861 | 1.563483 | 1.549586 | 1.275909 | 1.580107 | 0.484905 | 0.572851 | 0.552164 | 0.515132 |
| Elavl4 | 0.797835 | 0.871123 | 0.906207 | 1.10367 | 0.878817 | 0.826456 | 0.8261 | 0.869792 | 2.554782 | 2.802077 | 2.377365 | 2.509326 |
| Gdf11 | 0.903964 | 0.888645 | 1.424627 | 1.507544 | 2.262116 | 2.19071 | 3.107593 | 3.022939 | 4.346923 | 4.357806 | 4.507352 | 5.107631 |
| Krt42 | 3.22663 | 4.657664 | 2.153542 | 2.058849 | 1.227134 | 1.281306 | 0.907669 | 0.607874 | 0.941769 | 0.646649 | 1.114477 | 1.044648 |
| Tulp2 | 2.15406 | 2.288976 | 2.928365 | 2.509369 | 2.142066 | 2.052481 | 1.809656 | 1.937376 | 1.501959 | 1.277969 | 0.894407 | 0.742486 |
| Hist1h2ai | 3.119597 | 3.056163 | 0.743021 | 1.206397 | 2.717278 | 4.0729 | 3.355482 | 2.146584 | 2.316693 | 3.4907 | 2.719864 | 2.739333 |
| Ifitm5 | 1.649184 | 1.87459 | 0.614147 | 1.088767 | 0.7686 | 0.846972 | 0.595919 | 0.661965 | 1.190446 | 0.971564 | 0.299654 | 0.322587 |
| Arf2 | 9.669989 | 7.763492 | 8.212413 | 7.655965 | 9.824073 | 9.446505 | 10.32257 | 10.76534 | 12.7142 | 11.99274 | 24.84057 | 25.83959 |
| Wnt7b | 0.329754 | 0.401039 | 1.817694 | 1.503345 | 1.469861 | 1.324169 | 1.34461 | 1.100544 | 1.440271 | 1.844903 | 3.890517 | 3.714383 |
| Xlr | 0.130184 | 0.136624 | 0.446914 | 0.379483 | 0.606896 | 0.465402 | 0.577381 | 0.907105 | 0.819906 | 0.8931 | 1.203478 | 1.280289 |
| Urah | 6.28012 | 4.191891 | 3.987421 | 3.384213 | 2.986156 | 2.061329 | 2.549705 | 1.790428 | 1.380832 | 1.536889 | 1.22973 | 1.003722 |
| Lrrc8b | 2.534931 | 2.973637 | 2.068418 | 2.283473 | 2.643667 | 2.539887 | 3.583271 | 3.117874 | 6.46715 | 6.464319 | 7.405761 | 7.692987 |
| Runx1 | 0.468023 | 0.472457 | 0.476402 | 0.500223 | 0.659626 | 0.610474 | 1.044433 | 0.735672 | 1.378003 | 1.069733 | 1.793243 | 1.682584 |
| Gyg | 4.445485 | 4.628271 | 7.417871 | 6.859517 | 9.495365 | 9.154983 | 13.61815 | 11.72896 | 19.18859 | 19.0152 | 22.51866 | 23.05131 |
| Sigirr | 4.526321 | 4.9201 | 5.369279 | 5.113273 | 3.884271 | 4.064975 | 3.018588 | 3.202977 | 2.099288 | 2.377747 | 1.954785 | 1.523192 |
| Serpina3m | 0.728948 | 0.817498 | 0.359732 | 0.565988 | 0.485303 | 0.64983 | 0.71501 | 0.565004 | 2.044937 | 1.896285 | 1.179405 | 1.377876 |
| Sparc | 78.83125 | 72.75526 | 69.35686 | 74.05037 | 78.58219 | 74.98634 | 97.91562 | 103.5355 | 169.7343 | 155.3244 | 237.7123 | 248.4548 |
| Phldb2 | 2.034213 | 1.98854 | 3.465967 | 3.300026 | 4.954902 | 5.1371 | 7.251397 | 6.633022 | 9.064423 | 9.194962 | 10.41351 | 11.07139 |
| Atxn1 | 0.148532 | 0.213185 | 0.316149 | 0.331543 | 0.455531 | 0.466745 | 0.585375 | 0.424257 | 0.692945 | 0.785784 | 0.936962 | 1.112084 |
| Ubxn11 | 1.150441 | 1.408617 | 1.320194 | 1.471497 | 0.939845 | 0.923245 | 0.686937 | 0.722012 | 0.635977 | 0.53892 | 0.380828 | 0.438831 |
| Pdk1 | 97.87241 | 89.40818 | 32.85782 | 31.7727 | 24.85152 | 24.34471 | 18.73575 | 18.96377 | 14.36004 | 16.9758 | 9.620534 | 9.479903 |
| Amfr | 10.38265 | 9.359106 | 16.77161 | 15.86857 | 21.67191 | 21.31975 | 26.09985 | 28.26462 | 34.43837 | 34.32253 | 50.66806 | 53.15313 |
| Tmem88 | 4.799722 | 4.694113 | 10.37151 | 10.10568 | 9.55081 | 10.5879 | 10.72763 | 9.154191 | 3.819798 | 3.162694 | 3.852849 | 3.625887 |
| Alox5ap | 1.299969 | 1.420539 | 3.960165 | 3.104958 | 2.080988 | 2.773981 | 3.021954 | 2.805424 | 1.291678 | 1.368033 | 0.840181 | 0.927703 |
| Ppm1n | 3.320723 | 3.389068 | 2.650247 | 2.076557 | 2.790147 | 2.62351 | 1.660228 | 1.982728 | 1.177774 | 1.118169 | 1.127012 | 0.784725 |
| Sncb | 1.478308 | 1.140504 | 0.940293 | 1.592676 | 1.894819 | 1.377035 | 0.979182 | 1.110101 | 0.641781 | 0.623589 | 0.674851 | 0.477303 |
| Klf7 | 1.403418 | 1.533211 | 1.440984 | 1.54761 | 1.829708 | 1.861442 | 2.015894 | 1.914565 | 2.192318 | 2.282573 | 4.876348 | 5.161001 |
| Ripply1 | 12.19556 | 9.171495 | 4.089418 | 4.631871 | 5.220352 | 5.816847 | 15.41435 | 14.79405 | 8.773386 | 7.15194 | 14.41461 | 15.4457 |
| Enpp3 | 57.42045 | 56.61473 | 28.16598 | 27.87113 | 19.52189 | 18.69442 | 19.3314 | 18.29731 | 15.78456 | 16.02476 | 7.677985 | 8.358929 |
| Ptk2b | 1.995315 | 2.486808 | 1.880459 | 1.949423 | 1.531675 | 1.385567 | 1.653858 | 1.092094 | 0.620007 | 0.585247 | 1.083406 | 1.030081 |
| Tex12 | 0.588002 | 0.755084 | 0.867086 | 0.917374 | 1.568693 | 1.300932 | 1.089543 | 2.202829 | 1.403638 | 1.150181 | 2.476591 | 3.050075 |

| | | | | | | | | | | | | |
|---|---|---|---|---|---|---|---|---|---|---|---|---|
| Art1 | 0.852458 | 0.43242 | 1.272535 | 1.232161 | 1.861281 | 2.351706 | 1.996621 | 1.885268 | 1.188061 | 1.456273 | 0.563916 | 0.70844 |
| Stard8 | 1.255927 | 1.405848 | 1.06636 | 1.172996 | 1.415922 | 1.288257 | 1.900992 | 1.737567 | 3.248271 | 3.007053 | 3.660516 | 3.893557 |
| Igfbp4 | 3.809714 | 3.918172 | 2.770141 | 2.883261 | 4.74831 | 4.868186 | 6.345859 | 7.75613 | 10.46111 | 9.169329 | 10.43603 | 9.566539 |
| Hus1b | 1.511113 | 1.030084 | 1.716635 | 1.363029 | 0.863667 | 1.24634 | 0.886527 | 1.184509 | 0.443594 | 0.410815 | 0.780482 | 0.930168 |
| Pde3b | 3.134785 | 3.331853 | 2.792951 | 2.883229 | 4.12778 | 3.814478 | 5.065278 | 5.369073 | 8.808421 | 9.438819 | 9.880878 | 9.565561 |
| Rap1gap | 5.078034 | 5.535446 | 6.185278 | 6.460793 | 5.841524 | 6.393971 | 4.963938 | 5.445186 | 2.921068 | 2.834611 | 2.078992 | 1.948639 |
| Apobec2 | 2.117224 | 2.480992 | 2.684421 | 2.559137 | 1.604923 | 1.421425 | 1.118182 | 0.923489 | 1.116737 | 0.956949 | 0.800351 | 0.772535 |
| Acer2 | 6.863821 | 6.20048 | 7.620264 | 7.812074 | 5.709669 | 5.411659 | 4.439435 | 4.001934 | 2.93128 | 2.637227 | 2.759505 | 2.358741 |
| Mmp2 | 6.116946 | 7.094711 | 7.277595 | 7.181423 | 11.86667 | 11.13676 | 15.63341 | 16.47599 | 23.42958 | 23.76869 | 20.31172 | 19.59905 |
| Man1a | 0.647416 | 0.673688 | 0.985591 | 0.934278 | 1.121129 | 1.188738 | 1.394048 | 1.141096 | 1.532376 | 1.779524 | 2.981627 | 3.091965 |
| Ablim2 | 1.532385 | 1.364548 | 1.672137 | 1.545781 | 1.71208 | 1.538428 | 1.415853 | 1.91992 | 0.980242 | 1.287018 | 0.516193 | 0.580143 |
| Ctcfl | 0.294857 | 0.231973 | 1.144889 | 1.140337 | 1.276206 | 0.925324 | 0.819403 | 0.757566 | 0.417153 | 0.569956 | 0.542987 | 0.344758 |
| Tmem44 | 1.936203 | 2.112395 | 1.692952 | 1.971422 | 2.301509 | 2.597719 | 3.280598 | 3.367389 | 6.124559 | 6.513038 | 2.777569 | 2.730851 |
| Meiob | 1.39525 | 1.235662 | 0.680854 | 0.643505 | 0.685788 | 0.751085 | 0.417685 | 0.798089 | 0.4296 | 0.593636 | 0.219145 | 0.241613 |
| Pla1a | 2.400969 | 2.14612 | 0.642026 | 0.679912 | 0.377785 | 0.340786 | 0.314034 | 0.356494 | 0.289027 | 0.223173 | 0.388126 | 0.20613 |
| Tnnc1 | 0.560254 | 0.057463 | 0.174414 | 0.461697 | 0.474802 | 0.315764 | 0.62673 | 0.585026 | 0.382633 | 0.571862 | 1.149082 | 1.040548 |
| Sfrp4 | 3.580508 | 3.541319 | 0.854513 | 0.988216 | 0.573321 | 0.490529 | 0.421349 | 0.613863 | 0.447734 | 0.310432 | 0.34261 | 0.30011 |
| Ptprz1 | 0.27487 | 0.296885 | 0.764547 | 0.817769 | 1.338316 | 1.183729 | 2.06693 | 2.242278 | 2.467528 | 2.692247 | 1.258703 | 1.440898 |
| Mtfp1 | 9.993812 | 9.68226 | 8.845965 | 7.919475 | 7.794438 | 7.58796 | 6.44383 | 7.40706 | 4.201898 | 3.82275 | 3.194835 | 2.406204 |
| Syngr4 | 1.27302 | 0.961663 | 1.2696 | 0.875976 | 0.541175 | 0.476915 | 0.622363 | 0.82393 | 0.36784 | 0.346444 | 0.84155 | 1.13939 |
| Tcea3 | 33.42311 | 35.79272 | 12.42058 | 10.91976 | 7.328266 | 7.244553 | 6.666655 | 7.829913 | 3.609177 | 3.324341 | 3.087258 | 3.829253 |
| Hist1h2bf | 2.578636 | 3.160602 | 0.628801 | 2.025254 | 2.147839 | 3.698658 | 1.869619 | 3.456575 | 1.416023 | 2.429634 | 1.214329 | 1.126951 |
| Helz2 | 0.503924 | 0.566609 | 3.285815 | 3.546029 | 3.1178 | 3.272382 | 3.265671 | 2.92079 | 1.803496 | 2.147738 | 1.083845 | 1.080497 |
| H2-Q6 | 1.408826 | 1.138432 | 1.963779 | 2.263961 | 2.539429 | 2.969113 | 2.622915 | 2.944647 | 1.136054 | 1.168448 | 0.687759 | 0.905058 |
| Sox1 | 10.2033 | 10.21326 | 12.03475 | 12.45035 | 18.69633 | 19.84041 | 32.03653 | 26.98349 | 45.84569 | 40.84124 | 33.4413 | 32.41964 |
| Mt3 | 8.432666 | 6.640227 | 6.731051 | 6.819723 | 5.768892 | 5.193526 | 4.818131 | 5.226813 | 2.823106 | 2.28078 | 2.20414 | 2.080642 |
| Icam1 | 18.64312 | 19.04268 | 14.05065 | 13.39135 | 9.799386 | 8.681743 | 7.743838 | 8.081805 | 4.16519 | 4.086319 | 9.088902 | 8.037688 |
| Ceacam20 | 1.747588 | 2.487756 | 3.940605 | 3.85625 | 1.967274 | 1.969743 | 1.849199 | 2.070759 | 1.209731 | 1.47456 | 1.262993 | 1.177169 |
| Car14 | 4.159506 | 4.046272 | 0.626088 | 0.62124 | 0.65957 | 0.692842 | 0.568844 | 0.74044 | 2.176039 | 2.034449 | 1.377551 | 1.594549 |
| Fzd1 | 0.566161 | 0.425524 | 0.953845 | 1.011181 | 1.413167 | 1.412722 | 1.666601 | 1.857571 | 3.271498 | 3.201207 | 3.821232 | 3.30803 |
| Id1 | 15.64331 | 13.54222 | 5.986364 | 6.171548 | 5.140672 | 5.452052 | 6.958007 | 9.413821 | 19.01109 | 16.47876 | 19.95929 | 17.83357 |
| Cdk14 | 0.541098 | 0.724949 | 0.988872 | 1.252353 | 1.659644 | 1.588767 | 2.330662 | 2.05869 | 3.85232 | 3.759836 | 3.992821 | 4.095956 |
| Rbms1 | 12.92325 | 13.3016 | 23.52077 | 23.02312 | 31.65892 | 31.26797 | 40.48124 | 39.64051 | 68.51292 | 65.49881 | 70.55225 | 75.2938 |
| Spesp1 | 0.26721 | 0.267693 | 0.66921 | 0.771425 | 1.264921 | 1.234518 | 2.213369 | 2.519903 | 2.199828 | 2.120559 | 2.840302 | 2.190917 |
| Gpr124 | 0.587914 | 0.645943 | 6.503841 | 6.165327 | 7.826448 | 7.893673 | 11.21891 | 9.85348 | 8.05318 | 7.574147 | 20.25124 | 20.13397 |
| Slc22a15 | 0.12999 | 0.098358 | 0.256668 | 0.328957 | 0.437744 | 0.430393 | 0.556935 | 0.490755 | 0.567741 | 0.584758 | 1.006916 | 1.073801 |
| Cdk18 | 3.652119 | 4.177934 | 3.488598 | 3.413506 | 4.106421 | 3.493014 | 3.092739 | 3.766586 | 2.537891 | 2.598605 | 1.327403 | 1.154056 |
| Ankrd45 | 1.971781 | 2.463992 | 4.078356 | 3.470612 | 2.142746 | 2.209877 | 1.906203 | 1.574347 | 1.346116 | 1.063706 | 1.631941 | 1.589619 |
| St3gal2 | 5.684382 | 6.750835 | 10.62857 | 10.2197 | 12.74455 | 12.32756 | 15.50807 | 14.23284 | 23.96788 | 21.91311 | 31.59585 | 33.32491 |
| Ubash3b | 0.415696 | 0.387477 | 0.788643 | 0.779673 | 1.353702 | 1.490418 | 2.048096 | 1.718607 | 1.964326 | 2.140085 | 2.434493 | 2.540167 |
| Gprin1 | 0.280034 | 0.398156 | 0.938273 | 0.804539 | 0.999779 | 0.904623 | 0.99418 | 0.992947 | 1.327916 | 0.981345 | 2.744827 | 2.620889 |
| Morc1 | 50.02922 | 48.28068 | 32.01275 | 31.1767 | 23.97775 | 23.36062 | 20.16172 | 23.20305 | 11.41615 | 11.69139 | 9.000672 | 9.575664 |
| Fam57b | 1.011243 | 1.196422 | 0.620867 | 0.718006 | 0.829402 | 0.640534 | 0.858027 | 0.844452 | 0.391876 | 0.310378 | 0.372694 | 0.259403 |
| Ephx1 | 4.397628 | 5.012424 | 3.478113 | 3.367461 | 2.46444 | 2.471007 | 1.879008 | 2.23433 | 1.45578 | 1.468791 | 1.223251 | 1.034883 |
| Npm3-ps1 | 2.671064 | 2.233777 | 2.67641 | 1.771974 | 1.641788 | 2.417398 | 1.311511 | 2.130322 | 0.979829 | 0.743359 | 1.612367 | 1.002806 |
| Zbtb32 | 3.350947 | 3.305532 | 3.333444 | 3.659748 | 2.33637 | 2.791046 | 1.823354 | 2.376359 | 1.395999 | 1.640562 | 1.396556 | 1.125404 |
| Tshz3 | 0.312143 | 0.390287 | 0.674381 | 0.789669 | 1.031478 | 0.985439 | 1.481594 | 1.526076 | 2.508984 | 2.567376 | 2.256876 | 2.507792 |
| P2rx3 | 0.400131 | 0.524128 | 0.712832 | 0.985675 | 0.767767 | 0.881631 | 0.835915 | 0.75008 | 0.901248 | 0.985656 | 2.795479 | 2.437328 |
| Gli3 | 2.080925 | 2.599711 | 4.195304 | 4.432957 | 4.527104 | 4.731099 | 6.74481 | 5.116945 | 12.21137 | 11.9579 | 13.57297 | 14.39746 |
| Acp5 | 6.825059 | 6.386185 | 18.30127 | 17.23062 | 17.16685 | 15.52731 | 9.356646 | 10.69249 | 5.721312 | 5.636903 | 5.512472 | 5.313065 |
| Chst11 | 0.635133 | 0.618456 | 1.246005 | 1.351604 | 2.011763 | 2.015921 | 2.481827 | 2.560858 | 3.305005 | 3.206313 | 4.286466 | 4.381556 |
| 1700086O( | 0.901384 | 0.878039 | 1.459566 | 1.086066 | 1.416717 | 1.507139 | 2.043873 | 1.528275 | 1.812405 | 1.381495 | 3.420687 | 3.519995 |
| Aifm3 | 1.675557 | 1.66006 | 1.298286 | 1.517351 | 1.388613 | 1.368523 | 0.998755 | 1.30854 | 0.647246 | 0.82903 | 0.451777 | 0.468296 |
| Syt11 | 1.904228 | 2.025127 | 2.299581 | 2.373888 | 2.382826 | 2.481416 | 3.212482 | 3.192937 | 6.445369 | 5.970788 | 7.390462 | 7.691685 |
| Paqr4 | 0.867838 | 0.740761 | 1.208125 | 1.411208 | 1.720408 | 1.511931 | 2.131877 | 1.962922 | 1.96019 | 2.470071 | 4.470765 | 4.567105 |
| Opn3 | 1.12699 | 1.448465 | 1.312503 | 1.165363 | 1.63066 | 1.411295 | 1.825263 | 1.469956 | 3.39396 | 3.218521 | 3.610784 | 3.770121 |
| Gpa33 | 29.10822 | 28.74347 | 11.23261 | 10.37856 | 6.575514 | 6.508918 | 5.460846 | 6.056383 | 4.678622 | 4.698278 | 3.766626 | 3.209611 |
| Ptprn | 1.503295 | 1.596111 | 2.751554 | 2.424641 | 2.653071 | 2.646596 | 2.491786 | 2.650615 | 0.863553 | 0.900174 | 3.694542 | 2.909723 |
| Ngfr | 4.479148 | 5.053568 | 1.043446 | 1.034473 | 1.227057 | 0.933869 | 1.516587 | 1.441646 | 2.409912 | 2.15939 | 3.127604 | 3.0184 |
| Ppm1l | 0.747312 | 1.099109 | 1.962586 | 1.854089 | 2.364593 | 2.262589 | 3.190991 | 2.493407 | 4.484198 | 4.757732 | 5.570174 | 5.991781 |
| 4933406J0( | 2.143468 | 2.194287 | 3.263144 | 3.097373 | 1.846783 | 2.026105 | 1.787409 | 1.79491 | 0.77166 | 0.958825 | 1.661327 | 1.286182 |

| | | | | | | | | | | | | |
|---|---|---|---|---|---|---|---|---|---|---|---|---|
| Efemp2 | 1.912791 | 1.746582 | 2.099114 | 2.113274 | 2.137195 | 2.650532 | 3.082762 | 3.07886 | 5.419944 | 5.142087 | 7.435092 | 6.823224 |
| Chst2 | 0.09363 | 0.118042 | 0.331291 | 0.374004 | 0.632277 | 0.66532 | 0.918821 | 0.837826 | 0.951499 | 1.051719 | 1.414282 | 1.206062 |
| Rasal1 | 2.026763 | 1.831113 | 1.98457 | 1.772252 | 1.852069 | 1.771867 | 1.681092 | 1.499286 | 0.920716 | 0.890254 | 0.613762 | 0.551093 |
| Smagp | 23.29009 | 24.36957 | 16.66922 | 15.17523 | 9.785251 | 9.027954 | 7.325468 | 7.907786 | 5.566062 | 5.244793 | 6.335198 | 4.719329 |
| Qprt | 12.07357 | 11.8073 | 8.518458 | 8.742322 | 9.562055 | 8.846826 | 8.448993 | 8.984764 | 6.126811 | 6.152779 | 2.817277 | 2.794563 |
| Fam174b | 0.618195 | 0.602014 | 0.931209 | 0.86406 | 1.353832 | 1.111563 | 1.600208 | 2.030091 | 2.473143 | 2.441157 | 2.575424 | 2.777436 |
| Tubb4a | 1.213218 | 1.271156 | 1.273305 | 1.526684 | 2.64049 | 2.017279 | 4.233043 | 4.375384 | 5.415988 | 4.906569 | 3.43703 | 3.227772 |
| Pard6g | 4.294097 | 4.460211 | 4.158441 | 4.261232 | 5.621273 | 4.963691 | 7.269499 | 6.740033 | 14.2037 | 13.68828 | 12.91929 | 13.33472 |
| Irgm2 | 0.649587 | 0.634424 | 0.749993 | 0.77376 | 1.276386 | 1.359499 | 2.907034 | 2.485171 | 1.598783 | 1.589223 | 1.229044 | 1.215685 |
| Slc5a11 | 4.161727 | 4.756182 | 2.779047 | 2.524195 | 1.429188 | 1.391936 | 1.070203 | 1.271436 | 1.183891 | 0.785947 | 0.812099 | 0.921627 |
| Rab36 | 0.349471 | 0.349437 | 0.579 | 0.487615 | 0.489531 | 0.62698 | 0.676579 | 0.697808 | 1.118639 | 1.151034 | 1.722802 | 1.563786 |
| Smyd4 | 5.169465 | 5.484158 | 18.812 | 18.59594 | 15.9371 | 16.19789 | 15.86176 | 15.84446 | 7.432983 | 6.833884 | 6.167013 | 5.80138 |
| Galnt16 | 0.668064 | 0.577683 | 1.092364 | 1.222617 | 1.922078 | 1.742785 | 2.240815 | 2.116193 | 3.695994 | 3.884622 | 3.58188 | 3.914282 |
| Cth | 20.10338 | 17.2153 | 24.47093 | 22.34505 | 14.78006 | 14.85588 | 11.9874 | 11.31007 | 7.818745 | 7.352529 | 7.267876 | 6.980461 |
| Rarb | 0.272691 | 0.182691 | 7.091902 | 6.798285 | 11.28573 | 9.989661 | 15.55533 | 17.17739 | 21.63354 | 21.75318 | 20.01387 | 20.47563 |
| Rhobtb1 | 0.319436 | 0.268598 | 1.269126 | 1.071337 | 1.462245 | 1.71009 | 2.281328 | 2.670852 | 2.881955 | 3.042821 | 3.55303 | 3.427956 |
| Zfp600 | 10.34143 | 10.75453 | 11.1584 | 10.59815 | 6.514877 | 6.320208 | 5.193999 | 4.903155 | 3.426227 | 3.312578 | 3.348564 | 3.716086 |
| Zic3 | 46.74303 | 52.6255 | 34.27244 | 33.36313 | 29.02126 | 28.94436 | 27.94033 | 27.20166 | 16.80519 | 16.93253 | 9.721847 | 10.43027 |
| Fam151b | 0.437038 | 0.398692 | 0.334919 | 0.34899 | 0.540192 | 0.42091 | 0.563048 | 0.644804 | 0.954216 | 1.115289 | 0.813815 | 0.726908 |
| Bmi1 | 5.022663 | 4.74183 | 10.79656 | 10.66345 | 14.3513 | 14.42656 | 17.5979 | 20.14737 | 25.37845 | 26.05797 | 30.35048 | 34.03951 |
| Mesdc1 | 5.115792 | 5.314671 | 7.595832 | 6.93779 | 9.091465 | 9.225801 | 10.51913 | 11.77756 | 17.35534 | 14.85132 | 25.34353 | 22.12232 |
| Hpse | 0.155102 | 0.265111 | 0.4602 | 0.470607 | 0.949023 | 0.950622 | 1.14549 | 1.217913 | 1.50495 | 1.498286 | 1.201552 | 1.366194 |
| Ptpn13 | 7.480736 | 7.975995 | 7.560936 | 7.626831 | 10.11868 | 10.11411 | 14.83799 | 13.48844 | 23.83906 | 24.27511 | 17.02378 | 17.33172 |
| Slc4a4 | 0.277199 | 0.394479 | 0.909709 | 1.01512 | 1.67499 | 1.646047 | 2.510983 | 2.162824 | 2.722568 | 2.902002 | 2.878322 | 3.227504 |
| Mdfi | 5.839251 | 6.085355 | 5.420135 | 4.827392 | 6.863442 | 6.216134 | 5.85365 | 6.382251 | 4.564241 | 5.161351 | 2.409532 | 2.007786 |
| Gstt3 | 0.997715 | 1.104381 | 1.335542 | 1.708966 | 2.014314 | 1.61044 | 1.503421 | 1.620405 | 1.294306 | 1.292672 | 0.592867 | 0.53794 |
| Hipk2 | 2.67973 | 3.741717 | 5.628733 | 6.125895 | 6.984647 | 7.343037 | 9.863637 | 7.636335 | 18.27091 | 19.44594 | 19.13986 | 19.42915 |
| E130102H2 | 4.433832 | 4.453263 | 2.583028 | 2.288932 | 2.229886 | 2.608296 | 2.603065 | 0.82223 | 2.341555 | 2.075895 | 1.074708 | 1.270065 |
| Sema4a | 4.562758 | 4.921046 | 4.129861 | 4.123014 | 3.369787 | 3.353043 | 3.195052 | 3.451056 | 1.869168 | 2.097974 | 1.2536 | 1.30118 |
| Rab26 | 1.385332 | 1.439665 | 2.154573 | 2.083463 | 2.637873 | 2.718355 | 3.031285 | 2.713655 | 1.61589 | 1.360557 | 0.673885 | 0.858139 |
| Hist1h4k | 16.1466 | 17.45232 | 6.74276 | 15.75519 | 11.19021 | 21.6408 | 9.717931 | 11.71677 | 8.213264 | 7.491362 | 5.617342 | 6.837185 |
| Ret | 0.172991 | 0.272671 | 0.752579 | 0.850429 | 1.443138 | 1.509568 | 2.524445 | 2.623944 | 2.723586 | 2.690513 | 2.077112 | 2.003014 |
| Tmem229b | 2.429525 | 2.513866 | 11.03598 | 10.7526 | 14.75069 | 14.65336 | 14.70146 | 11.32824 | 6.233003 | 6.70258 | 4.524992 | 4.637635 |
| Abhd14b | 1.484575 | 1.229496 | 1.367637 | 1.440629 | 1.320743 | 1.428456 | 1.132092 | 1.457665 | 0.465987 | 0.482539 | 0.404945 | 0.461471 |
| Smad3 | 4.186962 | 4.895483 | 5.869183 | 6.002535 | 9.106859 | 9.087199 | 13.34528 | 11.46356 | 18.75209 | 18.77425 | 18.49466 | 18.96011 |
| Gna14 | 1.92418 | 1.564237 | 0.73778 | 0.646466 | 0.374966 | 0.340867 | 0.337443 | 0.277074 | 0.302598 | 0.349226 | 0.302801 | 0.204692 |
| Gm13580 | 2.029539 | 1.453463 | 0.790806 | 0.669322 | 0.462831 | 0.429172 | 0.666571 | 0.248412 | 0.769478 | 0.784537 | 0.865945 | 0.656306 |
| Lclat1 | 1.606545 | 1.75642 | 2.137636 | 1.864279 | 2.335602 | 2.116348 | 3.271857 | 2.706897 | 4.363754 | 4.881675 | 5.79486 | 5.887377 |
| Tmem42 | 1.737268 | 2.005038 | 2.394798 | 2.964178 | 2.468881 | 2.521202 | 2.154222 | 1.930349 | 1.835197 | 1.395581 | 4.721753 | 4.405501 |
| Bmp7 | 1.982247 | 2.149396 | 2.134345 | 2.214245 | 3.442094 | 3.00068 | 5.094603 | 4.444073 | 6.863962 | 6.989202 | 5.370137 | 4.924409 |
| Slc47a2 | 0.863957 | 0.725704 | 2.113498 | 2.242478 | 1.964818 | 1.886391 | 1.487741 | 1.557636 | 0.984628 | 1.103276 | 0.937158 | 0.710503 |
| Tspan6 | 6.969979 | 6.323264 | 9.884314 | 10.12825 | 14.53568 | 14.40293 | 18.29801 | 19.83553 | 27.36924 | 30.37892 | 29.42933 | 31.94502 |
| Zbtb8a | 14.46887 | 14.32083 | 12.16781 | 11.64371 | 7.57755 | 7.939274 | 6.385711 | 6.481834 | 4.595099 | 4.074701 | 3.533841 | 3.691997 |
| Acvr2a | 3.406473 | 3.349896 | 4.187152 | 3.950196 | 4.809014 | 4.633803 | 5.570756 | 5.563715 | 10.70759 | 10.94746 | 11.6595 | 12.45358 |
| Mgll | 0.399002 | 0.308679 | 0.582122 | 0.54356 | 0.842035 | 0.735482 | 0.978838 | 1.233035 | 1.49749 | 1.485597 | 1.998002 | 1.713383 |
| Mtss1 | 27.25509 | 30.70263 | 27.52964 | 25.32904 | 16.86638 | 17.13107 | 14.52529 | 12.37518 | 8.164037 | 8.0503 | 9.465911 | 9.763256 |
| Zfp459 | 12.09362 | 12.0088 | 2.455544 | 2.494612 | 1.635885 | 1.629802 | 1.064848 | 1.100763 | 0.855102 | 0.918023 | 0.718099 | 0.793424 |
| 1190007I07 | 15.54075 | 14.38628 | 14.86645 | 15.69977 | 11.84125 | 13.20791 | 9.394091 | 9.818524 | 6.43284 | 5.401558 | 6.264064 | 4.994287 |
| Ampd3 | 2.481271 | 2.594294 | 3.795875 | 3.419085 | 2.910954 | 2.815084 | 2.626794 | 2.449819 | 1.325288 | 1.325687 | 1.188006 | 1.087962 |
| Zmym3 | 11.88943 | 12.4252 | 30.00758 | 28.96074 | 26.59522 | 26.16426 | 22.66665 | 22.96542 | 13.48775 | 13.63423 | 9.171829 | 9.216098 |
| Slc18a1 | 0.517188 | 0.646281 | 1.373287 | 1.550356 | 1.26204 | 1.384955 | 1.18807 | 1.293923 | 0.462603 | 0.493816 | 0.439346 | 0.547765 |
| Cd47 | 1.627257 | 1.39373 | 1.611603 | 1.61423 | 3.012947 | 2.787897 | 3.870981 | 3.730183 | 5.042314 | 5.063913 | 4.888141 | 5.067107 |
| Cd37 | 1.554358 | 1.819714 | 1.644076 | 1.620696 | 1.078505 | 0.816356 | 0.65675 | 0.590219 | 0.682838 | 0.691214 | 0.651029 | 0.516496 |
| Efhd1 | 1.281313 | 1.129548 | 0.97145 | 0.860269 | 0.912634 | 1.141836 | 0.935094 | 1.061714 | 0.678076 | 0.879939 | 0.396961 | 0.364104 |
| Plekha4 | 2.36336 | 2.201596 | 2.40387 | 2.197408 | 1.361623 | 1.535837 | 1.128638 | 1.215725 | 0.78526 | 0.701258 | 0.96277 | 1.281988 |
| Shf | 4.661832 | 3.812243 | 3.10324 | 3.491145 | 4.707008 | 4.60534 | 5.925449 | 5.536661 | 9.612308 | 8.191136 | 10.1397 | 10.93327 |
| Apobec3 | 50.34331 | 49.45474 | 27.0295 | 28.05735 | 23.32715 | 22.23777 | 19.81401 | 23.13692 | 13.83118 | 15.63158 | 9.406014 | 8.961802 |
| Il17d | 11.92896 | 11.53504 | 6.059119 | 5.564195 | 3.728378 | 3.586831 | 2.69581 | 2.686893 | 2.088884 | 1.777723 | 2.295283 | 2.708583 |
| Irak3 | 6.737574 | 7.044258 | 6.230505 | 6.679615 | 4.658885 | 4.651452 | 4.600977 | 5.281438 | 2.373869 | 2.134572 | 2.976781 | 2.610468 |
| Syt17 | 0.685128 | 0.811398 | 1.942936 | 1.654727 | 2.466131 | 2.409653 | 3.046539 | 3.333157 | 1.671718 | 1.63408 | 1.396784 | 1.0661 |
| Slc30a2 | 3.38271 | 3.742193 | 8.293827 | 7.453695 | 5.927623 | 5.543218 | 4.363158 | 4.459285 | 2.317501 | 2.384613 | 3.803781 | 3.307684 |

| | | | | | | | | | | | | |
|---|---|---|---|---|---|---|---|---|---|---|---|---|
| Usp17ld | 0.263875 | 0.138317 | 1.186384 | 1.230255 | 2.303192 | 1.930835 | 4.493329 | 3.844658 | 3.938912 | 3.759206 | 3.582501 | 3.751745 |
| Rcbtb2 | 9.926302 | 8.928996 | 10.46593 | 10.79567 | 14.05385 | 14.02844 | 16.7142 | 18.19302 | 30.09063 | 29.9323 | 32.11273 | 33.72259 |
| Cilp2 | 0.359912 | 0.470605 | 1.995553 | 1.681438 | 1.954983 | 2.18107 | 1.852228 | 1.770824 | 0.92576 | 0.699355 | 1.172561 | 1.268561 |
| Tbc1d10c | 1.293562 | 0.93042 | 1.066932 | 0.885568 | 0.699571 | 0.570285 | 0.519257 | 0.586505 | 0.460996 | 0.284161 | 0.582038 | 0.463957 |
| Chst14 | 1.108297 | 1.342087 | 2.131433 | 1.70872 | 2.782969 | 2.948718 | 3.211909 | 3.139422 | 4.883488 | 4.7167 | 5.224974 | 5.320689 |
| Asgr2 | 0.176969 | 0.202187 | 0.665972 | 0.696336 | 0.851945 | 1.023645 | 1.111848 | 1.297123 | 0.556456 | 0.593406 | 0.715921 | 0.416707 |
| Onecut1 | 0.183792 | 0.375785 | 0.381365 | 0.43975 | 0.320779 | 0.350829 | 0.806978 | 0.544828 | 1.163991 | 1.091554 | 1.02182 | 0.794589 |
| Evpl | 0.439305 | 0.619945 | 1.836961 | 1.745545 | 1.658743 | 1.575024 | 1.22378 | 1.512315 | 0.645858 | 0.635292 | 0.694968 | 0.561306 |
| Asic2 | 0.310612 | 0.42593 | 0.297136 | 0.372279 | 0.680095 | 0.363389 | 0.580248 | 0.682001 | 1.308462 | 1.129575 | 1.113771 | 1.088929 |
| Pcsk9 | 2.203911 | 2.575324 | 1.872956 | 2.43162 | 3.98166 | 3.531603 | 5.665509 | 5.164564 | 7.517495 | 7.557468 | 3.236142 | 3.159307 |
| Popdc3 | 0.336517 | 0.376885 | 0.664848 | 0.667927 | 0.870628 | 1.092241 | 1.695766 | 2.073989 | 2.154196 | 1.996268 | 2.098471 | 2.014885 |
| F730043M1 | 2.130537 | 2.248709 | 2.028468 | 1.971431 | 2.157931 | 1.939775 | 1.649899 | 1.981097 | 1.11487 | 1.22149 | 0.400936 | 0.638719 |
| Srf | 15.01766 | 15.34491 | 18.06328 | 16.69671 | 17.76968 | 18.22766 | 19.76372 | 18.88195 | 20.95913 | 20.86569 | 52.65938 | 51.74597 |
| Sirpa | 1.742248 | 1.931512 | 1.721426 | 1.654434 | 2.65983 | 2.274901 | 3.725549 | 3.672158 | 5.049492 | 5.124811 | 3.49763 | 3.818073 |
| Cdkn1a | 20.05322 | 20.66848 | 20.31669 | 19.07342 | 25.59672 | 26.48387 | 32.66539 | 30.25504 | 29.21696 | 25.62846 | 58.56755 | 59.08089 |
| Aoah | 3.439922 | 3.887764 | 2.432692 | 2.020946 | 1.604549 | 1.625183 | 1.151916 | 1.221921 | 0.793502 | 0.748655 | 0.578523 | 0.65248 |
| Cmtm8 | 1.721856 | 1.354263 | 1.71075 | 1.47424 | 2.725568 | 2.947007 | 2.983433 | 3.171457 | 5.070795 | 4.562695 | 3.517414 | 3.410267 |
| Farp1 | 2.939348 | 3.442313 | 5.487618 | 5.624731 | 8.299193 | 7.541603 | 11.22318 | 10.17932 | 14.03254 | 14.14046 | 17.5148 | 17.40399 |
| Zap70 | 1.463773 | 1.349066 | 1.029122 | 1.216647 | 0.763383 | 0.869175 | 0.757363 | 0.60595 | 0.406358 | 0.393271 | 0.693857 | 0.628626 |
| 1700001L0 | 6.271293 | 6.25788 | 5.340801 | 6.198736 | 4.044234 | 4.182716 | 3.4942 | 3.633748 | 2.211198 | 2.004926 | 2.207573 | 2.104749 |
| Plek2 | 1.744955 | 1.623638 | 1.833861 | 1.944097 | 1.94553 | 1.83573 | 1.399557 | 1.746784 | 0.857981 | 0.894068 | 0.70483 | 0.629325 |
| Polr3gl | 13.70117 | 11.23672 | 9.347238 | 8.371251 | 7.817841 | 7.541817 | 6.714643 | 6.338716 | 4.570329 | 4.149344 | 2.678803 | 2.71003 |
| Rest | 55.98137 | 69.47088 | 75.76841 | 73.82703 | 51.03408 | 51.4297 | 41.4024 | 34.66853 | 25.36041 | 26.40681 | 24.09518 | 23.93577 |
| Tbc1d12 | 1.38798 | 1.213087 | 2.61352 | 2.284661 | 3.711898 | 3.922441 | 5.840829 | 6.024019 | 6.690699 | 7.0437 | 5.992973 | 6.191361 |
| Kdm4c | 34.29288 | 35.2638 | 24.5415 | 25.2184 | 21.74165 | 20.85628 | 17.57909 | 16.76537 | 11.4509 | 12.12412 | 7.976236 | 8.189548 |
| Lmna | 8.158267 | 8.271623 | 8.937019 | 7.673017 | 7.072249 | 6.578308 | 6.329155 | 5.940062 | 5.699899 | 5.200504 | 17.78339 | 16.01023 |
| Wfdc10 | 0.263469 | 0.31592 | 0.933355 | 1.054284 | 1.932257 | 1.259571 | 1.997874 | 1.531604 | 1.164603 | 0.725569 | 0.3804 | 0.497585 |
| Gfpt2 | 31.97696 | 34.965 | 29.77644 | 30.59313 | 26.56394 | 26.19222 | 22.94836 | 23.69595 | 10.80114 | 11.49674 | 11.05221 | 9.944475 |
| Pgc | 1.784402 | 1.590048 | 0.638813 | 0.806099 | 1.000841 | 0.625923 | 0.811439 | 0.371425 | 0.441192 | 0.424142 | 0.293148 | 0.262126 |
| Slc1a2 | 0.250649 | 0.32136 | 0.405698 | 0.479481 | 0.673817 | 0.756694 | 1.486847 | 1.172182 | 1.359608 | 1.473962 | 1.045341 | 1.225402 |
| Hoxb5 | 0.014369 | 0.083778 | 5.733292 | 6.066993 | 10.52641 | 10.89357 | 16.71457 | 15.22423 | 21.79526 | 18.645 | 18.88007 | 18.47251 |
| Adm | 1.523031 | 1.566882 | 0.401754 | 0.36185 | 0.550676 | 1.111575 | 0.816273 | 0.370891 | 0.55325 | 0.367965 | 0.703265 | 0.799696 |
| Stra6 | 0.491954 | 0.653963 | 10.16466 | 10.18404 | 15.00386 | 14.44724 | 27.38351 | 24.23319 | 33.65046 | 31.26551 | 25.406 | 24.78994 |
| D8Ertd82e | 2.928657 | 3.960656 | 4.055523 | 4.068505 | 4.940894 | 4.473047 | 6.341118 | 5.559339 | 9.57534 | 9.655194 | 13.08785 | 12.48951 |
| Phf2 | 2.828998 | 3.087393 | 4.407537 | 4.218922 | 5.565199 | 5.623039 | 6.060466 | 6.067773 | 10.11784 | 9.663173 | 14.00907 | 12.9499 |
| Aff3 | 2.914297 | 3.300127 | 3.122672 | 3.132393 | 3.997099 | 3.907132 | 6.322165 | 5.731666 | 9.811791 | 9.613512 | 6.567424 | 6.423426 |
| Phc2 | 9.648952 | 10.55581 | 18.24266 | 16.1041 | 24.20091 | 23.76865 | 35.60177 | 31.72341 | 51.85024 | 47.92299 | 50.72892 | 49.39128 |
| Rps6ka3 | 5.857423 | 6.609534 | 9.243087 | 9.207715 | 10.5999 | 10.23 | 13.14727 | 12.22142 | 18.53603 | 18.75694 | 25.90901 | 28.23944 |
| App | 8.125091 | 8.056442 | 8.736284 | 8.267537 | 12.62179 | 11.70835 | 15.9342 | 15.34268 | 23.82951 | 24.63913 | 24.57418 | 25.3455 |
| Frmd4b | 1.901451 | 2.076077 | 1.806065 | 1.99647 | 2.38272 | 2.208065 | 3.447915 | 3.500961 | 5.684447 | 6.118506 | 4.992463 | 5.212185 |
| Zdhhc2 | 4.011222 | 3.84778 | 4.475941 | 4.849865 | 7.529513 | 7.06922 | 9.031663 | 9.772641 | 14.18726 | 14.86298 | 14.35835 | 14.55526 |
| Kitl | 0.905721 | 0.837372 | 0.682284 | 0.867374 | 1.163594 | 0.974625 | 1.713565 | 1.876344 | 2.638227 | 2.656784 | 2.475189 | 2.591456 |
| Usp3 | 4.344213 | 4.743857 | 3.754633 | 4.001308 | 5.340944 | 5.016453 | 6.444654 | 6.285723 | 11.76273 | 12.25444 | 8.58823 | 9.121705 |
| Bcl3 | 14.85392 | 17.66318 | 31.34244 | 29.34127 | 25.08003 | 24.74647 | 20.51696 | 20.41885 | 9.779311 | 9.583482 | 21.30842 | 16.92384 |
| Nfix | 0.191751 | 0.155024 | 0.387678 | 0.503257 | 0.959426 | 0.837413 | 1.200268 | 1.069829 | 1.369306 | 1.211882 | 1.602952 | 1.54018 |
| Tuba3b | 1.440459 | 1.386019 | 5.138276 | 5.229517 | 7.832231 | 8.621957 | 13.54155 | 16.00082 | 13.29921 | 13.56663 | 17.30612 | 15.5945 |
| Ptges3l | 0.193635 | 0.245126 | 0.332173 | 0.643155 | 0.537036 | 0.47941 | 0.640639 | 0.800873 | 0.692065 | 0.841429 | 0.984932 | 1.466492 |
| Hoxb8 | 0.130404 | 0.137499 | 23.04855 | 22.81339 | 47.38126 | 48.86771 | 77.74716 | 69.687 | 86.8706 | 69.41341 | 70.1725 | 67.12927 |
| Dgkk | 0.154992 | 0.223203 | 0.583578 | 0.693332 | 1.016151 | 1.015887 | 1.470858 | 1.27075 | 1.428355 | 1.606558 | 1.903799 | 2.117557 |
| Fbn1 | 0.38611 | 0.503089 | 1.606364 | 1.532229 | 2.284986 | 2.112781 | 3.308756 | 2.533039 | 4.044712 | 3.862368 | 4.834183 | 4.679206 |
| Ccnb1ip1 | 20.80045 | 20.07315 | 27.1871 | 26.84988 | 18.84175 | 19.28492 | 14.16831 | 15.25978 | 7.245956 | 8.794173 | 9.521219 | 9.11027 |
| 1700001G1 | 0.395903 | 0.442283 | 1.258209 | 1.124125 | 0.684439 | 0.732675 | 0.825127 | 0.490755 | 0.308202 | 0.368431 | 0.435787 | 0.561599 |
| Loxl2 | 3.206846 | 3.559056 | 2.056864 | 2.357564 | 3.811508 | 3.744234 | 4.502031 | 3.905591 | 5.18378 | 5.716225 | 7.412709 | 7.190409 |
| Ctdsp1 | 9.403304 | 9.227912 | 12.59758 | 11.97981 | 15.57851 | 15.88744 | 17.04298 | 16.21408 | 22.31345 | 19.62216 | 38.10137 | 36.5343 |
| 1700067K0 | 0.592188 | 0.865444 | 1.357122 | 1.170314 | 0.936321 | 1.147151 | 0.912362 | 0.798665 | 0.621552 | 0.383891 | 0.621054 | 0.536516 |
| Prkd1 | 0.613555 | 0.905854 | 1.740465 | 1.714543 | 2.578824 | 2.293885 | 3.83221 | 3.0641 | 4.055568 | 4.090884 | 5.298703 | 5.225032 |
| Epha2 | 41.79122 | 46.49615 | 47.05984 | 44.52537 | 34.57636 | 32.90136 | 28.47383 | 26.85306 | 15.25323 | 14.61325 | 22.71682 | 23.45892 |
| Zfp493 | 2.567921 | 2.192608 | 3.321667 | 3.434177 | 2.261514 | 2.325389 | 1.777636 | 1.844882 | 1.067422 | 1.272946 | 1.131445 | 1.127651 |
| Lyz2 | 10.889 | 2.311272 | 1.415914 | 1.839473 | 1.388484 | 0.863597 | 1.441993 | 2.628407 | 0.970876 | 1.542132 | 3.884607 | 1.893462 |
| Tex14 | 6.813559 | 7.466415 | 10.61556 | 10.04801 | 7.521781 | 7.41639 | 6.607683 | 7.586506 | 3.439725 | 3.83671 | 3.6393 | 3.303324 |
| Ptges | 1.51428 | 1.462251 | 4.359884 | 3.740875 | 5.932537 | 6.083078 | 9.385267 | 9.316874 | 9.571454 | 8.409024 | 11.57995 | 11.37335 |

| | | | | | | | | | | | | |
|---|---|---|---|---|---|---|---|---|---|---|---|---|
| Draxin | 0.372314 | 0.403437 | 0.392265 | 0.402313 | 0.437979 | 0.360175 | 0.694681 | 0.657942 | 0.868898 | 0.970903 | 0.938676 | 1.094903 |
| Cldn12 | 9.372762 | 10.23214 | 11.25083 | 11.23634 | 12.2923 | 11.58225 | 13.85472 | 13.12828 | 17.67568 | 18.07638 | 32.39408 | 34.15468 |
| Cenpw | 16.8453 | 12.16663 | 20.0608 | 16.93175 | 18.27985 | 18.69157 | 14.66541 | 11.91852 | 7.179394 | 6.15048 | 8.62538 | 8.228531 |
| Otud7a | 0.590652 | 0.656111 | 0.486502 | 0.646412 | 0.930671 | 0.84803 | 1.283396 | 1.466699 | 1.734742 | 1.962322 | 1.785776 | 1.857345 |
| Slc39a6 | 6.42664 | 7.085913 | 8.508177 | 9.290993 | 10.42819 | 10.52471 | 12.88715 | 12.71279 | 21.46109 | 20.95425 | 27.26858 | 28.19974 |
| Grip1 | 1.527373 | 1.677868 | 2.091165 | 2.334796 | 4.13444 | 3.924154 | 5.503296 | 5.08995 | 7.240133 | 7.085572 | 6.375437 | 6.717242 |
| Tekt1 | 2.464969 | 2.631726 | 2.44076 | 1.950798 | 1.697336 | 1.272975 | 0.90269 | 1.13362 | 0.777277 | 0.928774 | 0.568809 | 0.643277 |
| Ramp2 | 1.678568 | 1.545542 | 0.398021 | 0.511307 | 0.872233 | 0.99141 | 1.028579 | 0.792745 | 1.93911 | 1.550399 | 1.234778 | 1.210497 |
| Grrp1 | 0.791388 | 0.464489 | 1.300403 | 1.043801 | 1.659344 | 1.550317 | 1.182559 | 1.529106 | 1.498469 | 0.874762 | 0.951755 | 0.511617 |
| Gata4 | 3.236393 | 3.207085 | 3.859783 | 3.395537 | 4.140865 | 4.355178 | 5.530744 | 5.210518 | 10.67421 | 8.597597 | 10.53421 | 10.28734 |
| Usp25 | 4.17229 | 4.449089 | 7.127165 | 7.721237 | 9.349665 | 9.547676 | 11.37768 | 11.1219 | 19.76583 | 20.00288 | 23.2307 | 23.39064 |
| Amot | 1.650172 | 1.854016 | 2.572913 | 2.379282 | 3.205836 | 3.06434 | 4.851687 | 4.068955 | 7.364159 | 7.206293 | 6.343293 | 6.503864 |
| Fzd5 | 16.19533 | 18.90962 | 13.49259 | 12.79344 | 8.914162 | 9.036998 | 7.746482 | 7.028369 | 4.855486 | 4.647518 | 4.776242 | 4.227425 |
| Epor | 0.451892 | 0.706527 | 0.816913 | 0.834341 | 1.212559 | 1.061863 | 0.669913 | 0.845606 | 0.518245 | 0.424834 | 0.339943 | 0.351202 |
| Gm13939 | 3.419941 | 2.9618 | 0.71416 | 0.646816 | 0.34471 | 0.514116 | 0.397015 | 0.382163 | 0.087645 | 0.279189 | 0.112499 | 0.213969 |
| Foxc2 | 0.037136 | 0.015543 | 0.253424 | 0.346017 | 0.664683 | 0.502697 | 0.482398 | 0.567565 | 0.572934 | 0.691804 | 1.272963 | 1.045166 |
| Akt3 | 6.551036 | 6.781099 | 5.986363 | 6.183737 | 9.866465 | 9.486385 | 12.8668 | 11.93172 | 16.86673 | 18.67527 | 14.70368 | 16.08596 |
| Fgf15 | 5.256285 | 5.142092 | 0.369964 | 0.445018 | 0.400635 | 0.381652 | 0.509229 | 0.699509 | 0.302793 | 0.291474 | 0.286561 | 0.231678 |
| Vnn1 | 0.367069 | 0.359181 | 1.058662 | 0.965065 | 0.833907 | 0.876402 | 1.299768 | 1.377303 | 1.69485 | 2.060151 | 2.420617 | 2.645259 |
| Hormad1 | 14.98156 | 12.77496 | 16.70164 | 15.64074 | 9.530257 | 10.93841 | 8.308729 | 8.777886 | 5.166389 | 5.185872 | 7.096958 | 9.967347 |
| Jam3 | 8.705847 | 7.5692 | 9.461015 | 8.79713 | 12.40037 | 13.24463 | 16.37696 | 17.94707 | 25.02515 | 26.5135 | 21.85254 | 25.84381 |
| Phactr2 | 0.752233 | 0.667866 | 1.726458 | 1.821746 | 2.52336 | 2.374762 | 2.859583 | 2.823114 | 4.95374 | 5.063656 | 4.951285 | 5.489591 |
| Gm13247 | 24.79859 | 25.03981 | 18.17471 | 18.15904 | 11.05937 | 11.04371 | 7.388294 | 8.531106 | 5.99293 | 6.030345 | 5.802823 | 6.766825 |
| Zhx2 | 0.364724 | 0.478561 | 0.781267 | 0.849711 | 1.136895 | 1.029943 | 1.5053 | 1.681475 | 2.537463 | 2.248571 | 3.143698 | 2.558564 |
| Arid3a | 2.931065 | 3.443827 | 3.539271 | 3.48641 | 4.996214 | 4.70552 | 6.308965 | 6.084591 | 8.390172 | 9.052718 | 11.51288 | 10.49027 |
| Igf1r | 4.297107 | 5.617545 | 6.391446 | 6.984005 | 7.893405 | 7.508407 | 9.981071 | 8.310582 | 14.1262 | 14.4441 | 20.75136 | 21.0142 |
| Egln1 | 8.136586 | 7.316005 | 7.999394 | 8.595617 | 11.73027 | 10.93882 | 13.643 | 13.02678 | 24.63466 | 23.62235 | 25.78728 | 25.862 |
| Csn3 | 0.127089 | 0.072606 | 7.444189 | 8.302921 | 10.21496 | 10.81566 | 13.33083 | 10.49823 | 4.210792 | 3.59525 | 7.904389 | 7.671383 |
| Tsacc | 2.421895 | 2.539661 | 2.149177 | 2.712863 | 2.114186 | 1.781111 | 2.224607 | 1.171677 | 1.185671 | 0.901947 | 0.749771 | 1.095755 |
| S100a6 | 4.513821 | 3.739881 | 12.92495 | 10.90389 | 6.858021 | 7.872889 | 7.974666 | 5.279881 | 4.224843 | 3.625514 | 13.85806 | 10.01386 |
| Ndrg2 | 12.30081 | 12.45814 | 3.527693 | 3.246613 | 2.997799 | 2.894311 | 2.388235 | 2.911539 | 1.904013 | 2.614572 | 1.555037 | 1.079612 |
| Cnr1 | 0.889456 | 0.872041 | 0.916511 | 0.849983 | 1.507746 | 1.501074 | 1.450692 | 1.363964 | 1.961051 | 2.060002 | 2.376675 | 2.552179 |
| Pdlim7 | 9.849867 | 10.18272 | 18.73727 | 17.48322 | 18.41155 | 19.00997 | 19.22908 | 17.13082 | 17.0444 | 16.2622 | 54.12729 | 48.79919 |
| Fzd8 | 0.927156 | 0.978643 | 1.735212 | 1.749073 | 2.630073 | 2.735116 | 3.787391 | 3.487718 | 4.163582 | 3.772568 | 5.901586 | 5.244392 |
| Dab2 | 5.778735 | 5.28609 | 5.748904 | 5.861625 | 6.969567 | 6.777753 | 9.534874 | 8.523116 | 19.71932 | 17.56077 | 14.9625 | 17.12869 |
| Pdzd3 | 0.572964 | 0.444527 | 0.323479 | 0.590992 | 0.581526 | 0.469543 | 0.491764 | 0.580198 | 1.18337 | 0.890686 | 1.635383 | 1.404793 |
| Vmn1r53 | 0.968558 | 0.86985 | 2.182723 | 2.151247 | 1.3646 | 0.937481 | 0.914899 | 1.21136 | 0.998856 | 0.719057 | 1.402621 | 0.986273 |
| Caps2 | 1.442044 | 1.272558 | 1.432971 | 1.371703 | 1.312673 | 1.529518 | 1.016768 | 1.222045 | 0.494926 | 0.511296 | 0.382779 | 0.540229 |
| Tmem82 | 0.876034 | 0.862902 | 1.248847 | 0.895824 | 0.681029 | 0.821881 | 0.761614 | 0.614544 | 0.346268 | 0.372113 | 0.447663 | 0.299603 |
| Hist1h1d | 7.024107 | 8.284545 | 1.939512 | 3.839625 | 4.09949 | 5.859075 | 3.779058 | 5.039472 | 1.959964 | 4.468578 | 1.906066 | 1.959655 |
| Rasl11b | 3.806063 | 3.323766 | 5.972708 | 5.624722 | 7.603158 | 8.073793 | 11.891 | 12.45473 | 12.11813 | 10.45141 | 17.40612 | 16.80725 |
| Pacsin1 | 2.662154 | 2.573305 | 3.840135 | 3.625298 | 4.219959 | 3.455897 | 3.623978 | 2.941777 | 1.968528 | 1.793507 | 1.311489 | 1.213288 |
| 9530068E0 | 16.34868 | 14.36341 | 13.94571 | 14.00535 | 17.6121 | 16.99754 | 22.74451 | 21.99265 | 40.18417 | 41.00461 | 41.70908 | 41.84701 |
| Padi3 | 0.743392 | 1.060638 | 1.31106 | 1.531066 | 1.685542 | 1.26415 | 1.516942 | 1.628313 | 1.122612 | 1.281583 | 0.454748 | 0.545082 |
| Cradd | 7.219383 | 6.976289 | 5.975765 | 6.337251 | 5.202802 | 5.005269 | 4.543885 | 3.891389 | 4.236956 | 3.521274 | 2.422122 | 2.121753 |
| Ppfia4 | 5.113387 | 6.000561 | 6.120978 | 5.476993 | 4.474648 | 4.730246 | 3.788512 | 4.642677 | 2.440512 | 2.614796 | 1.870889 | 1.834361 |
| Cela1 | 1.260981 | 1.038427 | 1.203879 | 1.438892 | 1.498781 | 1.271379 | 1.301132 | 1.317207 | 0.33899 | 0.483468 | 0.553863 | 0.482294 |
| Oas1g | 1.489025 | 1.621405 | 1.174068 | 1.249092 | 0.855957 | 0.812806 | 0.644448 | 0.663773 | 0.443724 | 0.554491 | 0.543998 | 0.418858 |
| Myc | 15.12142 | 17.79271 | 11.35457 | 10.51382 | 7.306181 | 6.761775 | 6.06921 | 5.824069 | 9.395886 | 7.250464 | 17.52279 | 17.34771 |
| Klhl29 | 0.234968 | 0.247804 | 0.473147 | 0.582938 | 0.820988 | 0.826997 | 0.940219 | 0.898602 | 1.664297 | 1.735613 | 1.689066 | 1.723158 |
| Pbx1 | 5.435179 | 6.170491 | 16.80879 | 18.94293 | 30.84168 | 29.65124 | 44.18797 | 38.70147 | 53.79461 | 56.3561 | 37.07272 | 39.35955 |
| Sowahc | 1.263741 | 1.03802 | 1.653974 | 1.391032 | 1.712641 | 1.583253 | 1.913632 | 2.070434 | 3.103194 | 2.931763 | 3.532594 | 4.137277 |
| Bnip3 | 76.42092 | 60.00303 | 13.10184 | 16.44353 | 21.58975 | 19.74412 | 14.10846 | 16.3074 | 17.65886 | 24.01343 | 7.722171 | 8.076023 |
| Hist3h2ba | 1.936077 | 1.445561 | 2.662535 | 2.105658 | 4.131959 | 4.898503 | 5.891337 | 6.259123 | 5.019671 | 3.189867 | 3.610393 | 3.535082 |
| Gpx8 | 3.708199 | 3.121935 | 6.763512 | 5.623697 | 9.480581 | 9.227448 | 10.86671 | 9.825482 | 13.48334 | 11.93868 | 15.96363 | 16.71122 |
| Fzd2 | 6.255542 | 6.08172 | 9.251935 | 8.622382 | 14.19487 | 14.08718 | 22.03176 | 20.382 | 26.30372 | 25.62068 | 24.36233 | 22.56288 |
| Lamb1 | 17.52273 | 20.43646 | 29.47457 | 28.50595 | 30.23528 | 28.78679 | 41.66369 | 36.40626 | 76.0094 | 71.84595 | 78.61609 | 84.69144 |
| Zfp90 | 0.390496 | 0.458788 | 0.629222 | 0.43532 | 0.680786 | 0.737898 | 0.945902 | 0.801299 | 0.951029 | 1.015199 | 1.059798 | 1.292854 |
| Il4ra | 0.43579 | 0.419406 | 0.698623 | 0.758075 | 1.170889 | 1.109347 | 1.910092 | 1.905241 | 1.871661 | 1.783313 | 2.627524 | 2.251117 |
| Atf4 | 70.48827 | 55.74878 | 141.0985 | 130.0107 | 97.88572 | 105.4469 | 78.08002 | 80.98437 | 65.76528 | 54.445 | 161.8118 | 161.6469 |
| Nck2 | 6.115722 | 7.515076 | 7.012523 | 7.614192 | 10.25152 | 9.928742 | 13.15165 | 13.45823 | 23.01914 | 22.60251 | 20.99858 | 22.52403 |

| | | | | | | | | | | | | |
|---|---|---|---|---|---|---|---|---|---|---|---|---|
| Egfr | 0.239006 | 0.296611 | 0.493604 | 0.433979 | 0.509203 | 0.460221 | 0.527671 | 0.426983 | 0.583589 | 0.618465 | 1.282838 | 1.267363 |
| Cd3e | 0.64429 | 0.855325 | 2.149254 | 2.074412 | 1.374389 | 1.422216 | 1.084902 | 1.30549 | 0.737533 | 0.734368 | 0.920142 | 0.698948 |
| Pml | 40.48398 | 48.0012 | 53.63305 | 50.35611 | 39.07487 | 35.65342 | 31.47282 | 29.73513 | 23.11459 | 22.5067 | 17.6415 | 16.97189 |
| Gulp1 | 1.55986 | 1.583682 | 2.737314 | 2.422205 | 4.160159 | 3.972859 | 4.873701 | 4.510472 | 6.950271 | 7.186086 | 3.153303 | 3.683333 |
| Ncmap | 0.231857 | 0.212689 | 1.50267 | 1.339533 | 1.157572 | 1.348548 | 0.993079 | 1.007642 | 0.439122 | 0.454946 | 0.721833 | 0.467985 |
| 6330403K0 | 0.997395 | 0.881288 | 0.80321 | 1.127768 | 1.456266 | 1.5809 | 2.40022 | 2.729208 | 3.422652 | 3.34233 | 2.792677 | 3.02178 |
| Rbp1 | 1.085395 | 0.869861 | 13.99575 | 15.34194 | 28.0841 | 29.34278 | 37.86788 | 45.41796 | 28.71444 | 26.19167 | 30.14592 | 29.03859 |
| 2410137M | 3.473946 | 3.91387 | 3.526094 | 4.300287 | 2.189902 | 1.886201 | 2.581443 | 2.654899 | 1.920548 | 1.918068 | 1.344492 | 1.453345 |
| Cd109 | 0.484897 | 0.55463 | 0.52231 | 0.531341 | 0.566946 | 0.658376 | 0.714372 | 0.61583 | 0.699618 | 0.755337 | 1.549699 | 1.571103 |
| Rcn2 | 12.77589 | 11.4736 | 12.18516 | 11.52204 | 14.87396 | 15.67669 | 18.13234 | 19.9946 | 28.84889 | 27.39582 | 33.05364 | 34.06416 |
| Hkdc1 | 0.349542 | 0.300537 | 0.356959 | 0.459068 | 0.555065 | 0.512397 | 0.552218 | 0.519709 | 1.084127 | 1.104122 | 1.046511 | 1.357066 |
| Dapk2 | 1.161914 | 1.058432 | 1.214878 | 1.024211 | 1.172277 | 1.268641 | 1.031878 | 1.190191 | 0.602203 | 0.521322 | 0.466892 | 0.429194 |
| Ophn1 | 0.334382 | 0.438464 | 0.512794 | 0.627594 | 0.882169 | 0.91294 | 1.237553 | 1.36614 | 1.836195 | 1.85503 | 1.413918 | 1.626736 |
| Parm1 | 0.764187 | 0.972772 | 1.51446 | 1.809953 | 2.106125 | 1.993877 | 2.750814 | 2.200941 | 4.896808 | 5.348717 | 5.510312 | 4.936902 |
| Sox10 | 0.081465 | 0.086712 | 0.879125 | 0.878522 | 1.587984 | 1.441819 | 1.598577 | 1.4511 | 1.9999 | 2.347696 | 0.938392 | 0.795339 |
| Fam20c | 0.831843 | 0.601217 | 1.508427 | 1.235113 | 1.517536 | 1.765163 | 1.870569 | 2.185037 | 2.320754 | 1.795756 | 3.677448 | 3.6414 |
| Acot11 | 0.644558 | 0.693824 | 1.059537 | 1.316906 | 1.31672 | 1.137404 | 1.183997 | 1.425208 | 0.821394 | 0.751571 | 0.369179 | 0.483445 |
| Anxa5 | 7.723352 | 6.592078 | 8.192965 | 7.989221 | 12.49625 | 12.18933 | 17.28582 | 18.2385 | 17.6876 | 17.08794 | 24.03273 | 23.5502 |
| Emx2 | 0.013729 | 0.028732 | 0.547476 | 0.582997 | 1.152873 | 1.381557 | 1.788372 | 1.526877 | 1.629818 | 1.281378 | 1.52986 | 1.717648 |
| Dmrta2 | 0.227294 | 0.275082 | 1.544224 | 0.921557 | 1.722294 | 2.175764 | 2.393551 | 1.395427 | 2.087516 | 1.859442 | 3.08682 | 2.714786 |
| 1110002L0 | 3.886236 | 4.34835 | 4.468303 | 4.53395 | 3.521464 | 4.266552 | 2.97075 | 3.085953 | 2.092772 | 2.486579 | 1.548576 | 1.539735 |
| Erf | 49.7699 | 48.25558 | 166.898 | 156.8252 | 165.4369 | 168.7679 | 152.4215 | 138.6891 | 63.46837 | 57.34279 | 69.94772 | 63.70876 |
| Tgfbr2 | 0.64868 | 0.653837 | 0.91449 | 0.966169 | 1.288278 | 1.151757 | 1.584059 | 1.390831 | 2.179717 | 1.999599 | 2.546907 | 2.842846 |
| Gpm6a | 0.888268 | 0.827014 | 0.970521 | 0.823904 | 1.721742 | 1.77878 | 2.543379 | 2.346164 | 2.306167 | 2.422481 | 1.579219 | 1.43597 |
| Crip1 | 29.04691 | 20.48658 | 14.77149 | 13.59535 | 15.66322 | 17.38804 | 12.71384 | 9.478737 | 7.264108 | 6.562133 | 9.840432 | 5.915668 |
| Gm12794 | 0.888776 | 1.044077 | 5.485647 | 5.655856 | 9.032026 | 8.204776 | 15.82206 | 16.62297 | 12.7842 | 13.44763 | 11.48207 | 9.186705 |
| Apoc1 | 19.03796 | 16.84059 | 17.89169 | 15.45329 | 16.84814 | 17.87806 | 10.53505 | 9.238687 | 8.505494 | 6.517139 | 7.091817 | 6.085427 |
| Tex21 | 2.331259 | 1.807799 | 1.607071 | 0.893537 | 0.812621 | 1.054904 | 0.605212 | 1.163991 | 0.351834 | 0.515548 | 0.413992 | 0.396207 |
| Ttc28 | 0.429156 | 0.582882 | 1.125038 | 1.054181 | 1.753691 | 1.519234 | 2.440012 | 1.952071 | 3.055593 | 3.095359 | 2.768412 | 2.75346 |
| Cdkn1c | 12.53487 | 12.0326 | 17.63867 | 15.67808 | 13.33109 | 13.64096 | 16.24328 | 16.38285 | 28.71689 | 30.9548 | 49.5586 | 40.04068 |
| Krt7 | 0.645715 | 0.39953 | 0.5831 | 0.723784 | 0.767793 | 0.750803 | 0.692948 | 0.778447 | 1.035513 | 1.379995 | 3.000131 | 2.124476 |
| Slc35g2 | 0.177119 | 0.213679 | 1.156871 | 1.16083 | 2.089243 | 1.933919 | 3.136443 | 2.564426 | 3.236264 | 3.403612 | 3.196747 | 3.404296 |
| Panx1 | 4.569529 | 4.317755 | 5.338462 | 4.934597 | 7.21124 | 7.142868 | 9.243317 | 10.54032 | 11.78814 | 12.29742 | 13.83581 | 14.45773 |
| Kirrel | 5.298266 | 6.876654 | 7.21577 | 7.368924 | 8.577316 | 8.817646 | 12.73181 | 10.70898 | 17.18169 | 17.11414 | 20.22111 | 21.57611 |
| Slc52a3 | 1.93427 | 2.206083 | 2.102682 | 1.800084 | 1.192438 | 1.35333 | 1.349798 | 1.475023 | 0.847373 | 0.729337 | 0.692769 | 0.614859 |
| Galt | 7.208253 | 7.512385 | 7.748369 | 7.091221 | 6.426143 | 6.605423 | 4.381526 | 5.596823 | 3.451863 | 3.332204 | 2.49364 | 2.424277 |
| Gfod1 | 2.529667 | 3.079103 | 1.060913 | 1.171529 | 0.768854 | 0.747764 | 0.560498 | 0.704345 | 0.43117 | 0.400682 | 0.693884 | 0.671634 |
| Oas1c | 2.241395 | 1.718469 | 1.001982 | 0.816524 | 1.318667 | 1.354489 | 1.168865 | 1.022852 | 0.834982 | 0.836683 | 0.678815 | 0.463314 |
| Rtkn | 1.066711 | 1.036858 | 1.356358 | 1.040131 | 1.458909 | 1.507637 | 1.988174 | 1.810774 | 2.588333 | 2.470098 | 3.59026 | 3.04059 |
| AU015836 | 1.681855 | 2.160503 | 3.637018 | 3.53181 | 4.396276 | 4.016347 | 5.453283 | 5.676594 | 9.273358 | 10.31756 | 8.253908 | 8.149042 |
| Sox4 | 7.544244 | 8.562418 | 15.59349 | 15.12456 | 24.25747 | 24.26829 | 33.18176 | 32.4893 | 44.00857 | 44.16005 | 43.24948 | 37.98339 |
| 1600016N2 | 2.238084 | 2.181776 | 2.400042 | 2.038674 | 2.351714 | 2.353392 | 1.842502 | 2.254309 | 1.526262 | 1.422619 | 0.853994 | 0.806194 |
| Sertad4 | 1.368624 | 1.244859 | 3.583913 | 3.23815 | 4.735622 | 4.732489 | 4.794091 | 5.384964 | 4.01724 | 3.630744 | 8.868848 | 9.451728 |
| Extl1 | 1.29974 | 1.577129 | 1.745546 | 1.385482 | 1.19641 | 1.173598 | 0.83074 | 0.999589 | 0.650743 | 0.609189 | 0.582217 | 0.474974 |
| Foxh1 | 30.31623 | 28.02037 | 39.26176 | 34.66919 | 33.5486 | 32.71548 | 29.28324 | 27.56043 | 13.50167 | 11.89203 | 14.93709 | 13.45531 |
| Gm1631 | 5.433463 | 4.39175 | 2.82974 | 2.989174 | 2.293203 | 2.931631 | 1.190172 | 2.632161 | 1.337031 | 1.07928 | 0.919852 | 1.025668 |
| Dusp2 | 0.374801 | 0.339932 | 0.37171 | 0.487286 | 0.410771 | 0.596311 | 0.849619 | 0.637647 | 0.950548 | 1.001579 | 1.720723 | 1.419927 |
| C330046G1 | 12.39959 | 11.69298 | 3.265854 | 2.515777 | 1.177937 | 1.501914 | 1.075823 | 1.826903 | 1.119656 | 1.052066 | 0.850616 | 0.863377 |
| Zfp41 | 10.13894 | 10.17135 | 11.51865 | 11.42748 | 13.78464 | 13.21767 | 16.70766 | 15.64327 | 33.19022 | 33.28625 | 29.48063 | 28.40998 |
| Jazf1 | 0.228664 | 0.305929 | 0.784656 | 0.699816 | 0.896615 | 0.935206 | 0.79677 | 1.021162 | 0.874327 | 0.922289 | 1.436108 | 2.035555 |
| Plcb3 | 26.84511 | 29.03038 | 67.70057 | 64.97139 | 62.63189 | 59.3142 | 59.99719 | 64.97356 | 37.64837 | 38.49042 | 22.65761 | 22.34685 |
| Mtap7d3 | 8.15092 | 7.286968 | 4.080767 | 3.993828 | 3.234267 | 3.297652 | 2.318099 | 3.080162 | 1.708265 | 1.723712 | 1.307134 | 1.373954 |
| Ttpa | 3.610686 | 3.576326 | 3.325126 | 3.94389 | 2.656983 | 2.403604 | 1.826791 | 1.815856 | 1.485333 | 1.669305 | 1.122261 | 1.35788 |
| 0610043K1 | 1.542161 | 1.414828 | 1.024068 | 0.644839 | 0.819006 | 0.90146 | 0.834793 | 0.610646 | 0.844219 | 0.555275 | 0.288785 | 0.310607 |
| C1qtnf6 | 0.788878 | 0.723318 | 1.141142 | 0.904547 | 1.477705 | 1.458329 | 2.266861 | 2.399938 | 2.724424 | 2.624315 | 1.783894 | 1.930625 |
| Pnma2 | 3.145759 | 3.74362 | 4.48496 | 4.62172 | 4.31135 | 4.060251 | 3.723061 | 3.482321 | 2.049996 | 2.24585 | 1.506454 | 1.59359 |
| Stmn2 | 137.3877 | 120.4389 | 56.31392 | 55.54966 | 43.79705 | 43.43034 | 37.93365 | 41.57485 | 23.10425 | 21.97277 | 20.63091 | 19.15614 |
| Cdk6 | 3.278309 | 4.490695 | 2.10004 | 2.195887 | 1.62936 | 1.526238 | 1.504885 | 1.555962 | 1.932161 | 1.862787 | 4.6276 | 4.424799 |
| Fez1 | 5.649369 | 5.204348 | 6.604599 | 6.127209 | 7.053706 | 7.137388 | 6.532068 | 6.844581 | 18.91493 | 15.47472 | 15.57991 | 17.76123 |
| Spock1 | 0.366697 | 0.304317 | 0.331681 | 0.346949 | 0.615536 | 0.719265 | 0.871171 | 1.005601 | 0.908967 | 0.893197 | 1.155228 | 0.953369 |
| Hoxb2 | 0.051627 | 0.05494 | 1.537022 | 1.254134 | 2.798038 | 2.803073 | 3.492873 | 3.633919 | 3.315043 | 2.43262 | 2.457396 | 2.025042 |

| | | | | | | | | | | | | |
|---|---|---|---|---|---|---|---|---|---|---|---|---|
| Bhmt | 0.229111 | 0.213511 | 0.72876 | 0.744247 | 0.965477 | 0.952207 | 1.1797 | 1.147138 | 2.151151 | 2.155921 | 2.134223 | 1.881793 |
| 5830418P1 | 0.73267 | 0.843993 | 1.773267 | 1.826616 | 1.396843 | 1.367463 | 1.343171 | 1.226549 | 0.652803 | 0.630714 | 0.568452 | 0.725885 |
| Vcan | 3.100455 | 3.817588 | 2.894687 | 2.826101 | 3.255258 | 3.13632 | 4.18831 | 3.910934 | 7.774234 | 8.180542 | 7.63065 | 8.111292 |
| Oplah | 1.879753 | 1.833557 | 1.618536 | 1.59406 | 1.982659 | 1.766522 | 2.174539 | 1.950405 | 3.710473 | 3.897963 | 4.176901 | 4.613064 |
| Letmd1 | 1.804934 | 2.164534 | 4.679419 | 4.979515 | 6.111068 | 6.341483 | 8.276447 | 8.813702 | 13.02501 | 13.14028 | 14.29299 | 14.40982 |
| Cox6b2 | 54.87496 | 49.16252 | 38.48951 | 41.27081 | 44.09671 | 44.10406 | 35.63969 | 33.8084 | 19.19453 | 16.36732 | 16.91061 | 15.24564 |
| Tdpoz3 | 0.603863 | 0.384051 | 1.275363 | 1.35867 | 1.668592 | 1.683298 | 3.903444 | 3.926933 | 3.305857 | 3.708284 | 3.04885 | 3.256593 |
| Plin2 | 13.90381 | 13.67403 | 30.00485 | 30.40048 | 33.91778 | 33.38443 | 44.50049 | 44.55849 | 45.24309 | 41.92405 | 93.26287 | 87.80913 |
| Parva | 1.608018 | 1.808655 | 5.289907 | 5.403185 | 7.603104 | 7.243679 | 10.89653 | 10.6527 | 11.81079 | 11.82043 | 15.72215 | 15.60441 |
| Zfp711 | 0.339513 | 0.274518 | 0.472064 | 0.468714 | 0.762693 | 0.582053 | 0.819424 | 0.887552 | 1.316956 | 1.353232 | 1.035187 | 1.235586 |
| Skap1 | 0.115581 | 0.191835 | 1.034689 | 0.87914 | 1.586791 | 1.32922 | 1.393067 | 1.476481 | 0.584128 | 0.697345 | 0.50763 | 0.511412 |
| Rgma | 0.444841 | 0.53277 | 4.706277 | 4.348055 | 6.787226 | 5.59423 | 9.282642 | 8.207622 | 13.28061 | 11.97621 | 13.29929 | 12.54864 |
| Ube2j1 | 5.652569 | 5.205349 | 6.934705 | 6.215821 | 8.839618 | 8.64421 | 9.803038 | 10.83387 | 14.42714 | 13.81338 | 17.60085 | 17.92422 |
| Rpl10l | 62.69416 | 65.73718 | 150.2669 | 137.4126 | 107.0976 | 105.5248 | 95.61874 | 100.8811 | 52.94032 | 47.65458 | 68.55102 | 58.62376 |
| Ubtd2 | 12.56684 | 13.98059 | 17.29581 | 15.35269 | 18.75084 | 19.40448 | 23.76085 | 20.74749 | 37.62544 | 36.69037 | 42.28989 | 44.2421 |
| Layn | 0.20329 | 0.191779 | 0.395021 | 0.355514 | 0.857268 | 0.801612 | 1.177269 | 1.023413 | 1.052478 | 0.877939 | 0.627753 | 0.99468 |
| Slco2a1 | 0.708599 | 0.557059 | 0.482733 | 0.548726 | 0.441101 | 0.558297 | 0.610902 | 0.579093 | 0.999469 | 1.163968 | 1.527052 | 1.579481 |
| Serpinf1 | 1.485327 | 1.08859 | 8.787544 | 8.373154 | 13.00115 | 12.24954 | 16.83345 | 19.57059 | 20.94731 | 18.60326 | 23.78302 | 24.09802 |
| Rhox10 | 0.396422 | 0.167736 | 1.028003 | 0.789933 | 1.400656 | 1.539535 | 1.645129 | 2.272716 | 1.342975 | 1.366577 | 1.788069 | 1.21691 |
| Snn | 8.499573 | 7.640999 | 7.502607 | 7.164392 | 11.13705 | 11.00239 | 15.40578 | 14.22809 | 20.50029 | 20.61236 | 14.21644 | 15.03718 |
| Gm13128 | 0.553142 | 0.535025 | 1.043116 | 0.980155 | 0.865809 | 1.051993 | 1.696462 | 2.363498 | 1.8897 | 1.994813 | 2.389858 | 2.81978 |
| 5430417L2 | 0.61957 | 0.62276 | 1.245487 | 1.571104 | 2.243922 | 2.329963 | 2.301572 | 2.410132 | 3.611507 | 3.411337 | 4.423192 | 4.518962 |
| 1500015O1 | 0.334716 | 0.397582 | 0.245752 | 0.570051 | 0.86925 | 1.212099 | 1.404792 | 1.298977 | 1.395959 | 1.639378 | 1.394922 | 1.4493 |
| Gpsm1 | 4.37065 | 4.494681 | 5.560239 | 6.010728 | 9.26675 | 9.148294 | 11.4086 | 12.34494 | 16.36144 | 16.83113 | 17.7068 | 17.27209 |
| Pik3ap1 | 0.293208 | 0.301572 | 0.730311 | 0.65685 | 0.746835 | 0.83962 | 1.094969 | 1.290413 | 2.015712 | 1.886729 | 1.000954 | 1.496944 |
| Sh3bp4 | 5.318063 | 5.811067 | 6.733113 | 6.642215 | 7.441091 | 7.468899 | 9.659961 | 8.763536 | 13.42675 | 12.9451 | 18.28107 | 19.07508 |
| Zfp113 | 1.652093 | 1.746813 | 1.831653 | 1.778557 | 2.188487 | 2.013868 | 3.027985 | 2.832308 | 4.26043 | 4.618897 | 4.826749 | 5.101357 |
| Glrb | 0.406041 | 0.364462 | 1.100553 | 1.031185 | 1.694738 | 1.65235 | 1.975562 | 2.614902 | 3.109696 | 2.957375 | 2.424227 | 2.7653 |
| Rbm46 | 0.371199 | 0.421402 | 0.941262 | 1.10519 | 1.076349 | 0.886277 | 1.52133 | 1.08581 | 1.3723 | 1.338384 | 2.262093 | 2.541443 |
| BC051019 | 0.505013 | 0.392856 | 0.951901 | 0.766306 | 0.741811 | 0.705579 | 0.804282 | 0.886092 | 0.630017 | 0.413264 | 1.454609 | 1.184857 |
| Mthfd2l | 2.056495 | 1.993882 | 1.8892 | 1.896377 | 2.343999 | 2.358371 | 2.689621 | 2.70508 | 4.235206 | 4.147178 | 4.721258 | 5.43521 |
| Igfals | 1.634058 | 1.538547 | 1.255037 | 1.321921 | 1.144559 | 0.847192 | 0.878389 | 0.999062 | 0.472514 | 0.57656 | 0.597314 | 0.461226 |
| Rhd | 1.89279 | 1.837507 | 5.596066 | 5.555647 | 3.870631 | 4.437175 | 3.808803 | 3.559918 | 2.074435 | 2.59251 | 1.722197 | 1.939155 |
| Samd12 | 1.360136 | 1.653979 | 2.926416 | 2.503997 | 2.665091 | 3.329944 | 2.999171 | 2.987721 | 2.555807 | 3.138381 | 1.222957 | 1.16282 |
| Ulk2 | 1.859776 | 1.91912 | 2.288795 | 2.334305 | 3.608523 | 3.639272 | 4.323186 | 4.390474 | 6.204817 | 6.466419 | 6.445758 | 6.678854 |
| Marcks | 19.55394 | 23.12822 | 25.474 | 24.39059 | 30.33958 | 30.23209 | 44.7999 | 36.75964 | 77.76439 | 68.85863 | 69.01635 | 69.78564 |
| Arnt2 | 3.3584 | 3.607778 | 4.736642 | 4.468282 | 6.676718 | 6.446143 | 7.755693 | 6.989364 | 10.98597 | 11.11835 | 11.64202 | 12.78413 |
| Ppef2 | 1.072657 | 1.371577 | 1.33285 | 1.215338 | 0.974382 | 1.087319 | 0.959264 | 0.762569 | 0.519496 | 0.556166 | 0.369099 | 0.424882 |
| Hoxa1 | 1.370886 | 1.37352 | 15.73437 | 16.3502 | 30.27032 | 30.85679 | 46.92411 | 46.76299 | 36.85101 | 35.38224 | 30.56537 | 30.15297 |
| Fgf5 | 2.266175 | 2.071347 | 0.16874 | 0.263065 | 0.161243 | 0.103848 | 0.212476 | 0.206133 | 0.307084 | 0.200532 | 0.255568 | 0.296988 |
| Cobl | 26.74311 | 32.50136 | 19.0979 | 19.2126 | 12.15253 | 11.83694 | 10.43772 | 9.774063 | 7.045869 | 6.822068 | 6.496529 | 6.720666 |
| Rapgef3 | 2.358683 | 2.417448 | 2.242976 | 1.999896 | 1.547365 | 1.256405 | 1.207288 | 1.705677 | 0.520176 | 0.699737 | 0.864998 | 1.051084 |
| Rnf103 | 2.391367 | 1.915371 | 3.447818 | 3.404214 | 5.558614 | 5.247386 | 6.209595 | 6.734327 | 7.811069 | 8.110955 | 9.656055 | 9.725616 |
| 1700007K1 | 10.69705 | 9.947647 | 9.286918 | 8.234351 | 6.62205 | 6.671518 | 5.683775 | 4.155703 | 3.250674 | 2.883244 | 4.663863 | 4.348021 |
| Hist2h2ac | 1.432797 | 1.960092 | 0.47547 | 1.12997 | 0.582416 | 0.780842 | 0.611498 | 0.630903 | 0.728673 | 0.873056 | 0.474991 | 0.395805 |
| Ankrd13b | 4.710592 | 4.909819 | 4.784853 | 4.833061 | 6.169007 | 6.550156 | 7.459075 | 7.066591 | 15.05066 | 13.78246 | 11.94905 | 13.58999 |
| Mkrn1 | 188.8267 | 184.9073 | 144.2398 | 138.1976 | 116.2946 | 110.1901 | 104.8054 | 104.4431 | 65.50267 | 61.91985 | 49.00751 | 48.47879 |
| Etv4 | 38.02594 | 38.14245 | 39.51199 | 35.85229 | 25.54517 | 25.15851 | 17.69796 | 17.78797 | 13.02866 | 12.58886 | 18.28324 | 16.5888 |
| Rara | 14.19283 | 17.89869 | 31.47012 | 30.39378 | 24.21098 | 23.16979 | 16.56033 | 16.28714 | 11.53312 | 11.07974 | 11.69831 | 10.67481 |
| Myh7b | 1.200165 | 1.198905 | 3.099097 | 2.857306 | 3.113946 | 3.015909 | 2.685351 | 2.50191 | 1.54202 | 1.49496 | 1.084607 | 1.059853 |
| Cd164l2 | 1.371138 | 0.949682 | 1.063557 | 0.705919 | 0.693975 | 0.958857 | 1.017754 | 1.093249 | 0.343304 | 0.993131 | 0.366334 | 0.384317 |
| Spn | 1.033143 | 1.192906 | 0.753375 | 0.599939 | 0.597379 | 0.327418 | 0.500061 | 0.525893 | 0.318068 | 0.363309 | 0.19869 | 0.211168 |
| Wdr1 | 41.34645 | 46.53921 | 45.68408 | 43.89894 | 44.0845 | 43.21746 | 53.66429 | 49.8241 | 54.97979 | 49.94626 | 124.8444 | 122.7599 |
| Cables1 | 1.374124 | 1.452763 | 1.430602 | 1.40153 | 1.941094 | 2.021005 | 2.216529 | 2.266648 | 3.579663 | 3.475154 | 3.961418 | 3.980424 |
| Fam114a1 | 2.10794 | 2.111747 | 2.07243 | 1.921129 | 3.669793 | 3.029067 | 4.412005 | 4.98143 | 5.609513 | 5.448862 | 3.113432 | 3.29138 |
| Nrxn2 | 1.355878 | 1.534027 | 3.217454 | 3.167669 | 3.422472 | 3.388185 | 3.522725 | 3.149311 | 1.911083 | 1.865973 | 1.263113 | 1.195059 |
| Kcnt1 | 0.20568 | 0.201648 | 0.702022 | 0.747709 | 1.000067 | 0.994843 | 0.955746 | 1.099755 | 0.503629 | 0.706814 | 0.455807 | 0.388297 |
| Rpa1 | 60.76045 | 59.83821 | 198.5874 | 189.5188 | 193.9065 | 195.5984 | 208.7342 | 215.4575 | 95.81314 | 98.76265 | 76.43917 | 76.0818 |
| Fam81a | 7.174251 | 7.327301 | 7.150292 | 6.956825 | 6.043264 | 5.424849 | 4.106885 | 4.45431 | 3.483008 | 3.714721 | 3.069426 | 2.457487 |
| Mxra7 | 5.806451 | 5.164159 | 6.131867 | 5.894677 | 8.760423 | 8.312701 | 11.43845 | 10.63199 | 18.72601 | 16.68472 | 13.66285 | 12.62 |
| 1700008I0 | 0.193791 | 0.22837 | 0.571258 | 0.466224 | 0.997855 | 0.857869 | 1.117118 | 1.319456 | 1.046921 | 1.129572 | 0.981375 | 0.743178 |

| | | | | | | | | | | | | |
|---|---|---|---|---|---|---|---|---|---|---|---|---|
| Rhox9 | 1.323683 | 1.246696 | 1.649162 | 1.666783 | 2.29542 | 2.041107 | 2.309224 | 2.825184 | 2.91362 | 2.436589 | 6.330646 | 4.713956 |
| Gpr4 | 2.117238 | 2.413385 | 2.043487 | 1.857392 | 1.522188 | 1.374111 | 1.109627 | 1.186303 | 0.908232 | 0.833752 | 0.619782 | 0.656964 |
| Bmper | 0.675041 | 0.632227 | 0.56475 | 0.723602 | 0.679865 | 0.770682 | 0.968005 | 0.803153 | 1.506936 | 1.280384 | 1.76731 | 2.044903 |
| Fblim1 | 73.3256 | 72.72014 | 76.90617 | 75.57462 | 63.52245 | 60.12575 | 52.21905 | 53.6833 | 27.76542 | 26.74713 | 39.90476 | 38.11944 |
| Tgfbr1 | 7.923834 | 8.118392 | 9.290414 | 10.46566 | 12.00002 | 11.70503 | 15.18577 | 14.50144 | 23.56179 | 25.11214 | 27.04497 | 29.56914 |
| Lrrc16b | 0.797172 | 0.7211 | 0.555197 | 0.5825 | 0.690695 | 0.621082 | 0.689892 | 1.064231 | 1.742852 | 1.645557 | 1.311813 | 1.575157 |
| Nyap1 | 1.577498 | 1.972784 | 3.326271 | 2.982863 | 3.2732 | 3.326225 | 2.95061 | 3.102908 | 1.687083 | 1.651391 | 1.35926 | 1.177697 |
| Arap3 | 1.219041 | 1.243522 | 1.942119 | 1.794633 | 2.189832 | 2.028014 | 2.425165 | 2.574819 | 1.716221 | 2.067279 | 5.766697 | 5.066186 |
| Bok | 9.097291 | 8.2178 | 7.050584 | 7.009738 | 7.888819 | 7.087099 | 8.798886 | 7.406788 | 11.57521 | 10.4502 | 21.48971 | 19.76935 |
| Hist1h3d | 6.827012 | 7.141356 | 1.625213 | 4.416742 | 3.714123 | 6.076342 | 4.368265 | 6.77899 | 3.406697 | 4.782801 | 2.638128 | 2.406441 |
| Bmp2k | 1.077511 | 1.556762 | 1.880989 | 1.920668 | 2.055843 | 2.135081 | 2.569961 | 2.39189 | 2.902242 | 2.759839 | 5.285441 | 5.409776 |
| Pced1b | 22.60534 | 23.20891 | 28.67829 | 27.31073 | 22.68676 | 21.46505 | 18.65815 | 18.69993 | 11.34398 | 10.76647 | 11.01901 | 9.699348 |
| Mcoln3 | 0.700637 | 0.720322 | 0.70264 | 0.814669 | 1.245119 | 1.129947 | 1.694997 | 1.7393 | 1.98157 | 2.293122 | 1.582101 | 1.668695 |
| Hspa4l | 1.799279 | 1.739311 | 2.582273 | 2.660828 | 3.556464 | 3.288951 | 3.84302 | 3.708231 | 5.448823 | 5.760823 | 7.239643 | 7.486782 |
| Pstpip2 | 3.974185 | 4.462377 | 4.414145 | 3.592266 | 3.694341 | 2.908635 | 3.111358 | 2.150175 | 1.75376 | 2.604579 | 1.181139 | 1.276922 |
| Camkv | 9.194084 | 9.622789 | 5.549033 | 5.783668 | 5.508523 | 5.748105 | 4.36412 | 5.102652 | 4.07492 | 3.74933 | 1.73187 | 2.058094 |
| Fhl3 | 4.55769 | 5.518249 | 7.554807 | 7.460011 | 7.951873 | 7.428893 | 8.741885 | 6.928668 | 7.830691 | 7.928938 | 21.29623 | 19.45367 |
| Thpo | 2.261557 | 2.756522 | 2.799988 | 2.800596 | 2.175677 | 2.328034 | 2.204547 | 2.1353 | 1.409515 | 1.200722 | 0.811828 | 0.998118 |
| Sh2d5 | 1.010872 | 1.031607 | 1.198898 | 1.306704 | 0.902363 | 1.031726 | 0.620883 | 0.624186 | 0.461337 | 0.567907 | 0.397678 | 0.465774 |
| Wfikkn1 | 0.427744 | 0.703203 | 1.109294 | 1.645309 | 2.806414 | 2.827607 | 4.455248 | 4.424102 | 3.265237 | 3.197054 | 1.242538 | 1.578377 |
| Zfyve28 | 3.066953 | 3.129813 | 8.663143 | 8.996512 | 9.147294 | 8.607398 | 8.230057 | 8.853453 | 4.867191 | 5.059182 | 3.402852 | 3.210883 |
| Klhl3 | 0.492236 | 0.60419 | 1.360851 | 1.153855 | 0.96924 | 0.980767 | 1.024171 | 0.876159 | 0.455325 | 0.599166 | 0.526742 | 0.411851 |
| Ptpn21 | 4.275734 | 4.613546 | 5.788223 | 5.859311 | 5.78295 | 5.669336 | 5.964304 | 5.651692 | 8.274235 | 8.584109 | 14.87407 | 15.83029 |
| Mmp15 | 1.939775 | 2.465502 | 4.832993 | 4.921023 | 6.987795 | 6.777741 | 9.078012 | 7.265046 | 12.12924 | 11.75652 | 14.26676 | 13.7694 |
| Tmem35 | 0.336709 | 0.342993 | 0.523412 | 0.593285 | 0.798058 | 0.892257 | 1.788537 | 1.588301 | 1.65177 | 1.659756 | 1.480155 | 1.499463 |
| Fbxo4 | 0.940423 | 0.997787 | 1.086822 | 1.087878 | 1.223458 | 0.923551 | 0.995476 | 1.167934 | 1.869287 | 1.761874 | 2.996225 | 2.582945 |
| Gstm5 | 10.04054 | 6.941991 | 6.047923 | 7.283803 | 11.27516 | 12.08245 | 13.97883 | 16.44459 | 20.23215 | 20.3688 | 17.33461 | 18.25828 |
| Cbs | 6.352154 | 6.245761 | 4.720404 | 4.158216 | 3.161347 | 2.917694 | 2.308269 | 2.721557 | 1.680732 | 1.775853 | 1.603155 | 1.48732 |
| Zfp667 | 3.295934 | 3.151456 | 2.972436 | 2.824573 | 2.748838 | 2.569599 | 2.224172 | 2.276583 | 1.87988 | 2.153327 | 1.051645 | 1.010666 |
| Zfp768 | 1.83013 | 1.922238 | 3.494801 | 3.557443 | 4.933311 | 4.823796 | 5.892731 | 5.511319 | 7.721924 | 7.08228 | 10.42516 | 9.940267 |
| 1110019D1 | 3.847755 | 3.237281 | 4.392249 | 4.238432 | 3.428983 | 3.084722 | 2.845731 | 2.938964 | 1.950381 | 2.304922 | 1.453019 | 1.516871 |
| Kifc3 | 8.50865 | 9.34012 | 9.177849 | 8.520879 | 6.829993 | 6.622528 | 5.715379 | 6.227865 | 4.058818 | 3.982056 | 3.843852 | 3.04955 |
| Srcin1 | 1.41076 | 1.630456 | 4.808262 | 4.540428 | 4.545338 | 4.454034 | 3.862034 | 4.331917 | 2.254851 | 2.435524 | 1.51016 | 1.625601 |
| Fam98c | 2.220712 | 2.317291 | 2.865481 | 2.112074 | 2.108718 | 2.303888 | 1.681178 | 1.815046 | 1.269145 | 1.108623 | 0.783215 | 0.824935 |
| Mettl7a1 | 5.640574 | 5.080848 | 3.708224 | 4.159058 | 4.600836 | 4.721631 | 5.036007 | 4.928138 | 3.252321 | 2.965548 | 1.850818 | 1.76586 |
| Ndst1 | 5.950192 | 6.808655 | 8.032037 | 8.198988 | 9.771488 | 9.154383 | 11.95828 | 10.85587 | 18.21237 | 18.92121 | 22.15899 | 22.86114 |
| Wnk2 | 0.907298 | 1.382684 | 1.672209 | 1.85939 | 2.667605 | 2.478071 | 3.191099 | 2.874406 | 5.039876 | 5.181226 | 4.203415 | 4.165831 |
| Msc | 3.657665 | 3.476351 | 1.57194 | 1.346795 | 0.855705 | 0.784361 | 1.173637 | 1.157544 | 0.918814 | 0.870983 | 1.915269 | 2.185027 |
| Gm5091 | 1.927407 | 1.836773 | 1.91089 | 1.866862 | 1.863031 | 1.610432 | 1.559134 | 1.431056 | 0.58371 | 0.901202 | 0.817629 | 0.670401 |
| Ptpla | 7.217811 | 5.471207 | 5.337156 | 5.488773 | 5.606431 | 6.317592 | 4.976746 | 6.493633 | 9.348437 | 10.33533 | 17.04135 | 15.26281 |
| Lrig3 | 6.470048 | 6.386741 | 4.896518 | 5.215015 | 6.162884 | 5.830114 | 7.20527 | 8.181489 | 14.3114 | 14.49828 | 8.955958 | 8.72739 |
| Gm2381 | 14.13303 | 16.31344 | 13.21409 | 15.03677 | 10.99647 | 10.91204 | 8.857982 | 8.883229 | 6.415812 | 7.744086 | 4.518659 | 5.412352 |
| Rnf219 | 3.104633 | 3.057906 | 4.270897 | 4.646436 | 5.716278 | 5.133491 | 6.074097 | 6.550001 | 11.31368 | 11.66229 | 11.63517 | 12.89992 |
| Podxl2 | 0.542206 | 0.631703 | 0.866163 | 1.131681 | 1.871612 | 1.638437 | 1.852562 | 2.818431 | 2.959834 | 3.140906 | 2.659576 | 2.63854 |
| Dusp4 | 14.7341 | 15.02304 | 12.87963 | 12.88633 | 15.07082 | 14.98169 | 18.69388 | 17.39556 | 35.34635 | 35.75582 | 35.50725 | 34.15009 |
| Cdkn2a | 4.005141 | 2.242132 | 7.649332 | 6.49422 | 10.84781 | 8.804485 | 13.65531 | 13.14373 | 15.75999 | 12.80162 | 17.82124 | 18.01468 |
| Snx32 | 1.563908 | 2.110676 | 3.595014 | 2.989714 | 3.320321 | 3.152557 | 2.644835 | 3.23809 | 1.179605 | 1.811733 | 1.345147 | 1.168008 |
| Bves | 0.512761 | 0.538853 | 1.527954 | 1.572601 | 1.894178 | 1.510872 | 2.457952 | 2.508133 | 3.557442 | 3.580839 | 3.839645 | 4.186413 |
| Mark1 | 2.160507 | 2.40125 | 2.428696 | 2.344099 | 3.572022 | 3.575475 | 4.16918 | 4.824857 | 6.401854 | 5.958027 | 6.329979 | 6.49098 |
| Ncs1 | 2.487413 | 3.11656 | 4.501518 | 4.150228 | 5.365255 | 4.897437 | 6.37284 | 5.157092 | 9.264896 | 9.225167 | 11.31923 | 11.4915 |
| Prrt3 | 0.574102 | 0.645255 | 5.142791 | 4.922615 | 4.156588 | 4.595859 | 4.524078 | 3.957602 | 1.994453 | 1.778836 | 2.717516 | 2.66796 |
| Itgb4 | 0.818791 | 0.980044 | 2.069971 | 1.87772 | 1.722532 | 1.805613 | 1.821155 | 1.819421 | 1.354991 | 1.27659 | 0.756204 | 0.678658 |
| Stard13 | 0.436415 | 0.485524 | 0.655502 | 0.856033 | 0.966354 | 0.797359 | 0.84303 | 0.828459 | 1.235306 | 1.008629 | 2.284882 | 2.206021 |
| Rgs10 | 1.374232 | 1.157188 | 3.792975 | 2.966064 | 5.428022 | 5.336787 | 6.675715 | 8.201741 | 6.76146 | 5.419749 | 4.957644 | 4.670875 |
| Zfp362 | 2.603882 | 3.540617 | 6.384448 | 6.764817 | 10.28582 | 9.860669 | 13.57818 | 12.45011 | 20.62688 | 18.70161 | 17.30287 | 16.45348 |
| Epm2a | 1.030744 | 0.778005 | 1.093276 | 0.929276 | 1.525872 | 1.531107 | 2.207496 | 2.095973 | 2.509014 | 2.519642 | 2.964652 | 2.568954 |
| Emilin1 | 1.50706 | 1.886736 | 5.071617 | 5.212005 | 9.486214 | 9.254589 | 15.90021 | 14.40193 | 14.25604 | 14.28426 | 7.314051 | 6.282275 |
| Itgb3 | 0.429265 | 0.688793 | 0.837247 | 0.68579 | 1.138605 | 1.078549 | 1.614284 | 1.482828 | 1.679381 | 1.894486 | 1.562033 | 1.506748 |
| Cyb5r1 | 39.59156 | 44.20894 | 40.74204 | 40.02049 | 30.7777 | 30.57333 | 24.46188 | 27.07446 | 17.43004 | 17.13338 | 13.73084 | 14.49244 |
| Hoxa2 | 0.575526 | 0.366704 | 1.810656 | 2.078794 | 3.691194 | 3.755067 | 4.416086 | 4.405116 | 5.732617 | 5.738106 | 5.422322 | 5.620269 |
| Rab20 | 0.696744 | 0.438074 | 0.42599 | 0.741986 | 0.934097 | 1.155917 | 1.05764 | 1.037169 | 2.002226 | 2.04675 | 1.09584 | 1.226956 |

| | | | | | | | | | | | | |
|---|---|---|---|---|---|---|---|---|---|---|---|---|
| Slc35e4 | 0.716842 | 0.930186 | 1.134254 | 1.231976 | 1.576852 | 1.431722 | 1.614414 | 1.882028 | 1.724158 | 1.674833 | 4.056614 | 3.397449 |
| Etv6 | 4.442977 | 5.429711 | 4.240088 | 4.471422 | 6.179017 | 5.593946 | 8.584926 | 7.119987 | 11.57001 | 11.57576 | 11.83862 | 12.32503 |
| Fstl1 | 20.43668 | 22.30952 | 15.16025 | 15.58397 | 22.26485 | 20.7445 | 34.20044 | 31.91267 | 41.79776 | 42.9469 | 31.38788 | 33.44493 |
| Pfn2 | 7.285997 | 6.397262 | 9.183644 | 9.095003 | 13.80635 | 13.67627 | 17.32017 | 16.31244 | 23.6746 | 21.96633 | 24.04977 | 25.04679 |
| Fsip1 | 0.22832 | 0.392119 | 0.726185 | 0.518879 | 0.746488 | 0.870487 | 0.880352 | 1.428704 | 0.799149 | 1.062825 | 0.917534 | 1.025849 |
| Klf6 | 9.12646 | 9.592807 | 11.23084 | 10.34425 | 12.60757 | 11.39928 | 12.10657 | 10.99159 | 12.80043 | 12.18724 | 25.47942 | 28.46939 |
| Dgka | 11.43031 | 11.11361 | 8.666157 | 8.558961 | 7.613699 | 7.813163 | 5.85526 | 6.225579 | 4.775995 | 4.554654 | 3.15121 | 3.110796 |
| Psors1c2 | 0.644722 | 1.607718 | 1.135245 | 1.26808 | 1.410862 | 0.733138 | 0.746153 | 1.223103 | 0.568137 | 0.460904 | 0.73755 | 0.925595 |
| Ltbp3 | 0.535656 | 0.50143 | 1.695696 | 1.405567 | 1.287838 | 1.063227 | 0.853221 | 1.119753 | 0.603793 | 0.64622 | 0.848437 | 0.510991 |
| Mobp | 1.563887 | 1.730906 | 0.574047 | 0.886412 | 0.864729 | 0.791736 | 0.884347 | 1.141013 | 1.810376 | 2.177728 | 1.745331 | 1.439432 |
| Nckap1l | 3.018575 | 3.788603 | 8.061691 | 7.920213 | 7.168014 | 7.208093 | 6.330674 | 6.905369 | 3.320211 | 3.306798 | 3.106186 | 2.879534 |
| Lasp1 | 48.7329 | 54.41081 | 107.1099 | 103.6411 | 82.68659 | 77.78897 | 75.14567 | 61.73481 | 38.71172 | 37.68289 | 54.20003 | 48.36773 |
| Rhou | 5.495455 | 6.108428 | 7.47425 | 7.706136 | 10.32396 | 9.917143 | 11.21245 | 11.039 | 15.77129 | 16.13856 | 21.82319 | 21.19227 |
| Nbl1 | 0.585161 | 0.329157 | 0.551586 | 0.821397 | 1.100512 | 1.070076 | 1.30919 | 1.20239 | 1.479955 | 1.23384 | 1.888003 | 2.258755 |
| Hsd11b2 | 0.852249 | 0.824906 | 0.568383 | 0.637416 | 0.972169 | 1.068332 | 2.019516 | 1.752753 | 1.612489 | 1.622323 | 0.852976 | 1.039163 |
| Zxdb | 0.290572 | 0.266186 | 0.493505 | 0.539867 | 0.648115 | 0.64171 | 0.718709 | 0.617221 | 1.30658 | 1.255166 | 1.260926 | 1.483494 |
| Sh3rf1 | 3.488084 | 4.203062 | 5.430413 | 5.264057 | 6.243516 | 6.133572 | 7.851938 | 6.585455 | 11.16965 | 12.43246 | 14.91147 | 14.46384 |
| Gys1 | 9.087336 | 9.878131 | 22.10422 | 22.21191 | 15.37916 | 15.14496 | 12.9455 | 13.0866 | 10.83427 | 10.13593 | 8.580027 | 8.08815 |
| Srgap1 | 0.979357 | 1.17665 | 1.444407 | 1.622192 | 2.296007 | 2.105834 | 2.691599 | 2.425509 | 3.409182 | 3.393917 | 4.374288 | 4.453916 |
| Map4k1 | 9.087969 | 9.698976 | 9.320974 | 9.159395 | 7.574947 | 7.864503 | 5.951249 | 7.067397 | 5.211479 | 4.797387 | 3.786671 | 3.338697 |
| Creb3 | 5.345849 | 4.201411 | 5.799223 | 5.784394 | 6.694502 | 6.811783 | 6.834811 | 7.96044 | 7.976367 | 8.901259 | 16.19966 | 15.86031 |
| Fbp2 | 1.201403 | 1.144464 | 6.22862 | 5.761251 | 4.743751 | 5.390734 | 6.272548 | 5.730165 | 2.57779 | 2.321822 | 2.034195 | 2.102397 |
| AI467606 | 3.214746 | 3.212213 | 3.194916 | 2.395912 | 1.943163 | 2.087578 | 1.463812 | 1.905961 | 1.040025 | 0.874366 | 1.071208 | 1.463978 |
| Ghr | 0.77755 | 0.811782 | 0.61467 | 0.710418 | 1.264024 | 1.098913 | 1.559089 | 1.627719 | 2.097598 | 1.9447 | 1.421175 | 1.47972 |
| Angptl2 | 0.820659 | 0.851782 | 0.972592 | 0.902895 | 1.250327 | 1.251967 | 1.537952 | 1.698751 | 2.429685 | 2.46959 | 1.491257 | 1.426398 |
| Rspo1 | 0.801577 | 0.865751 | 1.508506 | 2.050835 | 2.567743 | 2.198963 | 2.190946 | 2.555457 | 1.212229 | 1.090583 | 0.757824 | 0.934437 |
| Pik3r3 | 2.323411 | 2.029024 | 3.052609 | 3.292011 | 4.684468 | 4.955389 | 7.439728 | 7.26587 | 9.237314 | 8.994459 | 6.067291 | 6.106492 |
| Adam10 | 8.647349 | 9.455495 | 10.87947 | 11.6316 | 14.09198 | 14.18836 | 20.05707 | 18.27541 | 29.93216 | 31.75926 | 25.71836 | 27.83196 |
| Stx11 | 0.353773 | 0.333931 | 0.510619 | 0.568365 | 0.596685 | 0.728463 | 0.760821 | 0.958467 | 0.813586 | 0.869489 | 1.794668 | 1.551881 |
| Gprc5c | 2.157104 | 2.588426 | 7.745701 | 8.060053 | 12.98615 | 11.99614 | 22.12914 | 19.74629 | 20.81694 | 21.99923 | 18.30063 | 17.19437 |
| Cblc | 1.321781 | 1.460018 | 1.403008 | 1.389061 | 1.066544 | 1.063123 | 0.838787 | 0.930574 | 0.626891 | 0.509278 | 0.336132 | 0.560061 |
| Plxnd1 | 0.633694 | 0.650787 | 1.142709 | 1.254741 | 1.9011 | 1.794146 | 2.446399 | 2.402049 | 2.902566 | 3.18639 | 3.876324 | 3.420849 |
| Cpeb2 | 0.535028 | 0.667908 | 1.140846 | 1.043418 | 1.241899 | 1.297451 | 1.479868 | 1.579867 | 1.462906 | 1.411359 | 2.577525 | 2.844284 |
| Irf2bp2 | 8.294363 | 8.538235 | 8.610054 | 8.833795 | 9.430723 | 9.836118 | 10.83876 | 10.28677 | 15.25625 | 14.28157 | 25.38177 | 24.07324 |
| Satb2 | 0.735871 | 1.188635 | 1.684235 | 1.844486 | 2.21714 | 2.2691 | 3.063861 | 2.710332 | 4.261707 | 4.288786 | 4.985763 | 5.024627 |
| Slc22a18 | 1.402387 | 1.59495 | 1.137129 | 0.930352 | 0.843428 | 0.768987 | 0.471358 | 1.056925 | 0.544999 | 0.634804 | 0.494906 | 0.388002 |
| Cib2 | 0.857296 | 0.576014 | 0.876071 | 0.759279 | 1.658964 | 1.39378 | 2.354428 | 1.602573 | 2.203697 | 2.066696 | 1.539664 | 1.451673 |
| Pdk3 | 17.0267 | 16.74869 | 19.31798 | 20.26457 | 25.19863 | 22.11948 | 35.01168 | 32.59645 | 56.49605 | 55.14765 | 41.76712 | 45.3142 |
| Rassf4 | 0.419199 | 0.443889 | 0.645745 | 0.670482 | 0.841845 | 0.782394 | 1.162459 | 1.115531 | 1.239 | 1.411062 | 1.81923 | 1.821738 |
| Cercam | 3.008708 | 3.174872 | 3.172421 | 3.052344 | 4.064157 | 3.975199 | 5.57665 | 5.213209 | 7.885689 | 8.284376 | 8.613597 | 7.711938 |
| Snta1 | 9.345867 | 9.490461 | 8.187628 | 7.934549 | 7.88048 | 7.796531 | 7.581416 | 6.949789 | 4.916261 | 4.493775 | 3.381667 | 2.923799 |
| Tspan17 | 5.058647 | 5.386067 | 6.934889 | 5.849868 | 5.880187 | 5.526931 | 3.665469 | 4.169249 | 2.526017 | 2.412493 | 2.078373 | 2.156295 |
| Nlgn2 | 2.557423 | 3.551343 | 3.579569 | 3.416681 | 4.956777 | 4.613562 | 6.108478 | 6.026731 | 9.382509 | 9.263329 | 6.241183 | 5.876725 |
| Eras | 19.93327 | 17.91823 | 15.24259 | 12.22917 | 9.068797 | 8.474907 | 6.879794 | 7.049646 | 6.237792 | 7.239057 | 6.059883 | 4.510627 |
| Naglu | 1.156002 | 1.115116 | 1.546949 | 1.276035 | 2.570875 | 2.1479 | 2.858696 | 2.544737 | 3.403074 | 3.458102 | 2.81164 | 2.674828 |
| Jakmip2 | 0.390919 | 0.37448 | 0.529368 | 0.459697 | 0.728936 | 0.834813 | 0.994511 | 0.9878 | 1.062389 | 0.898801 | 1.042351 | 1.245785 |
| Pth1r | 2.642042 | 2.373962 | 4.165386 | 3.718333 | 4.765964 | 5.033577 | 4.930814 | 4.412 | 7.946651 | 7.975591 | 10.44381 | 10.07615 |
| Kndc1 | 1.909132 | 1.932904 | 0.840134 | 0.793246 | 0.709399 | 0.617643 | 0.653891 | 0.571472 | 0.373253 | 0.493995 | 0.359511 | 0.292895 |
| Extl2 | 2.487404 | 2.336055 | 2.195397 | 2.353109 | 3.050358 | 3.042843 | 3.660975 | 3.844383 | 5.781293 | 5.533015 | 5.443003 | 6.368893 |
| 9330182L0 | 0.584832 | 0.636828 | 0.981151 | 1.054659 | 1.662473 | 1.591785 | 1.851683 | 1.759515 | 2.526863 | 2.853312 | 1.531319 | 1.532498 |
| Nfkb2 | 3.203268 | 3.94776 | 5.551682 | 5.670402 | 5.713522 | 4.708396 | 5.495014 | 5.088845 | 4.592748 | 4.558288 | 14.31093 | 12.32697 |
| Hmgn3 | 2.347158 | 1.569736 | 3.104441 | 3.341522 | 6.371953 | 6.179058 | 7.383597 | 9.029748 | 9.337513 | 9.026754 | 8.923261 | 8.597627 |
| Ispd | 0.276558 | 0.275361 | 0.606153 | 0.723951 | 0.883602 | 0.905929 | 1.009498 | 1.045363 | 1.296799 | 1.459037 | 2.044669 | 1.956176 |
| Adap1 | 8.769729 | 8.824307 | 7.723002 | 6.645517 | 6.159052 | 5.892299 | 5.290644 | 4.857168 | 2.958914 | 2.460598 | 4.867381 | 4.527073 |
| Rasip1 | 0.944455 | 1.025824 | 2.128773 | 1.707142 | 1.220162 | 1.268791 | 1.261654 | 1.141672 | 0.716382 | 0.632691 | 0.987567 | 0.720999 |
| Sorbs2 | 0.342218 | 0.316845 | 0.417331 | 0.427484 | 0.505054 | 0.415458 | 0.422534 | 0.452258 | 0.751497 | 0.891621 | 0.977174 | 1.120925 |
| Dppa5a | 1426.002 | 1291 | 1092.597 | 1109.048 | 935.4223 | 964.1253 | 705.6972 | 750.4791 | 482.2662 | 483.2334 | 457.7247 | 411.0764 |
| Bend5 | 2.152488 | 1.748918 | 1.816266 | 1.778529 | 2.195283 | 2.732365 | 2.681772 | 3.439315 | 4.554631 | 4.79068 | 4.122689 | 4.35079 |
| Prph | 3.718567 | 3.525646 | 10.91318 | 10.95314 | 14.11667 | 13.79735 | 13.90252 | 17.38983 | 6.843008 | 8.146888 | 7.250498 | 6.456812 |
| Prr7 | 2.888599 | 3.088806 | 3.357013 | 3.571815 | 3.006586 | 3.380103 | 3.386393 | 3.032576 | 4.906248 | 4.837479 | 9.124371 | 8.166666 |
| Rxra | 3.597808 | 4.222089 | 5.523472 | 5.42584 | 7.429163 | 7.689363 | 10.16027 | 9.191874 | 14.13773 | 14.60414 | 15.30835 | 14.53891 |

| | | | | | | | | | | | | |
|---|---|---|---|---|---|---|---|---|---|---|---|---|
| Atp6v0a4 | 4.274658 | 3.665209 | 0.398315 | 0.25733 | 0.316429 | 0.304584 | 0.410156 | 0.439335 | 0.595875 | 0.692342 | 0.706156 | 0.53112 |
| Rec8 | 7.595254 | 7.700721 | 25.75286 | 25.42973 | 40.0742 | 39.92562 | 51.68479 | 55.25767 | 48.27999 | 46.21462 | 19.59214 | 20.54341 |
| Arntl2 | 0.972306 | 0.812369 | 3.213574 | 3.317236 | 2.445737 | 2.404684 | 2.347138 | 2.485255 | 1.180629 | 1.23429 | 2.326702 | 2.448268 |
| E230016M | 0.333989 | 0.367283 | 0.600733 | 0.819661 | 0.773673 | 0.715114 | 0.743877 | 0.682254 | 1.009747 | 0.858445 | 1.810326 | 1.833584 |
| Prickle2 | 0.556896 | 0.672971 | 0.516588 | 0.719837 | 1.11174 | 1.026635 | 1.774741 | 1.472795 | 2.011077 | 1.934038 | 1.712651 | 1.792601 |
| Mpped2 | 1.661492 | 2.038981 | 3.344588 | 3.393965 | 4.823959 | 5.152327 | 7.688948 | 7.191334 | 9.551931 | 9.115779 | 7.025301 | 7.706896 |
| Fgf3 | 0.72603 | 0.872861 | 0.445016 | 0.580646 | 0.41202 | 0.541745 | 0.450134 | 0.611576 | 0.938415 | 0.81653 | 1.797128 | 1.454919 |
| 1700019B2 | 0.323409 | 0.39565 | 1.346767 | 0.560787 | 0.862621 | 1.035245 | 1.138919 | 1.50577 | 1.021436 | 1.074823 | 1.181131 | 0.907011 |
| Cnksr2 | 0.277618 | 0.328222 | 0.497385 | 0.598356 | 0.80046 | 0.80176 | 1.048466 | 0.963806 | 1.480969 | 1.60601 | 1.273368 | 1.391026 |
| Gpr157 | 0.407647 | 0.342644 | 1.200561 | 1.09252 | 2.497616 | 2.515203 | 2.544394 | 2.410307 | 1.574608 | 1.638001 | 0.907893 | 0.937187 |
| Itgb1 | 85.39994 | 78.47484 | 76.80096 | 74.36837 | 91.12431 | 88.72838 | 107.1853 | 114.7537 | 132.4485 | 143.3666 | 183.4113 | 199.5692 |
| Med12l | 0.33643 | 0.511714 | 0.598273 | 0.668056 | 0.960976 | 0.935993 | 1.102834 | 0.926931 | 1.563204 | 1.658153 | 1.758106 | 1.79271 |
| Enpp4 | 3.352587 | 3.554586 | 7.495552 | 7.470155 | 5.57377 | 5.326811 | 3.778461 | 3.782256 | 2.771924 | 2.935585 | 2.986781 | 2.784412 |
| Crim1 | 1.33874 | 1.385311 | 2.13037 | 2.069092 | 2.398188 | 2.301333 | 3.129929 | 2.697105 | 4.063497 | 4.005768 | 5.458581 | 5.54982 |
| Prss41 | 4.78367 | 4.995317 | 4.079434 | 4.655803 | 3.847822 | 3.913223 | 2.457373 | 2.661986 | 1.723612 | 2.371202 | 1.348894 | 1.736089 |
| Tgm2 | 1.783612 | 2.082892 | 2.021779 | 1.99261 | 3.361488 | 2.994761 | 5.147264 | 5.343578 | 4.644644 | 4.22357 | 3.373408 | 3.025151 |
| Tha1 | 0.517612 | 0.559598 | 0.53643 | 0.609446 | 0.78023 | 0.790638 | 1.009163 | 0.977125 | 1.180975 | 1.213989 | 1.587501 | 1.633986 |
| BC029214 | 0.643137 | 0.604263 | 0.929745 | 0.904744 | 1.637953 | 1.624882 | 1.132261 | 1.345467 | 1.615767 | 1.288359 | 2.491738 | 2.425398 |
| Pkn3 | 6.615164 | 6.793402 | 6.765138 | 6.290557 | 6.186934 | 6.222023 | 5.042997 | 5.851614 | 4.522322 | 4.744795 | 2.522707 | 2.346864 |
| Gm13152 | 3.496183 | 3.875963 | 5.616483 | 5.202082 | 3.505213 | 3.53972 | 2.700399 | 2.740213 | 2.125733 | 1.94097 | 1.59021 | 2.26975 |
| Ube2l6 | 1.472181 | 1.577632 | 1.427787 | 1.550662 | 2.273988 | 2.238692 | 2.948583 | 3.454946 | 3.945728 | 4.155153 | 4.535806 | 3.115991 |
| Gbgt1 | 0.785705 | 1.279692 | 2.128239 | 1.756382 | 1.433179 | 1.414348 | 1.291802 | 1.263832 | 1.436056 | 1.502618 | 0.660902 | 0.655968 |
| Nop10 | 108.5841 | 79.38701 | 70.29282 | 74.67254 | 68.72789 | 81.54955 | 56.9461 | 46.23318 | 39.65186 | 33.19396 | 35.72662 | 30.48676 |
| Phldb1 | 3.304891 | 3.63012 | 5.880479 | 5.671231 | 7.295524 | 7.413391 | 8.489507 | 8.168264 | 9.74822 | 9.740615 | 16.68526 | 15.16983 |
| Slc16a3 | 22.38788 | 19.61231 | 7.318173 | 8.966836 | 11.61919 | 11.17182 | 9.505269 | 10.15183 | 10.72003 | 12.58936 | 4.921964 | 4.709903 |
| Ccrn4l | 9.052593 | 11.12073 | 12.38155 | 11.06649 | 9.426966 | 9.262327 | 9.677207 | 8.969663 | 7.474828 | 6.676137 | 18.33352 | 17.84239 |
| Akap12 | 96.64378 | 114.1843 | 81.35911 | 82.51463 | 87.79633 | 85.36231 | 111.561 | 100.8798 | 151.0919 | 152.4246 | 211.3925 | 220.4566 |
| Fahd1 | 6.671537 | 5.890784 | 6.262494 | 6.655257 | 6.042086 | 5.833871 | 4.895973 | 5.163041 | 2.901605 | 2.703656 | 2.593891 | 2.491493 |
| Aldh2 | 56.27266 | 54.11333 | 63.98702 | 60.04203 | 47.43928 | 46.39646 | 42.0065 | 35.71692 | 26.41843 | 23.21057 | 23.59583 | 22.4812 |
| Myadm | 32.0342 | 37.23374 | 41.85453 | 41.31052 | 36.57928 | 35.12316 | 34.96752 | 28.87612 | 25.95842 | 27.01891 | 67.26067 | 72.15083 |
| Sdc2 | 2.57678 | 2.75565 | 2.846833 | 3.122761 | 4.560345 | 4.402455 | 5.538997 | 5.431616 | 8.390997 | 8.334788 | 6.509791 | 7.030029 |
| Phlda2 | 36.33572 | 36.68974 | 23.50773 | 25.63464 | 18.16884 | 18.48289 | 11.55227 | 11.94894 | 9.30436 | 9.605401 | 24.86206 | 19.45097 |
| 2610034B1 | 1.769614 | 1.517358 | 2.694936 | 2.742384 | 3.653587 | 3.488541 | 4.86196 | 4.466473 | 5.501425 | 5.142395 | 6.373857 | 7.310767 |
| Ank2 | 1.078681 | 1.346465 | 1.981578 | 2.187954 | 1.753809 | 1.865421 | 1.996361 | 1.686927 | 3.808245 | 3.789869 | 4.156209 | 4.495805 |
| Mrc1 | 0.744984 | 0.644523 | 1.804869 | 1.850452 | 1.686047 | 1.460876 | 1.470056 | 1.532463 | 0.672165 | 0.750386 | 0.784345 | 0.694349 |
| Fam184b | 0.635393 | 0.421049 | 0.658252 | 0.601336 | 0.952841 | 0.832527 | 1.313018 | 1.102889 | 1.797015 | 1.601799 | 1.072492 | 1.121239 |
| Rftn1 | 0.540348 | 0.703396 | 0.817871 | 0.787973 | 1.420221 | 1.120771 | 1.704587 | 1.582011 | 1.901208 | 2.097387 | 1.943409 | 1.852245 |
| Smarca5-ps | 1.103833 | 1.129303 | 2.291817 | 2.48328 | 1.921626 | 2.078701 | 1.457893 | 1.680792 | 0.940324 | 0.933681 | 1.612918 | 1.778123 |
| Fgfr3 | 0.607814 | 0.542781 | 1.852262 | 1.837286 | 1.872925 | 1.876676 | 2.116405 | 2.108743 | 2.412697 | 2.454391 | 5.764402 | 4.885965 |
| EU599041 | 25.81025 | 24.99597 | 21.37229 | 22.36466 | 16.7091 | 17.24853 | 12.99832 | 15.2752 | 11.24465 | 12.24091 | 7.223361 | 8.410964 |
| Igf2bp2 | 34.32937 | 36.39018 | 15.77098 | 16.13243 | 19.7887 | 17.75068 | 20.85796 | 22.56283 | 43.21253 | 42.88692 | 39.3844 | 39.38882 |
| Snhg9 | 4.308338 | 3.235103 | 4.192953 | 3.480379 | 3.07869 | 5.444224 | 1.81775 | 2.048835 | 3.445785 | 2.090452 | 3.147929 | 3.085634 |
| Gm14327 | 1.31837 | 1.238945 | 3.863314 | 3.602517 | 2.636091 | 3.179997 | 3.003733 | 2.713188 | 1.568332 | 1.357939 | 1.152092 | 1.627328 |
| Yaf2 | 10.24085 | 8.664906 | 12.27911 | 11.59636 | 17.9037 | 19.04998 | 20.20752 | 20.72678 | 29.74052 | 30.03619 | 31.43016 | 30.75938 |
| Hook2 | 21.56577 | 23.43083 | 18.36924 | 17.22998 | 13.50625 | 13.15334 | 10.15274 | 11.55112 | 8.726507 | 8.962253 | 6.450692 | 6.498077 |
| Cdx1 | 0.069263 | 0.135483 | 6.184808 | 6.314112 | 5.684037 | 5.049621 | 4.453226 | 5.008083 | 2.616401 | 2.447838 | 4.017673 | 2.381693 |
| D1Pas1 | 12.4649 | 13.18657 | 32.76778 | 30.37906 | 23.30565 | 23.09005 | 23.82273 | 22.09115 | 12.39316 | 11.46206 | 19.33542 | 19.25959 |
| Gpr161 | 0.972974 | 0.997519 | 1.579848 | 1.779891 | 2.520606 | 2.568341 | 2.706102 | 2.832034 | 3.44682 | 4.194881 | 4.947453 | 4.713936 |
| Esx1 | 1.300318 | 1.708933 | 2.406066 | 2.188329 | 2.880155 | 2.822538 | 3.066846 | 2.686825 | 5.778943 | 5.241717 | 5.670571 | 5.795617 |
| H2-M5 | 20.13793 | 18.33062 | 15.63685 | 13.55167 | 8.165417 | 7.708492 | 6.899309 | 7.852477 | 5.684888 | 5.11976 | 5.473394 | 5.222991 |
| Socs5 | 2.332475 | 2.449605 | 3.255381 | 3.260462 | 3.705981 | 3.382526 | 4.239802 | 4.151024 | 4.708983 | 4.964208 | 8.493829 | 8.625407 |
| Rasgef1b | 0.431234 | 0.592461 | 0.967342 | 1.228694 | 1.198412 | 1.049608 | 1.623767 | 1.551787 | 2.023115 | 2.323706 | 2.603449 | 2.776222 |
| Tmem63a | 2.841695 | 2.555106 | 3.034691 | 3.289586 | 3.506388 | 3.462759 | 2.923202 | 2.877804 | 1.342789 | 1.409643 | 1.41895 | 1.310131 |
| Cep85l | 0.74951 | 0.580624 | 0.92433 | 0.871542 | 1.264765 | 1.063467 | 1.382551 | 1.497796 | 1.722897 | 2.303403 | 1.926229 | 2.251882 |
| B230208H1 | 0.722762 | 0.44174 | 1.707822 | 1.231695 | 1.195546 | 1.445069 | 1.550859 | 1.685867 | 0.883342 | 0.638185 | 0.912851 | 0.760501 |
| Chrna4 | 0.242623 | 0.372933 | 0.804478 | 0.968172 | 1.37604 | 1.253704 | 1.738005 | 1.622125 | 2.024448 | 1.745125 | 2.843163 | 2.556118 |
| Lefty2 | 22.44795 | 21.11838 | 7.662981 | 7.883509 | 7.605265 | 7.801969 | 5.75686 | 6.216096 | 3.020589 | 2.988927 | 3.008867 | 3.239895 |
| Hpdl | 6.75658 | 6.62371 | 4.077366 | 3.477116 | 2.923424 | 2.731648 | 2.099827 | 2.104343 | 1.986371 | 1.898701 | 1.552012 | 1.318746 |
| Tlx3 | 0.079068 | 0.152253 | 0.435182 | 0.410351 | 0.419809 | 0.400088 | 0.44718 | 0.458582 | 0.516782 | 0.478429 | 1.134414 | 1.054362 |
| Bcar3 | 16.9865 | 16.41236 | 17.77405 | 17.28011 | 20.18998 | 19.44234 | 20.32868 | 20.70508 | 11.57653 | 11.90314 | 7.658143 | 7.857554 |
| Gdpd2 | 0.338704 | 0.347444 | 0.431058 | 0.500606 | 1.015052 | 0.907013 | 1.117021 | 1.065907 | 1.431685 | 1.317795 | 0.746657 | 0.84466 |

| | | | | | | | | | | | | |
|---|---|---|---|---|---|---|---|---|---|---|---|---|
| Msi1 | 10.41302 | 13.18279 | 20.84966 | 22.32031 | 26.67151 | 25.90678 | 34.35055 | 27.71935 | 52.73787 | 49.85814 | 58.63213 | 58.72884 |
| Tmem132a | 4.573065 | 4.913521 | 5.859653 | 5.909098 | 8.676957 | 8.011015 | 10.62915 | 12.22515 | 16.78303 | 15.51498 | 13.29432 | 13.71583 |
| Dhrs3 | 1.155561 | 0.958147 | 18.78159 | 18.69528 | 30.25082 | 32.64286 | 46.88129 | 49.05302 | 38.35497 | 35.4931 | 23.2567 | 23.37862 |
| Chek2 | 23.4731 | 22.75113 | 21.3921 | 19.68478 | 17.01211 | 16.46696 | 16.27987 | 16.01866 | 15.22993 | 13.58388 | 7.441216 | 7.508833 |
| Timp2 | 1.740515 | 1.550754 | 3.37616 | 4.06764 | 4.977798 | 4.499923 | 6.108783 | 5.496679 | 8.631184 | 8.119271 | 11.19138 | 10.65428 |
| Ccdc141 | 1.387239 | 1.520957 | 0.644966 | 0.558289 | 0.451159 | 0.464087 | 0.423663 | 0.522545 | 0.341516 | 0.391413 | 0.172487 | 0.21321 |
| Sycp3 | 4.225495 | 3.73094 | 4.044055 | 3.823421 | 3.581294 | 3.500441 | 2.591409 | 2.445588 | 1.604391 | 1.522739 | 1.657534 | 1.461744 |
| 5930412G1 | 0.597328 | 0.393651 | 0.231659 | 0.426826 | 0.555407 | 0.492457 | 0.481166 | 0.501665 | 0.898648 | 1.116387 | 0.473212 | 0.446285 |
| Rex2 | 13.1395 | 13.24354 | 12.30926 | 11.28749 | 7.991479 | 7.330473 | 5.743729 | 6.432047 | 4.195169 | 4.316554 | 3.812694 | 4.446495 |
| Nin | 1.756442 | 1.998193 | 3.117675 | 3.129117 | 3.61903 | 3.74688 | 4.564623 | 4.675058 | 6.492634 | 6.451722 | 7.81786 | 8.177728 |
| Col5a1 | 0.844567 | 1.121223 | 1.757173 | 1.720508 | 2.37109 | 2.116336 | 3.007088 | 2.503369 | 3.408411 | 3.291918 | 4.683729 | 4.495976 |
| Igfbp2 | 392.4413 | 350.0666 | 144.8299 | 133.9792 | 170.353 | 163.9416 | 190.7911 | 220.3506 | 315.9715 | 350.0428 | 310.6019 | 274.449 |
| Dram1 | 0.963067 | 0.714237 | 0.449519 | 0.541745 | 0.911249 | 0.794319 | 1.281457 | 1.235541 | 1.455914 | 1.415226 | 1.352868 | 1.211365 |
| St6galnac6 | 10.89164 | 10.27383 | 9.388242 | 9.122712 | 8.960446 | 8.981055 | 7.149369 | 8.565634 | 6.876548 | 7.151705 | 3.268644 | 3.495565 |
| Klk10 | 0.81759 | 1.267868 | 0.990147 | 0.840819 | 0.579969 | 0.559883 | 0.659249 | 1.176869 | 0.856815 | 0.641268 | 2.285686 | 1.460242 |
| Il20rb | 0.389586 | 0.414535 | 0.60364 | 0.499313 | 0.662489 | 0.788354 | 1.052168 | 0.654428 | 1.074204 | 1.025517 | 1.176545 | 1.301701 |
| Ehhadh | 1.08927 | 1.058159 | 2.083404 | 2.032471 | 2.12928 | 2.150124 | 2.330773 | 2.624433 | 1.601577 | 1.743197 | 1.015839 | 1.006889 |
| Ntng2 | 0.288802 | 0.441539 | 0.43892 | 0.782775 | 0.895738 | 0.838294 | 0.85952 | 0.933989 | 1.388026 | 1.449468 | 2.21149 | 2.040038 |
| Syne2 | 2.257234 | 2.840918 | 2.180249 | 2.52187 | 3.155337 | 3.047178 | 4.049409 | 3.429089 | 5.62433 | 6.572048 | 4.509002 | 4.504016 |
| Tpm2 | 3.800305 | 3.961228 | 5.429019 | 5.78269 | 6.996361 | 6.71226 | 7.255223 | 7.908148 | 8.054863 | 6.730637 | 16.93718 | 15.05945 |
| Abcc2 | 0.377163 | 0.393836 | 1.066118 | 1.052549 | 0.995446 | 1.000697 | 0.82794 | 0.940028 | 0.416189 | 0.404241 | 0.463591 | 0.445795 |
| Tnni3 | 3.644248 | 3.349754 | 2.353445 | 2.095698 | 2.050628 | 2.175008 | 1.062068 | 2.36391 | 1.639937 | 1.223736 | 0.559491 | 0.907985 |
| Scd1 | 23.44756 | 26.96518 | 25.24931 | 26.18224 | 45.73237 | 41.37292 | 56.53876 | 51.89687 | 64.22709 | 68.13067 | 42.00737 | 40.5002 |
| Rassf1 | 2.162824 | 1.904244 | 3.312475 | 2.480445 | 2.9486 | 3.115982 | 3.096008 | 3.433631 | 3.07548 | 3.115745 | 6.563538 | 6.453576 |
| Gsta4 | 291.3702 | 278.7198 | 240.0412 | 238.035 | 210.4652 | 217.176 | 185.571 | 203.0417 | 135.4135 | 123.0782 | 82.56782 | 91.66732 |
| Plagl2 | 4.160655 | 5.115373 | 6.069861 | 6.40678 | 7.356977 | 6.853386 | 8.668928 | 7.680785 | 16.93837 | 16.63627 | 16.48193 | 16.57726 |
| Arhgdib | 3.5163 | 3.580751 | 7.956281 | 7.855924 | 6.295157 | 5.281173 | 4.599533 | 4.562909 | 3.171306 | 3.026519 | 3.393142 | 3.198748 |
| Arhgef19 | 1.364296 | 1.39861 | 1.166858 | 1.279974 | 1.238381 | 1.115179 | 0.911261 | 1.015865 | 0.599408 | 0.554502 | 0.552254 | 0.49318 |
| Olfr147 | 0.484722 | 0.383376 | 0.91948 | 1.130198 | 0.949019 | 1.251318 | 1.190169 | 1.085259 | 1.159673 | 1.064227 | 0.725479 | 0.482146 |
| Limd1 | 5.158726 | 5.478866 | 6.368799 | 6.356289 | 7.274985 | 7.389483 | 8.700221 | 8.215514 | 15.40511 | 16.16898 | 16.99612 | 16.49334 |
| Tubb3 | 23.67958 | 22.55303 | 12.55122 | 12.31667 | 12.77497 | 11.52518 | 11.96541 | 12.34401 | 10.77468 | 9.565271 | 25.96483 | 24.81655 |
| 2610018G0 | 4.987443 | 4.408934 | 4.260526 | 3.909878 | 4.062279 | 4.176299 | 3.222514 | 4.031162 | 2.297433 | 2.622835 | 1.458473 | 1.610051 |
| Lrp4 | 1.841972 | 2.016915 | 2.750753 | 2.738113 | 3.557271 | 3.327228 | 4.649533 | 4.440725 | 6.082993 | 6.358778 | 7.028785 | 7.096812 |
| Slc25a31 | 2.428483 | 2.004057 | 6.163447 | 6.111618 | 6.110433 | 7.000572 | 6.574479 | 8.219767 | 6.773485 | 6.926342 | 17.01934 | 15.83301 |
| Snai3 | 0.912922 | 0.881326 | 1.924863 | 1.356593 | 1.47036 | 1.457335 | 1.481642 | 1.622209 | 0.964148 | 0.716074 | 1.964965 | 1.853894 |
| Slc7a2 | 0.431349 | 0.510937 | 0.70464 | 0.702677 | 1.046257 | 0.959573 | 1.263285 | 1.078358 | 1.681157 | 1.81885 | 1.798878 | 1.789559 |
| B4galt2 | 4.396603 | 4.409143 | 3.705147 | 3.642816 | 4.574839 | 4.402784 | 5.495632 | 5.461911 | 9.333693 | 9.105357 | 10.05404 | 9.428106 |
| Vpreb3 | 2.370859 | 2.099116 | 2.803891 | 1.933015 | 2.071923 | 1.649445 | 2.109313 | 1.864397 | 1.371028 | 0.77495 | 1.014857 | 0.747033 |
| Mast1 | 1.12258 | 1.566378 | 1.574518 | 1.586914 | 1.264233 | 1.325245 | 1.053614 | 1.106263 | 0.558574 | 0.665627 | 0.510472 | 0.613472 |
| Zfp536 | 0.109434 | 0.137681 | 0.616689 | 0.532277 | 0.475114 | 0.421644 | 0.568773 | 0.486825 | 0.745287 | 0.726246 | 1.135966 | 1.089853 |
| Hist1h4n | 4.409947 | 4.413012 | 1.938115 | 1.548295 | 2.301741 | 4.001744 | 3.123501 | 2.615838 | 2.906882 | 2.729283 | 2.583081 | 2.334557 |
| Col2a1 | 1.59924 | 1.886635 | 3.74545 | 3.794705 | 4.820886 | 4.531597 | 5.271781 | 4.986056 | 10.52037 | 9.807045 | 9.159266 | 9.687 |
| Npr2 | 0.672531 | 0.662122 | 1.054062 | 1.057205 | 1.859107 | 1.785583 | 2.235946 | 2.253718 | 2.479393 | 2.655999 | 2.888783 | 2.731393 |
| Dmc1 | 10.65768 | 9.944819 | 8.074552 | 7.561629 | 7.41422 | 7.196227 | 5.932991 | 5.901148 | 4.103446 | 3.889488 | 2.816869 | 2.928302 |
| 2410141K0 | 30.17776 | 27.56283 | 27.21811 | 27.36681 | 20.78778 | 21.38102 | 16.16389 | 17.6155 | 10.28207 | 10.60406 | 10.76484 | 11.26624 |
| Ephx2 | 8.005871 | 8.163669 | 4.168612 | 4.025897 | 3.010193 | 2.825806 | 2.865241 | 3.316394 | 1.638205 | 1.739727 | 1.323711 | 1.560958 |
| Fam19a5 | 3.530185 | 2.728917 | 3.752973 | 3.70068 | 6.813506 | 6.690058 | 8.388142 | 9.064004 | 9.814977 | 8.90369 | 9.76583 | 9.54423 |
| Ly6k | 0.281521 | 0.270053 | 0.873048 | 0.713096 | 1.427087 | 1.55726 | 1.330351 | 1.125758 | 0.903167 | 0.603994 | 1.145767 | 0.767798 |
| Tjp3 | 4.344111 | 4.727558 | 2.694595 | 2.244061 | 2.279232 | 1.898138 | 1.582476 | 1.472346 | 1.318629 | 1.503755 | 0.894377 | 0.870414 |
| Zfp697 | 0.464958 | 0.499147 | 1.009778 | 1.060258 | 1.600415 | 1.536098 | 2.080344 | 1.488709 | 2.281643 | 2.490544 | 2.734006 | 2.733342 |
| Slc30a3 | 0.520744 | 0.702339 | 1.077458 | 1.109824 | 1.204583 | 1.288094 | 1.042517 | 1.099324 | 0.503544 | 0.599715 | 0.534105 | 0.499774 |
| Spag1 | 3.957697 | 3.612114 | 4.202483 | 3.934761 | 3.956701 | 3.857542 | 3.443243 | 3.584333 | 2.64746 | 3.048879 | 1.71334 | 1.527031 |
| Mfsd7b | 2.396003 | 2.729945 | 2.903284 | 3.175105 | 3.288731 | 3.065594 | 3.419559 | 3.531946 | 4.751332 | 4.749233 | 7.957641 | 7.892798 |
| Fam3b | 0.502936 | 0.551908 | 1.341848 | 1.325884 | 1.144036 | 1.341969 | 1.541346 | 2.077064 | 0.891159 | 1.182639 | 1.019334 | 0.806895 |
| S100a10 | 75.18968 | 58.4747 | 27.2979 | 24.9989 | 23.40928 | 24.69887 | 26.92636 | 26.88871 | 32.26464 | 31.439 | 62.40544 | 63.56299 |
| Grb14 | 0.591296 | 0.747735 | 1.300047 | 1.333411 | 1.30405 | 1.347478 | 1.649789 | 1.608595 | 1.039835 | 1.094225 | 0.803218 | 0.625294 |
| Neil2 | 5.96503 | 5.635867 | 2.68304 | 2.66588 | 2.178679 | 2.49278 | 2.136484 | 2.474349 | 1.79192 | 1.850948 | 0.683317 | 1.036439 |
| Gm14393 | 7.926043 | 7.935923 | 7.673186 | 7.412676 | 4.903349 | 5.538313 | 4.14622 | 4.79919 | 3.061009 | 3.615607 | 2.943539 | 2.882497 |
| Chrna7 | 2.407651 | 2.647234 | 3.040676 | 3.044756 | 3.006061 | 2.825143 | 3.24169 | 3.451713 | 5.770928 | 5.429185 | 6.156175 | 7.263731 |
| Nsmf | 3.04358 | 2.8702 | 4.263592 | 4.557779 | 7.37752 | 7.329227 | 9.012262 | 9.953511 | 11.34288 | 11.71798 | 11.19658 | 11.67776 |
| Fuom | 4.728727 | 3.961005 | 2.841062 | 2.817363 | 2.905096 | 3.192933 | 2.634707 | 3.087387 | 1.679174 | 1.591733 | 1.311694 | 1.241955 |

| | | | | | | | | | | | | |
|---|---|---|---|---|---|---|---|---|---|---|---|---|
| Tspan13 | 21.53593 | 20.10339 | 22.71656 | 22.05017 | 28.19782 | 27.54434 | 36.6851 | 35.95084 | 58.14154 | 56.64177 | 50.9272 | 51.19449 |
| Il1rap | 0.347938 | 0.267969 | 0.486646 | 0.561034 | 0.855459 | 0.741222 | 0.991413 | 1.036883 | 1.258719 | 1.440185 | 0.976167 | 1.215114 |
| Ppp1r14a | 30.58482 | 25.0225 | 30.60662 | 27.4576 | 30.01618 | 29.91198 | 22.82408 | 24.44856 | 14.19688 | 12.7 | 14.04803 | 11.65372 |
| Shank1 | 0.474968 | 0.527734 | 1.781664 | 1.876366 | 1.562742 | 1.78579 | 1.667147 | 1.651595 | 0.827164 | 0.798894 | 0.77429 | 0.731542 |
| Arhgap33 | 0.877083 | 0.753459 | 1.392903 | 1.222237 | 1.486115 | 1.380692 | 1.59546 | 1.576837 | 2.399793 | 2.568103 | 3.492716 | 3.134959 |
| Synj2 | 2.252032 | 2.721321 | 3.238173 | 3.212635 | 2.812696 | 2.937743 | 2.768433 | 2.735961 | 1.95971 | 1.976511 | 4.606831 | 5.068808 |
| Gm561 | 11.46788 | 7.331592 | 10.28153 | 9.82803 | 9.627035 | 10.46927 | 7.656083 | 6.666868 | 4.1859 | 4.082573 | 5.174244 | 5.237413 |
| Slc39a14 | 23.71877 | 25.68018 | 21.37475 | 21.87522 | 26.30548 | 27.61052 | 42.37345 | 36.8104 | 62.36197 | 56.09601 | 43.95551 | 50.73192 |
| Cadm4 | 2.324004 | 2.46477 | 1.868079 | 1.985302 | 3.122653 | 2.872473 | 3.867969 | 4.4026 | 4.839963 | 4.63693 | 5.46373 | 5.089592 |
| Hist1h4h | 4.774291 | 4.94129 | 1.834009 | 4.480921 | 3.199725 | 4.93464 | 3.048836 | 3.071627 | 2.459048 | 2.313563 | 1.453637 | 1.926168 |
| Ltbr | 2.400189 | 2.0514 | 3.52946 | 3.614885 | 4.493345 | 3.906461 | 5.013976 | 4.626651 | 6.434001 | 6.094488 | 9.908287 | 9.260778 |
| Mpdz | 4.472865 | 4.984922 | 5.965355 | 5.922692 | 8.077118 | 8.131452 | 9.928437 | 9.151488 | 14.52778 | 15.1717 | 13.27679 | 14.19131 |
| Hoxb7 | 0.280374 | 0.197554 | 6.433931 | 5.80231 | 10.12108 | 9.358315 | 12.7183 | 14.8479 | 13.71093 | 11.99195 | 12.14634 | 13.80876 |
| Smim3 | 2.356461 | 2.166068 | 3.09187 | 2.785254 | 3.259812 | 3.014199 | 3.358915 | 3.023683 | 4.553006 | 4.15644 | 7.517308 | 7.126002 |
| Mapkapk2 | 10.89696 | 11.62934 | 19.51445 | 17.92646 | 24.57725 | 23.7449 | 30.96992 | 29.54236 | 34.47809 | 32.64964 | 46.88926 | 45.86135 |
| Ak4 | 38.13023 | 35.8583 | 17.21132 | 16.17045 | 22.08674 | 19.97153 | 24.92106 | 26.05638 | 37.31981 | 41.35973 | 38.10886 | 35.11005 |
| Dusp14 | 1.912417 | 2.208914 | 3.995595 | 4.385207 | 4.821277 | 5.382934 | 4.661002 | 5.459375 | 3.580386 | 3.184919 | 8.666776 | 8.146091 |
| Apoe | 300.3455 | 285.7329 | 191.0596 | 157.8847 | 134.3803 | 138.918 | 110.496 | 114.4964 | 77.38658 | 66.87445 | 67.52772 | 61.7512 |
| Mocos | 1.02035 | 1.289631 | 0.943033 | 0.924373 | 0.844802 | 0.809322 | 0.704373 | 0.89379 | 0.25229 | 0.361546 | 0.477217 | 0.470663 |
| Col11a1 | 0.671137 | 0.723658 | 0.995244 | 1.026499 | 1.410795 | 1.309619 | 1.837224 | 1.717706 | 2.247954 | 2.62344 | 1.592309 | 1.554299 |
| Rnf144a | 3.791117 | 3.487529 | 3.608536 | 3.693534 | 4.856961 | 5.151214 | 6.150697 | 5.873137 | 10.61589 | 9.311119 | 9.402018 | 9.434038 |
| Zfp358 | 4.285514 | 4.047689 | 5.193717 | 5.00391 | 6.558606 | 6.547697 | 7.626393 | 8.627858 | 13.40559 | 12.7769 | 12.44034 | 12.03278 |
| Hist1h3e | 13.71869 | 14.25414 | 2.439942 | 6.260384 | 6.792287 | 10.36588 | 6.243474 | 9.698572 | 5.295304 | 7.363028 | 4.231281 | 4.061189 |
| Fam198b | 2.342039 | 2.618503 | 3.143464 | 3.304968 | 3.529421 | 3.490439 | 4.892009 | 4.172 | 8.689539 | 8.43496 | 7.404314 | 8.399613 |
| Lin28b | 9.080831 | 9.610719 | 9.53346 | 9.489221 | 9.972502 | 9.88061 | 11.53823 | 10.75809 | 24.62291 | 24.2009 | 18.50814 | 19.87197 |
| Zmat1 | 0.414586 | 0.425257 | 0.457217 | 0.641409 | 0.800176 | 0.767694 | 0.910895 | 1.365451 | 1.191825 | 1.32261 | 1.305632 | 1.634878 |
| Ddb2 | 4.63278 | 4.328858 | 4.207178 | 4.632863 | 3.630501 | 3.976379 | 3.252377 | 3.591035 | 3.513046 | 3.344091 | 1.880104 | 1.817849 |
| Mafa | 1.694544 | 2.191826 | 5.910797 | 5.110154 | 4.250753 | 4.369081 | 2.949665 | 4.035777 | 1.986487 | 2.007985 | 3.047695 | 2.451461 |
| Dusp3 | 3.751231 | 4.214812 | 6.326205 | 6.085873 | 5.343963 | 4.92263 | 4.675411 | 4.154248 | 4.654496 | 4.260109 | 10.01613 | 10.56949 |
| Dpysl2 | 2.601322 | 3.881694 | 3.64984 | 4.354738 | 6.014052 | 5.708373 | 7.707298 | 6.573835 | 10.7405 | 10.72095 | 9.953219 | 11.07399 |
| Sntb1 | 2.480841 | 2.380299 | 5.274944 | 4.907827 | 4.619478 | 4.121218 | 3.602289 | 3.442753 | 2.056346 | 1.9686 | 2.015853 | 1.930745 |
| Pde4a | 1.490404 | 1.917734 | 3.544144 | 3.744943 | 3.458161 | 2.958585 | 2.90741 | 2.709488 | 2.253505 | 2.011635 | 1.635862 | 1.473395 |
| Dppa4 | 105.1682 | 92.78333 | 87.19554 | 85.64491 | 75.16702 | 74.45566 | 67.53941 | 70.11732 | 39.66771 | 37.03027 | 34.25221 | 33.70952 |
| Cdyl2 | 12.98902 | 15.60859 | 11.61179 | 11.80375 | 8.241533 | 7.854161 | 6.27878 | 5.157872 | 4.346003 | 4.647356 | 4.963083 | 4.789041 |
| Mef2a | 3.591083 | 3.647817 | 4.480714 | 4.263971 | 4.769524 | 4.791186 | 5.064752 | 4.696864 | 7.002047 | 6.860811 | 10.31633 | 10.82953 |
| Tspan3 | 42.55543 | 37.47874 | 38.09933 | 37.52405 | 48.25937 | 46.40594 | 57.80718 | 62.84204 | 91.08933 | 88.44167 | 94.43149 | 95.28635 |
| 4833417C1 | 1.222655 | 1.803723 | 0.893552 | 1.210254 | 1.142554 | 1.093054 | 0.861125 | 0.849443 | 0.5081 | 0.724799 | 0.246501 | 0.476627 |
| Igsf1 | 1.021282 | 1.034017 | 0.43053 | 0.615742 | 0.705645 | 0.68093 | 0.698948 | 0.763273 | 0.528943 | 0.570788 | 0.312887 | 0.30084 |
| She | 0.714047 | 0.763413 | 0.692634 | 0.815351 | 1.275398 | 1.129074 | 1.514295 | 1.741268 | 2.104517 | 2.063459 | 2.170256 | 1.944733 |
| Zfp783 | 0.547203 | 0.531828 | 1.276739 | 1.483656 | 1.271239 | 1.224227 | 1.419184 | 1.452996 | 1.368887 | 1.348416 | 2.810759 | 3.097084 |
| Kctd10 | 6.111116 | 5.611817 | 7.790359 | 7.467921 | 8.601641 | 8.868678 | 10.09684 | 9.735093 | 10.92303 | 10.97882 | 20.85826 | 18.88539 |
| Hist1h2ah | 3.874224 | 2.770128 | 1.310449 | 2.06384 | 2.738739 | 3.454808 | 3.309444 | 4.397406 | 2.829448 | 2.931904 | 2.406982 | 1.74096 |
| Nab1 | 2.763623 | 2.292535 | 4.087364 | 3.566451 | 4.14398 | 3.968928 | 4.354784 | 4.185858 | 4.553837 | 4.271014 | 8.712657 | 9.005667 |
| Tmem62 | 0.462454 | 0.331911 | 0.993844 | 0.848582 | 0.930575 | 1.015482 | 0.933948 | 0.834334 | 1.213626 | 1.043039 | 1.894641 | 2.10457 |
| Nudt11 | 1.712261 | 1.805205 | 3.642954 | 3.502403 | 4.744556 | 4.789589 | 5.419706 | 5.881542 | 7.263253 | 8.026958 | 7.946277 | 8.833331 |
| Il17b | 2.038439 | 2.562577 | 1.42916 | 1.707967 | 0.902805 | 0.877876 | 0.953375 | 0.67727 | 0.654231 | 1.071384 | 0.88282 | 0.841179 |
| Plk2 | 12.35061 | 12.53377 | 17.71783 | 16.50219 | 18.37519 | 18.46286 | 21.83143 | 19.98793 | 17.79876 | 16.25337 | 39.24688 | 40.98101 |
| Tlr2 | 3.304695 | 3.356053 | 3.118274 | 3.142255 | 3.615496 | 3.629173 | 4.650109 | 4.713152 | 7.158681 | 7.546777 | 7.782717 | 7.92106 |
| Gca | 7.205773 | 7.571944 | 16.97338 | 17.01216 | 13.97822 | 12.72898 | 10.54212 | 11.04825 | 8.263964 | 8.47012 | 6.952681 | 6.751862 |
| Fam189a2 | 0.887392 | 0.902461 | 1.216769 | 1.513029 | 2.0792 | 2.25899 | 3.254126 | 3.091351 | 4.132301 | 3.522416 | 3.386415 | 3.809755 |
| Onecut2 | 0.313257 | 0.472669 | 0.386463 | 0.44631 | 0.45854 | 0.454231 | 0.689338 | 0.479486 | 0.592266 | 0.66182 | 1.075223 | 1.12357 |
| Wdr96 | 1.176084 | 1.307537 | 1.225252 | 1.333058 | 1.790303 | 1.543189 | 1.963593 | 1.789568 | 2.370883 | 2.467261 | 2.826214 | 3.355891 |
| Iqcg | 5.038898 | 5.346757 | 4.411975 | 4.400863 | 2.423074 | 2.633396 | 2.190132 | 2.298352 | 1.895322 | 1.883023 | 1.713471 | 1.748671 |
| Col5a2 | 0.611738 | 0.590861 | 0.831663 | 0.725871 | 0.94872 | 0.811207 | 1.438733 | 1.379484 | 1.776139 | 1.820683 | 1.682401 | 1.826424 |
| Loxl1 | 1.009499 | 0.7475 | 1.446916 | 1.334042 | 1.622642 | 1.465162 | 1.370927 | 1.263642 | 0.915778 | 0.99649 | 2.84849 | 2.506383 |
| Lgals9 | 3.315381 | 2.83533 | 6.652309 | 6.835549 | 3.516979 | 3.167572 | 2.572683 | 2.718646 | 2.454293 | 2.809686 | 3.916852 | 3.994749 |
| Zfp629 | 0.774055 | 0.891988 | 1.299759 | 1.432304 | 1.704231 | 1.698441 | 2.039677 | 1.945137 | 3.341434 | 3.477028 | 3.960334 | 3.600011 |
| Kcnq4 | 0.427046 | 0.339837 | 0.524293 | 0.535818 | 0.837669 | 0.713367 | 1.022277 | 0.944756 | 1.394987 | 1.236177 | 1.512819 | 1.346617 |
| Efcab4a | 1.519862 | 1.366434 | 1.336846 | 1.559951 | 1.256628 | 1.553155 | 1.028005 | 0.966047 | 0.714216 | 0.714468 | 0.730741 | 0.620721 |
| Zbtb42 | 0.457614 | 0.504968 | 1.354363 | 1.156922 | 1.755346 | 1.512811 | 1.857724 | 1.832641 | 2.594064 | 2.406266 | 3.332806 | 2.905392 |
| Peli2 | 1.648404 | 1.911885 | 1.830427 | 1.77945 | 2.538801 | 2.451925 | 3.458595 | 2.932699 | 4.532412 | 4.465074 | 3.243174 | 3.417079 |

| | | | | | | | | | | | | |
|---|---|---|---|---|---|---|---|---|---|---|---|---|
| Srcrb4d | 1.916061 | 1.653435 | 1.41397 | 1.276796 | 1.05728 | 1.071612 | 1.015939 | 0.853681 | 0.82537 | 0.509118 | 0.733813 | 0.906821 |
| Adamts20 | 0.266621 | 0.279787 | 0.384749 | 0.463606 | 0.616827 | 0.514038 | 0.598109 | 0.612553 | 1.132066 | 1.162372 | 0.828993 | 0.904125 |
| Rsad1 | 0.551646 | 0.599248 | 0.853488 | 0.819217 | 1.169048 | 1.108167 | 1.038335 | 1.137388 | 2.181249 | 1.761518 | 1.93886 | 2.051922 |
| Zfp422 | 8.74614 | 8.021615 | 11.48513 | 10.51103 | 12.61592 | 12.20114 | 13.072 | 13.70572 | 20.18011 | 20.89738 | 25.41842 | 26.32305 |
| H2-T22 | 1.315759 | 0.918165 | 0.954345 | 1.104815 | 1.841033 | 1.423629 | 2.049038 | 1.913255 | 2.671888 | 2.766248 | 2.354604 | 2.046367 |
| Cd59b | 1.365229 | 0.878506 | 0.765818 | 1.08386 | 0.910996 | 1.135645 | 1.148547 | 1.059809 | 0.759255 | 0.555367 | 0.391527 | 0.453947 |
| Pik3r1 | 2.550361 | 3.487624 | 3.761249 | 3.887518 | 4.694625 | 4.59611 | 5.684603 | 5.508349 | 8.693474 | 8.517882 | 9.424664 | 9.724909 |
| Cltb | 7.744914 | 7.271516 | 7.347205 | 6.606662 | 5.583144 | 5.441592 | 4.379783 | 4.651271 | 3.337301 | 2.641812 | 5.861453 | 5.162582 |
| Orai1 | 5.996233 | 6.189896 | 7.119597 | 6.02017 | 7.803666 | 7.11778 | 9.618388 | 9.152749 | 9.906395 | 9.79143 | 16.51466 | 15.05193 |
| Spg20 | 7.859158 | 7.140869 | 9.022052 | 8.739556 | 13.28815 | 12.60477 | 16.20398 | 15.22209 | 20.10957 | 21.83116 | 18.47842 | 18.51093 |
| Slc38a5 | 0.335906 | 0.214047 | 0.454664 | 0.50064 | 0.793032 | 0.76063 | 1.425017 | 0.992646 | 1.349953 | 1.250531 | 0.948628 | 0.8515 |
| Pdgfra | 2.086735 | 2.129429 | 2.341662 | 2.697572 | 2.88422 | 3.229057 | 4.550369 | 4.936381 | 7.310256 | 6.736512 | 5.416671 | 6.275739 |
| Sult2b1 | 0.470057 | 0.609359 | 0.794564 | 0.49431 | 0.989072 | 1.000272 | 0.911303 | 1.056585 | 1.127695 | 1.233695 | 0.748103 | 0.874948 |
| Msln | 0.458434 | 0.481269 | 1.232074 | 1.080526 | 0.504672 | 0.641519 | 0.65382 | 0.753775 | 0.666885 | 0.433216 | 0.996066 | 0.924756 |
| Rnd1 | 1.335133 | 1.521311 | 1.734751 | 1.71558 | 1.818684 | 1.693405 | 1.897429 | 1.808308 | 1.828022 | 2.281958 | 5.111606 | 4.223679 |
| Dnmt3l | 145.7487 | 136.0531 | 63.75456 | 60.41097 | 42.2335 | 43.99058 | 38.53204 | 41.28522 | 25.11841 | 24.22493 | 28.24949 | 27.07056 |
| Hs3st6 | 0.940747 | 0.589896 | 0.965189 | 1.032104 | 1.587497 | 1.359602 | 1.625283 | 1.576334 | 1.921626 | 1.909189 | 0.590402 | 0.765776 |
| Erap1 | 1.411845 | 1.195526 | 1.069971 | 1.388116 | 1.915497 | 1.662008 | 2.174952 | 2.393561 | 2.158308 | 2.362005 | 3.358636 | 3.46076 |
| Mdk | 162.6907 | 145.1281 | 392.027 | 398.2766 | 728.4435 | 754.1993 | 975.4216 | 933.9179 | 1156.268 | 988.2547 | 1069.681 | 992.1947 |
| Mrpl14 | 24.62655 | 20.23884 | 22.56011 | 19.94766 | 15.92525 | 16.63275 | 13.76465 | 11.26263 | 11.094 | 10.28679 | 12.72475 | 8.007278 |
| Arl2bp | 27.8446 | 23.37903 | 29.9511 | 29.18776 | 35.89368 | 36.36649 | 38.10901 | 41.51437 | 54.88207 | 48.49854 | 76.22195 | 72.71111 |
| Hmox1 | 35.81115 | 36.34767 | 68.65701 | 62.04376 | 51.14083 | 51.80755 | 50.37233 | 44.64922 | 42.29823 | 34.08635 | 86.62868 | 84.89754 |
| Mxd4 | 2.472181 | 2.334267 | 5.070964 | 4.957454 | 8.0166 | 8.247321 | 12.52095 | 12.34619 | 12.16682 | 11.92783 | 12.50872 | 12.22322 |
| Fzd6 | 3.275439 | 3.115713 | 2.858977 | 2.992193 | 3.266381 | 3.377769 | 3.713735 | 3.766823 | 5.93233 | 6.308538 | 6.727742 | 7.451638 |
| Hoxa4 | 0.354047 | 0.409816 | 5.340619 | 5.683026 | 8.758672 | 9.739883 | 11.29046 | 9.114679 | 4.278628 | 3.914802 | 4.870154 | 4.667473 |
| Spin4 | 0.951601 | 1.120068 | 1.217668 | 1.26036 | 1.16285 | 1.174618 | 1.179139 | 1.3028 | 1.793267 | 1.786047 | 2.323054 | 2.919969 |
| Fam219a | 0.468311 | 0.290727 | 0.935234 | 0.8483 | 0.906021 | 1.00862 | 1.082318 | 0.893504 | 1.007553 | 0.935325 | 2.007342 | 2.108612 |
| Tesk1 | 1.017641 | 1.01823 | 1.540657 | 1.453282 | 1.63155 | 1.926394 | 2.259956 | 2.196709 | 2.727959 | 2.554225 | 4.003372 | 3.612316 |
| Reep3 | 4.497921 | 4.847546 | 4.895724 | 4.947872 | 5.794962 | 5.169785 | 6.365815 | 5.497656 | 9.917138 | 10.41861 | 11.69698 | 12.29682 |
| Serf1 | 2.82308 | 1.586374 | 1.180977 | 1.387512 | 1.44001 | 2.223336 | 2.412342 | 2.885328 | 4.312713 | 3.44697 | 4.202539 | 3.264946 |
| 4930502E1 | 0.834212 | 0.728129 | 2.451085 | 2.304285 | 2.683123 | 2.416896 | 2.467898 | 2.594604 | 1.431236 | 1.39471 | 1.044503 | 1.044509 |
| Pdlim5 | 12.30851 | 13.21238 | 12.22128 | 12.97798 | 13.33371 | 13.51569 | 14.8015 | 14.49959 | 14.21986 | 14.73794 | 30.52495 | 32.23682 |
| Slc30a4 | 1.650868 | 1.66061 | 2.536011 | 2.568377 | 3.114583 | 2.97216 | 3.245839 | 3.177139 | 4.67428 | 4.967543 | 6.159204 | 6.379731 |
| Tmem119 | 0.611315 | 0.484392 | 0.98395 | 0.774385 | 1.347939 | 1.24747 | 1.728453 | 1.923181 | 1.13047 | 1.259926 | 0.980982 | 0.82717 |
| 4931428F0 | 1.964746 | 2.017836 | 2.135634 | 2.346573 | 2.267231 | 2.622858 | 2.552668 | 2.902804 | 3.216335 | 2.905295 | 6.239552 | 5.824124 |
| Pck2 | 4.674093 | 5.134397 | 12.26389 | 11.97629 | 11.25711 | 9.96598 | 7.244448 | 7.181692 | 5.259017 | 4.825695 | 8.757104 | 8.362587 |
| Shroom3 | 6.074868 | 7.634649 | 8.438789 | 8.153619 | 8.55488 | 8.182929 | 9.842754 | 8.590568 | 17.53412 | 17.22113 | 20.84277 | 20.22659 |
| Acbd4 | 5.273965 | 5.447874 | 4.499238 | 4.438492 | 4.788672 | 4.958606 | 4.700088 | 5.962448 | 4.045599 | 3.855151 | 2.505196 | 2.40446 |
| 1810062O1 | 0.483456 | 0.755816 | 1.007537 | 1.005544 | 0.670949 | 1.08285 | 0.664826 | 0.436844 | 0.913705 | 0.966085 | 0.763462 | 0.517834 |
| Kat2b | 19.09503 | 20.26861 | 23.20117 | 21.99246 | 17.93051 | 16.94882 | 15.42428 | 15.44106 | 10.18995 | 11.78819 | 8.55196 | 8.875533 |
| Irf8 | 0.307737 | 0.334287 | 0.549295 | 0.667602 | 0.789583 | 0.752557 | 0.696973 | 0.810616 | 1.280326 | 1.29772 | 1.522913 | 1.653694 |
| Stxbp6 | 0.75687 | 0.673753 | 1.161178 | 0.98515 | 1.512676 | 1.492362 | 1.658808 | 1.447546 | 2.549738 | 2.439517 | 1.80189 | 1.965594 |
| Lrrc8c | 1.248076 | 1.285057 | 1.308925 | 1.243016 | 1.661743 | 1.622699 | 2.487973 | 2.277515 | 2.705177 | 3.077342 | 2.496188 | 2.39343 |
| Wsb2 | 21.84294 | 23.85612 | 24.34229 | 23.61516 | 16.68679 | 16.63044 | 14.03252 | 14.25418 | 9.832991 | 9.544562 | 13.32554 | 13.54121 |
| Soga3 | 1.87506 | 2.260081 | 2.269102 | 2.134481 | 2.412761 | 2.50275 | 2.973538 | 3.062743 | 3.574217 | 3.306679 | 4.696835 | 5.280838 |
| Coro6 | 0.481652 | 0.437376 | 0.842998 | 0.69242 | 1.277401 | 1.329737 | 1.015711 | 1.337149 | 0.479309 | 0.54786 | 0.715472 | 0.540566 |
| Tex101 | 0.534646 | 0.359024 | 1.322541 | 1.081481 | 1.050632 | 1.009455 | 0.818464 | 1.289512 | 1.010534 | 0.605586 | 1.857484 | 1.497616 |
| Ogfr | 13.87213 | 13.39479 | 18.26208 | 16.38333 | 22.96326 | 22.63944 | 26.64728 | 27.15574 | 30.78969 | 28.1132 | 40.84086 | 40.51519 |
| Mapk8ip1 | 9.873787 | 9.417007 | 12.28667 | 11.053 | 13.25363 | 12.34792 | 11.41568 | 12.91207 | 14.75256 | 13.55064 | 29.03859 | 27.31811 |
| Cyp39a1 | 1.168935 | 1.402351 | 1.403269 | 1.629205 | 1.629296 | 1.729328 | 1.87601 | 2.37943 | 2.354478 | 2.933439 | 1.295343 | 1.18764 |
| Bmp4 | 11.9537 | 13.84078 | 3.526569 | 3.026342 | 1.863482 | 1.683684 | 1.304525 | 1.339708 | 1.412512 | 1.225476 | 1.409214 | 1.501474 |
| Epha1 | 4.310487 | 4.178971 | 2.573614 | 2.272559 | 2.087217 | 1.964325 | 1.461193 | 1.918787 | 0.834882 | 0.920583 | 0.880035 | 0.977532 |
| Irgm1 | 0.667445 | 0.639602 | 0.523959 | 0.495698 | 0.67284 | 0.733541 | 1.256946 | 1.223045 | 0.709174 | 0.745946 | 1.429061 | 1.199685 |
| Cask | 2.158082 | 2.248215 | 2.220862 | 1.998088 | 2.982764 | 2.851782 | 3.527999 | 3.193192 | 4.714313 | 4.927124 | 4.243465 | 4.457375 |
| Hivep1 | 1.903703 | 2.627847 | 3.3502 | 3.710739 | 4.597202 | 4.527167 | 6.219225 | 4.818963 | 7.101148 | 7.690864 | 9.134906 | 9.149381 |
| Cited2 | 25.71326 | 27.0582 | 19.17964 | 20.07336 | 23.32607 | 21.87825 | 32.45854 | 28.55368 | 52.93246 | 49.47147 | 43.86027 | 39.39235 |
| B4galt4 | 2.333121 | 2.202518 | 2.880771 | 2.953363 | 4.765158 | 4.377362 | 5.149441 | 5.573604 | 7.048949 | 6.592158 | 7.320967 | 7.27855 |
| Elk3 | 3.671364 | 4.01702 | 3.47244 | 3.64801 | 3.447116 | 3.732627 | 4.115795 | 3.326985 | 3.587768 | 3.9645 | 1.807317 | 1.609179 |
| Psma8 | 15.70148 | 14.43223 | 13.79581 | 14.07653 | 11.2694 | 11.10028 | 9.913082 | 10.10425 | 6.365299 | 6.492458 | 5.381842 | 5.721116 |
| Pkp2 | 12.83672 | 11.971 | 14.82259 | 13.83401 | 12.41226 | 12.56578 | 13.22739 | 15.03073 | 16.46109 | 16.36456 | 30.23363 | 30.91566 |
| Slc25a19 | 7.484345 | 7.601247 | 7.607623 | 7.380992 | 6.156873 | 6.051862 | 5.534362 | 4.876759 | 3.29757 | 3.687014 | 3.588866 | 3.001699 |

| | | | | | | | | | | | | |
|---|---|---|---|---|---|---|---|---|---|---|---|---|
| Uggt2 | 0.873891 | 1.104482 | 1.763653 | 1.904328 | 2.828723 | 2.693268 | 3.280461 | 3.324834 | 4.405801 | 4.681072 | 4.408755 | 4.605009 |
| Cmbl | 0.807489 | 0.664621 | 1.818229 | 1.947275 | 2.998986 | 3.41302 | 5.357268 | 4.785094 | 4.495358 | 4.405358 | 3.217591 | 2.885358 |
| Rras2 | 27.29311 | 23.92702 | 24.09028 | 24.21077 | 28.22378 | 27.78157 | 30.654 | 31.99421 | 37.3561 | 37.75905 | 59.72283 | 59.47275 |
| Neil1 | 4.273863 | 4.575881 | 5.150427 | 4.709885 | 3.747299 | 4.323407 | 3.79451 | 3.321944 | 3.116896 | 3.374509 | 1.538063 | 1.917955 |
| Mthfd2 | 27.56294 | 28.04653 | 65.37309 | 63.17488 | 52.11501 | 50.61547 | 36.18234 | 33.8522 | 28.66143 | 25.73555 | 64.48513 | 63.12719 |
| Zfp934 | 3.009276 | 3.045725 | 4.042059 | 3.853801 | 3.108415 | 3.496787 | 3.055255 | 3.434938 | 2.819403 | 2.513057 | 1.175348 | 1.570333 |
| Syngr1 | 8.190255 | 7.827027 | 4.992039 | 5.059073 | 5.116399 | 4.990193 | 3.797796 | 4.13731 | 2.800222 | 3.099046 | 2.141799 | 2.062576 |
| Zranb3 | 11.52236 | 11.59977 | 13.36361 | 14.19064 | 12.40013 | 12.081 | 11.64304 | 11.80327 | 10.68144 | 12.3915 | 5.819427 | 5.786978 |
| Habp4 | 5.074523 | 4.636989 | 4.750466 | 4.493682 | 6.670224 | 6.311541 | 7.265085 | 7.835534 | 10.22316 | 9.491725 | 11.355 | 11.01727 |
| Nfkbia | 10.80538 | 10.56529 | 10.79367 | 10.62121 | 11.58653 | 11.33887 | 12.25583 | 13.31418 | 9.969595 | 9.564479 | 28.21593 | 23.43698 |
| Rab39b | 0.314381 | 0.224429 | 0.505752 | 0.463935 | 0.538996 | 0.541267 | 0.866524 | 0.564996 | 0.530993 | 0.57178 | 1.113852 | 1.136299 |
| Adam23 | 38.79421 | 43.89475 | 20.41283 | 20.67889 | 16.90637 | 17.95522 | 16.61596 | 16.04538 | 12.2769 | 13.043 | 7.745194 | 8.446797 |
| Gpx3 | 23.7358 | 18.54587 | 24.18256 | 25.54501 | 34.78729 | 38.35666 | 44.21793 | 48.21959 | 65.67872 | 62.49082 | 59.92587 | 57.95306 |
| Eif4e3 | 1.498134 | 1.596892 | 1.499213 | 1.589215 | 2.426828 | 2.485887 | 2.939505 | 2.776619 | 2.54983 | 2.786193 | 3.79571 | 3.886212 |
| Rbm20 | 0.764431 | 0.81597 | 0.709077 | 0.856136 | 1.136165 | 1.059685 | 1.457673 | 1.571242 | 2.130881 | 2.090805 | 1.949823 | 2.089775 |
| Sema4c | 2.570219 | 2.810557 | 5.12272 | 5.308462 | 8.721089 | 8.962829 | 12.41402 | 11.41808 | 13.97284 | 12.95584 | 9.90003 | 10.09548 |
| Pawr | 29.71754 | 28.68164 | 37.49365 | 34.00848 | 35.85333 | 35.10028 | 38.96822 | 38.58587 | 48.30907 | 43.40265 | 82.92375 | 82.99447 |
| Nfkbie | 1.224396 | 1.391163 | 1.683507 | 1.990135 | 2.396392 | 2.565516 | 2.615542 | 2.341419 | 2.583554 | 1.875099 | 4.480458 | 4.574085 |
| Dleu7 | 0.953161 | 0.605047 | 5.796513 | 6.093534 | 6.342115 | 6.441888 | 5.435231 | 5.955292 | 2.928345 | 2.641069 | 3.233101 | 3.692406 |
| Rtn4rl2 | 0.511847 | 0.780107 | 2.043103 | 1.844895 | 2.36609 | 2.603057 | 3.44351 | 3.057331 | 1.935007 | 1.752561 | 1.125486 | 1.253715 |
| Ttyh1 | 2.459197 | 2.501165 | 1.524756 | 1.342326 | 1.337346 | 1.106998 | 1.113069 | 1.108962 | 0.790209 | 0.705387 | 0.51299 | 0.550488 |
| Cpd | 2.417837 | 2.741308 | 6.317317 | 6.813298 | 8.034661 | 8.163439 | 11.20364 | 10.02104 | 12.99189 | 14.03655 | 14.33514 | 16.61179 |
| Naa38 | 5.05341 | 3.69381 | 10.23143 | 8.847187 | 9.808404 | 9.203841 | 9.222923 | 5.308344 | 5.576839 | 3.775645 | 9.488226 | 7.391292 |
| Tmem108 | 0.571171 | 0.705314 | 0.818511 | 0.796957 | 1.05145 | 1.05454 | 1.574193 | 1.248347 | 2.114566 | 1.941931 | 1.274461 | 1.180389 |
| Gltscr1l | 13.54998 | 15.22065 | 10.03708 | 10.05222 | 6.812094 | 6.649545 | 5.606677 | 5.342805 | 5.092547 | 5.176007 | 4.176684 | 4.126375 |
| Tmem53 | 0.231956 | 0.311048 | 0.623993 | 0.593873 | 1.116559 | 1.402528 | 1.596795 | 1.299858 | 1.138998 | 1.446226 | 0.938334 | 1.119418 |
| D430019H1 | 0.679687 | 0.879672 | 1.127605 | 1.281804 | 2.487473 | 2.119567 | 2.642922 | 2.373325 | 2.514016 | 2.571455 | 0.922096 | 1.056447 |
| Gpr160 | 2.585019 | 2.701793 | 1.834913 | 1.592723 | 1.547895 | 1.453045 | 1.382553 | 1.560379 | 0.678511 | 0.655277 | 1.095297 | 1.127859 |
| Rnft2 | 2.29764 | 2.317072 | 2.72502 | 2.769116 | 2.776598 | 2.762005 | 2.446985 | 2.369268 | 2.177551 | 2.589789 | 1.136184 | 1.139433 |
| 2200002D0 | 1.772479 | 1.941729 | 2.848849 | 2.392509 | 2.928224 | 3.254719 | 3.367978 | 3.763768 | 3.236253 | 2.457131 | 5.582792 | 5.814342 |
| E130317F2 | 0.771263 | 0.718105 | 0.86106 | 0.790326 | 1.129406 | 1.264076 | 1.184569 | 1.220123 | 1.39458 | 1.259176 | 1.99928 | 1.919897 |
| Rab11fip5 | 1.817604 | 1.875544 | 2.442002 | 1.993673 | 2.772747 | 2.452122 | 3.336639 | 3.425957 | 3.379807 | 3.236607 | 4.642534 | 4.842295 |
| Ovgp1 | 0.392672 | 0.360927 | 0.61596 | 0.514782 | 0.638595 | 0.588396 | 0.797846 | 0.824115 | 1.239646 | 1.12528 | 0.511905 | 0.463367 |
| Sh2d3c | 0.875892 | 0.841297 | 0.985796 | 1.03831 | 1.121291 | 1.090786 | 1.292623 | 1.206748 | 1.660721 | 1.197539 | 2.318013 | 2.5206 |
| Arhgap8 | 4.869643 | 4.819663 | 3.618901 | 3.352056 | 2.241116 | 2.271362 | 1.714611 | 1.935231 | 3.262908 | 2.943368 | 4.40452 | 4.69451 |
| Gpr63 | 0.246921 | 0.25815 | 0.683496 | 0.925859 | 1.155124 | 1.148415 | 1.654258 | 1.048083 | 1.839927 | 2.146951 | 2.349338 | 2.245868 |
| Calb2 | 0.478246 | 0.38503 | 0.694584 | 1.179612 | 1.831053 | 1.476307 | 1.324992 | 1.638709 | 1.318693 | 0.845606 | 0.795407 | 0.675651 |
| Itpripl2 | 0.678768 | 0.545765 | 1.051427 | 1.003752 | 1.48732 | 1.317318 | 1.537015 | 1.22256 | 1.365177 | 1.321 | 2.710915 | 2.433993 |
| 2410012M0 | 6.178241 | 6.530716 | 12.52113 | 12.83277 | 12.90313 | 11.88546 | 11.73462 | 9.595243 | 6.576996 | 6.742309 | 6.306393 | 5.29223 |
| Itpk1 | 34.51384 | 35.53337 | 29.5413 | 29.01636 | 24.86393 | 24.39378 | 24.74415 | 25.15211 | 19.03895 | 18.83349 | 12.91521 | 11.97182 |
| Isyna1 | 90.36359 | 95.7308 | 103.8935 | 98.71762 | 83.28405 | 81.51056 | 69.47606 | 74.10412 | 58.58709 | 56.80574 | 44.33357 | 40.73317 |
| Gata6 | 2.443125 | 2.644833 | 5.008279 | 4.42734 | 4.75267 | 5.102063 | 7.005374 | 6.632412 | 11.15776 | 8.400475 | 10.46478 | 10.71672 |
| Met | 0.835078 | 0.999358 | 1.026117 | 1.08996 | 1.780576 | 1.592988 | 2.875914 | 2.395322 | 2.536391 | 2.505031 | 2.644285 | 2.63825 |
| Oaf | 0.843563 | 1.099239 | 1.616428 | 1.439761 | 2.082931 | 1.822816 | 2.303237 | 2.610269 | 2.019415 | 2.098502 | 3.978761 | 3.483316 |
| Map1a | 3.420144 | 5.126577 | 3.646596 | 3.452386 | 2.613892 | 2.38759 | 2.503994 | 1.934134 | 1.568982 | 1.428517 | 1.898465 | 1.788867 |
| Rbp4 | 0.923255 | 0.879972 | 1.179748 | 1.326415 | 2.300169 | 2.417174 | 2.37265 | 3.20491 | 2.700688 | 2.350664 | 2.415686 | 2.331619 |
| Akt1 | 20.78035 | 20.04316 | 26.51135 | 25.24108 | 32.1062 | 31.18888 | 35.07671 | 36.57027 | 51.1674 | 48.52642 | 60.09945 | 60.98369 |
| Dstn | 170.5186 | 156.2524 | 98.86134 | 89.83441 | 71.60328 | 70.8037 | 75.85998 | 78.84756 | 92.49847 | 92.60167 | 164.8197 | 170.9844 |
| Cacna2d2 | 0.961392 | 1.165245 | 1.460818 | 1.589992 | 1.916524 | 2.084155 | 2.194358 | 2.60552 | 1.701927 | 1.5312 | 1.046358 | 1.079221 |
| Kbtbd11 | 4.74029 | 4.989175 | 5.735281 | 5.183964 | 3.39564 | 3.298179 | 2.551255 | 2.325543 | 2.201683 | 2.147676 | 3.864566 | 3.60251 |
| Slc25a47 | 0.394735 | 0.683691 | 0.622285 | 0.806377 | 1.275177 | 1.319404 | 1.361144 | 1.246623 | 1.608822 | 1.835366 | 2.421186 | 1.94532 |
| Hist1h1e | 9.011753 | 10.04475 | 2.826747 | 5.886888 | 5.883864 | 10.44697 | 5.688648 | 9.899733 | 5.525122 | 6.845934 | 4.313971 | 4.331512 |
| Mmgt1 | 7.900373 | 7.52193 | 8.987874 | 8.835837 | 8.648033 | 8.614986 | 10.22683 | 9.242801 | 13.66268 | 14.23928 | 18.0837 | 20.77093 |
| 1700013H1 | 3.770564 | 3.136416 | 9.873528 | 9.71977 | 14.86208 | 15.30887 | 17.21917 | 23.43354 | 16.91687 | 17.66263 | 17.28705 | 16.28017 |
| Dusp16 | 6.843926 | 7.03242 | 7.588657 | 7.688063 | 7.631696 | 7.573012 | 8.381622 | 7.226567 | 11.8227 | 12.94005 | 16.29271 | 17.41342 |
| Mmd | 4.660669 | 4.319529 | 5.741402 | 6.749672 | 9.434235 | 8.986458 | 12.47417 | 11.46729 | 17.35083 | 16.2593 | 11.06653 | 11.8661 |
| Map6 | 0.918097 | 1.186271 | 3.056359 | 3.028855 | 2.770885 | 2.719468 | 2.609886 | 2.438613 | 1.866054 | 1.840009 | 4.621189 | 4.431974 |
| Endod1 | 0.613665 | 0.471687 | 1.166808 | 1.199371 | 1.356282 | 1.338897 | 1.560568 | 1.562236 | 1.533048 | 1.528589 | 2.980756 | 2.888113 |
| Hoga1 | 0.374841 | 0.392188 | 0.920046 | 0.544458 | 1.087224 | 0.945785 | 1.145563 | 0.927227 | 1.426781 | 1.310869 | 1.008018 | 1.014103 |
| Rbks | 2.340636 | 2.518292 | 5.657914 | 4.499164 | 5.413133 | 4.272046 | 4.489376 | 5.127494 | 2.134576 | 2.347556 | 2.68847 | 2.130993 |
| Zfp473 | 2.800871 | 3.186174 | 3.094031 | 3.045837 | 2.109085 | 2.092729 | 1.714961 | 1.944966 | 1.154645 | 1.453026 | 1.376801 | 1.266163 |

| | | | | | | | | | | | | |
|---|---|---|---|---|---|---|---|---|---|---|---|---|
| Cnpy1 | 55.20769 | 45.5198 | 14.80296 | 14.23578 | 13.98909 | 15.28843 | 10.92764 | 10.9835 | 9.307433 | 8.196406 | 5.717225 | 6.358519 |
| Fkbp6 | 1.589856 | 1.991052 | 6.065558 | 6.463449 | 4.449693 | 4.554842 | 4.641961 | 3.227576 | 3.297637 | 2.892393 | 7.282425 | 6.954088 |
| Vwa5a | 5.110032 | 4.411814 | 4.594619 | 4.730269 | 6.425211 | 5.920367 | 10.51114 | 10.73309 | 9.914092 | 10.53165 | 10.6551 | 11.37187 |
| Lhx2 | 0.326657 | 0.433456 | 0.959893 | 1.037158 | 1.780791 | 1.441402 | 2.26179 | 2.491877 | 1.887648 | 1.443413 | 1.750061 | 1.578989 |
| Creld1 | 7.630203 | 7.794591 | 8.570109 | 8.186498 | 6.987641 | 7.14716 | 6.975802 | 6.503672 | 4.194249 | 3.975741 | 3.46315 | 3.408164 |
| Lzts2 | 11.13597 | 11.73133 | 16.33769 | 16.09291 | 19.83152 | 18.80739 | 24.58938 | 24.17181 | 31.6972 | 29.9319 | 39.67158 | 38.64893 |
| Slc7a1 | 10.9352 | 13.31227 | 24.61824 | 23.80397 | 20.0308 | 19.00769 | 16.99728 | 13.76178 | 15.92516 | 15.43882 | 32.11416 | 33.04539 |
| Angptl4 | 8.739463 | 7.3275 | 6.402674 | 6.702011 | 5.114006 | 5.975894 | 4.913781 | 5.019829 | 3.106089 | 2.791977 | 3.773934 | 3.66345 |
| Hist1h3f | 2.87119 | 2.769225 | 0.645976 | 1.31544 | 2.141177 | 2.282878 | 2.162793 | 3.039729 | 2.671747 | 2.309645 | 1.908337 | 1.26639 |
| Atf5 | 5.075478 | 5.230356 | 18.70374 | 18.07895 | 19.64282 | 17.32496 | 17.45397 | 15.64819 | 11.99007 | 10.13204 | 26.90219 | 24.2987 |
| Rem1 | 0.108995 | 0.114884 | 0.503698 | 0.459991 | 0.448225 | 0.509482 | 0.844135 | 0.901832 | 1.019652 | 0.91191 | 1.155377 | 1.103085 |
| Hoxb4 | 0.321342 | 0.304814 | 15.38273 | 14.57807 | 28.73768 | 27.7592 | 38.10202 | 34.95854 | 21.43203 | 19.50376 | 19.02349 | 18.60786 |
| Ppm1e | 0.562935 | 0.759914 | 0.684049 | 0.922373 | 1.098154 | 1.040431 | 1.579068 | 1.470137 | 1.863075 | 2.211565 | 1.658186 | 1.750289 |
| Foxp1 | 12.39637 | 14.61579 | 27.56227 | 28.06214 | 21.48105 | 21.88108 | 17.10999 | 15.24797 | 11.78114 | 11.70572 | 11.73842 | 12.23667 |
| Bbc3 | 14.43238 | 13.67047 | 13.54175 | 12.29525 | 12.92447 | 13.30005 | 14.54794 | 13.84827 | 15.42764 | 12.6946 | 33.31285 | 29.46768 |
| Pmel | 3.707392 | 4.197101 | 1.767182 | 1.752546 | 1.654525 | 1.251729 | 1.460225 | 1.144173 | 1.085092 | 1.261014 | 0.649966 | 0.731565 |
| Got1l1 | 10.91012 | 10.83772 | 9.379864 | 9.628779 | 12.06803 | 10.24017 | 12.17776 | 14.11201 | 10.5553 | 10.6444 | 5.417732 | 5.892523 |
| Tram1l1 | 0.679388 | 0.455806 | 1.263151 | 1.524634 | 1.857585 | 1.652243 | 1.82431 | 2.423433 | 2.009291 | 1.963554 | 3.482736 | 3.64995 |
| Inha | 0.928632 | 1.018413 | 1.885791 | 1.597649 | 1.598001 | 1.6909 | 1.134219 | 1.429843 | 0.763879 | 0.706914 | 0.860382 | 1.076912 |
| Pmepa1 | 4.183932 | 4.63 | 8.476417 | 7.69742 | 6.209952 | 5.69553 | 4.758801 | 4.564842 | 3.625173 | 3.42152 | 3.366914 | 3.219228 |
| Morn2 | 3.478527 | 2.035881 | 2.416052 | 2.898658 | 2.441899 | 3.259328 | 3.156526 | 2.619899 | 1.540816 | 1.933079 | 1.686843 | 1.363675 |
| Hip1 | 6.324755 | 8.368923 | 8.852108 | 9.417338 | 12.35154 | 11.44419 | 15.57717 | 14.27506 | 20.71331 | 22.50692 | 15.30722 | 15.55291 |
| Sepn1 | 2.969134 | 3.081691 | 6.541561 | 6.638425 | 9.735511 | 8.984661 | 12.69497 | 11.03315 | 16.62037 | 15.86298 | 15.04538 | 15.75586 |
| Daam1 | 4.449665 | 5.106991 | 6.713095 | 6.694442 | 6.369758 | 6.496141 | 6.519704 | 6.416514 | 7.689341 | 7.494752 | 13.7688 | 15.33202 |
| Perp | 7.409336 | 7.31032 | 15.74751 | 16.04545 | 11.38189 | 12.37532 | 10.96011 | 9.960181 | 6.915473 | 6.721391 | 9.758859 | 10.09807 |
| Stox2 | 1.595631 | 1.678321 | 5.258032 | 5.678956 | 6.506877 | 6.476954 | 8.099226 | 6.859691 | 12.80002 | 13.18223 | 12.0149 | 13.55036 |
| Khdrbs3 | 16.62891 | 15.39859 | 16.85605 | 17.64558 | 20.52802 | 19.45427 | 22.31503 | 21.03679 | 33.1356 | 33.49593 | 42.78289 | 42.10135 |
| Bhlhe40 | 15.38711 | 16.05766 | 14.06898 | 14.95737 | 14.03936 | 12.08578 | 10.37494 | 11.99906 | 6.121804 | 6.84778 | 7.452882 | 6.268965 |
| Btg1 | 6.134822 | 7.064477 | 4.508772 | 5.023419 | 6.384773 | 5.845137 | 8.759555 | 7.427639 | 12.90369 | 11.98492 | 11.49596 | 11.71489 |
| Tmem173 | 0.419544 | 0.351175 | 0.793496 | 0.79375 | 1.022947 | 1.09823 | 1.685277 | 1.588025 | 1.931997 | 1.893562 | 1.380235 | 1.580574 |
| Rab27a | 8.820009 | 9.202866 | 4.096678 | 4.666403 | 3.826235 | 3.922284 | 3.499164 | 3.668888 | 2.793026 | 2.917696 | 1.858371 | 1.956466 |
| Shisa7 | 1.225336 | 1.673743 | 2.590299 | 2.241461 | 1.709951 | 1.801084 | 1.188975 | 1.404608 | 1.273521 | 1.152516 | 0.745218 | 0.940922 |
| Sox15 | 15.23509 | 12.94362 | 11.36019 | 8.71831 | 8.160521 | 6.814729 | 5.540929 | 6.259832 | 4.265311 | 4.894752 | 3.711819 | 3.663747 |
| Mctp2 | 1.98513 | 2.191114 | 1.353457 | 1.494354 | 1.520478 | 1.581023 | 1.370024 | 1.408684 | 0.796356 | 0.794742 | 0.558347 | 0.6645 |
| Laptm5 | 14.0611 | 18.04193 | 15.99474 | 13.66967 | 9.81602 | 9.500871 | 10.41525 | 9.517624 | 6.944185 | 5.746177 | 5.561348 | 6.023147 |
| Zswim5 | 1.903407 | 2.348408 | 3.716072 | 3.973093 | 5.147914 | 4.944156 | 6.901593 | 6.34746 | 9.094683 | 9.450561 | 6.965972 | 6.937054 |
| Whsc1 | 10.95808 | 12.464 | 19.12682 | 20.10748 | 24.18171 | 23.3511 | 29.65572 | 28.92926 | 41.91601 | 43.5495 | 46.08489 | 47.82279 |
| Ddit3 | 4.664128 | 4.552756 | 13.48147 | 13.26483 | 11.17618 | 10.10772 | 7.477337 | 8.319029 | 6.448061 | 5.578693 | 10.45611 | 11.08599 |
| Magel2 | 0.540208 | 0.487016 | 0.434334 | 0.549833 | 0.873044 | 0.801462 | 1.201147 | 1.30689 | 1.139747 | 1.043878 | 1.116338 | 1.139937 |
| Lpar1 | 3.230293 | 3.436031 | 3.349649 | 3.49487 | 4.047958 | 3.737875 | 4.955937 | 5.555794 | 8.416946 | 8.304942 | 6.307744 | 5.35999 |
| Fkbp11 | 32.79323 | 27.60598 | 19.96428 | 20.29337 | 25.05178 | 25.66504 | 19.42432 | 16.60544 | 15.85047 | 12.43902 | 10.97088 | 10.80311 |
| Snx33 | 0.551747 | 0.673296 | 1.986195 | 2.155149 | 2.85905 | 2.865878 | 3.745508 | 3.717328 | 4.274175 | 4.235449 | 4.916081 | 5.119639 |
| Tex19.2 | 0.394045 | 0.412658 | 0.965702 | 0.976888 | 0.678671 | 0.720009 | 1.021174 | 0.916268 | 0.931543 | 1.023148 | 1.927389 | 1.709791 |
| Gtsf1 | 4.673139 | 4.798085 | 9.696841 | 9.079516 | 8.066847 | 6.86047 | 7.145595 | 5.722477 | 3.395591 | 3.824231 | 5.221955 | 5.046328 |
| Upp1 | 330.307 | 258.2712 | 202.6765 | 196.0411 | 174.6344 | 173.3296 | 132.9318 | 148.4186 | 125.0608 | 131.8086 | 87.9485 | 82.57394 |
| Epas1 | 11.84889 | 13.60861 | 11.38668 | 11.31324 | 8.959111 | 8.870756 | 8.628964 | 7.018146 | 4.548391 | 4.767175 | 5.561132 | 6.039424 |
| Cgnl1 | 9.222593 | 10.39218 | 21.03567 | 21.24875 | 19.2272 | 17.17352 | 16.36068 | 16.55684 | 10.65036 | 11.30903 | 9.466649 | 8.954447 |
| L3hypdh | 1.549112 | 1.367409 | 1.248063 | 1.089533 | 1.227625 | 1.466451 | 0.908482 | 1.368557 | 0.799818 | 0.85202 | 0.610271 | 0.618263 |
| Nt5m | 1.070195 | 1.241322 | 2.037337 | 1.806576 | 1.978476 | 2.571647 | 3.091371 | 3.230029 | 3.747188 | 3.429681 | 4.673988 | 4.282849 |
| Hoxa6 | 0.022902 | 0.072161 | 1.475426 | 1.625779 | 1.520875 | 1.838578 | 2.144747 | 2.413065 | 3.045486 | 3.474572 | 3.189739 | 3.853286 |
| Lpin3 | 4.101664 | 3.925938 | 4.310777 | 3.929839 | 3.146978 | 3.524012 | 3.001116 | 2.795818 | 2.033602 | 1.658367 | 2.090917 | 2.438348 |
| Sh3kbp1 | 0.606164 | 0.681581 | 1.1282 | 1.298786 | 1.921291 | 1.587496 | 2.613036 | 2.600525 | 3.026139 | 3.076734 | 2.287417 | 2.29296 |
| Hist1h2bj | 1.568962 | 2.053225 | 0.639887 | 1.215221 | 1.308671 | 1.916114 | 1.78482 | 0.809799 | 1.575825 | 1.44727 | 1.214938 | 1.26297 |
| Fut9 | 2.930147 | 3.469752 | 0.648004 | 0.730215 | 0.640949 | 0.603542 | 0.564659 | 0.444183 | 1.283484 | 1.050752 | 0.615145 | 0.715071 |
| Htra1 | 10.77029 | 12.11557 | 11.27958 | 11.06329 | 11.61923 | 10.67819 | 12.80712 | 13.66287 | 20.35804 | 19.11852 | 25.69 | 25.24845 |
| 0610040B1 | 0.936876 | 0.854756 | 1.101241 | 0.934478 | 1.144145 | 1.33518 | 1.072794 | 2.187131 | 0.954939 | 0.9251 | 1.99897 | 1.84848 |
| Nfu1 | 45.44471 | 41.54734 | 55.91483 | 46.61606 | 35.73087 | 38.61613 | 30.75181 | 28.47323 | 21.29993 | 19.79307 | 21.75367 | 19.71806 |
| Gm20748 | 11.17601 | 10.20935 | 10.17139 | 8.922291 | 7.808194 | 8.106729 | 5.957229 | 6.456181 | 5.42228 | 3.775814 | 3.528416 | 4.084339 |
| Tmem200b | 0.325288 | 0.231524 | 5.543279 | 5.361144 | 7.317226 | 7.729359 | 8.90695 | 9.304999 | 5.448841 | 4.518652 | 4.599663 | 3.938128 |
| Gatsl3 | 21.2782 | 23.90402 | 11.94903 | 10.80791 | 10.82739 | 10.19439 | 9.435322 | 9.141167 | 7.0916 | 6.329765 | 4.332384 | 4.580313 |
| Gm11944 | 3.068249 | 5.096632 | 2.976209 | 3.44176 | 3.784745 | 4.150063 | 4.894275 | 4.294808 | 6.255382 | 6.886734 | 6.240434 | 8.120169 |

| | | | | | | | | | | | | |
|---|---|---|---|---|---|---|---|---|---|---|---|---|
| Tmem128 | 11.94185 | 10.76539 | 12.04867 | 12.38627 | 15.77556 | 14.37253 | 17.94442 | 15.28036 | 20.5358 | 19.58456 | 29.42418 | 29.21978 |
| Nab2 | 0.968929 | 1.122018 | 2.856211 | 3.000771 | 3.4239 | 3.696546 | 4.094268 | 3.230961 | 3.552592 | 3.239825 | 7.135642 | 7.076139 |
| Ypel1 | 2.298524 | 2.008438 | 2.250129 | 2.428237 | 2.891799 | 2.781506 | 3.062955 | 3.809185 | 5.219908 | 5.725345 | 3.655501 | 4.486047 |
| Mbd6 | 6.887175 | 9.183783 | 28.56079 | 27.37416 | 21.98022 | 22.3781 | 18.07929 | 15.21787 | 13.08376 | 11.61742 | 16.72554 | 15.31836 |
| Cldn6 | 10.87489 | 11.51553 | 10.29059 | 9.563326 | 6.897061 | 6.511137 | 5.070036 | 4.325714 | 3.942155 | 4.102847 | 5.204335 | 4.060597 |
| Hist1h3h | 8.074999 | 9.646238 | 2.717462 | 4.117221 | 5.839345 | 7.750927 | 6.915688 | 9.692707 | 4.565777 | 7.47138 | 4.756878 | 4.707667 |
| Taf9b | 2.684393 | 2.561812 | 2.99194 | 2.901786 | 3.365346 | 3.466569 | 4.120309 | 4.165172 | 4.252098 | 4.35746 | 6.281364 | 6.831278 |
| Slc22a7 | 1.065342 | 1.12785 | 0.935624 | 0.710918 | 0.892991 | 0.86497 | 0.557329 | 0.467036 | 0.3731 | 0.367437 | 0.268321 | 0.382576 |
| P4ha2 | 5.860828 | 6.339731 | 9.00008 | 8.462971 | 12.26537 | 11.2216 | 14.91294 | 17.2765 | 19.74631 | 19.91286 | 17.42959 | 17.41829 |
| Lingo1 | 0.440168 | 0.341326 | 0.506497 | 0.740697 | 0.861814 | 1.084458 | 1.455467 | 1.351251 | 1.917085 | 1.299466 | 1.761482 | 1.742428 |
| Pitpnc1 | 17.00376 | 20.40797 | 14.16701 | 15.14861 | 12.78241 | 12.31571 | 11.87626 | 10.56713 | 9.80978 | 9.767397 | 5.416545 | 6.439633 |
| Jph3 | 1.153682 | 1.249948 | 1.517845 | 1.793799 | 2.46979 | 2.518116 | 2.106097 | 2.023133 | 1.587414 | 1.594731 | 1.244359 | 1.070779 |
| 1700017B0 | 6.354692 | 8.253649 | 10.72542 | 11.29647 | 10.16899 | 9.64631 | 11.329 | 9.745771 | 13.73756 | 12.94092 | 22.57928 | 22.64585 |
| Copz2 | 2.179883 | 2.157291 | 2.033533 | 2.811184 | 4.452985 | 4.939547 | 5.654318 | 5.516413 | 7.08048 | 6.599356 | 4.899884 | 5.270422 |
| Sall2 | 7.449063 | 8.013085 | 11.23263 | 11.55259 | 15.76632 | 14.58029 | 21.93667 | 20.02222 | 27.68577 | 27.1065 | 17.79486 | 17.89946 |
| Mylpf | 65.92069 | 58.92042 | 66.31631 | 61.52654 | 51.93287 | 50.8063 | 33.52565 | 32.79473 | 25.94311 | 26.24154 | 28.73306 | 26.67322 |
| Wdr31 | 6.964097 | 6.91988 | 5.446907 | 5.117707 | 3.39794 | 3.317942 | 2.165151 | 2.794742 | 2.086973 | 2.182888 | 2.153971 | 2.228937 |
| Flna | 64.5409 | 78.73522 | 77.06715 | 75.49433 | 68.21321 | 66.84028 | 81.04482 | 68.26655 | 93.94207 | 94.92897 | 154.5259 | 156.6608 |
| Irf5 | 0.440889 | 0.579928 | 1.270685 | 0.926772 | 0.833984 | 0.726653 | 0.674653 | 0.439183 | 0.368868 | 0.395413 | 0.749069 | 0.744659 |
| Cacna1h | 0.379919 | 0.551205 | 0.769442 | 0.758737 | 1.266613 | 1.161951 | 1.447261 | 1.776735 | 1.000842 | 1.053499 | 1.132741 | 1.110422 |
| Agrp | 1.068991 | 1.125409 | 0.815056 | 0.998369 | 0.580605 | 0.912326 | 0.782009 | 0.568948 | 0.532165 | 0.63166 | 0.482902 | 0.426412 |
| Hoxa5 | 1.246064 | 1.003869 | 26.51179 | 26.03709 | 47.02764 | 51.03185 | 64.24551 | 60.94969 | 42.74617 | 37.91286 | 37.64819 | 40.5819 |
| Gfra4 | 0.595748 | 0.627956 | 1.531632 | 1.019607 | 1.820635 | 1.683469 | 1.30479 | 1.229901 | 1.419051 | 0.719461 | 1.268651 | 1.116645 |
| Alg2 | 5.705603 | 6.434845 | 9.235515 | 7.893747 | 9.756426 | 9.109726 | 13.59076 | 10.5959 | 15.42747 | 14.65775 | 17.49302 | 18.47037 |
| Creg1 | 6.425681 | 5.597652 | 6.110037 | 6.197815 | 9.538005 | 8.683965 | 11.27494 | 12.90514 | 14.67718 | 14.50149 | 10.36151 | 10.17358 |
| Setd7 | 5.060117 | 5.559064 | 7.879982 | 7.823291 | 10.15171 | 10.06249 | 12.16853 | 10.32214 | 16.70418 | 16.76316 | 16.81714 | 18.29018 |
| A330009N2 | 0.50768 | 0.671366 | 1.701157 | 1.623677 | 1.085953 | 1.475872 | 0.963876 | 1.178411 | 0.482753 | 0.694952 | 0.574044 | 0.703037 |
| Gbx1 | 0.197124 | 0.256479 | 0.677156 | 0.685315 | 0.788278 | 0.842109 | 0.963662 | 0.917111 | 1.118706 | 0.991424 | 1.679818 | 1.600971 |
| Tesc | 0.62504 | 0.75774 | 1.660079 | 1.489102 | 1.326584 | 1.434922 | 0.714946 | 1.363737 | 0.64402 | 0.770965 | 1.039396 | 0.637685 |
| Tet3 | 2.056361 | 3.06438 | 3.915324 | 4.623507 | 5.355323 | 5.257547 | 6.201441 | 5.252346 | 9.197551 | 9.375892 | 10.652 | 10.79398 |
| Lgi2 | 1.706527 | 1.852535 | 2.108479 | 2.210899 | 3.532589 | 3.348774 | 3.463576 | 3.052369 | 4.871329 | 5.159946 | 3.573629 | 3.224896 |
| Nav2 | 6.66239 | 9.832781 | 7.092486 | 6.959013 | 4.216576 | 3.884077 | 3.906606 | 3.520105 | 3.166642 | 3.623006 | 3.050294 | 2.981786 |
| Pnma5 | 10.98087 | 9.0631 | 21.28142 | 20.24691 | 24.31533 | 24.46936 | 26.57818 | 26.74033 | 12.94183 | 11.45882 | 13.08676 | 13.64768 |
| Klk8 | 8.217987 | 8.116606 | 6.710325 | 7.325005 | 7.180472 | 7.474173 | 6.72146 | 7.303041 | 4.103425 | 4.0065 | 2.714112 | 3.203995 |
| Fam101b | 0.665487 | 0.758967 | 1.718141 | 1.529973 | 2.771265 | 2.861568 | 3.655426 | 3.336211 | 3.617304 | 3.56208 | 3.636433 | 3.568274 |
| Tmem170b | 1.626101 | 2.039221 | 2.40144 | 2.419211 | 2.837066 | 2.948011 | 3.328422 | 2.500451 | 5.068076 | 5.428889 | 4.859536 | 5.640059 |
| BC021614 | 7.110733 | 6.641442 | 2.763639 | 2.858262 | 2.079624 | 2.591218 | 1.948298 | 2.316876 | 1.619954 | 1.226292 | 2.023761 | 1.630275 |
| Car4 | 10.08346 | 11.01094 | 8.649628 | 9.251488 | 7.938982 | 6.864415 | 7.600536 | 8.582349 | 6.246028 | 5.860176 | 4.461406 | 3.969679 |
| Ptprj | 1.949645 | 2.243979 | 2.054101 | 2.087861 | 2.98831 | 3.078665 | 4.023694 | 3.38887 | 4.055218 | 4.448151 | 4.523576 | 4.865428 |
| Fhod3 | 4.838109 | 4.923955 | 5.284445 | 5.717611 | 7.800257 | 7.154186 | 10.45855 | 10.3735 | 14.07203 | 13.31912 | 10.16371 | 10.53342 |
| Zmynd11 | 10.49782 | 10.51282 | 12.5199 | 12.74544 | 15.5479 | 14.73515 | 16.7854 | 17.14335 | 26.18673 | 26.31989 | 29.33662 | 29.68127 |
| Mif4gd | 28.36009 | 27.98759 | 34.55598 | 31.36503 | 28.02952 | 27.32023 | 24.18042 | 23.54675 | 17.81991 | 16.57648 | 14.02977 | 13.47397 |
| Cyp26c1 | 0.057925 | 0.193948 | 5.695222 | 6.091406 | 5.876077 | 6.650911 | 5.915753 | 6.277561 | 2.652155 | 2.858188 | 3.133719 | 2.948613 |
| Gcdh | 25.22012 | 22.68328 | 24.63787 | 24.15854 | 23.88451 | 22.96796 | 21.08769 | 22.75227 | 13.50792 | 14.07739 | 10.44863 | 10.38369 |
| 2210409E1 | 1.283166 | 1.109676 | 0.90556 | 1.222048 | 1.1661 | 1.222112 | 1.164944 | 2.167264 | 2.967318 | 2.38349 | 3.018811 | 2.84279 |
| St7 | 0.861654 | 0.952436 | 1.310417 | 1.383306 | 1.466258 | 1.65267 | 1.825661 | 1.72861 | 2.796884 | 2.474653 | 3.367892 | 3.217911 |
| Fam169a | 59.54252 | 64.53355 | 62.33545 | 62.3319 | 53.1866 | 51.08831 | 40.65877 | 43.53194 | 30.88078 | 34.33229 | 25.20439 | 26.79793 |
| Best1 | 0.435586 | 0.571472 | 0.905859 | 1.102373 | 1.287269 | 1.094894 | 1.177477 | 0.811161 | 0.804554 | 0.691896 | 0.262066 | 0.474084 |
| Gpr173 | 0.3028 | 0.257063 | 0.406532 | 0.426573 | 0.721166 | 0.462066 | 0.799203 | 0.598509 | 1.009513 | 0.991317 | 0.784655 | 0.650182 |
| Slitrk5 | 0.473755 | 0.461605 | 0.928232 | 1.00597 | 1.086102 | 1.165846 | 1.095972 | 1.345747 | 1.075126 | 0.9936 | 0.563882 | 0.579554 |
| Zfp334 | 1.216516 | 1.822822 | 2.013237 | 2.170725 | 2.41336 | 2.353105 | 2.971022 | 2.827448 | 4.655819 | 5.038531 | 5.231999 | 5.039605 |
| BC030307 | 0.604861 | 0.663277 | 1.3106 | 1.219617 | 1.097787 | 1.154579 | 1.073426 | 0.998543 | 0.835102 | 0.93323 | 0.482045 | 0.525437 |
| Ppfibp2 | 7.111667 | 7.140307 | 4.052845 | 3.757229 | 2.763572 | 2.941223 | 2.261074 | 2.32743 | 1.901689 | 1.849595 | 1.731729 | 1.619207 |
| Lims2 | 0.478276 | 0.385901 | 1.160419 | 0.968933 | 0.728462 | 0.635881 | 0.595318 | 0.944005 | 0.360409 | 0.443898 | 0.811445 | 0.417707 |
| Rab38 | 0.630663 | 0.490095 | 1.024426 | 1.098915 | 1.902915 | 1.556419 | 2.073139 | 2.548856 | 2.387218 | 2.014566 | 1.357509 | 1.2966 |
| Pld1 | 0.536576 | 0.595931 | 0.590774 | 0.655821 | 1.013303 | 1.021117 | 1.146066 | 1.391805 | 1.380881 | 1.520323 | 1.289336 | 1.17515 |
| Spire2 | 10.07358 | 10.25244 | 8.045461 | 7.704128 | 5.852445 | 5.207368 | 4.263212 | 5.672326 | 3.313045 | 3.323969 | 3.580347 | 4.219346 |
| H1fx | 8.437944 | 8.764233 | 20.08477 | 14.6676 | 25.96269 | 33.99169 | 32.05287 | 25.76342 | 22.23373 | 15.04058 | 26.55811 | 23.80255 |
| Camk1d | 5.584752 | 5.872346 | 5.745265 | 5.205322 | 3.666956 | 3.792558 | 3.249636 | 3.110111 | 2.041284 | 2.395927 | 2.545079 | 2.247234 |
| Zic5 | 13.91713 | 14.2339 | 10.4278 | 9.756509 | 8.476597 | 7.937219 | 6.24056 | 5.721973 | 5.590797 | 5.607349 | 4.280608 | 4.212177 |
| Fam89a | 0.950081 | 0.851754 | 1.616001 | 1.883458 | 2.60267 | 2.291655 | 2.885487 | 2.708668 | 1.513128 | 1.602426 | 1.270365 | 1.169825 |

| | | | | | | | | | | | | |
|---|---|---|---|---|---|---|---|---|---|---|---|---|
| Anxa11 | 11.40474 | 10.33245 | 9.781351 | 9.602578 | 9.455146 | 8.896363 | 8.176808 | 8.608591 | 5.233168 | 4.833446 | 4.661902 | 4.148264 |
| Crybg3 | 4.21781 | 4.758508 | 5.741551 | 5.723 | 5.931779 | 6.000537 | 6.317878 | 5.429451 | 8.237044 | 8.675394 | 12.37555 | 12.56731 |
| Tdrd1 | 0.486282 | 0.447845 | 1.113589 | 0.893936 | 0.698268 | 0.743496 | 0.998261 | 1.120401 | 1.221166 | 1.32954 | 1.993563 | 1.72084 |
| AU022751 | 2.425759 | 2.965901 | 9.821462 | 8.986087 | 12.62554 | 12.45883 | 16.53793 | 14.92677 | 15.98363 | 14.56947 | 22.30454 | 20.79521 |
| Cdkn2b | 1.144953 | 0.914781 | 1.358739 | 1.745007 | 2.038969 | 2.41156 | 2.489108 | 2.186907 | 1.902769 | 1.703377 | 3.482083 | 3.941498 |
| Phlpp2 | 1.795449 | 1.99359 | 2.562287 | 2.855993 | 3.187725 | 3.230086 | 3.722789 | 3.530264 | 5.233877 | 5.610872 | 6.195954 | 6.607191 |
| Camk2n2 | 1.718424 | 1.801504 | 4.110977 | 2.527288 | 3.368153 | 3.367188 | 3.752568 | 2.541205 | 3.438622 | 2.338493 | 5.59726 | 5.407452 |
| Prep | 26.43529 | 28.19982 | 24.80947 | 25.54636 | 30.71067 | 30.5702 | 37.26532 | 34.6394 | 58.35792 | 59.0626 | 57.44542 | 56.31407 |
| Gm14511 | 0.047097 | 0.148397 | 0.614348 | 0.964908 | 1.089384 | 0.862865 | 1.539968 | 0.441102 | 0.581342 | 0.417448 | 0.810265 | 0.664319 |
| Iglon5 | 1.514965 | 2.061187 | 1.991459 | 1.629347 | 3.005657 | 3.118725 | 3.691353 | 3.471636 | 4.113573 | 3.761612 | 3.26997 | 3.39765 |
| Sgcb | 2.097596 | 1.762629 | 2.379677 | 2.197305 | 3.453549 | 3.420816 | 4.527651 | 4.018262 | 5.520518 | 5.071961 | 4.494706 | 5.037044 |
| Aim2 | 17.31732 | 15.52 | 15.45384 | 14.36584 | 12.82205 | 12.95773 | 12.239 | 13.48729 | 8.883306 | 9.414894 | 5.693438 | 6.22416 |
| Sct | 0.86994 | 0.956944 | 1.101639 | 1.364822 | 1.469801 | 1.120017 | 1.183122 | 0.965427 | 1.175236 | 1.659402 | 2.385924 | 2.227704 |
| Endov | 2.249589 | 2.235623 | 2.505246 | 2.541924 | 2.569644 | 2.97077 | 3.279095 | 3.264056 | 4.187258 | 4.20392 | 6.04614 | 5.859255 |
| Bcor | 5.710312 | 6.556198 | 6.022737 | 5.726736 | 6.618834 | 5.789664 | 8.289231 | 7.339652 | 10.41443 | 10.17807 | 14.41443 | 13.18451 |
| Ppfia3 | 1.779906 | 1.853956 | 2.078723 | 2.027243 | 1.592945 | 1.54172 | 1.048964 | 1.178143 | 0.914913 | 0.909485 | 0.820658 | 0.880589 |
| Brsk2 | 0.974238 | 1.095903 | 1.316306 | 1.402496 | 1.784121 | 1.581438 | 1.544989 | 1.875571 | 3.173999 | 3.227042 | 3.196907 | 3.131095 |
| Dazl | 7.976172 | 8.85959 | 26.8784 | 25.6995 | 31.39734 | 30.15717 | 45.05998 | 41.49856 | 48.02034 | 50.14504 | 61.93068 | 59.12734 |
| Btbd17 | 0.507314 | 0.582771 | 1.036075 | 1.026468 | 1.027502 | 1.011724 | 1.422469 | 1.008729 | 1.273209 | 1.356707 | 2.608739 | 2.320719 |
| Atp2b4 | 0.44884 | 0.546898 | 0.489184 | 0.564093 | 0.611526 | 0.614425 | 0.852471 | 0.591585 | 0.98711 | 1.046985 | 1.289943 | 1.295222 |
| Maged2 | 8.390101 | 7.488412 | 8.303642 | 7.519175 | 11.7877 | 11.42518 | 16.07034 | 16.80072 | 18.3765 | 17.26076 | 16.50122 | 16.57371 |
| Prkar1b | 0.527652 | 0.617481 | 0.590854 | 0.74201 | 1.000856 | 0.924137 | 1.658411 | 1.581786 | 1.668957 | 1.62292 | 2.159611 | 1.702429 |
| Dock7 | 5.839535 | 6.819125 | 7.553211 | 7.97982 | 10.04284 | 9.569911 | 12.60611 | 11.65709 | 18.51865 | 18.30347 | 15.26561 | 16.05364 |
| Mbnl1 | 1.923371 | 2.231594 | 3.167361 | 3.252961 | 3.304982 | 3.254034 | 3.662724 | 3.124473 | 4.677915 | 4.953793 | 6.627391 | 7.166417 |
| Cdkn2c | 0.455991 | 0.248346 | 1.613018 | 1.240689 | 1.849028 | 1.87177 | 2.165498 | 2.845028 | 2.088572 | 2.206068 | 3.054069 | 2.705414 |
| Taf7l | 2.231523 | 1.872169 | 6.870633 | 5.82677 | 9.258048 | 9.254426 | 9.649392 | 10.9106 | 6.151759 | 5.845361 | 12.74192 | 13.35924 |
| Dnaja4 | 0.348826 | 0.343543 | 0.736737 | 0.639537 | 1.153382 | 1.126475 | 1.446517 | 1.420215 | 1.139782 | 1.464762 | 1.364739 | 1.465702 |
| Fam131c | 3.41813 | 3.911301 | 3.38233 | 3.450876 | 3.512318 | 3.133145 | 2.675147 | 2.649152 | 2.025978 | 2.373725 | 1.679341 | 1.505878 |
| 2310022B0 | 10.06983 | 9.787546 | 8.955633 | 9.090376 | 12.09418 | 11.87331 | 14.55295 | 13.87391 | 20.88605 | 20.82954 | 20.43392 | 20.24562 |
| Klhl36 | 2.077963 | 2.427392 | 4.678001 | 4.206908 | 5.539163 | 4.444585 | 5.83852 | 4.969094 | 3.191567 | 3.080261 | 2.603694 | 2.169467 |
| Bche | 1.074552 | 0.979877 | 1.957421 | 2.10344 | 2.366438 | 2.473743 | 3.029425 | 2.769762 | 4.732191 | 4.816326 | 4.244257 | 4.656625 |
| Hist1h1b | 4.848066 | 4.041013 | 0.846269 | 2.495437 | 3.099327 | 3.926448 | 2.354751 | 2.901596 | 1.906864 | 2.569397 | 2.255656 | 1.715664 |
| Triml2 | 66.117 | 64.55769 | 52.31962 | 49.08847 | 39.78893 | 37.176 | 28.6916 | 32.93795 | 21.12636 | 21.45795 | 23.88659 | 23.41551 |
| Lrrfip1 | 8.213712 | 8.705195 | 8.315241 | 8.340706 | 9.996661 | 9.536099 | 10.26817 | 10.51777 | 13.17809 | 12.7375 | 18.421 | 19.05651 |
| Nqo1 | 17.58345 | 15.56336 | 13.87284 | 12.62496 | 9.753403 | 10.60533 | 9.712374 | 10.20031 | 6.490824 | 6.32344 | 6.019066 | 5.527514 |
| Gpr153 | 2.431498 | 2.820699 | 3.736954 | 4.579248 | 6.977974 | 6.359406 | 9.078871 | 8.376973 | 9.759297 | 10.45823 | 7.43888 | 6.356297 |
| Ccdc8 | 1.408006 | 1.397364 | 1.854418 | 1.851322 | 2.035592 | 2.126323 | 2.357471 | 1.997869 | 2.941733 | 2.922477 | 4.671336 | 4.225094 |
| Smo | 14.39075 | 15.05986 | 18.87272 | 19.03299 | 24.57806 | 24.13042 | 32.17195 | 27.58201 | 43.97079 | 43.42089 | 41.11059 | 41.88414 |
| Fabp3 | 242.3816 | 202.7325 | 76.25215 | 66.72802 | 64.22135 | 66.81607 | 53.04686 | 44.64617 | 65.01622 | 57.90149 | 93.45477 | 101.8501 |
| Unc13a | 2.433952 | 2.719252 | 1.609955 | 1.54839 | 1.280518 | 1.177505 | 1.037781 | 1.041999 | 0.779665 | 0.863472 | 0.712451 | 0.679027 |
| Ephb3 | 5.692384 | 7.290629 | 6.895751 | 6.248534 | 7.566223 | 6.720091 | 9.532721 | 8.834466 | 12.34359 | 10.81454 | 14.07372 | 14.24767 |
| Kcnj12 | 1.816913 | 1.714544 | 3.437287 | 2.891939 | 4.700011 | 4.706539 | 6.206098 | 6.146192 | 5.736994 | 6.592999 | 4.552365 | 4.412426 |
| Card10 | 2.560641 | 3.401372 | 12.52924 | 11.90837 | 11.28231 | 10.61457 | 9.422436 | 9.183014 | 4.85859 | 5.223699 | 12.85556 | 11.51749 |
| Zswim6 | 5.484599 | 6.278414 | 6.421659 | 6.549884 | 7.230643 | 7.75902 | 9.660953 | 9.23426 | 13.48758 | 14.92382 | 11.66544 | 12.31701 |
| Creb3l1 | 2.540606 | 3.109332 | 3.869827 | 3.979093 | 6.205367 | 6.015092 | 7.617761 | 6.823906 | 5.737617 | 4.764849 | 10.35966 | 9.062342 |
| Hopx | 1.549202 | 0.980611 | 0.660896 | 0.45353 | 0.560132 | 0.602418 | 0.424722 | 0.743332 | 0.717252 | 0.573794 | 0.421842 | 0.326437 |
| Ccdc136 | 11.3582 | 12.42048 | 6.705168 | 6.763709 | 8.415325 | 8.428098 | 8.957202 | 9.909952 | 7.704633 | 9.148019 | 4.756434 | 4.354439 |
| BC068157 | 1.066311 | 1.529282 | 1.773576 | 1.969279 | 2.602732 | 2.619521 | 3.518704 | 2.670287 | 4.197241 | 4.481023 | 2.970601 | 3.149546 |
| Gskip | 2.648161 | 2.321867 | 3.019022 | 2.850922 | 3.309691 | 3.587563 | 4.123129 | 5.267483 | 6.265318 | 6.484481 | 5.948299 | 5.738203 |
| Prkar2b | 4.158232 | 4.2182 | 5.020677 | 4.817518 | 5.9269 | 6.177704 | 7.257669 | 7.31171 | 10.64741 | 10.95407 | 7.428641 | 7.856363 |
| Myh9 | 47.4372 | 63.95959 | 76.0982 | 74.20985 | 67.91664 | 65.12599 | 65.92618 | 56.36914 | 66.57027 | 68.85687 | 131.4771 | 128.1587 |
| Creb3l2 | 1.612135 | 2.139035 | 3.436721 | 3.001479 | 4.253145 | 4.083505 | 6.02134 | 4.917008 | 6.496722 | 6.8235 | 6.024576 | 5.907409 |
| Ttc9 | 1.09405 | 0.899753 | 1.351711 | 1.446044 | 1.647705 | 1.542053 | 1.844499 | 1.90498 | 2.692143 | 2.537588 | 3.531515 | 3.287282 |
| Sh2b2 | 1.182598 | 1.239644 | 1.254307 | 1.334424 | 1.711103 | 1.492282 | 1.413105 | 1.694983 | 3.162268 | 3.03112 | 2.662165 | 2.678492 |
| 1700042O1 | 1.385889 | 0.986678 | 1.558418 | 1.726067 | 2.848335 | 2.709049 | 3.859107 | 3.662173 | 2.164893 | 1.86432 | 3.677327 | 3.919329 |
| Hist1h2bm | 1.42578 | 1.988835 | 0.648936 | 0.87371 | 0.636025 | 1.105732 | 0.582062 | 0.984086 | 0.633783 | 0.738796 | 0.511114 | 0.487492 |
| 1810014B0 | 2.0632 | 2.113058 | 2.660125 | 2.314198 | 2.09793 | 1.957953 | 1.775097 | 1.439026 | 0.859006 | 1.020396 | 1.299879 | 1.497234 |
| Rgs2 | 0.29365 | 0.307516 | 0.292093 | 0.409688 | 0.513045 | 0.433635 | 0.554241 | 0.713633 | 1.204908 | 0.929133 | 0.60718 | 0.84227 |
| Atg2a | 5.806091 | 8.143736 | 13.73034 | 12.67598 | 9.524762 | 8.785547 | 8.568979 | 7.569649 | 6.14721 | 6.08074 | 5.451315 | 5.590147 |
| Spef1 | 0.457961 | 0.43292 | 1.066529 | 0.886966 | 1.37794 | 1.308148 | 1.397144 | 1.24378 | 2.07705 | 1.890751 | 2.157288 | 2.01029 |
| Ttc30b | 1.330707 | 0.727632 | 1.198499 | 1.097992 | 1.199425 | 1.344278 | 1.866851 | 2.208935 | 2.29579 | 2.300941 | 2.81892 | 2.488144 |

| | | | | | | | | | | | | |
|---|---|---|---|---|---|---|---|---|---|---|---|---|
| P4ha3 | 0.125988 | 0.12376 | 0.487844 | 0.399497 | 0.603708 | 0.393614 | 0.549931 | 0.526597 | 0.69641 | 0.600539 | 1.133405 | 0.891797 |
| Pea15a | 12.90552 | 11.8063 | 12.76728 | 14.83874 | 19.90934 | 19.24505 | 28.37707 | 28.35262 | 31.96483 | 29.91408 | 33.3137 | 33.59797 |
| Hist1h3g | 11.27843 | 10.78773 | 4.232922 | 7.566857 | 6.661681 | 10.45703 | 8.48407 | 8.341683 | 6.458062 | 7.89719 | 4.811657 | 4.619109 |
| Smc1b | 4.601963 | 5.210136 | 4.736356 | 4.401631 | 2.484422 | 2.629377 | 1.894312 | 2.205237 | 1.876753 | 1.944932 | 1.958447 | 2.043284 |
| Cobll1 | 0.644235 | 0.783754 | 1.358712 | 1.372039 | 2.016491 | 2.054926 | 2.641678 | 2.238844 | 2.703794 | 2.5983 | 3.313409 | 3.104032 |
| Efcab12 | 1.193212 | 1.451103 | 1.549067 | 1.256806 | 1.365284 | 1.202643 | 0.877503 | 1.139067 | 0.655222 | 0.812786 | 0.490038 | 0.555788 |
| Dppa3 | 13.85822 | 12.67796 | 4.48035 | 4.896357 | 3.651519 | 4.362006 | 3.239015 | 3.432044 | 1.733508 | 2.165639 | 2.013334 | 2.584568 |
| Bag2 | 6.048566 | 5.523335 | 6.048223 | 6.555349 | 7.340431 | 7.582537 | 9.957769 | 9.952957 | 14.50023 | 13.52842 | 16.11646 | 14.81725 |
| Mep1b | 5.932852 | 5.622419 | 4.778473 | 4.101738 | 2.774699 | 2.576298 | 2.607933 | 2.644731 | 2.662959 | 2.680264 | 1.435793 | 1.815559 |
| Apobec1 | 4.947763 | 4.841014 | 5.128617 | 4.859133 | 4.669968 | 4.381405 | 3.52837 | 4.076299 | 2.382926 | 2.330574 | 1.46329 | 2.151232 |
| Zfp282 | 1.791948 | 1.818398 | 4.047926 | 3.631969 | 4.615337 | 4.482802 | 5.493632 | 5.079441 | 6.023201 | 6.200625 | 8.432574 | 8.201953 |
| Gatsl2 | 1.484295 | 2.115904 | 2.459528 | 2.770457 | 3.814305 | 3.367043 | 4.217169 | 4.122421 | 4.339465 | 4.889035 | 2.221234 | 2.165285 |
| Aldh3b1 | 2.311227 | 1.930573 | 2.017752 | 2.211173 | 1.965917 | 2.05263 | 1.824288 | 1.449755 | 1.444459 | 1.492805 | 0.931004 | 0.979691 |
| Sema4g | 3.113211 | 3.728513 | 7.725502 | 7.739726 | 6.891398 | 6.612203 | 5.414689 | 5.508324 | 5.014693 | 5.081908 | 3.452751 | 3.429237 |
| Slit2 | 0.80676 | 1.022139 | 1.385403 | 1.613743 | 1.812275 | 1.545427 | 1.995936 | 1.907779 | 3.200641 | 3.484671 | 2.807957 | 2.927466 |
| Rela | 6.584106 | 7.099996 | 11.03934 | 10.56192 | 9.766041 | 9.180022 | 10.9586 | 10.19825 | 11.69351 | 10.94326 | 20.6471 | 20.69636 |
| 4933438K2 | 1.477059 | 1.917867 | 1.397449 | 1.466408 | 1.13048 | 1.206072 | 1.046643 | 0.913535 | 0.66197 | 1.020985 | 0.401322 | 0.650591 |
| Zkscan16 | 0.786256 | 0.647381 | 1.64321 | 1.63942 | 1.773229 | 1.244342 | 1.617347 | 1.794838 | 1.656159 | 1.526091 | 0.74838 | 0.796641 |
| Ypel3 | 7.992224 | 7.466879 | 9.290578 | 8.204056 | 7.167446 | 7.400482 | 7.63784 | 8.820657 | 7.006625 | 6.668364 | 3.78353 | 3.915145 |
| Crb3 | 4.626277 | 4.140963 | 2.610392 | 1.784167 | 1.569525 | 1.380754 | 1.506936 | 1.20471 | 0.780152 | 0.792252 | 1.01441 | 0.969743 |
| Rsph3a | 0.533404 | 0.658262 | 0.813043 | 0.708519 | 0.745962 | 0.678506 | 0.869383 | 0.984873 | 1.256635 | 1.526766 | 1.296017 | 1.178322 |
| Nlrc3 | 0.235454 | 0.318071 | 0.76255 | 0.699868 | 0.693397 | 0.680681 | 0.690133 | 0.65057 | 0.668854 | 0.526835 | 1.102322 | 1.185218 |
| Pet100 | 8.493082 | 5.164613 | 5.395752 | 5.207034 | 5.91258 | 8.054991 | 4.966015 | 4.572399 | 3.954374 | 3.586279 | 4.736914 | 3.947137 |
| Sirt5 | 3.626477 | 3.489264 | 3.332031 | 3.60435 | 2.636654 | 2.864927 | 2.184431 | 2.194216 | 1.806659 | 1.605245 | 1.926378 | 1.688836 |
| Snapc5 | 20.86953 | 17.93888 | 23.08247 | 20.30658 | 17.44598 | 16.92774 | 13.54747 | 11.39174 | 10.02662 | 9.044663 | 11.83122 | 10.77346 |
| Zbtb10 | 4.474939 | 5.766504 | 6.23101 | 5.87228 | 7.999415 | 7.449729 | 9.04756 | 7.604682 | 12.23502 | 13.18252 | 12.72068 | 13.13182 |
| Vash1 | 2.723686 | 3.294278 | 8.036138 | 8.264222 | 10.65123 | 10.03906 | 14.1521 | 12.74915 | 15.43899 | 15.9505 | 19.21441 | 18.54906 |
| Nat8l | 0.929737 | 1.178569 | 1.979305 | 1.91609 | 3.121985 | 2.661539 | 3.244297 | 2.871318 | 4.297154 | 4.299361 | 4.194491 | 4.147457 |
| Colec12 | 1.826703 | 1.620833 | 2.533083 | 2.477997 | 3.167129 | 3.071636 | 3.999453 | 3.992591 | 5.366717 | 5.560119 | 4.394949 | 5.29352 |
| Rras | 15.38249 | 15.396 | 6.946258 | 7.051341 | 9.654305 | 8.05152 | 9.252068 | 9.737993 | 11.08092 | 10.13472 | 18.82862 | 15.8216 |
| Ttc3 | 17.73663 | 19.18694 | 21.32558 | 21.89416 | 23.55358 | 23.44266 | 28.03343 | 29.05418 | 44.44756 | 49.11631 | 34.17144 | 37.57514 |
| Acadm | 61.67578 | 54.92139 | 50.69518 | 47.53708 | 42.98708 | 43.36485 | 38.96895 | 40.27319 | 25.75892 | 26.54342 | 22.14932 | 21.19311 |
| Tirap | 2.529978 | 3.010858 | 4.354371 | 3.916481 | 4.736625 | 4.674466 | 6.240618 | 5.709022 | 8.498674 | 8.583713 | 8.540554 | 8.784752 |
| Kalrn | 1.138361 | 1.196376 | 1.273138 | 1.144742 | 1.391766 | 1.536969 | 2.140935 | 2.04978 | 2.269434 | 2.555411 | 2.434034 | 2.567125 |
| Adcy6 | 0.797979 | 0.963193 | 2.092115 | 2.618766 | 3.833981 | 3.510695 | 5.127188 | 4.712413 | 5.865643 | 5.871608 | 5.691752 | 5.67661 |
| St14 | 13.1613 | 15.12765 | 11.3694 | 11.45684 | 8.47884 | 8.82749 | 7.3599 | 8.312113 | 5.149171 | 5.110428 | 6.285911 | 5.632373 |
| Atp2b1 | 11.96582 | 13.14344 | 15.89309 | 16.41375 | 18.335 | 18.2166 | 22.44041 | 22.77622 | 31.46546 | 31.9877 | 34.17715 | 36.77482 |
| Frmpd1 | 0.222839 | 0.360307 | 0.608399 | 0.517429 | 0.487664 | 0.511924 | 0.589857 | 0.572313 | 1.149762 | 1.079052 | 1.430539 | 1.146701 |
| Ttyh3 | 6.590172 | 8.103646 | 11.65963 | 12.05384 | 14.26187 | 13.86486 | 19.46383 | 16.57986 | 27.51104 | 26.71297 | 28.77612 | 26.98292 |
| Zfp800 | 1.939069 | 1.882481 | 3.149903 | 3.399949 | 4.077121 | 3.914201 | 4.181062 | 4.047959 | 6.22261 | 6.30341 | 7.466942 | 7.609943 |
| Wwtr1 | 9.531383 | 9.408853 | 11.587 | 10.74813 | 12.5959 | 12.37625 | 15.48637 | 14.67777 | 21.01347 | 21.78609 | 22.99792 | 24.05491 |
| Hist1h2ak | 7.473598 | 11.18218 | 3.248666 | 4.634431 | 4.621695 | 7.555611 | 5.308903 | 5.843479 | 5.713097 | 7.258688 | 2.839047 | 3.377487 |
| Col1a1 | 0.272193 | 0.356488 | 1.24687 | 1.233527 | 1.578445 | 1.241313 | 2.317666 | 1.588581 | 1.825337 | 1.714457 | 3.43866 | 2.759046 |
| Hmgcll1 | 0.571857 | 0.593651 | 0.74717 | 0.870612 | 1.099219 | 1.20396 | 1.313363 | 1.46734 | 1.757684 | 1.946713 | 1.245451 | 1.308709 |
| Eya2 | 0.312022 | 0.223818 | 0.75813 | 0.616345 | 0.835368 | 0.941981 | 0.931243 | 0.666808 | 0.995324 | 1.004812 | 1.276321 | 1.377177 |
| Fam126a | 4.669059 | 4.889802 | 7.840987 | 8.49261 | 8.421281 | 8.102251 | 9.416296 | 8.51376 | 11.87156 | 11.43205 | 17.46415 | 18.09519 |
| Armcx3 | 3.155778 | 2.97611 | 2.620301 | 3.093029 | 4.740419 | 4.706776 | 6.774688 | 6.906446 | 6.334475 | 6.817384 | 4.756188 | 5.879117 |
| Slc27a4 | 6.575375 | 6.757224 | 9.328576 | 8.903921 | 10.10875 | 10.3136 | 13.59063 | 13.22544 | 15.1663 | 14.73425 | 20.17145 | 19.88073 |
| Wdr45 | 9.495103 | 9.300747 | 10.84497 | 9.852754 | 8.123305 | 8.929383 | 8.937137 | 9.344126 | 9.848802 | 9.671102 | 19.50483 | 19.93501 |
| Thnsl2 | 1.776909 | 1.685053 | 4.789285 | 5.314189 | 6.003108 | 5.625979 | 5.927297 | 6.259969 | 3.94859 | 4.852779 | 3.176179 | 2.804035 |
| Nfatc2 | 0.581621 | 0.762415 | 1.455298 | 1.4273 | 1.438333 | 1.398638 | 1.276469 | 1.105097 | 0.677843 | 0.781223 | 1.865755 | 1.743703 |
| Trnp1 | 0.670474 | 0.752884 | 1.480662 | 1.108801 | 1.462638 | 1.496539 | 1.302409 | 0.963411 | 0.750273 | 0.685507 | 0.582122 | 0.670549 |
| Grasp | 11.19839 | 12.3383 | 37.45518 | 33.53948 | 33.12593 | 32.37279 | 31.20027 | 29.84779 | 20.33383 | 17.80192 | 16.61473 | 15.03021 |
| Cacng4 | 0.334242 | 0.41565 | 0.590709 | 0.585714 | 0.891677 | 0.902811 | 0.94085 | 1.044467 | 1.276761 | 1.238013 | 1.299593 | 1.306937 |
| Smarca2 | 0.399354 | 0.324579 | 0.832312 | 0.696888 | 0.878796 | 1.065934 | 1.308045 | 1.355365 | 1.543379 | 1.433419 | 1.535486 | 1.554466 |
| Gucy1a2 | 0.550739 | 0.675651 | 0.563883 | 0.573805 | 0.744392 | 0.74153 | 0.749731 | 0.835484 | 1.130565 | 1.279625 | 0.795067 | 0.80847 |
| Rwdd2a | 3.716245 | 3.99818 | 3.466911 | 3.676608 | 4.447337 | 3.693703 | 3.87382 | 4.876944 | 2.80862 | 2.993185 | 1.70458 | 2.187286 |
| Twf2 | 68.83404 | 61.86554 | 55.09938 | 51.09072 | 39.8805 | 39.98619 | 32.54742 | 35.20905 | 27.0148 | 24.12384 | 25.35846 | 22.91731 |
| Tada1 | 3.776357 | 4.005254 | 5.678377 | 5.37199 | 6.810422 | 7.276251 | 7.646424 | 8.019245 | 9.892373 | 9.088829 | 10.93077 | 11.9733 |
| AU040320 | 2.48994 | 2.84954 | 4.008106 | 3.828681 | 5.150722 | 4.690617 | 5.968989 | 6.008053 | 8.141167 | 8.531566 | 8.146816 | 8.431819 |
| Cyld | 2.706101 | 2.923199 | 3.963961 | 3.861172 | 4.062045 | 4.007484 | 4.08932 | 4.30579 | 3.427197 | 3.551131 | 7.309766 | 7.911907 |

| | | | | | | | | | | | | |
|---|---|---|---|---|---|---|---|---|---|---|---|---|
| Trim61 | 35.17336 | 28.78161 | 4.533749 | 4.454974 | 2.652894 | 2.685383 | 2.177531 | 1.999894 | 3.140869 | 2.930193 | 2.076152 | 2.555775 |
| Ankrd33b | 3.668253 | 4.059408 | 1.404506 | 1.272161 | 1.226449 | 1.314762 | 1.766894 | 1.442836 | 2.726242 | 2.56326 | 2.726599 | 2.833485 |
| Megf6 | 1.098307 | 1.412669 | 2.18059 | 2.197736 | 3.382649 | 3.109509 | 4.046408 | 3.358888 | 5.150718 | 4.891724 | 3.58427 | 3.457755 |
| Acss1 | 5.197582 | 5.755409 | 1.759707 | 1.528432 | 1.285052 | 1.696565 | 1.447139 | 1.891265 | 1.140177 | 1.319857 | 0.537059 | 0.85037 |
| Xylt1 | 0.789933 | 1.032672 | 1.805481 | 1.886278 | 1.930541 | 1.896981 | 2.931712 | 1.6156 | 2.797655 | 3.008423 | 3.362985 | 3.59292 |
| Golim4 | 4.667836 | 4.472807 | 5.819268 | 5.31953 | 7.225379 | 6.878682 | 7.847596 | 8.808862 | 11.99773 | 11.82903 | 11.13201 | 11.1677 |
| Dnmt3a | 28.5803 | 29.28307 | 25.89322 | 25.61364 | 24.22039 | 23.26713 | 24.87721 | 24.07957 | 19.30266 | 20.34929 | 11.4455 | 11.52335 |
| Cd97 | 14.03585 | 14.87158 | 12.16835 | 12.82254 | 8.97264 | 8.510625 | 6.262104 | 6.623882 | 5.755093 | 5.957438 | 6.093399 | 5.768796 |
| Ano10 | 1.97663 | 1.888207 | 1.927231 | 2.391939 | 2.782412 | 2.320357 | 3.466692 | 3.047533 | 5.291528 | 5.155794 | 4.880512 | 4.487568 |
| Pcsk4 | 1.556009 | 1.702956 | 0.911163 | 1.216551 | 0.944664 | 0.780913 | 0.742137 | 0.811551 | 1.064514 | 0.731548 | 0.463121 | 0.547643 |
| Ror2 | 2.282437 | 2.505832 | 5.463107 | 5.138934 | 6.844525 | 6.816496 | 8.617665 | 8.561918 | 11.76976 | 11.40197 | 8.96378 | 8.375919 |
| Olfm1 | 2.289791 | 2.879223 | 4.553546 | 4.364091 | 7.094412 | 6.807709 | 10.12026 | 9.655606 | 9.590025 | 9.528566 | 8.153069 | 7.807147 |
| Impa2 | 52.27174 | 52.68154 | 39.5143 | 38.49176 | 30.622 | 29.43285 | 26.61927 | 26.35392 | 25.11113 | 23.63456 | 19.89799 | 17.40409 |
| Gm20939 | 9.264371 | 8.340379 | 6.361762 | 5.855644 | 4.726775 | 4.499403 | 4.014185 | 3.458386 | 3.439901 | 3.302526 | 2.767894 | 2.648208 |
| Clcn5 | 3.158728 | 3.831146 | 8.798072 | 8.784793 | 10.47345 | 9.529652 | 12.54767 | 9.803545 | 14.76342 | 15.13683 | 17.79844 | 19.42314 |
| Spag9 | 6.152143 | 7.085701 | 7.070322 | 7.001886 | 6.966128 | 6.696569 | 8.344099 | 8.294326 | 12.30772 | 12.64875 | 13.5062 | 14.79679 |
| Pdcd4 | 9.688461 | 11.17931 | 10.02338 | 10.16811 | 13.38263 | 11.9416 | 16.8599 | 18.04519 | 22.05405 | 22.46185 | 15.09032 | 15.9336 |
| Mfhas1 | 0.688853 | 0.791981 | 1.945502 | 1.940938 | 2.646521 | 2.492335 | 2.777601 | 2.976733 | 3.418721 | 3.608644 | 4.238777 | 4.287466 |
| Pspn | 0.381817 | 0.625471 | 1.088198 | 1.016784 | 0.678655 | 1.560554 | 0.844494 | 0.791006 | 0.539393 | 0.706776 | 0.447584 | 1.122301 |
| Hist1h3i | 6.948773 | 7.907867 | 1.830657 | 4.840533 | 5.109369 | 7.61677 | 5.263136 | 5.245465 | 4.936706 | 7.273686 | 5.037572 | 3.449673 |
| Antxr1 | 0.899252 | 0.991755 | 1.89695 | 2.136605 | 3.187246 | 3.168797 | 4.122831 | 3.740106 | 4.683746 | 4.716259 | 4.504725 | 4.603738 |
| Nr4a2 | 0.248163 | 0.226571 | 0.451616 | 0.477012 | 0.530393 | 0.526996 | 0.847087 | 0.906342 | 1.055007 | 0.946115 | 0.982293 | 1.052829 |
| Orai2 | 0.492256 | 0.467663 | 0.580799 | 0.506718 | 0.859359 | 0.721611 | 0.846248 | 0.98706 | 1.084914 | 1.117407 | 0.837004 | 0.751665 |
| Rusc2 | 1.284099 | 1.455349 | 1.940501 | 1.750963 | 1.739342 | 2.010631 | 2.069315 | 2.094717 | 2.44069 | 2.206326 | 4.188725 | 3.859906 |
| Bmpr1a | 17.40289 | 18.74204 | 20.35422 | 20.51813 | 23.6357 | 22.84496 | 28.32284 | 27.67487 | 42.05446 | 45.22832 | 34.50288 | 38.73272 |
| Pnmal2 | 0.452572 | 0.483226 | 1.099894 | 0.919267 | 1.266993 | 1.056621 | 1.398945 | 1.164049 | 0.784162 | 0.781995 | 0.606809 | 0.528243 |
| 2410003L1 | 40.14841 | 40.33187 | 45.4407 | 41.48533 | 40.89173 | 42.54878 | 40.73615 | 36.13584 | 30.21698 | 26.56029 | 17.81828 | 19.32471 |
| 1700020DC | 0.571803 | 0.82874 | 1.181574 | 1.197388 | 0.945794 | 0.963938 | 0.721948 | 1.422997 | 0.69383 | 0.646506 | 0.812971 | 0.690587 |
| 1700101E0 | 2.093122 | 1.932471 | 1.133443 | 0.944421 | 0.874763 | 0.703295 | 0.654558 | 0.636435 | 0.461076 | 0.539445 | 0.648539 | 0.429334 |
| Accs | 0.730015 | 0.719132 | 1.028914 | 1.147909 | 1.49784 | 1.725253 | 1.839384 | 2.14667 | 2.585196 | 2.524653 | 1.862468 | 2.324577 |
| Zfand5 | 19.60683 | 19.65203 | 21.02165 | 19.95738 | 22.85669 | 22.6989 | 27.37804 | 26.06294 | 43.15558 | 41.90793 | 43.8812 | 43.87644 |
| Lace1 | 4.769341 | 4.598574 | 3.793486 | 3.987231 | 2.825246 | 2.866146 | 2.792744 | 3.268943 | 1.922488 | 1.945766 | 1.645973 | 1.814222 |
| Hook3 | 3.030361 | 3.560559 | 4.035719 | 3.987975 | 4.5103 | 4.460164 | 5.387223 | 4.92915 | 7.554543 | 8.761199 | 7.331172 | 7.439283 |
| Cds1 | 3.434401 | 3.696365 | 2.516006 | 2.613906 | 2.679024 | 2.418467 | 2.10417 | 2.090148 | 1.55936 | 1.927625 | 1.222509 | 1.190128 |
| Tdrkh | 1.779571 | 1.770759 | 2.493327 | 2.894596 | 3.3533 | 3.243093 | 3.187979 | 3.443454 | 5.23549 | 5.635845 | 6.721121 | 6.355984 |
| Faah | 0.695642 | 0.945919 | 1.636016 | 1.295671 | 1.469974 | 1.302504 | 1.133773 | 1.22733 | 0.562847 | 0.593568 | 0.630044 | 0.668902 |
| Vash2 | 3.875101 | 4.295255 | 5.20318 | 5.286154 | 6.790053 | 6.763282 | 8.356669 | 7.86548 | 10.97038 | 11.59109 | 8.212231 | 8.509684 |
| Ehd2 | 0.831129 | 0.456751 | 1.750281 | 1.886286 | 2.274474 | 2.356508 | 3.576696 | 3.299076 | 3.115469 | 2.963661 | 4.551228 | 4.134992 |
| Gas8 | 6.688859 | 6.978073 | 9.802839 | 9.341105 | 10.21092 | 10.83034 | 11.0914 | 10.77469 | 12.7556 | 10.78274 | 21.99878 | 20.46804 |
| Dnah8 | 1.631374 | 2.146457 | 0.833693 | 0.977907 | 1.04938 | 0.898665 | 1.042674 | 0.960931 | 0.67322 | 0.901335 | 0.470265 | 0.446339 |
| Acvr1 | 2.8958 | 3.421038 | 4.037618 | 3.442382 | 4.642002 | 4.825501 | 6.011817 | 6.091441 | 7.787381 | 7.324001 | 7.616505 | 7.539984 |
| Snrpn | 57.06888 | 56.39526 | 36.77091 | 36.23828 | 34.98145 | 33.85362 | 28.83955 | 31.23679 | 23.1405 | 22.44762 | 16.56423 | 16.55435 |
| Kmt2e | 5.495333 | 6.057863 | 6.042131 | 7.025503 | 8.248532 | 8.006701 | 9.82593 | 8.679091 | 14.93788 | 15.37525 | 11.31092 | 12.43688 |
| Rimbp3 | 0.459259 | 0.622887 | 1.451154 | 1.42873 | 0.957121 | 0.949898 | 0.906449 | 0.850362 | 0.833828 | 0.743954 | 0.827674 | 0.652998 |
| Arhgef3 | 4.833573 | 4.940442 | 2.420972 | 2.429705 | 1.904664 | 2.020967 | 1.867626 | 2.000185 | 4.038905 | 4.37584 | 4.139803 | 3.918132 |
| Armcx4 | 9.552152 | 10.46313 | 9.299837 | 9.884731 | 12.19133 | 11.78781 | 14.32873 | 14.65915 | 21.38436 | 21.60868 | 16.64982 | 17.93482 |
| Uqcrq | 72.76836 | 58.71556 | 57.49107 | 59.26426 | 60.79071 | 62.54941 | 49.35316 | 48.86996 | 34.30798 | 28.61421 | 46.23922 | 41.5413 |
| Bcar1 | 27.8281 | 31.02625 | 32.63696 | 30.26259 | 31.72779 | 31.34548 | 33.07943 | 34.37807 | 32.50551 | 31.80729 | 67.31694 | 66.14763 |
| S100a16 | 1.019765 | 1.054713 | 1.668081 | 1.430377 | 2.286337 | 2.038443 | 2.295667 | 2.661485 | 3.167003 | 2.407935 | 3.472285 | 3.12375 |
| Brsk1 | 3.923536 | 4.65985 | 8.776143 | 9.264031 | 7.812053 | 7.902081 | 6.366807 | 6.695299 | 4.649043 | 4.528243 | 4.305246 | 4.243226 |
| Ccdc85a | 1.194845 | 1.420072 | 0.562391 | 0.569803 | 0.745362 | 0.687192 | 0.948663 | 0.906103 | 1.344552 | 1.243876 | 0.756611 | 0.879724 |
| Dok3 | 0.417188 | 0.684886 | 0.838236 | 1.172666 | 1.133344 | 1.276237 | 1.199404 | 1.788589 | 0.931442 | 0.819339 | 1.990815 | 1.623208 |
| Pdha2 | 0.325758 | 0.379724 | 1.910305 | 1.900098 | 1.643649 | 1.705521 | 1.907102 | 1.873089 | 0.912588 | 0.870608 | 1.506772 | 1.562007 |
| Egln3 | 10.56232 | 8.649488 | 2.250749 | 2.927441 | 4.245156 | 3.527819 | 3.972035 | 4.383886 | 4.213742 | 5.103962 | 2.122752 | 2.33877 |
| Aldh1a2 | 0.480616 | 0.476017 | 1.17869 | 1.186797 | 1.227896 | 1.36114 | 1.18867 | 1.138556 | 1.118959 | 1.020924 | 0.546859 | 0.624045 |
| Cyp26a1 | 0.999753 | 0.880731 | 96.98327 | 91.71224 | 142.4395 | 147.0254 | 139.4969 | 140.7334 | 76.26971 | 67.42735 | 74.82755 | 73.95695 |
| Morc4 | 1.449322 | 1.851254 | 4.099096 | 4.078241 | 5.816262 | 4.991901 | 6.120125 | 5.51315 | 6.10499 | 6.128683 | 8.220859 | 8.890992 |
| Zfp618 | 0.905299 | 1.050079 | 3.281922 | 3.169834 | 4.523763 | 4.29863 | 6.306469 | 5.336667 | 6.878649 | 6.908134 | 5.035913 | 5.445947 |
| Dhtkd1 | 6.723156 | 7.195056 | 7.269813 | 6.700914 | 6.724846 | 6.742494 | 6.303942 | 5.68372 | 5.389754 | 5.378873 | 2.593962 | 3.094096 |
| Ifi27 | 0.246989 | 0.278472 | 0.593403 | 0.767483 | 0.90737 | 1.115844 | 1.209857 | 1.391142 | 1.205047 | 1.40199 | 1.849725 | 1.672222 |
| Shroom2 | 3.084996 | 3.763303 | 4.490978 | 4.42034 | 4.808696 | 4.517154 | 5.07492 | 4.409034 | 5.916241 | 6.076972 | 8.767334 | 9.605644 |

| | | | | | | | | | | | | |
|---|---|---|---|---|---|---|---|---|---|---|---|---|
| Tgfbr3 | 0.38433 | 0.383145 | 1.245178 | 1.282911 | 1.783479 | 1.775572 | 2.369241 | 1.732396 | 2.521987 | 2.352126 | 2.625041 | 2.794498 |
| Slc29a1 | 67.09456 | 63.22741 | 39.20542 | 37.36215 | 32.65052 | 32.90236 | 28.8897 | 28.86894 | 22.46468 | 22.40154 | 18.47185 | 17.15925 |
| Cbfb | 27.64762 | 29.29432 | 37.29085 | 35.57695 | 38.73147 | 37.57938 | 44.27229 | 38.52758 | 53.32278 | 53.97059 | 77.96527 | 77.45203 |
| Spint2 | 63.48931 | 68.29409 | 58.7977 | 53.07514 | 46.45601 | 45.13854 | 37.996 | 37.39088 | 27.70298 | 24.37998 | 32.87863 | 28.41324 |
| Atp2a3 | 4.844032 | 4.780218 | 3.486838 | 3.40038 | 3.271139 | 3.33103 | 3.274711 | 2.869414 | 1.939931 | 2.027457 | 1.682898 | 1.562446 |
| Zfp560 | 1.247368 | 1.150373 | 1.761312 | 1.873807 | 2.178303 | 2.200353 | 3.096628 | 2.979133 | 3.377468 | 3.467712 | 4.157626 | 4.077538 |
| Il13ra1 | 0.559756 | 0.598644 | 1.151844 | 0.996721 | 1.475788 | 1.303645 | 1.695853 | 1.81812 | 1.766235 | 2.062333 | 1.670605 | 2.168534 |
| Bcat2 | 38.74576 | 34.66672 | 21.5796 | 22.34018 | 17.64353 | 17.28671 | 16.60521 | 18.21983 | 11.89805 | 11.45739 | 9.593793 | 10.27012 |
| Otx2 | 24.46254 | 24.55604 | 2.130494 | 1.712522 | 1.507957 | 1.359326 | 1.214572 | 0.882349 | 1.843691 | 1.918947 | 1.693337 | 1.635981 |
| Mtus1 | 6.372011 | 7.625503 | 5.426569 | 5.340481 | 4.023459 | 4.245355 | 3.895566 | 3.593048 | 2.625605 | 2.857412 | 2.56628 | 2.455629 |
| Rnf19a | 6.672131 | 6.793893 | 6.892276 | 6.805568 | 8.726853 | 7.940964 | 9.154163 | 9.294502 | 10.51415 | 11.23839 | 14.26417 | 14.796 |
| Rock2 | 13.39535 | 15.62428 | 25.9324 | 26.05901 | 25.97491 | 24.21319 | 25.04639 | 25.31127 | 26.49112 | 26.96438 | 47.9714 | 52.63229 |
| Acsl4 | 15.07501 | 16.39997 | 20.06135 | 20.30061 | 26.78214 | 24.87033 | 32.92814 | 31.63529 | 43.1853 | 44.11956 | 29.31676 | 31.88278 |
| Nap1l2 | 0.515636 | 0.514708 | 0.856037 | 1.024998 | 1.079152 | 1.140972 | 1.079589 | 0.893612 | 0.565351 | 0.525119 | 0.701732 | 0.757457 |
| H2-DMa | 0.596595 | 0.915105 | 1.432378 | 1.098303 | 1.400182 | 1.125193 | 1.147174 | 1.356282 | 0.683022 | 0.786565 | 0.599263 | 0.62454 |
| Pcmtd1 | 26.0941 | 25.82562 | 18.36814 | 17.61163 | 15.42944 | 14.37026 | 12.21238 | 13.09627 | 11.8597 | 12.71828 | 7.370693 | 8.11298 |
| Pdgfrl | 1.373366 | 1.128995 | 1.069327 | 1.412544 | 1.967197 | 1.969445 | 3.390435 | 2.808854 | 3.206906 | 3.065015 | 3.178174 | 2.240069 |
| Tanc2 | 0.445606 | 0.544084 | 1.267176 | 1.381509 | 1.433687 | 1.353987 | 1.537609 | 1.367863 | 2.205567 | 2.081194 | 2.791529 | 2.937568 |
| Thy1 | 2.171808 | 2.186126 | 0.350122 | 0.599501 | 0.557041 | 0.475036 | 0.588447 | 0.540255 | 0.8603 | 1.030574 | 0.778526 | 0.702862 |
| Rassf5 | 0.549789 | 0.764847 | 1.107001 | 1.33245 | 1.184978 | 1.211097 | 1.227842 | 0.954824 | 0.784669 | 0.614301 | 0.680417 | 0.807141 |
| Sycp2 | 0.440763 | 0.351169 | 1.358571 | 1.310703 | 1.553047 | 1.450667 | 1.39298 | 1.516602 | 1.096128 | 0.907424 | 1.88582 | 1.968146 |
| Smug1 | 5.896835 | 5.407042 | 6.096768 | 5.931478 | 7.986928 | 7.594194 | 10.58565 | 9.556258 | 12.87319 | 12.86076 | 11.95358 | 12.71107 |
| 2500004C0 | 0.911939 | 0.781501 | 1.936032 | 1.720068 | 1.835965 | 2.210159 | 2.581025 | 2.469288 | 2.852756 | 2.072096 | 3.673996 | 3.728109 |
| Synpo2 | 0.210009 | 0.235229 | 0.595325 | 0.494438 | 0.548851 | 0.713491 | 0.709124 | 0.685153 | 0.661109 | 0.618582 | 0.985168 | 1.071553 |
| Arhgef28 | 1.578 | 1.854903 | 0.774728 | 0.83928 | 1.042557 | 1.154122 | 1.55494 | 1.804596 | 1.78552 | 1.744093 | 1.550375 | 1.818847 |
| Prkrir | 16.02523 | 15.95787 | 19.09531 | 17.47735 | 18.12232 | 17.76637 | 20.9135 | 20.56965 | 26.78936 | 26.83284 | 34.59742 | 37.85497 |
| Gpr125 | 17.02003 | 18.13495 | 21.97021 | 22.1635 | 27.94446 | 26.58721 | 32.76426 | 33.36727 | 47.0783 | 47.96485 | 39.45459 | 39.34177 |
| Pdgfrb | 0.502338 | 0.585065 | 1.679442 | 1.765463 | 1.989942 | 2.025868 | 2.086427 | 1.832164 | 1.48126 | 1.745772 | 1.1715 | 0.936448 |
| Manea | 4.445291 | 4.718077 | 6.056222 | 5.75993 | 7.363089 | 7.262743 | 8.835816 | 7.73254 | 11.53159 | 12.45988 | 8.934877 | 10.01083 |
| Dapk3 | 14.87876 | 14.14733 | 22.06057 | 20.62828 | 18.91418 | 17.47021 | 17.47754 | 18.41595 | 19.36695 | 17.94943 | 42.56262 | 37.77865 |
| Arhgef17 | 3.606778 | 4.646726 | 5.446488 | 5.470945 | 7.136443 | 6.929443 | 8.60142 | 8.010843 | 9.125099 | 9.587429 | 12.24883 | 11.82495 |
| Kctd15 | 7.509625 | 8.25866 | 11.74779 | 10.81226 | 12.00115 | 11.53719 | 12.02204 | 12.05694 | 16.88018 | 15.89887 | 24.16783 | 23.35355 |
| Pknox2 | 3.15838 | 3.345935 | 8.745312 | 9.544409 | 15.41072 | 13.74473 | 21.37864 | 20.11257 | 22.1489 | 20.61431 | 10.11865 | 10.10782 |
| Them6 | 9.658666 | 10.00125 | 9.240179 | 8.595144 | 8.345126 | 7.905086 | 6.512575 | 6.541427 | 4.593901 | 3.980035 | 4.675254 | 4.316538 |
| Dock11 | 3.523601 | 4.102017 | 4.935734 | 5.06819 | 6.015232 | 5.751103 | 7.280922 | 7.495922 | 10.6666 | 10.94507 | 9.376923 | 10.29032 |
| Slc39a4 | 11.86149 | 12.91393 | 10.3083 | 9.498037 | 7.486953 | 8.111754 | 7.124473 | 7.207079 | 6.367888 | 6.116115 | 4.833909 | 4.398139 |
| Zyg11a | 14.05732 | 16.39844 | 7.399956 | 6.604258 | 4.673715 | 4.226607 | 3.954471 | 4.381633 | 3.094364 | 3.693402 | 3.811402 | 3.059599 |
| Stmn3 | 6.452612 | 6.540744 | 4.540543 | 4.033891 | 3.173517 | 2.858679 | 1.934902 | 2.929193 | 1.880467 | 1.86889 | 2.494292 | 2.72803 |
| Msh6 | 173.8864 | 191.3982 | 193.8205 | 184.9857 | 150.7562 | 143.025 | 134.4731 | 129.3858 | 120.3908 | 122.4474 | 87.07212 | 85.79834 |
| H2-Bl | 2.5288 | 2.596448 | 44.47608 | 36.37076 | 43.45963 | 43.46995 | 42.67645 | 50.37004 | 31.76191 | 29.17121 | 29.79051 | 23.36764 |
| Tnrc18 | 5.782478 | 7.206801 | 9.319538 | 9.003655 | 11.37539 | 11.06802 | 14.85602 | 13.77127 | 17.50394 | 16.8626 | 21.80121 | 19.40381 |
| Slco4c1 | 2.121065 | 2.179217 | 0.928484 | 0.844733 | 0.858241 | 0.77891 | 0.849256 | 0.534751 | 0.786773 | 0.787562 | 0.492766 | 0.392126 |
| Mup6 | 9.288863 | 7.378753 | 2.808148 | 3.267465 | 2.67396 | 2.339828 | 2.535639 | 3.542869 | 1.743906 | 1.680434 | 1.224154 | 1.644756 |
| Fmn2 | 0.811577 | 1.072441 | 1.037875 | 1.094593 | 0.792458 | 0.884289 | 0.975017 | 0.947546 | 1.861243 | 1.904512 | 1.815909 | 1.579047 |
| Cxcl14 | 0.784759 | 0.680558 | 1.224149 | 1.170933 | 2.034543 | 1.921274 | 2.027452 | 2.51946 | 2.674595 | 2.372645 | 2.189072 | 2.071816 |
| Notch2 | 3.143139 | 4.553177 | 7.09323 | 7.210481 | 7.579744 | 7.441766 | 9.460325 | 7.186644 | 14.50565 | 15.03428 | 15.20579 | 15.46306 |
| Lhfpl4 | 0.779198 | 0.873058 | 1.034384 | 1.197263 | 1.231199 | 1.308706 | 1.356656 | 1.210553 | 1.497187 | 1.463952 | 2.283929 | 2.575676 |
| Sptbn4 | 3.43167 | 3.529108 | 3.782091 | 3.581883 | 2.889155 | 3.027815 | 2.265522 | 2.432566 | 1.730659 | 1.666326 | 1.564471 | 1.698033 |
| Sidt2 | 8.950377 | 10.82809 | 13.11007 | 12.47585 | 13.53367 | 13.32633 | 16.3758 | 15.1541 | 17.25071 | 16.85993 | 27.9414 | 26.81346 |
| Tcerg1l | 0.856452 | 0.726888 | 1.502394 | 1.452701 | 1.469039 | 1.662261 | 2.029004 | 2.192612 | 1.82492 | 1.550264 | 1.035106 | 1.020283 |
| Csad | 22.68992 | 21.60513 | 15.99279 | 14.95526 | 16.2795 | 15.04412 | 14.41872 | 16.20174 | 10.77363 | 11.21449 | 6.7922 | 7.539156 |
| Vezf1 | 11.29189 | 12.6098 | 20.73953 | 22.10252 | 24.95733 | 24.52676 | 30.79522 | 29.7176 | 47.01678 | 47.49659 | 43.30636 | 45.35876 |
| Gm13212 | 3.581283 | 3.112681 | 2.986963 | 2.651634 | 2.361581 | 2.02438 | 1.644339 | 1.79907 | 1.519318 | 1.452971 | 1.22217 | 1.234034 |
| 3632451O( | 5.879089 | 5.665759 | 2.501114 | 2.541242 | 3.219564 | 3.344881 | 3.520143 | 3.737152 | 4.862688 | 5.458423 | 3.261605 | 3.593864 |
| Myl12a | 73.65068 | 71.80898 | 93.87865 | 89.35556 | 88.61962 | 87.10034 | 92.60375 | 80.18458 | 88.13334 | 74.14321 | 174.8127 | 159.2216 |
| Adrbk2 | 1.637626 | 1.905755 | 2.631284 | 2.661431 | 3.444514 | 3.402975 | 3.363389 | 3.22628 | 2.089103 | 2.280012 | 1.757755 | 1.585447 |
| Maml1 | 6.934803 | 8.140274 | 11.05199 | 10.31963 | 11.56975 | 11.84111 | 13.09886 | 11.23352 | 16.06706 | 15.62484 | 23.48349 | 22.14787 |
| Twist1 | 1.44611 | 1.191641 | 2.809372 | 2.611856 | 3.876119 | 3.875603 | 3.704381 | 4.08927 | 4.904601 | 4.796194 | 6.684232 | 5.60435 |
| Rnf24 | 1.393889 | 1.49753 | 2.483702 | 2.459797 | 2.886734 | 2.98374 | 3.889863 | 3.410369 | 5.348878 | 5.276186 | 4.768206 | 4.981165 |
| Cd63 | 102.9245 | 88.13181 | 43.29445 | 42.90273 | 49.30776 | 45.05041 | 47.06321 | 52.71298 | 67.11989 | 61.41783 | 94.87515 | 92.0168 |
| Commd3 | 30.39121 | 22.87815 | 29.39924 | 28.55595 | 45.08709 | 44.01589 | 53.83507 | 51.56379 | 63.32903 | 55.89929 | 57.24015 | 61.22829 |

| | | | | | | | | | | | | |
|---|---|---|---|---|---|---|---|---|---|---|---|---|
| F2r | 17.56127 | 18.19019 | 20.83741 | 20.41424 | 25.31322 | 25.9542 | 38.23017 | 32.89341 | 44.34418 | 43.77009 | 36.14263 | 38.84932 |
| Sobp | 0.307648 | 0.394819 | 0.677525 | 0.680991 | 1.055146 | 0.971229 | 1.410021 | 1.025806 | 1.317562 | 1.211245 | 1.324069 | 1.458704 |
| Dkk3 | 0.989871 | 0.697325 | 0.888257 | 1.192946 | 2.057 | 1.800036 | 2.531053 | 2.554864 | 2.42839 | 2.538518 | 2.001226 | 1.957662 |
| Gm13034 | 1.179489 | 1.265886 | 3.787874 | 3.13528 | 2.326285 | 2.366202 | 2.801961 | 3.186431 | 1.710021 | 1.488744 | 2.939716 | 2.837152 |
| Fads1 | 21.7973 | 23.96757 | 30.58266 | 29.8699 | 40.67336 | 38.15595 | 49.01088 | 44.53458 | 63.07492 | 63.91967 | 54.28646 | 53.22752 |
| D10Bwg13 | 0.582303 | 0.742205 | 2.204854 | 2.431753 | 1.957759 | 1.775393 | 1.879964 | 1.517362 | 1.076763 | 1.136822 | 1.506361 | 1.478823 |
| Lphn1 | 21.46958 | 25.8615 | 23.31082 | 22.73298 | 20.65303 | 20.51467 | 18.13662 | 17.04883 | 16.18382 | 16.64651 | 10.20883 | 10.63374 |
| Akr1b8 | 25.76812 | 22.05543 | 15.642 | 14.27487 | 14.36083 | 14.06098 | 15.49084 | 16.70373 | 10.41208 | 8.098377 | 6.272435 | 7.814207 |
| Acan | 1.14056 | 1.14618 | 3.035589 | 2.849727 | 2.828293 | 3.004515 | 2.876147 | 2.759137 | 1.615572 | 1.653433 | 1.459565 | 1.405619 |
| Ttc12 | 3.413688 | 3.193733 | 2.179603 | 1.989019 | 2.420451 | 2.274334 | 2.439046 | 3.126195 | 1.656478 | 1.911805 | 1.271343 | 1.463407 |
| Mlkl | 0.442755 | 0.574346 | 0.969419 | 0.756264 | 1.175446 | 1.087765 | 1.337623 | 1.614493 | 1.021157 | 1.034169 | 0.782408 | 0.989674 |
| Mir22hg | 1.185015 | 1.242078 | 3.240937 | 2.379837 | 4.046214 | 3.520516 | 3.540942 | 3.825339 | 2.643443 | 2.129356 | 4.053087 | 4.54489 |
| Lrrc34 | 21.81217 | 18.36487 | 2.561995 | 2.721608 | 1.972087 | 1.827999 | 1.658013 | 1.487057 | 2.055793 | 2.164657 | 1.294507 | 1.275187 |
| Rfx3 | 1.248321 | 1.631142 | 2.137726 | 2.022169 | 2.441563 | 2.293537 | 3.057594 | 2.562163 | 4.074097 | 4.314427 | 3.65231 | 3.834997 |
| Npnt | 1.042867 | 0.830727 | 1.578156 | 1.668318 | 2.567953 | 2.203025 | 2.806509 | 2.983097 | 3.757548 | 3.558521 | 2.874876 | 2.75704 |
| Llgl2 | 38.79501 | 40.90595 | 89.08232 | 81.96147 | 86.76838 | 84.76481 | 84.42991 | 88.24559 | 50.81083 | 50.10354 | 46.04523 | 41.38057 |
| Gabbr1 | 20.11176 | 22.9849 | 18.26623 | 17.66522 | 14.36455 | 15.83533 | 13.39774 | 15.94981 | 11.41578 | 12.34168 | 8.082535 | 8.28782 |
| Nhs | 1.763955 | 2.09147 | 2.273536 | 2.333911 | 2.443165 | 2.183374 | 2.375463 | 2.059916 | 4.294614 | 4.39054 | 3.079351 | 3.351695 |
| Cpeb1 | 2.287134 | 2.26066 | 5.378267 | 5.38058 | 6.048382 | 5.824599 | 5.33412 | 5.574035 | 2.818281 | 2.732839 | 3.020881 | 3.250483 |
| Cxx1c | 0.402797 | 0.476437 | 0.67504 | 0.877653 | 1.856826 | 1.487587 | 2.394676 | 1.870009 | 2.540361 | 1.196347 | 1.828564 | 1.683411 |
| Scd2 | 88.35567 | 101.2015 | 127.8537 | 139.2805 | 214.5753 | 197.2301 | 253.5955 | 230.6483 | 282.1027 | 296.6974 | 203.9688 | 197.4196 |
| Noxred1 | 4.189524 | 4.038307 | 3.869876 | 4.809054 | 3.823445 | 2.856125 | 3.072534 | 3.150013 | 1.960587 | 2.257758 | 2.346784 | 2.577611 |
| Micall1 | 1.293944 | 1.410348 | 1.598342 | 1.634322 | 2.383191 | 2.273969 | 2.485471 | 2.2311 | 2.923049 | 2.962829 | 3.58067 | 3.480978 |
| B3gnt3 | 0.345293 | 0.493637 | 1.236493 | 1.181189 | 0.632167 | 0.800011 | 0.695058 | 1.002862 | 0.442542 | 0.555178 | 0.573996 | 0.556034 |
| Sh3pxd2a | 2.963642 | 4.025614 | 3.005275 | 3.065698 | 3.849331 | 3.671065 | 4.778052 | 4.228071 | 6.163339 | 6.520463 | 5.710833 | 5.324049 |
| AI661453 | 0.452145 | 0.608896 | 1.106456 | 1.063072 | 0.753193 | 0.687638 | 0.631157 | 0.529735 | 0.494057 | 0.504327 | 1.017746 | 1.072638 |
| Zmiz1 | 5.495724 | 7.162673 | 21.95219 | 23.95745 | 33.37462 | 32.09557 | 46.09188 | 33.96329 | 53.04273 | 50.91076 | 40.90903 | 39.22915 |
| Alox12 | 0.409829 | 0.407434 | 0.58595 | 0.608685 | 0.967629 | 0.874616 | 1.016191 | 1.186904 | 0.571105 | 0.605034 | 0.54724 | 0.558563 |
| Tead2 | 59.91043 | 53.7097 | 39.17861 | 39.58256 | 53.06629 | 50.48389 | 58.81727 | 68.13393 | 86.30538 | 84.09216 | 77.6367 | 75.18083 |
| Tor1aip1 | 3.530966 | 3.665236 | 4.363946 | 4.759861 | 4.78566 | 5.150009 | 5.36574 | 6.264197 | 8.53356 | 8.242244 | 9.891268 | 10.1096 |
| Pmm1 | 139.1112 | 128.6003 | 114.571 | 104.0978 | 102.6011 | 97.90146 | 83.30932 | 92.29212 | 64.86346 | 59.88027 | 52.47329 | 49.05085 |
| Fignl2 | 6.3123 | 7.49352 | 19.45149 | 17.99918 | 31.97569 | 32.65762 | 44.78833 | 38.1979 | 40.59684 | 36.1147 | 34.41145 | 31.76925 |
| Galnt4 | 0.753276 | 0.740886 | 1.410626 | 1.242511 | 1.376688 | 1.362744 | 1.768682 | 1.759012 | 1.909912 | 1.903954 | 2.605111 | 2.636138 |
| Stag3 | 21.92654 | 23.51618 | 19.93797 | 19.41995 | 15.38615 | 14.61224 | 12.10634 | 13.48094 | 8.80682 | 9.52792 | 10.38433 | 9.155509 |
| Arhgef26 | 1.270306 | 1.363859 | 3.059919 | 2.806489 | 3.15486 | 3.220077 | 4.57243 | 4.646808 | 6.132057 | 5.952423 | 5.205514 | 4.77139 |
| Lonrf1 | 4.847838 | 5.790473 | 6.035802 | 5.709594 | 6.415505 | 6.328926 | 6.703023 | 6.594784 | 8.686836 | 9.370538 | 12.00277 | 12.10317 |
| Isoc2a | 37.46942 | 34.34516 | 38.9495 | 33.22176 | 29.3573 | 29.05494 | 22.75743 | 24.85507 | 19.23853 | 16.70199 | 16.42094 | 15.67246 |
| Zfpm1 | 1.68378 | 2.10422 | 3.646071 | 3.146497 | 3.945023 | 4.149824 | 4.952969 | 4.520808 | 5.565819 | 5.116346 | 7.99368 | 6.666871 |
| Epb4.1l4b | 1.491406 | 1.620755 | 1.473614 | 1.502551 | 1.811729 | 1.74778 | 1.93774 | 2.02169 | 2.030964 | 2.079804 | 3.02565 | 3.182877 |
| Col22a1 | 0.272072 | 0.322552 | 0.60321 | 0.640891 | 0.93523 | 0.913867 | 1.007137 | 1.143878 | 1.248826 | 1.357565 | 1.338809 | 1.174438 |
| Ube2h | 8.597818 | 9.612236 | 12.66732 | 12.09323 | 16.32768 | 16.22545 | 21.8807 | 19.53382 | 23.67738 | 25.61128 | 20.7728 | 20.91022 |
| Fam53b | 1.586591 | 1.735982 | 1.669266 | 1.819522 | 2.406727 | 2.320369 | 3.1844 | 2.324059 | 4.00728 | 3.851713 | 3.300259 | 3.037325 |
| Purb | 12.92451 | 14.98811 | 21.62718 | 20.71502 | 18.74106 | 17.90952 | 19.17925 | 15.2111 | 21.06053 | 22.0329 | 30.62549 | 32.19652 |
| Fign | 0.38325 | 0.504819 | 0.856825 | 0.998542 | 1.048947 | 1.145892 | 1.511712 | 1.198881 | 2.052274 | 2.113128 | 1.645189 | 1.856302 |
| Crtc3 | 0.982559 | 0.77393 | 1.712234 | 1.934658 | 2.46332 | 2.203451 | 2.751714 | 2.18988 | 3.220734 | 3.30682 | 4.179802 | 4.09288 |
| Casp7 | 2.417819 | 2.739841 | 4.720896 | 4.41332 | 3.770361 | 3.92283 | 4.045027 | 3.330555 | 2.955132 | 3.162343 | 7.42641 | 6.687427 |
| Ank | 3.174811 | 3.833446 | 4.329889 | 4.785952 | 6.920296 | 5.765303 | 7.795423 | 7.891571 | 9.816845 | 10.11914 | 8.902892 | 9.042098 |
| Bcl2l1 | 14.45835 | 14.74985 | 16.52691 | 14.36441 | 14.34598 | 14.62397 | 15.43464 | 17.50216 | 18.26569 | 16.26854 | 30.47711 | 30.36964 |
| Chrnb1 | 1.412217 | 1.238599 | 2.218476 | 2.210435 | 2.046982 | 2.122106 | 1.971014 | 1.848586 | 1.799302 | 1.716426 | 3.351553 | 3.628334 |
| Mob1b | 3.90748 | 4.609299 | 6.583177 | 6.479062 | 6.745551 | 6.529517 | 7.707835 | 6.432603 | 9.749119 | 9.272836 | 12.06959 | 13.58344 |
| St5 | 4.315903 | 4.750123 | 4.539225 | 4.759271 | 4.681949 | 4.587613 | 5.60616 | 4.943965 | 7.606257 | 6.932193 | 9.373772 | 9.685946 |
| Foxa1 | 0.164641 | 0.145269 | 1.486251 | 1.387587 | 2.179507 | 1.969688 | 2.625168 | 2.293371 | 3.294468 | 2.929295 | 1.749804 | 1.513512 |
| Frmd6 | 3.487479 | 3.874646 | 5.353288 | 4.997485 | 5.850707 | 5.500163 | 6.484493 | 7.030063 | 7.540624 | 7.736673 | 10.55725 | 10.54781 |
| Spata33 | 0.737948 | 0.697659 | 1.361953 | 0.967203 | 1.940799 | 1.501985 | 1.214299 | 1.266678 | 1.139318 | 0.711821 | 0.995517 | 0.87221 |
| Dmrtb1 | 1.201204 | 1.149545 | 2.940022 | 3.278462 | 4.688622 | 5.043942 | 5.240182 | 5.893014 | 3.06696 | 2.792885 | 3.590608 | 3.771375 |
| Aard | 68.01441 | 59.7563 | 147.1225 | 134.0751 | 174.6062 | 168.4864 | 157.5486 | 189.3458 | 95.83279 | 93.55121 | 88.86812 | 89.7449 |
| Vldlr | 0.412352 | 0.391226 | 0.805136 | 0.867269 | 1.012472 | 0.958058 | 0.80549 | 0.746722 | 0.751411 | 0.794047 | 1.361784 | 1.574584 |
| Ptpru | 1.488526 | 1.572983 | 4.894443 | 4.583766 | 4.953149 | 4.925316 | 4.711697 | 4.760206 | 3.304876 | 3.647479 | 2.289054 | 2.33594 |
| Adcyap1r1 | 0.422676 | 0.377742 | 0.525267 | 0.597809 | 0.913565 | 0.712257 | 0.938762 | 0.713712 | 1.291047 | 1.087677 | 1.139247 | 1.260316 |
| Rai1 | 4.434662 | 5.356922 | 6.204899 | 6.481976 | 6.641543 | 5.979023 | 6.696818 | 6.30805 | 11.53127 | 10.59363 | 12.87715 | 12.59999 |
| Dpp7 | 8.147855 | 7.107011 | 8.897319 | 8.324372 | 6.28783 | 6.215121 | 5.996578 | 5.809841 | 4.707937 | 3.953428 | 3.641502 | 4.106411 |

| | | | | | | | | | | | | |
|---|---|---|---|---|---|---|---|---|---|---|---|---|
| Parp12 | 5.999963 | 5.850227 | 6.60738 | 6.213548 | 4.927943 | 4.248346 | 4.079736 | 3.829437 | 2.631942 | 2.951612 | 5.601444 | 5.161717 |
| Sike1 | 10.99992 | 10.38351 | 10.65412 | 11.00495 | 11.90347 | 12.49045 | 12.17356 | 11.90858 | 17.06354 | 15.32331 | 22.64852 | 23.16081 |
| Uqcrb | 103.4205 | 73.87518 | 65.28939 | 72.86927 | 84.65333 | 97.74988 | 72.82358 | 67.10704 | 57.83669 | 49.68667 | 50.65426 | 46.46967 |
| Dopey2 | 3.089616 | 4.180966 | 7.614058 | 7.235518 | 6.618186 | 6.625385 | 7.082063 | 6.709947 | 4.050919 | 4.136016 | 3.562919 | 3.439762 |
| 3830417A1 | 0.868072 | 0.771087 | 1.62863 | 1.67835 | 1.686949 | 1.310135 | 2.110273 | 2.074771 | 2.157794 | 2.75405 | 3.424896 | 2.524447 |
| Calca | 17.35684 | 13.3848 | 7.167795 | 8.383036 | 8.833032 | 8.096522 | 7.001019 | 7.606453 | 4.051159 | 3.988705 | 4.045887 | 4.497908 |
| Rfx2 | 10.14244 | 11.45359 | 6.844988 | 6.511436 | 5.819302 | 4.895006 | 4.620325 | 4.500806 | 4.578389 | 4.738347 | 3.346438 | 3.104835 |
| Aldoc | 8.115545 | 7.622411 | 1.98225 | 2.001174 | 2.515285 | 2.167834 | 2.164445 | 1.985038 | 3.063898 | 3.800277 | 2.224901 | 1.812214 |
| Radil | 5.222337 | 5.443841 | 3.872403 | 4.00655 | 4.603689 | 3.862901 | 4.01475 | 4.245593 | 2.569842 | 2.840066 | 2.032247 | 2.025621 |
| Ina | 22.03538 | 23.91432 | 19.202 | 19.78232 | 17.55405 | 17.41878 | 16.85983 | 14.95697 | 10.37113 | 9.443102 | 9.946167 | 10.29807 |
| Mpp3 | 1.355961 | 1.512575 | 1.440443 | 1.500099 | 1.778222 | 1.820475 | 1.871748 | 1.694539 | 3.116682 | 3.140568 | 3.235164 | 3.029438 |
| Tcf7l2 | 2.057417 | 2.732375 | 3.730271 | 3.471346 | 3.926013 | 3.912707 | 5.128479 | 4.293689 | 6.934729 | 6.841065 | 6.801493 | 7.265651 |
| Gstm6 | 1.213651 | 1.36586 | 1.433609 | 1.485881 | 1.946996 | 1.402101 | 1.552059 | 1.590062 | 0.638521 | 1.350048 | 0.304066 | 0.759798 |
| Hhat | 0.329736 | 0.353145 | 0.872011 | 0.722792 | 1.179113 | 1.184834 | 1.405456 | 1.039566 | 1.173038 | 1.255533 | 1.782226 | 1.512141 |
| Hfe2 | 0.498649 | 0.589525 | 1.124251 | 1.415595 | 1.513869 | 1.275885 | 1.361931 | 1.414593 | 1.377532 | 1.020071 | 0.887122 | 0.67674 |
| C1rl | 0.528857 | 0.837677 | 0.999867 | 0.944322 | 1.190785 | 0.852642 | 0.831271 | 1.105503 | 0.844026 | 0.809352 | 0.489781 | 0.528549 |
| Dhodh | 19.40795 | 18.35265 | 13.32047 | 12.75114 | 11.36784 | 11.65371 | 9.757067 | 10.19332 | 8.370746 | 7.86813 | 5.908662 | 6.096457 |
| Prrt4 | 0.396593 | 0.415824 | 1.552801 | 1.193693 | 1.729881 | 1.999633 | 1.961674 | 1.552618 | 1.360161 | 1.211633 | 0.962247 | 0.956328 |
| Syk | 12.59299 | 13.64789 | 8.784068 | 8.544235 | 6.707403 | 6.621809 | 6.687353 | 6.335752 | 4.111957 | 4.374731 | 3.998929 | 4.087052 |
| Cacfd1 | 4.275333 | 4.501047 | 6.515582 | 6.584178 | 8.446768 | 8.289849 | 10.3593 | 9.760346 | 13.4421 | 11.8294 | 13.86478 | 13.75999 |
| Baz2b | 4.726941 | 5.52785 | 6.519577 | 6.74245 | 7.594084 | 7.624006 | 9.34108 | 9.670233 | 13.15492 | 14.08949 | 10.57443 | 11.01324 |
| Tubb6 | 55.37833 | 54.23717 | 76.28883 | 74.30044 | 93.11347 | 86.79951 | 104.881 | 100.1094 | 97.52037 | 95.08593 | 163.3728 | 155.2457 |
| Vsig10 | 5.757863 | 6.039797 | 5.856322 | 5.721461 | 4.749507 | 4.78819 | 3.968086 | 3.503757 | 2.738202 | 2.738352 | 4.111296 | 3.706186 |
| Mirg | 7.486727 | 7.339871 | 4.138508 | 4.523241 | 5.192399 | 5.882453 | 6.128157 | 6.523624 | 4.86491 | 4.572039 | 3.546602 | 3.124036 |
| 5730405O1 | 1.209777 | 0.981529 | 0.81528 | 0.826682 | 0.759424 | 0.766681 | 0.632388 | 0.723506 | 0.381353 | 0.444266 | 0.342432 | 0.395925 |
| Tst | 14.54253 | 14.66733 | 9.309464 | 8.261486 | 9.852319 | 10.36663 | 8.808737 | 8.338579 | 5.852941 | 6.538393 | 5.09892 | 4.967928 |
| Reep5 | 7.509802 | 8.228418 | 10.61887 | 11.21998 | 15.16124 | 14.16901 | 18.86174 | 19.57924 | 21.02517 | 19.39194 | 23.42267 | 23.4077 |
| Cebpb | 1.771363 | 1.565666 | 4.774452 | 4.041749 | 5.330432 | 4.875241 | 4.269084 | 2.773385 | 3.410543 | 2.748091 | 7.34415 | 5.73111 |
| Aqp11 | 1.250815 | 1.439585 | 1.742872 | 1.616166 | 2.590881 | 2.162372 | 2.846964 | 2.517068 | 1.649632 | 1.679456 | 1.365258 | 1.207067 |
| Nr2c2ap | 1.511642 | 0.862733 | 0.60587 | 0.893562 | 1.185029 | 0.80365 | 0.816064 | 0.63819 | 0.415434 | 0.428515 | 0.309716 | 0.450623 |
| Glb1 | 28.72161 | 29.79088 | 21.50656 | 21.78603 | 18.5096 | 17.42201 | 16.98344 | 18.25762 | 16.12072 | 16.43516 | 10.34848 | 10.45302 |
| Cldn7 | 3.906443 | 4.697043 | 2.607223 | 2.443992 | 1.613331 | 1.947623 | 1.490309 | 1.825535 | 1.535116 | 1.35268 | 1.654419 | 1.172937 |
| Zswim8 | 5.739362 | 6.481064 | 13.35376 | 12.18373 | 14.92881 | 15.16307 | 20.15043 | 17.71409 | 21.31189 | 19.03816 | 26.51504 | 25.38124 |
| Gm9 | 0.27154 | 0.597855 | 1.575736 | 1.7632 | 1.97364 | 2.258936 | 3.894848 | 3.671351 | 3.54372 | 3.554506 | 3.897035 | 3.043532 |
| Fbxl8 | 0.221911 | 0.274576 | 1.075187 | 1.209594 | 1.631295 | 1.678359 | 1.755272 | 1.469956 | 1.237675 | 0.806196 | 1.345594 | 0.974632 |
| Ppm1h | 0.674783 | 0.721009 | 0.701441 | 0.690641 | 0.860091 | 0.875377 | 1.106878 | 1.044925 | 1.391723 | 1.437552 | 1.1308 | 1.083062 |
| Nrk | 0.639278 | 0.875791 | 1.677748 | 1.582162 | 1.429942 | 1.187656 | 1.201114 | 1.054699 | 1.93858 | 2.195205 | 2.184332 | 1.703687 |
| Gna12 | 10.70012 | 12.01821 | 13.55108 | 12.93997 | 14.5795 | 14.69062 | 18.1139 | 15.55834 | 26.03798 | 26.9301 | 23.794 | 21.58984 |
| Fndc4 | 5.309522 | 4.621528 | 5.12732 | 5.965509 | 7.232815 | 7.751346 | 8.384735 | 7.545166 | 11.85661 | 10.2831 | 11.25974 | 12.40912 |
| Pfkp | 59.9987 | 56.55785 | 23.14136 | 23.04377 | 22.73867 | 20.26082 | 15.82258 | 17.12664 | 14.18895 | 16.34546 | 12.19725 | 11.08015 |
| Krtcap3 | 4.274843 | 3.877969 | 2.43508 | 2.899152 | 1.997455 | 2.220071 | 1.991449 | 1.593826 | 1.44434 | 1.439416 | 1.785918 | 1.395281 |
| Pgam2 | 1.491636 | 1.960912 | 3.539711 | 4.092008 | 3.429974 | 4.648411 | 4.882296 | 3.961385 | 2.698384 | 2.638291 | 1.585913 | 2.237795 |
| Car12 | 1.269654 | 1.365224 | 0.594235 | 0.759256 | 0.975063 | 0.939282 | 1.070228 | 1.109877 | 1.440234 | 1.576313 | 0.882849 | 0.96431 |
| Helb | 21.29789 | 22.21597 | 17.20829 | 16.01295 | 14.46498 | 13.65743 | 12.88996 | 13.91399 | 8.89377 | 9.872484 | 7.639099 | 7.716392 |
| Senp7 | 1.071541 | 1.206768 | 1.909248 | 1.944401 | 2.339026 | 2.131184 | 2.82828 | 2.908197 | 3.499576 | 4.033493 | 2.399695 | 2.410149 |
| Prps2 | 35.46513 | 36.09564 | 33.27897 | 32.24968 | 28.83199 | 26.14671 | 27.34808 | 25.22638 | 22.8019 | 21.79281 | 13.94737 | 15.55232 |
| Hist1h2ad | 1.824145 | 2.152672 | 0.727365 | 1.19405 | 1.132898 | 1.617022 | 1.846529 | 2.024496 | 1.841998 | 2.359264 | 1.771826 | 1.137845 |
| Slc37a1 | 4.344728 | 4.71891 | 3.819923 | 3.734244 | 2.649211 | 2.600183 | 2.392745 | 1.80258 | 1.952727 | 1.907815 | 2.041454 | 1.987178 |
| Mfsd4 | 1.528102 | 2.204492 | 2.01799 | 2.281533 | 1.893045 | 2.0058 | 2.444157 | 2.337653 | 3.149674 | 3.51527 | 3.720616 | 4.153981 |
| 4921507P0 | 0.279525 | 0.285744 | 0.763658 | 0.962811 | 0.972886 | 1.114776 | 1.488486 | 1.754067 | 1.905502 | 1.993942 | 1.478243 | 1.515333 |
| Top2b | 22.50853 | 23.62941 | 26.94988 | 26.83598 | 33.81734 | 31.75012 | 37.76217 | 38.20579 | 53.79905 | 55.54451 | 47.82004 | 52.46614 |
| Slc37a2 | 0.82862 | 0.919675 | 2.412283 | 2.279023 | 2.63491 | 2.807134 | 3.253958 | 2.7827 | 1.805079 | 1.604261 | 1.390588 | 1.356395 |
| Fras1 | 0.184805 | 0.198721 | 0.397155 | 0.523001 | 0.675008 | 0.746124 | 1.015986 | 0.66539 | 1.03997 | 1.082264 | 0.719595 | 0.639491 |
| Msl3l2 | 1.478496 | 1.493594 | 2.761903 | 2.336191 | 2.889268 | 2.576575 | 3.060229 | 3.788788 | 3.226705 | 2.474091 | 5.869582 | 4.833558 |
| 5031439G0 | 6.721441 | 7.270623 | 7.666316 | 7.031252 | 9.414381 | 9.105354 | 10.46112 | 10.48946 | 13.14406 | 13.62114 | 15.52562 | 14.54748 |
| 4933416C0 | 0.151706 | 0.111514 | 0.639242 | 0.643207 | 0.883602 | 0.817906 | 1.011078 | 1.330743 | 1.229805 | 1.082621 | 1.623979 | 1.236835 |
| Hoxa10 | 0.235519 | 0.123301 | 2.294306 | 2.472805 | 3.905072 | 4.136458 | 4.127421 | 3.599539 | 2.88386 | 2.000775 | 1.846679 | 2.157179 |
| Nlrp5-ps | 0.909585 | 1.146979 | 2.227538 | 2.145364 | 1.745123 | 1.525231 | 1.204927 | 1.796194 | 1.439183 | 1.35536 | 1.216916 | 1.037747 |
| Smim19 | 4.602345 | 4.161589 | 3.93686 | 4.213853 | 5.595109 | 4.447154 | 6.173503 | 5.91464 | 7.241685 | 7.077919 | 9.683664 | 8.710506 |
| Mbp | 1.08519 | 1.344528 | 1.378747 | 1.27422 | 1.38596 | 1.525529 | 2.009208 | 1.739075 | 2.51445 | 2.633086 | 1.862601 | 1.973612 |
| Cfl2 | 24.39951 | 19.94948 | 26.98513 | 27.05729 | 32.94887 | 33.6292 | 38.78066 | 36.32997 | 45.1794 | 44.06541 | 51.28499 | 55.8776 |

| | | | | | | | | | | | | |
|---|---|---|---|---|---|---|---|---|---|---|---|---|
| Snx19 | 3.399034 | 3.844818 | 5.656514 | 5.753865 | 7.20065 | 6.703129 | 8.307608 | 7.671621 | 8.285464 | 8.129574 | 12.82741 | 11.88198 |
| Magee2 | 0.478107 | 0.494553 | 0.956203 | 1.095784 | 0.843428 | 1.018451 | 1.105038 | 1.092756 | 0.522689 | 0.559756 | 0.496972 | 0.530912 |
| Gfra1 | 0.4652 | 0.590311 | 5.645252 | 5.627376 | 7.676717 | 7.925053 | 13.48818 | 11.61326 | 10.98004 | 10.0173 | 7.48907 | 7.927413 |
| Kdelr3 | 25.54858 | 24.90119 | 7.916748 | 7.715002 | 9.916302 | 10.31919 | 11.2865 | 13.85463 | 16.08575 | 15.92121 | 11.68278 | 12.27676 |
| Smarce1 | 36.78743 | 34.07888 | 36.47366 | 36.04028 | 41.78113 | 40.86011 | 50.09421 | 52.70104 | 69.7283 | 68.17592 | 75.18348 | 74.36123 |
| H2-Q7 | 0.982168 | 0.7525 | 2.621628 | 2.474921 | 4.752495 | 4.578366 | 4.430747 | 4.565338 | 3.13218 | 3.41825 | 2.475834 | 2.219595 |
| Rab6b | 1.257717 | 1.511918 | 1.200878 | 1.264317 | 1.971632 | 1.805672 | 2.405353 | 1.900144 | 3.003405 | 2.607711 | 1.356132 | 1.453358 |
| Klhl23 | 1.736013 | 1.69398 | 3.672422 | 3.817165 | 4.349359 | 4.316896 | 4.452299 | 3.887827 | 5.275607 | 4.895718 | 6.838968 | 7.869258 |
| Topaz1 | 0.259599 | 0.302492 | 1.083518 | 1.014697 | 0.830838 | 0.839165 | 0.904907 | 0.867734 | 0.52448 | 0.618724 | 0.573431 | 0.492332 |
| Fanci | 18.87368 | 19.2646 | 19.88803 | 19.26947 | 16.60831 | 15.20223 | 14.90285 | 16.24469 | 12.56354 | 14.27471 | 8.611725 | 9.352026 |
| Cox7c | 435.061 | 282.8581 | 241.7373 | 259.1351 | 310.3722 | 354.0061 | 261.6869 | 251.4561 | 191.8123 | 172.8919 | 182.6969 | 171.8357 |
| Mcf2l | 4.167661 | 4.994784 | 2.901147 | 2.912291 | 2.917705 | 3.078577 | 3.058262 | 3.152218 | 2.103342 | 2.115257 | 1.331611 | 1.532128 |
| Tdrd7 | 0.684017 | 0.485017 | 1.268766 | 1.343949 | 2.243932 | 2.122364 | 2.569881 | 2.762081 | 2.519552 | 2.411142 | 2.414964 | 2.59701 |
| Dlc1 | 24.68322 | 30.88378 | 17.71915 | 17.99377 | 12.55419 | 12.3639 | 10.76121 | 9.64226 | 9.20929 | 9.722753 | 8.92108 | 8.758589 |
| Hspg2 | 24.21714 | 33.29931 | 42.79068 | 40.76851 | 35.21673 | 32.12104 | 37.97548 | 26.87747 | 39.28328 | 38.17661 | 58.1836 | 55.19509 |
| Il34 | 0.134514 | 0.199163 | 0.458087 | 0.452605 | 0.582307 | 0.58043 | 0.896586 | 0.782858 | 0.921642 | 0.692053 | 1.494691 | 0.929165 |
| Rpl39 | 635.6662 | 423.1302 | 210.8574 | 300.7036 | 366.6708 | 476.8192 | 308.3887 | 293.2256 | 314.7854 | 282.1436 | 283.2011 | 232.264 |
| Fam161a | 0.852964 | 1.160395 | 1.870439 | 1.543022 | 1.720965 | 1.43995 | 1.37081 | 1.227171 | 1.094959 | 0.996858 | 0.755508 | 0.751743 |
| Asnsd1 | 2.673498 | 2.199627 | 1.152126 | 1.891789 | 2.476993 | 2.474201 | 2.098006 | 1.842441 | 1.883664 | 2.091085 | 1.33772 | 1.205442 |
| Nudt10 | 0.896509 | 0.91631 | 2.205875 | 2.31158 | 2.732588 | 3.005032 | 3.414063 | 3.115671 | 4.613247 | 3.927691 | 4.973785 | 4.743437 |
| Robo1 | 3.980557 | 4.894582 | 3.837317 | 4.020594 | 4.495785 | 4.369245 | 5.869722 | 5.199192 | 8.25125 | 8.249228 | 7.598443 | 7.164414 |
| Mapk1ip1 | 9.29938 | 10.36337 | 7.940532 | 8.324655 | 7.416556 | 7.418041 | 5.268854 | 5.973501 | 4.450014 | 4.935157 | 3.41666 | 4.057686 |
| Aldh1b1 | 5.12822 | 4.728029 | 2.672169 | 2.662827 | 2.821789 | 2.131158 | 2.245662 | 1.549999 | 1.33728 | 1.298247 | 1.516401 | 1.736382 |
| Gm14420 | 7.696095 | 7.721089 | 15.7066 | 14.82416 | 11.03634 | 12.13364 | 9.555371 | 10.03655 | 7.326353 | 8.24347 | 6.152384 | 7.231773 |
| Cpsf4l | 12.25206 | 12.17702 | 5.468856 | 4.886598 | 3.894572 | 3.754363 | 3.331042 | 3.101749 | 2.413487 | 2.384496 | 2.933856 | 2.582187 |
| Slc12a5 | 0.442437 | 0.539155 | 1.577696 | 1.503646 | 1.574819 | 1.719266 | 1.589657 | 1.587133 | 0.965579 | 0.845999 | 0.868247 | 0.838945 |
| Kif1b | 8.38891 | 10.01923 | 14.51306 | 15.24134 | 17.77796 | 17.12143 | 20.54616 | 18.61768 | 24.70313 | 26.67293 | 28.88693 | 31.23402 |
| Plscr3 | 3.699621 | 3.683533 | 4.241428 | 3.736854 | 5.778839 | 5.311288 | 6.036141 | 5.67117 | 8.555741 | 7.657309 | 6.719513 | 6.184709 |
| 2610305D1 | 70.63835 | 64.89143 | 54.21035 | 53.81317 | 43.56463 | 41.95946 | 33.44092 | 35.90012 | 28.11589 | 27.66298 | 24.49725 | 26.26199 |
| Gm4984 | 2.243354 | 2.535842 | 3.04914 | 1.774638 | 1.788813 | 1.477731 | 2.028217 | 2.213168 | 1.428741 | 1.200366 | 1.059268 | 1.08026 |
| Rnf144b | 0.251396 | 0.179576 | 0.78578 | 0.719007 | 1.065439 | 0.930725 | 0.931289 | 0.891048 | 0.945634 | 0.793591 | 1.372608 | 1.472905 |
| Zhx1 | 2.038103 | 2.054411 | 4.569576 | 4.264652 | 5.675412 | 5.274747 | 6.509809 | 7.013878 | 7.491369 | 7.661469 | 7.909658 | 8.735844 |
| Mrc2 | 1.148098 | 1.406188 | 3.322764 | 3.507393 | 4.729879 | 4.789102 | 7.180161 | 7.036129 | 7.44676 | 7.182977 | 4.486401 | 4.335829 |
| D17H6S53E | 1.271013 | 1.325208 | 1.94735 | 1.572529 | 1.748179 | 1.906714 | 2.420603 | 2.494327 | 2.894375 | 2.825342 | 3.186085 | 3.220446 |
| Ano6 | 6.492862 | 6.8761 | 7.364029 | 7.41956 | 8.077392 | 7.833669 | 10.03404 | 9.697499 | 12.64864 | 12.64288 | 13.80798 | 15.19298 |
| Tm6sf1 | 0.984525 | 0.925712 | 1.172627 | 1.230808 | 1.755913 | 1.821856 | 1.995614 | 2.097299 | 2.217943 | 2.519662 | 1.494908 | 1.704965 |
| Apbb1 | 6.056714 | 5.79406 | 8.909347 | 8.898267 | 13.90805 | 13.46573 | 15.71894 | 18.20619 | 16.05986 | 15.25266 | 14.06178 | 13.86846 |
| Dgkz | 20.19959 | 21.23662 | 39.30551 | 38.69385 | 57.99211 | 55.05664 | 77.30793 | 79.1648 | 78.21844 | 78.79553 | 63.3507 | 58.05126 |
| Vangl2 | 6.466401 | 7.859498 | 9.008317 | 8.664217 | 9.812188 | 9.404104 | 10.94996 | 9.72808 | 18.56135 | 17.72503 | 15.54629 | 15.90468 |
| Cacnb1 | 4.063742 | 4.031416 | 3.118075 | 2.985072 | 3.048696 | 3.460725 | 2.462108 | 2.732533 | 2.23114 | 1.972924 | 1.802344 | 1.692173 |
| Serping1 | 4.64024 | 3.974003 | 7.323787 | 7.31929 | 8.659847 | 8.725796 | 13.91224 | 14.18207 | 15.49509 | 14.96662 | 13.64181 | 14.10336 |
| Auts2 | 3.529124 | 4.748879 | 4.022078 | 4.250077 | 4.535949 | 4.501233 | 6.005024 | 5.098512 | 8.468579 | 8.152672 | 8.926867 | 8.690479 |
| Mrpl24 | 47.75091 | 45.89319 | 47.87049 | 46.54444 | 42.1187 | 40.38942 | 35.06968 | 40.3568 | 28.36513 | 27.19674 | 21.31928 | 22.77133 |
| Krt17 | 2.063979 | 2.000231 | 0.845157 | 0.447502 | 0.364033 | 0.331143 | 0.334058 | 0.45326 | 0.353161 | 0.314402 | 0.870885 | 0.642309 |
| Rhox13 | 4.666745 | 3.982961 | 8.382159 | 8.2037 | 13.19465 | 12.58386 | 13.39248 | 16.74589 | 12.30417 | 11.72173 | 14.47682 | 13.27494 |
| Tle3 | 10.04841 | 12.97623 | 10.12887 | 10.2007 | 10.65468 | 10.27639 | 13.88386 | 12.66355 | 21.09396 | 19.74907 | 22.93924 | 20.81671 |
| Mpv17l | 4.702243 | 4.454586 | 4.035162 | 3.635879 | 3.632057 | 3.603578 | 2.855091 | 2.913073 | 2.584478 | 2.767754 | 1.636814 | 1.782383 |
| Lrrc73 | 0.277605 | 0.263785 | 0.364997 | 0.602525 | 0.495724 | 0.823319 | 0.870694 | 0.982042 | 0.7002 | 0.567187 | 1.704664 | 1.156836 |
| Mllt3 | 2.199126 | 2.264338 | 2.882651 | 3.147191 | 3.78305 | 3.588695 | 4.377333 | 4.332252 | 6.249906 | 6.416977 | 3.554599 | 3.782377 |
| Kcnk6 | 1.118957 | 1.311963 | 3.277694 | 3.395861 | 4.879291 | 5.016096 | 5.767539 | 5.169249 | 3.649595 | 3.567925 | 2.503802 | 2.53624 |
| Shc2 | 3.753389 | 4.138136 | 4.59539 | 4.485934 | 5.403018 | 5.274953 | 6.297431 | 6.268695 | 7.506699 | 7.319579 | 9.067619 | 9.141623 |
| Narf | 6.867209 | 6.095531 | 5.74396 | 6.030533 | 7.198425 | 6.440551 | 6.709456 | 7.09138 | 6.072708 | 7.093212 | 3.425987 | 3.483346 |
| Nova1 | 1.759179 | 1.961204 | 2.329047 | 2.11616 | 2.931152 | 2.796174 | 3.075975 | 2.778549 | 4.499584 | 4.242763 | 3.660734 | 4.306651 |
| Aldh1l2 | 0.451673 | 0.583416 | 1.07933 | 1.135191 | 0.851895 | 0.774023 | 0.590213 | 0.684305 | 0.580984 | 0.557808 | 0.833839 | 0.983414 |
| Slc17a9 | 7.0947 | 7.602162 | 1.273236 | 0.908612 | 0.973687 | 0.596119 | 0.973845 | 1.009445 | 0.928792 | 1.21291 | 0.701287 | 0.767673 |
| Aldh1l1 | 2.355956 | 2.258811 | 1.30336 | 1.156733 | 1.412983 | 1.030901 | 1.090057 | 1.29095 | 0.829743 | 1.075404 | 0.812495 | 0.634624 |
| Atxn7 | 1.290024 | 1.330116 | 2.167549 | 2.107465 | 2.423439 | 2.337343 | 2.596741 | 2.416253 | 2.854845 | 2.956714 | 4.297436 | 4.286232 |
| Rcan2 | 0.52716 | 0.442707 | 0.755478 | 0.864951 | 1.426436 | 1.265622 | 2.028432 | 1.742012 | 1.682781 | 1.758259 | 1.453955 | 1.262557 |
| Rab11fip2 | 1.301075 | 1.576711 | 1.921594 | 2.077129 | 2.264477 | 2.338253 | 2.541595 | 2.481867 | 3.630674 | 3.382692 | 3.770524 | 4.221805 |
| Fbxl17 | 1.93633 | 2.214965 | 1.93712 | 2.072284 | 2.538293 | 2.646585 | 3.375135 | 2.845332 | 3.63388 | 3.240416 | 4.128257 | 4.211549 |
| Pim3 | 69.47344 | 69.75841 | 61.86788 | 54.7481 | 43.25489 | 42.84775 | 36.07666 | 36.30795 | 26.7431 | 26.93914 | 36.00797 | 34.52396 |

| | | | | | | | | | | | | |
|---|---|---|---|---|---|---|---|---|---|---|---|---|
| Sh3bp5 | 1.037032 | 1.081441 | 2.280734 | 2.786403 | 3.683583 | 3.28323 | 3.997133 | 3.701871 | 5.25298 | 5.045793 | 5.44614 | 5.660682 |
| E330009J0 | 3.235949 | 3.673895 | 4.302709 | 4.658659 | 5.328452 | 5.418778 | 6.752515 | 5.728045 | 8.132222 | 8.977935 | 4.082534 | 4.420151 |
| Folr2 | 0.989638 | 1.061525 | 2.676638 | 2.793243 | 2.146765 | 2.163193 | 2.337679 | 1.867734 | 1.831848 | 1.376004 | 1.6128 | 1.383849 |
| Clip1 | 4.684849 | 5.192401 | 7.079657 | 7.148856 | 8.781425 | 8.993768 | 10.6625 | 9.819698 | 13.90489 | 14.30471 | 13.98943 | 14.51094 |
| Mov10 | 32.3511 | 41.34388 | 23.67566 | 22.00098 | 18.79616 | 18.21503 | 19.60177 | 17.10561 | 15.83626 | 15.89663 | 11.68369 | 10.83934 |
| Zfyve9 | 3.861692 | 3.933925 | 4.809009 | 4.891527 | 6.007331 | 5.665465 | 6.293762 | 6.610299 | 8.430615 | 9.256726 | 9.044645 | 9.927204 |
| Prr5 | 10.50299 | 11.64861 | 11.75664 | 9.984684 | 11.93012 | 12.49702 | 13.39076 | 12.01129 | 18.72825 | 16.70845 | 21.48783 | 20.26245 |
| P2rx7 | 1.138775 | 1.320003 | 2.484222 | 2.371709 | 1.623978 | 1.598427 | 1.65299 | 1.851516 | 1.940015 | 1.834884 | 2.997469 | 3.242895 |
| Klhl34 | 0.249989 | 0.281206 | 0.928583 | 0.79978 | 0.864178 | 0.864611 | 1.346799 | 0.847725 | 1.028301 | 1.535538 | 1.327493 | 1.622331 |
| Cd74 | 2.007758 | 1.080053 | 1.912241 | 1.602337 | 1.930538 | 1.990233 | 2.392289 | 3.091234 | 2.078459 | 1.691247 | 4.330532 | 3.249829 |
| Pgbd5 | 2.446484 | 2.460256 | 5.0825 | 4.363553 | 6.539708 | 7.332639 | 6.901794 | 6.138396 | 3.532843 | 3.615619 | 4.695111 | 3.628569 |
| Hykk | 1.101033 | 1.651624 | 2.217364 | 2.26787 | 2.096817 | 1.782071 | 1.8878 | 1.586694 | 1.496273 | 1.560124 | 1.143325 | 1.119125 |
| Arl4a | 11.84094 | 9.844555 | 5.500789 | 5.662438 | 4.88884 | 5.082476 | 4.693924 | 5.036783 | 6.788861 | 6.808945 | 9.51681 | 10.20382 |
| Rsph3b | 0.94787 | 0.941254 | 1.473777 | 1.305554 | 1.683517 | 2.312091 | 2.194974 | 2.148801 | 2.252183 | 2.188625 | 2.625105 | 2.643056 |
| Fam212b | 0.725559 | 0.641132 | 1.20287 | 1.533893 | 1.426665 | 1.714942 | 2.305882 | 2.302978 | 1.948901 | 1.753281 | 3.355424 | 3.103899 |
| Impad1 | 15.49298 | 17.55503 | 18.5744 | 17.75296 | 20.90398 | 20.55246 | 24.6948 | 20.80833 | 33.65304 | 35.6389 | 34.08245 | 35.91972 |
| Zfand3 | 16.95834 | 16.19254 | 23.46612 | 23.35055 | 28.92274 | 28.77271 | 34.31369 | 35.09657 | 41.20295 | 40.75456 | 49.94288 | 47.23357 |
| Reep2 | 2.384264 | 2.544168 | 3.074303 | 3.44627 | 4.714959 | 4.939172 | 5.816168 | 6.18081 | 7.509812 | 5.459961 | 6.538049 | 6.970187 |
| Syngr3 | 2.431594 | 2.171304 | 1.332011 | 1.430556 | 1.936968 | 1.640905 | 1.678326 | 2.122733 | 1.141187 | 1.346728 | 1.062729 | 1.049696 |
| Mapkapk3 | 13.89521 | 14.989 | 8.705775 | 8.093751 | 8.010991 | 7.383566 | 7.367723 | 7.025665 | 4.648993 | 4.004913 | 4.515241 | 4.604752 |
| Prob1 | 0.280678 | 0.263884 | 0.465948 | 0.578752 | 0.696381 | 0.645543 | 0.917565 | 1.083023 | 1.143022 | 1.169511 | 1.096514 | 1.062179 |
| Ppp3cc | 3.410988 | 2.610195 | 4.340564 | 4.062267 | 5.750418 | 5.74019 | 6.246926 | 8.011674 | 8.686648 | 8.036942 | 6.86606 | 8.207902 |
| Galnt12 | 1.267063 | 1.258349 | 2.03921 | 1.926788 | 3.141946 | 2.689616 | 2.993963 | 3.891935 | 3.043084 | 2.726522 | 2.174473 | 2.073824 |
| Ccnjl | 1.526015 | 2.050088 | 2.771982 | 2.977088 | 4.071148 | 4.190386 | 5.252221 | 4.332399 | 6.335015 | 6.013412 | 5.362942 | 5.073058 |
| Nrg2 | 0.671716 | 0.826076 | 1.4425 | 1.203181 | 1.295342 | 1.125138 | 1.604333 | 1.327458 | 1.508924 | 1.750993 | 1.9476 | 2.272522 |
| Wbscr27 | 5.972367 | 5.620078 | 4.860258 | 4.974754 | 5.774762 | 5.878204 | 5.450972 | 6.342412 | 4.406799 | 5.09061 | 2.701053 | 3.140876 |
| Sorbs3 | 1.81448 | 1.955989 | 2.146059 | 2.557432 | 3.227148 | 3.612593 | 4.343837 | 3.731354 | 5.701268 | 5.163777 | 5.321715 | 5.021818 |
| Ifitm1 | 85.79429 | 78.94554 | 16.45208 | 15.78527 | 12.10324 | 9.721544 | 8.669611 | 9.139614 | 20.0422 | 18.45252 | 17.6379 | 16.38796 |
| Hmgn5 | 28.01158 | 26.6138 | 25.1165 | 24.24744 | 25.5794 | 23.71814 | 19.47424 | 26.43847 | 19.91337 | 18.99975 | 11.74645 | 13.09933 |
| Nmrk1 | 3.947648 | 3.913085 | 3.249628 | 3.601289 | 3.706522 | 3.850444 | 3.528879 | 3.708709 | 2.950428 | 2.991986 | 1.937083 | 1.907817 |
| Tmem28 | 2.559018 | 2.745436 | 2.653247 | 2.638702 | 2.586296 | 2.208702 | 1.777776 | 1.462723 | 1.915866 | 1.712947 | 1.241559 | 1.307641 |
| Bai2 | 1.977443 | 2.370025 | 3.338145 | 3.260588 | 3.912723 | 3.626341 | 4.002382 | 3.861776 | 2.844752 | 2.731441 | 2.170476 | 1.913764 |
| Aff1 | 7.058768 | 9.879811 | 9.118844 | 8.952678 | 8.609192 | 7.899158 | 11.65614 | 9.285574 | 13.38575 | 13.22385 | 16.66709 | 15.92646 |
| 5730508B0 | 1.33054 | 1.114874 | 2.048993 | 2.100061 | 2.715372 | 2.704973 | 2.158209 | 2.105072 | 1.579116 | 1.341722 | 1.546405 | 2.053696 |
| Tomm5 | 181.7327 | 135.4743 | 98.48756 | 100.8556 | 92.49208 | 97.30614 | 69.98098 | 71.25322 | 58.72207 | 50.0283 | 57.48471 | 55.39809 |
| Mafk | 2.734365 | 3.109559 | 3.209876 | 3.2246 | 2.587643 | 2.696485 | 2.820782 | 2.860149 | 3.278032 | 2.658172 | 5.640046 | 5.356482 |
| Ptplad2 | 0.340557 | 0.337694 | 0.723232 | 0.68453 | 0.744975 | 0.812505 | 1.060045 | 1.011448 | 1.135577 | 1.378625 | 1.003434 | 1.258905 |
| Slc9b1 | 12.28369 | 13.24442 | 11.26683 | 10.37947 | 8.503623 | 8.358437 | 7.482655 | 8.568672 | 5.425463 | 5.15405 | 6.300919 | 6.530136 |
| Nfkbid | 4.130133 | 4.425463 | 2.527618 | 2.242618 | 1.651421 | 2.138863 | 1.469955 | 2.183606 | 1.598297 | 1.11411 | 1.301935 | 1.329156 |
| Ubtfl1 | 0.084065 | 0.112693 | 0.713165 | 0.60883 | 0.779167 | 0.764393 | 1.048026 | 1.225399 | 1.149549 | 1.186854 | 0.980551 | 0.829089 |
| Caskin1 | 0.388992 | 0.404719 | 1.3931 | 1.233733 | 1.925379 | 2.281527 | 2.41388 | 2.483075 | 2.02974 | 1.891692 | 1.60779 | 1.735653 |
| Ppp1r18 | 8.956212 | 9.817953 | 19.28751 | 18.50667 | 18.09555 | 18.80755 | 15.80659 | 14.73574 | 10.11211 | 9.347753 | 18.20701 | 17.26082 |
| Tmem43 | 10.39563 | 11.80377 | 12.77024 | 12.78589 | 15.50198 | 14.41619 | 17.43053 | 15.71397 | 20.56123 | 19.33236 | 23.79625 | 25.72233 |
| Sall3 | 1.039594 | 1.194698 | 1.945722 | 1.930642 | 2.015617 | 2.000992 | 2.896435 | 2.59314 | 4.469548 | 3.88089 | 3.746811 | 3.883813 |
| Fbxo44 | 3.515273 | 3.825598 | 1.692091 | 1.686604 | 1.299004 | 1.20558 | 1.156432 | 0.999353 | 0.701676 | 0.838867 | 1.212869 | 0.908131 |
| Specc1 | 2.731793 | 2.872957 | 4.759681 | 5.111401 | 5.999082 | 5.331472 | 6.278516 | 6.479069 | 7.789343 | 8.261471 | 10.4678 | 10.27227 |
| Tmem38b | 7.458748 | 7.473744 | 8.100487 | 7.98286 | 7.613634 | 6.97047 | 6.593246 | 6.370125 | 4.964155 | 5.08728 | 3.676254 | 3.974275 |
| Rnpepl1 | 4.204187 | 5.091103 | 6.775954 | 7.427708 | 9.373256 | 8.785329 | 10.6607 | 10.3442 | 12.94299 | 12.9015 | 16.09925 | 14.91775 |
| Tmem198 | 0.36856 | 0.517345 | 0.78627 | 0.709528 | 0.728994 | 0.833936 | 0.530393 | 0.553398 | 0.732375 | 0.509296 | 1.11651 | 1.022719 |
| H2bfm | 0.267963 | 0.182485 | 0.690897 | 0.976197 | 0.863851 | 1.540892 | 2.210796 | 1.96022 | 1.346768 | 1.258726 | 1.136041 | 1.03297 |
| Elf1 | 2.099315 | 2.600384 | 3.483487 | 3.16414 | 3.776626 | 3.385155 | 4.971338 | 4.378855 | 4.833475 | 5.188971 | 6.349377 | 6.353189 |
| Bmpr2 | 2.88948 | 3.580067 | 4.137459 | 4.552196 | 5.300211 | 4.986741 | 5.995044 | 5.075845 | 8.287927 | 9.137552 | 8.013139 | 8.787879 |
| Fgfr2 | 7.226449 | 7.331487 | 13.14807 | 13.17856 | 17.24592 | 17.59498 | 20.35897 | 19.12143 | 26.69136 | 26.43954 | 24.6669 | 25.49498 |
| Art5 | 0.494503 | 0.542865 | 0.888407 | 0.94138 | 1.457759 | 1.210952 | 2.01962 | 1.887816 | 1.50901 | 1.789075 | 1.011138 | 0.965467 |
| Gnaz | 1.261178 | 1.476329 | 1.99757 | 2.103264 | 2.896687 | 2.354216 | 2.658847 | 3.04989 | 2.208584 | 2.456518 | 1.812289 | 1.521099 |
| Txlng | 18.41397 | 18.43323 | 13.42579 | 13.56223 | 11.46876 | 11.66023 | 10.11868 | 10.4342 | 9.786578 | 10.41423 | 6.362901 | 6.764387 |
| Nfil3 | 2.039485 | 1.972552 | 3.329717 | 2.831943 | 3.890497 | 4.400888 | 5.530393 | 4.603431 | 6.11071 | 5.677697 | 5.380289 | 5.516384 |
| Rmnd5b | 24.47306 | 26.37833 | 21.67028 | 20.92651 | 17.41879 | 16.35659 | 13.96531 | 15.07646 | 11.48208 | 10.43902 | 13.40764 | 12.02172 |
| Tomm7 | 49.35032 | 34.91448 | 36.46529 | 36.04695 | 37.388 | 39.92887 | 32.14334 | 25.91229 | 21.95665 | 19.91906 | 25.79138 | 25.37831 |
| Nr2f6 | 5.305668 | 5.487604 | 10.56969 | 10.04727 | 8.29617 | 8.821508 | 8.243704 | 8.304223 | 9.599317 | 8.487594 | 19.87357 | 16.64277 |
| C77370 | 4.253176 | 4.550421 | 3.150542 | 3.314942 | 2.498227 | 2.372465 | 1.830899 | 2.017458 | 1.6615 | 1.654205 | 1.854929 | 1.797534 |

| | | | | | | | | | | | | |
|---|---|---|---|---|---|---|---|---|---|---|---|---|
| Fndc3b | 3.282934 | 3.603883 | 5.542973 | 6.034041 | 6.786969 | 6.728147 | 8.181807 | 7.103347 | 10.23026 | 10.62876 | 12.65168 | 12.08968 |
| Mertk | 0.914214 | 1.19852 | 2.079409 | 2.044623 | 2.227773 | 1.844625 | 1.685697 | 1.68861 | 1.144908 | 1.020607 | 0.957869 | 1.103652 |
| Ppp1r26 | 1.05984 | 1.163626 | 1.821815 | 1.784586 | 1.878881 | 1.75202 | 1.787931 | 1.81775 | 2.441925 | 2.156698 | 3.146385 | 3.506382 |
| Sccpdh | 5.522324 | 4.4918 | 6.121358 | 5.848853 | 8.007295 | 7.965673 | 8.242631 | 8.818015 | 9.075918 | 9.223873 | 11.3831 | 11.70536 |
| Spink2 | 0.533096 | 0.98085 | 0.755784 | 0.731249 | 1.504632 | 1.286659 | 1.375244 | 0.728882 | 1.359627 | 1.458282 | 0.56175 | 1.061455 |
| Bnc2 | 6.652699 | 8.293086 | 3.174201 | 3.085388 | 2.001806 | 2.122501 | 1.915597 | 1.542203 | 1.715024 | 1.817226 | 1.576397 | 1.648369 |
| Slc2a8 | 9.131454 | 8.634182 | 8.548993 | 8.31463 | 8.597034 | 8.244936 | 8.166631 | 8.24843 | 6.759405 | 6.498747 | 4.514417 | 4.156944 |
| Ptpn14 | 5.391628 | 6.303636 | 8.669028 | 9.328848 | 10.42245 | 10.40071 | 13.47548 | 10.16763 | 13.15485 | 13.53956 | 17.72682 | 18.6534 |
| Zdhhc15 | 3.010978 | 3.529213 | 3.204699 | 3.34788 | 4.021863 | 3.825948 | 5.483592 | 5.453223 | 6.414627 | 6.69072 | 5.721508 | 6.470267 |
| Tspyl4 | 0.654628 | 0.649539 | 1.928668 | 2.002696 | 2.336112 | 2.547269 | 2.485459 | 2.573522 | 1.998144 | 1.692499 | 3.270329 | 3.38056 |
| Tmem246 | 1.71426 | 1.632745 | 1.963857 | 1.831235 | 2.855748 | 2.635289 | 3.616078 | 3.24573 | 3.604637 | 3.65746 | 2.686702 | 3.195962 |
| Mybl2 | 167.0972 | 182.1569 | 99.04533 | 90.34235 | 64.90644 | 59.88326 | 51.75787 | 51.96329 | 51.16527 | 52.20688 | 47.08299 | 45.24122 |
| Zfp579 | 1.061426 | 1.438796 | 2.133134 | 1.845606 | 2.022402 | 2.051073 | 2.02426 | 1.901798 | 2.488774 | 2.409126 | 3.52148 | 3.685008 |
| 2410004N( | 51.73941 | 39.56089 | 30.59345 | 35.24861 | 28.71922 | 39.72524 | 23.44114 | 29.86415 | 24.67496 | 19.89742 | 22.91184 | 24.51706 |
| 4930519F1 | 2.020084 | 1.913895 | 0.581112 | 0.678874 | 0.410083 | 0.495919 | 0.292787 | 0.340439 | 0.284341 | 0.378225 | 0.271846 | 0.348762 |
| A2m | 7.209033 | 5.832861 | 1.266541 | 1.196854 | 1.673303 | 1.63267 | 2.279461 | 2.37848 | 2.316245 | 2.385512 | 1.773569 | 1.706553 |
| Apod | 1.20478 | 0.989755 | 0.989668 | 0.853862 | 0.766923 | 0.937182 | 1.007388 | 0.926495 | 0.591008 | 0.470239 | 0.527705 | 0.517873 |
| Notch1 | 4.180665 | 6.297547 | 6.733744 | 6.93609 | 6.207537 | 6.219301 | 7.860799 | 6.159703 | 11.36128 | 11.19058 | 12.48947 | 12.27369 |
| Slc16a2 | 0.995347 | 1.063484 | 4.621727 | 4.766489 | 4.997428 | 4.89205 | 5.996623 | 5.294371 | 3.396168 | 3.411501 | 2.244116 | 2.659155 |
| D630045J1 | 1.096511 | 1.614328 | 2.283986 | 2.423402 | 2.905789 | 2.78628 | 3.179408 | 2.679218 | 4.279068 | 4.824656 | 4.072823 | 4.014319 |
| Zbtb39 | 4.016496 | 4.413153 | 5.800333 | 5.99362 | 6.384259 | 6.21433 | 7.454699 | 6.660392 | 10.77684 | 10.86858 | 11.81202 | 11.93179 |
| Nek8 | 6.066923 | 5.962656 | 6.318852 | 5.642435 | 6.041486 | 6.053981 | 4.978082 | 5.415247 | 3.558891 | 3.628892 | 3.022204 | 3.042268 |
| Fndc5 | 1.094012 | 1.126162 | 2.424563 | 2.634333 | 4.547324 | 4.027468 | 4.801112 | 5.241379 | 5.027845 | 5.146076 | 5.136147 | 5.11575 |
| Cyp27a1 | 1.652312 | 1.793276 | 2.307164 | 2.739566 | 3.968942 | 3.285843 | 3.363327 | 3.765221 | 2.922079 | 2.785926 | 1.709212 | 1.892467 |
| Pbx3 | 5.305252 | 5.384187 | 7.443619 | 7.484399 | 9.253543 | 9.447823 | 11.60555 | 11.06903 | 14.58411 | 14.8882 | 11.93783 | 12.49037 |
| Usp29 | 0.679036 | 0.557545 | 1.443005 | 1.336618 | 1.127712 | 1.035171 | 0.912989 | 1.059448 | 0.89052 | 1.172671 | 0.450185 | 0.672067 |
| Prex2 | 3.343966 | 4.238849 | 9.736384 | 10.43274 | 8.649541 | 8.500476 | 9.305095 | 8.056407 | 6.746159 | 6.808626 | 5.492985 | 5.246101 |
| Lgals1 | 39.27245 | 35.25851 | 48.91892 | 47.62038 | 65.82303 | 67.04897 | 76.40838 | 70.18347 | 69.52497 | 62.3248 | 114.1384 | 94.69171 |
| Smtn | 5.652071 | 6.911552 | 7.635645 | 6.934903 | 7.381718 | 7.037194 | 7.530137 | 8.138762 | 6.675481 | 6.464942 | 14.14027 | 12.85259 |
| Snx1 | 20.59198 | 18.09582 | 21.31637 | 21.11916 | 28.32414 | 26.93162 | 34.47194 | 38.04984 | 41.95861 | 41.97001 | 38.01398 | 39.69095 |
| Cdc42ep4 | 18.95377 | 19.59421 | 8.39991 | 7.711337 | 11.29463 | 10.34737 | 14.26674 | 13.10352 | 15.79866 | 15.32298 | 11.46302 | 10.96297 |
| Fah | 15.33947 | 15.15807 | 9.188374 | 9.277892 | 9.689752 | 10.02979 | 9.274913 | 8.289417 | 7.461438 | 7.569227 | 5.052908 | 5.047645 |
| Traf3 | 3.464399 | 3.820803 | 5.95926 | 5.904415 | 6.970597 | 6.604328 | 7.280751 | 6.897295 | 8.448368 | 8.582826 | 12.01389 | 11.72857 |
| Osbpl10 | 8.882307 | 8.764687 | 6.580001 | 7.063924 | 6.124249 | 6.29443 | 6.320921 | 6.184558 | 4.490133 | 4.807231 | 3.515792 | 3.558757 |
| Rnf125 | 10.38291 | 10.63457 | 5.501145 | 4.546116 | 2.824195 | 2.99859 | 2.108343 | 2.42416 | 2.860477 | 2.291301 | 2.664446 | 2.853411 |
| Nxt2 | 18.09084 | 16.88486 | 16.99438 | 16.86143 | 17.06333 | 16.80538 | 16.20434 | 16.65647 | 11.53612 | 11.7405 | 7.041529 | 8.502726 |
| Mgat5 | 1.545312 | 2.230484 | 2.923017 | 2.788582 | 3.086833 | 2.910566 | 3.413013 | 3.111139 | 5.149366 | 5.525813 | 4.363044 | 4.654345 |
| Sh3bp5l | 5.8724 | 5.322102 | 7.392129 | 7.16146 | 7.651258 | 7.095903 | 7.996652 | 7.91371 | 7.554301 | 7.277313 | 13.19423 | 14.06046 |
| Pvt1 | 4.796286 | 5.113494 | 3.179493 | 2.720594 | 1.770695 | 2.207575 | 1.759254 | 2.019281 | 1.429811 | 1.374879 | 1.664994 | 1.694271 |
| Sik2 | 2.753442 | 3.392817 | 4.232705 | 4.568738 | 4.466994 | 4.296571 | 4.746607 | 4.233663 | 4.275972 | 4.691017 | 7.395197 | 8.37565 |
| Zfp52 | 2.252293 | 2.309435 | 1.718868 | 2.264201 | 1.629322 | 1.617541 | 1.32706 | 2.068616 | 1.596407 | 1.398973 | 0.97614 | 1.144549 |
| Atg4d | 8.976152 | 9.279504 | 18.09913 | 16.76786 | 14.72102 | 14.70558 | 13.12855 | 13.02171 | 8.282319 | 8.478228 | 9.468825 | 9.538951 |
| Ttc39c | 1.09882 | 1.017407 | 1.835022 | 2.235862 | 3.049389 | 2.448435 | 3.251806 | 3.901978 | 3.776959 | 4.421715 | 2.923218 | 2.944163 |
| Arsa | 4.646243 | 5.303136 | 4.423386 | 3.943997 | 4.32413 | 4.280421 | 4.149835 | 3.878748 | 3.98979 | 3.764719 | 2.144263 | 2.165743 |
| Ier5l | 8.889341 | 9.329378 | 48.22367 | 42.57545 | 63.35339 | 64.05184 | 58.16536 | 56.52699 | 38.23274 | 32.43437 | 47.67957 | 41.07519 |
| Gramd3 | 3.413119 | 3.777987 | 4.122954 | 4.161261 | 5.992487 | 5.559061 | 7.162078 | 7.826174 | 8.138658 | 8.215847 | 6.181835 | 6.805629 |
| Aqp3 | 15.78442 | 17.1735 | 133.6292 | 126.2885 | 121.045 | 111.8311 | 99.18146 | 109.4008 | 77.17673 | 76.88346 | 74.71186 | 64.03181 |
| Shank2 | 0.186141 | 0.218245 | 0.69767 | 0.923861 | 1.161168 | 1.268328 | 1.524382 | 1.110669 | 0.762765 | 0.782023 | 0.543372 | 0.643539 |
| Abcc4 | 18.65778 | 23.11464 | 16.23599 | 16.05655 | 13.28859 | 13.94688 | 13.57619 | 12.41008 | 11.40356 | 11.43002 | 8.106731 | 8.147459 |
| Slc16a14 | 0.308173 | 0.36246 | 1.3432 | 1.269625 | 1.267308 | 1.346822 | 1.200551 | 0.920682 | 0.681577 | 0.683608 | 0.922895 | 1.095416 |
| Man2a2 | 3.953339 | 4.522061 | 5.726548 | 5.583818 | 6.487247 | 6.080528 | 7.517306 | 7.652646 | 10.27866 | 11.0005 | 7.896379 | 7.759134 |
| Ddit4l | 4.085005 | 3.818025 | 4.702146 | 4.413925 | 6.939383 | 6.546077 | 8.408881 | 8.695033 | 6.531388 | 6.394713 | 8.307386 | 8.207429 |
| Tmcc2 | 2.441922 | 2.815169 | 4.702046 | 4.564977 | 6.510049 | 6.1256 | 6.673247 | 6.777234 | 8.015003 | 8.558241 | 9.7143 | 8.98883 |
| Acyp1 | 12.63847 | 9.657596 | 10.44546 | 10.53274 | 10.48961 | 10.56041 | 10.7603 | 8.672006 | 7.079285 | 6.815472 | 5.894086 | 5.36381 |
| Zscan29 | 1.95152 | 2.249255 | 2.601256 | 2.389269 | 2.87181 | 2.933892 | 3.286606 | 3.594499 | 4.389564 | 4.701954 | 4.841835 | 4.446748 |
| Marveld1 | 6.710929 | 6.917418 | 13.4743 | 12.26895 | 17.31613 | 17.14423 | 23.41546 | 22.75332 | 15.91769 | 15.6778 | 12.29183 | 11.57056 |
| Ogdhl | 38.67236 | 41.85694 | 35.68808 | 33.48615 | 29.65749 | 29.83569 | 28.84678 | 27.82946 | 24.70731 | 24.28219 | 18.0435 | 17.04282 |
| Fhdc1 | 0.527823 | 0.619639 | 0.796853 | 0.767842 | 1.214613 | 1.238172 | 1.268044 | 1.283721 | 1.699857 | 1.507886 | 1.239014 | 1.214954 |
| Serpinh1 | 40.53859 | 38.02048 | 87.1347 | 91.05936 | 129.4447 | 123.6694 | 168.1827 | 160.3399 | 187.4968 | 178.7844 | 188.6462 | 177.8795 |
| Rhof | 0.701793 | 0.806543 | 1.19887 | 1.10996 | 1.478506 | 1.493353 | 1.797677 | 1.861187 | 1.098546 | 1.141681 | 1.076562 | 0.948051 |
| Manba | 44.5134 | 44.22542 | 21.08232 | 19.03555 | 11.55466 | 10.18278 | 9.298002 | 10.73798 | 9.383713 | 9.698237 | 9.270192 | 10.00993 |

| | | | | | | | | | | | | |
|---|---|---|---|---|---|---|---|---|---|---|---|---|
| Etv5 | 96.99145 | 106.6268 | 45.71402 | 43.51586 | 35.29078 | 34.4039 | 35.49812 | 29.86524 | 28.41935 | 27.57565 | 22.42499 | 22.18227 |
| Smyd2 | 24.9461 | 22.49655 | 54.45393 | 55.24267 | 48.80788 | 48.5491 | 42.5867 | 45.11095 | 29.86501 | 28.1635 | 30.44617 | 30.94884 |
| Nanos3 | 2.761395 | 2.200902 | 1.176666 | 1.763004 | 0.967571 | 1.594685 | 1.45104 | 0.898821 | 1.16691 | 1.183923 | 1.585704 | 1.718705 |
| Actb | 631.7439 | 820.0361 | 899.7038 | 809.3032 | 761.3373 | 679.7096 | 798.3019 | 659.874 | 786.791 | 745.8954 | 1326.009 | 1293.541 |
| Gabrp | 1.720668 | 1.933514 | 0.97279 | 0.91964 | 0.982222 | 0.760389 | 0.853006 | 0.743015 | 0.653477 | 0.734277 | 0.485936 | 0.469759 |
| Gramd4 | 4.522551 | 4.57711 | 5.414307 | 5.400648 | 6.799373 | 6.695528 | 7.204914 | 8.156667 | 10.05087 | 10.5691 | 9.531932 | 9.133053 |
| Prkch | 0.581029 | 0.499966 | 2.248064 | 1.885168 | 2.737985 | 2.423354 | 3.645794 | 3.688552 | 3.638829 | 3.640841 | 4.551483 | 3.65585 |
| Pik3c2a | 6.88356 | 8.097085 | 9.707279 | 10.23071 | 10.37509 | 10.00591 | 11.40597 | 10.98538 | 14.63825 | 16.15028 | 16.86276 | 19.5758 |
| Dcaf12l1 | 5.169683 | 5.714301 | 15.03119 | 14.56812 | 12.62518 | 12.01165 | 13.10291 | 12.28775 | 7.399476 | 7.449305 | 10.54745 | 9.487585 |
| Atp5l | 222.5132 | 165.2597 | 149.4042 | 139.4181 | 151.2318 | 159.5988 | 126.2244 | 135.7403 | 96.76124 | 81.62093 | 96.64169 | 97.55374 |
| Ccnj | 3.723486 | 3.768963 | 5.372943 | 5.821089 | 5.869539 | 5.616428 | 5.765904 | 5.841512 | 8.733548 | 8.823241 | 10.06942 | 10.98173 |
| Atf7ip2 | 7.621632 | 8.151133 | 8.495825 | 8.01112 | 6.437969 | 6.240212 | 5.514066 | 5.820427 | 4.102892 | 4.653825 | 3.794102 | 4.101738 |
| Dynll2 | 42.62671 | 43.30665 | 74.72411 | 67.88564 | 83.70609 | 81.21824 | 95.29644 | 80.97634 | 86.56608 | 75.46453 | 146.1571 | 132.5856 |
| Foxd3 | 14.70307 | 13.04496 | 1.146412 | 0.839179 | 0.467606 | 0.514033 | 0.858218 | 0.64533 | 0.979626 | 1.003917 | 1.288042 | 0.83691 |
| Msrb2 | 8.930811 | 9.438582 | 5.127181 | 4.510465 | 4.549736 | 4.444442 | 4.425614 | 4.506451 | 2.887108 | 2.751388 | 2.316062 | 2.310002 |
| Pex2 | 6.739032 | 5.802273 | 9.877141 | 9.10666 | 10.94899 | 11.80675 | 12.01701 | 13.51722 | 15.5174 | 14.83663 | 16.67175 | 17.77768 |
| Bicc1 | 0.738294 | 0.776709 | 1.67501 | 1.670092 | 2.139841 | 2.298013 | 3.009437 | 2.735568 | 3.56606 | 3.257463 | 2.6887 | 2.535564 |
| Ifrd2 | 48.68381 | 49.5258 | 48.02078 | 46.62306 | 40.52911 | 41.23719 | 34.9325 | 34.6059 | 30.30919 | 27.48139 | 23.96055 | 23.91493 |
| Zfp385a | 10.90169 | 13.11137 | 18.04321 | 17.62381 | 23.71806 | 21.40487 | 30.03641 | 24.44617 | 38.07449 | 32.12409 | 36.00348 | 34.3563 |
| Aig1 | 0.921827 | 1.060971 | 2.059723 | 2.05656 | 2.360375 | 2.374739 | 2.981804 | 2.295618 | 3.432238 | 3.055328 | 4.356071 | 4.007226 |
| Tmem164 | 5.331387 | 5.863532 | 7.93331 | 8.448676 | 11.3835 | 11.03871 | 14.52379 | 13.78919 | 15.18442 | 16.44886 | 10.08822 | 9.964999 |
| Cblb | 3.364603 | 4.232235 | 4.225455 | 4.429631 | 5.297823 | 5.122518 | 6.233715 | 5.188512 | 8.169504 | 8.623626 | 6.583434 | 6.258653 |
| Tmx4 | 1.7117 | 1.770396 | 2.478261 | 2.548921 | 2.919902 | 3.125199 | 3.834259 | 3.048752 | 4.3334 | 4.291299 | 4.469867 | 4.960151 |
| Gnpda1 | 62.95737 | 63.67121 | 43.52723 | 39.95059 | 33.66958 | 33.26762 | 30.75219 | 28.84916 | 24.51449 | 23.79837 | 21.12943 | 20.53153 |
| Ndufb3 | 49.76591 | 40.55351 | 42.0055 | 42.23183 | 43.99739 | 45.90347 | 39.56986 | 37.53053 | 25.10987 | 23.59578 | 25.35569 | 25.57446 |
| Lrp12 | 2.95312 | 2.936267 | 4.138328 | 4.033749 | 4.905154 | 4.729301 | 6.002873 | 5.234449 | 8.14222 | 7.845296 | 7.681774 | 7.776885 |
| Amotl1 | 3.781866 | 4.531856 | 8.147624 | 8.043843 | 9.521336 | 9.026929 | 10.54287 | 9.842246 | 13.37188 | 14.25764 | 15.20174 | 15.64045 |
| Ubqln2 | 21.03266 | 24.61091 | 25.71872 | 25.01499 | 29.26254 | 26.02398 | 34.70437 | 31.12793 | 49.17831 | 47.08061 | 51.40797 | 48.63333 |
| Agtrap | 22.04933 | 21.9699 | 21.84025 | 20.34337 | 19.28777 | 18.19628 | 17.84172 | 15.75138 | 11.8827 | 11.47335 | 10.72263 | 10.46444 |
| Mgat1 | 5.09118 | 5.48985 | 5.787546 | 5.921153 | 6.399365 | 6.589244 | 7.682782 | 8.314409 | 9.905055 | 9.366411 | 12.45142 | 11.50746 |
| Amhr2 | 3.098226 | 3.121333 | 1.78499 | 1.817002 | 2.036867 | 2.093677 | 1.820765 | 2.17649 | 1.18362 | 1.186509 | 0.812044 | 1.120078 |
| 2210016L2 | 14.11142 | 13.52667 | 14.29048 | 14.37545 | 15.154 | 14.00369 | 13.25647 | 14.20215 | 9.791805 | 9.249189 | 7.046571 | 7.398184 |
| Hal | 0.764815 | 0.713958 | 2.918053 | 2.758024 | 2.783895 | 2.8262 | 2.479288 | 3.101563 | 1.572356 | 1.597292 | 1.709793 | 1.785572 |
| Fig4 | 28.91231 | 27.22247 | 28.74922 | 27.5687 | 25.59401 | 25.0795 | 24.79253 | 24.97759 | 18.58177 | 19.82786 | 13.98087 | 14.19795 |
| Phf20l1 | 5.359382 | 5.070541 | 6.294611 | 6.161311 | 6.63505 | 6.603147 | 6.366744 | 6.478479 | 8.253366 | 8.180331 | 10.86688 | 11.95996 |
| Ager | 2.638836 | 0.685875 | 1.200919 | 1.511548 | 1.997134 | 2.711483 | 1.938895 | 2.848 | 1.885825 | 1.634681 | 1.290621 | 1.46729 |
| Acbd6 | 24.79033 | 22.83581 | 45.27439 | 41.21577 | 47.11372 | 50.56856 | 54.62555 | 54.61528 | 32.0674 | 28.13818 | 33.60326 | 30.92904 |
| Slc35f2 | 17.60723 | 15.94811 | 18.35623 | 17.22529 | 13.15513 | 12.95393 | 10.55467 | 11.38826 | 8.680887 | 8.877856 | 9.635354 | 9.424491 |
| Hltf | 22.2708 | 22.9294 | 16.86695 | 16.82727 | 14.96334 | 14.172 | 13.09793 | 14.40536 | 11.2953 | 12.50074 | 8.259027 | 8.681486 |
| Metrn | 0.65106 | 0.758257 | 1.671681 | 1.752577 | 2.128852 | 2.407562 | 2.541854 | 2.75082 | 3.029467 | 2.780651 | 4.364245 | 3.396716 |
| Ubr7 | 82.5232 | 76.73848 | 63.65287 | 61.18056 | 56.33849 | 56.55145 | 47.42344 | 48.6569 | 35.49714 | 36.1226 | 31.31916 | 31.57259 |
| Ankzf1 | 7.682154 | 7.411321 | 6.482023 | 5.858903 | 5.701334 | 5.647109 | 4.772779 | 4.952098 | 4.088545 | 4.289154 | 3.043933 | 3.023619 |
| Lpgat1 | 6.659569 | 7.393318 | 9.733645 | 10.02403 | 11.95111 | 11.67788 | 15.70974 | 12.98516 | 16.87424 | 16.5071 | 18.84899 | 19.4219 |
| Stxbp4 | 2.973893 | 3.295751 | 4.359385 | 4.549411 | 5.256397 | 5.138104 | 6.422945 | 5.912223 | 7.827943 | 7.956413 | 4.491722 | 4.106491 |
| Atp7a | 4.578354 | 5.394592 | 6.172908 | 6.185596 | 6.608379 | 6.344008 | 8.413705 | 7.211793 | 11.14183 | 11.98428 | 10.01821 | 10.15779 |
| Pou4f1 | 1.301779 | 1.351566 | 1.934345 | 1.971718 | 2.410741 | 2.157082 | 2.464536 | 2.593769 | 3.128672 | 3.016395 | 3.743417 | 3.81851 |
| Elk4 | 2.694581 | 3.290544 | 3.262781 | 3.294591 | 3.117303 | 3.463336 | 3.505945 | 3.139979 | 4.906996 | 5.702944 | 5.602381 | 6.077856 |
| Zcchc12 | 0.338997 | 0.228814 | 0.614108 | 0.718181 | 1.008532 | 1.096051 | 1.382374 | 1.365268 | 0.895565 | 1.120629 | 1.088477 | 1.390107 |
| Ppp2r3a | 1.956487 | 2.222575 | 2.633672 | 2.60771 | 3.235757 | 2.964594 | 3.289179 | 3.418687 | 4.397114 | 4.881905 | 4.966761 | 5.04695 |
| Ino80d | 4.538863 | 6.578001 | 10.751 | 11.14099 | 7.862286 | 7.716373 | 7.223766 | 5.757712 | 6.716835 | 7.143668 | 8.929185 | 9.318012 |
| Zkscan2 | 0.80666 | 0.878402 | 1.161079 | 1.191806 | 1.197641 | 1.33773 | 1.358711 | 1.330372 | 2.032583 | 2.305916 | 1.506814 | 1.623334 |
| Ptar1 | 2.776422 | 3.514936 | 5.016731 | 6.109293 | 4.914454 | 4.610045 | 5.889406 | 4.927402 | 5.764616 | 5.939876 | 7.986001 | 8.915073 |
| Hist1h2bg | 7.433294 | 6.024249 | 1.671201 | 2.730977 | 4.145218 | 5.280344 | 2.359561 | 4.32317 | 2.618361 | 2.767905 | 2.240894 | 2.877713 |
| Prtg | 8.498656 | 11.23219 | 32.72901 | 36.96143 | 54.1625 | 52.24324 | 74.11601 | 53.32745 | 71.12327 | 71.45941 | 57.32638 | 60.7568 |
| Setd4 | 5.585395 | 5.77907 | 5.539932 | 5.378646 | 4.879456 | 4.053506 | 3.21288 | 3.653377 | 2.491802 | 2.782845 | 2.611189 | 2.801901 |
| Barx1 | 0.259127 | 0.271806 | 0.434211 | 0.827002 | 1.309733 | 1.345166 | 1.74667 | 1.57452 | 1.71342 | 1.431098 | 1.439 | 1.598121 |
| C1ql4 | 0.668294 | 0.461946 | 1.304718 | 1.02625 | 2.164571 | 1.982679 | 1.876694 | 1.823205 | 1.72269 | 1.859817 | 1.468847 | 1.458705 |
| Dcaf4 | 31.80444 | 32.29569 | 30.44229 | 28.46099 | 25.25574 | 23.41339 | 22.45876 | 22.54395 | 20.0472 | 19.25221 | 16.03331 | 14.73358 |
| Pir | 11.72105 | 10.70475 | 7.869844 | 6.715441 | 7.44937 | 7.795133 | 6.787529 | 6.608018 | 4.827779 | 4.61745 | 3.828822 | 4.03566 |
| Upk3bl | 1.223323 | 1.079941 | 0.613334 | 0.649557 | 0.306651 | 0.559312 | 0.314109 | 0.376947 | 0.298923 | 0.357872 | 0.187171 | 0.336414 |
| Dennd5b | 2.224527 | 2.96737 | 3.594461 | 3.875918 | 4.543975 | 4.494692 | 6.116952 | 5.224934 | 6.97642 | 7.482947 | 5.429049 | 5.490289 |

| | | | | | | | | | | | | |
|---|---|---|---|---|---|---|---|---|---|---|---|---|
| Vangl1 | 16.37846 | 16.77118 | 7.475328 | 7.60154 | 7.483237 | 7.253147 | 9.997734 | 9.735941 | 14.40068 | 13.70611 | 14.16477 | 14.0028 |
| Rdh10 | 1.433275 | 1.527277 | 2.270488 | 1.794274 | 2.067217 | 2.118102 | 2.229355 | 1.907609 | 2.019696 | 2.182862 | 3.783154 | 3.463588 |
| Bst2 | 84.43289 | 74.11324 | 42.00644 | 36.82557 | 39.6188 | 37.77056 | 36.15933 | 38.95367 | 27.82511 | 25.1735 | 22.62873 | 20.18698 |
| Poc1b | 5.191669 | 5.127253 | 6.171226 | 5.611204 | 7.864778 | 7.184709 | 8.606562 | 8.221104 | 10.97747 | 10.82555 | 9.761348 | 9.90475 |
| Fxyd6 | 40.95341 | 37.67609 | 12.77429 | 11.93921 | 13.86631 | 12.93119 | 13.77566 | 14.5233 | 21.01789 | 21.6415 | 20.42853 | 23.03252 |
| Pnldc1 | 13.78391 | 14.69755 | 12.67628 | 12.97046 | 9.103253 | 9.09108 | 7.48594 | 8.074792 | 6.811868 | 6.729651 | 8.463079 | 9.237868 |
| Apcdd1 | 1.418931 | 1.476484 | 2.167224 | 2.301824 | 2.675886 | 2.932976 | 3.584511 | 3.466055 | 4.834308 | 4.436154 | 4.601066 | 3.871246 |
| Maged1 | 63.16981 | 61.92399 | 94.61384 | 92.38701 | 132.5932 | 128.3515 | 156.2771 | 176.5804 | 177.0627 | 178.0138 | 153.5306 | 152.1874 |
| Cdhr1 | 1.387315 | 1.507752 | 0.379924 | 0.2356 | 0.224885 | 0.213721 | 0.206796 | 0.326752 | 0.296621 | 0.229879 | 0.387233 | 0.411694 |
| Actn2 | 1.345647 | 1.483152 | 2.560124 | 2.484566 | 2.440813 | 2.569658 | 2.56078 | 2.722118 | 3.719871 | 4.242774 | 4.566972 | 4.784542 |
| Zkscan1 | 2.745124 | 3.471794 | 2.998869 | 3.506227 | 3.276322 | 3.451458 | 3.976153 | 3.630909 | 5.991327 | 6.643717 | 5.563852 | 5.677106 |
| Dzip1 | 3.582585 | 3.922738 | 4.412867 | 4.51943 | 5.570591 | 5.475191 | 6.606085 | 7.23529 | 8.640511 | 8.696409 | 7.848163 | 8.195678 |
| Hspbap1 | 16.60462 | 17.39146 | 13.37323 | 12.4269 | 9.522127 | 8.988574 | 7.805494 | 7.872446 | 7.522167 | 7.279548 | 6.251669 | 6.46131 |
| Rbm43 | 2.535051 | 2.885679 | 1.864249 | 1.878154 | 1.816994 | 1.684472 | 1.55797 | 1.646802 | 1.151133 | 1.130411 | 2.193452 | 2.172104 |
| Mcam | 26.69216 | 25.57369 | 12.22456 | 12.35299 | 12.67735 | 12.8517 | 13.80871 | 14.73906 | 7.400296 | 7.672442 | 10.77308 | 10.96345 |
| Snhg4 | 35.34071 | 33.27638 | 26.89957 | 28.2215 | 24.93745 | 26.23925 | 17.57277 | 20.00569 | 19.37737 | 18.46676 | 13.14023 | 14.69723 |
| Sync | 0.337425 | 0.643911 | 0.898315 | 0.62704 | 0.976247 | 0.782667 | 0.864362 | 0.905158 | 1.075585 | 0.997914 | 1.039639 | 1.20397 |
| AI314180 | 7.661435 | 8.24526 | 11.81654 | 12.00728 | 13.47455 | 13.61624 | 15.79478 | 14.42269 | 20.75997 | 21.99537 | 21.27615 | 23.03698 |
| Glt1d1 | 13.48073 | 13.19853 | 9.648291 | 9.52713 | 8.308089 | 7.849924 | 6.552702 | 6.930221 | 5.524583 | 5.720357 | 5.311356 | 4.966673 |
| Prr24 | 1.571832 | 1.449082 | 2.960384 | 2.633511 | 3.356415 | 3.372779 | 3.45069 | 2.87511 | 2.537468 | 1.971623 | 3.967593 | 3.781298 |
| Iqgap3 | 8.303534 | 10.03581 | 22.4379 | 20.86398 | 15.95003 | 16.04197 | 13.95543 | 12.9223 | 10.24973 | 10.87955 | 12.08164 | 12.25815 |
| Gnmt | 1.617268 | 1.366845 | 1.254147 | 1.454101 | 1.26493 | 1.241898 | 1.239048 | 0.81508 | 1.106208 | 0.758442 | 0.43203 | 0.792864 |
| Zfp12 | 2.913065 | 2.914671 | 3.789619 | 3.590477 | 3.905066 | 3.784097 | 4.637858 | 4.459717 | 6.303707 | 6.527639 | 6.698011 | 6.882509 |
| Tmem127 | 5.664472 | 5.595618 | 6.809882 | 6.938418 | 8.068121 | 7.863237 | 9.360768 | 9.225345 | 10.48219 | 10.56964 | 13.1099 | 13.29929 |
| Fggy | 3.64739 | 3.386089 | 3.285225 | 3.315882 | 3.654117 | 3.199542 | 2.817932 | 3.132703 | 2.338168 | 2.189185 | 1.82482 | 1.73021 |
| Rab25 | 2.386579 | 1.988779 | 0.85179 | 0.951822 | 0.968208 | 0.568827 | 0.518016 | 0.693685 | 0.660875 | 0.652773 | 0.525382 | 0.496715 |
| Gm3264 | 0.665387 | 0.548564 | 1.150289 | 1.146572 | 1.073502 | 0.908286 | 0.732944 | 0.797059 | 1.269518 | 1.114714 | 1.565263 | 1.525077 |
| Man2a1 | 8.223655 | 10.00093 | 36.84783 | 36.63623 | 38.41259 | 38.09047 | 43.83947 | 39.85479 | 29.72854 | 31.61917 | 21.32048 | 20.83757 |
| Chd3 | 3.442815 | 3.807473 | 5.757315 | 5.660088 | 7.97242 | 7.463295 | 10.36264 | 9.804587 | 11.47643 | 10.82284 | 9.381966 | 9.804848 |
| Tmem8 | 10.55233 | 11.08071 | 5.262135 | 4.685214 | 5.704714 | 4.868119 | 5.275332 | 5.790737 | 4.030239 | 4.080064 | 3.22848 | 3.028732 |
| Anxa8 | 2.004597 | 2.224909 | 0.977284 | 0.8281 | 0.761102 | 0.836857 | 0.799356 | 0.69802 | 0.455871 | 0.500832 | 0.573507 | 0.437821 |
| Nav1 | 0.565138 | 0.723151 | 1.110422 | 0.880997 | 0.867861 | 1.08433 | 0.883883 | 0.683775 | 1.45658 | 1.306188 | 0.938095 | 0.878539 |
| Ero1lb | 0.994311 | 1.408356 | 2.779368 | 2.647512 | 2.881083 | 2.818295 | 2.786829 | 2.864899 | 1.674581 | 1.901788 | 1.462963 | 1.500141 |
| Oxld1 | 3.418817 | 2.895251 | 2.653147 | 2.359864 | 3.270224 | 2.615975 | 2.079241 | 2.882355 | 1.679201 | 1.509298 | 2.181915 | 2.46931 |
| Cuedc1 | 6.896374 | 7.223905 | 14.92805 | 14.54043 | 18.37106 | 18.34506 | 19.32428 | 20.088 | 26.45529 | 23.76899 | 28.52579 | 27.76335 |
| Nap1l5 | 0.681429 | 0.783565 | 1.228417 | 1.220592 | 1.659546 | 1.800027 | 2.094271 | 2.327262 | 2.1444 | 1.713184 | 1.776624 | 2.307782 |
| B4galnt1 | 2.428335 | 2.778354 | 3.875185 | 3.95994 | 5.022876 | 4.87738 | 5.704095 | 5.94837 | 7.768648 | 7.550059 | 6.561581 | 7.461646 |
| Trio | 6.132671 | 7.613545 | 8.152644 | 8.693397 | 9.575842 | 9.808085 | 11.41161 | 10.21013 | 12.75374 | 12.72574 | 16.17677 | 16.57053 |
| Slco3a1 | 0.747265 | 0.603012 | 1.124011 | 1.210648 | 1.641342 | 1.508058 | 1.504284 | 1.409383 | 1.817341 | 1.827138 | 2.532681 | 2.307032 |
| Rad54b | 15.8438 | 17.50582 | 14.65416 | 15.7443 | 13.52648 | 13.60552 | 12.42853 | 14.02747 | 10.35087 | 11.82646 | 7.646171 | 8.262174 |
| Cdon | 3.02104 | 3.595613 | 8.018876 | 8.345275 | 10.13379 | 9.732214 | 12.911 | 11.48445 | 14.44751 | 15.89901 | 14.73656 | 14.60809 |
| Lrrc48 | 2.133733 | 1.669241 | 0.780711 | 1.158866 | 1.138599 | 1.189078 | 0.635279 | 1.211086 | 0.52558 | 0.635831 | 0.617397 | 0.651536 |
| 0610009B2 | 28.3391 | 22.28678 | 25.0037 | 24.45624 | 23.21407 | 22.25532 | 18.98624 | 16.49806 | 14.18833 | 13.89415 | 12.10955 | 12.83977 |
| Tead1 | 18.89982 | 24.94214 | 30.75416 | 31.07257 | 29.92042 | 28.28096 | 33.11908 | 26.99891 | 38.01268 | 39.0755 | 48.74385 | 51.39294 |
| Lmbrd1 | 2.777767 | 2.60313 | 3.082592 | 3.596434 | 4.187423 | 4.26355 | 5.044684 | 4.482315 | 5.792878 | 6.159855 | 6.111155 | 6.844387 |
| Pkd1 | 2.755376 | 3.339903 | 3.440531 | 3.577052 | 3.63004 | 3.830981 | 3.841482 | 3.469554 | 4.049248 | 4.346374 | 6.563979 | 6.602469 |
| Dnajc6 | 7.807403 | 8.28036 | 1.941201 | 2.060803 | 1.717803 | 1.547172 | 1.274291 | 1.100787 | 1.6047 | 1.749076 | 1.060464 | 1.083028 |
| Phlda3 | 28.97882 | 27.39139 | 38.74749 | 35.78819 | 44.35095 | 40.86901 | 53.73725 | 48.07551 | 49.34083 | 42.48301 | 77.23413 | 68.09825 |
| Scamp1 | 5.595884 | 5.896473 | 7.537845 | 7.717294 | 10.85676 | 10.42776 | 13.66056 | 13.08405 | 14.52131 | 13.95321 | 14.54623 | 14.68396 |
| Rps27 | 3747.92 | 2501.503 | 1271.491 | 1758.547 | 2468.358 | 2740.397 | 1941.11 | 2301.45 | 1842.019 | 1598.609 | 1597.912 | 1440.501 |
| Rapgefl1 | 0.374206 | 0.471457 | 0.985162 | 1.020804 | 0.991286 | 1.073351 | 1.259417 | 1.11979 | 0.876332 | 0.917824 | 0.598539 | 0.588714 |
| Mmp11 | 15.98556 | 14.17702 | 11.23196 | 10.66856 | 9.778167 | 10.1734 | 7.834371 | 9.443063 | 6.124638 | 5.610581 | 6.750163 | 6.010308 |
| Fam120c | 0.364086 | 0.342169 | 0.888008 | 0.73668 | 1.186014 | 1.145485 | 1.432809 | 1.400459 | 1.055837 | 1.289147 | 0.944756 | 1.006576 |
| Chst8 | 0.373408 | 0.437655 | 0.567005 | 0.690354 | 0.773075 | 0.964073 | 1.025756 | 1.126233 | 0.762188 | 0.592483 | 0.845946 | 0.633697 |
| Slc1a5 | 16.34162 | 16.67418 | 32.47038 | 32.35674 | 39.29021 | 37.72889 | 49.6926 | 46.11575 | 24.75165 | 24.26075 | 28.33316 | 27.37903 |
| 9430020K0 | 2.091156 | 2.191608 | 2.551708 | 2.551503 | 3.359029 | 2.950079 | 3.357621 | 3.300892 | 3.804875 | 4.0266 | 4.973574 | 4.847591 |
| Wbp1 | 10.00548 | 9.144027 | 8.938251 | 9.580577 | 13.46943 | 13.09679 | 13.25505 | 16.53863 | 20.12887 | 18.19338 | 16.7918 | 15.05014 |
| 1110007C0 | 14.21217 | 11.64356 | 9.31841 | 10.98522 | 12.52809 | 12.31391 | 10.2575 | 10.78409 | 7.163212 | 6.484518 | 7.753002 | 8.299083 |
| Arap2 | 1.91095 | 2.123504 | 3.200903 | 2.977232 | 3.04359 | 3.017973 | 3.395102 | 3.313194 | 3.422663 | 3.580571 | 5.015684 | 5.653037 |
| Atrnl1 | 1.581139 | 1.988336 | 2.312483 | 2.445012 | 3.023611 | 2.984054 | 3.724297 | 3.446843 | 4.321841 | 4.642126 | 4.213824 | 3.97223 |
| Ston1 | 3.523578 | 3.846342 | 6.284712 | 6.956233 | 6.765386 | 6.851774 | 8.275061 | 7.499875 | 8.492391 | 8.522863 | 12.29514 | 13.00528 |

| | | | | | | | | | | | | |
|---|---|---|---|---|---|---|---|---|---|---|---|---|
| Npr1 | 3.002746 | 3.361464 | 10.01815 | 9.071758 | 10.12397 | 10.22826 | 11.68541 | 11.67643 | 7.569784 | 7.748187 | 6.169939 | 6.152868 |
| Mtss1l | 3.28562 | 3.910004 | 4.899101 | 5.356067 | 6.259913 | 5.793119 | 6.799535 | 6.209865 | 8.085979 | 8.019642 | 10.06504 | 10.1589 |
| Grik5 | 4.739865 | 5.489416 | 6.913265 | 7.071932 | 9.392095 | 9.322496 | 11.74379 | 12.16883 | 12.64082 | 12.93476 | 7.368464 | 6.820687 |
| Zdhhc7 | 7.203875 | 7.333789 | 11.10815 | 9.802936 | 12.88359 | 12.31628 | 14.41737 | 15.85692 | 15.91463 | 15.83378 | 20.28671 | 18.58947 |
| Atp6v0e2 | 3.655047 | 4.075653 | 6.417582 | 5.43895 | 6.125578 | 5.559823 | 7.113019 | 6.344453 | 8.705518 | 8.178009 | 10.48123 | 10.31133 |
| Spsb2 | 2.12395 | 2.110973 | 2.65703 | 3.052599 | 3.716264 | 3.066877 | 3.584905 | 2.268906 | 2.902484 | 2.559582 | 4.675607 | 4.301209 |
| H2-M3 | 0.463214 | 0.77954 | 1.114635 | 1.222347 | 1.39993 | 1.535269 | 1.464538 | 1.597001 | 1.223437 | 1.607395 | 0.887896 | 0.848028 |
| Sptbn2 | 1.334839 | 2.01402 | 2.567229 | 2.381132 | 2.418331 | 2.267448 | 2.544877 | 2.226627 | 3.810505 | 3.91638 | 4.119335 | 4.219684 |
| Sos2 | 2.643409 | 3.239274 | 5.482734 | 5.574502 | 6.676812 | 7.278896 | 8.103648 | 7.706698 | 9.816291 | 10.55559 | 9.161325 | 9.768129 |
| Gnas | 1.619716 | 2.028639 | 2.69251 | 2.500985 | 2.705061 | 3.098099 | 3.469324 | 3.087267 | 4.216198 | 3.473816 | 5.903986 | 4.735165 |
| Timm8b | 87.4372 | 66.64144 | 62.54335 | 64.16822 | 66.63049 | 70.0889 | 52.53653 | 50.69116 | 45.89874 | 37.04317 | 40.12646 | 39.08917 |
| Lemd2 | 8.654485 | 8.541638 | 11.06702 | 9.132787 | 11.20523 | 11.42642 | 11.24093 | 11.19753 | 13.09335 | 12.54984 | 18.74152 | 17.27803 |
| Slc1a6 | 0.366238 | 0.407256 | 0.664226 | 0.677381 | 0.925775 | 0.914904 | 0.967775 | 1.076889 | 0.831064 | 1.05709 | 0.557272 | 0.569375 |
| Zeb2 | 2.367548 | 2.482153 | 6.599441 | 6.77329 | 9.71194 | 10.31691 | 11.90885 | 11.2012 | 12.40833 | 12.80671 | 9.984762 | 10.98508 |
| Leprel2 | 1.286435 | 1.514076 | 1.752317 | 2.085275 | 2.900854 | 2.421694 | 3.236069 | 3.085699 | 4.010165 | 3.942651 | 3.682347 | 3.441163 |
| Mbnl3 | 1.093425 | 1.225818 | 1.726142 | 1.873813 | 1.853519 | 1.815487 | 1.903988 | 1.552491 | 2.103754 | 2.141285 | 2.618908 | 2.935038 |
| Lama5 | 14.38065 | 18.35646 | 30.58987 | 29.08771 | 23.12213 | 22.26566 | 22.87043 | 20.42924 | 17.49018 | 17.70593 | 16.42932 | 15.38751 |
| Neurod1 | 3.123054 | 3.151703 | 1.392576 | 1.428043 | 0.723204 | 0.761685 | 0.668493 | 0.755644 | 0.916859 | 0.779364 | 1.283294 | 1.25596 |
| Mgat4b | 8.598144 | 9.646957 | 12.57274 | 10.94237 | 15.80777 | 15.31185 | 17.97341 | 18.29148 | 20.7442 | 20.27303 | 20.72458 | 20.67846 |
| Ankrd37 | 3.453324 | 2.615647 | 1.080663 | 1.283446 | 1.847544 | 1.577314 | 1.545165 | 0.960598 | 1.298319 | 1.590971 | 0.727093 | 0.841906 |
| Pycr2 | 69.4269 | 60.10142 | 55.35048 | 50.48928 | 59.11999 | 59.08504 | 47.79485 | 50.38652 | 31.6937 | 31.32947 | 32.68805 | 31.27622 |
| Gse1 | 8.431257 | 11.4073 | 20.59809 | 21.97569 | 26.41103 | 25.95385 | 30.37843 | 25.81734 | 15.38206 | 15.00385 | 15.75697 | 13.73852 |
| Magee1 | 0.584124 | 0.458415 | 1.488401 | 1.523536 | 1.974823 | 1.816433 | 2.459268 | 2.415965 | 2.619446 | 2.877887 | 2.537604 | 2.336084 |
| Nox4 | 0.724544 | 0.923295 | 1.019785 | 1.029991 | 1.300111 | 1.229762 | 1.313962 | 1.93636 | 1.56076 | 1.666244 | 1.640401 | 1.94551 |
| Sema4f | 0.473208 | 0.542494 | 1.311936 | 1.202502 | 1.452926 | 1.253813 | 1.468778 | 1.250248 | 0.750337 | 0.743215 | 0.753918 | 0.663928 |
| Kctd1 | 0.892417 | 0.973873 | 2.219629 | 2.026065 | 2.639968 | 2.257708 | 3.101217 | 2.780021 | 3.448839 | 3.243541 | 3.52161 | 3.824824 |
| Pdhb | 155.0411 | 127.6792 | 123.5212 | 120.5449 | 111.9689 | 107.5557 | 89.91567 | 101.5296 | 61.06293 | 63.86698 | 65.57392 | 66.3107 |
| Ldhb | 375.1447 | 333.7509 | 95.07589 | 98.32286 | 82.13298 | 73.80914 | 57.40442 | 71.45075 | 58.83432 | 62.272 | 59.51659 | 52.10831 |
| Zc3hav1l | 4.992831 | 6.233329 | 11.66975 | 12.03263 | 13.75161 | 13.55988 | 16.90054 | 14.13327 | 22.27095 | 22.70423 | 22.07501 | 22.41134 |
| Kit | 18.30748 | 21.39551 | 38.68711 | 37.49334 | 39.20055 | 35.72286 | 36.54039 | 36.45814 | 32.50865 | 34.40534 | 21.42754 | 19.87306 |
| Fbxw4 | 2.601798 | 2.532868 | 3.055946 | 3.013852 | 3.445652 | 3.43726 | 3.609608 | 3.286135 | 4.480386 | 3.968382 | 5.228512 | 5.68487 |
| Ajuba | 39.64631 | 41.91892 | 63.79818 | 59.90938 | 56.70507 | 55.63041 | 57.12351 | 54.62698 | 61.4013 | 57.25175 | 101.0656 | 103.0256 |
| Sox13 | 13.34773 | 14.37642 | 18.9988 | 19.7395 | 26.95405 | 26.5313 | 33.62498 | 33.64329 | 40.27162 | 37.22342 | 31.0926 | 29.71087 |
| Igf2r | 12.37315 | 16.04059 | 23.64532 | 24.07327 | 21.72381 | 20.88029 | 22.7426 | 18.34182 | 26.35183 | 27.21689 | 33.95994 | 34.58527 |
| F2rl1 | 47.01393 | 46.78111 | 24.98668 | 25.16781 | 26.45506 | 26.37451 | 28.97946 | 26.29254 | 13.37843 | 13.98804 | 25.18465 | 24.7774 |
| Prrg3 | 1.09046 | 1.404445 | 1.590586 | 1.761366 | 2.591217 | 2.258356 | 2.572893 | 2.449637 | 3.236826 | 3.190415 | 2.90826 | 3.32072 |
| Tnfsf9 | 0.540189 | 0.706264 | 0.622098 | 0.747902 | 1.081016 | 0.923853 | 1.19177 | 1.409823 | 1.199526 | 0.913781 | 0.715117 | 0.979292 |
| Lmtk3 | 1.352766 | 2.034482 | 3.740877 | 3.501662 | 3.557653 | 3.446157 | 3.115111 | 3.915643 | 2.406819 | 2.149453 | 2.191039 | 2.077679 |
| Hpcal1 | 12.2114 | 11.58479 | 14.1035 | 13.70176 | 18.68512 | 19.90261 | 24.09382 | 24.64951 | 27.76559 | 25.81271 | 24.64786 | 25.34954 |
| Dusp6 | 38.06548 | 39.43944 | 64.07795 | 62.66123 | 90.52401 | 89.54078 | 112.9317 | 109.5745 | 115.8191 | 109.4275 | 111.2622 | 117.9746 |
| Agpat6 | 16.26981 | 18.17351 | 22.12227 | 20.60159 | 23.78871 | 22.37503 | 27.34404 | 28.30563 | 33.41796 | 32.92334 | 38.32951 | 38.78173 |
| Lrpap1 | 20.27489 | 19.05022 | 24.12678 | 22.66395 | 23.01328 | 21.9124 | 23.79516 | 25.23492 | 30.95488 | 27.45149 | 40.97865 | 41.24528 |
| Htatip2 | 10.76915 | 9.515837 | 10.73863 | 10.06868 | 9.950136 | 9.344468 | 8.357934 | 8.861559 | 5.286677 | 5.384024 | 6.088866 | 5.350719 |
| Sirt3 | 12.2722 | 10.81707 | 9.281425 | 8.195223 | 7.168111 | 6.646351 | 6.202794 | 5.838337 | 4.239945 | 4.355363 | 4.493635 | 4.836759 |
| Clcn4-2 | 2.020124 | 2.329763 | 3.234386 | 3.588273 | 4.034279 | 3.257826 | 3.830194 | 4.064271 | 5.447236 | 5.511647 | 5.773172 | 6.128666 |
| Tcp11l1 | 2.62089 | 2.688417 | 3.262574 | 3.709588 | 4.353722 | 4.146827 | 5.095925 | 5.000069 | 7.271584 | 6.977671 | 4.687399 | 4.62969 |
| Bex2 | 24.89582 | 20.52973 | 32.55207 | 31.52743 | 37.44579 | 38.30497 | 41.93702 | 40.3275 | 46.74902 | 36.18645 | 59.20019 | 59.29763 |
| Ypel2 | 1.926022 | 2.24881 | 2.675832 | 2.620306 | 2.833835 | 3.075667 | 3.784902 | 3.896727 | 4.544298 | 4.925176 | 3.845252 | 3.934518 |
| Zbtb46 | 1.563774 | 1.692556 | 2.288011 | 2.361884 | 2.787842 | 2.700219 | 3.328908 | 3.751839 | 3.96288 | 4.438661 | 3.752224 | 4.155072 |
| Rsg1 | 0.839573 | 0.806486 | 1.694851 | 2.095603 | 1.491272 | 1.949974 | 1.498681 | 1.365819 | 1.569977 | 1.409577 | 1.023148 | 1.115569 |
| Magi3 | 2.727638 | 2.894455 | 5.165852 | 5.240281 | 6.099955 | 5.965537 | 6.973492 | 7.015947 | 7.053018 | 7.647647 | 9.469363 | 9.840525 |
| Rrp9 | 69.18073 | 65.90881 | 66.52284 | 59.19064 | 49.59504 | 50.02817 | 43.22185 | 42.44512 | 34.27614 | 31.52104 | 45.2012 | 40.04971 |
| Npm3 | 104.0536 | 87.44595 | 83.62747 | 79.90367 | 72.09205 | 63.97664 | 55.6522 | 58.79498 | 47.2755 | 42.56995 | 50.59517 | 50.11877 |
| Itga3 | 6.376173 | 8.155172 | 38.78146 | 37.36894 | 33.55029 | 31.34258 | 32.21859 | 29.62818 | 29.39161 | 29.56927 | 62.14752 | 55.49337 |
| Gap43 | 5.450279 | 5.63876 | 6.656963 | 6.569257 | 7.607748 | 7.097502 | 12.143 | 12.04682 | 15.21092 | 12.3276 | 11.05899 | 11.39651 |
| Homer3 | 7.198933 | 7.192255 | 8.448103 | 8.259005 | 10.1308 | 10.04326 | 11.39234 | 12.81297 | 16.67704 | 15.49777 | 14.98123 | 14.00049 |
| Gm6880 | 0.674396 | 0.70853 | 3.045875 | 3.694516 | 5.492776 | 5.939878 | 5.481019 | 6.802082 | 3.748755 | 4.09752 | 4.636463 | 3.626188 |
| Zbtb33 | 3.573528 | 3.986152 | 5.269924 | 5.231747 | 5.786914 | 5.818375 | 6.395796 | 6.130603 | 8.417293 | 9.810828 | 9.020812 | 9.362142 |
| Dnajb14 | 1.319185 | 1.705625 | 1.725703 | 1.865503 | 1.886547 | 1.774145 | 2.284361 | 2.027594 | 2.124291 | 2.131148 | 3.268815 | 3.326074 |
| Pitpnm1 | 1.655979 | 1.915476 | 2.288902 | 1.942434 | 2.368683 | 2.13496 | 2.513931 | 2.747478 | 3.476715 | 3.607111 | 1.55371 | 1.926932 |
| Moxd1 | 0.646161 | 0.6235 | 1.296796 | 1.342366 | 2.053132 | 2.058484 | 2.227035 | 2.214432 | 2.162684 | 2.26455 | 2.074295 | 2.512552 |

| | | | | | | | | | | | | |
|---|---|---|---|---|---|---|---|---|---|---|---|---|
| Slc16a6 | 4.224447 | 3.791133 | 6.547896 | 6.64055 | 6.968061 | 6.689801 | 6.901537 | 7.487181 | 5.836017 | 5.470313 | 10.03649 | 10.23177 |
| Marveld2 | 1.206115 | 1.231443 | 1.763774 | 1.875033 | 1.711214 | 1.881199 | 1.880895 | 1.515793 | 1.539738 | 1.819702 | 3.04961 | 2.83481 |
| Clic4 | 42.00139 | 45.63805 | 55.20824 | 53.97787 | 57.73486 | 54.9217 | 67.62391 | 64.17344 | 74.40188 | 76.60554 | 100.0417 | 100.9001 |
| Ifit2 | 0.770231 | 0.737326 | 1.190772 | 1.09103 | 1.613026 | 1.508211 | 1.578834 | 1.338008 | 2.198996 | 2.039327 | 1.280276 | 1.744284 |
| Ifngr1 | 2.996984 | 2.653396 | 8.331974 | 8.205055 | 8.496305 | 9.050146 | 8.534766 | 8.831961 | 5.729275 | 4.842063 | 6.407324 | 6.169924 |
| Ick | 3.923141 | 4.256452 | 5.525158 | 5.321747 | 6.081702 | 6.297841 | 7.605988 | 6.837165 | 9.937778 | 9.944033 | 8.016558 | 8.458934 |
| Ddc | 7.065717 | 6.897661 | 10.23259 | 9.608395 | 11.95742 | 11.54542 | 17.04503 | 17.95256 | 15.31891 | 15.98372 | 14.40857 | 15.86908 |
| Spry4 | 28.7289 | 37.42038 | 13.18495 | 13.2206 | 9.780492 | 9.721275 | 10.38027 | 9.4102 | 8.004961 | 8.530479 | 7.191673 | 7.078729 |
| Hps6 | 1.570278 | 1.577626 | 1.891092 | 1.611811 | 2.285525 | 1.820031 | 2.611499 | 2.329447 | 2.294042 | 2.530022 | 3.211753 | 3.008231 |
| 1700001K1 | 0.486364 | 0.73301 | 0.730578 | 0.852074 | 0.885956 | 0.778211 | 1.255056 | 1.129919 | 0.806099 | 0.67942 | 0.943108 | 1.268006 |
| Hist1h2bh | 2.638177 | 2.717061 | 0.807536 | 2.869929 | 2.755949 | 3.451955 | 3.067969 | 2.536621 | 2.401204 | 2.54447 | 2.661108 | 1.849756 |
| Pbxip1 | 3.613451 | 3.942411 | 4.693059 | 4.347679 | 5.161611 | 5.268744 | 6.13946 | 6.539837 | 6.794148 | 7.077451 | 8.527995 | 8.11016 |
| Man1a2 | 6.439656 | 7.115664 | 10.48681 | 10.45288 | 11.8923 | 11.46036 | 13.93816 | 13.34439 | 16.26562 | 16.38343 | 19.75273 | 19.4984 |
| Tlcd2 | 2.951245 | 2.630021 | 4.022555 | 3.901506 | 5.453338 | 5.577085 | 6.921626 | 7.275984 | 7.745127 | 7.227607 | 3.632719 | 4.496625 |
| Ankrd44 | 0.69021 | 0.782659 | 5.084887 | 5.304384 | 4.01438 | 3.864657 | 3.390519 | 2.906486 | 2.947634 | 2.845283 | 3.939661 | 3.874883 |
| 1700007J1( | 0.726919 | 1.317278 | 0.882501 | 1.747566 | 1.288278 | 1.822743 | 1.357652 | 1.138984 | 1.518157 | 1.759228 | 1.617227 | 2.123253 |
| Glce | 7.294006 | 7.818275 | 9.356342 | 9.380449 | 10.60374 | 10.01968 | 11.83033 | 10.60066 | 15.16437 | 14.73519 | 16.27748 | 17.48596 |
| Lag3 | 1.046445 | 1.11145 | 0.5507 | 0.716968 | 0.612996 | 0.731503 | 0.568415 | 0.51528 | 0.374719 | 0.44182 | 0.402622 | 0.392431 |
| Ptpn6 | 10.12131 | 9.444013 | 6.316735 | 6.24383 | 7.944774 | 7.630469 | 7.869256 | 7.980401 | 5.752842 | 5.757034 | 4.39161 | 4.282583 |
| Bcl2l11 | 7.754237 | 9.151017 | 15.18117 | 14.69205 | 14.6296 | 13.23459 | 14.8892 | 13.46638 | 19.79059 | 18.93834 | 24.51567 | 24.66082 |
| Mfap5 | 0.525543 | 0.452604 | 1.417822 | 1.245461 | 1.018313 | 0.817356 | 0.868136 | 1.053616 | 0.628763 | 0.668454 | 0.732014 | 0.934923 |
| Apol7b | 0.172967 | 0.163524 | 0.770598 | 0.760865 | 0.802263 | 0.774366 | 1.183615 | 1.166376 | 0.741206 | 0.626108 | 0.886399 | 0.632458 |
| Slc12a8 | 2.692069 | 3.084677 | 1.390885 | 1.421923 | 1.189596 | 1.11357 | 1.152882 | 1.302686 | 0.835614 | 0.763837 | 0.829194 | 0.809306 |
| Pafah1b2 | 17.16221 | 17.99437 | 19.45633 | 19.90201 | 20.78104 | 20.59992 | 23.57723 | 21.93614 | 35.59562 | 36.2841 | 36.5143 | 37.03544 |
| Map1lc3a | 12.382 | 13.43992 | 17.13195 | 16.91018 | 17.06789 | 18.21385 | 20.15181 | 17.65945 | 21.70596 | 19.57875 | 35.39101 | 31.46249 |
| Slc43a3 | 3.360038 | 2.898292 | 7.311629 | 7.26461 | 10.2833 | 10.35986 | 12.87604 | 12.33815 | 13.49729 | 13.51286 | 9.240951 | 9.589251 |
| Hist1h4d | 11.63732 | 12.48245 | 2.977506 | 6.168067 | 6.526628 | 9.512481 | 6.795864 | 8.824846 | 5.425462 | 6.582826 | 6.355886 | 5.114168 |
| Fam83g | 0.801259 | 1.186263 | 2.339759 | 2.22465 | 2.242133 | 2.064232 | 2.428231 | 2.138348 | 2.007807 | 1.811333 | 3.801681 | 3.368437 |
| Gm2694 | 2.176348 | 1.69762 | 1.617451 | 1.726649 | 1.976582 | 2.004859 | 2.012835 | 2.207655 | 1.547633 | 1.452964 | 1.008015 | 1.187479 |
| Pltp | 11.14621 | 13.06694 | 7.720474 | 5.934705 | 6.035112 | 5.662249 | 5.63569 | 5.60047 | 4.065388 | 4.554036 | 3.985892 | 3.192575 |
| Folr1 | 23.70766 | 23.61833 | 69.60979 | 66.76212 | 59.4978 | 63.46347 | 56.17352 | 61.26949 | 40.51386 | 35.92166 | 42.97168 | 44.66667 |
| Disp1 | 1.889058 | 2.308486 | 3.939974 | 3.732141 | 3.891223 | 3.77293 | 4.765655 | 4.258103 | 5.735059 | 5.91512 | 7.507912 | 6.936144 |
| Dars2 | 24.08386 | 21.95968 | 21.96147 | 21.68689 | 20.0753 | 19.366 | 18.00539 | 19.89079 | 14.91077 | 16.12416 | 11.02008 | 11.67175 |
| Piwil2 | 3.061344 | 3.238365 | 7.875851 | 8.218578 | 5.835715 | 5.434703 | 6.402761 | 6.343703 | 4.2619 | 4.423614 | 4.994108 | 5.11776 |
| Irx2 | 0.913553 | 1.130476 | 1.846436 | 2.24251 | 3.45257 | 2.964353 | 3.40235 | 3.481555 | 3.935135 | 3.672245 | 4.703254 | 4.165915 |
| Gal3st1 | 0.375359 | 0.471383 | 1.761635 | 1.480965 | 1.533142 | 1.512198 | 1.494195 | 1.471612 | 0.880398 | 0.978932 | 0.871125 | 0.814535 |
| Pgm2 | 24.2019 | 19.64607 | 9.816626 | 9.926953 | 13.80982 | 13.13264 | 12.64325 | 15.32515 | 11.99332 | 13.80751 | 8.321855 | 8.257041 |
| Oprl1 | 0.506809 | 0.342529 | 1.368051 | 1.473481 | 2.201689 | 2.2538 | 2.449074 | 2.734669 | 2.286269 | 2.032746 | 1.799189 | 2.055452 |
| Nr5a2 | 12.46839 | 15.61996 | 6.355686 | 6.22848 | 4.784542 | 4.06638 | 4.016218 | 3.813732 | 4.157861 | 4.906554 | 3.562583 | 3.364284 |
| Adamts4 | 0.656217 | 0.719021 | 0.996512 | 0.909889 | 1.05493 | 1.173227 | 1.381065 | 1.345467 | 0.993535 | 1.057322 | 0.857556 | 0.727194 |
| Atxn1l | 5.003882 | 5.255186 | 6.891051 | 6.704398 | 7.217036 | 7.496211 | 8.927032 | 7.298463 | 10.10426 | 10.54725 | 12.66714 | 12.40043 |
| Mpzl2 | 5.162653 | 5.544193 | 3.785401 | 3.480266 | 2.675943 | 2.945142 | 2.361876 | 1.885544 | 2.094253 | 1.883317 | 3.115723 | 3.010853 |
| Kank3 | 15.19973 | 13.94643 | 9.536004 | 9.746574 | 7.898278 | 7.994429 | 6.646268 | 7.493079 | 5.778634 | 5.518475 | 4.809535 | 5.277244 |
| Arrb1 | 3.383331 | 3.855364 | 4.064218 | 4.301439 | 5.243871 | 4.670096 | 6.282487 | 5.491365 | 5.80716 | 6.334183 | 8.050873 | 7.943689 |
| Emilin3 | 0.562909 | 0.511105 | 1.054 | 1.156856 | 1.198624 | 1.026491 | 1.422071 | 1.360956 | 1.15561 | 1.02493 | 0.705308 | 0.73716 |
| Ccdc138 | 4.208963 | 4.532803 | 2.82935 | 3.036972 | 2.417997 | 2.480066 | 2.360268 | 1.947172 | 2.612332 | 2.47882 | 1.877597 | 1.64534 |
| Cers6 | 2.932333 | 3.721745 | 7.081168 | 7.187395 | 6.465625 | 6.536833 | 8.535509 | 6.306214 | 11.09902 | 11.63892 | 9.73259 | 11.46513 |
| Fam168b | 25.06475 | 25.21504 | 29.8839 | 29.39331 | 29.71979 | 28.89316 | 34.03721 | 31.35592 | 46.5834 | 47.96406 | 54.17686 | 53.29911 |
| Scube3 | 2.137851 | 2.786297 | 9.687059 | 9.700863 | 8.982523 | 8.121546 | 8.066523 | 7.136055 | 6.325479 | 6.867888 | 5.583751 | 5.259003 |
| Spata5l1 | 3.144233 | 3.437322 | 4.491942 | 4.085947 | 4.075403 | 3.533277 | 3.738672 | 4.299733 | 5.251952 | 4.943926 | 8.038448 | 6.515271 |
| Slc27a2 | 14.58815 | 13.40252 | 2.974361 | 2.944564 | 2.775934 | 3.037909 | 2.614512 | 2.698311 | 1.769315 | 1.841023 | 1.878937 | 1.647584 |
| Lrrc58 | 7.109405 | 9.740398 | 10.29525 | 11.01498 | 11.38122 | 11.12016 | 13.33437 | 10.57832 | 16.51387 | 18.21399 | 17.21548 | 19.50048 |
| Itm2c | 15.70855 | 13.9217 | 17.2799 | 17.77088 | 24.08253 | 24.09798 | 32.01429 | 29.27617 | 32.33724 | 32.74313 | 30.26208 | 28.83748 |
| Purg | 3.197682 | 3.895376 | 5.994369 | 6.140209 | 7.000199 | 6.135169 | 8.007218 | 8.193912 | 6.751463 | 7.012933 | 4.565331 | 4.44731 |
| Slc45a3 | 0.990166 | 0.763725 | 1.161442 | 1.069229 | 1.561664 | 1.278306 | 1.714808 | 1.908361 | 2.030213 | 1.968033 | 1.358482 | 1.35613 |
| Otud1 | 2.901681 | 3.084043 | 3.70968 | 3.739853 | 3.348754 | 3.510686 | 3.823249 | 3.193979 | 4.192715 | 3.849691 | 6.592805 | 5.875646 |
| Hist1h1a | 8.967618 | 9.537037 | 3.233674 | 5.167633 | 6.857236 | 8.527104 | 5.743862 | 8.030187 | 4.681086 | 6.570806 | 6.433709 | 4.635856 |
| Dstyk | 2.315606 | 2.492635 | 3.192065 | 3.605396 | 3.901696 | 3.465701 | 4.133013 | 4.103295 | 4.83591 | 5.309598 | 6.102619 | 6.373456 |
| Bend7 | 0.852291 | 1.127409 | 1.680556 | 1.700048 | 2.148097 | 2.067307 | 2.379297 | 2.687483 | 3.119974 | 3.095343 | 2.553803 | 3.126141 |
| Dmrt3 | 0.211005 | 0.335542 | 1.115658 | 0.917882 | 1.053215 | 1.148462 | 1.263121 | 1.117751 | 1.076956 | 1.474676 | 1.198364 | 0.802128 |
| Arhgap1 | 13.44397 | 14.22561 | 20.05719 | 17.71368 | 19.97659 | 18.83428 | 21.34981 | 19.85372 | 21.18269 | 19.11355 | 32.52065 | 32.55072 |

| | | | | | | | | | | | | |
|---|---|---|---|---|---|---|---|---|---|---|---|---|
| Tfpi | 15.40044 | 13.11151 | 7.409522 | 7.411292 | 6.858696 | 6.973269 | 7.541262 | 8.527732 | 10.60028 | 10.56588 | 11.8803 | 12.81023 |
| Wdyhv1 | 14.08175 | 13.91675 | 13.97684 | 13.59617 | 12.70923 | 12.43088 | 12.5203 | 13.57798 | 11.05094 | 10.81871 | 7.116796 | 7.404637 |
| Tmem65 | 5.917208 | 6.651465 | 8.521761 | 8.217838 | 9.754031 | 9.468984 | 9.921058 | 9.906254 | 11.34566 | 11.76749 | 14.57108 | 15.08195 |
| Dab1 | 5.277365 | 4.880978 | 1.798678 | 1.693664 | 2.553099 | 2.609589 | 3.095332 | 3.107897 | 2.377596 | 2.026159 | 1.74245 | 2.048636 |
| Casd1 | 5.223938 | 5.309623 | 9.511774 | 8.741026 | 10.22066 | 10.02136 | 11.57311 | 11.64035 | 15.28711 | 16.02842 | 12.67496 | 13.82889 |
| Zfp865 | 4.192338 | 4.927745 | 6.250658 | 5.524051 | 6.781469 | 6.759285 | 6.823681 | 6.933861 | 8.756163 | 8.55587 | 11.52918 | 10.12856 |
| Kcnk5 | 17.52047 | 17.27548 | 11.68932 | 12.25588 | 11.36838 | 10.4701 | 11.21719 | 11.21522 | 8.798092 | 9.15427 | 7.598464 | 6.68431 |
| Syne4 | 10.42634 | 9.961844 | 9.543445 | 9.32975 | 7.254055 | 6.760115 | 5.605177 | 7.194358 | 6.402554 | 6.103522 | 4.98288 | 5.092756 |
| Abhd14a | 19.89021 | 20.39205 | 16.40765 | 15.56132 | 16.39149 | 15.43546 | 13.98121 | 14.2315 | 12.5619 | 12.61538 | 9.46105 | 8.495271 |
| Gpc1 | 6.994743 | 8.221327 | 10.6582 | 10.93686 | 14.4817 | 13.0188 | 16.32943 | 14.72453 | 17.73964 | 17.57041 | 22.57342 | 20.03373 |
| Trim35 | 17.1433 | 18.01296 | 22.12392 | 23.00262 | 27.75596 | 26.74033 | 30.77641 | 31.09182 | 41.81852 | 41.74153 | 44.84919 | 42.12402 |
| Gnb5 | 3.644959 | 3.324858 | 4.418936 | 4.22928 | 5.408551 | 5.415414 | 5.736895 | 6.509318 | 7.901558 | 7.742167 | 5.944699 | 5.772463 |
| Pim1 | 23.01977 | 23.9126 | 54.38706 | 53.42624 | 55.76523 | 54.29101 | 49.31785 | 45.07271 | 28.96567 | 29.66943 | 35.3213 | 30.66355 |
| Anks6 | 0.364341 | 0.351171 | 0.845613 | 0.966923 | 1.634605 | 1.491245 | 1.852768 | 1.648591 | 1.589445 | 1.515066 | 2.037468 | 1.769121 |
| Mob3c | 3.019503 | 3.095322 | 1.518019 | 1.422505 | 1.23558 | 1.264099 | 1.350967 | 1.283807 | 1.584217 | 1.591845 | 2.009291 | 2.312633 |
| 4930506M( | 1.201127 | 1.262519 | 2.119941 | 2.19516 | 3.196783 | 3.031177 | 3.668627 | 3.870181 | 4.248002 | 4.015963 | 3.467228 | 3.806023 |
| Pom121 | 14.66217 | 21.06252 | 28.9039 | 28.14332 | 27.76797 | 25.63689 | 30.03872 | 23.62672 | 33.98504 | 33.78019 | 47.92845 | 43.15555 |
| Pde1b | 5.081929 | 4.724229 | 2.338514 | 2.124501 | 2.222631 | 1.998727 | 2.222864 | 2.030801 | 2.415237 | 2.590904 | 3.349446 | 3.650306 |
| Tmem91 | 1.022193 | 1.539786 | 0.795 | 0.829067 | 0.68745 | 0.935144 | 0.508168 | 0.644031 | 0.417 | 0.635091 | 0.311601 | 0.512348 |
| Slc43a2 | 2.551037 | 3.105779 | 4.787899 | 4.776022 | 4.95107 | 4.297845 | 4.957417 | 4.486647 | 3.898399 | 3.768529 | 7.233123 | 6.877813 |
| Usmg5 | 204.07 | 131.1732 | 126.9664 | 122.5576 | 135.5399 | 147.9696 | 103.3329 | 128.7814 | 100.0131 | 81.07933 | 117.2552 | 111.8397 |
| Furin | 3.766503 | 4.354086 | 5.138708 | 5.191487 | 5.946074 | 6.105092 | 6.638917 | 5.466193 | 7.357551 | 7.511727 | 10.48274 | 9.47414 |
| Zdhhc14 | 0.479259 | 0.418828 | 0.799887 | 0.639753 | 0.885032 | 1.167475 | 1.213848 | 0.827509 | 0.991419 | 0.866706 | 0.694129 | 0.864383 |
| Cachd1 | 4.21315 | 5.012962 | 5.362059 | 5.621148 | 6.571147 | 6.48012 | 8.972864 | 8.519811 | 9.485319 | 9.081426 | 10.60952 | 10.25722 |
| Esco2 | 65.12475 | 61.99513 | 43.71451 | 41.08121 | 34.89216 | 33.22619 | 29.76536 | 33.73851 | 24.81931 | 27.11615 | 20.73622 | 22.51627 |
| Nipsnap1 | 53.47982 | 53.98401 | 45.32111 | 41.03225 | 39.06994 | 39.1585 | 36.38548 | 36.11151 | 30.57365 | 30.27134 | 21.86968 | 22.49025 |
| Hs6st1 | 19.11817 | 23.61577 | 26.69135 | 25.92993 | 25.75732 | 23.79466 | 27.09449 | 23.32643 | 32.37915 | 32.0506 | 44.44194 | 42.55535 |
| Lxn | 1.871792 | 1.990882 | 3.235499 | 3.180293 | 3.674933 | 4.130472 | 4.316488 | 3.751053 | 4.960037 | 4.310386 | 5.982285 | 5.797573 |
| Hoxd13 | 0.267705 | 0.332432 | 1.183747 | 1.108303 | 1.559053 | 1.687377 | 1.561809 | 2.019645 | 1.540101 | 1.377942 | 2.08213 | 1.521043 |
| Nr4a1 | 13.4009 | 15.78721 | 9.305435 | 8.515145 | 6.739801 | 7.00193 | 5.518888 | 5.891538 | 5.067731 | 4.672846 | 7.190671 | 7.017511 |
| Rhox2a | 0.456349 | 0.235674 | 1.485296 | 1.792266 | 1.975289 | 2.344566 | 2.419258 | 1.650665 | 1.60021 | 1.28664 | 1.341127 | 1.313028 |
| Lrp11 | 17.26976 | 17.36391 | 13.30919 | 13.04732 | 12.54313 | 12.43793 | 11.05017 | 12.0282 | 8.661448 | 8.870924 | 7.955442 | 7.161813 |
| Psmb10 | 14.57865 | 13.67094 | 11.01128 | 11.03616 | 11.29683 | 12.78314 | 11.13437 | 11.4769 | 7.096817 | 7.017627 | 7.515792 | 7.243327 |
| Gal3st3 | 1.664588 | 1.952103 | 3.399199 | 2.743035 | 2.910806 | 2.871639 | 3.100012 | 2.845709 | 2.973996 | 3.473919 | 1.99569 | 1.907178 |
| Spsb1 | 2.640874 | 2.64903 | 9.413614 | 9.503288 | 12.67761 | 13.06663 | 17.61125 | 17.3099 | 16.78197 | 16.6555 | 16.05949 | 16.77718 |
| Gm13251 | 18.65344 | 15.38856 | 12.91977 | 13.19699 | 10.83331 | 10.76825 | 8.500038 | 10.10475 | 7.964534 | 7.741856 | 6.602899 | 7.246022 |
| Dnajc25 | 1.682614 | 1.764083 | 2.286143 | 2.365282 | 2.349162 | 2.273781 | 2.821367 | 2.341607 | 3.119846 | 2.921376 | 4.227796 | 4.140913 |
| Rpp40 | 9.023374 | 8.744481 | 7.950524 | 7.956523 | 7.196686 | 6.977674 | 6.296495 | 6.420149 | 5.594003 | 5.409228 | 4.260361 | 4.370283 |
| Pdlim4 | 1.427366 | 1.692604 | 1.896518 | 2.105031 | 3.060003 | 2.74187 | 3.526032 | 3.83147 | 3.587735 | 3.322121 | 3.11723 | 2.496632 |
| 4632415L0 | 4.231003 | 4.192383 | 4.751675 | 4.945127 | 5.235526 | 5.554743 | 5.323715 | 6.570966 | 6.926737 | 6.114169 | 8.537692 | 9.000137 |
| Wdr81 | 2.509296 | 2.887713 | 6.469587 | 6.152618 | 6.449533 | 6.274756 | 7.874623 | 7.503619 | 4.388645 | 4.50278 | 4.504453 | 4.123826 |
| Hyi | 2.422893 | 2.103234 | 2.812309 | 2.837321 | 3.76411 | 3.821079 | 3.9813 | 4.990477 | 5.252263 | 4.747746 | 5.241682 | 5.160423 |
| Brdt | 0.429048 | 0.338406 | 1.379033 | 1.156964 | 1.659995 | 1.403082 | 2.150736 | 1.876659 | 1.731489 | 1.535676 | 2.230261 | 2.10424 |
| Mdfic | 0.92021 | 0.794703 | 1.413728 | 1.800567 | 2.060859 | 1.990267 | 2.28374 | 2.269218 | 2.42616 | 2.74462 | 2.820248 | 3.274338 |
| Asxl3 | 0.227857 | 0.28483 | 0.752753 | 0.765735 | 0.671437 | 0.78655 | 0.886766 | 0.681149 | 1.110871 | 1.121494 | 0.988534 | 1.238411 |
| Reck | 2.730004 | 3.124169 | 4.572582 | 4.755507 | 5.653201 | 5.731116 | 6.080163 | 7.108879 | 4.866798 | 5.681004 | 3.500981 | 3.910871 |
| Abcd4 | 10.20435 | 10.75976 | 10.56608 | 9.592106 | 9.746454 | 9.505171 | 9.028077 | 9.180534 | 6.72632 | 7.025509 | 5.03441 | 5.277779 |
| Zfp750 | 1.345728 | 1.392332 | 0.509121 | 0.682832 | 0.661011 | 0.699918 | 0.652471 | 0.594539 | 0.548422 | 0.38518 | 0.70134 | 0.590077 |
| Ppp2r2c | 1.702445 | 2.037737 | 2.794114 | 2.728452 | 4.675566 | 3.803717 | 4.526571 | 4.780718 | 5.154834 | 4.957553 | 4.771781 | 4.265332 |
| Atn1 | 7.368188 | 9.477968 | 17.66018 | 17.93018 | 18.75287 | 17.11869 | 18.91014 | 16.48225 | 22.87062 | 21.10647 | 32.73346 | 29.94773 |
| Trim44 | 9.295155 | 9.857849 | 18.84947 | 18.46731 | 20.06828 | 18.75053 | 21.97803 | 21.70484 | 26.71831 | 26.92288 | 32.72864 | 33.50656 |
| Zfp316 | 0.840795 | 0.942033 | 1.568184 | 1.514769 | 2.232961 | 1.996812 | 2.613392 | 2.747686 | 2.742903 | 2.730397 | 2.763783 | 2.458962 |
| Tnks1bp1 | 4.516447 | 5.240685 | 8.772007 | 8.811109 | 7.691652 | 7.699519 | 7.456921 | 6.641827 | 7.637807 | 7.232397 | 12.59896 | 12.0475 |
| Zfp672 | 1.334337 | 1.361049 | 2.40563 | 2.669091 | 2.158016 | 2.394951 | 2.704385 | 3.119473 | 3.19271 | 2.984476 | 4.847177 | 4.344095 |
| Tigd5 | 0.784401 | 0.542138 | 1.057233 | 1.307116 | 1.071161 | 1.133193 | 1.245167 | 1.119155 | 1.366199 | 1.521381 | 1.998332 | 2.029882 |
| Hmgn2 | 12.50588 | 10.87719 | 18.66556 | 17.89172 | 19.40299 | 21.50437 | 21.97218 | 17.64842 | 21.96298 | 18.54383 | 32.89334 | 32.0018 |
| Pim2 | 43.10459 | 40.30263 | 7.705463 | 7.682605 | 5.850472 | 6.205486 | 6.414746 | 5.660719 | 4.606489 | 4.548162 | 4.518326 | 4.236868 |
| Slc38a6 | 2.206912 | 2.411176 | 2.88534 | 3.153486 | 3.026241 | 3.448773 | 4.81527 | 4.893177 | 5.410188 | 4.681318 | 5.591268 | 5.717087 |
| Dnajc1 | 2.946342 | 3.157796 | 3.888697 | 3.813042 | 4.342023 | 4.386164 | 5.31006 | 5.200823 | 6.382266 | 6.549271 | 7.182219 | 6.902921 |
| Smad1 | 6.563141 | 7.550647 | 9.027112 | 9.212377 | 9.604462 | 10.10453 | 11.77634 | 11.52123 | 14.66185 | 14.68432 | 16.24964 | 16.67712 |
| C2cd2 | 1.313051 | 1.115054 | 2.129484 | 2.086141 | 2.371478 | 2.438723 | 3.226931 | 3.052974 | 3.272931 | 3.774847 | 3.328789 | 3.489007 |

| | | | | | | | | | | | | |
|---|---|---|---|---|---|---|---|---|---|---|---|---|
| Tmem150a | 8.587239 | 8.390102 | 10.1498 | 10.10372 | 13.322 | 13.29107 | 14.39514 | 14.36934 | 19.92765 | 17.61243 | 17.01811 | 18.27922 |
| Ntrk2 | 0.200472 | 0.165202 | 0.719463 | 0.63702 | 0.747864 | 0.715786 | 1.005595 | 1.152334 | 0.925353 | 1.016295 | 1.038595 | 1.043891 |
| Flt4 | 1.118956 | 1.229393 | 1.740924 | 1.619732 | 2.395534 | 2.384269 | 2.834184 | 2.929661 | 2.457671 | 2.609408 | 1.983691 | 1.770707 |
| Porcn | 5.657445 | 5.145629 | 15.23299 | 14.15118 | 20.66284 | 20.39927 | 24.28592 | 25.57849 | 19.16463 | 15.96771 | 19.99777 | 21.79105 |
| Gatm | 1.973851 | 1.842989 | 3.089657 | 3.512257 | 5.389628 | 5.013105 | 4.915096 | 5.1595 | 6.399971 | 6.345049 | 5.240157 | 5.851383 |
| Plekhg5 | 8.041956 | 10.32216 | 6.785542 | 5.832354 | 4.622684 | 4.431043 | 4.26082 | 4.421209 | 3.609728 | 3.424268 | 3.720569 | 3.229276 |
| Erbb2 | 2.679493 | 3.24237 | 5.825294 | 6.143859 | 7.329291 | 7.046375 | 8.771803 | 9.154605 | 9.554458 | 9.649712 | 11.29223 | 11.09403 |
| Hoxa3 | 0.801076 | 0.768989 | 9.702914 | 9.303479 | 14.59102 | 16.79886 | 18.55317 | 15.26662 | 13.31763 | 11.8789 | 16.19555 | 14.23681 |
| Epha4 | 9.22436 | 12.00136 | 4.580836 | 4.680754 | 4.703168 | 4.830289 | 6.921056 | 5.776917 | 8.246476 | 8.450114 | 8.560437 | 8.255615 |
| Zfp428 | 37.19864 | 30.97622 | 22.84707 | 23.19482 | 19.35083 | 20.40851 | 17.55466 | 18.79442 | 16.12928 | 12.85308 | 21.04435 | 17.2931 |
| Klf9 | 48.47709 | 50.02116 | 36.64265 | 34.89686 | 30.08449 | 28.06613 | 25.62292 | 26.7299 | 20.21068 | 22.34598 | 20.3827 | 19.33871 |
| Rtn4rl1 | 0.22824 | 0.383437 | 1.208719 | 0.931003 | 1.130353 | 0.917384 | 1.163481 | 1.076946 | 0.806751 | 0.597206 | 0.778683 | 0.647069 |
| Hp1bp3 | 12.58543 | 12.72214 | 17.77876 | 17.22718 | 18.58456 | 18.40401 | 21.40056 | 21.31066 | 28.51161 | 27.65278 | 31.42371 | 31.05832 |
| Efna2 | 3.566586 | 3.707743 | 6.666359 | 6.190442 | 7.862335 | 7.835442 | 8.990295 | 8.621257 | 11.6321 | 11.08141 | 11.92519 | 11.15924 |
| Mutyh | 14.89414 | 14.95734 | 12.65633 | 11.9387 | 11.58854 | 11.27176 | 8.606832 | 11.31038 | 9.517312 | 9.133794 | 6.477636 | 6.623621 |
| Adc | 9.657556 | 8.82272 | 5.712402 | 6.390157 | 7.165699 | 7.307954 | 6.771474 | 7.996711 | 4.084728 | 4.439224 | 4.191974 | 4.594141 |
| Dlg4 | 1.55191 | 2.014699 | 2.807792 | 2.982008 | 2.83345 | 2.880965 | 3.350845 | 3.701104 | 2.98627 | 2.750079 | 2.295987 | 2.054866 |
| Metrnl | 0.446675 | 0.289497 | 1.165369 | 1.003423 | 1.323615 | 1.140268 | 1.777026 | 1.80644 | 0.813648 | 1.172965 | 1.730957 | 1.474048 |
| Csrnp1 | 9.393298 | 9.836835 | 21.5215 | 18.89443 | 17.43188 | 17.39234 | 15.85633 | 14.28875 | 10.46073 | 10.49654 | 17.77398 | 16.85836 |
| Slc4a3 | 1.471782 | 1.392755 | 3.524195 | 3.535352 | 3.769195 | 3.895922 | 3.294037 | 3.530367 | 2.48734 | 2.164751 | 2.522485 | 2.179949 |
| Slc46a1 | 3.954201 | 4.226809 | 6.023609 | 5.323649 | 6.789638 | 6.654166 | 8.362098 | 8.624682 | 8.407234 | 8.091281 | 10.08027 | 9.580352 |
| Abtb2 | 0.770524 | 0.954048 | 1.393056 | 1.365808 | 1.489324 | 1.595988 | 1.707912 | 1.501228 | 1.510299 | 1.630404 | 3.039825 | 2.457606 |
| Eva1b | 4.008447 | 3.382525 | 11.36258 | 9.802921 | 10.43106 | 10.8542 | 12.32071 | 11.48785 | 7.181545 | 6.386945 | 7.804563 | 6.467288 |
| Fam46a | 0.528644 | 0.534777 | 0.798909 | 0.992232 | 1.248319 | 1.356339 | 1.734035 | 1.784069 | 1.676501 | 1.362771 | 1.306476 | 1.482138 |
| Gm6406 | 0.252446 | 0.382498 | 2.397537 | 1.889236 | 2.010648 | 1.54469 | 2.167319 | 1.649126 | 1.275504 | 1.050805 | 2.106671 | 1.552237 |
| Lif | 1.30247 | 1.304015 | 1.745488 | 1.763017 | 1.795794 | 1.695552 | 1.682614 | 1.746166 | 1.712357 | 1.761716 | 3.450551 | 3.048394 |
| Sh3glb1 | 12.91779 | 12.71448 | 16.49554 | 16.0851 | 18.51501 | 17.7552 | 20.08392 | 19.20257 | 25.70262 | 25.28725 | 28.13166 | 28.91835 |
| Pemt | 2.794192 | 2.675412 | 3.878216 | 3.708622 | 4.516328 | 4.615996 | 5.117996 | 4.999779 | 4.244485 | 3.571369 | 5.851942 | 6.417351 |
| Bcl6 | 0.307835 | 0.464931 | 0.739201 | 0.619048 | 0.619839 | 0.723652 | 0.940459 | 0.846062 | 1.018797 | 1.110821 | 0.863573 | 0.687727 |
| Clybl | 19.28312 | 15.81354 | 11.45538 | 11.14178 | 11.1674 | 10.85922 | 10.47178 | 12.48276 | 10.67969 | 9.646096 | 7.727582 | 6.956755 |
| Gnaq | 7.057325 | 7.581193 | 9.298395 | 9.543504 | 10.9161 | 10.43271 | 11.80434 | 10.11412 | 13.33662 | 13.5932 | 16.68078 | 17.12327 |
| Vegfc | 8.593342 | 9.154137 | 6.469077 | 6.232317 | 3.897658 | 4.017104 | 3.781926 | 3.687575 | 5.847903 | 5.411561 | 6.354869 | 6.615888 |
| Renbp | 24.25307 | 22.96285 | 21.32936 | 19.17313 | 18.32748 | 17.85378 | 15.89279 | 17.37942 | 12.26832 | 11.58308 | 10.59135 | 10.69023 |
| Mettl5 | 17.46592 | 14.72099 | 14.34181 | 13.93083 | 11.9219 | 11.95942 | 9.269235 | 10.33767 | 8.249226 | 8.030617 | 7.423929 | 7.769449 |
| Cep112 | 0.705264 | 0.818609 | 0.885595 | 0.84842 | 1.063047 | 1.28137 | 1.244638 | 1.277582 | 1.500857 | 1.520942 | 0.845386 | 1.117531 |
| Fam171b | 1.332734 | 1.437916 | 3.536186 | 3.611842 | 4.463063 | 3.973526 | 4.923647 | 4.691718 | 6.517592 | 6.474828 | 4.352868 | 4.597072 |
| Gxylt2 | 0.298712 | 0.266485 | 0.889456 | 1.002511 | 1.071977 | 0.960775 | 0.875686 | 0.949662 | 0.574425 | 0.60031 | 0.825808 | 0.559301 |
| Gcnt1 | 4.009292 | 4.037626 | 4.153655 | 4.720935 | 5.522324 | 5.638448 | 6.387848 | 6.206888 | 5.646016 | 6.365463 | 8.110532 | 8.456229 |
| Zfp964 | 0.469036 | 0.47531 | 0.87305 | 0.937192 | 1.57866 | 1.261904 | 1.653284 | 1.621813 | 1.615713 | 1.678456 | 1.025282 | 1.274287 |
| Fads2 | 42.58741 | 47.36287 | 53.98687 | 55.04851 | 74.22929 | 67.53447 | 86.56816 | 82.73222 | 95.55042 | 98.5688 | 70.54526 | 67.96261 |
| Mdga1 | 0.384587 | 0.439827 | 1.055877 | 0.923748 | 0.886708 | 1.022694 | 1.035289 | 1.208349 | 0.922937 | 0.883311 | 0.831627 | 0.675231 |
| Prkar2a | 16.32714 | 18.65189 | 21.442 | 21.87001 | 20.44966 | 20.14217 | 23.35572 | 21.22415 | 31.3701 | 30.10305 | 34.69236 | 36.03856 |
| 0610011F0 | 8.565877 | 7.739558 | 11.9826 | 13.14484 | 15.50588 | 17.76721 | 17.66933 | 18.13473 | 14.59538 | 14.78425 | 10.60006 | 10.13782 |
| Amigo3 | 0.515891 | 0.615504 | 0.715784 | 0.616827 | 0.895035 | 0.858629 | 0.636195 | 0.903062 | 0.602057 | 0.707786 | 1.157929 | 1.10326 |
| AU018091 | 45.51703 | 57.09398 | 39.3024 | 37.46763 | 30.08971 | 27.13084 | 27.57138 | 25.61832 | 22.88087 | 22.35387 | 21.98471 | 20.95474 |
| Lats1 | 5.095654 | 5.932867 | 7.052184 | 7.145229 | 7.190508 | 7.43439 | 7.187179 | 6.472493 | 9.323846 | 9.604668 | 10.97963 | 11.57035 |
| Plk3 | 12.92748 | 13.60838 | 53.0585 | 49.29142 | 53.13903 | 53.185 | 61.76776 | 58.50923 | 35.09465 | 32.75648 | 44.05688 | 41.36117 |
| Mfap4 | 3.716011 | 3.075585 | 1.517285 | 1.640084 | 1.73015 | 1.438465 | 2.500679 | 2.515465 | 2.621191 | 2.569092 | 1.484711 | 1.630371 |
| Lmx1b | 0.168487 | 0.254685 | 0.671467 | 0.725553 | 1.018093 | 0.941164 | 1.117688 | 0.998005 | 1.119151 | 1.295704 | 1.59311 | 1.227916 |
| Map7d1 | 17.31317 | 18.30374 | 26.58411 | 24.83555 | 31.51432 | 30.0606 | 38.54384 | 37.28668 | 36.81103 | 34.68005 | 49.23982 | 44.33664 |
| Prkcb | 1.751235 | 1.966486 | 2.629709 | 2.663731 | 3.378157 | 3.724894 | 3.64166 | 4.309294 | 4.250539 | 4.754795 | 3.510251 | 3.762413 |
| Zfp423 | 7.903532 | 10.02597 | 23.2816 | 22.62918 | 19.98919 | 19.18771 | 20.14605 | 17.25878 | 15.79157 | 15.29444 | 13.70479 | 12.68102 |
| Fsd1l | 4.155527 | 5.098403 | 8.508325 | 8.496837 | 6.319916 | 5.937372 | 5.804137 | 5.579864 | 4.818043 | 4.76405 | 5.375696 | 5.360621 |
| Pja2 | 7.961048 | 9.043672 | 10.61265 | 10.51821 | 13.6119 | 12.73798 | 15.16057 | 15.00933 | 18.42516 | 18.75774 | 18.006 | 18.38414 |
| Pnpo | 10.0713 | 9.311722 | 7.732963 | 7.146289 | 7.185922 | 7.237279 | 6.170653 | 6.630245 | 5.47928 | 5.164175 | 4.155373 | 4.058901 |
| Pigyl | 22.99286 | 17.96921 | 16.45343 | 14.39378 | 17.44132 | 17.48095 | 13.80897 | 12.54285 | 11.59697 | 9.805952 | 11.95471 | 10.58295 |
| Ccdc50 | 12.61041 | 14.02489 | 17.2878 | 17.37133 | 18.02502 | 17.08045 | 19.71973 | 18.47181 | 28.64736 | 30.43771 | 26.99162 | 30.3743 |
| E2f6 | 14.08928 | 13.30085 | 19.00681 | 20.01439 | 22.12238 | 21.42251 | 23.51228 | 24.02398 | 26.21693 | 26.86477 | 35.76731 | 35.65586 |
| Acy1 | 25.01843 | 22.9392 | 19.2055 | 18.84727 | 17.29191 | 17.88741 | 14.60724 | 14.37092 | 13.11927 | 12.81805 | 10.5367 | 10.58065 |
| Ephb6 | 0.777337 | 1.047257 | 1.352967 | 1.091179 | 1.622455 | 1.596177 | 1.968835 | 1.943452 | 1.751158 | 1.904267 | 1.683804 | 1.711569 |
| Bahd1 | 2.549494 | 3.157409 | 4.341729 | 4.398958 | 5.254411 | 5.199777 | 5.856116 | 5.774221 | 7.367775 | 6.734251 | 8.70748 | 7.832093 |

| | | | | | | | | | | | | |
|---|---|---|---|---|---|---|---|---|---|---|---|---|
| Enc1 | 43.33095 | 47.05084 | 33.43053 | 32.47026 | 25.80537 | 25.06357 | 23.50921 | 21.63477 | 19.0152 | 18.24566 | 25.07158 | 24.64066 |
| Cyp7b1 | 1.033344 | 0.906904 | 1.023406 | 1.142561 | 1.693199 | 1.700102 | 2.259527 | 1.774285 | 1.932492 | 2.032638 | 1.742686 | 1.830394 |
| Asap1 | 6.463363 | 7.27692 | 9.473208 | 9.553804 | 11.20892 | 10.25714 | 12.56629 | 11.06288 | 15.40211 | 15.16349 | 18.17885 | 16.99595 |
| Rhoq | 3.750565 | 4.137493 | 4.914471 | 4.905334 | 5.938996 | 5.906133 | 6.568894 | 6.650136 | 8.307235 | 8.719756 | 6.987172 | 7.869739 |
| Rpl35 | 53.15317 | 33.99169 | 33.18115 | 33.48157 | 32.5547 | 37.0093 | 22.61887 | 24.6721 | 22.17799 | 20.82479 | 29.74027 | 28.27389 |
| Vamp8 | 2.843262 | 2.17075 | 3.489745 | 3.52661 | 4.764544 | 5.038257 | 6.201419 | 5.73422 | 6.24224 | 5.098143 | 6.781754 | 6.262692 |
| Siah2 | 1.838202 | 1.852277 | 4.228059 | 3.973324 | 5.219513 | 5.522114 | 6.258256 | 5.412766 | 6.95314 | 6.305963 | 8.313662 | 7.055239 |
| Map3k14 | 0.55575 | 0.695416 | 0.912613 | 0.963231 | 0.972406 | 0.974137 | 0.962426 | 1.170022 | 1.126696 | 0.920662 | 1.690998 | 1.6346 |
| Nudt15 | 0.846779 | 1.030133 | 1.904471 | 1.296284 | 1.469357 | 1.4391 | 2.064648 | 1.7539 | 2.104144 | 1.850709 | 2.267604 | 2.301136 |
| Qsox1 | 3.884019 | 3.292161 | 5.736704 | 5.738166 | 8.779154 | 8.160098 | 9.958874 | 10.18606 | 9.999422 | 9.184762 | 10.56923 | 10.08096 |
| Il17rc | 1.436644 | 1.340685 | 1.868471 | 2.383465 | 3.016866 | 2.804775 | 2.892767 | 3.196409 | 2.111109 | 2.147638 | 2.074976 | 1.800851 |
| Garnl3 | 4.728244 | 5.325746 | 4.579454 | 4.025413 | 4.202454 | 4.284458 | 4.144963 | 4.568272 | 3.678192 | 3.369011 | 2.229431 | 2.575325 |
| Ldoc1l | 2.09145 | 2.221933 | 4.200745 | 3.974804 | 4.562436 | 4.209494 | 5.348907 | 5.040383 | 5.054344 | 5.169665 | 8.090945 | 7.049939 |
| Esrrb | 107.5169 | 115.1521 | 54.08483 | 49.24714 | 38.57346 | 36.58211 | 34.26152 | 36.07676 | 35.91383 | 35.94783 | 31.09529 | 27.77739 |
| Maml2 | 0.914701 | 0.800137 | 0.425261 | 0.575537 | 0.771234 | 0.68783 | 1.067743 | 0.848602 | 1.00532 | 1.020322 | 0.772287 | 0.616155 |
| Bdh1 | 5.97249 | 5.595863 | 4.320219 | 3.699677 | 2.981148 | 3.327416 | 2.790132 | 2.732162 | 2.660974 | 2.087023 | 2.130782 | 2.223261 |
| H2afy2 | 8.700622 | 9.418659 | 16.2126 | 14.93493 | 17.90458 | 16.72288 | 20.05646 | 18.94077 | 25.51367 | 21.36634 | 26.90848 | 26.46785 |
| Ntrk1 | 0.582865 | 0.760094 | 1.550813 | 1.250533 | 1.673837 | 1.444957 | 1.582522 | 2.046361 | 1.459608 | 1.335086 | 1.366673 | 1.154817 |
| Mcur1 | 2.094564 | 1.973307 | 2.236778 | 2.29299 | 2.919052 | 3.07699 | 3.331779 | 2.844348 | 2.832067 | 2.647565 | 4.429028 | 4.061014 |
| Ppp3ca | 5.616557 | 5.924906 | 6.779552 | 6.465018 | 8.279467 | 8.051705 | 9.070848 | 9.663013 | 10.81432 | 10.5752 | 12.13428 | 11.44989 |
| Greb1l | 2.81518 | 3.64723 | 14.49625 | 15.83561 | 18.62207 | 17.98772 | 24.43404 | 21.2779 | 26.57938 | 28.02219 | 16.11502 | 17.05486 |
| Lrfn3 | 1.112897 | 1.362261 | 1.68742 | 1.569589 | 2.244994 | 1.599388 | 2.193479 | 2.489968 | 2.611712 | 2.776206 | 2.123954 | 2.448704 |
| Gm973 | 0.566683 | 0.571002 | 1.236762 | 1.333146 | 0.907871 | 0.886246 | 0.698322 | 0.964719 | 0.805624 | 0.832632 | 0.742197 | 0.753866 |
| Crebrf | 4.481284 | 5.043577 | 3.890112 | 3.715797 | 2.930657 | 2.835516 | 2.784949 | 3.127822 | 2.446496 | 2.983172 | 1.76337 | 2.102615 |
| Klhl5 | 3.586136 | 3.187757 | 4.317404 | 4.454972 | 6.098022 | 5.355046 | 7.682895 | 7.064737 | 8.016315 | 7.871581 | 5.497473 | 5.817694 |
| Pitpnm2 | 2.491854 | 3.124515 | 4.070943 | 3.864129 | 3.904057 | 4.345536 | 4.383214 | 3.781996 | 4.811773 | 4.8007 | 6.600066 | 6.680707 |
| Cebpa | 0.543346 | 0.306211 | 1.403153 | 1.510927 | 2.362841 | 1.979274 | 3.001061 | 2.668969 | 2.812494 | 2.462387 | 3.459896 | 2.482795 |
| Rab11fip3 | 6.702808 | 7.277627 | 8.844019 | 9.512587 | 11.32208 | 11.32458 | 13.07971 | 13.0416 | 16.1771 | 16.44485 | 16.80488 | 16.79187 |
| H2-D1 | 8.217961 | 7.058514 | 11.81309 | 10.52362 | 13.77445 | 13.29756 | 12.49129 | 14.04675 | 14.14854 | 13.14895 | 20.20684 | 18.56365 |
| Cd99l2 | 2.410542 | 2.413952 | 2.97571 | 3.237631 | 4.308056 | 4.274372 | 5.151239 | 4.908026 | 5.46849 | 5.709996 | 4.695529 | 4.8249 |
| Ksr1 | 2.510602 | 3.079367 | 6.988153 | 6.36664 | 6.003125 | 5.624978 | 4.87536 | 4.827438 | 3.863953 | 4.000539 | 3.947752 | 3.610209 |
| Hipk1 | 8.09102 | 10.54262 | 11.71013 | 12.05457 | 11.97487 | 11.63449 | 13.50102 | 11.07478 | 17.73723 | 18.57947 | 18.97069 | 19.52907 |
| Mob3b | 1.769463 | 2.165685 | 1.912639 | 2.198949 | 2.83223 | 2.629833 | 3.355551 | 3.054522 | 4.229119 | 3.87343 | 2.55567 | 2.508083 |
| Pfkm | 26.2807 | 24.0641 | 18.11572 | 17.92048 | 18.95231 | 19.04179 | 19.3031 | 21.17156 | 17.38373 | 17.12374 | 11.96532 | 12.02595 |
| Prelid2 | 28.27477 | 23.24254 | 8.931974 | 8.849729 | 10.54664 | 10.46209 | 6.710591 | 9.515832 | 10.95696 | 12.60559 | 8.140186 | 7.164264 |
| Zfp276 | 1.749859 | 1.958484 | 3.193271 | 3.194738 | 3.060734 | 3.12952 | 3.103612 | 3.454325 | 2.507059 | 2.399359 | 4.178169 | 4.220799 |
| Rdm1 | 32.66639 | 31.04823 | 22.56587 | 21.29197 | 18.88922 | 17.91673 | 17.64371 | 19.01369 | 16.12378 | 15.62577 | 12.14439 | 12.10691 |
| Sorcs2 | 0.978792 | 1.102305 | 3.699661 | 3.759341 | 4.466422 | 4.520443 | 6.254793 | 5.075147 | 6.37054 | 6.328411 | 7.212077 | 6.611255 |
| Plekhb1 | 3.327906 | 3.58284 | 6.592707 | 6.67489 | 7.270575 | 7.281318 | 7.374908 | 6.544277 | 5.327682 | 5.447857 | 3.90873 | 4.143719 |
| Stoml1 | 21.51723 | 19.53722 | 16.68351 | 15.26811 | 14.6821 | 14.25802 | 12.49081 | 12.64256 | 9.192947 | 8.694176 | 11.31039 | 11.90381 |
| Dyrk1a | 9.704712 | 12.40194 | 17.23169 | 16.66736 | 15.03862 | 15.32108 | 16.87506 | 13.57428 | 18.23328 | 18.15797 | 23.3642 | 23.82073 |
| Tfeb | 11.73641 | 12.17955 | 7.138747 | 6.199727 | 5.951514 | 5.864809 | 5.110533 | 5.783343 | 3.143416 | 3.535397 | 4.170646 | 3.831466 |
| Wdfy2 | 3.019632 | 3.884575 | 5.199813 | 5.277116 | 4.635767 | 4.527297 | 5.204415 | 5.197096 | 7.160848 | 7.935692 | 7.714222 | 6.746637 |
| Dsc2 | 0.78394 | 0.731608 | 3.749575 | 3.61733 | 5.340532 | 4.825361 | 6.278655 | 6.194959 | 5.344145 | 5.295069 | 6.006597 | 6.339884 |
| 6030419C1 | 1.819059 | 1.675085 | 3.507551 | 3.312171 | 4.321068 | 3.947862 | 3.618886 | 4.469793 | 4.83204 | 4.387112 | 6.218016 | 5.804939 |
| 2700089E2 | 4.80478 | 4.808519 | 9.37253 | 9.68722 | 9.018169 | 9.175296 | 8.397731 | 8.190546 | 7.772583 | 7.451502 | 12.76484 | 13.05359 |
| Trim36 | 2.776323 | 2.758728 | 0.758011 | 0.785369 | 0.873083 | 0.9322 | 1.178334 | 1.021092 | 1.049681 | 1.253318 | 1.393941 | 1.375594 |
| Il4 | 2.230936 | 2.504946 | 3.61462 | 3.009558 | 4.127507 | 2.739429 | 4.706257 | 3.950779 | 3.771643 | 4.796787 | 2.523804 | 3.355293 |
| Rnf38 | 10.41943 | 12.29885 | 14.46283 | 13.86644 | 15.48786 | 14.88568 | 17.52747 | 15.70584 | 22.47547 | 24.27238 | 20.14587 | 20.62117 |
| Zfp651 | 1.146375 | 1.086877 | 1.673472 | 1.60335 | 1.793918 | 1.771668 | 1.672339 | 1.630352 | 2.662141 | 2.50549 | 3.154137 | 2.804364 |
| Mrpl36 | 47.15763 | 41.69489 | 40.31232 | 40.93164 | 39.18579 | 37.93995 | 32.77536 | 35.03302 | 25.23074 | 24.5246 | 23.43368 | 23.40488 |
| Hells | 120.0906 | 116.8347 | 113.5323 | 113.2941 | 93.26696 | 90.19105 | 76.7368 | 86.39422 | 70.16078 | 73.78563 | 60.37781 | 64.79203 |
| Traf1 | 0.337409 | 0.455502 | 0.754478 | 0.862987 | 0.898548 | 0.870962 | 1.094449 | 1.265563 | 1.057417 | 0.986802 | 1.033707 | 0.723995 |
| Asrgl1 | 9.281806 | 9.125033 | 7.413012 | 7.32626 | 7.2179 | 7.429657 | 7.040076 | 7.418295 | 7.292643 | 6.865453 | 4.486358 | 4.250673 |
| Slc35c1 | 2.782537 | 3.266227 | 3.979046 | 3.968505 | 4.286465 | 4.485644 | 5.273696 | 3.657992 | 5.282982 | 5.308231 | 6.508333 | 6.388728 |
| Plekhh3 | 0.54091 | 0.509026 | 1.221242 | 1.298153 | 1.568275 | 1.190015 | 1.092148 | 1.150207 | 1.114713 | 0.934695 | 1.868457 | 1.632443 |
| Tnk1 | 1.124531 | 1.385723 | 1.168519 | 0.979517 | 0.964048 | 0.932157 | 0.485257 | 0.725445 | 0.504175 | 0.637956 | 0.456826 | 0.560919 |
| Parvb | 29.15746 | 32.25333 | 23.69639 | 21.4504 | 19.2972 | 17.63169 | 15.69209 | 16.21274 | 14.36093 | 14.629 | 12.91003 | 12.28887 |
| Actn1 | 56.96544 | 68.33758 | 80.96945 | 81.76514 | 78.15925 | 73.51087 | 86.10248 | 79.51988 | 79.67588 | 77.89511 | 141.5358 | 128.2883 |
| Ap3b2 | 1.977612 | 1.77719 | 5.14705 | 5.553006 | 7.623513 | 7.837194 | 9.140005 | 9.548369 | 5.790753 | 5.612589 | 5.219134 | 5.472594 |
| Sgk1 | 76.12296 | 79.05443 | 39.121 | 38.10565 | 29.51368 | 27.93573 | 27.34472 | 27.33914 | 25.77345 | 27.96736 | 25.2513 | 21.86227 |

| | | | | | | | | | | | | |
|---|---|---|---|---|---|---|---|---|---|---|---|---|
| Fam43a | 13.27972 | 13.26636 | 25.89742 | 23.21159 | 22.42983 | 21.58537 | 22.04086 | 20.98731 | 18.6832 | 18.56294 | 33.61313 | 32.33915 |
| Jup | 33.47422 | 41.80199 | 23.62531 | 20.04366 | 16.69904 | 16.04419 | 17.41666 | 14.72548 | 12.77215 | 11.50555 | 18.13384 | 16.15655 |
| AA414768 | 1.794318 | 1.417349 | 4.978435 | 4.870076 | 4.455048 | 3.30639 | 4.262943 | 3.355639 | 3.068647 | 2.79607 | 3.739929 | 3.65741 |
| Skida1 | 2.362475 | 2.135374 | 3.402697 | 3.41646 | 5.173137 | 4.751025 | 5.194079 | 5.044424 | 6.62358 | 5.94898 | 4.866831 | 5.008272 |
| 2610008E1 | 2.770064 | 2.841514 | 3.905039 | 4.326478 | 4.357684 | 4.337701 | 4.777242 | 4.865134 | 6.642268 | 6.83452 | 6.170141 | 7.532713 |
| Armc7 | 1.739996 | 1.949819 | 3.817721 | 3.380709 | 3.325002 | 3.015444 | 3.543282 | 3.338762 | 2.198588 | 2.179627 | 4.27059 | 3.793851 |
| Asna1 | 219.4519 | 187.2484 | 173.698 | 177.1263 | 157.4351 | 152.6582 | 131.4793 | 142.5884 | 108.3976 | 101.7697 | 114.4394 | 112.5906 |
| Smox | 3.014164 | 3.06554 | 5.405991 | 5.374024 | 7.120572 | 6.826098 | 7.290994 | 7.787398 | 8.764023 | 8.699005 | 9.516112 | 9.34118 |
| Trim68 | 2.075791 | 1.986922 | 3.132481 | 3.440831 | 3.515689 | 3.396378 | 3.132426 | 3.786249 | 4.380642 | 4.270685 | 5.425858 | 5.903151 |
| Stk40 | 14.53667 | 14.98095 | 21.47944 | 21.27247 | 22.33095 | 21.04329 | 20.40075 | 19.87602 | 16.63377 | 16.65258 | 30.00312 | 28.93781 |
| Rit1 | 3.123278 | 2.472742 | 3.721449 | 3.364466 | 4.0429 | 4.329127 | 4.29828 | 4.972078 | 5.097225 | 5.195986 | 6.626154 | 5.846244 |
| Grtp1 | 30.25696 | 27.91484 | 25.15415 | 24.61349 | 24.89423 | 23.86382 | 22.68467 | 25.11504 | 19.82072 | 19.51106 | 13.29858 | 14.4536 |
| Plekhh2 | 2.119682 | 2.269319 | 2.770621 | 3.060093 | 3.941623 | 4.08555 | 4.412645 | 4.097297 | 4.560395 | 4.976025 | 2.47619 | 2.863819 |
| Dlgap4 | 11.68776 | 13.13553 | 14.82602 | 14.59327 | 16.98778 | 17.4279 | 18.60038 | 19.32265 | 22.11175 | 19.79584 | 25.24638 | 25.35521 |
| St6gal1 | 3.262279 | 3.668805 | 5.155457 | 5.448053 | 7.998363 | 7.791735 | 10.98366 | 9.464947 | 9.081569 | 8.724711 | 5.42708 | 5.650741 |
| Mcee | 11.14788 | 8.581504 | 7.278928 | 5.788451 | 7.092162 | 7.464964 | 5.755815 | 6.029019 | 5.678071 | 5.319175 | 3.409123 | 4.29699 |
| Zmat4 | 3.53636 | 3.311602 | 0.679391 | 0.515922 | 0.549289 | 0.635391 | 0.764891 | 0.806894 | 0.993155 | 0.895833 | 0.624779 | 0.61745 |
| Slc5a3 | 1.214049 | 1.766069 | 1.747646 | 2.010982 | 1.898532 | 1.799091 | 2.415046 | 1.893923 | 2.727223 | 2.972511 | 3.082293 | 3.123536 |
| Stat5b | 2.847465 | 2.752455 | 3.810668 | 3.798833 | 5.439764 | 5.355508 | 6.263292 | 6.55402 | 6.540003 | 6.594684 | 6.363129 | 6.188822 |
| Gm13375 | 1.560284 | 1.512078 | 1.672214 | 2.107455 | 2.402535 | 2.668555 | 2.646522 | 2.19518 | 2.455268 | 2.722718 | 3.363727 | 3.658402 |
| Schip1 | 32.80162 | 31.46211 | 13.28831 | 13.17227 | 12.73243 | 12.73399 | 12.45624 | 13.43623 | 12.13929 | 13.58847 | 21.96101 | 22.09809 |
| Batf3 | 0.737184 | 0.446785 | 2.361924 | 1.990182 | 4.04546 | 3.452897 | 3.329198 | 3.422877 | 2.101999 | 2.597584 | 3.809449 | 2.766433 |
| Gm13157 | 22.98991 | 22.06576 | 15.40117 | 16.51113 | 12.80684 | 12.0013 | 10.54186 | 11.764 | 10.89975 | 12.20744 | 9.488668 | 9.517256 |
| Muc1 | 1.088091 | 0.979033 | 2.171363 | 1.946314 | 1.391732 | 1.476042 | 1.269976 | 1.21244 | 1.236304 | 1.280278 | 2.296966 | 2.102974 |
| Tmcc1 | 2.68575 | 2.778113 | 3.960941 | 4.369274 | 5.156288 | 5.072672 | 5.83543 | 5.939767 | 6.675973 | 7.578447 | 6.930175 | 6.928896 |
| Ypel4 | 0.856405 | 0.955761 | 1.007526 | 1.376918 | 1.490022 | 2.074497 | 1.943153 | 1.912942 | 2.173257 | 2.38689 | 1.597694 | 1.445039 |
| Dyrk2 | 6.711426 | 7.227479 | 7.844527 | 8.827891 | 9.525837 | 9.951421 | 11.21341 | 11.05335 | 15.6529 | 15.29472 | 14.88378 | 15.17554 |
| Ngfrap1 | 140.1357 | 120.384 | 175.0832 | 170.5442 | 186.9672 | 192.1997 | 185.0239 | 163.3567 | 192.6827 | 156.1638 | 295.2644 | 270.3058 |
| Ech1 | 54.25405 | 43.16539 | 42.5008 | 41.36832 | 40.48729 | 39.97061 | 32.47774 | 37.3975 | 26.81501 | 24.65569 | 24.66512 | 23.90265 |
| Cpxm1 | 4.30932 | 4.158234 | 7.642885 | 7.297694 | 9.883629 | 9.192866 | 14.36898 | 12.20208 | 12.61642 | 12.62985 | 12.51316 | 11.89855 |
| Prkce | 0.797434 | 0.843507 | 1.376878 | 1.685719 | 1.861322 | 1.645534 | 2.181965 | 1.955918 | 2.711194 | 2.846841 | 1.988592 | 2.146134 |
| E130307A1 | 1.3086 | 1.96397 | 2.38009 | 2.308331 | 2.386062 | 2.309456 | 2.610765 | 2.625092 | 3.565683 | 3.992101 | 2.514716 | 3.294526 |
| Chn2 | 1.222646 | 1.448219 | 3.323983 | 3.379851 | 3.903504 | 4.060751 | 3.630599 | 4.681996 | 3.144102 | 2.832843 | 3.109746 | 2.707956 |
| 1700018B2 | 2.38465 | 1.971323 | 3.701191 | 3.70895 | 4.357327 | 5.745244 | 5.199594 | 6.412558 | 5.653308 | 5.53863 | 5.134878 | 5.082634 |
| 2700081O1 | 5.888549 | 5.667097 | 8.923481 | 8.84972 | 9.806565 | 10.04151 | 10.70646 | 10.66229 | 13.60818 | 12.76318 | 15.05101 | 15.29999 |
| Fam199x | 3.093236 | 3.652212 | 4.485802 | 4.241364 | 5.308051 | 5.031664 | 5.737568 | 4.636448 | 7.168232 | 7.326709 | 5.830074 | 6.614842 |
| Neo1 | 8.535457 | 10.53368 | 14.89668 | 14.99945 | 16.05495 | 16.23385 | 20.33634 | 17.27617 | 25.38604 | 25.90975 | 25.79254 | 25.70392 |
| Cbfa2t2 | 24.29134 | 29.9826 | 19.17765 | 18.77628 | 15.28276 | 14.19215 | 13.44642 | 12.81907 | 13.42093 | 14.92972 | 11.12914 | 10.86983 |
| Zdhhc21 | 4.356152 | 5.341058 | 7.185057 | 7.451051 | 6.909463 | 6.630403 | 6.816964 | 5.717325 | 7.402698 | 8.325653 | 9.328368 | 9.87381 |
| Tmem74 | 0.436649 | 0.547934 | 0.666072 | 0.87032 | 0.931076 | 0.842915 | 1.139364 | 1.305401 | 1.014688 | 1.315665 | 0.557307 | 0.761912 |
| 1810037I1 | 60.8082 | 49.8453 | 47.07998 | 42.47096 | 46.50959 | 44.65881 | 41.10063 | 42.72323 | 27.52858 | 25.86491 | 29.20289 | 27.87581 |
| Mboat1 | 0.745509 | 1.099725 | 1.427089 | 1.218942 | 1.418293 | 1.389621 | 1.380777 | 1.375588 | 1.568126 | 1.644774 | 1.809396 | 2.10457 |
| Sgpp1 | 1.686685 | 1.865684 | 2.918861 | 3.060181 | 3.860254 | 3.986627 | 4.339412 | 4.187583 | 4.791062 | 5.282908 | 5.297033 | 5.23051 |
| Tab2 | 24.79239 | 27.56569 | 29.53162 | 30.36761 | 33.05045 | 33.49015 | 37.10121 | 36.30594 | 51.29222 | 52.41415 | 44.22343 | 45.37144 |
| Slc36a4 | 0.779135 | 0.837956 | 1.988112 | 2.056712 | 2.846185 | 2.753571 | 3.109041 | 2.974484 | 3.760463 | 3.549627 | 3.025602 | 3.217631 |
| Rasl10b | 1.384837 | 1.27727 | 12.99005 | 12.78115 | 19.21737 | 19.11265 | 18.85172 | 18.2029 | 11.44748 | 11.07523 | 11.74669 | 11.46504 |
| Rgs22 | 1.233677 | 1.486469 | 0.937492 | 0.666943 | 0.71396 | 0.588409 | 0.535119 | 0.503067 | 0.545886 | 0.588352 | 0.511233 | 0.386489 |
| 4933426M | 14.17545 | 15.27265 | 18.42684 | 19.33504 | 22.72667 | 22.79552 | 29.72034 | 26.39027 | 33.3948 | 33.3564 | 28.77376 | 30.23572 |
| Ccnd1 | 24.62832 | 24.56645 | 15.36797 | 14.86512 | 10.76172 | 11.04584 | 10.28433 | 8.621539 | 8.670105 | 8.633814 | 8.850403 | 9.92539 |
| Dnajb3 | 0.536437 | 0.62506 | 1.439886 | 1.435695 | 1.725105 | 1.606345 | 2.408624 | 2.229643 | 1.758631 | 1.324074 | 2.757541 | 2.282361 |
| Gnao1 | 1.07147 | 1.260345 | 1.395096 | 1.577801 | 2.320499 | 2.141246 | 2.990299 | 2.423226 | 2.78825 | 2.719658 | 2.68105 | 2.523762 |
| Cabp1 | 5.236144 | 5.802101 | 6.698023 | 6.902314 | 6.136126 | 6.09122 | 4.521047 | 4.857292 | 4.466043 | 5.094567 | 10.5138 | 8.371781 |
| Ddhd1 | 3.385377 | 3.796247 | 4.238632 | 4.524904 | 4.995373 | 5.152341 | 5.69976 | 5.909817 | 6.987894 | 7.791025 | 7.395137 | 6.978811 |
| Rassf10 | 0.611278 | 0.580718 | 1.363241 | 1.066805 | 1.158376 | 1.167796 | 1.353055 | 1.836611 | 1.780596 | 1.796874 | 2.039926 | 1.691202 |
| Cbx2 | 7.426264 | 8.086858 | 13.37366 | 13.64365 | 16.18826 | 14.67956 | 18.01843 | 17.19814 | 23.60278 | 23.47975 | 24.4419 | 22.63863 |
| Tmem9b | 6.052859 | 4.64025 | 6.815539 | 7.61795 | 8.941128 | 8.784308 | 9.96908 | 10.85009 | 10.15228 | 10.07823 | 13.6528 | 13.10776 |
| Adnp | 1.067436 | 1.546846 | 2.812089 | 2.250966 | 2.110963 | 2.693326 | 2.418448 | 1.586943 | 2.250798 | 1.709707 | 2.898651 | 2.730445 |
| Hspb2 | 1.55981 | 1.159674 | 1.230644 | 1.033435 | 1.26961 | 1.028668 | 0.713149 | 0.836912 | 0.792841 | 1.0785 | 1.828797 | 1.43989 |
| Tceal5 | 0.328334 | 0.326314 | 1.680229 | 1.789939 | 1.852049 | 1.595009 | 1.972568 | 1.436539 | 1.069648 | 1.040505 | 1.147343 | 1.060123 |
| Bckdha | 40.02886 | 36.29619 | 36.02515 | 32.05638 | 26.33594 | 26.86895 | 23.11711 | 26.64525 | 18.83207 | 18.63875 | 19.99856 | 19.71211 |
| Acot13 | 36.13493 | 29.67007 | 19.53468 | 18.83832 | 19.77658 | 19.82249 | 17.08787 | 16.70574 | 20.77261 | 18.96805 | 11.57409 | 11.53031 |

| | | | | | | | | | | | | |
|---|---|---|---|---|---|---|---|---|---|---|---|---|
| Tap1 | 1.345459 | 1.483718 | 1.652388 | 1.596266 | 2.29122 | 2.697857 | 2.72313 | 2.744199 | 2.76473 | 2.578811 | 2.436112 | 2.707103 |
| Tspyl5 | 0.90428 | 0.643908 | 1.128469 | 1.140051 | 1.280122 | 1.250985 | 1.249606 | 1.216112 | 1.282349 | 1.249304 | 2.073661 | 1.958937 |
| Myo5b | 0.492743 | 0.593318 | 0.569436 | 0.628086 | 0.719853 | 0.61358 | 0.840762 | 0.903295 | 1.183879 | 0.971568 | 1.206261 | 1.053939 |
| Ggct | 4.111252 | 3.173831 | 2.823619 | 2.528221 | 2.512755 | 2.679585 | 2.242828 | 2.390695 | 2.087711 | 2.003822 | 1.701869 | 1.560101 |
| Pag1 | 0.50631 | 0.624631 | 0.88397 | 1.036386 | 1.453396 | 1.41343 | 1.931441 | 1.73708 | 1.553782 | 1.778849 | 1.407852 | 1.375257 |
| Alg8 | 4.336858 | 4.337315 | 5.57232 | 5.876099 | 6.353321 | 5.379642 | 6.823142 | 6.912137 | 8.823119 | 9.232709 | 9.368237 | 8.235646 |
| Kdm8 | 2.35681 | 2.336945 | 3.827636 | 3.301504 | 3.463951 | 3.524592 | 3.787719 | 3.144123 | 3.721051 | 3.098513 | 5.787047 | 5.316564 |
| Rab11fip1 | 3.10396 | 3.620086 | 5.67741 | 5.80142 | 6.067233 | 5.568219 | 6.652444 | 5.925318 | 7.010089 | 7.027261 | 9.450182 | 9.549618 |
| Ezh1 | 7.13953 | 8.664354 | 6.130341 | 6.383808 | 5.614959 | 5.511758 | 5.550229 | 5.91548 | 5.213602 | 5.674401 | 3.590684 | 3.722921 |
| Rab8b | 3.9994 | 4.570988 | 5.541374 | 5.838728 | 6.822772 | 6.484976 | 8.171765 | 7.233125 | 8.958463 | 8.993791 | 9.27931 | 10.01019 |
| Ubxn8 | 6.213432 | 5.546896 | 11.75654 | 10.94247 | 10.26706 | 10.14591 | 10.19439 | 10.54801 | 8.772449 | 7.99022 | 14.67619 | 13.69145 |
| Fam65a | 4.299786 | 5.471337 | 6.701921 | 7.076495 | 7.992901 | 7.65017 | 9.669818 | 7.676211 | 10.73976 | 10.95116 | 12.6859 | 12.12384 |
| Gm14458 | 0.470596 | 0.396903 | 2.128935 | 1.096933 | 1.458493 | 1.56101 | 2.366649 | 1.806792 | 1.570074 | 1.56086 | 1.94522 | 1.879122 |
| Ccdc120 | 5.22921 | 6.517331 | 11.42631 | 11.75849 | 9.957039 | 9.269501 | 9.347714 | 9.621975 | 7.196735 | 6.86473 | 8.649486 | 8.530079 |
| Abca5 | 0.693173 | 0.862441 | 1.848139 | 1.793958 | 2.336132 | 2.256931 | 2.779684 | 2.357305 | 2.53161 | 2.823888 | 2.532832 | 3.070162 |
| Lancl2 | 8.020169 | 8.204485 | 14.3942 | 14.67597 | 13.86024 | 13.72435 | 14.62613 | 13.09439 | 12.60149 | 11.62712 | 20.89624 | 19.88054 |
| Tnfaip8l1 | 0.532872 | 0.585947 | 1.073336 | 0.832962 | 1.125396 | 0.783974 | 1.189244 | 1.339623 | 1.190247 | 0.952393 | 1.050449 | 1.170873 |
| C130074G1 | 0.645414 | 0.623109 | 1.068172 | 1.15508 | 1.539238 | 1.489526 | 1.373154 | 1.775853 | 1.040164 | 1.039787 | 1.681883 | 1.615516 |
| Cers4 | 3.216855 | 3.956921 | 1.667582 | 1.684704 | 2.040952 | 1.903277 | 2.286472 | 2.394055 | 2.174165 | 2.237723 | 1.444426 | 1.402249 |
| Slc13a4 | 0.506284 | 0.437523 | 0.666732 | 0.918131 | 1.051561 | 0.948067 | 1.586419 | 1.340232 | 1.836763 | 1.566951 | 1.310664 | 0.921883 |
| Fam184a | 0.822034 | 0.901351 | 1.73624 | 2.018263 | 2.347076 | 2.412833 | 2.306021 | 2.82087 | 2.365423 | 2.640967 | 1.548485 | 1.652999 |
| Foxj2 | 1.175749 | 1.368018 | 1.348334 | 1.477677 | 1.803024 | 1.571403 | 1.912449 | 1.596834 | 1.900369 | 2.069283 | 2.502264 | 2.520637 |
| Znrd1as | 13.90654 | 12.67795 | 11.376 | 10.92229 | 10.54632 | 10.56149 | 8.915938 | 8.224715 | 7.359142 | 7.835838 | 6.120295 | 6.408513 |
| Arhgef5 | 0.807635 | 0.949768 | 1.54879 | 1.839915 | 1.318827 | 1.500583 | 1.240434 | 1.079768 | 1.204033 | 1.384943 | 1.716561 | 1.553137 |
| Syce2 | 40.27401 | 34.61289 | 22.84904 | 21.13248 | 18.92368 | 20.83897 | 14.81797 | 15.01081 | 14.06176 | 14.14795 | 12.48738 | 12.40433 |
| Sema3e | 1.962344 | 2.296814 | 4.058486 | 3.96218 | 3.668385 | 3.422244 | 4.018246 | 4.00638 | 3.43786 | 3.300069 | 5.406518 | 5.618718 |
| Ntpcr | 12.70301 | 11.55764 | 9.661928 | 8.668807 | 8.225001 | 7.429842 | 6.422633 | 7.826835 | 6.74032 | 5.591955 | 5.214428 | 5.092316 |
| Man1c1 | 1.296397 | 1.826311 | 2.684301 | 2.279571 | 3.087791 | 3.010935 | 3.21381 | 3.031699 | 3.64701 | 3.87958 | 2.630864 | 2.631312 |
| Evl | 11.46375 | 11.8991 | 15.95105 | 14.77965 | 19.03248 | 19.22488 | 20.13079 | 20.99457 | 23.90525 | 19.76888 | 23.69077 | 25.13177 |
| Tln1 | 13.0724 | 18.55076 | 60.93217 | 57.25865 | 48.23605 | 46.32018 | 49.6656 | 39.37327 | 33.25737 | 33.67664 | 55.44124 | 53.73524 |
| 1600014K2 | 0.482098 | 0.356163 | 0.765782 | 0.62753 | 0.988215 | 0.959411 | 0.717419 | 0.898052 | 1.093968 | 0.84043 | 1.19806 | 1.066854 |
| Trap1 | 130.01 | 117.3447 | 107.8615 | 104.5966 | 94.97111 | 91.63301 | 80.3105 | 93.69378 | 69.09398 | 70.16023 | 62.60498 | 61.52507 |
| Cdc14a | 2.394905 | 2.095618 | 3.358978 | 3.259624 | 4.174929 | 4.329067 | 4.7549 | 5.181286 | 5.001722 | 5.414485 | 5.096829 | 5.541107 |
| Neu3 | 1.725282 | 1.976634 | 3.642761 | 3.566613 | 3.876544 | 4.314777 | 4.940007 | 4.26447 | 3.22282 | 2.77968 | 2.50236 | 2.539506 |
| Atrn | 2.472644 | 2.971534 | 4.159533 | 4.301491 | 4.166102 | 4.074442 | 4.496837 | 4.351753 | 5.17976 | 5.67215 | 6.644327 | 6.922444 |
| Fam63a | 21.88698 | 22.49614 | 17.17462 | 15.04293 | 12.6198 | 12.02541 | 11.35835 | 12.72069 | 9.58738 | 8.860905 | 9.942466 | 9.867528 |
| Ralgds | 5.941903 | 7.09661 | 7.563233 | 7.030905 | 9.107804 | 8.237823 | 9.849522 | 9.907037 | 10.44006 | 10.5664 | 12.4674 | 11.93317 |
| Ssbp3 | 35.32038 | 36.9519 | 53.86225 | 53.35009 | 71.44473 | 71.07422 | 97.31141 | 83.15791 | 96.06873 | 90.53988 | 87.38274 | 84.42336 |
| Pfkl | 87.58529 | 83.02134 | 43.7041 | 43.50706 | 51.16184 | 45.33539 | 48.34261 | 49.85754 | 53.58335 | 59.96278 | 38.14026 | 35.37273 |
| Rasl11a | 3.239036 | 3.434343 | 6.121709 | 5.475735 | 6.367625 | 6.005752 | 8.125693 | 7.159094 | 9.900126 | 8.54155 | 9.104756 | 9.282151 |
| Gm1821 | 103.5405 | 85.84809 | 50.28949 | 81.58461 | 100.8421 | 108.9349 | 104.2097 | 64.32338 | 95.20411 | 84.28967 | 99.74376 | 73.15274 |
| Zmym6 | 9.714016 | 9.142822 | 6.842873 | 7.412828 | 7.10803 | 6.935441 | 6.715096 | 7.558849 | 7.128572 | 7.881541 | 4.360942 | 4.654462 |
| Camta2 | 2.777416 | 3.578837 | 5.226428 | 6.140831 | 5.751373 | 5.38073 | 5.33624 | 5.059055 | 4.122968 | 4.285025 | 3.919342 | 3.626831 |
| Usp47 | 18.20786 | 18.30355 | 25.53528 | 25.68796 | 31.61398 | 30.61901 | 34.98402 | 36.10627 | 39.14742 | 43.47076 | 38.8388 | 40.9974 |
| Serpini1 | 1.83731 | 1.562146 | 0.89613 | 1.002678 | 0.753361 | 0.806621 | 0.977598 | 0.95086 | 0.731666 | 0.91724 | 0.563257 | 0.592519 |
| Neurl1b | 0.149751 | 0.177307 | 1.009806 | 1.068723 | 1.299625 | 1.064833 | 0.9869 | 0.98349 | 0.608276 | 0.631631 | 0.775416 | 0.725497 |
| Phb | 158.8777 | 139.6582 | 131.8432 | 125.0524 | 117.5597 | 114.0198 | 104.1888 | 112.2049 | 82.7874 | 81.80126 | 75.15055 | 73.92184 |
| Col18a1 | 35.43275 | 42.41208 | 51.47891 | 48.14123 | 56.77236 | 55.36973 | 69.91932 | 64.08176 | 83.13328 | 79.12803 | 85.59233 | 81.43578 |
| Cdkl5 | 1.886812 | 1.933327 | 2.313462 | 2.365927 | 2.699682 | 2.85956 | 3.031085 | 2.653937 | 3.514244 | 3.829151 | 3.846443 | 4.000828 |
| Itgav | 4.989126 | 6.056945 | 7.521668 | 7.381467 | 8.420722 | 8.275834 | 10.76464 | 7.886481 | 12.24175 | 12.47458 | 11.25019 | 11.97451 |
| Atp6v0a2 | 7.444545 | 8.414022 | 9.908813 | 10.53063 | 9.733253 | 9.284567 | 10.18306 | 10.85131 | 10.27512 | 10.78573 | 15.09776 | 15.68978 |
| Lama4 | 0.18664 | 0.22995 | 0.484385 | 0.651389 | 0.932024 | 0.818695 | 1.189767 | 1.100749 | 1.000018 | 0.872444 | 0.852027 | 0.77492 |
| Tpi1 | 336.1283 | 262.9679 | 110.503 | 114.2878 | 127.0291 | 115.6212 | 110.5354 | 117.6781 | 138.2435 | 161.1494 | 104.0809 | 95.38225 |
| E130311K1 | 1.585479 | 1.124803 | 2.182118 | 2.188 | 2.64189 | 2.333639 | 2.285138 | 2.514819 | 2.441303 | 2.207086 | 3.767343 | 3.696631 |
| Cnih2 | 4.074344 | 4.158888 | 5.961818 | 6.053764 | 7.131678 | 7.598132 | 8.038008 | 6.706124 | 6.796758 | 5.507861 | 9.606326 | 9.304756 |
| Igfbp3 | 8.127379 | 6.874399 | 11.17132 | 11.51925 | 14.86577 | 15.11281 | 18.41376 | 19.45536 | 14.43192 | 15.3056 | 14.06321 | 12.18657 |
| Sema6b | 0.570536 | 0.942626 | 1.070983 | 0.977338 | 1.15125 | 1.113503 | 1.029485 | 1.023352 | 0.935954 | 0.799692 | 0.777006 | 0.659439 |
| Usp22 | 35.98782 | 40.9447 | 47.78374 | 46.54835 | 53.04072 | 50.91467 | 60.24018 | 54.04408 | 68.53655 | 70.50433 | 77.20706 | 78.52814 |
| Wfdc2 | 26.64302 | 23.98545 | 18.14348 | 15.09122 | 17.41325 | 16.77552 | 13.96971 | 12.3654 | 13.20963 | 12.88152 | 14.2646 | 9.945798 |
| Ankrd50 | 4.577782 | 5.237016 | 6.285784 | 6.349945 | 7.549381 | 7.055957 | 8.866925 | 7.712306 | 9.731951 | 10.7103 | 10.14594 | 9.978295 |
| 2610015P0 | 4.854026 | 4.747495 | 3.416278 | 3.854528 | 3.18767 | 2.90874 | 2.614823 | 3.286525 | 3.051987 | 3.012513 | 2.317235 | 2.285574 |

| | | | | | | | | | | | | |
|---|---|---|---|---|---|---|---|---|---|---|---|---|
| Arfgef1 | 8.636422 | 9.858801 | 11.75406 | 11.84928 | 13.14087 | 12.22546 | 14.87136 | 14.08431 | 18.52575 | 19.98269 | 19.36302 | 19.73184 |
| Des | 3.276942 | 3.95002 | 5.734735 | 5.574707 | 7.74875 | 6.970441 | 9.577751 | 8.791449 | 6.521857 | 6.987562 | 11.28382 | 9.399002 |
| Gtf2ird2 | 1.661087 | 1.738586 | 2.116741 | 2.127865 | 2.151937 | 2.029971 | 2.055661 | 1.911294 | 2.081462 | 2.009042 | 3.444276 | 3.220526 |
| Slc18a2 | 0.482983 | 0.610827 | 0.564385 | 0.655757 | 0.72588 | 0.634015 | 0.912995 | 0.628509 | 0.979452 | 1.058623 | 0.936458 | 0.981092 |
| Capg | 9.54234 | 10.15563 | 4.958609 | 4.788526 | 3.956471 | 4.232983 | 4.094568 | 3.715891 | 3.384668 | 2.844628 | 3.744515 | 3.688946 |
| Zbtb7c | 0.873445 | 1.207486 | 3.984367 | 3.996184 | 3.129568 | 3.278646 | 3.677963 | 3.179864 | 3.268375 | 2.967504 | 5.338261 | 4.993793 |
| Zglp1 | 3.494063 | 3.794568 | 3.536914 | 2.723271 | 3.169946 | 3.210652 | 2.679644 | 3.062602 | 1.602832 | 1.937231 | 2.088663 | 1.908132 |
| Vmac | 7.965351 | 7.00961 | 5.506038 | 5.505008 | 5.947689 | 6.104449 | 5.550708 | 6.618306 | 4.307656 | 3.934386 | 3.14346 | 4.007296 |
| Gsk3b | 7.680198 | 10.01688 | 12.20255 | 12.72623 | 13.71957 | 13.48268 | 16.60742 | 14.04376 | 20.71272 | 21.39871 | 19.49895 | 21.04745 |
| Abhd5 | 2.751877 | 2.593743 | 4.383045 | 4.055841 | 3.981477 | 4.06258 | 4.434284 | 4.216108 | 4.191049 | 4.052437 | 6.39814 | 6.812031 |
| Gchfr | 3.182621 | 2.490943 | 1.394525 | 2.345482 | 3.092523 | 2.741169 | 3.413327 | 3.487783 | 4.261371 | 3.940094 | 3.705783 | 3.910225 |
| Sohlh2 | 31.15107 | 29.1945 | 62.23621 | 61.90007 | 74.72602 | 73.65682 | 92.27384 | 93.3575 | 101.6101 | 103.9515 | 97.52698 | 91.3648 |
| 2310003H0 | 4.639355 | 5.09343 | 7.080271 | 6.014161 | 6.127762 | 6.514688 | 7.645701 | 7.578879 | 6.993755 | 6.694162 | 11.44441 | 10.09834 |
| Mbd5 | 2.057722 | 2.268231 | 3.412384 | 3.519995 | 3.761684 | 3.758755 | 5.068414 | 4.423962 | 5.642325 | 5.909928 | 3.975126 | 4.329502 |
| Fbxo10 | 2.231422 | 2.216195 | 3.625061 | 3.530848 | 3.849966 | 3.979506 | 4.724618 | 4.823215 | 6.094711 | 5.927091 | 4.559123 | 4.449667 |
| Hid1 | 0.984391 | 0.988034 | 1.147042 | 1.304809 | 1.868339 | 1.513678 | 1.975901 | 1.80258 | 1.651871 | 1.477 | 1.204686 | 1.073863 |
| 9530082P2 | 4.036181 | 4.088524 | 5.32388 | 5.053285 | 7.183867 | 6.897489 | 7.854739 | 8.478171 | 9.29656 | 8.013542 | 8.449729 | 8.346002 |
| Dennd2a | 12.34599 | 13.89608 | 24.1771 | 23.92482 | 29.83859 | 28.85908 | 36.90836 | 36.43004 | 28.52095 | 28.2539 | 22.46481 | 21.71701 |
| Scrn1 | 2.925069 | 3.365635 | 4.207466 | 4.531945 | 4.764664 | 4.305346 | 4.61941 | 4.16055 | 4.64622 | 4.791916 | 6.381135 | 6.977269 |
| Mrpl15 | 39.28469 | 33.53978 | 28.30242 | 27.64045 | 24.42314 | 24.00616 | 21.73922 | 19.9935 | 18.21239 | 17.63604 | 16.90033 | 16.48557 |
| Rpl30 | 163.9988 | 119.1724 | 75.51376 | 90.96492 | 103.2215 | 119.0988 | 89.1878 | 93.11239 | 82.72733 | 80.32241 | 77.87851 | 71.04473 |
| Fbxl15 | 1.359917 | 1.708591 | 3.032706 | 2.791452 | 2.540307 | 2.534444 | 3.439797 | 2.955269 | 2.118552 | 1.763172 | 2.758257 | 2.933736 |
| Abhd3 | 0.698982 | 0.75789 | 1.813128 | 1.692377 | 1.345291 | 1.761848 | 2.203383 | 1.791555 | 1.466368 | 1.412481 | 1.358557 | 1.06915 |
| Kif16b | 3.692212 | 4.217115 | 7.926681 | 7.329576 | 6.507468 | 5.790842 | 6.392094 | 5.932729 | 5.367049 | 5.672479 | 4.455354 | 4.374651 |
| Tnfaip1 | 13.91858 | 14.34612 | 18.03122 | 17.02761 | 18.77718 | 17.4146 | 21.36326 | 20.63495 | 24.80154 | 24.33353 | 28.94617 | 28.52878 |
| Herpud2 | 9.849947 | 9.949296 | 13.96542 | 13.05759 | 16.19128 | 16.11379 | 18.26012 | 17.18232 | 18.65738 | 19.07523 | 23.29432 | 21.87343 |
| Sema7a | 1.198984 | 1.123668 | 3.367869 | 3.089414 | 3.085361 | 2.713367 | 3.304471 | 3.654382 | 2.460161 | 2.181929 | 2.276938 | 2.303749 |
| Adrb3 | 5.936987 | 6.609754 | 5.971815 | 5.127626 | 4.199672 | 4.321449 | 3.744795 | 4.594484 | 2.935072 | 3.062422 | 4.640341 | 4.657636 |
| Slc12a2 | 9.37706 | 10.88933 | 12.12794 | 12.35488 | 12.82716 | 13.01847 | 15.63559 | 14.10326 | 16.10341 | 16.46826 | 20.35122 | 20.68453 |
| Cotl1 | 40.40455 | 38.06328 | 104.9286 | 98.74833 | 148.8207 | 143.4142 | 160.4569 | 162.5378 | 115.9143 | 107.9479 | 108.0681 | 97.1029 |
| Cxx1a | 2.124474 | 2.534497 | 2.698629 | 2.741801 | 3.608919 | 3.664517 | 4.585691 | 4.588661 | 3.68272 | 4.14043 | 4.538495 | 3.739792 |
| Dvl3 | 7.309392 | 9.123324 | 14.63476 | 14.73413 | 15.12962 | 14.70934 | 17.27551 | 15.4042 | 20.68289 | 20.22584 | 24.88331 | 24.61004 |
| Glrx | 7.43298 | 6.183905 | 4.155102 | 2.569771 | 3.519961 | 3.363139 | 4.127668 | 3.902687 | 3.904745 | 4.299214 | 3.558276 | 3.541022 |
| Clip2 | 1.942757 | 2.406784 | 3.881381 | 4.185485 | 5.71734 | 5.601077 | 7.246324 | 6.999569 | 6.686291 | 6.895898 | 6.400713 | 6.510629 |
| Ecm1 | 1.146701 | 1.308869 | 3.151167 | 2.432933 | 1.89679 | 2.162764 | 1.810931 | 1.455346 | 1.566339 | 1.472057 | 2.692083 | 2.049786 |
| Nkx3-1 | 0.719073 | 0.708481 | 1.190803 | 1.280358 | 1.449737 | 1.274249 | 1.366117 | 1.613577 | 1.036221 | 1.309103 | 1.0273 | 0.965238 |
| Tspan8 | 0.781411 | 0.669677 | 0.393512 | 0.627239 | 0.886655 | 0.831418 | 0.899406 | 1.046965 | 0.71045 | 0.746299 | 1.045441 | 1.048451 |
| Asz1 | 0.343802 | 0.225567 | 0.97619 | 0.816247 | 0.884968 | 0.916428 | 1.058377 | 1.199158 | 1.10061 | 0.906198 | 1.521503 | 1.364304 |
| Slc39a3 | 1.316278 | 1.366168 | 1.736701 | 1.551768 | 1.931747 | 2.045171 | 2.371576 | 2.592753 | 2.513354 | 2.179656 | 2.786913 | 2.472977 |
| Adcy1 | 1.549498 | 1.789468 | 3.177942 | 3.258666 | 3.77456 | 3.833519 | 3.886562 | 3.543992 | 2.778515 | 3.084824 | 2.325369 | 2.294563 |
| Gm15401 | 0.690869 | 1.183545 | 1.40171 | 1.465074 | 0.892069 | 1.251597 | 1.208824 | 1.100763 | 0.913722 | 0.999193 | 0.958242 | 0.877289 |
| Dusp18 | 0.77946 | 1.012952 | 1.428313 | 1.271581 | 1.365052 | 1.493565 | 2.184399 | 1.902205 | 1.866201 | 2.122192 | 1.74959 | 1.821745 |
| Eif4a2 | 445.9706 | 393.9022 | 265.8609 | 266.2235 | 292.3873 | 277.305 | 219.1129 | 260.8842 | 213.1706 | 227.4791 | 151.3873 | 166.1947 |
| Flywch1 | 9.113129 | 9.511396 | 14.54432 | 13.39227 | 16.8586 | 16.08128 | 19.70361 | 20.3018 | 20.50061 | 20.14036 | 21.76751 | 22.34019 |
| Sh3d21 | 0.667376 | 0.557876 | 1.657586 | 1.54783 | 1.039562 | 1.069479 | 1.282105 | 1.022269 | 1.144454 | 1.075149 | 1.379362 | 1.70514 |
| 2810008D0 | 9.558846 | 8.182986 | 7.419902 | 6.698402 | 6.370721 | 6.170009 | 5.026642 | 4.266581 | 4.577703 | 4.01902 | 5.823882 | 5.728722 |
| Fam122a | 7.919743 | 8.583173 | 10.84168 | 10.7891 | 13.52584 | 12.70067 | 13.34626 | 17.97942 | 16.71768 | 14.75466 | 20.6717 | 17.81457 |
| Rel | 0.655732 | 0.635856 | 1.318155 | 1.091994 | 1.159766 | 0.998255 | 1.205838 | 1.18884 | 1.03945 | 0.988665 | 1.573058 | 1.646827 |
| Kcng1 | 0.567366 | 0.671129 | 1.514778 | 1.723178 | 2.343721 | 1.903501 | 2.169539 | 1.959959 | 2.838904 | 2.869829 | 2.734826 | 2.337461 |
| Sesn3 | 5.923116 | 7.146038 | 10.21946 | 10.64255 | 11.80334 | 11.05269 | 15.99331 | 14.459 | 16.23854 | 17.7235 | 11.97445 | 11.42131 |
| Zfp647 | 3.873925 | 3.942836 | 5.726236 | 5.256138 | 6.03705 | 6.086424 | 6.616319 | 6.995088 | 8.78383 | 8.000571 | 8.528264 | 8.749638 |
| Obsl1 | 10.23794 | 11.00428 | 25.2184 | 23.29768 | 24.71266 | 23.6795 | 23.74316 | 24.13915 | 16.97374 | 16.29814 | 16.61409 | 14.50449 |
| Csnk1g3 | 10.9053 | 10.96762 | 13.95427 | 14.05638 | 16.4147 | 16.05195 | 17.3185 | 18.45245 | 22.66484 | 23.38989 | 17.80046 | 19.94751 |
| Cldn4 | 17.70168 | 19.18045 | 11.51606 | 9.93162 | 9.779175 | 8.057304 | 7.49729 | 7.850043 | 8.487301 | 8.241508 | 15.83311 | 13.05731 |
| Snx9 | 9.101422 | 8.552688 | 15.39213 | 16.29532 | 15.63476 | 15.4667 | 16.60234 | 17.04788 | 15.99524 | 17.12241 | 27.05275 | 25.7213 |
| Map4k5 | 8.300794 | 7.851947 | 11.85936 | 11.64851 | 12.33169 | 12.69613 | 13.28853 | 14.28519 | 18.41151 | 18.86723 | 18.76027 | 19.35261 |
| Tfe3 | 15.35204 | 16.61664 | 12.74649 | 12.10582 | 10.28758 | 10.29732 | 10.07878 | 9.820759 | 7.69724 | 7.287459 | 9.239111 | 9.407016 |
| Dzip1l | 3.689103 | 3.976267 | 5.865608 | 5.977617 | 7.781746 | 7.552864 | 8.140859 | 8.457624 | 8.169853 | 8.907869 | 9.945724 | 9.925666 |
| Gan | 3.505756 | 4.298754 | 6.518393 | 6.044233 | 6.41589 | 6.056657 | 6.586909 | 5.385878 | 6.193619 | 6.814619 | 9.238418 | 8.941355 |
| Tex13 | 1.915559 | 1.95437 | 6.811327 | 6.305961 | 6.91236 | 7.321441 | 9.810084 | 8.460883 | 8.670882 | 8.571682 | 10.19822 | 10.46766 |
| Gm15455 | 0.196744 | 0.240526 | 0.808758 | 0.958327 | 1.134049 | 0.796696 | 1.264191 | 1.263708 | 1.291729 | 1.322349 | 1.842986 | 1.234515 |

| | | | | | | | | | | | | |
|---|---|---|---|---|---|---|---|---|---|---|---|---|
| Gpr98 | 1.079029 | 1.176758 | 1.605245 | 1.824327 | 1.552937 | 1.684619 | 1.922141 | 1.77648 | 2.582727 | 2.795905 | 2.481948 | 2.721753 |
| Chst3 | 0.296803 | 0.257165 | 0.888798 | 0.749193 | 1.081854 | 0.985277 | 1.243909 | 1.243037 | 1.186061 | 1.241338 | 1.120427 | 1.13516 |
| Gyk | 6.895049 | 6.436171 | 5.179777 | 4.735947 | 4.814985 | 4.648391 | 3.946689 | 3.617621 | 3.381415 | 3.302254 | 2.87763 | 2.854539 |
| Fam217b | 0.675087 | 0.84012 | 1.579455 | 1.805889 | 2.218053 | 2.083241 | 2.331357 | 2.497652 | 2.117667 | 2.185672 | 1.339022 | 1.505748 |
| Efr3b | 1.265909 | 1.56609 | 2.074788 | 2.214048 | 2.7369 | 2.721855 | 3.340986 | 3.307255 | 3.32677 | 3.268389 | 3.493232 | 3.670149 |
| Satb1 | 5.309984 | 6.511863 | 7.282315 | 7.973568 | 7.907299 | 7.578586 | 8.445432 | 7.867485 | 11.99849 | 12.56014 | 7.414543 | 8.06373 |
| Ust | 1.233634 | 1.300912 | 1.425435 | 1.758488 | 2.327393 | 2.240175 | 2.947204 | 2.297406 | 2.743399 | 2.91363 | 2.411431 | 2.718392 |
| Tmem129 | 4.242114 | 4.191046 | 5.3674 | 5.33753 | 5.817564 | 6.07235 | 5.990502 | 6.513133 | 8.049742 | 7.621239 | 8.785878 | 8.843631 |
| Tsks | 1.181411 | 0.967189 | 0.504235 | 0.598877 | 0.404008 | 0.477346 | 0.549022 | 0.534025 | 0.33227 | 0.627102 | 0.259428 | 0.378525 |
| Atp11a | 5.08037 | 5.247297 | 7.09734 | 7.402664 | 7.631203 | 7.509517 | 8.696736 | 8.766238 | 11.31413 | 12.2631 | 9.75462 | 10.3381 |
| Tgfa | 0.442942 | 0.475384 | 0.960628 | 1.132821 | 1.672254 | 1.184328 | 1.573325 | 1.451783 | 1.828232 | 1.875369 | 1.681865 | 1.816259 |
| Zfp442 | 1.711419 | 1.854539 | 2.007424 | 2.295559 | 2.437775 | 2.32223 | 2.661727 | 2.919168 | 3.451555 | 3.798205 | 2.613163 | 2.889002 |
| Rad51b | 3.425205 | 3.014716 | 2.753052 | 2.698838 | 2.106495 | 2.747506 | 2.263732 | 2.717705 | 2.429567 | 2.484911 | 1.565745 | 1.660669 |
| Gng11 | 0.596765 | 0.311264 | 0.503791 | 0.934865 | 1.373396 | 1.167281 | 1.564335 | 1.544484 | 1.003691 | 1.229998 | 1.504585 | 1.240527 |
| Tmem219 | 2.940445 | 2.147783 | 2.333525 | 1.930354 | 2.000172 | 2.143035 | 1.369733 | 1.718739 | 1.60439 | 1.710991 | 1.662512 | 1.297268 |
| 4933404O1 | 9.970709 | 11.2327 | 8.233985 | 8.306178 | 7.387482 | 7.39184 | 7.095157 | 6.237919 | 6.906861 | 7.333841 | 4.814671 | 5.028691 |
| Srrd | 4.717075 | 4.025058 | 6.713034 | 5.819237 | 5.226328 | 6.171257 | 5.55513 | 5.531161 | 6.664956 | 6.355691 | 10.43547 | 9.133943 |
| Ndnl2 | 3.921533 | 4.238492 | 5.116006 | 4.722094 | 5.549425 | 5.227586 | 5.49333 | 5.154907 | 7.435567 | 6.547304 | 8.747418 | 7.794942 |
| Atpif1 | 66.79291 | 41.07702 | 19.18696 | 24.20039 | 30.19147 | 33.92914 | 21.97566 | 23.94196 | 24.97237 | 23.81188 | 22.80841 | 20.55554 |
| Emc9 | 0.993844 | 0.918808 | 1.755069 | 2.32195 | 2.190772 | 1.934188 | 1.663783 | 2.229112 | 1.763992 | 1.622836 | 1.709921 | 1.408276 |
| Ctdspl | 1.826031 | 2.203504 | 4.803506 | 4.729572 | 6.228108 | 6.072587 | 7.307246 | 6.607759 | 7.49233 | 7.699091 | 7.540939 | 7.797946 |
| Csgalnact2 | 2.38946 | 2.00028 | 3.320933 | 3.081258 | 3.692476 | 4.008416 | 4.080332 | 4.167352 | 3.887087 | 4.02223 | 4.819407 | 5.080164 |
| Xlr3b | 1.481813 | 1.482666 | 2.884763 | 2.580049 | 3.052453 | 3.305756 | 3.598211 | 4.252754 | 3.415833 | 3.448581 | 2.464201 | 2.917677 |
| Armcx2 | 15.25523 | 14.42251 | 22.26002 | 22.53349 | 32.94803 | 31.74333 | 37.24753 | 37.1409 | 27.39815 | 28.77049 | 22.16164 | 23.24885 |
| Klhl9 | 5.990843 | 6.836626 | 10.53002 | 10.23936 | 11.26695 | 10.39054 | 12.17612 | 10.82412 | 13.25808 | 14.77482 | 15.70663 | 16.87457 |
| Sec63 | 10.49545 | 11.3409 | 12.97555 | 12.96504 | 15.72402 | 15.25906 | 17.56978 | 17.34984 | 21.46185 | 21.35233 | 17.46744 | 18.96944 |
| 5330426P1 | 0.515309 | 0.611691 | 1.264437 | 1.323923 | 1.502249 | 0.984941 | 1.028717 | 1.343221 | 0.829102 | 0.815884 | 1.07628 | 1.009415 |
| Slc6a15 | 3.734569 | 3.657222 | 1.177762 | 0.853232 | 0.817332 | 0.759787 | 0.696029 | 0.656114 | 1.066567 | 1.080085 | 0.834545 | 0.706072 |
| Tmem158 | 2.001736 | 2.156936 | 2.867354 | 2.827478 | 3.742284 | 3.546694 | 4.366208 | 4.654238 | 4.443526 | 3.606306 | 4.209859 | 3.790072 |
| Lrrn4 | 2.986148 | 3.201092 | 4.234763 | 4.168555 | 5.421402 | 4.919413 | 6.285318 | 5.967196 | 6.485443 | 6.855 | 6.712968 | 5.689833 |
| Hnf1b | 0.489608 | 0.520595 | 2.108259 | 2.189957 | 2.912282 | 2.544215 | 4.319978 | 3.601241 | 3.556567 | 2.97874 | 2.641686 | 2.609444 |
| Nme7 | 32.46897 | 28.21834 | 22.1449 | 22.74953 | 21.42699 | 21.12194 | 20.05814 | 23.82691 | 17.48937 | 17.82624 | 14.29651 | 14.49333 |
| 2610307P1 | 0.604775 | 0.837513 | 0.961999 | 1.053692 | 1.055413 | 1.287714 | 1.520382 | 1.578945 | 1.896794 | 1.732042 | 0.899748 | 1.710942 |
| Arid1b | 3.560953 | 4.790559 | 5.812754 | 6.341842 | 6.996593 | 6.60396 | 8.046525 | 7.035236 | 8.993871 | 9.14387 | 10.98617 | 10.42169 |
| Gls2 | 12.81029 | 12.2015 | 8.619345 | 8.928931 | 9.067083 | 8.326377 | 7.391 | 8.105657 | 6.690895 | 6.533102 | 4.967429 | 5.437074 |
| Lcor | 2.789942 | 3.954504 | 3.911918 | 4.67957 | 3.821764 | 4.036991 | 4.096268 | 3.829265 | 5.042712 | 5.335528 | 5.930317 | 6.287835 |
| Camk2a | 0.409513 | 0.4837 | 1.103031 | 1.313648 | 1.256454 | 1.197033 | 1.49393 | 1.49937 | 1.345651 | 1.155469 | 1.814827 | 1.89683 |
| Slc25a30 | 7.901157 | 9.185359 | 9.682645 | 10.87714 | 12.3366 | 12.98205 | 14.35729 | 15.02714 | 17.01877 | 17.85181 | 15.57023 | 16.14292 |
| E130308A1 | 2.240845 | 2.696436 | 3.002387 | 3.363514 | 3.509252 | 3.562653 | 4.056912 | 3.293993 | 5.174378 | 5.403623 | 4.215195 | 4.307678 |
| Hdhd3 | 1.605511 | 1.489578 | 2.448046 | 2.323965 | 3.210202 | 3.308848 | 4.064984 | 3.811597 | 3.311805 | 3.515499 | 2.002549 | 2.486417 |
| Sgsh | 0.587723 | 0.558837 | 1.098735 | 1.174269 | 1.501762 | 1.50998 | 1.642654 | 1.925455 | 1.840411 | 1.704244 | 1.807592 | 1.4565 |
| Rhog | 5.203608 | 6.215752 | 8.243347 | 7.114827 | 9.513624 | 9.908832 | 11.39448 | 10.57259 | 12.52573 | 11.66563 | 13.22915 | 11.26613 |
| Nek6 | 7.376172 | 7.763554 | 10.97579 | 10.31097 | 10.59881 | 10.90153 | 10.61609 | 11.25406 | 9.987614 | 9.356495 | 15.47121 | 15.33519 |
| Clptm1l | 23.16085 | 23.38097 | 28.17029 | 27.28197 | 28.13915 | 27.03585 | 32.88656 | 33.50465 | 33.85045 | 32.28378 | 46.84943 | 44.29975 |
| Wdr34 | 18.98881 | 17.03897 | 14.92944 | 13.90545 | 12.0589 | 12.70943 | 11.56885 | 11.36798 | 9.974388 | 10.26972 | 8.999295 | 8.490865 |
| Fam162a | 108.5793 | 86.07308 | 57.63593 | 56.76238 | 65.44636 | 63.60626 | 66.87777 | 69.96851 | 66.15391 | 63.96475 | 49.04658 | 42.72708 |
| Slc25a18 | 0.840728 | 0.940584 | 1.280102 | 1.275795 | 1.606221 | 1.393847 | 2.018591 | 1.919939 | 1.482203 | 1.711069 | 2.03343 | 2.088914 |
| Eea1 | 3.534992 | 4.289136 | 5.099291 | 5.381996 | 5.609387 | 5.279349 | 5.698359 | 5.376916 | 6.61325 | 7.141166 | 8.50943 | 8.643997 |
| Speg | 0.66082 | 0.780024 | 1.367935 | 1.460781 | 2.019207 | 1.981797 | 2.061818 | 2.317103 | 1.333884 | 1.415377 | 1.633565 | 1.515011 |
| Slmap | 22.25771 | 25.02122 | 34.53933 | 34.25026 | 33.82157 | 33.1397 | 36.73163 | 37.58097 | 47.94987 | 47.49868 | 51.60483 | 54.25094 |
| 2010204K1 | 18.75736 | 16.36288 | 11.36283 | 11.43969 | 9.489687 | 10.38646 | 9.076819 | 10.53848 | 7.584996 | 7.631374 | 8.008876 | 6.989037 |
| Svip | 3.109227 | 3.091599 | 1.94331 | 1.905483 | 2.121258 | 2.147762 | 1.702702 | 1.884489 | 1.505754 | 1.653085 | 0.954762 | 1.312608 |
| Tnfrsf10b | 8.903459 | 11.43892 | 14.12418 | 15.1855 | 14.41014 | 14.98795 | 17.03681 | 14.49909 | 17.10354 | 17.26936 | 24.52625 | 23.72054 |
| Asb1 | 1.896508 | 2.336095 | 3.84836 | 3.680104 | 3.86759 | 3.772418 | 3.424697 | 3.056899 | 2.333823 | 2.306483 | 3.228929 | 3.117069 |
| Tmem206 | 3.528171 | 3.708936 | 4.402375 | 4.5468 | 5.417395 | 5.229335 | 5.428327 | 5.112394 | 6.827753 | 7.256518 | 7.64791 | 7.435325 |
| Lgr4 | 31.2263 | 34.58439 | 15.7069 | 15.79396 | 15.94879 | 15.66684 | 17.53221 | 15.87553 | 26.28955 | 25.60872 | 20.72801 | 21.5192 |
| Homer2 | 1.255689 | 1.123216 | 2.057022 | 2.163444 | 2.611208 | 2.626841 | 2.841643 | 2.386178 | 3.646871 | 3.454443 | 3.378851 | 3.535877 |
| Nubpl | 6.354384 | 5.701559 | 4.561848 | 4.986867 | 4.475311 | 4.733556 | 4.373758 | 4.095183 | 3.667933 | 4.108556 | 3.07901 | 3.05213 |
| Asph | 6.450984 | 6.875995 | 7.895611 | 8.328886 | 9.51252 | 9.7438 | 11.06693 | 10.95012 | 14.22395 | 13.60797 | 11.52083 | 11.78409 |
| Dpf1 | 4.578091 | 4.493049 | 3.650289 | 3.558942 | 2.924229 | 2.782592 | 2.16551 | 2.960027 | 2.285111 | 2.178929 | 2.802759 | 2.52864 |
| Zfp703 | 0.27464 | 0.274348 | 33.81526 | 32.88309 | 50.07758 | 49.37889 | 70.65654 | 51.88856 | 36.49622 | 31.76969 | 48.82633 | 43.87694 |

| | | | | | | | | | | | | |
|---|---|---|---|---|---|---|---|---|---|---|---|---|
| Tenm3 | 4.59591 | 5.705897 | 7.786363 | 8.017333 | 8.225951 | 8.064454 | 9.738151 | 8.03262 | 13.02096 | 13.09399 | 10.9966 | 11.51959 |
| Dennd2c | 11.98808 | 13.2178 | 9.903757 | 9.86475 | 7.420722 | 7.167015 | 6.935809 | 6.494937 | 5.981325 | 6.26154 | 5.761221 | 6.040606 |
| Slc27a3 | 6.897378 | 6.63147 | 12.69346 | 10.68503 | 12.14507 | 12.81301 | 15.25495 | 15.6789 | 15.23173 | 13.54258 | 18.37966 | 17.44942 |
| Gm15800 | 3.518671 | 4.730376 | 7.186757 | 7.247127 | 5.819383 | 6.215438 | 5.886832 | 5.400522 | 4.262272 | 4.67743 | 4.546514 | 4.438433 |
| Nfxl1 | 11.25263 | 11.60478 | 13.78484 | 14.02797 | 17.05811 | 16.91941 | 17.68071 | 18.81249 | 23.10315 | 22.90405 | 16.31567 | 17.49008 |
| Arhgef40 | 1.421435 | 1.699754 | 3.300103 | 3.376331 | 4.620475 | 4.289887 | 5.239459 | 5.511323 | 4.666686 | 4.825305 | 5.578586 | 5.432217 |
| Ldha | 728.6949 | 577.8184 | 199.9506 | 214.233 | 225.8081 | 206.0582 | 178.2348 | 205.6738 | 280.8384 | 335.6616 | 231.6148 | 215.9116 |
| Psen1 | 13.26389 | 15.1068 | 17.76292 | 17.9626 | 19.66127 | 19.08913 | 23.1508 | 19.922 | 26.77614 | 25.29425 | 31.04575 | 29.31394 |
| Rom1 | 0.804685 | 1.221801 | 1.508484 | 1.669846 | 1.981189 | 2.224993 | 2.787181 | 2.364853 | 2.682671 | 2.646042 | 2.820373 | 2.724886 |
| Ly75 | 8.554126 | 10.1395 | 5.722535 | 6.097429 | 5.246313 | 5.193927 | 5.259383 | 5.195381 | 5.840855 | 5.881702 | 3.80721 | 3.737554 |
| Rab32 | 0.602301 | 0.485265 | 1.08089 | 1.153128 | 1.166164 | 1.404896 | 1.433779 | 1.000483 | 0.942961 | 0.86137 | 1.440219 | 1.117777 |
| Aldh6a1 | 9.470831 | 9.412194 | 7.655935 | 7.587555 | 6.892174 | 6.622292 | 6.746229 | 6.889743 | 5.654777 | 6.343131 | 4.230773 | 4.6527 |
| Cdc42bpa | 3.939146 | 4.815225 | 6.522281 | 7.037708 | 7.912647 | 7.411806 | 8.820943 | 8.211077 | 9.222468 | 9.637331 | 10.76907 | 11.46803 |
| Mpzl1 | 26.64923 | 26.02552 | 36.45332 | 34.08901 | 41.73509 | 42.36545 | 48.99917 | 48.81657 | 54.91053 | 55.54198 | 50.64362 | 49.70394 |
| Ndufb4 | 49.85001 | 36.38375 | 20.00377 | 24.59581 | 26.22948 | 28.97773 | 22.69339 | 19.2102 | 20.71919 | 20.17697 | 19.54104 | 17.79338 |
| H1f0 | 15.46996 | 13.95157 | 32.73105 | 31.25828 | 45.67436 | 43.50222 | 51.49194 | 50.90418 | 41.39145 | 39.9446 | 52.3099 | 47.82447 |
| Fndc9 | 0.661023 | 0.944449 | 1.861513 | 1.422444 | 1.239945 | 1.458419 | 1.218403 | 1.023299 | 0.903313 | 0.895878 | 1.047824 | 0.911666 |
| Cttnbp2nl | 7.248157 | 8.453215 | 9.341527 | 9.723045 | 10.97735 | 10.15679 | 11.3991 | 10.3191 | 10.76636 | 10.77886 | 15.93704 | 15.82817 |
| Smpd1 | 3.15019 | 3.442691 | 5.533265 | 4.815588 | 5.773464 | 5.277888 | 6.486312 | 5.928852 | 7.548299 | 7.837715 | 8.692861 | 7.532092 |
| Tmem256 | 37.08062 | 27.12992 | 16.40282 | 17.31634 | 22.85553 | 25.40986 | 17.77438 | 25.52363 | 17.78114 | 17.09028 | 16.31979 | 15.68921 |
| Ankrd27 | 23.98396 | 26.79161 | 19.90781 | 18.75161 | 19.0038 | 17.8788 | 15.76626 | 17.44935 | 14.88856 | 15.8839 | 11.89287 | 11.53482 |
| Ncf2 | 0.584283 | 0.639725 | 0.643407 | 0.778575 | 0.678171 | 0.862645 | 0.858345 | 0.794357 | 1.01179 | 1.26565 | 1.225889 | 1.100406 |
| Lat2 | 0.576516 | 0.322097 | 1.093331 | 1.083781 | 1.190853 | 1.042253 | 1.125052 | 0.871946 | 1.22945 | 1.22491 | 1.059818 | 1.417184 |
| Zfp516 | 11.12048 | 12.83707 | 14.21591 | 14.25301 | 15.7184 | 15.37483 | 18.19103 | 16.35646 | 22.74021 | 23.16218 | 18.54874 | 18.47526 |
| Coq3 | 10.65796 | 8.123264 | 7.403328 | 7.030654 | 7.347917 | 7.476578 | 6.637364 | 6.912498 | 4.378759 | 4.600927 | 5.600383 | 4.973249 |
| Uvssa | 0.936021 | 1.151026 | 1.562646 | 1.495354 | 1.681945 | 1.57769 | 1.704162 | 1.695301 | 2.455596 | 2.428136 | 2.153557 | 2.287672 |
| Tmed3 | 5.159215 | 4.5217 | 7.397275 | 6.856814 | 9.213943 | 8.821217 | 9.921175 | 11.01184 | 10.61859 | 11.13052 | 11.92792 | 10.93996 |
| Ccdc71l | 2.34777 | 2.313311 | 4.126134 | 3.964518 | 5.386146 | 5.40371 | 5.762317 | 5.316762 | 5.462722 | 5.062677 | 6.451807 | 6.435344 |
| Suv39h1 | 10.64549 | 10.12896 | 14.77914 | 14.5505 | 16.11012 | 15.88265 | 17.67961 | 17.20666 | 22.17212 | 21.86138 | 24.18191 | 23.61342 |
| Pqlc1 | 3.059717 | 3.189297 | 2.544748 | 2.147668 | 1.693424 | 1.538647 | 1.413108 | 1.323514 | 1.29814 | 1.50145 | 1.626808 | 1.581852 |
| Amigo1 | 1.044118 | 1.187859 | 1.617425 | 1.44269 | 1.571541 | 1.724141 | 1.680565 | 1.820994 | 2.351449 | 2.338782 | 1.925415 | 1.605802 |
| Yod1 | 0.881969 | 0.906453 | 1.438578 | 1.310542 | 1.340297 | 1.464454 | 1.502019 | 1.208388 | 2.012576 | 1.95827 | 1.888532 | 1.918892 |
| Mettl18 | 10.23585 | 7.974939 | 6.505568 | 6.270677 | 6.789823 | 5.772339 | 4.793546 | 7.124541 | 5.678005 | 5.712374 | 4.924898 | 4.400216 |
| Ring1 | 4.476847 | 3.702184 | 6.44942 | 5.822736 | 6.016194 | 6.316578 | 7.001227 | 6.648075 | 8.281036 | 7.082394 | 10.72735 | 9.423304 |
| 1810022K0 | 96.72929 | 68.87651 | 41.18164 | 45.21814 | 47.01539 | 56.5108 | 40.90115 | 41.55762 | 40.1295 | 36.45942 | 33.93715 | 34.93037 |
| Raet1e | 2.258024 | 1.996843 | 4.323601 | 4.494844 | 4.644024 | 4.63131 | 4.849332 | 4.778722 | 3.351674 | 3.463688 | 2.833582 | 2.954239 |
| Elovl4 | 3.534127 | 3.406923 | 6.461781 | 5.965187 | 8.503374 | 8.041504 | 8.955109 | 9.385912 | 10.54346 | 9.646963 | 7.687302 | 8.767871 |
| Tmem37 | 2.932308 | 3.196898 | 3.91656 | 4.200526 | 5.499332 | 5.819749 | 6.922039 | 6.792332 | 6.622782 | 6.424694 | 5.298655 | 5.003376 |
| BC005764 | 1.658413 | 1.975812 | 2.593489 | 3.303557 | 3.183712 | 2.945957 | 3.560989 | 3.794658 | 5.517265 | 4.763159 | 3.604892 | 3.402436 |
| Sertm1 | 0.359342 | 0.384656 | 1.001075 | 0.993131 | 1.221031 | 1.078343 | 1.297679 | 1.155335 | 0.860443 | 1.069879 | 0.715142 | 0.714642 |
| Tgfbrap1 | 2.031866 | 2.257719 | 3.218632 | 3.161634 | 3.376551 | 3.279882 | 3.789434 | 3.619306 | 4.586137 | 4.419134 | 5.188516 | 5.108955 |
| Cerk | 1.459499 | 1.413281 | 1.913952 | 2.173742 | 2.749433 | 2.671926 | 2.552658 | 3.070531 | 3.632414 | 3.512184 | 3.359796 | 3.44847 |
| Ephb2 | 2.061149 | 2.504584 | 4.697876 | 4.5928 | 5.770497 | 5.337405 | 6.604128 | 5.670307 | 6.83297 | 7.020279 | 7.310976 | 7.420293 |
| Iqsec1 | 4.425648 | 5.779988 | 8.397013 | 7.815173 | 8.905023 | 8.424295 | 9.193138 | 8.405746 | 11.18837 | 11.67972 | 12.84832 | 12.62096 |
| Osgep | 18.28969 | 16.69148 | 13.79014 | 14.39605 | 14.28727 | 14.08572 | 13.49392 | 14.49726 | 11.84161 | 11.55498 | 8.87965 | 8.979328 |
| Zfp821 | 6.073251 | 6.120192 | 7.842322 | 7.423441 | 8.191971 | 7.966053 | 8.856556 | 9.34257 | 10.70608 | 10.24016 | 12.27579 | 11.98416 |
| Dmtn | 12.2203 | 12.59083 | 2.621147 | 2.35141 | 1.957374 | 1.751423 | 1.475505 | 1.600667 | 1.539148 | 1.456753 | 2.047789 | 1.877649 |
| Ugcg | 12.13787 | 12.88657 | 16.80498 | 17.58024 | 20.3415 | 19.46035 | 22.23932 | 21.3542 | 25.64559 | 28.36689 | 24.9939 | 27.40776 |
| Jak1 | 10.98649 | 12.03199 | 16.49754 | 16.05071 | 18.58875 | 17.83844 | 23.58578 | 23.89209 | 23.37982 | 23.72369 | 24.87828 | 25.89162 |
| Cdk5r1 | 0.523281 | 0.741532 | 0.882378 | 1.013308 | 0.808851 | 0.900004 | 0.914687 | 1.10355 | 1.156936 | 1.171643 | 1.339008 | 1.451161 |
| Tktl1 | 0.399528 | 0.385155 | 1.640169 | 1.607358 | 1.702639 | 1.719835 | 2.250552 | 2.293133 | 0.943977 | 1.422761 | 1.543258 | 1.47524 |
| Slc5a5 | 0.619436 | 0.757676 | 3.259737 | 3.327641 | 2.90183 | 2.563434 | 3.232015 | 3.105265 | 2.221194 | 2.068945 | 3.299718 | 2.880299 |
| Efnb1 | 3.978859 | 3.854735 | 12.8187 | 13.25711 | 17.56879 | 16.34709 | 19.01075 | 16.80195 | 21.46568 | 19.95214 | 22.64872 | 21.30337 |
| Atl1 | 1.286737 | 1.311649 | 1.640554 | 1.474215 | 1.751518 | 1.467368 | 1.763432 | 2.012391 | 1.860428 | 1.846473 | 2.817312 | 2.357201 |
| Emb | 49.7325 | 45.51655 | 21.76181 | 22.0402 | 20.96878 | 19.90088 | 23.66294 | 25.10811 | 32.82473 | 31.94746 | 26.71197 | 30.55223 |
| Cox7b | 177.9368 | 128.1801 | 96.56784 | 100.3703 | 107.2003 | 117.3941 | 94.62165 | 82.21639 | 85.38491 | 73.13949 | 79.41074 | 78.58634 |
| Zfx | 6.349778 | 7.276102 | 9.554311 | 9.101 | 8.772354 | 9.020036 | 9.556151 | 9.430981 | 12.04155 | 13.64108 | 12.86412 | 14.46798 |
| Zfp809 | 21.88335 | 26.39385 | 64.9355 | 64.51352 | 61.52406 | 61.00334 | 62.6016 | 62.07679 | 39.33775 | 42.76148 | 39.53377 | 40.2305 |
| Rc3h2 | 6.60428 | 8.447567 | 11.5709 | 11.91322 | 11.3258 | 11.29471 | 12.14573 | 9.595497 | 11.00901 | 12.2933 | 13.33425 | 15.38197 |
| Fam117b | 4.908118 | 6.222375 | 6.983418 | 6.711696 | 8.185168 | 7.603133 | 9.497682 | 8.429637 | 11.16215 | 10.65318 | 10.52255 | 10.75792 |
| Kcnip3 | 1.718913 | 1.665356 | 2.244555 | 2.410371 | 3.523793 | 3.370308 | 3.765804 | 3.84269 | 4.153518 | 3.861446 | 3.337072 | 3.581903 |

| | | | | | | | | | | | | |
|---|---|---|---|---|---|---|---|---|---|---|---|---|
| Mplkip | 13.28194 | 12.14704 | 10.52556 | 10.5322 | 10.31046 | 10.14336 | 9.092593 | 8.033356 | 7.115342 | 6.599779 | 6.050875 | 6.575842 |
| Col7a1 | 0.248703 | 0.350423 | 0.723072 | 0.628843 | 0.783424 | 0.797092 | 0.721178 | 0.790407 | 0.740707 | 0.745983 | 1.100955 | 1.006765 |
| Utp14a | 43.21075 | 43.13593 | 29.04087 | 27.5329 | 20.50454 | 20.50235 | 15.96537 | 17.19893 | 18.23644 | 17.75249 | 16.6304 | 18.17375 |
| Rapgef2 | 3.324989 | 4.174909 | 4.766469 | 5.351248 | 5.391319 | 5.735685 | 6.598281 | 5.825409 | 7.523896 | 8.177819 | 8.493981 | 8.565105 |
| Dpy30 | 214.5468 | 172.2553 | 103.9467 | 109.1972 | 90.95093 | 96.70794 | 80.09007 | 84.40769 | 77.92499 | 69.97657 | 73.46639 | 68.26498 |
| Dnmt3b | 81.98438 | 87.9515 | 36.06464 | 34.71169 | 28.83059 | 27.55011 | 29.60194 | 27.04622 | 24.96707 | 24.11824 | 22.15293 | 21.71757 |
| Zfp57 | 154.8857 | 162.3243 | 99.45829 | 91.37838 | 78.95823 | 78.92213 | 78.63738 | 87.68856 | 72.74865 | 71.37248 | 54.85353 | 57.18503 |
| Ckmt1 | 7.157273 | 7.707674 | 2.790592 | 2.231615 | 2.06355 | 2.506592 | 1.702099 | 2.285494 | 1.665533 | 1.568664 | 1.596618 | 1.917599 |
| Zfp251 | 5.025405 | 5.22406 | 6.903238 | 7.097719 | 8.133348 | 7.938428 | 8.875545 | 7.679394 | 10.20351 | 10.21674 | 10.08714 | 11.34062 |
| Plxnb1 | 4.58567 | 4.976689 | 6.952209 | 6.743565 | 6.195183 | 6.541566 | 7.135703 | 7.202353 | 9.90941 | 9.622631 | 10.45769 | 10.44834 |
| Spns2 | 0.721648 | 0.631092 | 1.173902 | 1.292962 | 1.181369 | 1.269216 | 1.120153 | 1.229158 | 1.070405 | 1.110097 | 1.653453 | 1.772943 |
| Midn | 30.8135 | 35.22456 | 49.1825 | 46.48995 | 47.15127 | 46.28466 | 51.52623 | 45.23234 | 55.77772 | 52.61784 | 76.3631 | 72.22807 |
| Asphd2 | 1.578349 | 1.628701 | 3.386878 | 4.037873 | 5.161801 | 5.417545 | 5.133431 | 6.006325 | 5.299607 | 5.680179 | 4.494687 | 3.762352 |
| B3galt1 | 0.441838 | 0.471275 | 0.771291 | 0.789724 | 0.901905 | 0.898816 | 1.047519 | 1.139425 | 1.170554 | 1.260542 | 0.754456 | 0.872763 |
| Hes6 | 20.72174 | 18.85931 | 9.296729 | 8.616934 | 8.744843 | 8.655615 | 8.77499 | 8.90862 | 9.829108 | 9.479292 | 16.19928 | 13.75011 |
| Dlgap3 | 8.749742 | 11.40102 | 7.610234 | 7.062057 | 6.183022 | 5.103955 | 5.036597 | 5.369607 | 4.151302 | 4.425767 | 5.963329 | 5.345538 |
| Jund | 12.53827 | 12.08709 | 34.51836 | 30.95857 | 34.74398 | 33.76429 | 31.89827 | 32.73205 | 25.61137 | 21.17328 | 35.14196 | 31.29123 |
| Baz1a | 8.474994 | 9.234937 | 15.0661 | 14.70033 | 17.15611 | 15.90811 | 19.44643 | 21.54008 | 21.96828 | 23.40489 | 20.65869 | 21.48001 |
| Atxn7l3b | 16.80164 | 17.39961 | 27.77555 | 28.46379 | 30.07803 | 28.34798 | 33.53746 | 27.45523 | 32.71715 | 32.47532 | 43.94438 | 43.7107 |
| Letm2 | 0.681763 | 0.686814 | 0.993584 | 0.996482 | 1.257891 | 1.009987 | 1.095535 | 1.586065 | 1.249926 | 1.331277 | 1.492668 | 1.541698 |
| Dcbld2 | 7.11281 | 8.200985 | 11.77989 | 11.44275 | 13.19447 | 12.55585 | 15.17132 | 14.05084 | 15.51619 | 16.73397 | 17.41597 | 18.20536 |
| G6pdx | 24.65947 | 25.59399 | 20.40453 | 20.67007 | 19.60199 | 18.96702 | 17.948 | 18.0258 | 16.22607 | 14.79294 | 12.31068 | 13.00714 |
| Fbxl16 | 2.02941 | 2.701546 | 3.07643 | 3.292307 | 4.45056 | 3.907601 | 4.736982 | 4.379472 | 4.904551 | 5.147693 | 5.054593 | 5.227748 |
| Csrp2 | 191.6189 | 151.1056 | 80.86507 | 73.92275 | 65.55242 | 65.24445 | 54.90989 | 53.32155 | 50.70647 | 46.55647 | 47.22092 | 46.86038 |
| Zfp46 | 1.003433 | 0.996634 | 1.487275 | 1.520686 | 1.771371 | 1.702269 | 2.126509 | 1.666969 | 2.096907 | 2.41427 | 2.273549 | 2.252397 |
| Zfand1 | 11.08521 | 8.871565 | 8.128112 | 7.150799 | 8.035391 | 7.491669 | 7.120637 | 8.122962 | 7.438135 | 7.442137 | 4.911903 | 5.117264 |
| Spats1 | 3.802922 | 4.101576 | 2.528323 | 2.565344 | 1.87365 | 2.082273 | 1.846145 | 1.616107 | 1.83552 | 1.679806 | 1.402071 | 1.71977 |
| Ncoa6 | 7.128824 | 8.1959 | 12.31138 | 12.65416 | 13.33682 | 13.42748 | 15.46344 | 13.93935 | 18.57032 | 19.12252 | 20.23034 | 20.08388 |
| Mkl2 | 3.396529 | 4.532803 | 4.864657 | 5.41425 | 4.985872 | 5.366553 | 6.472653 | 4.627384 | 6.278471 | 6.88314 | 7.222177 | 7.34076 |
| Spidr | 21.09721 | 18.72829 | 13.71946 | 13.6838 | 14.23983 | 13.6264 | 14.10126 | 15.26504 | 13.39879 | 14.90728 | 8.944239 | 9.625407 |
| Fads3 | 0.647432 | 0.671739 | 1.482679 | 1.736778 | 2.187679 | 2.135086 | 2.653565 | 1.949809 | 2.430362 | 2.321243 | 2.848337 | 2.754103 |
| Polr2k | 17.23171 | 11.4546 | 10.02941 | 10.47863 | 10.92387 | 11.25274 | 10.97323 | 7.096348 | 7.846841 | 7.543615 | 7.7532 | 8.40043 |
| Tmem205 | 13.21976 | 9.969876 | 9.279897 | 7.939392 | 9.042325 | 8.788511 | 9.027704 | 9.24293 | 7.514435 | 5.829672 | 5.866443 | 5.860195 |
| Enho | 2.350669 | 2.001067 | 2.794883 | 3.121572 | 4.219728 | 4.526927 | 4.894759 | 4.948928 | 4.209388 | 4.333125 | 4.861114 | 4.631001 |
| Phox2a | 0.713933 | 0.774514 | 0.972317 | 0.717847 | 1.202206 | 1.099826 | 0.758919 | 1.135995 | 0.698396 | 0.852569 | 1.748281 | 0.723655 |
| Josd2 | 6.559503 | 4.91133 | 7.898382 | 8.534608 | 12.14321 | 13.16684 | 13.27522 | 13.50337 | 10.49694 | 10.33668 | 10.64751 | 9.541594 |
| Acot9 | 34.64166 | 31.91061 | 15.70958 | 15.80253 | 13.70562 | 12.50482 | 12.34759 | 13.37531 | 11.12222 | 11.7987 | 17.51764 | 18.66554 |
| Zfp410 | 10.25198 | 9.907582 | 14.86469 | 13.00171 | 13.43574 | 12.90401 | 15.48184 | 14.92726 | 17.33044 | 16.25793 | 20.87753 | 20.40754 |
| Plekhg4 | 0.453951 | 0.434847 | 1.093709 | 1.16806 | 1.291033 | 1.353745 | 1.508208 | 1.599394 | 1.187556 | 1.444404 | 1.146937 | 1.011439 |
| AI504432 | 0.249401 | 0.309659 | 0.594767 | 0.657927 | 0.78062 | 0.783034 | 0.913259 | 0.732167 | 0.892799 | 0.783066 | 1.027474 | 1.039971 |
| Islr2 | 2.609622 | 2.191286 | 10.68938 | 9.822471 | 12.05309 | 12.5793 | 12.84812 | 12.64529 | 9.623082 | 9.488683 | 7.080186 | 8.000049 |
| Tnfrsf1a | 3.554889 | 3.205225 | 6.727448 | 5.710823 | 7.097732 | 5.957107 | 8.110953 | 7.956102 | 5.235614 | 5.0338 | 8.958215 | 7.926921 |
| 4930402H2 | 3.185405 | 3.50293 | 4.400508 | 4.34303 | 5.344138 | 5.087063 | 5.576576 | 5.480084 | 5.800765 | 6.177331 | 6.940803 | 6.863944 |
| Lrrc29 | 0.839514 | 0.824192 | 1.435005 | 1.310785 | 1.424373 | 1.527215 | 1.834076 | 1.731302 | 1.112493 | 1.273741 | 1.309281 | 1.095463 |
| Lrrc14 | 3.420517 | 3.447632 | 5.602653 | 5.126298 | 5.125635 | 4.875676 | 5.066095 | 4.96797 | 5.436598 | 5.592336 | 8.112712 | 7.703617 |
| Uhmk1 | 8.965719 | 10.36523 | 14.57176 | 14.80628 | 14.62073 | 13.77226 | 15.4037 | 12.8074 | 14.59819 | 16.39473 | 19.20152 | 20.23273 |
| Mum1l1 | 2.954838 | 3.009966 | 3.423729 | 3.953467 | 4.800863 | 4.420778 | 5.547911 | 5.736124 | 6.135291 | 6.245343 | 4.413706 | 5.038802 |
| Dusp9 | 27.71191 | 28.06536 | 113.3769 | 107.7176 | 97.75111 | 91.03159 | 74.59472 | 75.36269 | 67.7823 | 68.1898 | 75.17488 | 68.2188 |
| Pfdn4 | 46.85017 | 34.22868 | 31.44732 | 32.25766 | 34.53871 | 37.12331 | 29.86789 | 29.80405 | 24.84088 | 24.85651 | 23.00628 | 23.50101 |
| Nkd1 | 0.639915 | 0.661515 | 0.962453 | 1.236962 | 1.647658 | 1.662413 | 1.91219 | 1.629593 | 2.001677 | 1.931245 | 2.079309 | 1.953843 |
| Slc16a13 | 1.760739 | 1.988045 | 4.507078 | 4.445555 | 4.720192 | 4.640993 | 5.003359 | 5.295926 | 3.45613 | 3.353653 | 2.962092 | 3.356944 |
| Afap1l1 | 1.557122 | 1.044502 | 2.332578 | 2.552198 | 2.995214 | 3.286983 | 3.692941 | 4.028066 | 2.884089 | 2.947459 | 2.648868 | 2.83571 |
| Agap1 | 3.208038 | 4.34472 | 5.259664 | 5.167148 | 5.716038 | 5.701804 | 6.420569 | 5.31142 | 7.912475 | 8.152946 | 8.427974 | 7.991158 |
| Camsap1 | 7.583242 | 9.690368 | 12.76992 | 12.63 | 11.62812 | 11.23977 | 12.73897 | 11.65658 | 15.97905 | 15.92406 | 17.3102 | 17.72799 |
| Slc22a21 | 0.360579 | 0.404631 | 0.973649 | 1.047799 | 1.722145 | 1.385304 | 1.4745 | 1.652062 | 1.614093 | 1.579012 | 1.417035 | 1.481172 |
| Ttc17 | 3.4079 | 3.232863 | 5.412172 | 4.888965 | 4.866321 | 5.015714 | 5.283728 | 5.309561 | 5.012006 | 4.982385 | 7.510957 | 7.706209 |
| Rps6ka4 | 2.395683 | 2.593452 | 7.610014 | 7.11536 | 6.661797 | 6.39105 | 5.888446 | 6.777238 | 4.525469 | 4.517977 | 5.591351 | 5.019322 |
| Sfi1 | 10.14769 | 10.95487 | 17.89182 | 18.74428 | 20.71193 | 21.73946 | 23.03795 | 26.07248 | 19.55262 | 20.27768 | 15.95253 | 16.58272 |
| Adamts9 | 1.653538 | 1.68891 | 2.094214 | 2.402347 | 2.702278 | 2.622168 | 3.101684 | 2.724559 | 3.468245 | 3.650097 | 3.643052 | 3.776184 |
| Fam63b | 0.691282 | 0.724642 | 1.176628 | 1.282758 | 1.597102 | 1.602182 | 1.84414 | 1.826894 | 1.678955 | 1.946134 | 1.904872 | 2.015871 |
| Mecp2 | 2.666733 | 2.994781 | 3.85634 | 3.97682 | 4.024171 | 4.118135 | 4.738536 | 3.969897 | 5.989412 | 6.235474 | 5.898849 | 6.208472 |

| | | | | | | | | | | | | |
|---|---|---|---|---|---|---|---|---|---|---|---|---|
| Smad4 | 22.10843 | 26.58926 | 31.36877 | 29.18243 | 29.47456 | 28.2732 | 34.38032 | 32.71279 | 40.71156 | 40.0986 | 44.95489 | 44.38642 |
| Zcchc14 | 9.074397 | 11.07645 | 15.18519 | 14.80053 | 16.00778 | 15.41128 | 17.74087 | 15.88469 | 22.06276 | 23.23463 | 21.49986 | 20.26364 |
| R3hcc1 | 13.42179 | 14.19596 | 17.80257 | 18.30487 | 20.01721 | 19.24747 | 20.76082 | 21.63833 | 24.93127 | 23.55846 | 29.15907 | 28.7311 |
| Cic | 16.08379 | 19.9758 | 35.61135 | 35.60277 | 31.89773 | 31.00627 | 30.00476 | 28.42468 | 22.86397 | 22.6927 | 26.70132 | 24.38612 |
| Hibch | 7.122034 | 6.442152 | 5.426797 | 5.219063 | 5.316443 | 4.986672 | 4.444392 | 5.078824 | 4.728027 | 4.992274 | 2.956493 | 3.327833 |
| Tmod2 | 1.243277 | 1.382562 | 3.193194 | 3.278284 | 3.821928 | 3.701012 | 4.511393 | 3.577281 | 4.455649 | 4.808291 | 5.052439 | 5.137224 |
| Rhox1 | 1.163656 | 1.026551 | 3.065584 | 3.396121 | 3.693965 | 3.345059 | 3.602153 | 3.290781 | 2.261347 | 2.168589 | 3.791209 | 3.285234 |
| Chpf2 | 0.950699 | 0.938218 | 1.674416 | 1.702797 | 1.812338 | 1.913496 | 2.088443 | 1.929513 | 2.285559 | 1.998492 | 2.797354 | 2.665013 |
| Ttbk2 | 1.077074 | 1.366434 | 1.750332 | 1.912854 | 2.110827 | 2.151625 | 2.276739 | 2.185338 | 2.731417 | 2.99358 | 2.626039 | 2.792903 |
| Lysmd1 | 1.029342 | 1.304754 | 1.537592 | 1.472545 | 1.378459 | 1.578714 | 2.059753 | 1.820651 | 2.207226 | 2.304387 | 1.881872 | 1.764092 |
| Stx1b | 0.89788 | 1.097919 | 1.788417 | 1.8618 | 2.353586 | 2.268432 | 2.405391 | 2.717382 | 2.475169 | 2.662634 | 1.748468 | 1.73655 |
| N4bp2 | 12.12019 | 15.57651 | 17.19225 | 17.91878 | 20.8345 | 19.50728 | 23.71715 | 21.28299 | 26.69725 | 28.01057 | 24.84388 | 26.32861 |
| l7Rn6 | 27.75831 | 24.07874 | 21.11425 | 21.60234 | 20.02998 | 18.93331 | 18.41997 | 21.02111 | 16.10996 | 14.91178 | 13.45703 | 13.82013 |
| Mapre3 | 1.478371 | 1.542834 | 2.219592 | 2.310499 | 3.208839 | 2.732246 | 3.014794 | 3.089927 | 2.500708 | 2.276968 | 2.084113 | 1.977099 |
| Notum | 5.925215 | 5.151167 | 2.124681 | 1.712393 | 1.283851 | 1.536898 | 1.573511 | 1.377197 | 1.849346 | 1.56465 | 2.057362 | 2.150857 |
| Hrsp12 | 9.893775 | 9.115515 | 7.827628 | 7.019889 | 7.940251 | 7.567363 | 6.699893 | 7.267067 | 6.225132 | 6.007309 | 4.523466 | 4.846372 |
| Alg13 | 94.60282 | 95.97085 | 41.42142 | 41.68046 | 33.69298 | 31.70843 | 30.62423 | 32.72703 | 31.02707 | 32.05313 | 26.61684 | 26.71118 |
| Arl10 | 9.305211 | 8.95904 | 14.68161 | 13.53322 | 12.90065 | 15.11707 | 14.34736 | 15.50288 | 16.38801 | 17.29069 | 21.63496 | 21.1087 |
| Ak2 | 129.5961 | 115.2288 | 102.238 | 93.99321 | 90.06775 | 87.23834 | 78.62215 | 80.97138 | 70.14248 | 67.55479 | 65.10661 | 60.29189 |
| Epb4.1l5 | 5.621785 | 6.750907 | 8.015072 | 8.59096 | 8.563424 | 8.72096 | 10.41607 | 9.496214 | 12.19068 | 12.23585 | 13.01505 | 13.39239 |
| Sh3bp2 | 1.673714 | 1.490632 | 2.907014 | 2.555014 | 2.64271 | 2.605499 | 2.953822 | 3.290274 | 3.751753 | 3.982822 | 3.754812 | 3.806254 |
| Nampt | 59.75149 | 55.56399 | 33.60187 | 34.24598 | 36.28812 | 35.11286 | 35.28175 | 36.2197 | 40.78464 | 44.5579 | 25.95548 | 28.58446 |
| Nxf2 | 0.353402 | 0.317422 | 0.765108 | 0.71369 | 0.68956 | 0.82124 | 0.924402 | 1.018424 | 0.758845 | 0.84243 | 0.913585 | 1.112483 |
| Tfrc | 90.44507 | 85.49123 | 33.83708 | 33.30194 | 31.81299 | 29.76237 | 26.32655 | 25.52292 | 36.09329 | 39.78326 | 24.75501 | 28.16642 |
| Arhgef12 | 6.740532 | 8.255989 | 12.22542 | 13.07599 | 12.95535 | 13.05282 | 15.38316 | 14.04269 | 17.7648 | 18.72323 | 20.18266 | 20.3449 |
| Tubb2a | 13.07696 | 12.34691 | 24.51959 | 22.97146 | 26.3955 | 27.35943 | 33.38569 | 35.80125 | 26.33967 | 24.79207 | 32.26256 | 32.15443 |
| Ankib1 | 4.872291 | 5.966031 | 8.343869 | 8.150883 | 7.65203 | 7.539867 | 7.937601 | 7.834816 | 8.475979 | 9.030049 | 11.71808 | 11.74875 |
| Mib1 | 12.39241 | 15.75683 | 19.49562 | 20.04885 | 21.39574 | 20.21295 | 22.25307 | 18.9145 | 26.46449 | 29.47132 | 26.29954 | 27.85134 |
| Abl1 | 20.93362 | 23.26765 | 45.36086 | 45.1239 | 50.12691 | 50.41593 | 55.88818 | 55.0935 | 42.04741 | 43.14195 | 36.66828 | 35.36337 |
| Shmt1 | 58.24469 | 58.73569 | 44.06946 | 42.62561 | 33.93084 | 34.54739 | 28.39373 | 29.6158 | 28.90054 | 27.36818 | 27.91453 | 27.72809 |
| Mprip | 12.36912 | 13.67568 | 18.8673 | 18.17884 | 19.62183 | 18.89824 | 21.74344 | 22.90959 | 23.62484 | 23.74902 | 29.40304 | 28.30878 |
| Mroh2a | 1.058424 | 1.381202 | 2.334214 | 2.280785 | 2.619877 | 2.519139 | 2.680007 | 2.721355 | 2.528804 | 2.442891 | 1.836651 | 1.748099 |
| Lrrk1 | 0.300164 | 0.294161 | 0.97135 | 1.209262 | 1.436035 | 1.288203 | 1.88638 | 1.53817 | 1.735224 | 1.881603 | 1.690004 | 1.580526 |
| Acsl6 | 0.802372 | 0.842955 | 1.338977 | 1.393506 | 1.681882 | 1.628185 | 1.612571 | 1.517309 | 1.280094 | 1.376695 | 0.965099 | 1.047024 |
| Map3k11 | 6.831361 | 7.421752 | 10.3751 | 9.790506 | 11.33151 | 10.97468 | 13.59444 | 12.62612 | 13.61427 | 13.59307 | 16.16046 | 15.222 |
| Ddx4 | 4.731265 | 5.030046 | 7.778356 | 8.247273 | 8.222309 | 7.552006 | 7.609504 | 8.603541 | 7.988288 | 9.404082 | 12.05869 | 11.73902 |
| Smad7 | 14.97065 | 19.11238 | 7.606 | 7.748661 | 6.549602 | 6.456704 | 6.648631 | 5.981634 | 9.248567 | 8.58963 | 10.5244 | 9.29664 |
| Hhip | 0.202832 | 0.198029 | 1.063859 | 1.258484 | 1.370802 | 1.47964 | 1.943086 | 1.769534 | 1.17816 | 1.138668 | 1.654111 | 1.623266 |
| Hspa2 | 1.552529 | 1.592625 | 1.853164 | 1.935208 | 2.572319 | 2.412205 | 2.843711 | 2.547933 | 2.599707 | 2.727939 | 3.466944 | 3.005188 |
| Ppapdc2 | 0.839365 | 0.713156 | 1.113802 | 1.279206 | 1.339357 | 1.355332 | 1.271061 | 1.295033 | 1.220493 | 1.430387 | 1.966334 | 1.986145 |
| B3gnt1 | 4.15724 | 3.965201 | 5.957536 | 5.746387 | 7.233056 | 6.557746 | 7.552185 | 6.540665 | 9.536454 | 8.298175 | 8.713335 | 8.920352 |
| Kif21b | 2.312128 | 2.711779 | 3.460877 | 3.729862 | 4.840139 | 4.408083 | 4.93882 | 4.948038 | 5.555515 | 5.789455 | 3.953826 | 4.045149 |
| Adprm | 19.08251 | 17.30041 | 13.62112 | 13.57601 | 15.48687 | 14.9826 | 14.67104 | 13.86727 | 10.43732 | 10.5056 | 9.077034 | 9.656847 |
| Acaca | 10.36301 | 13.03624 | 19.34023 | 19.62252 | 21.24013 | 19.53982 | 23.36857 | 19.74596 | 23.54433 | 25.61145 | 16.98921 | 16.51724 |
| Trim16 | 0.392169 | 0.472031 | 1.1754 | 1.463658 | 1.206525 | 1.216866 | 1.48704 | 1.535839 | 1.416222 | 1.432615 | 1.920461 | 1.886138 |
| Shroom4 | 1.50298 | 1.932327 | 1.988494 | 2.216438 | 2.614314 | 2.538555 | 3.093199 | 2.571661 | 3.211756 | 3.433666 | 2.616027 | 2.874004 |
| Emp3 | 2.482266 | 2.482578 | 2.8735 | 2.643985 | 3.018647 | 2.867866 | 3.895359 | 3.573095 | 2.375512 | 2.884063 | 4.982208 | 4.094073 |
| Dcaf10 | 1.855199 | 2.129032 | 3.24243 | 3.353091 | 3.525283 | 3.477053 | 4.011041 | 3.671794 | 4.098951 | 4.260558 | 4.868095 | 5.18856 |
| Ctbp2 | 93.54541 | 100.0222 | 58.01297 | 54.73982 | 44.63499 | 43.09081 | 37.61035 | 40.58607 | 35.52558 | 35.37659 | 38.80848 | 38.50145 |
| Tm6sf2 | 0.445654 | 0.646294 | 1.642069 | 1.458972 | 1.184421 | 1.316321 | 1.14586 | 1.815046 | 1.213899 | 1.173481 | 1.295572 | 1.325786 |
| Oaz1-ps | 1.449629 | 1.209373 | 2.040672 | 2.041281 | 1.802669 | 1.643325 | 2.467975 | 2.339913 | 2.77347 | 2.54112 | 3.294993 | 2.343733 |
| BC026585 | 3.038817 | 3.213987 | 5.759351 | 5.536741 | 6.12273 | 6.080949 | 6.811181 | 7.733939 | 5.424902 | 5.405319 | 4.753421 | 5.001605 |
| Sipa1 | 5.579947 | 6.176091 | 18.60817 | 16.96853 | 17.9328 | 17.61632 | 17.48833 | 17.69027 | 11.80365 | 11.4409 | 12.83202 | 11.69896 |
| Slc25a29 | 0.531391 | 0.876454 | 0.869399 | 0.889542 | 0.69298 | 0.916539 | 1.030154 | 0.592954 | 1.323456 | 0.909849 | 1.179652 | 0.819158 |
| Fut8 | 5.960148 | 6.139069 | 7.974793 | 8.123481 | 9.724529 | 8.528637 | 10.76239 | 9.965188 | 12.69006 | 12.53458 | 12.2232 | 12.52271 |
| Abhd2 | 3.680716 | 4.922555 | 7.62046 | 7.452679 | 6.532286 | 6.624008 | 8.375794 | 6.646605 | 8.167622 | 8.429328 | 9.969985 | 10.20833 |
| Wipf1 | 1.834791 | 2.278235 | 4.727661 | 4.838899 | 5.656946 | 5.525317 | 6.617622 | 5.471792 | 7.104671 | 6.844734 | 8.22527 | 7.4571 |
| Znrd1 | 80.3683 | 62.45087 | 59.77205 | 56.54049 | 56.82719 | 59.10481 | 50.35973 | 54.32035 | 38.61579 | 38.36405 | 43.12333 | 41.79939 |
| Sun2 | 3.626216 | 3.906255 | 5.697848 | 5.875761 | 6.698924 | 6.716055 | 7.203394 | 7.367803 | 5.628298 | 5.122431 | 8.358577 | 7.890353 |
| Farp2 | 0.614691 | 0.833844 | 1.37729 | 1.610562 | 1.560261 | 1.653692 | 1.858258 | 1.345182 | 1.855541 | 2.071326 | 2.157371 | 2.045867 |
| H2-K2 | 1.941534 | 1.428473 | 1.558434 | 1.735732 | 1.247469 | 1.335333 | 1.316287 | 1.519048 | 0.737867 | 1.128129 | 1.072864 | 1.210448 |

| | | | | | | | | | | | | |
|---|---|---|---|---|---|---|---|---|---|---|---|---|
| Qpct | 0.616986 | 0.562535 | 1.060183 | 0.912565 | 1.036208 | 0.957597 | 1.55966 | 1.120523 | 1.587248 | 1.403217 | 0.949146 | 1.047306 |
| Ndufaf1 | 20.47045 | 17.42025 | 14.19609 | 14.9882 | 15.97373 | 15.49557 | 15.08961 | 17.50381 | 13.0524 | 13.14506 | 10.14775 | 11.38389 |
| Adck2 | 1.383121 | 1.505598 | 3.077402 | 2.79685 | 3.547678 | 3.291449 | 3.917418 | 4.30034 | 3.617344 | 3.298521 | 2.978459 | 2.901771 |
| Kcna3 | 0.267129 | 0.331934 | 0.866439 | 0.839764 | 0.992879 | 1.114061 | 0.977998 | 1.190729 | 1.155309 | 1.039479 | 1.264316 | 1.290406 |
| Lamc1 | 42.94507 | 54.30802 | 79.49209 | 78.56815 | 70.37217 | 68.59163 | 92.18057 | 74.01458 | 106.3506 | 102.9417 | 99.77189 | 105.3336 |
| Helz | 6.590729 | 9.727962 | 14.06875 | 14.2493 | 12.23665 | 11.91207 | 13.42069 | 10.16695 | 10.06153 | 10.31543 | 9.608155 | 9.282008 |
| Fkrp | 7.481947 | 6.588765 | 11.24344 | 10.07295 | 11.34717 | 11.70566 | 12.1959 | 11.58719 | 12.52849 | 11.94227 | 16.11548 | 15.46347 |
| Pgap3 | 1.581953 | 1.750861 | 2.603706 | 2.68705 | 2.572096 | 2.666525 | 3.257648 | 3.061182 | 2.497357 | 2.979513 | 3.893051 | 4.093483 |
| Lnx2 | 4.22786 | 4.840325 | 8.849872 | 8.933657 | 6.735104 | 6.757643 | 6.913664 | 6.52046 | 5.767966 | 6.038907 | 5.367609 | 5.820787 |
| Trpc1 | 0.535156 | 0.573959 | 1.233795 | 1.212595 | 1.607293 | 1.5342 | 1.784161 | 1.860703 | 1.655788 | 1.804844 | 1.601143 | 1.679293 |
| Hdx | 0.827369 | 1.02203 | 1.267148 | 1.421785 | 1.677347 | 1.649849 | 1.82882 | 1.843419 | 1.937993 | 2.181204 | 1.268231 | 1.544425 |
| St3gal5 | 3.818559 | 3.828423 | 5.608674 | 5.390668 | 6.748121 | 6.841476 | 7.379924 | 7.186671 | 7.793514 | 8.17292 | 8.287983 | 8.269746 |
| Nr6a1 | 11.83648 | 14.74433 | 28.62597 | 28.0398 | 29.95433 | 26.52146 | 36.31624 | 30.97342 | 37.2482 | 40.66151 | 32.7135 | 32.61507 |
| Ecel1 | 0.35885 | 0.292335 | 0.780046 | 0.668322 | 1.089311 | 0.851202 | 0.937188 | 0.834429 | 0.993885 | 1.023536 | 1.004558 | 0.859512 |
| P2ry1 | 0.549859 | 0.457756 | 0.652705 | 0.752072 | 0.92751 | 1.000259 | 1.11656 | 0.844235 | 0.94113 | 0.943209 | 1.36617 | 1.15132 |
| Rtn1 | 1.615557 | 1.901323 | 2.490875 | 2.467276 | 2.862716 | 2.885083 | 3.716437 | 3.486942 | 3.318163 | 2.970894 | 3.616056 | 3.77659 |
| Zfp319 | 0.250575 | 0.30364 | 0.662166 | 1.054075 | 1.110741 | 1.24183 | 1.14768 | 0.937342 | 1.609254 | 1.434279 | 1.197915 | 1.394098 |
| Sh3tc2 | 0.244297 | 0.359785 | 1.172385 | 1.194091 | 1.114064 | 1.111239 | 1.294637 | 1.219206 | 1.162718 | 1.162439 | 2.051287 | 1.699994 |
| Elf3 | 5.791406 | 7.033539 | 12.10289 | 10.51532 | 11.91267 | 11.24281 | 14.06158 | 16.0847 | 16.21022 | 15.43339 | 14.94415 | 14.55298 |
| Hspb8 | 15.45671 | 13.6012 | 6.73232 | 7.002243 | 7.743469 | 7.232932 | 7.417205 | 8.439173 | 8.370532 | 8.62779 | 11.48601 | 10.71038 |
| Rc3h1 | 3.09575 | 4.38167 | 7.026997 | 7.343065 | 6.203739 | 6.259998 | 7.18612 | 5.727443 | 6.718811 | 6.882769 | 8.302297 | 8.756347 |
| Tbc1d2b | 2.612907 | 2.942659 | 4.036706 | 3.978137 | 4.466097 | 4.447184 | 5.181076 | 4.744939 | 5.512372 | 5.541627 | 5.96467 | 6.080674 |
| Ano3 | 0.851307 | 0.946814 | 1.217506 | 1.166419 | 1.408742 | 1.287606 | 1.771846 | 1.64961 | 1.406604 | 1.42522 | 0.689991 | 1.079288 |
| Nkain4 | 2.670607 | 2.486904 | 1.314757 | 1.380979 | 2.032532 | 1.623555 | 2.18758 | 1.628015 | 1.559391 | 1.715191 | 2.083658 | 2.110589 |
| Cntfr | 12.56711 | 14.64965 | 66.30734 | 67.86142 | 72.03959 | 70.66315 | 70.41445 | 67.14092 | 56.77164 | 54.01127 | 49.55187 | 46.24695 |
| Cyb5d1 | 14.07208 | 13.68711 | 20.68358 | 19.80664 | 19.53307 | 18.62883 | 24.64864 | 23.75083 | 24.63328 | 23.34991 | 28.5278 | 28.45958 |
| Il17ra | 1.330365 | 1.512708 | 2.204996 | 2.182503 | 2.475472 | 2.405624 | 2.976706 | 2.986744 | 2.54596 | 2.34624 | 3.181318 | 3.334181 |
| Trim13 | 27.42372 | 24.63217 | 16.70966 | 16.08932 | 14.96194 | 13.89689 | 13.44293 | 12.48836 | 9.841805 | 10.53265 | 9.121261 | 10.59428 |
| Pbx2 | 20.44202 | 21.15624 | 62.43683 | 58.92642 | 73.16549 | 68.35126 | 75.68428 | 70.25481 | 61.53018 | 59.05183 | 46.36715 | 46.01542 |
| Aebp1 | 0.979079 | 1.078024 | 2.044636 | 1.915658 | 2.381341 | 2.403766 | 2.783262 | 2.923841 | 2.768937 | 2.770247 | 2.567221 | 2.843349 |
| Pfn4 | 0.524506 | 0.385625 | 0.948203 | 0.956685 | 1.126306 | 1.096231 | 1.224939 | 1.256009 | 1.055429 | 0.834792 | 0.938813 | 1.273707 |
| Pptc7 | 8.414995 | 8.470014 | 17.67981 | 17.58436 | 15.09098 | 15.69896 | 13.6501 | 14.6939 | 11.03439 | 11.52508 | 12.82889 | 13.63254 |
| Clasp1 | 6.919161 | 8.758886 | 9.935416 | 11.05352 | 11.71824 | 11.63373 | 14.48502 | 12.55572 | 15.21018 | 16.86101 | 13.47797 | 13.55754 |
| Ccdc172 | 1.649344 | 1.952518 | 4.592579 | 4.471691 | 5.026791 | 5.58939 | 6.183152 | 6.818601 | 4.806148 | 4.809616 | 3.621543 | 4.84541 |
| Prmt2 | 6.48706 | 6.360028 | 11.86091 | 11.75348 | 14.76142 | 13.68519 | 17.29516 | 17.92108 | 13.55318 | 13.705 | 12.6797 | 11.78152 |
| Gm364 | 3.635089 | 2.747413 | 6.434841 | 5.631678 | 6.343077 | 7.055498 | 9.658585 | 7.716984 | 7.652633 | 7.40591 | 7.801258 | 8.586322 |
| Ppp2r5b | 6.629854 | 7.167498 | 13.47031 | 12.26032 | 11.91169 | 11.56464 | 11.25687 | 11.16127 | 9.227855 | 8.041875 | 9.764888 | 10.2523 |
| Zbtb12 | 6.117917 | 6.218596 | 9.241542 | 9.167031 | 8.898911 | 9.147135 | 9.816466 | 8.298855 | 14.12919 | 12.64689 | 12.27718 | 10.74433 |
| Zkscan7 | 0.525066 | 0.700455 | 0.961444 | 1.007007 | 0.840486 | 0.901371 | 1.123036 | 1.26213 | 0.736566 | 0.829193 | 1.116257 | 1.035 |
| Strip2 | 0.251033 | 0.267815 | 1.027218 | 0.981975 | 1.049955 | 0.893129 | 0.800607 | 1.029854 | 0.675981 | 0.695844 | 0.528592 | 0.67663 |
| Stac2 | 3.569045 | 3.501474 | 2.135631 | 2.190942 | 2.193095 | 1.790776 | 1.760334 | 1.598091 | 1.695429 | 1.561119 | 1.696458 | 1.440556 |
| Clip3 | 4.498826 | 5.676429 | 7.964037 | 8.73796 | 9.764657 | 9.434027 | 11.65852 | 9.897398 | 14.43827 | 13.28766 | 10.08558 | 9.890996 |
| Pcnx | 3.803953 | 4.92923 | 5.192255 | 5.575835 | 6.468305 | 6.29791 | 7.670878 | 6.358851 | 7.62181 | 8.477145 | 7.478474 | 7.632991 |
| Vegfa | 11.61074 | 10.38898 | 18.9346 | 18.19625 | 20.03787 | 19.53423 | 18.64832 | 20.2217 | 23.83763 | 23.7877 | 29.31287 | 27.65284 |
| Nlrp4c | 0.886402 | 0.782265 | 1.922659 | 2.04867 | 2.658543 | 2.3694 | 2.906023 | 3.112369 | 2.665112 | 2.706327 | 1.929398 | 2.130062 |
| Fam131a | 0.712618 | 0.703601 | 1.141961 | 1.347806 | 1.407949 | 1.58896 | 1.724481 | 1.830746 | 1.473443 | 1.397064 | 1.794524 | 2.04713 |
| Elmo1 | 6.987153 | 6.972363 | 3.30919 | 3.284425 | 2.983613 | 3.063369 | 3.595305 | 3.513781 | 4.461822 | 4.634054 | 4.486043 | 4.650315 |
| Phactr4 | 8.485136 | 10.5486 | 10.49549 | 11.63544 | 12.86175 | 12.43814 | 14.70086 | 13.18454 | 18.58501 | 17.65528 | 16.30692 | 16.24654 |
| Mlxip | 5.297224 | 6.775504 | 6.870329 | 7.165935 | 7.814106 | 7.611396 | 9.887664 | 8.118153 | 10.75676 | 10.87222 | 8.837962 | 8.837068 |
| Ncoa2 | 6.366096 | 7.439983 | 9.037028 | 9.687731 | 10.82782 | 9.859089 | 12.51367 | 10.88078 | 13.93608 | 13.76294 | 14.63135 | 14.69802 |
| Zfp787 | 4.265093 | 4.01599 | 5.760162 | 6.072031 | 7.985809 | 7.439792 | 8.063688 | 8.267759 | 10.35702 | 9.109715 | 10.09287 | 9.211587 |
| Repin1 | 3.795599 | 3.982899 | 8.336223 | 8.180465 | 8.100583 | 8.084234 | 8.544207 | 8.752108 | 9.738003 | 9.513843 | 12.32509 | 12.25566 |
| Usp28 | 57.6357 | 64.44513 | 25.08527 | 25.91569 | 21.378 | 21.08247 | 22.00345 | 21.87871 | 23.42895 | 23.72021 | 17.05943 | 17.09598 |
| C030039L0 | 2.954864 | 3.040468 | 2.211379 | 2.30229 | 1.834678 | 1.652614 | 1.336657 | 1.89013 | 1.934112 | 2.013185 | 2.303731 | 2.504443 |
| Gmfg | 2.467268 | 1.99588 | 1.680134 | 1.696908 | 2.078156 | 1.783026 | 1.597342 | 1.696838 | 0.922994 | 1.230114 | 1.012397 | 1.177695 |
| 1500011K1 | 10.3769 | 6.512343 | 6.466228 | 7.19395 | 7.924146 | 7.171045 | 5.23721 | 6.327057 | 5.572847 | 4.75282 | 6.747936 | 6.038418 |
| Immp2l | 3.826633 | 2.836813 | 2.239214 | 1.807779 | 2.372669 | 1.74268 | 1.87501 | 2.043029 | 1.865923 | 1.710902 | 1.458798 | 1.349849 |
| Zfp287 | 1.027907 | 1.018609 | 2.251497 | 2.246689 | 2.578575 | 2.658232 | 2.284755 | 3.220298 | 2.85554 | 3.035623 | 3.360886 | 3.400398 |
| Adam9 | 7.556379 | 7.394207 | 12.20507 | 11.57736 | 13.63672 | 13.75866 | 15.77101 | 16.6534 | 18.05093 | 17.51391 | 15.80145 | 16.7668 |
| Ccl25 | 3.782783 | 3.962665 | 5.600541 | 5.575583 | 5.972359 | 5.284484 | 5.417273 | 6.001016 | 8.22486 | 7.993549 | 6.050481 | 7.208897 |
| Plekhm2 | 3.068101 | 3.147168 | 6.668829 | 5.566229 | 6.385551 | 6.244734 | 6.463136 | 6.927394 | 7.499419 | 7.753843 | 8.869847 | 8.412997 |

| | | | | | | | | | | | | |
|---|---|---|---|---|---|---|---|---|---|---|---|---|
| Arhgap42 | 3.5548 | 4.224705 | 5.245999 | 4.804833 | 5.901119 | 5.718409 | 7.551159 | 6.743466 | 6.572066 | 7.261247 | 6.180147 | 6.42299 |
| Pdzd8 | 7.892439 | 9.552567 | 12.71633 | 12.16092 | 12.32019 | 11.81024 | 13.23571 | 11.8956 | 13.76287 | 13.58077 | 17.97559 | 17.84303 |
| Fam134a | 8.590162 | 8.989361 | 12.69621 | 12.4071 | 14.25295 | 14.27915 | 16.10505 | 17.8794 | 16.86898 | 16.4523 | 19.25414 | 18.74443 |
| Dok1 | 1.043721 | 1.106408 | 1.923273 | 1.894033 | 2.40739 | 2.240963 | 2.433377 | 2.860657 | 3.15654 | 2.754645 | 2.822843 | 2.582259 |
| Gm14305 | 5.988943 | 5.518218 | 5.046307 | 4.661381 | 4.002998 | 3.964661 | 3.454953 | 4.014739 | 4.169341 | 4.676143 | 2.880774 | 3.096169 |
| Tgif1 | 42.25561 | 45.22592 | 20.2022 | 20.08508 | 20.91619 | 20.76975 | 23.98406 | 22.60648 | 26.19031 | 25.35118 | 17.83979 | 16.78935 |
| Kif13b | 7.211578 | 9.615778 | 18.48374 | 19.30449 | 17.56309 | 18.27492 | 21.05807 | 19.80747 | 18.02645 | 19.28157 | 13.15326 | 13.11914 |
| Syt14 | 2.772348 | 3.843777 | 4.171766 | 4.605041 | 4.083256 | 4.367922 | 4.845683 | 4.629851 | 4.618984 | 4.479694 | 7.166771 | 6.59151 |
| Tsc22d2 | 3.76238 | 4.658351 | 6.296048 | 6.265758 | 5.961068 | 5.891561 | 6.091281 | 5.09598 | 7.149825 | 6.853052 | 8.025765 | 7.68972 |
| Smad5 | 8.223332 | 9.144665 | 13.21691 | 13.76733 | 15.16009 | 14.91446 | 17.52341 | 15.63897 | 19.84215 | 20.75325 | 18.94787 | 19.63026 |
| Zfp42 | 421.8086 | 400.611 | 160.1308 | 147.5283 | 119.6588 | 114.0758 | 111.2242 | 119.2507 | 126.5385 | 130.4503 | 98.99495 | 97.88795 |
| Stard10 | 3.780264 | 4.473769 | 5.013764 | 4.694554 | 6.594752 | 6.503838 | 6.551609 | 7.072225 | 8.066347 | 6.961573 | 7.176446 | 6.828745 |
| Nrip2 | 0.526003 | 0.772352 | 1.088915 | 1.353687 | 1.149167 | 1.102972 | 1.468937 | 1.307852 | 1.379045 | 1.322689 | 1.736989 | 1.661499 |
| Cand2 | 6.807443 | 8.16412 | 14.79906 | 15.0129 | 16.38934 | 16.4848 | 21.11393 | 19.94531 | 23.39908 | 22.60213 | 18.63456 | 18.02187 |
| Gm5635 | 0.952178 | 1.032161 | 2.300024 | 2.682315 | 2.804333 | 2.612562 | 2.893203 | 3.430649 | 2.629553 | 2.279568 | 2.633218 | 2.479931 |
| Spred3 | 2.099094 | 2.442653 | 3.122531 | 3.282198 | 3.645333 | 3.688973 | 4.126177 | 3.847816 | 4.248937 | 4.421899 | 4.41083 | 4.937666 |
| Prps1 | 76.23024 | 75.66673 | 57.86876 | 56.46296 | 50.10886 | 48.06353 | 45.15699 | 45.7461 | 47.60297 | 48.45953 | 38.33046 | 37.53632 |
| Mfn2 | 5.767543 | 7.071821 | 11.62002 | 11.20155 | 11.17296 | 10.75902 | 11.93901 | 11.12027 | 10.67982 | 10.28648 | 15.32428 | 15.46785 |
| Syde2 | 3.548303 | 4.391995 | 5.142599 | 5.406033 | 5.46023 | 5.315306 | 5.263931 | 5.288006 | 7.577711 | 7.595267 | 7.965523 | 7.950288 |
| Six5 | 1.448829 | 1.766889 | 2.936663 | 3.266687 | 4.412893 | 4.098842 | 4.591648 | 4.784901 | 4.904484 | 4.770165 | 5.052714 | 4.910343 |
| Eps8 | 10.19441 | 9.968589 | 6.206705 | 6.249831 | 6.492514 | 5.983641 | 5.960667 | 5.931504 | 4.090928 | 4.338157 | 4.312425 | 4.163489 |
| Prdm14 | 11.00583 | 9.904561 | 3.143951 | 2.8602 | 3.024969 | 2.884015 | 3.360849 | 4.277262 | 4.522651 | 4.292292 | 4.130996 | 3.512345 |
| Smad6 | 1.581666 | 1.658756 | 4.518263 | 3.757192 | 4.569052 | 4.246861 | 4.301822 | 4.590482 | 6.130983 | 5.635119 | 4.672578 | 4.482398 |
| Rab43 | 1.856329 | 1.737077 | 3.243662 | 2.594326 | 3.904281 | 3.860269 | 4.547037 | 3.806361 | 3.013664 | 3.477176 | 4.031519 | 3.889517 |
| Car13 | 15.63674 | 13.54192 | 8.965204 | 8.51285 | 10.34048 | 10.78557 | 11.30277 | 12.47615 | 10.76537 | 10.93981 | 7.679919 | 8.322541 |
| Sec24b | 11.29317 | 13.58171 | 16.68947 | 17.8613 | 18.05112 | 17.10925 | 19.91888 | 18.06869 | 22.19808 | 23.36364 | 26.32729 | 25.64472 |
| Wiz | 8.112436 | 9.909429 | 11.87513 | 11.76454 | 11.35659 | 11.49558 | 13.11599 | 10.7504 | 15.91321 | 14.595 | 16.77815 | 16.11154 |
| Ttyh2 | 2.540172 | 2.905065 | 3.779231 | 3.830302 | 5.136179 | 4.92179 | 5.327892 | 4.934085 | 4.844574 | 4.894774 | 3.628791 | 3.293303 |
| Slfn3 | 0.333117 | 0.515123 | 0.672529 | 0.92672 | 0.798292 | 0.906833 | 1.062919 | 1.006507 | 0.731157 | 0.750967 | 1.290414 | 1.124762 |
| Nynrin | 1.700741 | 2.283172 | 3.23741 | 3.084881 | 3.672308 | 3.500351 | 4.649301 | 4.325026 | 4.52904 | 4.618831 | 4.005797 | 4.226461 |
| Serinc2 | 1.971904 | 1.923434 | 3.161448 | 3.571491 | 4.024656 | 3.264691 | 4.106718 | 4.310477 | 3.892328 | 3.92867 | 4.936982 | 4.885295 |
| Grina | 12.79712 | 13.51958 | 17.43042 | 15.92062 | 21.45651 | 20.36722 | 24.64687 | 22.31239 | 27.17401 | 23.81148 | 23.00598 | 23.71016 |
| Spats2 | 32.00466 | 32.18839 | 36.48966 | 40.05747 | 48.19004 | 46.4311 | 53.68695 | 55.53712 | 64.33373 | 59.88726 | 46.08364 | 44.47692 |
| Slc25a51 | 10.07009 | 10.4738 | 15.55784 | 15.58726 | 15.67743 | 16.36737 | 18.41281 | 16.0307 | 19.5295 | 19.80412 | 22.49314 | 23.30021 |
| Cxx1b | 2.502049 | 2.336934 | 3.487793 | 3.640635 | 4.383925 | 5.228023 | 5.725046 | 5.43993 | 4.517995 | 4.67585 | 4.921329 | 4.329929 |
| Dgat2 | 1.298265 | 1.260218 | 3.285682 | 3.758201 | 4.249697 | 4.292577 | 5.166331 | 4.372318 | 3.36212 | 3.481959 | 5.59734 | 5.202574 |
| Igsf10 | 0.622177 | 0.783904 | 1.174577 | 1.259288 | 1.389521 | 1.261672 | 1.741338 | 1.282757 | 1.604755 | 1.880783 | 1.460973 | 1.570915 |
| Ifitm2 | 434.2596 | 370.6809 | 208.4315 | 215.2151 | 201.6371 | 208.0397 | 186.9947 | 156.9711 | 167.8065 | 154.2329 | 141.4083 | 144.1197 |
| Rab3b | 0.351372 | 0.373738 | 0.769414 | 0.972008 | 1.114079 | 1.005254 | 1.334567 | 1.45095 | 1.294263 | 1.174971 | 0.950931 | 0.99105 |
| Fgfr1 | 25.75988 | 27.02949 | 40.83636 | 40.71613 | 50.3616 | 48.20841 | 58.96065 | 60.77567 | 55.42477 | 53.97389 | 47.12639 | 45.70935 |
| Dusp22 | 3.385394 | 3.713811 | 5.84907 | 5.815075 | 7.338887 | 7.990459 | 8.389397 | 8.679091 | 8.62845 | 8.466833 | 8.86967 | 8.202144 |
| Slc9a1 | 7.268161 | 8.984149 | 16.03271 | 15.67408 | 14.67893 | 13.65435 | 15.14183 | 12.25801 | 10.73371 | 10.50249 | 12.32695 | 11.82763 |
| Prdm11 | 0.556393 | 0.570385 | 1.445355 | 1.321324 | 1.187739 | 1.059045 | 1.188055 | 0.97924 | 1.125803 | 1.110614 | 1.502035 | 1.461342 |
| Rere | 12.4858 | 15.91037 | 19.21658 | 20.07379 | 19.06755 | 18.89632 | 21.60117 | 17.47813 | 26.69434 | 26.08245 | 24.85162 | 23.69697 |
| Scmh1 | 3.962681 | 4.39876 | 6.408734 | 6.905147 | 8.06844 | 7.556773 | 9.045919 | 7.51143 | 8.469891 | 8.151571 | 9.756506 | 10.29268 |
| Pcyt1a | 6.597791 | 6.705674 | 10.36917 | 10.4754 | 10.64045 | 10.31921 | 11.49707 | 10.30868 | 11.18252 | 10.98703 | 14.64802 | 15.35693 |
| Suox | 1.801059 | 1.9024 | 3.07602 | 3.083465 | 3.972343 | 3.563627 | 3.910197 | 4.588945 | 3.936323 | 3.950222 | 3.744054 | 3.610681 |
| Ttc39a | 0.63271 | 0.845301 | 1.006015 | 1.109532 | 1.546769 | 1.323591 | 1.742222 | 1.65109 | 1.405577 | 1.6381 | 1.008285 | 1.429228 |
| Camsap2 | 5.666156 | 6.741192 | 8.917253 | 9.471251 | 10.22487 | 10.16304 | 11.52428 | 10.45418 | 12.67212 | 14.09188 | 14.09999 | 13.98615 |
| Mrpl42 | 170.9475 | 142.5334 | 113.1003 | 109.8909 | 115.043 | 117.1535 | 95.51849 | 93.87492 | 97.98884 | 89.49606 | 84.01219 | 78.81756 |
| Otud6a | 1.843814 | 2.020229 | 7.983723 | 8.098412 | 9.900956 | 8.719396 | 12.21949 | 12.02546 | 8.539408 | 10.76488 | 9.805887 | 8.746637 |
| Fbln1 | 7.740773 | 8.701732 | 15.66856 | 14.40832 | 15.30898 | 14.30851 | 19.91898 | 18.45736 | 20.36251 | 19.72228 | 14.80492 | 13.28242 |
| Abat | 1.200173 | 1.221404 | 1.906615 | 1.869273 | 1.998235 | 1.813325 | 2.455071 | 2.052204 | 2.570615 | 2.692459 | 2.387001 | 2.58188 |
| Bcl9l | 1.901173 | 2.655677 | 4.935161 | 4.810689 | 5.468 | 5.190991 | 6.616451 | 4.718519 | 5.547417 | 5.511316 | 7.813692 | 7.004144 |
| Otud7b | 6.801136 | 7.51065 | 9.657614 | 10.35378 | 12.01939 | 11.67745 | 13.14501 | 11.0558 | 11.72144 | 12.00998 | 15.14023 | 15.36095 |
| Ahi1 | 4.678231 | 4.761383 | 7.009142 | 7.442107 | 9.10483 | 8.040914 | 10.12689 | 10.0112 | 8.414784 | 9.144054 | 6.300559 | 6.751342 |
| Slc18b1 | 1.184188 | 0.916028 | 1.667363 | 1.803057 | 1.827903 | 1.72372 | 2.409492 | 2.555188 | 2.939626 | 2.164203 | 1.890381 | 1.801452 |
| Ier3 | 4.733955 | 5.276859 | 2.321783 | 2.781509 | 3.256817 | 2.760971 | 2.933134 | 2.444879 | 2.447372 | 2.119043 | 2.492313 | 3.139564 |
| Sufu | 7.368135 | 8.564139 | 12.07546 | 12.23794 | 13.6704 | 13.25874 | 16.35982 | 13.00712 | 15.28521 | 14.6051 | 18.8553 | 18.12752 |
| 1700019G1 | 1.562319 | 1.708778 | 3.181262 | 2.442692 | 2.440655 | 2.415665 | 3.038017 | 2.406227 | 1.761015 | 1.649321 | 2.056264 | 2.037191 |
| Elavl2 | 10.39745 | 9.593104 | 4.995723 | 5.361169 | 4.902411 | 4.605436 | 4.569024 | 4.137634 | 5.11693 | 5.96046 | 5.834422 | 6.126429 |

| | | | | | | | | | | | | |
|---|---|---|---|---|---|---|---|---|---|---|---|---|
| Stxbp1 | 4.947086 | 6.268084 | 7.594409 | 7.917675 | 10.41063 | 9.973408 | 11.84429 | 11.7228 | 11.29871 | 11.23179 | 8.465876 | 8.542324 |
| N4bp3 | 27.41555 | 27.64447 | 16.10863 | 14.34665 | 11.46283 | 10.64479 | 8.377548 | 12.01873 | 9.860759 | 9.69103 | 12.33156 | 13.4572 |
| Dnajb2 | 2.932025 | 2.876896 | 4.362965 | 4.097547 | 4.668549 | 4.254721 | 4.421437 | 4.616393 | 4.228787 | 3.689131 | 5.908819 | 5.459152 |
| Alg10b | 15.14046 | 17.76563 | 26.35443 | 26.20455 | 24.49481 | 24.3221 | 25.8802 | 23.76678 | 30.40842 | 32.10394 | 32.87729 | 35.15968 |
| Amigo2 | 2.013045 | 2.139699 | 2.37718 | 2.749457 | 3.874706 | 3.652259 | 4.672202 | 4.064753 | 3.503672 | 3.377634 | 3.046863 | 3.58684 |
| Ube2t | 14.85341 | 13.11182 | 9.631531 | 9.635767 | 9.419807 | 9.240516 | 8.910274 | 9.217387 | 8.396711 | 8.520846 | 6.771796 | 6.518826 |
| Ipo13 | 12.13984 | 13.08478 | 24.86929 | 23.54959 | 19.95452 | 19.83894 | 20.17092 | 20.24606 | 19.25418 | 18.53475 | 28.55946 | 27.39336 |
| Ubn1 | 8.143865 | 9.797259 | 12.63045 | 12.16791 | 12.09678 | 11.82928 | 13.29184 | 13.25057 | 13.66356 | 15.30043 | 18.00305 | 17.48057 |
| 4930430F0 | 6.961513 | 5.527799 | 5.32549 | 5.125262 | 5.928862 | 5.323211 | 4.515654 | 5.734873 | 4.73189 | 5.101149 | 3.440131 | 3.881802 |
| Ccdc74a | 1.027943 | 1.133371 | 2.277095 | 1.831145 | 1.627022 | 1.892513 | 1.410038 | 1.583099 | 1.362995 | 1.353132 | 2.027572 | 1.998687 |
| Dyrk1b | 4.042492 | 4.819263 | 7.705647 | 8.305571 | 8.949142 | 8.803274 | 12.16455 | 11.89248 | 10.6825 | 9.423246 | 8.610995 | 8.05201 |
| Cenpb | 13.30831 | 12.67839 | 22.4738 | 19.83153 | 24.0206 | 23.59343 | 26.32259 | 25.6781 | 28.54171 | 26.36346 | 31.64769 | 29.2843 |
| Cep170 | 7.001609 | 7.792737 | 9.891345 | 10.80061 | 11.43301 | 11.11759 | 11.88959 | 11.86894 | 14.47465 | 15.30375 | 14.88073 | 15.94717 |
| Rfxap | 3.907509 | 4.134937 | 5.37195 | 5.138977 | 6.566401 | 7.360858 | 7.047557 | 6.93685 | 8.029569 | 7.381817 | 7.629722 | 7.582485 |
| Socs1 | 2.196699 | 2.073118 | 3.095815 | 2.933211 | 3.532807 | 3.97865 | 4.365391 | 3.53424 | 3.980804 | 3.265895 | 4.606386 | 4.325311 |
| Paqr3 | 1.281487 | 1.657416 | 2.063442 | 2.175088 | 2.140139 | 1.978879 | 2.464297 | 1.828679 | 2.583997 | 2.676841 | 2.713817 | 2.695991 |
| Fam20b | 8.272066 | 8.622756 | 12.22727 | 12.43758 | 14.32719 | 12.94592 | 15.83541 | 15.90296 | 18.13063 | 18.33339 | 17.09887 | 17.69267 |
| Slc43a1 | 2.60569 | 2.956073 | 4.558132 | 4.392115 | 4.128571 | 3.949428 | 4.517919 | 4.451598 | 5.249868 | 5.82157 | 5.890871 | 5.798369 |
| Padi2 | 1.524489 | 1.715903 | 0.941154 | 0.877067 | 1.143039 | 0.982995 | 0.948757 | 0.736286 | 0.841 | 0.947947 | 0.660432 | 0.666985 |
| Ppp1r13b | 1.825227 | 2.052995 | 4.161043 | 4.114626 | 4.364148 | 4.11009 | 4.285772 | 4.023669 | 3.176986 | 2.792009 | 4.532628 | 4.027188 |
| Ubc | 1.088132 | 1.255167 | 1.51187 | 1.59765 | 1.823075 | 1.841985 | 2.044437 | 1.575364 | 1.826566 | 1.893647 | 2.294769 | 2.321459 |
| Fut4 | 0.302437 | 0.364748 | 3.745134 | 3.512449 | 4.621338 | 4.410459 | 5.278792 | 4.847879 | 3.179774 | 3.290959 | 3.57813 | 3.467679 |
| Rnf215 | 3.839562 | 3.581439 | 5.013515 | 5.160635 | 5.869677 | 5.607188 | 5.289374 | 5.95217 | 6.595466 | 5.479217 | 8.267219 | 7.595304 |
| Kank4 | 2.994925 | 3.508742 | 7.183754 | 7.265625 | 7.217757 | 6.55837 | 6.829285 | 6.734395 | 6.419368 | 7.327845 | 4.585827 | 4.979164 |
| Tmem260 | 0.92909 | 0.797427 | 1.604352 | 1.517257 | 1.851825 | 1.994607 | 2.295741 | 1.85145 | 2.26664 | 2.232513 | 2.151338 | 1.859095 |
| Glra1 | 0.422127 | 0.540685 | 0.488715 | 0.918702 | 0.73371 | 0.882658 | 0.837106 | 0.862825 | 1.265465 | 1.268938 | 1.41543 | 1.098257 |
| Ncdn | 7.280032 | 8.616662 | 11.45275 | 10.55575 | 10.51381 | 10.68094 | 11.56372 | 10.04568 | 11.22311 | 9.809937 | 16.39766 | 14.41631 |
| Clec2l | 0.352763 | 0.398784 | 1.144827 | 1.608828 | 1.71992 | 1.726002 | 2.120425 | 2.363047 | 2.285098 | 1.8587 | 1.401093 | 1.836495 |
| Chd7 | 13.6006 | 17.68652 | 33.14745 | 34.91555 | 30.37016 | 29.90067 | 32.40689 | 27.71379 | 40.47393 | 40.68117 | 38.39075 | 39.24746 |
| Kif13a | 8.361792 | 9.802693 | 12.96308 | 12.71464 | 12.93826 | 11.90317 | 12.47958 | 12.15705 | 15.55662 | 15.73101 | 18.24984 | 17.45921 |
| Zfp532 | 35.32051 | 40.51977 | 23.27306 | 24.37039 | 16.76364 | 17.89293 | 16.47999 | 17.01625 | 17.7308 | 18.09981 | 16.50154 | 16.61903 |
| Slc35d1 | 4.074729 | 4.014755 | 7.961254 | 8.205656 | 7.462665 | 8.314282 | 8.025309 | 7.697925 | 6.647737 | 6.512136 | 9.240775 | 9.546684 |
| Rbpms2 | 117.4842 | 121.3135 | 81.20288 | 77.73128 | 74.68679 | 77.63332 | 78.51552 | 76.05349 | 68.64579 | 62.59649 | 55.42293 | 53.06864 |
| Ski | 27.67943 | 32.45444 | 86.9095 | 85.84067 | 97.64175 | 91.82723 | 94.51059 | 91.01823 | 72.91173 | 71.63831 | 64.07258 | 62.71038 |
| Gns | 12.22648 | 13.99836 | 17.07637 | 16.72846 | 18.46837 | 17.00776 | 20.93754 | 19.65865 | 25.58019 | 24.48393 | 22.5959 | 23.49578 |
| Kcnj10 | 0.190187 | 0.27135 | 0.647275 | 0.604515 | 0.730175 | 0.625634 | 1.114334 | 0.884121 | 0.740288 | 0.737272 | 0.726173 | 0.784259 |
| Slc35e3 | 6.81912 | 7.11353 | 9.41026 | 9.562196 | 11.08806 | 10.33635 | 13.46459 | 13.20408 | 13.92599 | 13.22446 | 13.77898 | 13.96671 |
| Mog | 0.863057 | 1.608551 | 0.459059 | 0.758729 | 0.68491 | 0.55416 | 0.839775 | 0.770116 | 0.429935 | 0.644449 | 0.353066 | 0.527266 |
| Dnajb12 | 7.126816 | 7.681595 | 10.46001 | 10.135 | 11.08553 | 10.42262 | 12.31555 | 10.93057 | 12.41797 | 12.08957 | 14.60192 | 14.79749 |
| F8a | 0.973741 | 0.895362 | 1.665153 | 1.724972 | 2.2824 | 2.143194 | 2.604893 | 2.24513 | 2.175874 | 2.093828 | 2.456403 | 2.518348 |
| Zfp61 | 3.267238 | 2.81563 | 5.330301 | 4.844069 | 6.057962 | 5.963859 | 6.790261 | 7.067859 | 7.191957 | 7.049036 | 5.907551 | 6.333934 |
| Sptb | 2.89868 | 3.634176 | 1.506196 | 1.503503 | 1.506952 | 1.329491 | 1.263708 | 1.068645 | 0.941425 | 1.050408 | 1.017028 | 1.030845 |
| Arsb | 0.171933 | 0.195383 | 0.830352 | 0.883611 | 1.20167 | 1.175654 | 1.28827 | 1.186657 | 1.368751 | 1.288021 | 0.945688 | 1.095212 |
| Adcy9 | 0.838991 | 1.053729 | 2.032449 | 1.783035 | 1.995498 | 1.721554 | 1.719566 | 1.60181 | 0.985524 | 1.274063 | 1.140966 | 1.223837 |
| Rian | 46.39423 | 41.65144 | 27.67757 | 27.92635 | 31.40344 | 31.56968 | 38.16418 | 37.2202 | 32.87069 | 33.8101 | 23.00379 | 25.59219 |
| Mmgt2 | 1.584917 | 1.996247 | 2.493685 | 2.678959 | 2.95097 | 2.786912 | 3.229886 | 3.391697 | 3.742218 | 3.894346 | 3.02287 | 3.249378 |
| Tctn1 | 2.947357 | 3.082763 | 6.519609 | 6.374045 | 5.740184 | 6.14944 | 5.23373 | 5.274647 | 4.007082 | 4.386531 | 4.343935 | 4.908767 |
| Dlx3 | 0.061743 | 0.105068 | 1.296682 | 1.187035 | 1.537469 | 1.269223 | 0.997505 | 0.880893 | 1.217146 | 1.279694 | 1.096978 | 0.919169 |
| Optn | 5.637537 | 6.30293 | 3.105013 | 3.165991 | 3.114023 | 3.183499 | 3.134682 | 2.958328 | 2.479657 | 2.191603 | 2.562351 | 2.660087 |
| Slc22a17 | 4.562614 | 4.827963 | 8.963327 | 8.046233 | 11.40673 | 11.40399 | 11.23309 | 11.68613 | 10.42606 | 9.61772 | 8.727971 | 8.218482 |
| Pvrl1 | 7.047454 | 8.996073 | 15.6142 | 15.00315 | 12.05028 | 11.79963 | 13.26457 | 10.33969 | 10.33489 | 10.4165 | 11.28335 | 10.81991 |
| Tmem80 | 2.201867 | 2.087313 | 3.575606 | 3.800721 | 4.762976 | 4.199032 | 4.349961 | 4.154423 | 2.96782 | 2.894858 | 3.955521 | 3.627136 |
| Csk | 14.02037 | 16.02556 | 22.89506 | 20.96475 | 24.91249 | 23.92089 | 28.76706 | 26.14529 | 31.08454 | 30.40949 | 33.97264 | 30.08561 |
| Nanog | 212.3403 | 198.7739 | 97.90982 | 96.77117 | 96.63439 | 92.13134 | 99.19887 | 110.0561 | 119.2337 | 133.586 | 105.0152 | 103.8516 |
| Epb4.1 | 12.4641 | 13.63736 | 18.42418 | 18.7489 | 21.27283 | 20.80963 | 23.33265 | 22.76605 | 26.1833 | 27.16483 | 21.18005 | 20.10768 |
| Msantd3 | 44.98188 | 37.8537 | 20.35606 | 20.36663 | 21.72422 | 21.30447 | 21.25382 | 23.1254 | 27.47539 | 25.60399 | 29.86814 | 29.49909 |
| Fam185a | 9.632197 | 9.333172 | 6.344297 | 5.920168 | 5.414145 | 5.117716 | 4.111837 | 4.087516 | 4.860444 | 5.06343 | 4.644463 | 4.62407 |
| Calcoco2 | 8.995987 | 7.321503 | 29.32459 | 27.98867 | 31.93027 | 30.48811 | 34.10236 | 35.47558 | 37.80354 | 40.53669 | 38.58473 | 35.78864 |
| Tmcc3 | 0.671034 | 0.698455 | 2.104457 | 1.738195 | 2.382502 | 2.056204 | 2.354253 | 2.517093 | 2.09401 | 2.048548 | 2.043186 | 2.055048 |
| Mast2 | 10.35859 | 12.5658 | 16.79636 | 16.28821 | 16.70377 | 16.10415 | 18.87977 | 17.01727 | 19.67269 | 20.50689 | 24.14544 | 23.31327 |
| Stx18 | 3.001664 | 3.297977 | 5.141144 | 5.158151 | 6.341368 | 6.278975 | 6.395697 | 7.353231 | 6.345826 | 6.023396 | 7.549068 | 7.467129 |

| | | | | | | | | | | | | |
|---|---|---|---|---|---|---|---|---|---|---|---|---|
| Ppp1r1a | 8.064954 | 7.619223 | 2.602151 | 2.974305 | 2.985222 | 3.020909 | 2.749622 | 2.967646 | 3.573066 | 3.618601 | 4.718974 | 4.296042 |
| Fbxo11 | 12.61028 | 15.37989 | 19.75725 | 19.79603 | 20.01073 | 19.9192 | 22.28213 | 21.16541 | 26.4193 | 27.01729 | 27.18231 | 28.64373 |
| Zfp296 | 19.73677 | 25.56783 | 9.268227 | 9.127063 | 7.199847 | 6.55603 | 7.770902 | 7.903078 | 7.242185 | 7.20363 | 7.656369 | 6.309471 |
| Man2b2 | 3.65665 | 3.971689 | 5.81504 | 5.871797 | 7.228027 | 7.019144 | 7.683201 | 8.205259 | 8.423702 | 8.491441 | 7.568163 | 7.292769 |
| Lrrc41 | 6.867692 | 7.220989 | 8.647425 | 9.122023 | 9.443318 | 9.367687 | 10.08601 | 9.449873 | 12.47817 | 12.46391 | 14.07337 | 13.18343 |
| Blvrb | 25.82692 | 22.03019 | 15.06313 | 15.35059 | 16.23664 | 16.44765 | 14.14474 | 15.13606 | 12.3745 | 12.39536 | 10.43069 | 11.382 |
| Trp53inp2 | 6.462392 | 7.091342 | 11.56297 | 11.17545 | 13.02151 | 11.75897 | 15.2249 | 12.42643 | 13.25914 | 12.72545 | 16.91518 | 16.14706 |
| Mrps31 | 53.16384 | 41.71318 | 37.53874 | 36.14575 | 32.31032 | 30.88618 | 26.71632 | 30.35987 | 25.82998 | 25.01734 | 29.5639 | 28.96457 |
| Fam110b | 3.976025 | 3.936689 | 3.065161 | 2.607859 | 2.398758 | 2.397755 | 2.343455 | 2.542918 | 2.114113 | 1.85734 | 1.84686 | 1.805435 |
| Slc4a2 | 3.62268 | 3.912428 | 7.815919 | 7.467069 | 8.220614 | 8.386877 | 9.63964 | 10.02411 | 7.442541 | 6.940159 | 7.532716 | 7.575209 |
| Sertad2 | 4.451881 | 5.667406 | 9.332718 | 9.515077 | 9.534315 | 8.641189 | 10.25072 | 8.291167 | 9.185027 | 9.298276 | 10.96697 | 11.9668 |
| Crebbp | 6.763178 | 9.863649 | 15.10652 | 16.20519 | 13.61883 | 14.91776 | 14.63704 | 11.23802 | 13.0141 | 13.13091 | 14.2753 | 14.50684 |
| Ttpal | 2.430121 | 2.470711 | 4.741765 | 4.357147 | 4.858154 | 4.262092 | 5.546913 | 4.726093 | 4.116158 | 4.123546 | 5.613083 | 5.946011 |
| Aldh18a1 | 12.3787 | 13.62749 | 24.77045 | 24.07549 | 24.32061 | 22.97288 | 24.23673 | 20.88888 | 23.8584 | 23.53806 | 30.90737 | 30.09276 |
| Sun1 | 41.74737 | 41.15767 | 29.41627 | 28.27559 | 26.944 | 26.81676 | 24.24898 | 26.92534 | 22.20773 | 23.01305 | 20.02692 | 19.63359 |
| Zfp386 | 3.420067 | 3.294016 | 5.117408 | 4.915888 | 5.317247 | 5.162185 | 5.052217 | 5.383978 | 7.401173 | 7.077841 | 5.772419 | 6.460014 |
| Cenpv | 77.29008 | 65.09974 | 43.09121 | 42.08771 | 46.95143 | 48.75286 | 37.20968 | 40.96329 | 39.53816 | 36.67909 | 38.7145 | 33.88149 |
| Kdm6b | 6.070004 | 9.2421 | 21.03342 | 20.95816 | 21.05763 | 19.74304 | 19.93048 | 15.51671 | 16.24411 | 16.57648 | 15.5377 | 14.59318 |
| Ube2r2 | 20.16807 | 20.73367 | 24.73945 | 25.66283 | 30.80045 | 27.80382 | 35.16879 | 34.90296 | 40.51589 | 36.84634 | 37.2714 | 36.03593 |
| Rbm15b | 11.10003 | 11.78698 | 16.12669 | 15.47881 | 16.26398 | 15.91562 | 18.49205 | 16.36137 | 19.81193 | 19.14319 | 22.84214 | 22.21525 |
| Stim2 | 2.931031 | 3.256855 | 5.102101 | 5.140696 | 5.664546 | 5.318107 | 6.286835 | 6.447028 | 6.120624 | 7.034196 | 7.962492 | 7.376011 |
| Hectd1 | 10.47013 | 12.99245 | 19.6964 | 19.91015 | 19.21284 | 18.62295 | 21.19248 | 18.89349 | 24.64681 | 25.82137 | 25.00307 | 26.71889 |
| 2310030G( | 3.262425 | 3.000136 | 1.305971 | 1.71711 | 1.18516 | 1.352086 | 0.970242 | 1.584877 | 1.66311 | 1.511485 | 2.006797 | 1.939661 |
| Pcolce2 | 4.115962 | 4.055595 | 1.931644 | 1.694817 | 1.415691 | 1.61834 | 1.333141 | 1.510138 | 1.140298 | 1.181416 | 1.369137 | 1.310126 |
| AA415398 | 0.444023 | 0.412618 | 0.950156 | 0.920762 | 0.991539 | 1.074531 | 1.097372 | 0.957473 | 0.795092 | 0.749733 | 1.113575 | 0.98189 |
| Usp49 | 1.136077 | 1.283509 | 1.642225 | 1.772516 | 1.874002 | 1.8716 | 1.953338 | 1.675447 | 2.495155 | 2.400715 | 1.938213 | 1.899053 |
| Cyp2s1 | 1.886849 | 1.841009 | 0.47403 | 0.545079 | 0.490899 | 0.541395 | 0.554612 | 0.607893 | 0.709723 | 0.689181 | 0.886234 | 0.77556 |
| Cdh24 | 2.394027 | 2.259793 | 5.428591 | 4.738039 | 4.966546 | 5.194718 | 5.707968 | 6.786107 | 6.841528 | 6.725705 | 6.674986 | 6.401906 |
| Gcnt2 | 10.10376 | 10.49212 | 3.983497 | 4.106081 | 3.456943 | 3.479279 | 3.272536 | 3.576247 | 3.200218 | 3.508919 | 4.762485 | 4.982827 |
| Cox6c | 95.52612 | 68.18899 | 47.02325 | 50.3476 | 51.89454 | 58.09966 | 45.2077 | 49.05772 | 45.81236 | 42.40305 | 41.1708 | 40.56977 |
| Rtn2 | 7.664196 | 8.177768 | 5.146594 | 5.177659 | 5.282681 | 4.535802 | 3.833032 | 4.21708 | 4.905575 | 4.361674 | 5.450307 | 6.037457 |
| Gemin6 | 42.13572 | 37.76965 | 28.51874 | 27.28801 | 28.33318 | 28.2471 | 22.97975 | 26.90518 | 20.98497 | 20.44704 | 20.0052 | 19.7369 |
| Sh3gl2 | 9.257387 | 9.217194 | 4.109817 | 3.872252 | 3.558207 | 3.494236 | 3.27292 | 3.162187 | 2.983936 | 2.840587 | 2.761901 | 2.706615 |
| Wt1 | 7.603951 | 7.978422 | 18.18822 | 18.18722 | 19.74395 | 20.19775 | 26.93191 | 26.01199 | 23.44488 | 23.06718 | 22.05338 | 22.58922 |
| Baiap2l1 | 1.770435 | 1.928782 | 5.326292 | 5.58411 | 7.102797 | 6.355992 | 7.739385 | 6.740662 | 7.458396 | 7.98511 | 6.153418 | 6.179957 |
| Cyth3 | 5.135072 | 5.36719 | 7.292368 | 6.921429 | 8.225452 | 7.827611 | 9.365882 | 9.328281 | 10.3164 | 9.892781 | 9.265889 | 9.886869 |
| Mapk4 | 5.876526 | 6.741971 | 3.696897 | 3.595216 | 4.125103 | 3.585148 | 3.621018 | 2.892791 | 2.487486 | 2.667012 | 3.891091 | 3.809081 |
| Aak1 | 2.145437 | 2.755885 | 4.084771 | 4.238481 | 4.707815 | 4.708752 | 5.7989 | 4.069017 | 4.873592 | 5.81077 | 4.775485 | 4.904601 |
| Socs2 | 39.67673 | 36.60006 | 15.91792 | 14.83721 | 14.54945 | 14.30045 | 14.98238 | 15.87648 | 17.7593 | 17.8188 | 12.71902 | 12.48399 |
| Zak | 4.144301 | 4.937812 | 6.205568 | 6.781015 | 6.896694 | 7.078054 | 8.014135 | 6.88622 | 7.383745 | 7.834235 | 9.022867 | 9.663996 |
| Cep70 | 13.53425 | 12.83151 | 10.2973 | 9.799275 | 9.115091 | 9.144207 | 8.185786 | 10.05322 | 7.556627 | 9.076182 | 6.6544 | 7.058037 |
| Fbln2 | 0.504661 | 0.629552 | 1.298634 | 1.315849 | 1.425607 | 1.492571 | 1.552687 | 1.204482 | 1.287368 | 1.259504 | 1.177372 | 1.048084 |
| Ly6e | 40.75944 | 42.39001 | 20.20699 | 20.8908 | 22.01142 | 20.80765 | 20.28576 | 21.985 | 21.18809 | 21.17938 | 15.91977 | 15.43955 |
| Als2 | 8.614889 | 10.42871 | 18.05266 | 18.25907 | 16.70993 | 15.63057 | 18.5174 | 17.11513 | 14.05322 | 14.6997 | 12.45915 | 12.83353 |
| Dnajc21 | 48.75802 | 49.69228 | 32.37409 | 31.40665 | 28.65243 | 26.66701 | 25.0232 | 28.95783 | 25.23538 | 24.53879 | 22.50952 | 22.07777 |
| Nrip1 | 1.672381 | 2.251495 | 19.12441 | 19.89851 | 22.1978 | 21.77417 | 25.96759 | 21.22362 | 19.08001 | 19.88335 | 28.62236 | 28.28362 |
| Cml1 | 0.47378 | 0.578943 | 1.283877 | 1.148003 | 1.065754 | 1.024081 | 0.7643 | 0.963966 | 0.635439 | 0.807664 | 0.954777 | 0.919157 |
| Rap1gap2 | 0.718682 | 0.909647 | 1.167972 | 1.150446 | 1.530876 | 1.635084 | 1.749377 | 1.584602 | 1.350824 | 1.564113 | 1.296871 | 1.248983 |
| Spen | 10.44112 | 13.9313 | 21.26101 | 21.06754 | 18.36879 | 17.72537 | 18.41734 | 14.83102 | 16.14847 | 15.77252 | 19.15413 | 18.38362 |
| Scap | 7.510497 | 9.049279 | 13.61786 | 13.56279 | 14.63903 | 13.49992 | 17.30584 | 16.31414 | 18.75795 | 16.78612 | 20.08864 | 19.16624 |
| Tbc1d4 | 0.314456 | 0.476765 | 0.892622 | 0.859294 | 1.04001 | 0.89763 | 1.105376 | 0.968717 | 1.264648 | 1.219038 | 1.017457 | 1.099168 |
| Lrba | 3.518818 | 4.490457 | 6.946903 | 6.826474 | 7.098021 | 6.797235 | 7.795836 | 6.549631 | 6.504888 | 6.929019 | 4.850243 | 4.885651 |
| Plxnc1 | 0.36044 | 0.440697 | 0.926397 | 0.996613 | 1.100609 | 0.962121 | 1.148241 | 1.363769 | 1.322585 | 1.141792 | 0.917551 | 1.035776 |
| Uchl3 | 128.6127 | 103.5672 | 86.79683 | 83.27952 | 84.29567 | 84.66589 | 68.05312 | 73.55459 | 67.49062 | 66.17161 | 59.06396 | 59.73228 |
| Bcr | 3.874941 | 5.523353 | 8.485639 | 8.513495 | 9.394162 | 8.597864 | 10.70274 | 9.678839 | 11.41301 | 10.90658 | 12.30206 | 12.06698 |
| Fam83d | 9.614412 | 9.927418 | 15.70585 | 15.88238 | 18.67627 | 17.0736 | 20.59806 | 19.95513 | 15.70324 | 14.09567 | 18.68091 | 17.23464 |
| Atp1b1 | 43.0321 | 47.41535 | 26.37447 | 25.19429 | 21.47984 | 21.04013 | 20.05162 | 21.05342 | 21.12431 | 22.07677 | 18.25511 | 17.80209 |
| Sac3d1 | 15.05964 | 14.4779 | 21.12198 | 18.4457 | 21.23536 | 22.38727 | 24.31935 | 25.30959 | 25.16173 | 20.94386 | 30.38409 | 26.08858 |
| Slc25a10 | 5.191132 | 4.939709 | 6.716708 | 7.847438 | 7.923821 | 8.747003 | 10.08884 | 11.09705 | 11.09894 | 10.6415 | 10.38765 | 9.495329 |
| Gfod2 | 2.176518 | 2.543848 | 2.920442 | 3.178011 | 3.429819 | 3.187305 | 3.745318 | 4.490711 | 4.611553 | 4.360836 | 4.201924 | 4.264452 |
| Foxo6 | 0.631811 | 0.849071 | 0.846865 | 0.85198 | 1.3777 | 1.172449 | 1.485686 | 1.147034 | 0.89731 | 0.974106 | 1.382472 | 1.203319 |

| | | | | | | | | | | | | |
|---|---|---|---|---|---|---|---|---|---|---|---|---|
| Tmco3 | 2.944664 | 3.032827 | 4.893892 | 5.342046 | 5.78762 | 5.417564 | 6.090139 | 6.834138 | 5.183073 | 5.127304 | 4.949269 | 4.843161 |
| Lrp1 | 4.577385 | 5.859418 | 11.24802 | 11.8765 | 13.35873 | 12.37886 | 17.38452 | 14.06831 | 14.34376 | 14.73406 | 11.00612 | 10.45055 |
| Sgk3 | 3.006711 | 2.832958 | 1.286536 | 1.411562 | 1.823225 | 1.639593 | 1.890182 | 1.550248 | 1.639256 | 1.724739 | 2.102504 | 1.989407 |
| Whsc1l1 | 5.078493 | 5.914761 | 7.723586 | 7.917163 | 7.566263 | 7.821928 | 7.955284 | 7.513576 | 9.658207 | 10.58138 | 10.25056 | 10.51724 |
| Foxp4 | 17.04164 | 20.94157 | 31.37667 | 31.13384 | 30.04538 | 28.53462 | 33.2729 | 29.8618 | 36.84184 | 35.03683 | 43.50941 | 40.16867 |
| Sec24d | 11.2412 | 12.40711 | 18.51519 | 18.1154 | 23.75951 | 23.60928 | 27.50709 | 25.48763 | 19.9629 | 19.97192 | 18.10759 | 18.66585 |
| Aldoa | 829.0662 | 829.0274 | 432.5061 | 427.6225 | 408.7707 | 391.1101 | 350.5247 | 361.7147 | 428.661 | 453.4896 | 360.3011 | 322.5904 |
| Gypc | 0.258749 | 0.300924 | 1.237794 | 1.165295 | 1.581476 | 1.425419 | 1.733191 | 1.547904 | 1.294221 | 1.140272 | 0.833427 | 1.101569 |
| Tulp4 | 4.052821 | 4.698666 | 6.168557 | 6.56017 | 7.067825 | 7.307586 | 8.078937 | 6.816943 | 8.786232 | 8.820655 | 8.593126 | 9.213917 |
| Kat6a | 11.985 | 14.50552 | 18.72309 | 19.76795 | 18.13599 | 18.26657 | 19.33234 | 17.38809 | 22.48513 | 23.7183 | 24.16788 | 24.41625 |
| Insr | 2.214239 | 3.012308 | 3.776438 | 4.027967 | 4.570528 | 4.241961 | 4.777353 | 4.198249 | 4.684229 | 4.978379 | 3.421728 | 3.552182 |
| Ppp1r9b | 7.389598 | 9.345213 | 23.46232 | 21.74311 | 23.07507 | 21.63112 | 24.15782 | 23.08007 | 24.08045 | 24.16684 | 33.75381 | 30.29812 |
| Thap2 | 2.021576 | 2.33275 | 3.399168 | 4.091439 | 3.907138 | 4.253361 | 4.786973 | 4.194384 | 4.965604 | 5.728665 | 5.463311 | 5.671717 |
| Trmt12 | 0.921461 | 1.049151 | 1.833865 | 1.745854 | 1.681218 | 1.880029 | 2.071674 | 2.154548 | 1.851914 | 1.539127 | 2.434901 | 2.123141 |
| Gm608 | 2.535894 | 3.258324 | 4.025711 | 4.436646 | 4.555395 | 4.471118 | 4.952699 | 4.071551 | 5.11013 | 5.697532 | 4.49439 | 4.721145 |
| Ncoa7 | 3.231564 | 3.790126 | 5.473257 | 5.345943 | 5.313143 | 5.503748 | 6.463559 | 6.132657 | 6.616674 | 6.872389 | 7.314877 | 7.480744 |
| Pgk1 | 444.0882 | 360.7283 | 159.7992 | 161.3659 | 173.2856 | 164.4678 | 153.9835 | 160.7092 | 193.3273 | 224.7406 | 171.0059 | 168.7953 |
| Tmem60 | 11.83062 | 9.683278 | 9.482721 | 8.873038 | 10.55284 | 9.563618 | 10.54669 | 8.253838 | 7.158268 | 7.877509 | 5.783545 | 6.839113 |
| Axin2 | 1.360141 | 1.496058 | 1.683132 | 1.86916 | 2.37358 | 2.436067 | 3.363885 | 2.299348 | 2.898052 | 2.61322 | 2.383665 | 2.490503 |
| Prrx2 | 2.152661 | 2.442706 | 5.174006 | 4.632854 | 5.198628 | 5.838405 | 7.189111 | 6.472277 | 6.742916 | 5.945522 | 7.063712 | 6.455194 |
| Fgf13 | 1.040268 | 1.154115 | 1.68962 | 1.987682 | 2.610056 | 2.31709 | 2.722755 | 2.775642 | 2.663258 | 2.624711 | 2.520398 | 2.260244 |
| Pld2 | 1.189672 | 1.311379 | 3.103073 | 3.316072 | 3.735828 | 3.639154 | 4.354347 | 4.444624 | 3.460233 | 3.850728 | 3.459757 | 3.184179 |
| Lrp2 | 6.537849 | 9.334888 | 18.19278 | 18.63501 | 15.97069 | 15.22125 | 19.60153 | 13.96009 | 17.27431 | 18.41625 | 17.71537 | 19.48118 |
| Setd1b | 5.604269 | 7.780894 | 12.11995 | 12.61555 | 10.97848 | 11.02201 | 12.31717 | 10.73362 | 10.63803 | 10.03085 | 14.73809 | 13.99796 |
| Ccdc160 | 3.007204 | 3.271397 | 6.19446 | 5.791123 | 5.037368 | 4.824402 | 4.289037 | 4.515839 | 4.704593 | 4.404619 | 4.779575 | 4.152891 |
| Map6d1 | 0.678749 | 0.677811 | 0.835513 | 0.924601 | 1.390632 | 1.097492 | 1.075558 | 1.288738 | 1.142088 | 1.137387 | 1.141446 | 0.962165 |
| Zfhx3 | 1.958239 | 3.034123 | 6.747627 | 8.265229 | 7.312374 | 7.748535 | 8.70117 | 6.545364 | 7.328212 | 7.166529 | 5.766936 | 5.930357 |
| Klhl18 | 2.173335 | 2.409404 | 4.083636 | 3.96277 | 4.577635 | 4.449925 | 4.844727 | 5.042406 | 4.175997 | 4.453899 | 5.520451 | 5.52173 |
| Cables2 | 2.772194 | 2.846656 | 3.998828 | 4.177239 | 4.773381 | 4.041416 | 5.206299 | 5.631038 | 5.152475 | 5.563839 | 5.614067 | 5.568694 |
| Simc1 | 3.832832 | 4.390494 | 5.485823 | 5.634658 | 6.284741 | 5.805253 | 6.002922 | 5.397642 | 7.141215 | 7.328826 | 7.704499 | 7.519863 |
| Rcsd1 | 1.117958 | 1.041597 | 3.345638 | 3.120265 | 2.927092 | 3.047147 | 4.22053 | 4.054793 | 3.506307 | 3.424111 | 3.735142 | 2.911514 |
| Col27a1 | 0.336514 | 0.455037 | 0.948111 | 1.121636 | 1.280751 | 1.289996 | 1.094138 | 1.391482 | 1.147828 | 1.30753 | 0.873415 | 0.999858 |
| Fastkd2 | 17.53585 | 15.02493 | 13.99829 | 13.67597 | 10.7426 | 10.3708 | 9.12959 | 10.30115 | 8.755261 | 9.838738 | 10.35953 | 11.12783 |
| Ctdnep1 | 12.41507 | 13.70842 | 19.5471 | 19.59139 | 20.79956 | 19.26048 | 22.74398 | 21.37393 | 25.04703 | 21.26025 | 26.01862 | 26.75846 |
| 2010315B0 | 0.445678 | 0.45141 | 0.897915 | 1.151274 | 1.028426 | 1.194075 | 1.150918 | 1.161403 | 0.898842 | 0.860009 | 0.969556 | 1.072458 |
| Mast4 | 1.409554 | 1.68675 | 1.973323 | 2.125995 | 2.08839 | 2.119607 | 2.551834 | 2.026971 | 2.835475 | 2.812467 | 2.574574 | 2.607218 |
| Gata2 | 0.36121 | 0.403399 | 1.745394 | 1.379074 | 1.398366 | 1.287065 | 1.48448 | 1.315923 | 1.047687 | 1.059938 | 2.131978 | 1.470276 |
| Nipa1 | 0.925026 | 0.846242 | 1.55488 | 1.367051 | 1.492473 | 1.529392 | 1.881336 | 1.26628 | 1.336602 | 1.425515 | 1.518291 | 1.754932 |
| Zbtbd6 | 0.999962 | 0.98881 | 2.415919 | 2.21074 | 1.972434 | 2.218174 | 2.244771 | 2.404649 | 1.597836 | 1.791447 | 2.955384 | 2.481598 |
| Mall | 0.321522 | 0.462653 | 1.132657 | 1.268054 | 1.215368 | 1.450196 | 1.279654 | 1.051355 | 1.12594 | 1.047458 | 1.259861 | 1.210769 |
| Fgd6 | 1.795771 | 2.234105 | 3.603888 | 3.999356 | 4.12017 | 4.06519 | 4.707117 | 4.323145 | 3.927367 | 4.204698 | 2.987427 | 3.124613 |
| H2-K1 | 15.04983 | 13.434 | 20.53862 | 21.61029 | 30.26856 | 25.33919 | 25.75413 | 28.91054 | 24.00338 | 24.20544 | 36.24595 | 29.89016 |
| Cnppd1 | 3.558379 | 3.456164 | 6.486666 | 6.443681 | 7.28488 | 6.648981 | 7.391273 | 7.613188 | 7.351765 | 6.815541 | 9.659509 | 8.902609 |
| Dync1h1 | 24.6809 | 35.7396 | 54.15592 | 53.35931 | 45.41732 | 42.45481 | 49.46541 | 38.62238 | 43.95755 | 47.00854 | 44.48417 | 43.24581 |
| Wbp1l | 11.0774 | 12.80644 | 18.13458 | 17.32882 | 18.97872 | 18.67519 | 22.48597 | 23.8988 | 21.64159 | 21.37659 | 20.37574 | 19.5394 |
| 3110062M( | 0.613561 | 0.725876 | 1.165599 | 1.012806 | 1.225441 | 1.050661 | 1.333552 | 1.094966 | 0.969139 | 1.089969 | 0.705505 | 0.794196 |
| Chmp4c | 13.32898 | 11.3952 | 5.340067 | 4.978051 | 4.274985 | 4.48765 | 3.744507 | 3.610988 | 3.843438 | 3.649494 | 3.580048 | 3.631472 |
| Xpr1 | 3.021934 | 3.726209 | 7.775801 | 7.735498 | 7.991971 | 7.49452 | 8.718606 | 7.865542 | 6.464887 | 7.178133 | 5.845983 | 5.705658 |
| Foxm1 | 15.30415 | 17.67868 | 24.21619 | 24.4755 | 26.81506 | 24.58673 | 27.26478 | 25.40987 | 29.12511 | 28.97557 | 34.37943 | 33.72712 |
| Sult5a1 | 1.000338 | 0.95996 | 2.068414 | 1.538216 | 2.119321 | 1.99123 | 2.264927 | 2.119208 | 1.853222 | 1.740907 | 1.227423 | 1.677706 |
| Pecr | 23.77463 | 20.35219 | 14.28024 | 14.08545 | 16.83133 | 15.97284 | 15.17676 | 14.73982 | 16.07488 | 15.88485 | 11.99593 | 11.60499 |
| Nudt4 | 67.25074 | 66.34849 | 119.3398 | 113.6341 | 128.6364 | 124.7227 | 143.4519 | 141.2685 | 151.7834 | 156.2142 | 122.3449 | 123.7097 |
| Sort1 | 8.804263 | 9.109077 | 14.59137 | 15.22938 | 18.35814 | 17.00754 | 20.56317 | 18.27513 | 20.2556 | 20.93324 | 18.65707 | 18.96922 |
| Wasf1 | 13.70867 | 16.68207 | 20.14793 | 20.27188 | 22.37266 | 19.99363 | 25.04518 | 23.23532 | 28.5961 | 26.46814 | 21.63398 | 19.26007 |
| Dcaf5 | 3.913756 | 4.284908 | 7.652559 | 7.614784 | 9.313539 | 9.041604 | 10.33989 | 9.567226 | 9.831685 | 10.45855 | 9.362233 | 9.26114 |
| Syvn1 | 7.249474 | 8.34328 | 15.10704 | 14.76111 | 14.11677 | 13.31873 | 15.21165 | 13.87633 | 11.87649 | 10.75016 | 12.59539 | 12.99802 |
| Abca3 | 2.024299 | 2.320925 | 5.899291 | 5.716192 | 7.228667 | 6.941094 | 8.258349 | 7.208095 | 7.434246 | 7.846151 | 7.582921 | 7.174033 |
| Plekhh1 | 4.77995 | 5.941325 | 7.796374 | 8.106305 | 8.945193 | 8.774011 | 8.65526 | 8.496117 | 8.823318 | 9.358575 | 11.76663 | 11.12499 |
| Pkm | 1552.424 | 1361.566 | 755.6665 | 734.7251 | 777.0955 | 738.841 | 705.2931 | 806.2411 | 853.7057 | 942.7619 | 726.6405 | 686.9824 |
| Caskin2 | 2.870976 | 3.738939 | 7.674288 | 8.080406 | 7.876119 | 7.489711 | 9.289194 | 8.433984 | 9.123012 | 9.095138 | 10.81147 | 10.27461 |
| Dgke | 5.084867 | 5.498531 | 3.760334 | 3.512069 | 2.709912 | 2.70377 | 2.30397 | 2.560368 | 2.876456 | 2.721741 | 2.689048 | 2.776865 |

| | | | | | | | | | | | | |
|---|---|---|---|---|---|---|---|---|---|---|---|---|
| Phip | 10.62338 | 13.31021 | 20.05246 | 20.72526 | 20.15819 | 18.61386 | 20.27207 | 17.43355 | 23.17305 | 23.87712 | 21.8361 | 23.32871 |
| Phf13 | 11.04302 | 12.70441 | 16.96566 | 16.91461 | 19.34931 | 18.75059 | 22.97889 | 20.76483 | 23.2937 | 23.15871 | 18.50588 | 18.14067 |
| Pde4dip | 2.604743 | 3.139161 | 5.95229 | 6.200548 | 6.127036 | 5.927503 | 6.010425 | 5.433136 | 4.260707 | 4.530519 | 4.801826 | 4.572139 |
| Tktl2 | 0.371367 | 0.420747 | 1.562538 | 1.44619 | 1.424265 | 1.317554 | 1.801193 | 1.802909 | 1.373332 | 1.509159 | 1.677533 | 1.666624 |
| Apbb2 | 2.896381 | 3.655533 | 5.077546 | 5.14983 | 4.364443 | 4.29536 | 5.307037 | 4.059523 | 5.153484 | 5.302834 | 5.894225 | 5.552112 |
| Med14 | 11.57598 | 14.11567 | 27.36339 | 28.60071 | 31.28851 | 31.00151 | 36.81512 | 32.22743 | 26.03086 | 27.09462 | 22.33764 | 23.56926 |
| Casp8 | 0.608386 | 0.909148 | 2.002808 | 1.666279 | 1.769444 | 1.838551 | 2.168454 | 1.903692 | 1.851433 | 1.392573 | 1.079358 | 1.510465 |
| Rpl26 | 1947.75 | 1540.458 | 931.6502 | 976.7419 | 1197.257 | 1202.474 | 1047.551 | 1334.606 | 1104.817 | 1088.857 | 1000.362 | 976.8869 |
| Tpd52l1 | 15.50006 | 13.63573 | 8.517322 | 9.47215 | 10.90127 | 9.732822 | 10.00804 | 9.950471 | 7.826045 | 7.284525 | 8.327712 | 7.44833 |
| 1700048O2 | 0.919517 | 0.65184 | 1.902966 | 2.009062 | 2.47716 | 2.549013 | 2.389385 | 2.711555 | 2.050641 | 1.985894 | 2.633387 | 2.50327 |
| Cdc42ep5 | 9.475512 | 8.147709 | 3.115005 | 3.209392 | 3.12573 | 3.595664 | 4.042071 | 3.977987 | 3.963454 | 2.914521 | 2.158942 | 3.053262 |
| Pde8a | 4.5868 | 4.749501 | 1.699208 | 1.731647 | 1.647397 | 1.724964 | 1.801001 | 1.787409 | 2.011824 | 2.353501 | 2.28926 | 2.223371 |
| 9130023H2 | 0.405247 | 0.308796 | 0.811263 | 1.09336 | 0.798977 | 0.988457 | 1.052783 | 1.060105 | 0.991345 | 0.876818 | 1.20749 | 1.195428 |
| Med13 | 5.752368 | 8.345346 | 12.73263 | 13.13616 | 12.26701 | 11.88468 | 12.55036 | 10.27912 | 12.7333 | 13.15083 | 13.52349 | 14.01229 |
| Vps37d | 1.330969 | 1.386157 | 1.858099 | 1.92987 | 2.243138 | 2.386003 | 2.448022 | 2.549764 | 2.816987 | 2.041363 | 2.957739 | 2.629582 |
| Inpp5e | 2.843675 | 3.40647 | 5.069285 | 5.205414 | 5.604959 | 5.909011 | 6.870389 | 6.600043 | 6.93025 | 7.08917 | 6.734475 | 6.261634 |
| Zhx3 | 0.459437 | 0.502147 | 0.774522 | 0.892305 | 1.025788 | 1.163893 | 1.290887 | 1.156607 | 1.145279 | 1.214456 | 1.070306 | 1.139342 |
| Plekho2 | 2.183421 | 2.246359 | 3.969637 | 3.898306 | 4.648529 | 4.420268 | 4.928727 | 4.72826 | 4.099107 | 3.693001 | 5.448671 | 5.026178 |
| Gcat | 26.18294 | 24.22844 | 17.10375 | 16.8957 | 15.07636 | 15.43705 | 13.17062 | 14.66454 | 20.23143 | 19.93847 | 17.82764 | 17.09265 |
| Lyn | 1.211488 | 1.088758 | 2.427485 | 3.050482 | 3.176664 | 3.381344 | 3.682134 | 3.924764 | 3.594549 | 4.147067 | 3.877708 | 3.693243 |
| Tcl1 | 31.65067 | 34.00786 | 3.938121 | 3.075458 | 2.464947 | 2.449402 | 2.311082 | 2.547853 | 3.004858 | 3.164826 | 2.485694 | 2.329796 |
| Dock3 | 1.058012 | 1.191121 | 2.249853 | 2.378569 | 2.491282 | 2.676385 | 2.442292 | 2.344907 | 2.221258 | 2.246249 | 1.926321 | 1.971043 |
| 2010012O0 | 2.157609 | 2.248286 | 3.758413 | 3.385911 | 4.405009 | 4.407464 | 4.028391 | 3.618415 | 4.45135 | 3.827886 | 4.550733 | 4.596763 |
| Zfp746 | 7.265456 | 6.749566 | 12.48852 | 12.04713 | 11.47058 | 11.17563 | 11.47779 | 11.50176 | 12.25008 | 11.80715 | 15.82969 | 15.16165 |
| Atat1 | 4.033404 | 4.395399 | 5.907026 | 6.373932 | 6.622413 | 6.375981 | 7.052752 | 7.580763 | 8.65811 | 8.646267 | 6.291202 | 7.04818 |
| Myb | 2.273066 | 2.667396 | 0.544981 | 0.654296 | 0.469097 | 0.497642 | 0.452758 | 0.482381 | 0.384562 | 0.553537 | 0.697012 | 0.551893 |
| Eepd1 | 0.851066 | 1.077287 | 1.945136 | 1.71324 | 1.599105 | 1.719527 | 2.148168 | 2.323561 | 1.818507 | 2.028119 | 1.900035 | 1.751007 |
| Cux1 | 3.049438 | 3.895426 | 5.410786 | 5.918842 | 5.899573 | 5.870517 | 6.192595 | 5.276283 | 6.574505 | 6.79079 | 7.072335 | 7.154819 |
| C2cd2l | 1.474122 | 1.761363 | 3.109608 | 2.881139 | 3.114475 | 3.310372 | 3.0712 | 3.48205 | 2.705427 | 2.568601 | 3.03258 | 2.880434 |
| Eif4g3 | 7.387043 | 8.607075 | 18.02785 | 18.2961 | 22.3183 | 22.06655 | 26.53681 | 24.79335 | 23.39215 | 23.1408 | 25.15258 | 24.62533 |
| Gltscr1 | 2.770722 | 3.758801 | 5.147254 | 5.149006 | 5.503066 | 5.362421 | 5.464941 | 4.966512 | 6.798823 | 6.729545 | 6.039172 | 5.409047 |
| Nfam1 | 0.559871 | 0.603058 | 1.213318 | 1.331007 | 1.657635 | 1.418567 | 1.517689 | 1.333947 | 1.445812 | 1.690542 | 2.259973 | 1.801942 |
| Hoxa7 | 0.337412 | 0.249787 | 4.462275 | 4.620858 | 5.859202 | 6.069734 | 7.748713 | 6.25356 | 5.394629 | 4.699347 | 6.048123 | 6.184874 |
| Fasn | 41.87138 | 58.84173 | 76.51404 | 76.70019 | 78.63089 | 70.46466 | 90.93529 | 74.27263 | 89.59796 | 92.62642 | 73.29601 | 68.45714 |
| Stx3 | 15.22635 | 13.02855 | 5.592328 | 5.7802 | 5.685017 | 5.393285 | 5.718885 | 5.218605 | 5.24621 | 5.548992 | 5.925371 | 7.053373 |
| Zfp398 | 0.92732 | 1.230997 | 2.454993 | 2.859161 | 2.412379 | 2.670104 | 2.475281 | 2.358738 | 2.823827 | 2.708723 | 3.006945 | 3.185291 |
| Bcorl1 | 1.98319 | 2.904494 | 4.294936 | 4.648208 | 5.25448 | 4.575166 | 6.002392 | 5.34383 | 5.667192 | 6.17798 | 6.045252 | 5.799388 |
| Ell2 | 19.42448 | 19.52167 | 14.44841 | 13.98176 | 13.25609 | 12.77432 | 13.57377 | 12.86371 | 10.36602 | 11.62712 | 9.505585 | 10.3563 |
| Adcy7 | 0.727829 | 0.743604 | 1.720192 | 1.605201 | 1.824836 | 1.799645 | 1.92051 | 1.738087 | 1.776125 | 1.666174 | 1.429546 | 1.334822 |
| 4933427D0 | 3.177921 | 3.404289 | 6.45789 | 6.305282 | 7.760004 | 7.398073 | 8.31847 | 7.53029 | 5.641151 | 6.461644 | 5.986091 | 5.586858 |
| Cacna2d1 | 0.69081 | 0.812594 | 1.262547 | 1.446374 | 1.592157 | 1.438237 | 1.804133 | 1.492822 | 1.152038 | 1.107603 | 1.127703 | 1.221025 |
| Rem2 | 0.588179 | 0.577416 | 0.922037 | 0.949047 | 1.197899 | 1.028369 | 1.187837 | 1.144313 | 0.704236 | 0.912267 | 0.853812 | 0.851051 |
| Ggcx | 2.946917 | 3.866631 | 6.394309 | 6.570361 | 6.120792 | 6.377922 | 6.391706 | 6.208409 | 5.643515 | 5.866284 | 4.821405 | 4.891045 |
| Gria3 | 0.643322 | 0.67887 | 1.450577 | 1.405179 | 1.419555 | 1.25344 | 1.432496 | 1.422397 | 1.237077 | 1.183427 | 0.882416 | 1.058953 |
| Etv3 | 3.556847 | 3.710594 | 5.835546 | 5.777706 | 6.367204 | 6.418029 | 7.536876 | 6.923914 | 7.104748 | 7.084586 | 8.133401 | 7.756139 |
| Zcchc2 | 3.522177 | 4.735984 | 5.586313 | 5.592735 | 5.853666 | 5.460168 | 6.229315 | 5.240165 | 6.559794 | 6.809508 | 7.238175 | 7.032056 |
| Tagap1 | 5.126123 | 3.559125 | 3.49634 | 3.501165 | 3.081898 | 2.711459 | 2.496403 | 2.901864 | 3.587871 | 3.054105 | 3.336534 | 3.637304 |
| Fkbp3 | 195.7178 | 159.6791 | 113.6997 | 103.5907 | 108.2843 | 106.6495 | 94.18397 | 107.1955 | 144.2862 | 135.0714 | 133.8639 | 138.9261 |
| Sema3f | 1.1534 | 1.177834 | 1.55916 | 1.913135 | 2.318225 | 2.07734 | 2.332972 | 2.486318 | 1.810293 | 2.065215 | 1.649898 | 1.854756 |
| Celsr1 | 2.365892 | 3.176994 | 4.995857 | 5.36866 | 5.057993 | 4.460789 | 4.635813 | 4.139573 | 4.232164 | 4.005144 | 5.33118 | 5.127085 |
| Irak4 | 0.897512 | 1.220222 | 2.451417 | 2.543388 | 2.302627 | 2.34381 | 2.791659 | 2.727209 | 2.117796 | 2.527862 | 2.14748 | 2.034808 |
| Rpl34-ps1 | 974.2445 | 835.8874 | 444.0016 | 534.1062 | 624.8086 | 626.5102 | 655.4388 | 656.1393 | 610.2271 | 597.4564 | 530.4794 | 489.9845 |
| Ddx58 | 7.134066 | 8.115406 | 2.029651 | 1.759643 | 1.336159 | 1.35545 | 1.543055 | 1.814678 | 1.494692 | 1.681869 | 1.62363 | 1.599276 |
| Adam11 | 0.485992 | 0.717793 | 1.21611 | 1.125822 | 1.424168 | 1.285664 | 1.441598 | 1.506508 | 1.243276 | 1.206351 | 1.098378 | 1.197725 |
| Mrm1 | 7.487963 | 7.022156 | 5.852183 | 4.884887 | 5.13906 | 4.973214 | 4.409163 | 4.541018 | 3.460811 | 3.718899 | 4.583525 | 4.320612 |
| Tpcn1 | 4.31702 | 5.009562 | 9.110338 | 9.221612 | 10.30819 | 9.706478 | 11.31896 | 10.01916 | 8.42525 | 8.251112 | 8.185551 | 7.494785 |
| Mb21d1 | 0.462564 | 0.41818 | 0.846997 | 0.860756 | 0.795575 | 0.752673 | 0.763507 | 0.925998 | 0.79415 | 0.850373 | 1.297683 | 1.005278 |
| Gfpt1 | 7.221507 | 8.677296 | 13.63258 | 13.42903 | 15.77217 | 14.90276 | 16.33684 | 15.58229 | 15.06926 | 16.69344 | 12.09073 | 12.50305 |
| Armc5 | 2.44948 | 2.899657 | 5.507038 | 5.496629 | 5.026135 | 4.453726 | 5.201938 | 5.024093 | 4.838148 | 4.411165 | 6.767249 | 5.883992 |
| Kcnh2 | 3.549726 | 3.773641 | 5.966109 | 6.079042 | 7.210225 | 7.012448 | 7.772355 | 8.094316 | 6.54523 | 6.079188 | 8.024749 | 7.34294 |
| Ammecr1l | 5.666618 | 5.97003 | 10.15125 | 10.14437 | 9.606201 | 9.809269 | 11.15914 | 10.51537 | 10.70618 | 10.40745 | 12.74705 | 13.0596 |

| | | | | | | | | | | | | |
|---|---|---|---|---|---|---|---|---|---|---|---|---|
| Stau1 | 14.36199 | 17.23091 | 23.14992 | 22.45275 | 25.03694 | 23.5669 | 27.65245 | 27.2178 | 29.79159 | 29.85903 | 29.83433 | 29.48069 |
| Ebp | 5.702225 | 6.382482 | 9.260663 | 9.032533 | 9.197886 | 9.379228 | 11.93856 | 11.34705 | 11.31736 | 10.70991 | 11.30161 | 12.01153 |
| Tmem106a | 3.202558 | 3.039937 | 1.336099 | 1.465198 | 1.368329 | 1.395357 | 1.552981 | 1.486312 | 1.266092 | 1.428736 | 2.055626 | 1.855194 |
| Trrap | 12.34389 | 16.02833 | 25.33811 | 26.17417 | 22.75833 | 22.97545 | 26.28435 | 20.76138 | 21.23377 | 22.92418 | 20.67802 | 19.68658 |
| Ticam1 | 1.091953 | 1.216141 | 1.574352 | 1.810476 | 1.791275 | 1.929819 | 2.046185 | 2.239971 | 2.248763 | 2.405624 | 2.015037 | 2.059791 |
| Stra13 | 44.95566 | 35.22367 | 23.0331 | 25.51965 | 25.47868 | 26.42139 | 22.95232 | 24.15829 | 21.1128 | 20.75654 | 21.0907 | 19.89079 |
| Dagla | 0.411976 | 0.497462 | 0.861425 | 0.888067 | 0.882861 | 0.818987 | 0.862237 | 0.806906 | 0.772352 | 0.702613 | 1.152638 | 0.933079 |
| Tmc3 | 0.223754 | 0.21697 | 0.904857 | 1.137896 | 0.967733 | 1.056353 | 1.209971 | 1.069784 | 0.798233 | 0.857142 | 0.914188 | 1.061148 |
| Ptpdc1 | 1.632105 | 1.71038 | 3.603151 | 3.192605 | 3.597954 | 3.490447 | 3.362233 | 3.631928 | 3.916238 | 3.772732 | 4.092963 | 4.235414 |
| Camk2b | 10.60546 | 10.58586 | 5.138183 | 4.865952 | 6.462883 | 6.454766 | 6.848888 | 6.248945 | 6.434692 | 6.091406 | 4.844512 | 4.931561 |
| Zfp628 | 2.380878 | 3.398366 | 5.665476 | 5.652712 | 5.536809 | 4.774478 | 5.674969 | 4.947377 | 6.072328 | 5.694693 | 6.697699 | 6.332258 |
| Dysf | 0.66732 | 0.792771 | 0.968438 | 0.963589 | 1.045089 | 0.928993 | 0.951493 | 0.994109 | 0.760719 | 0.753033 | 1.435934 | 0.998075 |
| Keap1 | 2.811528 | 3.053732 | 5.202215 | 4.896533 | 5.270281 | 5.405981 | 5.213097 | 4.686709 | 4.931021 | 5.236323 | 6.725227 | 6.21113 |
| Gba2 | 5.078936 | 5.460056 | 9.35991 | 9.582681 | 9.784526 | 9.347052 | 10.20544 | 10.44724 | 7.979176 | 8.298086 | 7.695593 | 7.886789 |
| Celsr2 | 1.682856 | 2.251068 | 3.305957 | 3.24173 | 3.450671 | 3.312512 | 3.898814 | 3.400716 | 4.119014 | 4.291576 | 3.68186 | 3.732972 |
| Tepp | 0.6633 | 0.652667 | 0.938705 | 1.478634 | 1.599327 | 1.349338 | 1.392351 | 1.11715 | 1.168779 | 1.167771 | 1.591201 | 1.273217 |
| Nhp2 | 232.3323 | 196.5885 | 157.4988 | 151.835 | 134.8574 | 138.8948 | 112.1757 | 119.8664 | 128.6457 | 114.7287 | 160.929 | 150.0449 |
| Slc26a2 | 4.550022 | 5.587726 | 10.41304 | 10.2789 | 9.138744 | 9.35927 | 8.806023 | 7.767197 | 8.206613 | 7.91341 | 8.375528 | 8.581481 |
| Guca1a | 1.781342 | 1.775382 | 2.478919 | 2.854192 | 4.072581 | 3.467042 | 3.951248 | 3.766586 | 3.442309 | 3.322865 | 4.213704 | 2.849319 |
| Fzd4 | 1.056497 | 1.080884 | 2.089789 | 2.011326 | 2.524419 | 2.411122 | 2.643003 | 2.658133 | 2.431624 | 2.350181 | 2.239456 | 2.390763 |
| Bace1 | 2.707045 | 3.409098 | 5.185748 | 5.42684 | 6.40765 | 5.835597 | 6.920577 | 5.577942 | 6.078013 | 6.175404 | 6.933331 | 7.170141 |
| Pcdhga12 | 16.72144 | 18.03501 | 23.92033 | 24.53512 | 27.76857 | 28.03855 | 33.62033 | 28.74761 | 31.63964 | 30.09075 | 32.46571 | 32.40944 |
| Uxt | 16.87827 | 12.05887 | 7.259043 | 8.985781 | 9.236985 | 9.322738 | 8.035115 | 10.28398 | 8.789923 | 7.786613 | 8.32428 | 8.198722 |
| Slc41a3 | 0.635194 | 0.691493 | 1.210644 | 1.038808 | 1.377405 | 1.237397 | 1.588186 | 1.360156 | 1.152981 | 1.312451 | 0.719233 | 1.029953 |
| Tmem181a | 7.733881 | 7.989936 | 12.68938 | 11.95504 | 13.73594 | 13.66903 | 15.38835 | 15.12084 | 15.42256 | 15.14851 | 16.84174 | 15.78761 |
| Rpl31 | 728.6614 | 535.7074 | 357.4105 | 396.6135 | 479.6036 | 523.5576 | 434.8384 | 447.8036 | 445.993 | 409.7238 | 448.4475 | 405.2457 |
| Slc25a36 | 54.31968 | 58.59555 | 35.24267 | 35.41757 | 31.79317 | 30.61955 | 32.96386 | 31.46291 | 33.44396 | 36.20601 | 24.57048 | 27.44659 |
| Mier3 | 6.464436 | 7.072346 | 11.31005 | 11.38636 | 11.61617 | 11.24751 | 11.49758 | 11.4349 | 13.04964 | 13.38271 | 13.62347 | 14.83253 |
| Qdpr | 88.54513 | 79.19507 | 48.60273 | 48.59467 | 46.69189 | 43.81569 | 42.67346 | 43.21815 | 41.00531 | 38.33587 | 38.13593 | 36.86723 |
| Iqsec2 | 0.526811 | 0.766785 | 1.374787 | 1.508796 | 1.780575 | 1.76405 | 2.184468 | 1.900543 | 1.827306 | 1.624458 | 1.571613 | 1.442468 |
| 1700001O2 | 1.059979 | 1.177061 | 2.172232 | 2.232937 | 2.540352 | 2.75243 | 2.904119 | 2.690933 | 2.854518 | 2.358158 | 2.248773 | 2.089064 |
| Igf2bp1 | 42.26233 | 59.18376 | 83.27474 | 82.60019 | 79.7676 | 74.98791 | 92.15129 | 66.33444 | 78.78907 | 81.11519 | 86.8296 | 87.34629 |
| Itga5 | 18.56035 | 21.74856 | 35.30517 | 33.22574 | 29.95162 | 29.09651 | 31.07848 | 29.37889 | 27.41501 | 27.53656 | 38.58867 | 36.25759 |
| Zfp637 | 4.030156 | 4.024671 | 6.041939 | 5.630165 | 6.231829 | 5.672617 | 6.927832 | 6.813887 | 7.478726 | 7.052889 | 8.17089 | 7.407679 |
| Wdr91 | 8.822167 | 9.422669 | 19.95774 | 19.92885 | 19.79879 | 19.48788 | 19.10327 | 19.327 | 15.69631 | 15.15216 | 16.68692 | 15.53301 |
| Slc51a | 0.390076 | 0.357644 | 1.591913 | 1.289826 | 1.309431 | 1.353196 | 1.780033 | 1.435305 | 1.60796 | 1.173251 | 1.076744 | 1.091536 |
| Mageb16 | 8.724632 | 7.97542 | 4.376693 | 5.072286 | 6.091323 | 6.148105 | 6.302937 | 4.676597 | 6.121583 | 5.824024 | 5.764135 | 4.825195 |
| Zfyve16 | 3.208772 | 3.301657 | 5.448503 | 5.711888 | 5.881528 | 5.858475 | 6.54623 | 6.706185 | 6.690865 | 7.100882 | 6.957029 | 7.504631 |
| Raet1d | 1.258536 | 1.398212 | 2.970545 | 2.838834 | 3.165792 | 3.038163 | 3.650794 | 3.321808 | 2.674296 | 2.936576 | 2.390509 | 2.528625 |
| Snrk | 2.952969 | 2.994594 | 4.525578 | 4.580748 | 5.079139 | 5.130878 | 5.518674 | 5.458894 | 5.4364 | 5.774203 | 6.876699 | 6.012768 |
| Sgpl1 | 10.90622 | 11.01745 | 16.45666 | 16.01998 | 18.28668 | 17.64016 | 22.64403 | 20.26859 | 19.57605 | 21.01798 | 20.51814 | 20.11433 |
| Rpl22 | 328.1049 | 240.0558 | 159.9818 | 168.3553 | 191.0395 | 200.5939 | 168.6399 | 200.082 | 167.7912 | 171.8633 | 154.262 | 152.9303 |
| Mpp2 | 1.044917 | 1.222003 | 2.133524 | 2.043197 | 2.275002 | 2.086701 | 2.490984 | 2.143347 | 2.15605 | 2.16746 | 1.709725 | 1.653931 |
| Efs | 7.859673 | 9.616421 | 13.19446 | 13.09454 | 12.32939 | 12.51185 | 13.67608 | 13.58371 | 13.20928 | 12.55017 | 17.64789 | 16.39517 |
| Sepw1 | 39.87266 | 38.62794 | 67.15063 | 65.7285 | 69.47187 | 71.92621 | 68.8303 | 74.334 | 81.16525 | 58.0642 | 77.13337 | 76.0838 |
| Aldh3a2 | 10.67214 | 9.839467 | 19.41253 | 19.54021 | 22.04762 | 21.97165 | 27.02733 | 25.57903 | 22.26013 | 21.12055 | 21.55889 | 22.07957 |
| Fbxo22 | 45.88447 | 44.13631 | 31.82008 | 27.96998 | 24.94159 | 26.3097 | 26.44393 | 23.81079 | 21.86328 | 21.39191 | 23.42886 | 22.58097 |
| Eda | 0.612266 | 0.724221 | 1.368631 | 1.263425 | 0.994968 | 1.036723 | 1.198164 | 1.138397 | 1.240585 | 1.148569 | 0.843152 | 0.966855 |
| Rnpep | 7.562776 | 8.264586 | 12.49513 | 12.21851 | 13.69963 | 13.88976 | 15.4985 | 15.95693 | 16.56544 | 14.51081 | 15.98038 | 14.40782 |
| Dip2b | 6.499983 | 8.796858 | 11.75589 | 12.66771 | 11.91803 | 11.67471 | 12.31802 | 9.753023 | 11.67666 | 12.42373 | 13.08318 | 12.7273 |
| Fbrs | 6.615488 | 8.25438 | 12.67759 | 11.63341 | 10.60866 | 11.12243 | 12.39316 | 12.30924 | 11.36843 | 10.66455 | 13.74905 | 13.91628 |
| Prdx1 | 711.4905 | 565.8295 | 386.4081 | 395.5651 | 424.3384 | 415.5898 | 397.1008 | 446.4427 | 381.1849 | 377.5945 | 332.898 | 342.2464 |
| Sema5b | 4.391064 | 4.91485 | 3.000802 | 2.800558 | 2.292632 | 2.379579 | 2.198959 | 2.148663 | 2.866801 | 2.387097 | 2.845256 | 2.764129 |
| Nbea | 1.641567 | 1.963286 | 3.637577 | 3.598624 | 3.817471 | 3.431972 | 3.99618 | 3.531258 | 3.242333 | 3.200516 | 2.738763 | 2.761711 |
| Zcchc11 | 14.50796 | 18.55031 | 25.47799 | 26.3873 | 30.8049 | 29.36727 | 33.80699 | 32.07307 | 30.17554 | 31.20536 | 22.89711 | 24.65491 |
| Rab13 | 3.21956 | 4.211889 | 8.569389 | 8.305414 | 10.27884 | 9.65422 | 10.88387 | 8.710986 | 9.98204 | 8.460041 | 8.427451 | 7.425785 |
| Mgme1 | 21.29032 | 19.20913 | 14.27311 | 14.06854 | 12.09087 | 11.98207 | 10.82853 | 11.43623 | 10.65612 | 10.82219 | 11.8218 | 11.71414 |
| Klf2 | 53.31161 | 58.04633 | 19.1252 | 17.68005 | 15.21265 | 14.38268 | 14.24846 | 15.1487 | 14.41025 | 13.60766 | 18.74036 | 14.66712 |
| Tbc1d9b | 4.640181 | 5.441781 | 8.137712 | 8.712318 | 10.08094 | 9.768436 | 11.88373 | 10.76154 | 11.26522 | 10.8725 | 9.366029 | 8.368262 |
| 3110021A1 | 0.442914 | 0.647653 | 0.86711 | 0.962122 | 1.276801 | 1.138718 | 1.092319 | 1.198793 | 1.006862 | 1.004416 | 0.907738 | 0.92311 |
| Prkd2 | 1.846648 | 2.152711 | 2.921682 | 3.066428 | 3.383237 | 3.432759 | 3.537048 | 3.232213 | 3.065693 | 3.414898 | 3.746709 | 3.980899 |

| | | | | | | | | | | | | |
|---|---|---|---|---|---|---|---|---|---|---|---|---|
| Arhgef10 | 4.179589 | 4.166909 | 9.525921 | 9.62916 | 10.35762 | 10.18518 | 11.2476 | 10.66548 | 8.493028 | 8.95527 | 8.218286 | 8.215778 |
| Nabp1 | 15.10975 | 16.12767 | 9.810367 | 10.06328 | 8.450816 | 8.063409 | 7.41111 | 8.150373 | 7.361544 | 8.14313 | 7.941956 | 7.753422 |
| Fam222b | 6.43934 | 7.378381 | 11.51659 | 11.78969 | 11.02799 | 10.76254 | 12.63816 | 10.78864 | 13.34183 | 12.69741 | 14.54734 | 13.95679 |
| Prdm5 | 21.82693 | 22.85055 | 12.45781 | 11.41179 | 10.16564 | 11.04029 | 10.37034 | 10.60607 | 10.10881 | 10.92548 | 8.289065 | 8.80242 |
| Secisbp2l | 5.560482 | 6.367178 | 9.100553 | 9.075812 | 9.935731 | 9.655377 | 10.55798 | 9.922281 | 9.84986 | 10.64545 | 11.8235 | 11.75636 |
| Eno1 | 1088.026 | 1031.578 | 652.7336 | 643.8577 | 601.0698 | 570.9142 | 498.963 | 551.2837 | 613.9343 | 631.5167 | 522.8475 | 497.193 |
| Tor3a | 9.875588 | 10.26899 | 3.398258 | 3.370859 | 3.250977 | 3.329937 | 3.262566 | 3.556346 | 4.315914 | 4.308384 | 3.693654 | 3.565443 |
| Gch1 | 0.762888 | 0.8605 | 1.373336 | 1.329344 | 1.581613 | 1.535491 | 1.726476 | 1.570681 | 1.314607 | 1.214369 | 1.430412 | 1.440358 |
| Crat | 2.752149 | 3.665789 | 5.108589 | 4.987531 | 5.808368 | 5.69812 | 6.752899 | 6.446444 | 5.930987 | 5.585831 | 4.900532 | 5.006716 |
| Ankrd52 | 6.487193 | 8.268291 | 10.7969 | 10.75299 | 11.02207 | 11.70913 | 11.84042 | 11.15748 | 13.10344 | 12.61167 | 14.33724 | 13.89657 |
| Glis2 | 9.03215 | 11.05901 | 33.30432 | 33.43411 | 39.84777 | 38.84546 | 46.62616 | 41.54452 | 39.62456 | 39.21862 | 45.82763 | 43.18746 |
| Nme4 | 47.977 | 39.69042 | 24.25943 | 23.1688 | 25.6759 | 26.88098 | 23.30882 | 21.48885 | 28.88205 | 27.68037 | 26.99611 | 25.19564 |
| Zfp592 | 4.127263 | 5.412306 | 7.958504 | 7.897618 | 7.837622 | 7.546939 | 8.050385 | 6.88523 | 7.835081 | 7.89793 | 9.167543 | 8.857886 |
| Ptger1 | 0.734074 | 0.714282 | 1.431832 | 1.48599 | 1.942587 | 1.789009 | 1.907893 | 1.818971 | 1.562298 | 1.472672 | 1.978485 | 1.8939 |
| Ak3 | 9.171116 | 9.274589 | 17.3967 | 16.9778 | 18.29228 | 17.63381 | 20.06892 | 20.08828 | 20.66864 | 21.83149 | 18.6196 | 20.203 |
| Pgm2l1 | 3.022091 | 3.401747 | 6.32507 | 6.10245 | 6.577307 | 6.097281 | 6.978055 | 7.217092 | 7.542779 | 7.790297 | 6.268117 | 6.059653 |
| Snx10 | 30.2526 | 29.27793 | 13.03613 | 11.85026 | 13.70004 | 13.1645 | 14.7113 | 14.02233 | 15.08191 | 15.22499 | 13.6236 | 13.09788 |
| Gbf1 | 9.541588 | 12.00996 | 20.17495 | 19.46089 | 19.26927 | 18.93787 | 20.44235 | 17.88942 | 14.65641 | 15.15305 | 15.7981 | 15.38424 |
| Katnal1 | 1.621747 | 1.742329 | 2.837993 | 3.23598 | 3.034981 | 3.166548 | 3.026911 | 3.08614 | 3.091611 | 3.606693 | 3.291114 | 2.808926 |
| Zbtb7a | 6.23434 | 7.255523 | 4.531717 | 4.02185 | 3.639041 | 3.818975 | 3.366552 | 3.433342 | 3.099562 | 3.134926 | 3.906197 | 3.310937 |
| Ptprs | 17.53074 | 20.13636 | 27.71236 | 29.18811 | 31.76451 | 32.18982 | 32.11115 | 31.67545 | 36.28478 | 37.40842 | 38.19768 | 37.14276 |
| Chsy1 | 3.339805 | 3.231704 | 5.078204 | 5.740909 | 6.597855 | 6.643463 | 7.780052 | 6.534624 | 7.254842 | 7.350473 | 6.576266 | 6.636115 |
| Fhod1 | 3.271347 | 3.915215 | 7.892348 | 7.588361 | 8.01369 | 7.723992 | 7.88247 | 8.090671 | 6.43254 | 6.334915 | 7.459199 | 7.020742 |
| Hspa12b | 0.502903 | 0.516477 | 1.017318 | 1.171809 | 1.404708 | 1.248793 | 1.61356 | 1.455585 | 1.413233 | 1.399731 | 1.181849 | 1.140277 |
| Fcgrt | 1.86782 | 2.156391 | 3.029127 | 3.036005 | 2.827688 | 2.705533 | 2.572401 | 3.074136 | 4.223057 | 3.431935 | 2.969299 | 3.453425 |
| Pcgf3 | 8.241058 | 10.49049 | 15.19327 | 15.00264 | 15.6239 | 14.30288 | 17.12404 | 14.06378 | 17.67075 | 17.93895 | 16.24387 | 16.48648 |
| Cdyl | 38.87232 | 33.58528 | 19.77972 | 19.74308 | 19.78409 | 18.71393 | 17.54044 | 18.06513 | 15.44146 | 15.48548 | 17.51941 | 17.00584 |
| Inpp1 | 0.824954 | 0.915218 | 1.424403 | 1.29696 | 1.665113 | 1.520848 | 1.573922 | 1.373424 | 1.250606 | 1.215064 | 1.48239 | 1.548605 |
| Agpat3 | 7.020291 | 7.4246 | 14.84662 | 15.02961 | 19.1083 | 19.08834 | 21.42806 | 19.15438 | 16.25714 | 15.73068 | 19.33311 | 18.60991 |
| Zfp277 | 25.91637 | 23.12497 | 14.55864 | 14.05405 | 15.02227 | 13.79597 | 13.33287 | 15.35146 | 12.51365 | 14.36859 | 10.1633 | 12.05589 |
| Ptpn23 | 12.2197 | 15.15492 | 25.87442 | 24.00819 | 21.07655 | 19.6193 | 23.34831 | 18.87507 | 21.33331 | 20.36586 | 24.49462 | 23.36117 |
| Prr12 | 9.347731 | 13.71767 | 20.86319 | 19.88833 | 18.61783 | 16.80152 | 19.11331 | 15.64962 | 17.80488 | 17.39754 | 19.95301 | 17.69183 |
| Rad1 | 14.91255 | 12.3387 | 8.700553 | 8.891004 | 8.286327 | 8.337708 | 7.942398 | 8.245755 | 8.276764 | 8.182715 | 6.909877 | 7.007218 |
| Zfp59 | 13.2499 | 12.45674 | 5.469814 | 5.514026 | 5.256105 | 5.350534 | 5.289923 | 5.947964 | 6.14118 | 6.786261 | 5.606891 | 6.622947 |
| AI848285 | 1.911674 | 2.552393 | 5.926425 | 5.241908 | 6.316149 | 5.877674 | 5.678399 | 6.157151 | 5.375914 | 4.855639 | 6.929553 | 5.794723 |
| Cacybp | 225.769 | 189.3611 | 145.8949 | 138.2947 | 127.811 | 126.0249 | 111.4437 | 123.9817 | 132.8649 | 132.8411 | 103.733 | 109.0771 |
| Fbxo46 | 3.165435 | 3.269151 | 5.355932 | 4.597872 | 4.847574 | 4.857349 | 5.587476 | 4.810127 | 5.520406 | 4.760172 | 6.438075 | 5.821795 |
| Mical1 | 8.125162 | 8.021159 | 4.936495 | 4.513444 | 4.482252 | 4.110636 | 3.708593 | 3.857374 | 3.587792 | 3.839399 | 3.701511 | 3.566076 |
| Kif3c | 1.645547 | 1.978912 | 2.61785 | 2.976315 | 3.561518 | 3.465727 | 3.559129 | 3.756758 | 3.20263 | 3.533858 | 3.47775 | 3.472088 |
| Hexb | 18.34836 | 16.80002 | 10.993 | 11.79002 | 12.13269 | 11.33047 | 10.94138 | 10.75678 | 10.64829 | 10.22581 | 8.910847 | 9.343957 |
| Csrnp2 | 3.479248 | 4.141362 | 6.990442 | 7.645241 | 8.474321 | 8.359452 | 9.395672 | 9.416582 | 9.155229 | 9.646448 | 9.665663 | 9.291024 |
| Tex19.1 | 14.74194 | 14.2005 | 32.17175 | 29.86648 | 29.73473 | 30.77708 | 37.3165 | 33.89471 | 25.59315 | 26.89016 | 37.69252 | 33.26663 |
| Zfp612 | 1.33276 | 1.368276 | 3.22911 | 3.339589 | 3.176455 | 3.361384 | 3.171283 | 3.340375 | 3.144481 | 2.849707 | 3.703883 | 3.591294 |
| Bbx | 8.091134 | 9.753623 | 12.5687 | 12.93582 | 14.509 | 13.62154 | 16.37927 | 13.60502 | 15.15836 | 16.29319 | 12.15851 | 13.25164 |
| Tle4 | 39.34704 | 44.44806 | 23.36715 | 22.77735 | 20.44529 | 18.91612 | 19.93088 | 18.08833 | 21.59138 | 20.96022 | 18.51734 | 18.84449 |
| Wdfy3 | 5.715654 | 7.299114 | 10.08319 | 10.65445 | 9.790318 | 10.39206 | 11.63194 | 9.800915 | 11.22514 | 11.61223 | 12.44562 | 12.33887 |
| Zfp691 | 1.999005 | 2.105548 | 3.732973 | 3.855971 | 3.535542 | 3.696121 | 4.351304 | 3.063078 | 3.293202 | 3.433547 | 3.017453 | 3.086154 |
| Gnl3 | 184.9283 | 164.2883 | 119.521 | 114.4254 | 106.7242 | 106.2031 | 92.08493 | 99.68279 | 133.2821 | 125.4811 | 110.31 | 117.6762 |
| Gdpd5 | 1.334458 | 1.259751 | 8.843492 | 9.76851 | 8.767527 | 8.375573 | 8.903996 | 10.11398 | 8.28442 | 8.067265 | 10.23308 | 10.14836 |
| Rasal2 | 2.850146 | 3.57183 | 4.392279 | 4.629639 | 4.494502 | 4.722577 | 5.014856 | 4.921348 | 5.430141 | 5.821021 | 5.74903 | 5.756833 |
| Spry2 | 40.13388 | 42.44398 | 18.85168 | 18.9026 | 17.64541 | 17.18863 | 20.52676 | 17.66788 | 17.39024 | 16.96843 | 15.20671 | 15.05172 |
| C2cd3 | 4.301696 | 5.21397 | 7.551675 | 7.87021 | 7.201392 | 7.04511 | 8.640271 | 7.213556 | 6.986515 | 7.232848 | 6.112947 | 6.273731 |
| Mitf | 3.386987 | 3.824198 | 1.734925 | 1.537316 | 1.745594 | 1.405989 | 1.333422 | 1.392282 | 1.393796 | 1.433586 | 0.985748 | 1.226107 |
| Cpeb4 | 2.517665 | 2.876553 | 5.094414 | 5.216482 | 5.132713 | 5.209837 | 5.934686 | 4.892677 | 5.528326 | 5.532195 | 5.944081 | 6.10895 |
| Vps9d1 | 3.579333 | 4.047013 | 6.652082 | 6.436517 | 6.720839 | 6.584634 | 6.083604 | 6.088518 | 6.600319 | 6.529174 | 8.023712 | 7.595518 |
| Pnpla3 | 1.43754 | 1.654133 | 0.771725 | 0.717185 | 0.907069 | 0.775755 | 0.564047 | 0.747449 | 0.636171 | 0.622037 | 0.786888 | 0.631697 |
| Plxna1 | 13.80427 | 17.86773 | 22.31006 | 22.0914 | 25.26879 | 24.79029 | 29.58116 | 24.81229 | 24.38915 | 25.26342 | 28.26468 | 27.54286 |
| Rnls | 1.199585 | 1.321405 | 2.1341 | 2.093563 | 2.056128 | 2.187542 | 2.609115 | 2.347757 | 1.815418 | 1.956257 | 1.78009 | 2.435257 |
| Rgs9 | 0.908254 | 0.842311 | 1.485976 | 1.527381 | 2.039572 | 1.726463 | 2.061485 | 1.664161 | 1.503834 | 1.688688 | 1.592386 | 1.38722 |
| Lrp5 | 4.939801 | 6.967973 | 8.853616 | 9.071166 | 9.533994 | 8.843883 | 10.43494 | 10.11334 | 9.771343 | 9.751278 | 8.372086 | 8.133913 |
| Dap | 4.289688 | 5.043241 | 7.950029 | 6.959416 | 9.220883 | 8.211214 | 9.929095 | 8.650067 | 9.093634 | 7.495789 | 6.557061 | 7.382819 |

| | | | | | | | | | | | | |
|---|---|---|---|---|---|---|---|---|---|---|---|---|
| Trip11 | 2.838289 | 3.441406 | 5.421745 | 5.771472 | 5.741293 | 5.513062 | 5.961841 | 5.853046 | 6.562546 | 6.484152 | 6.443553 | 6.847672 |
| Map3k5 | 0.973939 | 1.343919 | 3.102515 | 2.987977 | 3.507475 | 3.250483 | 4.357192 | 3.70637 | 2.943986 | 3.104598 | 3.267992 | 3.06819 |
| Sat1 | 29.66084 | 23.49891 | 16.38145 | 14.59395 | 14.3172 | 14.72055 | 13.58005 | 13.41536 | 16.16123 | 16.64009 | 14.08641 | 13.92969 |
| Cul7 | 12.27006 | 14.78052 | 25.41456 | 23.92087 | 24.78081 | 22.79342 | 25.80261 | 24.45903 | 21.84396 | 21.77926 | 29.4396 | 26.99582 |
| Sugt1 | 99.20259 | 82.60479 | 55.52976 | 55.61059 | 49.66085 | 48.58822 | 43.89888 | 51.98379 | 42.94513 | 45.40361 | 44.6355 | 44.88945 |
| Adam22 | 1.406356 | 1.567773 | 2.64925 | 2.466789 | 2.618015 | 2.724326 | 2.912546 | 2.514449 | 2.66479 | 2.903265 | 2.130656 | 2.344513 |
| Klc3 | 3.399679 | 3.009673 | 2.112372 | 2.209517 | 2.300041 | 1.987309 | 1.508828 | 2.070929 | 1.667068 | 1.915526 | 1.577291 | 1.785893 |
| Zfp69 | 1.629781 | 1.788425 | 2.709865 | 2.988674 | 3.495709 | 3.181504 | 3.525806 | 3.492213 | 3.623377 | 3.695994 | 3.19796 | 3.157374 |
| Rcor1 | 8.513899 | 9.535819 | 15.15328 | 14.82714 | 16.57868 | 16.46449 | 18.12752 | 15.89721 | 17.84019 | 18.17979 | 17.30975 | 18.28713 |
| Rnf165 | 4.591628 | 6.128863 | 9.05306 | 9.038435 | 9.788979 | 9.257302 | 11.4606 | 8.750663 | 10.46548 | 10.36422 | 8.733308 | 8.409354 |
| Klhdc8b | 1.603111 | 1.498039 | 2.913439 | 3.393213 | 3.163969 | 3.318415 | 3.443113 | 3.755297 | 3.093014 | 3.132727 | 3.386548 | 3.853364 |
| Ildr1 | 9.982249 | 11.36254 | 3.840367 | 3.496964 | 3.880502 | 3.512663 | 3.662885 | 3.574347 | 3.864739 | 3.701938 | 3.474369 | 3.011732 |
| Phka1 | 1.631982 | 1.741999 | 3.075684 | 3.154678 | 3.460709 | 3.222543 | 3.872847 | 3.732506 | 3.53722 | 3.545308 | 2.980413 | 3.045618 |
| Tmem86b | 1.142331 | 1.68334 | 2.330611 | 3.135105 | 2.969946 | 3.19763 | 3.250182 | 2.727511 | 2.73504 | 2.620895 | 2.459005 | 2.61625 |
| Akap13 | 2.853108 | 3.871387 | 5.771986 | 6.217603 | 6.343168 | 6.147194 | 7.495501 | 6.261595 | 6.561525 | 7.060903 | 5.68544 | 5.779216 |
| Hmbox1 | 6.754309 | 8.095275 | 11.35462 | 11.41126 | 12.1406 | 11.92975 | 13.27633 | 11.23228 | 13.67987 | 13.72256 | 12.93464 | 13.62819 |
| Pard3 | 11.80313 | 13.53502 | 20.48708 | 19.87323 | 20.92915 | 20.21702 | 21.99941 | 20.64086 | 22.87049 | 23.37179 | 25.81713 | 24.17652 |
| Tbc1d8b | 0.585645 | 0.756606 | 0.892494 | 0.901127 | 1.209853 | 1.035744 | 0.983746 | 1.094853 | 1.020432 | 1.09075 | 0.831747 | 0.944798 |
| BC017158 | 1.692619 | 1.66438 | 3.254603 | 3.357811 | 3.030461 | 2.985524 | 3.351879 | 2.838709 | 3.354854 | 3.153475 | 3.844486 | 3.448616 |
| Dmxl2 | 3.907506 | 4.253648 | 6.848559 | 7.001577 | 7.584825 | 7.202663 | 7.916334 | 7.245986 | 6.482106 | 7.236433 | 5.802249 | 5.971469 |
| Zfp777 | 8.808867 | 8.390905 | 18.80334 | 17.32994 | 18.22482 | 17.77974 | 17.01963 | 16.94222 | 18.08186 | 16.91508 | 21.67404 | 20.52404 |
| Med21 | 21.95301 | 16.5427 | 13.32236 | 13.86539 | 14.16562 | 13.12087 | 11.1264 | 13.29679 | 11.99652 | 11.54091 | 11.00388 | 11.44517 |
| Rragb | 21.6992 | 20.61336 | 13.13314 | 13.07417 | 12.09695 | 11.07372 | 11.26281 | 12.76495 | 13.53908 | 13.37095 | 10.40765 | 11.70405 |
| Ssh2 | 1.717892 | 2.227225 | 5.386887 | 5.393534 | 5.898039 | 6.15357 | 7.248323 | 5.9286 | 5.406996 | 5.523539 | 5.060008 | 5.104373 |
| Nf1 | 3.571338 | 5.389823 | 7.131029 | 8.051468 | 7.463674 | 8.067065 | 8.056337 | 6.898118 | 7.703039 | 8.309455 | 7.928692 | 8.155668 |
| Gas2l1 | 5.772648 | 6.464784 | 10.67984 | 10.00153 | 10.15739 | 9.593597 | 10.74867 | 10.237 | 10.17885 | 8.986967 | 12.78043 | 10.82292 |
| Pramel1 | 0.974513 | 1.164245 | 2.393856 | 2.154805 | 2.191231 | 2.017575 | 2.666153 | 2.264516 | 1.908628 | 1.919171 | 2.715001 | 2.307226 |
| Nckap5l | 1.901015 | 2.687231 | 5.947197 | 5.407192 | 5.572434 | 5.012221 | 5.986796 | 5.639889 | 5.947401 | 5.168928 | 5.158936 | 4.693895 |
| Fam193a | 8.791379 | 11.10573 | 16.03504 | 15.65279 | 15.62553 | 15.45736 | 17.36412 | 15.12852 | 17.96146 | 18.16965 | 17.87593 | 18.03052 |
| Synj1 | 3.066611 | 3.765885 | 5.894672 | 5.936592 | 6.091124 | 5.651171 | 6.189735 | 4.944511 | 5.068032 | 5.245215 | 5.796031 | 5.879159 |
| Kremen1 | 3.758401 | 4.627228 | 6.262197 | 6.253932 | 6.718576 | 6.853398 | 8.130468 | 7.21923 | 6.991267 | 7.24838 | 7.613426 | 7.505708 |
| Garem | 2.614994 | 3.501671 | 4.731772 | 5.253355 | 5.100155 | 4.954698 | 5.277072 | 4.379212 | 4.859299 | 5.223493 | 5.685041 | 5.163697 |
| Zfp97 | 13.45322 | 11.72314 | 7.157455 | 7.588441 | 6.928113 | 6.327864 | 6.313634 | 6.94974 | 7.413798 | 7.478641 | 5.839192 | 6.718104 |
| Atad2b | 7.626773 | 8.719369 | 14.62772 | 15.19129 | 15.6005 | 14.57545 | 14.4838 | 15.04586 | 15.5808 | 17.00439 | 17.18657 | 17.47603 |
| Ntan1 | 28.45947 | 22.07026 | 16.97163 | 16.82332 | 17.35019 | 17.74993 | 16.0684 | 17.43057 | 14.56549 | 14.80393 | 13.69142 | 15.45112 |
| Cnnm4 | 1.185148 | 1.809958 | 3.426142 | 3.541563 | 4.512707 | 4.042674 | 4.95245 | 4.01983 | 3.311814 | 3.934305 | 3.774896 | 3.384061 |
| Fam53c | 10.23452 | 12.06282 | 18.26843 | 17.15867 | 17.393 | 17.86559 | 21.08343 | 18.14987 | 18.68385 | 18.90193 | 21.71835 | 20.46647 |
| Tspan33 | 1.272589 | 1.085427 | 1.818127 | 2.276937 | 2.756943 | 2.276754 | 2.604831 | 2.713496 | 2.498235 | 2.391189 | 2.907399 | 2.694098 |
| Ctdsp2 | 23.74945 | 25.82347 | 45.3293 | 44.73843 | 54.24154 | 53.1461 | 58.3918 | 52.99878 | 46.23689 | 47.16264 | 43.77459 | 44.61257 |
| Tradd | 2.420572 | 2.651798 | 3.792153 | 4.158139 | 4.578174 | 4.124713 | 4.666454 | 4.883723 | 4.62784 | 4.302598 | 4.999657 | 4.101975 |
| Mark4 | 6.250884 | 8.171575 | 11.77808 | 12.22467 | 14.18978 | 13.26913 | 15.39988 | 13.98336 | 14.06675 | 13.55196 | 13.88958 | 11.74715 |
| Ehd3 | 1.020444 | 1.127494 | 2.213604 | 2.328438 | 2.642503 | 2.657426 | 2.917762 | 2.48557 | 2.858436 | 2.756073 | 2.852361 | 2.766416 |
| Zfp960 | 11.2673 | 9.599439 | 6.638062 | 6.377392 | 5.886124 | 5.739448 | 5.501978 | 6.045099 | 6.659185 | 6.805294 | 5.416253 | 6.099554 |
| Stk17b | 11.70485 | 10.99065 | 6.099802 | 6.340102 | 6.483852 | 6.685444 | 6.833184 | 6.988078 | 5.515253 | 6.679321 | 6.238053 | 5.898823 |
| Camk1 | 1.388973 | 1.256393 | 3.710684 | 3.069633 | 3.478369 | 3.451981 | 3.719835 | 3.635558 | 4.262 | 3.500314 | 3.312523 | 3.434986 |
| Zfp574 | 7.416356 | 8.526622 | 14.51406 | 14.49447 | 13.466 | 12.94231 | 13.84798 | 13.59648 | 13.56025 | 12.24027 | 15.83169 | 14.14657 |
| Elf4 | 1.990882 | 2.694575 | 3.806676 | 3.575262 | 3.945954 | 3.426556 | 4.277094 | 3.87993 | 3.766176 | 4.048487 | 4.15977 | 4.045797 |
| Mex3a | 17.02729 | 18.97082 | 27.26966 | 27.70207 | 32.27602 | 32.39061 | 34.84648 | 29.24935 | 32.70218 | 32.69942 | 30.54446 | 31.38552 |
| Add1 | 23.75037 | 24.48653 | 43.99369 | 43.51006 | 45.04542 | 43.53353 | 47.90401 | 47.67106 | 40.91031 | 40.45281 | 41.17597 | 41.13925 |
| Hectd2 | 0.983802 | 1.004413 | 1.631806 | 1.78352 | 1.940865 | 1.896377 | 1.722955 | 1.872306 | 2.00661 | 2.035408 | 1.957241 | 2.095627 |
| Mkl1 | 5.510932 | 7.374689 | 10.34707 | 10.22002 | 10.07885 | 9.516774 | 11.03408 | 9.561228 | 9.985469 | 10.0498 | 11.44211 | 11.1742 |
| Parp4 | 1.29131 | 1.611809 | 2.877623 | 3.124876 | 3.606773 | 3.070362 | 3.815345 | 3.588227 | 2.842292 | 3.096096 | 3.806592 | 3.596626 |
| Ciz1 | 4.37853 | 5.293278 | 9.003748 | 8.248765 | 8.527667 | 8.360542 | 8.518704 | 8.660076 | 9.611136 | 9.484922 | 9.823274 | 9.661858 |
| Hcfc2 | 2.839036 | 2.792446 | 6.113994 | 6.020433 | 7.199705 | 6.222878 | 6.208597 | 5.902898 | 5.103145 | 5.314263 | 5.393559 | 5.707733 |
| Fam222a | 1.14286 | 1.584246 | 2.053597 | 1.93601 | 1.842237 | 2.034741 | 1.762484 | 1.919471 | 1.90245 | 2.105378 | 2.480763 | 1.7994 |
| Rundc3a | 4.196171 | 4.181263 | 2.823152 | 2.539111 | 2.94414 | 2.656722 | 2.038272 | 2.90915 | 2.208741 | 2.535554 | 3.107951 | 2.95952 |
| Arf3 | 36.92434 | 43.27821 | 64.78154 | 63.80448 | 68.08259 | 64.65843 | 73.41042 | 66.71915 | 64.80688 | 63.11801 | 75.43859 | 73.57669 |
| Nr2c1 | 2.764337 | 2.783891 | 5.793484 | 5.200179 | 5.164645 | 4.760448 | 5.635896 | 5.173966 | 5.318871 | 5.540428 | 5.498559 | 5.362017 |
| Efnb3 | 1.754064 | 1.859955 | 3.207528 | 3.19478 | 3.506543 | 3.205773 | 3.475641 | 3.209873 | 4.012066 | 3.712954 | 3.740943 | 3.365521 |
| Gstm1 | 20.69039 | 24.64846 | 10.19417 | 9.457891 | 10.40802 | 9.320335 | 9.928158 | 8.144475 | 10.00904 | 9.31331 | 8.307247 | 8.296359 |
| Sec22c | 2.724269 | 3.050386 | 5.973742 | 5.767285 | 6.134406 | 5.656852 | 5.739603 | 4.988329 | 5.4896 | 5.387364 | 5.237748 | 5.294534 |

| | | | | | | | | | | | | |
|---|---|---|---|---|---|---|---|---|---|---|---|---|
| Nit2 | 46.40172 | 42.95677 | 32.72577 | 31.86213 | 32.53886 | 33.1385 | 29.34215 | 29.6947 | 29.36645 | 28.82384 | 23.05539 | 28.68928 |
| Cdc42bpb | 13.26474 | 15.88262 | 23.1653 | 22.9251 | 22.91003 | 21.73495 | 24.36744 | 23.7126 | 22.81791 | 22.87582 | 26.68315 | 25.08286 |
| Tex11 | 3.314101 | 3.159606 | 6.925549 | 6.622811 | 7.195333 | 6.181796 | 6.467063 | 7.120137 | 5.584918 | 6.605236 | 5.804441 | 6.298746 |
| Zfp324 | 1.355966 | 1.477172 | 2.335533 | 2.416787 | 2.819099 | 2.453582 | 2.719802 | 2.335705 | 2.224048 | 2.132777 | 2.276615 | 2.229248 |
| Abcg2 | 53.79743 | 48.5139 | 25.83157 | 27.4323 | 26.95415 | 27.35594 | 26.22553 | 28.27428 | 26.53606 | 29.24144 | 30.29771 | 31.46521 |
| Tpd52 | 63.00913 | 61.05204 | 32.78668 | 31.30518 | 29.03161 | 29.53976 | 28.12285 | 27.28681 | 29.53083 | 30.71523 | 28.76375 | 28.9984 |
| Lrp10 | 7.015787 | 7.809042 | 12.80281 | 12.31003 | 13.64299 | 13.72408 | 15.12743 | 13.34278 | 12.74934 | 11.99069 | 13.70119 | 13.48658 |
| Fbxo6 | 18.8589 | 15.3252 | 9.784993 | 9.4368 | 9.78436 | 10.15197 | 8.995527 | 9.141259 | 8.929149 | 9.21293 | 11.88998 | 10.4404 |
| Cep350 | 3.402471 | 4.18934 | 6.317699 | 6.506113 | 7.161545 | 6.85614 | 7.71608 | 6.618704 | 7.028538 | 7.429927 | 7.17399 | 7.190472 |
| Col9a2 | 2.365806 | 2.352978 | 5.283366 | 4.582882 | 4.930581 | 4.029937 | 3.289359 | 4.4559 | 4.329811 | 4.013416 | 4.948013 | 4.262674 |
| Mycbp2 | 5.33175 | 6.621125 | 10.16536 | 10.74383 | 10.70527 | 10.47578 | 11.35705 | 9.488486 | 9.852013 | 10.63576 | 8.945588 | 9.419724 |
| Kif7 | 2.576456 | 3.471166 | 5.972856 | 5.649748 | 6.137619 | 5.688367 | 6.033142 | 6.195016 | 5.66942 | 5.433545 | 6.417542 | 6.068684 |
| Acox1 | 6.500462 | 7.11199 | 13.00085 | 12.39501 | 12.26355 | 12.40295 | 14.26582 | 12.06061 | 10.7745 | 10.88418 | 11.01241 | 11.00823 |
| Arhgef11 | 5.258927 | 6.478885 | 8.965186 | 9.756662 | 9.798587 | 9.696444 | 11.2133 | 10.63778 | 9.336406 | 9.338137 | 10.64843 | 10.57241 |
| Snap91 | 0.394844 | 0.498181 | 0.814611 | 0.792333 | 0.993168 | 0.899908 | 1.087577 | 0.883789 | 0.848858 | 0.838646 | 0.887697 | 0.869544 |
| Tacc1 | 6.25421 | 8.007488 | 11.95903 | 11.7145 | 11.96674 | 12.05403 | 12.6385 | 11.66348 | 10.65834 | 11.30896 | 12.75153 | 12.7782 |
| Capn15 | 2.979479 | 3.673607 | 6.347074 | 6.25745 | 5.771828 | 5.754041 | 5.397693 | 6.111362 | 5.92367 | 5.745138 | 7.257321 | 6.482997 |
| Invs | 1.588476 | 1.848311 | 2.976736 | 3.119129 | 3.25067 | 2.924338 | 3.395469 | 3.294757 | 2.868945 | 3.136068 | 3.109323 | 3.091886 |
| Tram2 | 1.649578 | 2.070629 | 3.536965 | 3.722792 | 3.902247 | 3.771086 | 4.308804 | 3.703453 | 3.92144 | 3.865213 | 3.360744 | 3.431999 |
| Cog1 | 6.019475 | 7.238626 | 11.19233 | 10.87622 | 10.0146 | 9.895664 | 10.35654 | 9.754548 | 9.737747 | 9.66538 | 12.20958 | 10.6199 |
| Acot2 | 1.562074 | 1.606437 | 3.388357 | 3.263087 | 3.291736 | 3.201098 | 3.472845 | 2.902516 | 2.512181 | 2.955261 | 3.332163 | 3.088647 |
| Tmem181b | 8.825131 | 10.2517 | 26.05516 | 25.76364 | 25.84269 | 26.02682 | 26.60904 | 25.65519 | 25.44792 | 25.65807 | 21.26422 | 23.18795 |
| Inpp5a | 7.956609 | 8.99791 | 13.47719 | 14.0503 | 14.55302 | 14.36516 | 14.42358 | 15.50627 | 13.5199 | 14.79307 | 16.27445 | 15.74211 |
| Strn | 3.906063 | 4.632108 | 7.173125 | 7.219881 | 7.369683 | 7.340557 | 8.210687 | 7.136789 | 7.457149 | 7.994049 | 7.471549 | 7.754041 |
| Cd55 | 3.265964 | 2.626542 | 1.631401 | 1.719785 | 2.057132 | 1.881802 | 1.714785 | 1.735264 | 1.685316 | 1.72095 | 1.832791 | 1.926136 |
| Dag1 | 35.2068 | 42.66478 | 86.08678 | 83.76288 | 87.38538 | 82.46921 | 100.6527 | 82.74722 | 84.92543 | 83.45073 | 76.73006 | 74.88941 |
| Zfp874a | 2.590345 | 2.346375 | 1.859362 | 1.54817 | 1.596647 | 1.552474 | 1.339448 | 1.389285 | 1.280762 | 1.400622 | 1.285214 | 1.434592 |
| Synrg | 4.204783 | 5.151248 | 9.533499 | 9.869301 | 9.863456 | 9.996057 | 10.76792 | 9.676424 | 8.534131 | 9.118746 | 8.976156 | 8.952944 |
| Daglb | 1.901493 | 2.130204 | 3.818544 | 3.999966 | 3.564304 | 4.077678 | 4.234557 | 3.995514 | 4.633666 | 4.46092 | 4.488666 | 4.18775 |
| Bend4 | 14.1366 | 18.06506 | 34.0882 | 33.12458 | 35.35642 | 33.26627 | 38.6462 | 33.75494 | 32.05162 | 33.9907 | 30.38883 | 30.52156 |
| Nacc2 | 1.220434 | 1.330605 | 2.762084 | 2.828597 | 3.243071 | 2.776018 | 3.388827 | 3.05426 | 2.663165 | 2.757297 | 2.638443 | 3.055951 |
| Afap1 | 3.144892 | 4.071537 | 5.615547 | 5.888527 | 6.854664 | 6.391897 | 7.127162 | 6.458565 | 5.954958 | 6.070075 | 6.660955 | 6.488953 |
| Ssh1 | 3.380788 | 4.213222 | 7.334708 | 7.419628 | 7.759233 | 7.548876 | 8.373392 | 7.327844 | 6.7156 | 6.855979 | 7.37816 | 7.249473 |
| Zdhhc8 | 4.632762 | 5.603231 | 8.475819 | 8.469637 | 9.266236 | 8.736008 | 9.920928 | 9.06142 | 9.934185 | 9.241612 | 9.071858 | 8.817822 |
| Inppl1 | 15.43505 | 18.25114 | 28.38698 | 27.40587 | 29.97148 | 28.00451 | 32.63233 | 28.26351 | 27.09131 | 26.76311 | 27.69471 | 25.90783 |
| Tspan9 | 8.738798 | 10.03297 | 26.0382 | 25.40864 | 27.83549 | 26.99686 | 26.52611 | 25.48912 | 24.15718 | 24.83019 | 28.76975 | 26.8609 |
| Zfp963 | 0.694103 | 0.775238 | 1.114565 | 1.113115 | 1.213336 | 1.123467 | 1.075435 | 1.186509 | 1.420307 | 1.174983 | 1.101447 | 1.130207 |
| Wdr6 | 22.36832 | 25.11538 | 51.80081 | 46.14376 | 48.53312 | 45.26137 | 56.34422 | 47.37104 | 49.93 | 44.66387 | 47.07746 | 45.2947 |
