## Supplementary Data 2 for "Inferring the Hoxa1 Gene Regulatory Network in Mouse Embryonic Stem Cells: Time-Series RNA-seq Data and Computational Modeling Approach"

TF list for NARROMI

| Sample ID | Avg. FC Wt RA/KO RA |
|---|---|
| Lmx1a | 2088.52 |
| Spz1 | 1567.61 |
| Thrsp | 805.953 |
| Onecut1 | 458.903 |
| Ascl1 | 370.332 |
| Sox10 | 9.314410574 |
| Otx2 | 5.821502095 |
| Polr2k | 5.821496506 |
| Pitx2 | 5.667863168 |
| Irx5 | 5.239363703 |
| Prrx2 | 5.239356186 |
| Batf3 | 4.657211316 |
| 2410141K09Rik | 4.657198456 |
| Hes6 | 4.233815392 |
| Hoxc5 | 4.075052432 |
| Lmo1 | 4.075047903 |
| Rxra | 3.962723478 |
| Hoxa10 | 3.955838543 |
| Ssbp2 | 3.694071573 |
| Meox1 | 3.672020707 |
| Nr2f1 | 3.664405678 |
| Elavl2 | 3.490959007 |
| Mafb | 3.420124518 |
| Brdt | 3.201839689 |
| Snai1 | 3.201835926 |
| Cebpa | 3.104805314 |
| Snai3 | 3.104799563 |
| Pax6 | 3.068012103 |
| Nfkbie | 2.993923518 |
| Zfp61 | 2.910761241 |
| Sox21 | 2.910752504 |
| Maf | 2.910747685 |
| Pou5f1 | 2.885214789 |
| Sox3 | 2.818832787 |
| Hoxb5 | 2.75045088 |
| Pou3f3 | 2.716709665 |
| Fah | 2.716708432 |
| Hsf2bp | 2.716694597 |
| Lmo2 | 2.585534762 |
| Hoxb2 | 2.49664486 |
| Gbx2 | 2.481803147 |
| Gata6 | 2.43133325 |
| Zfp97 | 2.4298404 |
| Hoxb6 | 2.395126952 |
| Zfp42 | 2.377519684 |

| | |
|---|---|
| Foxh1 | 2.34767893 |
| Cdkn2a | 2.339053266 |
| Prdm5 | 2.328611124 |
| Rem2 | 2.328609415 |
| Ebf3 | 2.328607307 |
| Gsc | 2.328606441 |
| Emx2 | 2.328605752 |
| Tlx3 | 2.328604233 |
| Irx2 | 2.328601177 |
| Phox2b | 2.328600954 |
| Hoxb13 | 2.328598656 |
| Foxb1 | 2.328597307 |
| Cdkn2c | 2.328597024 |
| Znrd1 | 2.253480916 |
| Ring1 | 2.231573286 |
| Bcl11a | 2.217509297 |
| Ndn | 2.177936549 |
| 3632451O06Rik | 2.173364643 |
| Hoxa7 | 2.116907833 |
| Tead2 | 2.11029315 |
| Barx1 | 2.095742916 |
| Hoxd8 | 2.037528093 |
| Nr2f6 | 2.037527222 |
| Zim1 | 2.037522931 |
| Pbx1 | 0.500833813 |
| Klf7 | 0.488256305 |
| Runx1 | 0.482197678 |
| Mkl1 | 0.477510954 |
| Nr6a1 | 0.471580064 |
| Msx2 | 0.465721456 |
| Sncaip | 0.465719826 |
| Tead1 | 0.458529196 |
| Rfx3 | 0.44883474 |
| Bach2 | 0.446552668 |
| Epas1 | 0.44305382 |
| Notch4 | 0.436614427 |
| Nfatc2 | 0.430559051 |
| Zfp423 | 0.428954168 |
| Rest | 0.426690449 |
| Msx1 | 0.407935168 |
| Rab8b | 0.406581296 |
| Hoxc8 | 0.404974799 |
| Notch2 | 0.401080337 |
| Rara | 0.40004122 |
| Nrip1 | 0.390627772 |
| Fosl2 | 0.390510538 |
| Sox6 | 0.390007925 |

| | |
|---|---|
| Hoxa1 | 0.385294008 |
| Etv6 | 0.383834693 |
| Foxp1 | 0.371665351 |
| Satb2 | 0.369167871 |
| Bbx | 0.36527085 |
| Crebbp | 0.352731736 |
| Tle3 | 0.348263271 |
| Nr4a2 | 0.347224823 |
| Sall3 | 0.344977845 |
| Dlx3 | 0.343998873 |
| Helz | 0.333650813 |
| Lbh | 0.332657475 |
| Nfatc1 | 0.309217607 |
| Mitf | 0.296190596 |
| Tle2 | 0.291075769 |
| Pou2f2 | 0.291075367 |
| Hoxa6 | 0.291074582 |
| Nfix | 0.282857727 |
| Esx1 | 0.265542961 |
| Gli3 | 0.250387426 |
| Sox9 | 0.194050391 |
| Rab25 | 0.194049612 |
| Gas7 | 0.179123364 |
| Msc | 0.166328706 |
| Ets1 | 0.126429117 |
| Gata3 | 0.122557923 |
| Mid1 | 0.107624485 |
| Cdx1 | 0.078935708 |
| Pou4f2 | 0.004470273 |
| Nhlh2 | 0.003499881 |
| Nhlh1 | 0.003461262 |
| Lef1 | 0.002470814 |
