## Supplementary Data 3 for "Inferring the Hoxa1 Gene Regulatory Network in Mouse Embryonic Stem Cells: Time-Series RNA-seq Data and Computational Modeling Approach"

NARROMI results: first neighbors of Hoxa1 in GRN

| TF | Target | Strenght | p value |
|---|---|---|---|
| Hoxa1 | 1700001K19Rik | 0.309774 | 1.91E-05 |
| Hoxa1 | 1700001O22Rik | 0.819606 | 0.008612 |
| Hoxa1 | 1700042O10Rik | 0.468963 | 0.000182 |
| Hoxa1 | 2810008D09Rik | 0.941307 | 1.26E-06 |
| Hoxa1 | 2810459M11Rik | 0.361172 | 0.001035 |
| Hoxa1 | 3110021A11Rik | 0.563199 | 0.002839 |
| Hoxa1 | 4933427D06Rik | 0.540192 | 0.001766 |
| Hoxa1 | Acaca | 0.438748 | 0.001826 |
| Hoxa1 | Adamts4 | 0.510268 | 5.35E-08 |
| Hoxa1 | Add1 | 0.555761 | 0.003604 |
| Hoxa1 | Adrb3 | 0.554763 | 0.003785 |
| Hoxa1 | Ager | 0.074576 | 1.19E-06 |
| Hoxa1 | Agpat3 | 0.781601 | 1.17E-08 |
| Hoxa1 | Ahi1 | 0.790524 | 7.72E-14 |
| Hoxa1 | Alox12 | 0.409041 | 5.02E-07 |
| Hoxa1 | Ano3 | 0.371816 | 3.84E-05 |
| Hoxa1 | Ap3b2 | 0.783975 | 1E-32 |
| Hoxa1 | Apol7b | 0.694203 | 5.13E-05 |
| Hoxa1 | Arhgap8 | 0.40332 | 8.2E-05 |
| Hoxa1 | Arhgef10 | 0.518222 | 0.003802 |
| Hoxa1 | Armcx2 | 0.850624 | 4.71E-16 |
| Hoxa1 | Armcx3 | 0.833394 | 4.47E-08 |
| Hoxa1 | Arsb | 0.913268 | 0.002408 |
| Hoxa1 | Art5 | 0.961787 | 1E-32 |
| Hoxa1 | Asgr2 | 0.483418 | 0.000277 |
| Hoxa1 | Asphd2 | 0.935709 | 1.41E-08 |
| Hoxa1 | Axin2 | 0.808095 | 1.27E-11 |
| Hoxa1 | B3galt1 | 0.774165 | 0.001862 |
| Hoxa1 | Batf3 | 0.486537 | 0.001184 |
| Hoxa1 | Bbx | 0.650948 | 0.00559 |
| Hoxa1 | BC026585 | 0.604092 | 1.33E-05 |
| Hoxa1 | Bicc1 | 0.872697 | 0.005306 |
| Hoxa1 | C030039L03Rik | 0.651774 | 0.000599 |
| Hoxa1 | C130074G19Rik | 0.40332 | 0.005003 |
| Hoxa1 | C1ql4 | 0.714648 | 1.41E-07 |
| Hoxa1 | Cacna1h | 0.985352 | 1E-32 |
| Hoxa1 | Cacna2d2 | 0.341205 | 3.73E-07 |
| Hoxa1 | Cand2 | 0.890511 | 0.004947 |
| Hoxa1 | Ccdc172 | 0.782524 | 1.1E-08 |
| Hoxa1 | Ccnd1 | 0.911835 | 0.003495 |
| Hoxa1 | Cd99l2 | 0.918258 | 6.23E-05 |
| Hoxa1 | Chn2 | 0.463831 | 0.001248 |
| Hoxa1 | Chrm4 | 0.809388 | 3.23E-06 |
| Hoxa1 | Chst8 | 0.647029 | 1E-32 |
| Hoxa1 | Cib2 | 0.724029 | 0.000249 |

| | | | |
|---|---|---|---|
| Hoxa1 | Cldn4 | 0.500339 | 0.000309 |
| Hoxa1 | Cltb | 0.563957 | 0.007033 |
| Hoxa1 | Crat | 0.965623 | 2E-15 |
| Hoxa1 | Ctdsp2 | 0.724337 | 0.007439 |
| Hoxa1 | Cxcl14 | 0.929324 | 0.004466 |
| Hoxa1 | Cxx1c | 0.746209 | 1.33E-09 |
| Hoxa1 | Cyp26a1 | 0.426254 | 0.000108 |
| Hoxa1 | Cyp7b1 | 0.944444 | 1.05E-13 |
| Hoxa1 | D430019H16Rik | 0.502422 | 1E-32 |
| Hoxa1 | Dap | 0.624471 | 8.91E-06 |
| Hoxa1 | Ddit4l | 0.779188 | 1.61E-05 |
| Hoxa1 | Dennd2a | 0.925766 | 1E-32 |
| Hoxa1 | Dhrs3 | 1.140224 | 1E-32 |
| Hoxa1 | Dlg4 | 0.42014 | 4.6E-05 |
| Hoxa1 | Dlgap3 | 0.74247 | 0.000292 |
| Hoxa1 | Dmrtb1 | 0.618722 | 1.88E-08 |
| Hoxa1 | Dnaja4 | 0.981227 | 4.83E-05 |
| Hoxa1 | Dusp18 | 0.880968 | 0.001469 |
| Hoxa1 | E330009J07Rik | 0.314175 | 0.00022 |
| Hoxa1 | Ebp | 0.815713 | 0.006158 |
| Hoxa1 | Ednra | 0.460559 | 0.005253 |
| Hoxa1 | Elovl4 | 0.997048 | 6.17E-12 |
| Hoxa1 | Emilin3 | 0.411143 | 6.05E-05 |
| Hoxa1 | Enc1 | 0.962234 | 1.1E-07 |
| Hoxa1 | Fads2 | 0.740723 | 1.11E-05 |
| Hoxa1 | Fam114a1 | 0.552014 | 7.07E-06 |
| Hoxa1 | Fam184a | 0.653698 | 0.000389 |
| Hoxa1 | Fam217b | 0.735083 | 3.53E-07 |
| Hoxa1 | Fam3b | 0.32997 | 0.000123 |
| Hoxa1 | Fam83d | 0.648938 | 7.26E-07 |
| Hoxa1 | Fasn | 0.416233 | 0.001065 |
| Hoxa1 | Fastkd2 | 0.999559 | 2.02E-07 |
| Hoxa1 | Fbln1 | 0.717031 | 0.004012 |
| Hoxa1 | Fgd6 | 0.555841 | 0.00015 |
| Hoxa1 | Fignl2 | 0.962937 | 1E-32 |
| Hoxa1 | Folr4 | 0.459268 | 8.23E-06 |
| Hoxa1 | Foxa1 | 0.748393 | 3.95E-05 |
| Hoxa1 | Foxo6 | 0.361338 | 3.29E-07 |
| Hoxa1 | Fras1 | 0.687192 | 0.007457 |
| Hoxa1 | Fsip1 | 0.693882 | 1.57E-16 |
| Hoxa1 | Galnt12 | 0.857565 | 1E-32 |
| Hoxa1 | Gatsl2 | 0.474486 | 1E-32 |
| Hoxa1 | Gdpd2 | 0.688703 | 0.000403 |
| Hoxa1 | Gfpt1 | 0.660782 | 0.006902 |
| Hoxa1 | Gm13119 | 0.763552 | 0.00027 |
| Hoxa1 | Gm2022 | 0.773512 | 9.62E-08 |
| Hoxa1 | Gm364 | 0.863065 | 0.000123 |

| | | | |
|---|---|---|---|
| Hoxa1 | Gm4340 | 0.711668 | 0.000857 |
| Hoxa1 | Gm44 | 0.285896 | 0.00237 |
| Hoxa1 | Gm5039 | 0.696925 | 1E-32 |
| Hoxa1 | Gm6880 | 0.749666 | 0.007174 |
| Hoxa1 | Gm7104 | 0.920629 | 0.00941 |
| Hoxa1 | Gm8300 | 0.917822 | 7.1E-06 |
| Hoxa1 | Gm9 | 0.948276 | 0.00042 |
| Hoxa1 | Gnao1 | 0.924628 | 0.001272 |
| Hoxa1 | Gnaz | 0.450761 | 1.56E-05 |
| Hoxa1 | Gng11 | 0.775564 | 1.5E-13 |
| Hoxa1 | Gpm6a | 0.916432 | 1E-32 |
| Hoxa1 | Gpr153 | 0.866956 | 3.75E-06 |
| Hoxa1 | Gpr157 | 0.497044 | 0.00039 |
| Hoxa1 | Greb1l | 0.787721 | 0.009984 |
| Hoxa1 | Grifin | 0.80578 | 3.64E-10 |
| Hoxa1 | Grik5 | 0.680232 | 3.1E-10 |
| Hoxa1 | Guca1a | 0.688286 | 4.58E-05 |
| Hoxa1 | Gypc | 0.596409 | 0.000124 |
| Hoxa1 | H1f0 | 0.931005 | 0.000403 |
| Hoxa1 | H1fx | 0.465987 | 0.001996 |
| Hoxa1 | H2-Q7 | 0.585804 | 1.65E-08 |
| Hoxa1 | Hdhd3 | 0.823464 | 1E-32 |
| Hoxa1 | Hdx | 0.667598 | 0.000398 |
| Hoxa1 | Hid1 | 0.54461 | 0.00072 |
| Hoxa1 | Hist3h2ba | 0.913016 | 1E-32 |
| Hoxa1 | Hoxa1 | 4.6 | 6.99E-15 |
| Hoxa1 | Hoxa10 | 0.517379 | 0.000533 |
| Hoxa1 | Hoxa3 | 0.872904 | 7.95E-09 |
| Hoxa1 | Hoxa4 | 0.443988 | 2.05E-05 |
| Hoxa1 | Hoxa5 | 0.949754 | 1E-32 |
| Hoxa1 | Hoxa7 | 0.823118 | 7.38E-07 |
| Hoxa1 | Hoxb1 | 0.403251 | 1.34E-06 |
| Hoxa1 | Hoxb4 | 0.641277 | 1.7E-11 |
| Hoxa1 | Hoxc4 | 0.80911 | 0.002447 |
| Hoxa1 | Hs3st6 | 0.291741 | 1E-32 |
| Hoxa1 | Hsd11b2 | 0.499422 | 1E-32 |
| Hoxa1 | Icam1 | 0.809782 | 0.001068 |
| Hoxa1 | Il17b | 0.842598 | 0.004462 |
| Hoxa1 | Il4 | 0.359603 | 3.14E-16 |
| Hoxa1 | Insr | 0.487379 | 4.56E-07 |
| Hoxa1 | Irgm2 | 0.784114 | 1E-32 |
| Hoxa1 | Josd2 | 0.898249 | 1E-32 |
| Hoxa1 | Kbtbd11 | 0.643122 | 9.21E-08 |
| Hoxa1 | Kcnj10 | 0.992216 | 3.11E-15 |
| Hoxa1 | Kcnk6 | 0.549055 | 2.84E-11 |
| Hoxa1 | Kif13b | 0.391314 | 5.68E-05 |
| Hoxa1 | Kif21b | 0.744302 | 6.23E-11 |

| | | | |
|---|---|---|---|
| Hoxa1 | Kif3c | 0.938596 | 0.002278 |
| Hoxa1 | Klhl5 | 0.895307 | 2.24E-06 |
| Hoxa1 | Krt42 | 0.892748 | 0.005114 |
| Hoxa1 | Lgals4 | 0.887579 | 0.007462 |
| Hoxa1 | Lmo2 | 0.245657 | 1.18E-05 |
| Hoxa1 | Lonrf3 | 0.928814 | 1E-32 |
| Hoxa1 | Lrp1 | 0.8452 | 1.41E-07 |
| Hoxa1 | Lrp5 | 0.634494 | 0.000517 |
| Hoxa1 | Lrrc29 | 0.443406 | 1.33E-08 |
| Hoxa1 | Lyn | 0.872905 | 0.009587 |
| Hoxa1 | Maged1 | 1.036585 | 0.000961 |
| Hoxa1 | Magel2 | 0.82995 | 1.58E-05 |
| Hoxa1 | Map3k5 | 0.758769 | 0.001117 |
| Hoxa1 | Map6d1 | 0.592295 | 5.79E-07 |
| Hoxa1 | Marveld1 | 0.850316 | 1E-32 |
| Hoxa1 | Mdga1 | 0.430086 | 0.006156 |
| Hoxa1 | Med14 | 0.53875 | 0.008118 |
| Hoxa1 | Metrnl | 0.554397 | 0.000791 |
| Hoxa1 | Mgme1 | 0.890354 | 0.00332 |
| Hoxa1 | Mlkl | 0.769073 | 1E-32 |
| Hoxa1 | Mogat2 | 0.610176 | 6.7E-11 |
| Hoxa1 | Moxd1 | 0.924463 | 0.003099 |
| Hoxa1 | Ms4a4d | 0.686971 | 1E-32 |
| Hoxa1 | Msx1 | 0.588769 | 0.000879 |
| Hoxa1 | Mum1l1 | 0.838622 | 1.06E-05 |
| Hoxa1 | Nhp2 | 0.851169 | 2.49E-14 |
| Hoxa1 | Nkx3-1 | 0.400432 | 1.11E-05 |
| Hoxa1 | Nphs1 | 0.875336 | 0.005608 |
| Hoxa1 | Nr4a1 | 0.908934 | 1.12E-05 |
| Hoxa1 | Ntrk1 | 0.553819 | 5.77E-09 |
| Hoxa1 | Nudt4 | 0.861541 | 2.54E-06 |
| Hoxa1 | Olfr815 | 0.904109 | 8.66E-05 |
| Hoxa1 | Olig3 | 0.184231 | 3.58E-10 |
| Hoxa1 | Parp12 | 0.399311 | 1.27E-07 |
| Hoxa1 | Pcsk9 | 0.35838 | 0.000491 |
| Hoxa1 | Pdgfrl | 0.654526 | 0.006899 |
| Hoxa1 | Phf13 | 0.932484 | 1.43E-06 |
| Hoxa1 | Phka1 | 0.912466 | 0.000583 |
| Hoxa1 | Phlda2 | 0.971085 | 1E-32 |
| Hoxa1 | Pknox2 | 0.896126 | 7.77E-16 |
| Hoxa1 | Plekhg4 | 0.875593 | 3.63E-05 |
| Hoxa1 | Plekhh2 | 0.49515 | 1E-32 |
| Hoxa1 | Plxnc1 | 0.843108 | 3.91E-05 |
| Hoxa1 | Prkcb | 0.824422 | 0.000504 |
| Hoxa1 | Prrg4 | 0.159405 | 0.00301 |
| Hoxa1 | Prtg | 0.934075 | 0.004519 |
| Hoxa1 | Ptprz1 | 0.757131 | 0.000111 |

| | | | |
|---|---|---|---|
| Hoxa1 | Purg | 0.532171 | 3.35E-08 |
| Hoxa1 | Qpct | 0.547728 | 0.001437 |
| Hoxa1 | Rab38 | 0.986889 | 1E-32 |
| Hoxa1 | Rab43 | 0.685409 | 0.000554 |
| Hoxa1 | Rab6b | 0.273112 | 0.002403 |
| Hoxa1 | Raet1d | 0.584063 | 0.008538 |
| Hoxa1 | Rap1gap2 | 0.894178 | 1E-32 |
| Hoxa1 | Rasl10b | 0.504464 | 3.74E-05 |
| Hoxa1 | Rbm43 | 0.433775 | 7.22E-07 |
| Hoxa1 | Rbp1 | 1.174151 | 9.14E-10 |
| Hoxa1 | Rec8 | 0.787209 | 1E-32 |
| Hoxa1 | Reck | 0.520946 | 1.35E-05 |
| Hoxa1 | Rem2 | 0.357234 | 7.74E-05 |
| Hoxa1 | Rhof | 0.489096 | 0.004432 |
| Hoxa1 | Rhox10 | 0.932905 | 1E-32 |
| Hoxa1 | Rnls | 0.438613 | 0.001095 |
| Hoxa1 | Rnpep | 0.986325 | 2.27E-05 |
| Hoxa1 | Rtn2 | 0.713882 | 0.003534 |
| Hoxa1 | Rtn4rl2 | 0.458552 | 3.21E-05 |
| Hoxa1 | Sec24d | 0.810433 | 1E-32 |
| Hoxa1 | Sema3f | 0.733452 | 2.14E-11 |
| Hoxa1 | Sesn3 | 0.750516 | 0.000111 |
| Hoxa1 | Sfi1 | 0.742659 | 0.008958 |
| Hoxa1 | Sh3kbp1 | 0.830345 | 0.003755 |
| Hoxa1 | Shank2 | 0.446504 | 8.21E-05 |
| Hoxa1 | Shisa3 | 0.973549 | 1E-32 |
| Hoxa1 | Shmt1 | 0.978944 | 0.000566 |
| Hoxa1 | Slc18a3 | 0.905319 | 1.6E-13 |
| Hoxa1 | Slc18b1 | 0.709312 | 8.51E-10 |
| Hoxa1 | Slc1a5 | 0.388027 | 1.82E-08 |
| Hoxa1 | Slc1a6 | 0.609337 | 4.97E-08 |
| Hoxa1 | Slc22a17 | 0.781752 | 7.45E-05 |
| Hoxa1 | Slc22a21 | 0.926415 | 0.000173 |
| Hoxa1 | Slc25a10 | 0.972148 | 2.61E-05 |
| Hoxa1 | Slc38a5 | 0.852087 | 0.000663 |
| Hoxa1 | Slc41a3 | 0.448042 | 9.74E-05 |
| Hoxa1 | Slc45a3 | 0.722188 | 3.16E-05 |
| Hoxa1 | Slc4a2 | 0.725947 | 2.9E-06 |
| Hoxa1 | Slitrk5 | 0.343829 | 0.004625 |
| Hoxa1 | Smc1b | 0.851937 | 0.001249 |
| Hoxa1 | Snhg9 | 0.167978 | 1.08E-06 |
| Hoxa1 | Sox10 | 0.538287 | 0.006874 |
| Hoxa1 | Sox13 | 0.867179 | 0.000768 |
| Hoxa1 | Speg | 0.629151 | 8.78E-06 |
| Hoxa1 | Spink2 | 0.130995 | 0.004238 |
| Hoxa1 | Ssbp3 | 0.7626 | 0.003556 |
| Hoxa1 | St6gal1 | 0.93929 | 1E-32 |

| | | | |
|---|---|---|---|
| Hoxa1 | Stc2 | 0.462033 | 1.73E-15 |
| Hoxa1 | Stmn3 | 0.938911 | 0.005629 |
| Hoxa1 | Stx1b | 0.823738 | 1.8E-08 |
| Hoxa1 | Stxbp4 | 0.434833 | 1.08E-06 |
| Hoxa1 | Sult2b1 | 0.490109 | 1.73E-07 |
| Hoxa1 | Sult5a1 | 0.421464 | 0.006769 |
| Hoxa1 | Syngr4 | 0.246085 | 6.52E-11 |
| Hoxa1 | Syt17 | 0.437329 | 3.18E-05 |
| Hoxa1 | Tap1 | 0.987834 | 1.26E-05 |
| Hoxa1 | Tbc1d8b | 0.441419 | 0.000183 |
| Hoxa1 | Tcerg1l | 0.520568 | 2.91E-05 |
| Hoxa1 | Tdpoz3 | 0.90671 | 4.26E-08 |
| Hoxa1 | Tgm2 | 0.942478 | 1E-32 |
| Hoxa1 | Tktl1 | 0.482086 | 1.9E-06 |
| Hoxa1 | Tlcd2 | 0.700638 | 2.06E-13 |
| Hoxa1 | Tm6sf1 | 0.734418 | 8.62E-05 |
| Hoxa1 | Tmco3 | 0.760774 | 0.000611 |
| Hoxa1 | Tmem119 | 0.776612 | 1E-32 |
| Hoxa1 | Tmem132e | 0.952473 | 2.3E-09 |
| Hoxa1 | Tmem164 | 0.832527 | 3E-05 |
| Hoxa1 | Tmem200b | 0.614816 | 1.02E-08 |
| Hoxa1 | Tmem246 | 0.887856 | 0.009265 |
| Hoxa1 | Tmem74 | 0.531483 | 6.36E-11 |
| Hoxa1 | Tnfaip8l1 | 0.619925 | 2.3E-06 |
| Hoxa1 | Tnfsf9 | 0.672274 | 1E-32 |
| Hoxa1 | Traf1 | 0.947882 | 1E-32 |
| Hoxa1 | Ttc39a | 0.877411 | 1E-32 |
| Hoxa1 | Ttc39c | 0.817279 | 7.36E-05 |
| Hoxa1 | Ttyh2 | 0.615294 | 5.67E-05 |
| Hoxa1 | Tubb2a | 0.835226 | 0.000104 |
| Hoxa1 | Usp17lb | 0.86664 | 1E-32 |
| Hoxa1 | Vegfc | 0.689818 | 5.27E-06 |
| Hoxa1 | Vstm2b | 0.957586 | 4.92E-07 |
| Hoxa1 | Wasf1 | 0.533302 | 0.007053 |
| Hoxa1 | Wfdc1 | 0.42515 | 1E-32 |
| Hoxa1 | Wfikkn1 | 0.84086 | 1E-32 |
| Hoxa1 | Xlr3b | 0.969768 | 4.56E-12 |
| Hoxa1 | Ypel4 | 0.606954 | 5.61E-06 |
| Hoxa1 | Zadh2 | 1.312515 | 1E-32 |
| Hoxa1 | Zcchc11 | 0.764068 | 3.79E-08 |
| Hoxa1 | Zcchc12 | 0.965198 | 1.9E-14 |
| Hoxa1 | Zdhhc14 | 0.518182 | 1.04E-08 |
| Hoxa1 | Zfp618 | 0.899059 | 5.08E-05 |
| Hoxa1 | Zfp69 | 0.926461 | 0.000534 |
| Hoxa1 | Zfp703 | 0.756471 | 2.11E-15 |
| Hoxa1 | Zscan4c | 0.958682 | 1E-32 |
| Hoxa1 | Zscan4d | 0.983301 | 1E-32 |

| | | | |
|---|---|---|---|
| Hoxa1 | Zscan4f | 0.971404 | 1E-32 |
