## Supplementary figures and images for "Inferring the Hoxa1 Gene Regulatory Network in Mouse Embryonic Stem Cells: Time-Series RNA-seq Data and Computational Modeling Approach"

### Supplementary Data 4

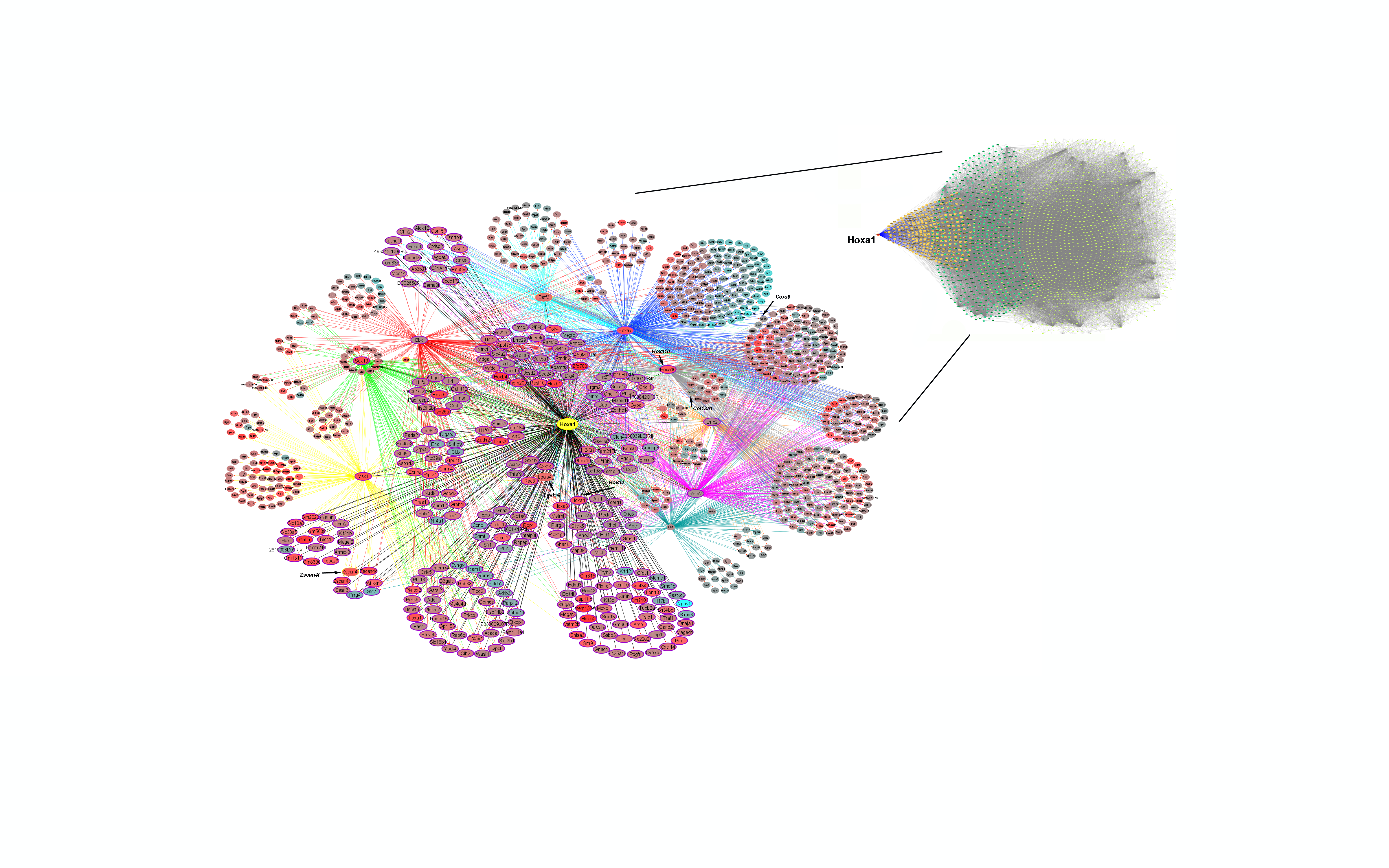
